# Diminutive, degraded but dissimilar: *Wolbachia* genomes from filarial nematodes do not conform to a single paradigm

**DOI:** 10.1101/2020.06.18.160200

**Authors:** Emilie Lefoulon, Travis Clark, Ricardo Guerrero, Israel Cañizales, Jorge Manuel Cardenas-Callirgos, Kerstin Junker, Nathaly Vallarino-Lhermitte, Benjamin L. Makepeace, Alistair C. Darby, Jeremy M. Foster, Coralie Martin, Barton E. Slatko

**Author notes:** present address (Emilie Lefoulon). corresponding author: Barton Slatko, Laboratory.

## Abstract

*Wolbachia* are alpha-proteobacteria symbionts infecting a large range of arthropod species and two different families of nematodes. Interestingly, these endosymbionts are able to induce diverse phenotypes in their hosts: they are reproductive parasites within many arthropods, nutritional mutualists within some insects and obligate mutualists within their filarial nematode hosts. Defining *Wolbachia* “species” is controversial and so they are commonly classified into 16 different phylogenetic lineages, termed supergroups, named A to S. However, available genomic data remains limited and not representative of the full *Wolbachia* diversity; indeed, of the 24 complete genomes and 55 draft genomes of *Wolbachia* available to date, 84% belong to supergroups A and B, exclusively composed of *Wolbachia* from arthropods.

For the current study, we took advantage of a recently developed DNA enrichment method to produce four complete genomes and two draft genomes of *Wolbachia* from filarial nematodes. Two complete genomes, *w*Ctub and *w*Dcau, are the smallest *Wolbachia* genomes sequenced to date (863,988bp and 863,427bp, respectively), as well as the first genomes representing supergroup J. These genomes confirm the validity of this supergroup, a controversial clade due to weaknesses of the multi-locus system typing (MLST) approach. We also produced the first draft *Wolbachia* genome from a supergroup F filarial nematode representative (*w*Mhie), two genomes from supergroup D (*w*Lsig and *w*Lbra) and the complete genome of *w*Dimm from supergroup C.

Our new data confirm the paradigm of smaller *Wolbachia* genomes from filarial nematodes containing low levels of transposable elements and the absence of intact bacteriophage sequences, unlike many *Wolbachia* from arthropods, where both are more abundant. However, we observe differences among the *Wolbachia* genomes from filarial nematodes: no global co-evolutionary pattern, strong synteny between supergroup C and supergroup J *Wolbachia*, and more transposable elements observed in supergroup D *Wolbachia* compared to the other supergroups. Metabolic pathway analysis indicates several highly conserved pathways (haem and nucleotide biosynthesis for example) as opposed to more variable pathways, such as vitamin B biosynthesis, which might be specific to certain host-symbiont associations. Overall, there appears to be no single *Wolbachia-* filarial nematode pattern of co-evolution or symbiotic relationship.

**Graphical abstract:** 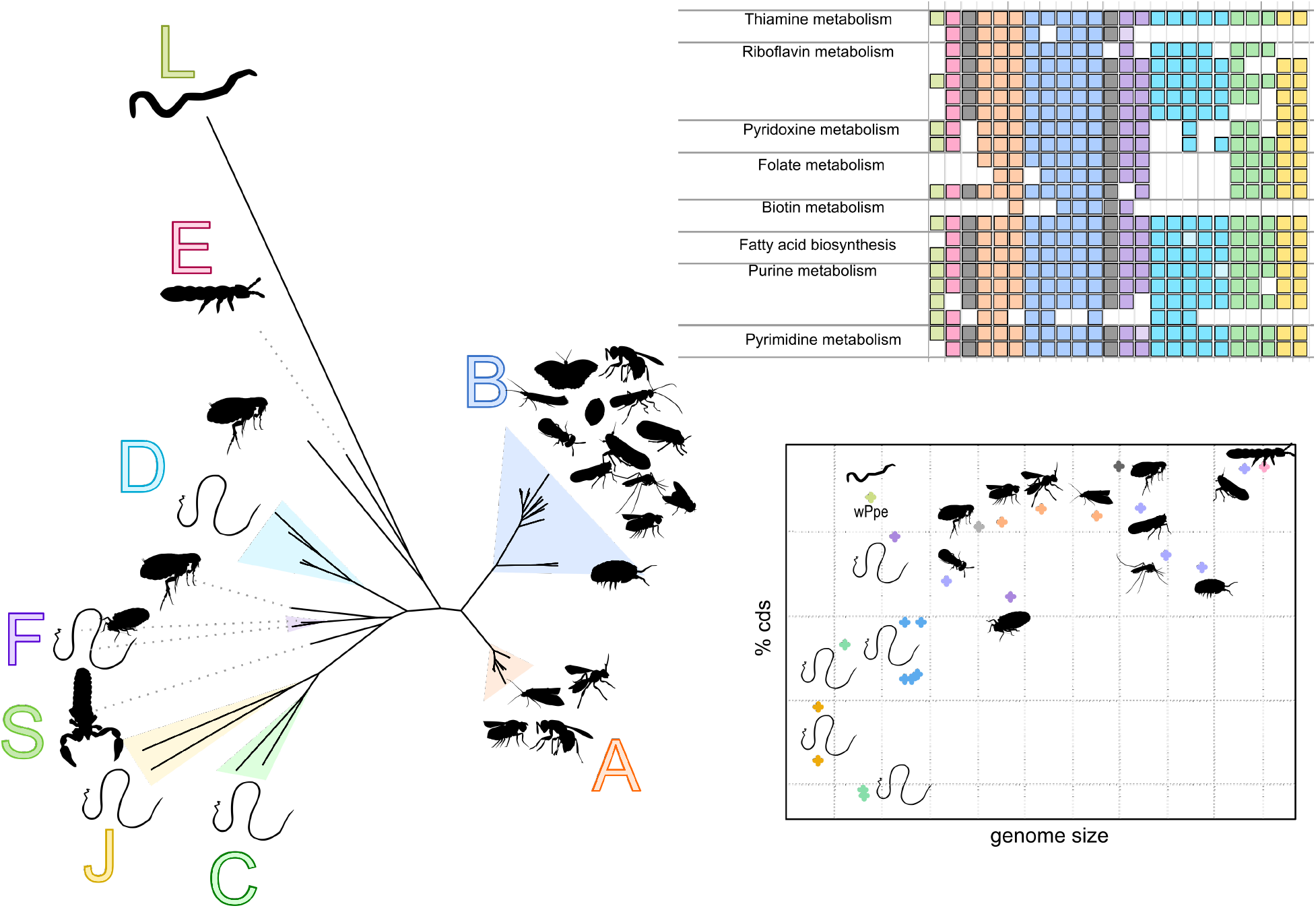

**Repositories:** Data generated are available in GenBank: BioProject PRJNA593581; BioSample SAMN13482485 for *w*Lsig, *Wolbachia* endosymbiont of *Litomosoides sigmodontis* (genome: CP046577); Biosample SAMN15190311 for the nematode host *Litomosoides sigmodontis* (genome: JABVXW000000000); BioSample SAMN13482488 for *w*Dimm*, Wolbachia* endosymbiont of *Dirofilaria* (*D*.) *immitis* (genome: CP046578); Biosample SAMN15190314 for the nematode host *Dirofilaria* (*D*.) *immitis* (genome: JABVXT000000000); BioSample SAMN13482046 for *w*Ctub, *Wolbachia* endosymbiont of *Cruorifilaria tuberocauda* (genome: CP046579); Biosample SAMN15190313 for the nematode host *Cruorifilaria tuberocauda* (genome: JABVXU000000000); BioSample SAMN13482057 for *w*Dcau, *Wolbachia* endosymbiont of *Dipetalonema caudispina* (genome: CP046580); Biosample SAMN15190312 for the nematode host *Dipetalonema caudispina* (genome: JABVXV000000000); BioSample SAMN13482459 for *w*Lbra, *Wolbachia* endosymbiont of *Litomosoides brasiliensis* (genome: WQM000000000); Biosample SAMN15190311 for the nematode host *Litomosoides brasiliensis* (genome: JABVXW000000000); BioSample SAMN13482487 for *w*Mhie, *Wolbachia* endosymbiont of *Madathamugadia hiepei* (genome: WQMP00000000); Biosample SAMN15190315 for the nematode host *Madathamugadia hiepei* (genome: JABVXS000000000). The raw data are available in GenBank as Sequence Read Archive (SRA): SRR10903008 to SRR10903010; SRR10902913 to SRR10902914; SRR10900508 to SRR10900511; SRR10898805 to SRR10898806.

**Data summary:** The authors confirm all supporting data, code and protocols have been provided within the article or through supplementary data files. Eleven Supplementary tables and two supplementary files are available with the online version of this article.

**Impact Statement:** *Wolbachia* are endosymbiotic bacteria infecting a large range of arthropod species and two different families of nematodes, characterized by causing diverse phenotypes in their hosts, ranging from reproductive parasitism to mutualism. While available *Wolbachia* genomic data are increasing, they are not representative of the full *Wolbachia* diversity; indeed, 84% of *Wolbachia* genomes available on the NCBI database to date belong to the two main studied clades (supergroups A and B, exclusively composed of *Wolbachia* from arthropods). The present study presents the assembly and analysis of four complete genomes and two draft genomes of *Wolbachia* from filarial nematodes. Our genomics comparisons confirm the paradigm that smaller *Wolbachia* genomes from filarial nematodes contain low levels of transposable elements and the absence of intact bacteriophage sequences, unlike many *Wolbachia* from arthropods. However, data show disparities among the *Wolbachia* genomes from filarial nematodes: no single pattern of co-evolution, stronger synteny between some clades (supergroups C and supergroup J) and more transposable elements in another clade (supergroup D). Metabolic pathway analysis indicates both highly conserved and more variable pathways, such as vitamin B biosynthesis, which might be specific to certain host-symbiont associations. Overall, there appears to be no single *Wolbachia*-filarial nematode pattern of symbiotic relationship.

## Introduction

The endosymbiotic alpha-proteobacterium *Wolbachia* represents a striking model for studies of symbioses. These bacteria have been detected in a large proportion of arthropods, where they are considered one of the most widespread symbionts (1, 2) and in only two divergent families of parasitic nematodes (filarial nematodes in vertebrates and pratylenchid nematodes feeding on plants) (3, 4). The nature of the relationships with their hosts is particularly fascinating; in arthropods, some *Wolbachia* are reproductive parasites inducing different phenotypes such as cytoplasmic incompatibility, male-killing, parthenogenesis or feminization of genetic males (5); others are nutritional mutualists, as in the case of bedbug symbionts (6), while in the case of the filarial nematodes, their *Wolbachia* are obligate mutualists (7). Numerous *Wolbachia* genomes have been subjected to genomic analyses to determine the nature of the symbiosis (8–12). Some candidate genes have been identified as being involved in cytoplasmic incompatibility (CI) (13, 14), male-killing (15), feminization (16) and nutritional supplementation (17, 18). The CI phenotype is being used as a method to directly suppress pest insect populations or as a means to drive population replacement of mosquito vectors of arboviruses (and potentially other human pathogens, such as *Plasmodium*) with *Wolbachia*-infected individuals that exhibit greatly reduced vector competence (19). With respect to the filarial nematodes, much of the research effort has been focused on drug screening or various treatments targeting *Wolbachia* to kill the parasitic species responsible for human diseases (10, 20–22). Nonetheless, the mechanisms underpinning the obligate mutualism between *Wolbachia* and their filarial hosts remain largely unknown.

To date, three complete genomes of *Wolbachia* from filarial nematodes have been published: first, the symbiont of the human parasite *Brugia malayi*, *w*Bm (23), followed by the symbiont of the bovine parasite *Onchocerca ochengi, w*Oo (24) and lastly, the symbiont of the human parasite *Onchocerca volvulus*, *w*Ov (25). The most striking difference between these genomes and the *Wolbachia* genomes from arthropods appears to be their smaller size [between 957,990 bp for *w*Oo to 1,080,084 bp for *w*Bm versus 1,267,782 bp for *w*Mel (from *Drosophila melanogaster*) or 1,801,626 bp for *w*Fol (from *Folsomia candida*)], as well as the presence of fewer transposable elements [as insertion sequences elements (ISs) and group II intron-associated genes], prophage-related genes and repeat-motif proteins (as ankyrin domains) (24, 26). The hypothesis of potential provisioning of resources [such as haem, riboflavin, flavin adenine dinucleotide (FAD) or nucleotides] by *Wolbachia* to their host nematodes has been suggested, beginning with the first comparative genomic analysis (23). However, the analysis of the highly reduced *w*Oo genome did not strongly support the hypothesis of provisioning of vitamins or cofactors by this strain (riboflavin metabolism and FAD pathways are incomplete) and transcriptomic analysis suggested more of a role in energy production and modulation of the vertebrate immune response (24). Alternatively, the relationship between filarial nematodes and *Wolbachia* may represent a “genetic addiction” rather than genuine mutualism (27).

The notion of *Wolbachia* species remains under debate within the scientific community (28–31). It is commonly accepted to describe the various *Wolbachia* strains as belonging to different phylogenetic lineages as “supergroups” (currently A – S). The first appearance of the “supergroups” designation dates to 1998 (32) but the concept was popularized later by Lo et al. (33). Most of the molecular characterizations of *Wolbachia* strains have been based on either single gene or multi-locus phylogenies (33–37). The supergroups A, B, E, H, I, K, M, N, O, P, Q and S are exclusively composed of symbionts of arthropods (33, 35, 37–42). In contrast, supergroups C, D and J are restricted to filarial nematodes (4, 36, 43, 44), whereas supergroup L is found only in plant-parasitic nematodes (3, 45). Supergroup F is, so far, the only known clade comprising symbionts of filarial nematodes as well as arthropods (33, 34, 46). Initially, the delimitation of these supergroups was defined arbitrarily by a threshold of 2.5% divergence of the *Wolbachia* surface protein gene (*wsp*) (32). However, after it was demonstrated that *wsp* could recombine between *Wolbachia* strains (47), a multi-locus sequence typing approach for *Wolbachia* was proposed (48). These typing methods were developed based almost exclusively on analyses of supergroup A and B *Wolbachia* (48), as they constituted the majority of genomes sequenced at that time. Recently, there has been an effort to revisit the MLST typing paradigm (49) and attempts to classify *Wolbachia* based on genomics (30). However, increased genomic information is needed to appraise the phylogenetic diversity of *Wolbachia* representatives from filarial nematodes. The presently available *Wolbachia* genomic information is not fully representative of *Wolbachia* diversity.

Recently, a method based on biotinylated probes was developed to capture large fragments of *Wolbachia* DNA for sequencing, using PacBio technology (LEFT-SEQ) (50) adapted from previous capture methods using Illumina technology (51, 52). We used this enrichment method to produce draft or complete genomes of *Wolbachia* from a diversity of filarial nematodes species: *Cruorifilaria tuberocauda* and *Dipetalonema caudispina* both parasites of the capybara, a cavy rodent; *Litomosoides brasiliensis*, a parasite of bats; *Litomosoides sigmodontis*, a parasite of cricetid rodents; *Dirofilaria* (*Dirofilaria*) *immitis*, a parasite of canines; and *Madathamugadia hiepei*, a parasite of geckos. These species had been previously characterized as positive for *Wolbachia* infections (two supergroup J, two supergroup D, one supergroup C and one supergroup F, respectively) (36). In the present study, we took advantage of this newly explored diversity to draw a more comprehensive picture of symbiosis between *Wolbachia* and their filarial nematode hosts.

## Methods

### Materials

Eight specimens belonging to six filarial nematode species were studied (Table 1): *Cruorifilaria tuberocauda* (two samples), *Dipetalonema caudispina* (two samples), *Litomosoides brasiliensis*, *Litomosoides sigmodontis*, *Dirofilaria* (*D*.) *immitis* and *Madathamugadia hiepei*. Most of the samples were collected as described in a previous study (53) and all procedures were conducted in compliance with the rules and regulations of the respective national ethical bodies. The *D.* (*D*.) *immitis* specimen was provided by the NIAID/NIH Filariasis Research Reagent Resource Center (MTA University of Wisconsin Oshkosh; www.filariasiscenter.org), and the *L. sigmodontis* specimen was provided by the National Museum of Natural History (MNHN, Paris), where the experimental procedures were carried out in strict accordance with the EU Directive 2010/63/UE and the relevant national legislation (ethical statement n°13845). Supplementary File S1 lists the author(s) and year of parasite and host species collection. In accordance with the generally used nomenclature of *Wolbachia* strains, we have named these newly typed strains after their hosts: *w*Ctub for the symbiont of *C. tuberocauda; w*Dcau for the symbiont of *D. caudispina; w*Lbra for the symbiont of *L. brasiliensis*; *w*Lsig for the symbiont of *L. sigmodontis*; *w*Dimm for the symbiont of *D*. (*D*.) *immitis* and *w*Mhie for the symbiont of *M. hiepei.* In the cases of *w*Lsig and *w*Dimm, they have been referred to as *w*Ls and *w*Di, respectively, in the prior literature (25, 54). However, the lack of a consensus on the nomenclature of *Wolbachia* strains has already led to some confusion, as *w*Di can refer to *Wolbachia* from *Dirofilaria* (*D*.) *immitis* (26) as well as *Wolbachia* from *Diaphorina citri* (55). Therefore, to avoid confusion, we use the longer strain abbreviations in this paper.

**Table 1.**
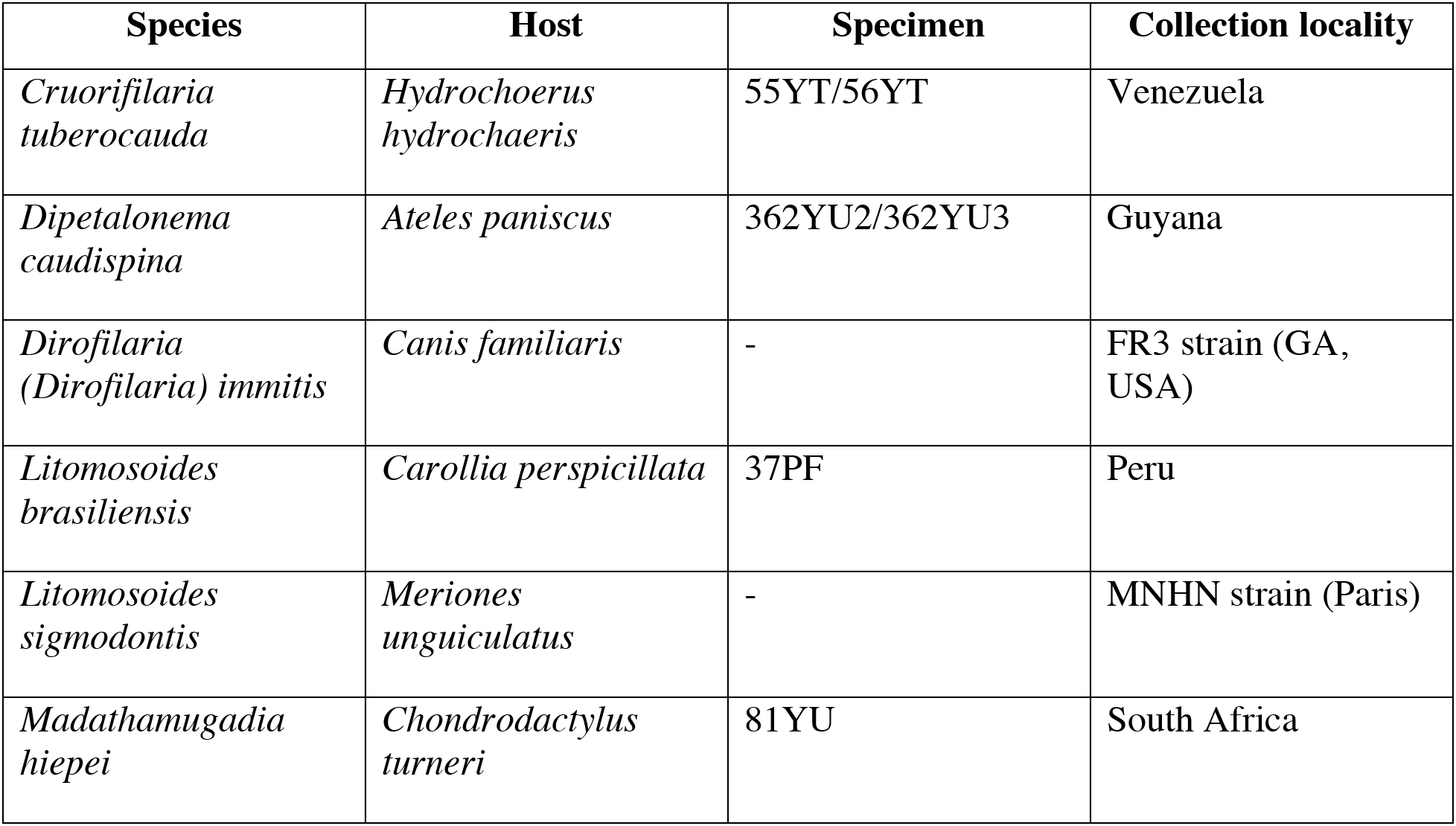
Information on species studied.

The DNA of the *M. hiepei* sample 81YU had been extracted previously and conserved at −20°C for 8 years (53). The DNA of the *D. caudispina* sample 362YU2 had been extracted in January 2015 and then conserved at −20°C for 4 years (unpublished data). A new fragment of a specimen from the same lot (362YU3) was also obtained for new DNA extraction. DNA was extracted from all samples using the DNeasy kit (Qiagen), following the manufacturer’s recommendations, including overnight incubation at 56°C with proteinase K.

### Library preparations

According to the amount and quality of DNA of each sample, different library preparation protocols were utilized (Table 2). We used capture enrichment methods for either Illumina or PacBio sequencing based on the use of biotinylated probes to capture *Wolbachia* DNA (probes designed by Roche NimbleGen) based on 25 complete or draft sequences as described by Lefoulon et al. (50). The Large Enriched Fragment Targeted Sequencing (LEFT-seq) method (50), developed for PacBio sequencing, was used for the freshly extracted DNA of *D*. (*D*.) *immitis, L. sigmodontis, C. tuberocauda* (55YT) and the 4-year old extracted DNA of *D. caudispina* (362YU2). We used 1 μg of DNA for each sample. Regarding the enriched libraries from *D*. (*D*.) *immitis* and *L. sigmodontis*, the last steps were modified, compared to the previously described protocol [Lefoulon et al. (50)]: after the second PCR amplification of the enriched DNA, the libraries were prepared with the SMRTbell Express Template kit v2.0 (PacBio) to perform PacBio Sequel sequencing.

**Table 2.**
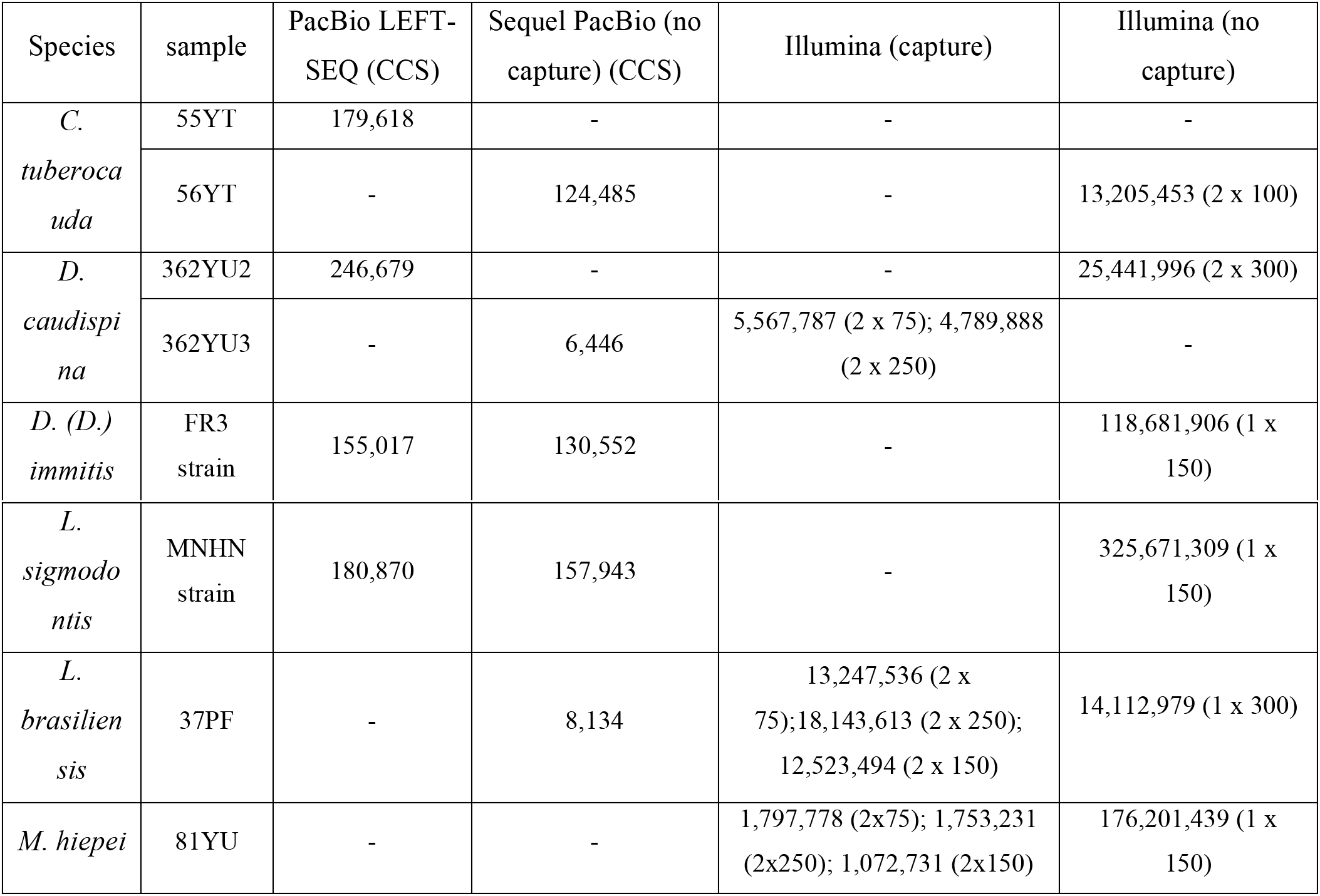
Sequencing data information. PacBio data are the number of circular consensus sequences (CCS) produced with three full passes and a minimum predicted accuracy superior of 90%. Illumina data are number of reads after filtering. Parenthesis indicate length of end-sequence protocol used.

For *M. hiepei* and *L. brasiliensis*, the DNA concentrations were either of low concentration or too fragmented to be used for the LEFT-SEQ protocol. For these, the enrichment method adapted for Illumina sequencing was a hybrid protocol between the method described by Geniez et al. (52) and Lefoulon et al. (50). We followed the same procedure for the freshly extracted DNA of *D. caudispina* (362YU2). DNA samples (50 ng for 81YU, 75 ng for 37PF and 100 ng for 362YU3) were fragmented using the NEBNext® Ultra™ II FS DNA kit (New England Biolabs, USA) at 37°C for 20 minutes (resulting in DNA fragments with an average size of 350 bp). The sheared DNA samples were independently ligated to SeqCap barcoded adaptors (Nimblegen, Roche), to enable processing of multiple samples simultaneously. The ligated DNAs were amplified by PCR and hybridized to the biotinylated probes, according to the SeqCap EZ HyperCap protocol (NimbleGen Roche User’s guide v1.0). For each sample, a library without the enrichment method was also processed using NEBNext® Ultra™ II FS DNA Library Prep kit following the manufacturer’s recommendations (New England Biolabs, USA). The library preparation for *D. caudispina* sample 362YU2 was performed in 2016 at the University of Liverpool. Supplementary libraries without enrichment for PacBio sequencing were also processed for *C. tuberocauda*, *D. caudispina* and *L. brasiliensis* using the SMRTbell Express Template Prep Kit v2.0, following the manufacturer’s recommendations (PacBio, USA). The produced data are summarized in Table 2.

### *De novo* assembly pipeline

The bioinformatics pipeline was slightly different for the six samples because of the variations in sequence protocols (as described above). However, the pre-processing of the reads was similar for both Illumina and PacBio data. The Illumina reads were filtered using the wrapper Trim Galore! (https://www.bioinformatics.babraham.ac.uk/projects/trim_galore/). For PacBio, circular consensus sequences (CCS) were generated using the SMRT® pipe RS_ReadsOfInsert Protocol (PacBio) with a minimum of three full passes and minimum predicted accuracy greater than 90%. The adapter and potential chimeric reads were removed using seqkt (github.com/lh3/seqtk) as described by Lefoulon et al. (50) (analyses were performed with an in-house shell script). When sufficient PacBio reads were obtained, a *de novo* long-read assembly was done using canu (56) according to Lefoulon et al. (50). Otherwise, a first hybrid *de novo* assembly was done using Spades (57). The contigs belonging to *Wolbachia* were detected by nucleotide similarity using blastn (similarity greater than 80%, bitscore greater than 50) (58) and isolated. The remaining contigs were manually curated to eliminate potential non-*Wolbachia* sequence contaminations. To improve the complete assembly of the *Wolbachia* genomes, a selection of reads mapping to the first *Wolbachia* draft genome was produced. The Illumina reads were merged with PEAR (59) (in the case of end-paired reads) and mapped against this contig selection using Bowtie2 (60). The PacBio reads were mapped against this contig selection using ngmlr (with the PacBio preset settings) (61). A second hybrid *de novo* assembly was then performed with this new selection of reads using Unicycler (62). A second selection of *Wolbachia* contigs using blastn was then performed and manually curated to eliminate potential contaminations. Assembly statistics were calculated using QUAST (63). PCR primers were designed to confirm the sites of circularization of the single contigs when applicable (Supplementary file S2).

In addition, contigs representative of filarial nematode genomes were isolated from the first *de novo* assembly. Mapped reads were selected, as described above, and *de novo* assemblies of the host genomes were produced using Unicycler (62).

The completeness of the draft genomes was studied using BUSCO v3 which analyzes the gene content compared to a selection of near-universal single-copy orthologous genes. This analysis was based on 221 genes common among proteobacteria for the draft genomes of *Wolbachia* (proteobacteria_odb9) while it was based on 982 genes common among nematodes for the host draft genomes (nematoda_odb9).

### Comparative genomic analyses and annotation

We used different comparative analyses between the produced draft genomes and a set of eight available complete genomes and seven draft genomes of *Wolbachia* (Table S1): *w*Mel, *Wolbachia* from *Drosophila melanogaster* (NC_002978) *w*Cau, *Wolbachia* from *Carposina sasakii* (CP041215) and *w*Nfla, *Wolbachia* from *Nomada flava* (LYUW00000000) for supergroup A; *w*Pip, *Wolbachia* from *Culex quinquefasciatus* (NC_010981), *w*Tpre, *Wolbachia* from *Trichogramma pretiosum* (NZ_CM003641), *w*Lug, *Wolbachia* from *Nilaparvata lugens* and *w*Stri (MUIY01000000), *Wolbachia* from *Laodelphax striatella* (LRUH01000000) for supergroup B; *w*VulC, *Wolbachia* from *Armadillidium vulgare* (ALWU00000000), closely related to the supergroup B; *w*Ppe, *Wolbachia* from *Pratylenchus penetrans* for supergroup L (NZ_MJMG01000000); *w*Cle, *Wolbachia* from *Cimex lectularius* for supergroup F (NZ_AP013028); *w*Fol*, Wolbachia* from *Folsomia candida* for supergroup E (NZ_CP015510); *w*Bm, *Wolbachia* from *Brugia malayi* (NC_006833), *w*Bp, *Wolbachia* from *Brugia pahangi* (NZ_CP050521) and *w*Wb, *Wolbachia* from *Wuchereria bancrofti* (NJBR02000000) for supergroup D; *w*Ov *Wolbachia* from *Onchocerca volvulus* (NZ_HG810405) and *w*Oo, *Wolbachia* from *Onchocerca ochengi* (NC_018267) for supergroup C; *w*CfeT (NZ_CP051156.1) and *w*CfeJ (NZ_CP051157.1) both *Wolbachia* from *Ctenocephalides felis* [not described as belonging to any supergroup (64)].

We calculated the Average Nucleotide Identity (ANI) between the different *Wolbachia* genomes using ANI Calculator (65) and an in-silico genome-to-genome comparison was done to calculate a digital DNA-DNA hybridization (dDDH) using GGDC (66). The calculation of dDDH allows analysis of species delineation as an alternative to the wet-lab DDH used for current taxonomic techniques. GGDC uses a Genome Blast Distance Phylogeny approach to calculate the probability that an intergenomic distance yielded a DDH larger than 70%, representing a novel species-delimitation threshold (66). We used formula 2 to calculate the dDDH because it is more robust using incomplete draft genomes (67).

The *Wolbachia* genomes were analyzed using the RAST pipeline (68). In order to compare the nature of these genomes using RAST pipeline, we identified the percentage of coding regions of genes (CDS), the presence of mobile elements, the group II intron-associated genes, the phage-like genes and the ankyrin-repeat protein genes. The presence of potential insertion sequences (ISs) was detected using ISSAGA (69) (degraded sequences were not manually curated) and the presence of prophage regions was detected using PHASTER (70). The *Wolbachia* genomes were annotated using Prokka (71). We also examined the correlation between the size of the genome and the previously described genetic characteristics using the Spearman’s rank correlation or Pearson rank correlation tests (if applicable after Shapiro-Wilk test) in the R environment (72). KEGG Orthology (KO) assignments were generated using KASS (KEGG Automatic Annotation Server) (73). KASS assigned orthologous genes by a BLAST comparison against the KEGG genes database using the BBH (bi-directional best hit) method. The same assignment analysis was performed for the newly produced genomes and the set of 15 *Wolbachia* genomes from the NCBI database. The assigned KO were ordered in 160 different KEGG pathways (Table S2). Several pathways which showed differences in the number of assigned genes between *Wolbachia* genomes were selected. A list of genes assigned to these pathways was compiled to study potential losses/acquisitions of these genes across the various *Wolbachia* (Table S3).

### Phylogenomic analyses

Single-copy orthologue genes were identified from a selection of *Wolbachia* genomes using Orthofinder (74). Three phylogenomic studies were performed: the first included only 11 *Wolbachia* genomes from filarial nematodes; the second included only 25 complete genomes and the third included 49 complete or draft genomes (Table S1). The supermatrix of orthologue sequence alignments was generated by Orthofinder (implemented as functionality). The poorly aligned positions of the produced orthologous gene alignments were eliminated using Gblocks (75). The phylogenetic analyses were performed with Maximum Likelihood inference using IQ-TREE (76). The most appropriate model of evolution was evaluated by Modelfinder (implemented as functionality of IQ-TREE) (76). The robustness of each node was evaluated by a bootstrap test (1,000 replicates). The phylogenetic trees were edited in FigTree (https://github.com/rambaut/figtree/) and Inkscape (https://inkscape.org/). To study the evolution of the filarial hosts infected with *Wolbachia*, the same workflow was applied to the amino-acid files previously produced by the BUSCO analysis (Augustus implemented as functionality) based on the set of 982 orthologue genes common among nematodes (nematoda_odb9) (Table S4). The six produced filarial nematode draft genomes were analyzed with five draft genomes available in the database (AAQA00000000 for *B. malayi*; JRWH00000000 for *B. pahangi*; CAWC010000000 for *O. ochengi*, CBVM000000000 for *O. volvulus*, LAQH01000000 for *W. bancrofti*).

### Synteny and coevolutionary analyses

The potential positions of the origin of replication (ORI) were identified based on the ORI position in the *w*Mel and *w*Bm genomes according to Ioannidis et al. (77) for the complete genomes generated for *w*Dimm, *w*Lsig, *w*Dcau and *w*Ctub, as well as available complete genomes of *Wolbachia* from supergroup C (*w*Oo and *w*Ov), supergroup D (*w*Bp), supergroup F (*w*Cle) and *w*CfeJ. The genome sequences were reorganized to start at the potential ORI position to study the genome rearrangement. Then, a pairwise genome alignment of these genomes was produced and plotted using MUMmer v3 (78).

Two global-fit methods were used to study the cophylogenetic pattern between filarial nematodes and their *Wolbachia* symbionts: PACo application (79) and Parafit function (80) both in the R environment (72). For these analyses, we independently produced two ML phylogenies: the phylogenetic tree of the 11 *Wolbachia* from filarial nematodes and the phylogenetic tree of the 11 filarial nematodes as described above. The global-fit method estimates the congruence between two phylogenetic trees changing the ML phylogenies into matrices of pairwise patristic distance, themselves transformed into matrices of principal coordinates (PCo). Then, PACo analysis transformed the symbiont PCo using least-squares superimposition (Procrustes analysis) to minimize the differences with the filarial PCo. The global fit was plotted in an ordination graph.

The congruence of the phylogenies was calculated by the residual sum of squares value (m^2^_XY_) of the Procrustean fit calculation. Subsequently, the square residual of each single association and its 95% confidence interval were estimated for each host-symbiont association and plotted in a bar chart (79). A low residual value represented a strong congruence between symbiont and filarial host. In addition, the global fit was estimated using Parafit function (80) in the R environment (72). The Parafit analysis tests the null hypothesis (H0), that the evolution of the two groups has been independent, by random permutations (1,000,000 permutations) of host-symbiont association (80). This test is based on analysis of the matrix of patristic distances among the hosts and the symbionts as described above for PACo.

## Results

### *De novo* assembly and completeness of draft genomes

We were able to produce complete circular assemblies for four of the genomes, *w*Ctub, *w*Dcau, *w*Lsig, *w*Dimm, as well as a 41-contig draft genome for *w*Lbra and a 208-contig draft genome for *w*Mhie (Table 3). The circularization of *w*Ctub, *w*Dcau, *w*Lsig and *w*Dimm was confirmed by PCR amplification of the sites of circularization of the single contigs (Supplementary file S2). The two supergroup J genomes, *w*Ctub and *w*Dcau, are the smallest observed among all sequenced *Wolbachia* comprising 863,988bp and 863,427 bp, respectively (Table 3). The genome of *w*Dimm displayed a total size of 920,122bp and the total length of *w*Lsig was 1,045,802bp. The draft genome of *w*Lbra had a total length of 1,046,149 bp, very close to the size observed for *w*Lsig. Although the assembly remains fragmented, the draft genome of *w*Mhie has a total length of 1,025,329 bp (Table 3). Varying success in producing complete genome sequences was attributed to DNA quantity and quality (Table 2); for example, the low quantity and quality of DNA obtained for *M. hiepei* and *L. brasiliensis* limited sequencing. The *de novo* assembly was successful with production of a circularized genome in the case of *w*Lsig based on 180,870 CCS PacBio reads, *w*Dimm based on 155,017 CCS PacBio reads and *w*Ctub based on 179,618 CCS PacBio reads. Of the sequenced CCS reads, 94.67% mapped to the draft genome for wLsig, 90.57% for *w*Dimm, but only 35% for *w*Ctub. However, for *w*Dcau, the analysis of the sequenced 246,679 CCS PacBio reads using canu did not produce an accurate draft genome (< 100,000 bp total length). In contrast, a hybrid *de novo* assembly, based on all the sequenced data, produced a draft genome containing one large *Wolbachia* contig which was circularized using minimus2. Only 1.8% of the CCS reads produced with LEFT-SEQ (4,466) mapped to this *w*Dcau draft and the low efficiency was likely due to the 4-year old extracted DNA that was used.

**Table 3.** Draft genomes information

|  | Size (bp) | % GC content | contigs | N50 (bp) | Complete BUSCOs (%) |
| --- | --- | --- | --- | --- | --- |
| nDimm | 83,711,400 | 27.72 | 1,294 | 126,493 | 94.9 |
| wDimm | 920,122 | 32.7 | 1 | 920,122 | 76.0 |
| nLsig | 64,161,459 | 33.96 | 839 | 135,680 | 94.1 |
| wLsig | 1,045,802 | 32.1 | 1 | 1,045,802 | 76.1 |
| nCtub | 75,522,022 | 30.29 | 1,888 | 105,487 | 95.6 |
| wCtub | 863,988 | 32.3 | 1 | 863,988 | 77.4 |
| nDcau | 81,590,899 | 30.44 | 2,615 | 173,032 | 97.7 |
| wDcau | 863,427 | 28 | 1 | 863,427 | 73.4 |
| nLbra | 65,202,511 | 31.74 | 1,026 | 147,890 | 93.4 |
| wLbra | 1,046,149 | 34.5 | 41 | 52,864 | 76.5 |
| nMhie | 77,701,753 | 33.59 | 13,165 | 17,407 | 86.4 |
| wMhie | 1,025,329 | 36.1 | 208 | 6,845 | 75.6 |

Among 221 single-copy orthologous genes conserved among proteobacteria (BUSCO database), 171 and 162 are present in the *w*Ctub and *w*Dcau complete genomes, respectively, suggesting 77.4% and 73.4% BUSCO completeness (Table 3, Table S4). The other two complete genomes, *w*Lsig and *w*Dimm, have 76.1% and 76% BUSCOs completeness with 168 genes identified. The draft genome *w*Lbra has 76.5% complete BUSCOs with 169 genes identified. The draft genome of *w*Mhie has 75.6% BUSCO completeness with 167 genes identified. These levels of completeness are similar to most *Wolbachia* genomes from filarial nematodes. For example, *Wolbachia* from *B. malayi*, *w*Bm, has a higher level with 175 complete genes identified (79.2%) and *Wolbachia* from *O. ochengi, w*Oo, has a lower level of completeness with 165 complete genes identified (74.7%) (Table S4). In general, *Wolbachia* genomes from arthropods present higher levels of complete BUSCOs [*e.g*.: *Wolbachia* from *D. melanogaster, w*Mel, has 180 BUSCO genes (81.4%)]. The higher BUSCOs in these genomes could be because these genomes are less degraded than those of filarial *Wolbachia.*

Along with the assembly of the *Wolbachia* genome, draft genomes of the nematode hosts were produced (Table 3): a 1,888-contig draft genome of *C. tuberocauda*, nCtub, of 75,522,022 bp total length; an 839-contig draft genome of *L. sigmodontis*, nLsig, of 64,161,459 bp total length; a 1,026-contig draft genome of *L. brasiliensis*, nLbra, of 65,202,511 bp total length; a 1,294-contig draft genome of *D.* (*D*.) *immitis*, nDimm, of 83,711,400 bp total length, a 2,615-contig draft genome of *D. caudispina*, nDcau, of 81,590,899 bp total length and a 13,165-contig draft genome of *M. hiepei*, nMhie, of 77,701,753 bp total length. Among 982 single-copy orthologous genes conserved among nematodes (BUSCO database), the draft genome of *D. caudispina* shows the highest level of completeness with 959 genes detected (97.7%). The draft genomes of *C. tuberocauda* and *L. brasiliensis* show similar results with 939 (95.6%) and 917 (93.4%) genes detected. The draft genome of *M. hiepei* has the lowest level of completeness with 849 (86.4%) genes identified (Table S4). Draft genomes of nDimm and nLsig have previously been published with total lengths similar to these results, with respectively 84,888,114bp (ASM107739v1) and 64,813,410bp (ASM90053727v1), respectively but with lower N50 values (15,147 and 45,863, respectively) (81).

### Average Nucleotide Identity and digital DNA-DNA hybridization

The Average Nucleotide Identity (ANI) calculation indicates that *w*Ctub and *w*Dcau are divergent from other *Wolbachia*. For both, the most similar genome is *w*Ov with 83% and 84% identity, respectively (Figure 1). The draft genome *w*Lbra shows a stronger similarity of 90% with the representatives of supergroup D: *w*Bm, *w*Bp, *w*Wb and *w*Lsig. The draft genome *w*Mhie is most similar to *w*Cle from supergroup F with 95% identity (Figure 1). Typically, strains representative of the same supergroup share strong identity: 99% for *w*Oo and *w*Ov (from supergroup C), 99% for *w*Bm and *w*Bp (from supergroup D), 97% for *w*Bm or wBp and *w*Wb (from supergroup D), 95% for *wPip* and *wstri* (from supergroup B) and 97% for *wMel* and *wCau* (from supergroup A). A digital DNA-DNA hybridization (dDDH) (66) metric higher than 70 indicates that the two strains might belong to the same species (see Materials and Methods). The present *in-silico* genome-to-genome comparison shows only five cases which might be considered as similar strains of *Wolbachia*: *w*Cau and *w*Mel; *w*Bm and *w*Bp; *w*Bm and *w*Wb; *w*Bp and *w*Wb; and *w*Oo and *w*Ov (Figure 1). These proximities had been previously suggested (30). Both ANI and dDDH analyses suggest that the four newly sequenced *Wolbachia* genomes are divergent from published *Wolbachia* genomes.

**Figure 1.**
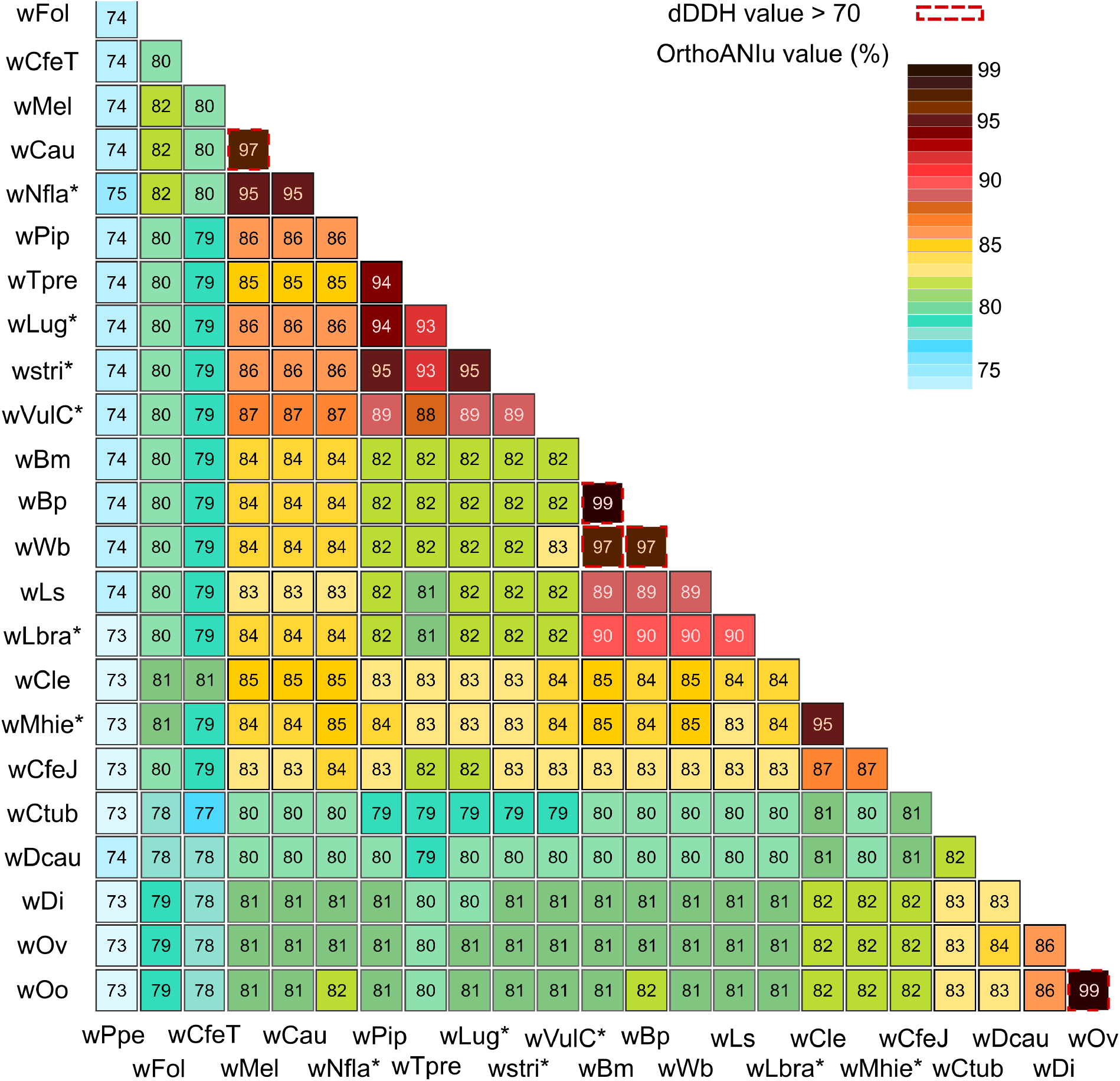
Graphic representation of ANI and dDDH calculations for *Wolbachia* genomes. The Average Nucleotide Identity (ANI) between 23 complete genomes of *Wolbachia* evaluated using the ANI Calculator and the probability of a digital DNA-DNA hybridization (DDH) greater than 70 using GGDC.

### Phylogenomic analyses

A total of 367 single-copy orthologous genes were identified from among the available 25 complete *Wolbachia* genomes. The newly sequenced *w*Ctub, *w*Dcau, *w*Lsig and *w*Dimm genomes were included in the Maximum Likelihood phylogenetic analyses based on these 367 orthologous genes (Figure 2A). This phylogenetic analysis confirms that *w*Lsig belongs to supergroup D and *w*Dimm belongs to supergroup C, as had been previously described (4). *w*Ctub and *w*Dcau were grouped in the same clade, a sister taxon of the supergroup C. *w*Ctub and *w*Dcau had been previously described as representatives of supergoup J (36). The present phylogenomic analysis supports the hypothesis that *Wolbachia* supergroup J is a clade distinct from *Wolbachia* supergroup C, although the two clades are closely related. A total of 160 single-copy orthologous genes were identified among the 49 complete and draft *Wolbachia* genomes (Figure 2B). The two phylogenomic analyses indicate the same topologies for the complete genomes *w*Ctub, *w*Dcau, *w*Lsig and *w*Dimm. In addition, the phylogenomic analysis based on 160 genes shows that *w*Lbra is closely related to *w*Lsig, as representative of supergroup D, and *w*Mhie is closely related to *w*Cle as a representative of supergroup F. The draft genome *w*Mhie is the first representative of supergroup F infecting a filarial nematode and the phylogenomic analysis confirms the evolutionary history of *w*Lbra and *w*Mhie, as previously deduced from multi-locus phylogenies (36, 46).

**Figure 2.**
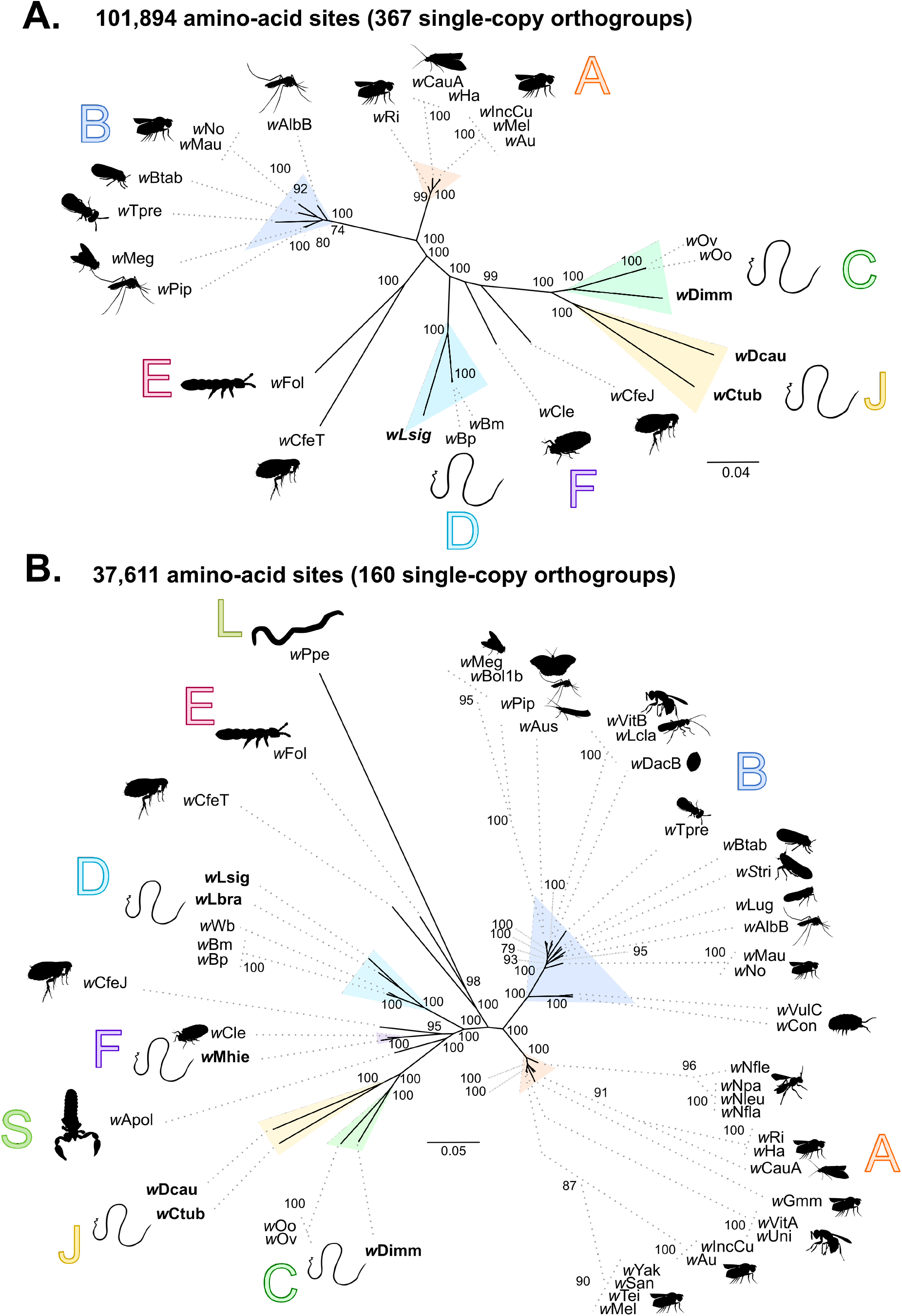
Phylogenomics analyses of *Wolbachia*. The topologies were inferred using Maximum Likelihood (ML) inference using IQTREE. Nodes are associated with bootstrap values based on 1,000 replicates; only bootstrap values superior to 70 are indicated. The *Wolbachia* supergroups (A–L) are indicated. A) Analysis based on concatenation of 365 single-copy orthogroups representing a 101,894 amino-acid matrix. The best-fit model calculated using ModelFinder according to the BIC index was JTT+F+I+G4. B) Analysis based on concatenation of 160 single-copy orthogroups representing a 37,611 amino-acid matrix. The best-fit model calculated using ModelFinder according to the BIC index was JTT+F+I+G4. The *Wolbachia* supergroups (A–L) are indicated and associated with different colors: orange for supergroup A, dark blue for B, light green for C, light blue for D, pink for E, purple for F, yellow for J, khaki green for L.

The validity of supergroup J had been previously discussed; some studies using multi-locus phylogenies suggest that *Wolbachia* from *Dipetalonema gracile* (historically, the only known representative of supergroup J) belongs to supergroup C (35, 82, 83). Interestingly, the *ftsZ* gene used in these MLST studies could not be detected in the *w*Dcau and *w*Ctub genomes. In addition, some multi-locus studies had observed PCR amplification of this gene to be unsuccessful within supergroup J *Wolbachia* (36, 84) while other studies included an *ftsZ* sequence from *Wolbachia* from *D. gracile* (34). To resolve this contradiction, we compared the 33 sequences of *Wolbachia* from *D. gracile* available on the NCBI database and our complete *w*Dcau genome, which should be closely related, using nblast. For six of these 33 sequences, from the *ftsZ* gene, it appears unlikely that they belong to *Wolbachia* from *D. gracile*, only having between 72.37% to 89.91% identity with *w*Dcau (while all other PCR sequences show 95.36-99.98% identity) (Table S5). Four of these six sequences are identical to genes of *Wolbachia* from *Drosophila* spp., one sequence is closely related to genes of *Wolbachia* from supergroup B (82) and one sequence is closely related to genes of *Wolbachia* from supergroup C (the *ftsZ* sequence) (34) (Table S5). Thus, our data suggest that the variable position of *Wolbachia* from *D. gracile* in previous multi-locus phylogenies might be linked to contamination or errors of sequence submission.

### Synteny conservation and co-evolutionary analysis

Strong conservation of synteny among supergroup C genomes and supergroup J genomes was observed (Figure 3). It had been previously shown that the supergroup C genomes *w*Oo and *w*Dimm exhibit a low level of intra-genomic recombination (26). Our results indicate a similar pattern of strong conservation of synteny among supergroup J genomes (*w*Dcau and *w*Ctub) and, more interestingly, between supergroup J genomes and *w*Dimm in supergroup C (Figure 3). This is in contrast to alignment of the complete genomic assemblies between the supergroup D genomes which show more rearrangement. Of further interest is the observation that a different level of rearrangement can be observed between *w*Bm and *w*Lsig or *w*Bp and *w*Lsig, even when *w*Bm and *w*Bp show less rearrangement between them. While *w*Bm and *w*Bp are characterized by a strong identity as described above (Figure 1), similar to that observed between *w*Oo and *w*Ov, they show more rearrangement (Figure 3).

**Figure 3.**
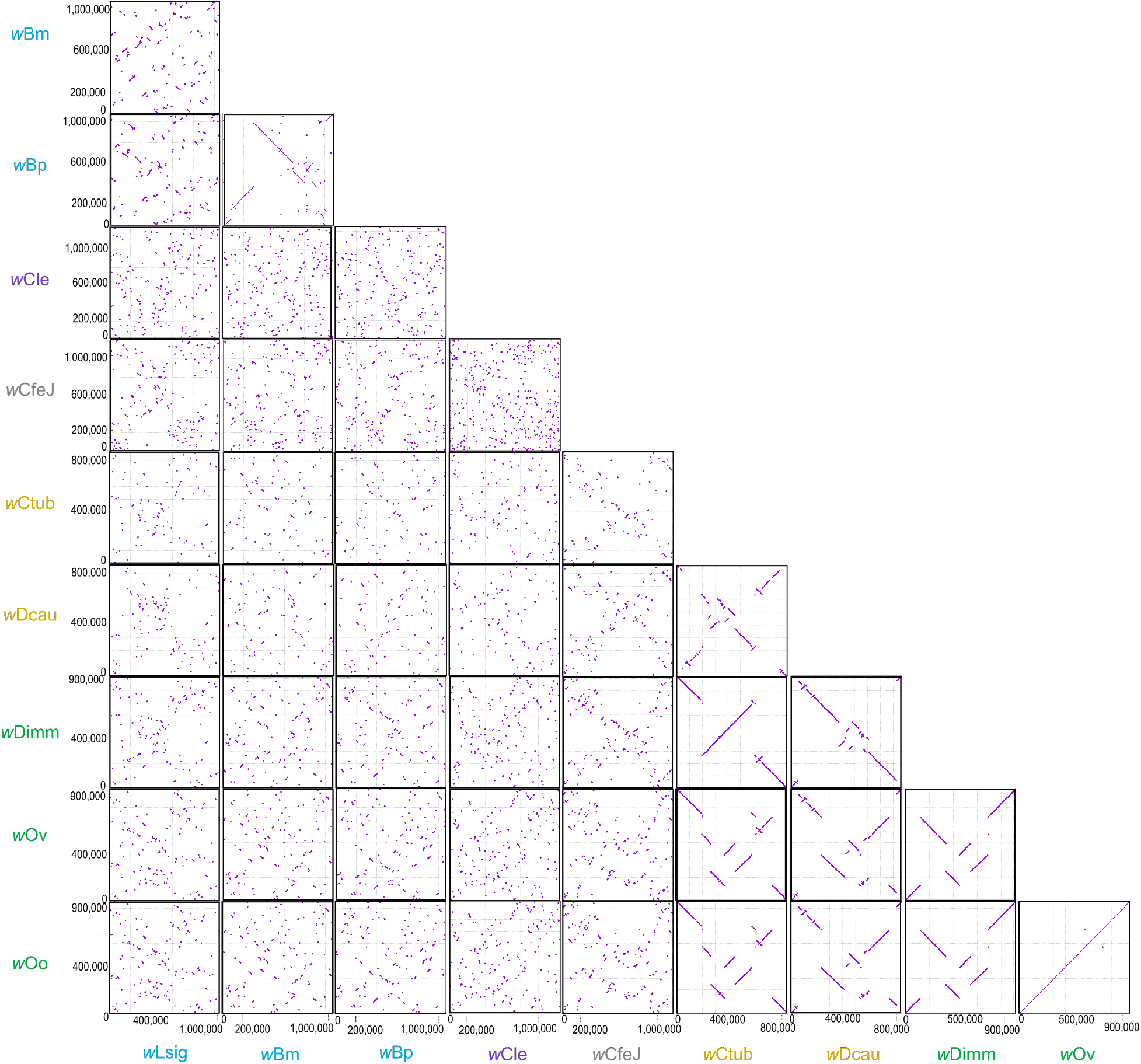
Pairwise complete genome alignment from *Wolbachia* supergroups C, D, J, F and *w*CfeJ produced by MUMmer. The *Wolbachia* supergroups are indicated by different colors: light green for C, light blue for D, purple for F, yellow for J and grey when no supergroup is assigned.

The global-fit analyses do not show a global coevolution pattern between filariae and their *Wolbachia* symbionts (PACo m^2^_XY_=0.038 with p-value=1; ParaFitGlobal= 0.0048 with p-value= 0.057; both 1e+06 permutations). The superimposition plot indicates at least five groups of associations and shows strong inequality (Figure 4A). The filarial nematodes *D.* (*D*.) *immitis* and *Onchocerca* spp. with their symbionts (supergroup C) show lower squared residuals and consequently strong coevolution. By contrast, *M. hiepei* and its symbiont (supergroup F) show high squared residual and consequently a weak coevolution (Figure 4B). The global-fit analysis confirms two different groups of association for *Wolbachia* from supergroup D and their filarial nematode hosts: on one hand, *Brugia* and *Wuchereria* species and their symbionts and on the other hand, *Litomosoides* species and their symbionts (Figure 4A). The same trend is observed for *Wolbachia* from supergroup J; the filarial nematodes *D. caudispina* and *C. tuberocauda* and their symbionts present a higher squared residual than the residual sum of squares value (m^2^_XY_) suggesting a low congruence of the phylogenies (Figure 4B). These results support the hypothesis of local patterns of co-evolution with multiple horizontal transmission events of *Wolbachia* among the filarial nematodes as part of the evolutionary history of this host-endosymbiont system, as previously described (36).

**Figure 4.**
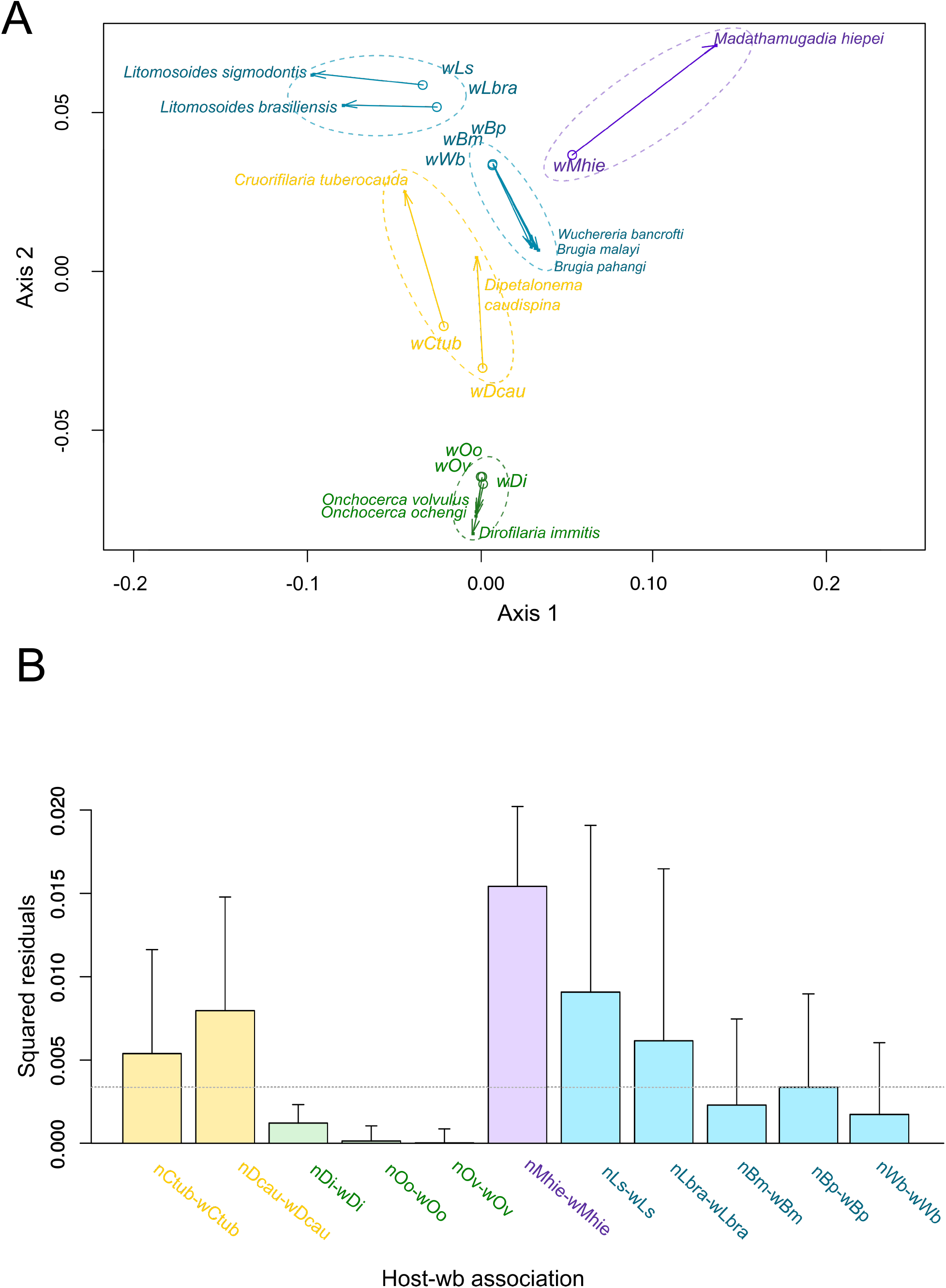
Coevolutionary analysis between filariae and *Wolbachia*. A PACo global-fit analysis of *Wolbachia* and their filarial host phylogenies was performed. (A) Representative plot of a Procrustes superimposition analysis which minimizes differences between the two partners’ principal correspondence coordinates of patristic distances. For each vector, the start point represents the configuration of *Wolbachia* and the arrowhead the configuration of filarial hosts. The vector length represents the global fit (residual sum of squares) which is inversely proportional to the topological congruence. (B) Contribution of each *Wolbachia*-filariae association to a general coevolution. Each bar represents a Jackknifed squared residual and error bars represent upper 95% confidence intervals. The *Wolbachia* supergroups are indicated by different color: light green for C, light blue for D, purple for F.

### Comparative genomics

We observed a positive correlation between *Wolbachia* genome size and the percentage of CDS. Indeed, *w*Dcau and *w*Ctub, despite having the smallest genomes, have a low percentage of CDS (71.45% and 74.63%, respectively) (Figure 5). Similarly, a positive correlation was seen between *Wolbachia* genome size and transposable elements as ISs group II intron associated genes and mobile elements (Figure 5). Interestingly, amongst the *Wolbachia* from filarial nematodes, supergroup C and supergroup J *Wolbachia* are all characterized by the absence or very low levels of transposable elements, unlike supergroup D *Wolbachia* and *w*Mhie (supergroup F) (Figure 6, Table S6, Table S7 and Table S8). We also observed a positive correlation between *Wolbachia* genome size and the amount of insertion of phage DNA, as recently described (85) (Figure 5). We studied phage DNA by two types of analyses: we used RAST annotation (68) to detect phage or phage-like genes and PHASTER (70) to detect prophage regions. None of the genomes of *Wolbachia* from filarial nematodes have significant prophage regions (Table S9) but supergroup D (*w*Bm, *w*Bp and *w*Wb), as well as the supergroup F (*w*Mhie) *Wolbachia* genomes, contain phage-like gene sequences inserted in their genomes. In the case of *w*Bm, *w*Bp and *w*Wb, mainly phage major capsid protein and some uncharacterized phage proteins were detected, representing fourteen (total 2,592 bp), eight (total 1,197 bp) and four regions (total 957 bp), respectively (Table S10). The closely related *w*Lsig and *w*Lbra, also belonging to supergroup D, do not appear to have phage protein sequences. In the case of *w*Mhie, one phage major capsid protein and eight other phage proteins were detected, representing 5,187 bp (Table S10).

**Figure 5.**
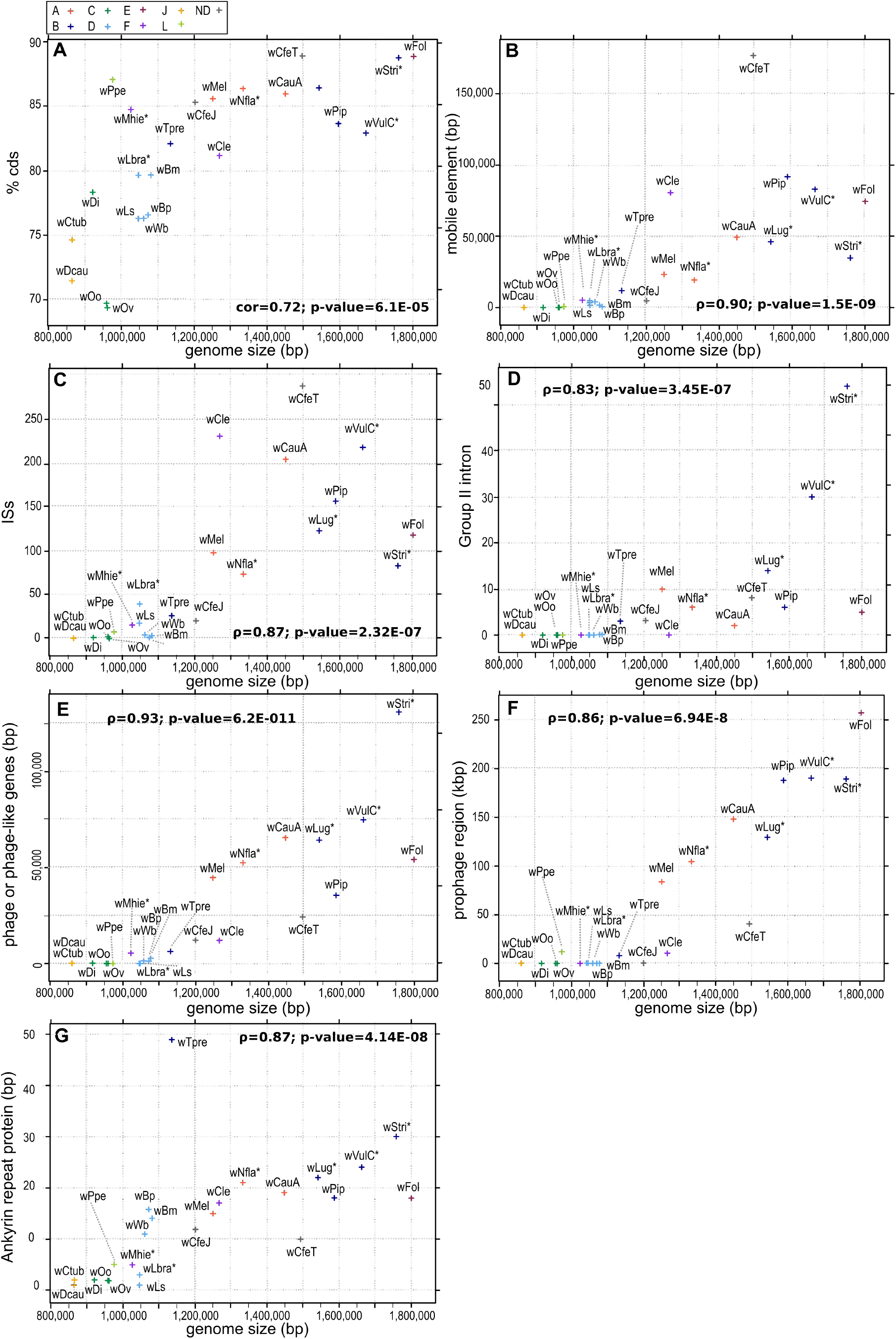
Graphic representation of the relationship between genome size of *Wolbachia* and different evolutionary factors: A) percentage of coding regions (CDS), B) regions identified as mobile elements, C) regions identified as insertion sequences (ISs), D) regions identified as group II intron-associated genes, E) regions identified as phage-like genes, F) regions identified as potential prophages and G) regions identified as ankyrin repeat. The *Wolbachia* supergroups (A–L) are indicated by different color: orange for supergroup A, dark blue for B, light green for C, light blue for D, pink for E, purple for F, yellow for J, khaki green for L, and grey when the strain is not assigned to a supergroup.

**Figure 6.**
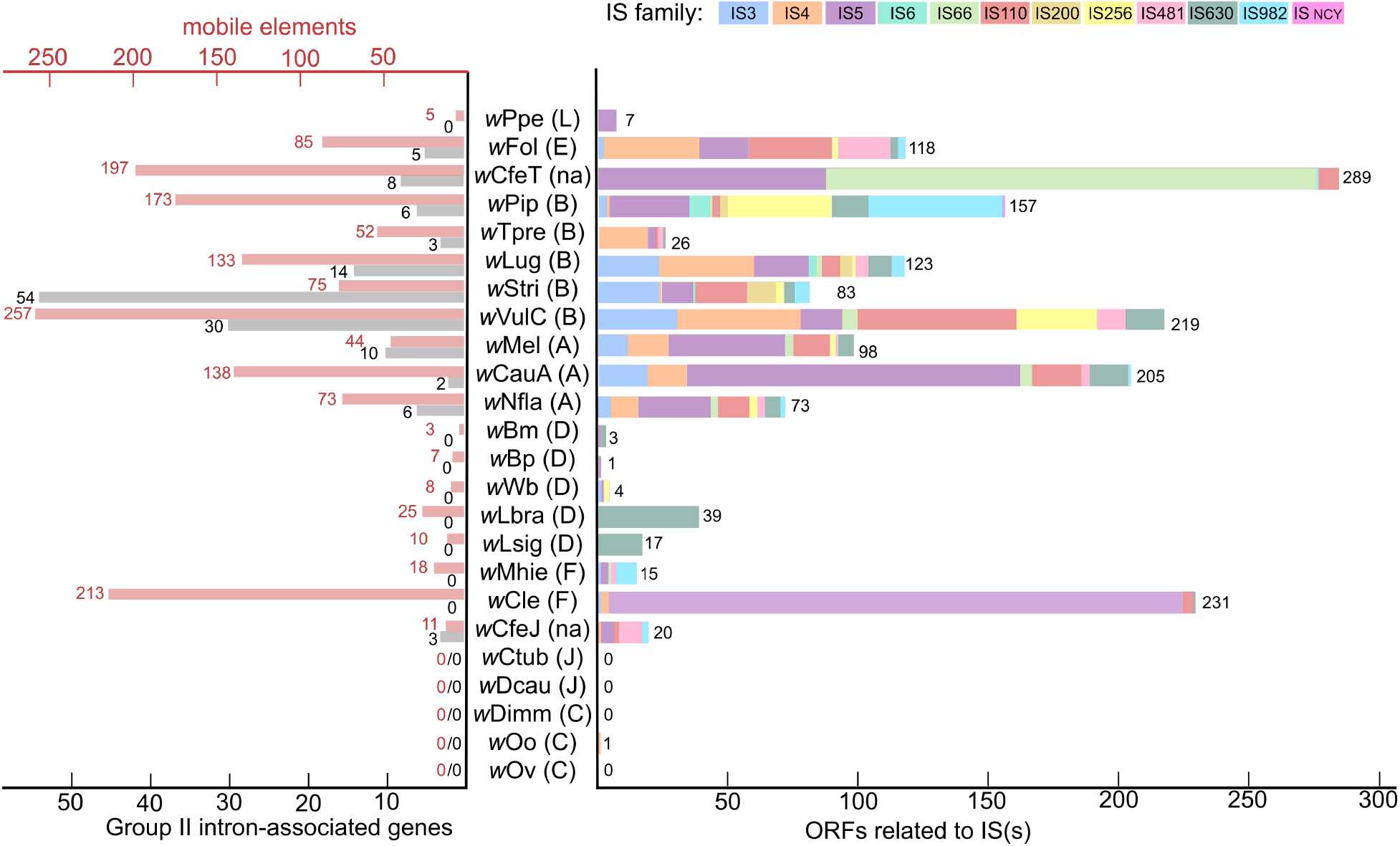
Graph of IS elements, mobile elements and group II intron-associated genes identified in *Wolbachia* genomes. The *Wolbachia* supergroups are indicated in bracket: A to L and “na” for *Wolbachia* without supergroup assignment.

A positive correlation was also observed between *Wolbachia* genome size and ankyrin repeat proteins (Figure 5). It has been suggested that *Wolbachia* from filarial nematodes are characterized by a low level of ankyrin repeat genes, suspected to have evolved as result of their mutualistic lifestyle (86–88). Most of the *Wolbachia* from filarial nematodes have 1 to 3 copies of ankyrin repeat genes, with the exception of *w*Mhie (supergroup F) with five copies and the supergroup D strains, *w*Bm, *w*Bp and *w*Wb, containing 14, 16 and 11 copies, respectively (Table S11).

### Metabolic pathways

Using KAAS, we assigned genes from the 24 studied *Wolbachia* genomes (including the seven draft genomes of *w*Nfla, *w*Lug, *w*Stri. *w*VulC. *w*Wb, *w*Lbra and *w*Mhie) to 160 different KEGG pathways (Table S2). Among these 160 KEGG pathways, 15 were selected based on strong variability among the genomes or because they had previously been suggested as being involved in symbiosis mechanisms (23, 24) (Figure 7 and TableS3).

In the context of a nutritional provisioning hypothesis, we observed variability among genomes in the vitamin B metabolism pathways (Figure 7). The thiamine metabolism (vitamin B1) pathway appears conserved, with the exception of the *tenA* gene, detected only in some *Wolbachia* genomes from arthropods: *w*Mel, *w*CauA, *w*Nfla (supergroup A); *w*Pip, *w*Lug, *w*Stri, *w*VulC (supergroup B); *w*Fol (supergroup E); *w*Cle (supergroup F); *w*CfeT and *w*CfeJ (no supergroup designated). As previously reported by Darby et al. (24), the riboflavin metabolism (vitamin B2) is incomplete for both *w*Oo and *w*Ov (supergroup C; between 3 to 4 genes not identified), but only *ribE* is missing for *w*Dimm, another representative of supergroup C. Similarly, a single gene (*ribA*) is missing for *w*Dcau and *w*Ctub (supergroup J), as is the case for *w*Mhie (supergroup F) and *w*Lbra (supergroup D). The folate (vitamin B9) and pyridoxine (vitamin B6) metabolisms appear incomplete for representatives of supergroup D, but mainly conserved for other *Wolbachia* strains (except for *w*Ppe, supergroup L, *w*Fol, supergroup E and *w*CfeT, in which only the *folC* gene is present in the folate pathway; and the absence of the *pdxJ* gene from *w*Ov in the pyridoxine pathway) (Figure 7). As described by previous authors, we note that only some *Wolbachia* have a complete biotin metabolism pathway (vitamin B7): *w*Cle (supergroup F) (17), *w*Nfla (supergroup A) (89), *w*Stri and *w*Lug (supergroup B) (18) and wCfeT (no supergroup designated) (64). In addition, we observed a complete biotin metabolism pathway in *w*VulC (supergroup B). Interestingly, it has previously been suggested that supplementation of biotin by *Wolbachia* increases the fitness of insect hosts in the case of *w*Cle from supergroup F (17), as well as *w*Stri and *w*Lug from supergroup B (18). These genes could not be detected in the newly produced supergroup F genome, *w*Mhie, from a filarial nematode host.

A further set of pathways previously considered of symbiotic interest, the *de novo* biosynthesis of purines and pyrimidines, has been identified in *Wolbachia* genomes, but was absent in other proteobacteria such as *Rickettsia* (23, 24). The pyrimidine metabolism pathway was complete for most of the *Wolbachia* genomes analyzed in the present study, with the exception of *w*Ppe (supergroup L) (Figure 7). The purine metabolism pathway was almost complete for the entire genome set as well, with the exception of the *purB* gene, which could not be identified in a large number of symbionts of filarial nematodes (*w*Lsig and *w*Lbra, supergroup D; all representatives of supergroups C and J; *w*Mhie, supergroup F) and some symbionts of insects (*w*Nfla, supergroup A; *w*Lug and *w*Stri, supergroup B; *w*Cle, supergroup F; *w*CfeJ and *w*CfeT). The *purB* gene encoding for the adenylosuccinate lyase protein is involved in the second step of the sub-pathway that synthesizes AMP from IMP.

Another important pathway, the haem metabolism, suggested to be involved in symbiotic mechanisms by several genome analyses, was complete in many of the current genomes. Only one gene, *bfr* (coding for bacterioferritin, a haem-storage protein), was not detected in *w*Bm and *w*Wb (supergroup D) or any representatives of supergroups C and J. The oxidative phosphorylation metabolism pathway also appears highly conserved, although the cytochrome bd ubiquinol oxidase genes (*cydA* and *cydB*) were detected only for *Wolbachia* belonging to supergroup A and *w*CfeJ (no supergroup assigned).

With regard to potential host interaction systems, the different secretion system pathways are very conserved (Figure 7). However, the type II secretion system gene encoding the general secretion pathway protein D (*gspD*) was neither identified in *Wolbachia* belonging to supergroups C and J, nor in *w*Bm, *w*Bp and *w*Wb (supergroup D). Similarly, the gene secE involved in the type Sec-SRP pathway was absent in wNfla (supergroup A), *w*Stri, *w*Lug (supergroup B), *w*CfeT and *w*CfeJ (no supergroups assigned).

A number of additional interesting variations among the studied *Wolbachia* genomes were noted, in particular for the cell cycle pathway, the homologous recombination (HR) pathway, the ATP binding cassette (ABC) transporter genes and glycerophospholipid metabolism (Figure 7). Regarding the cell cycle pathway, representatives of supergroup J showed losses of most cell division proteins (only *ftsQ* was detected in *w*Ctub), one gene of the two-component system (*pleD*) as well as the aspartyl protease family protein gene *perP.* The *ftsW* cell division protein gene was identified in only a few genomes: *w*Mel, *w*CauA, *w*Nfla (supergroup A); *w*Fol (supergroup E); *w*Cle, *w*Mhie (supergroup F); *w*Lbra (supergroup D); *w*CfeT and *w*CfeJ. Similarly, losses of numerous genes were detected in the HR metabolism pathway involved in repair of DNA damage. *Wolbachia* belonging to supergroup C, *w*Lsig and *w*Lbra (supergroup D) and *w*Dcau (supergroup J) showed losses of numerous genes within this set (5-9 genes) (Figure 7). We detected no recombination protein *rec* or Holliday junction DNA helicase *ruv* genes in the *w*Dcau, *w*Oo or *w*Ov genomes. Another pronounced difference observed among the studied *Wolbachia* genomes was the presence of genes encoding ATP binding cassette (ABC) transporters. These membrane transporters appear largely depleted in the *Wolbachia* genome representatives of supergroups J and C, as well as in *w*Lsig and *w*Lbra (supergroup D) (Figure 7). The haem exporter, phosphate transport system, lipoprotein-releasing system and zinc transport system appeared to be very conserved, unlike the biotin transport system, iron(III) transport system and phospholipid transport systems.

Regarding the glycerophospholipid metabolism, our results suggest that some genes are limited to a few genomes from arthropods. For example, the diacylglycerol kinase (ATP) gene (*dgkA*) was present in *w*Mel, *w*CauA, *w*Nfla (supergroup A), *w*Pip, *w*stri, *w*Lug (supergroup B), *w*VulC (closely related to supergroup B), *w*Fol (supergroup E) and wCfeT, while the phospholipase D gene (*pld*) was only detected in *w*VulC, *w*Fol, *w*CfeJ, *w*Lsig and *w*Dimm (Figure 7).

## Discussion

The LEFT-SEQ method was applied to four invertebrate DNA samples, enabling us to produce four complete *Wolbachia* genomes: *w*Lsig, *w*Dimm, *w*Dcau and *w*Ctub. For *w*Lsig and *w*Dimm, draft genomes had previously been sequenced and analyzed (26, 90) but not submitted to the NCBI database. The complete genomes of *w*Dcau and *w*Ctub are, so far, the smallest *Wolbachia* genomes. Using the enrichment method associated with Illumina sequencing (52), two draft genomes were sequenced, a 41-contig *w*Lbra draft genome and a 208-contig *w*Mhie draft genome.

Our data confirmed that *w*Lsig and *w*Lbra belong to supergroup D, *w*Dimm resides in supergroup C, *w*Mhie belongs to supergroup F, and *w*Dcau and *w*Ctub form a well-supported clade, supergroup J. Thus, *w*Dcau and *w*Ctub are now the first representative genomes of supergroup J, while *w*Mhie constitutes the first representative genome of supergroup F from a filarial nematode.

The ANI and dDDH index indicate that *w*Dcau, *w*Ctub, *w*Dimm, *w*Lsig and *w*Lbra are clearly divergent from other studied *Wolbachia* genomes (all ≤90% ANI and <70% dDDH) (Figure 1). Regarding *w*Mhie, the ANI is 95% with *w*Cle, suggesting a close genetic proximity, although the dDDH is equal to 58.12 (Model-based confidence intervals: 76 – 82.5%), below the threshold of 70. Our data suggest that these two *Wolbachia* are very similar, despite the fact that one infects a filarial nematode and the other infects bedbugs.

The analysis of supergroup J *Wolbachia* further highlights limitations of the current MLST system. In the past, the only representative identified from this supergroup was the symbiont of *D. gracile*, a filarial nematode parasite of monkeys. This symbiont was first described as a deep branch within supergroup C (44), and subsequently as the divergent clade J (39, 84). The latter phylogenetic position of this *Wolbachia* has been questioned by some authors and is often retained as belonging to supergroup C (35, 82, 83). More recently, using a concatenation of seven genes and newly studied *Wolbachia*, Lefoulon et al. (36), demonstrated the validity of supergroup J, as distinct from supergroup C; a phylogenetic position confirmed by the present study (Figure 2). Our analyses show that the *ftsZ* gene is not present (or is highly degenerate) in the *w*Dcau and *w*Ctub genomes, while previous *Wolbachia* phylogenies have been based on this marker. Our analyses (Table S5) suggest that the variable position of *Wolbachia* from *D. gracile* in some phylogenetic analyses is linked to the fact that some database sequences likely do not belong to this strain.

The two complete genomes, *w*Dcau and *w*Ctub, are divergent from supergroup C *Wolbachia*. In addition, they are highly divergent from each other, with an ANI of 81%, despite the fact they form their own clade (Figure 1). These divergences have been suggested by earlier multi-locus phylogenies with *Wolbachia* from *Cruorifilaria* and *Yatesia* species forming one subgroup and *Wolbachia* from *Dipetalonema* spp. forming another subgroup within supergroup J (36). Our data suggest that the use of the *ftsZ* gene for MLST studies is not appropriate for *Wolbachia* that are highly divergent. The fact that the MLST system was designed on the basis of supergroups A and B *Wolbachia* (48), which have low genetic diversity (30), is a source of concern for its general use when studying divergent phylogenies. Moreover, the risk of erroneous data finding their way into databases (e.g. through contamination, misidentification), combined with the fact that sequences used to build concatenated matrices very often do not originate from the same specimen, weakens multi-locus phylogenies, unless potential confounding factors are taken into consideration.

The symbiosis between *Wolbachia* and filarial nematodes was often considered and analyzed as a uniform pattern of association, but our results reveal strong disparities. Indeed, the genomes of supergroup J present a strong synteny pattern, as was previously described for representatives of supergroup C, unlike those of supergroup D (26) (Figure 3). We even observe a strong synteny pattern between *w*Dimm (supergroup C) and *w*Ctub or *w*Dcau (supergroup J). Interestingly, the smaller *Wolbachia* genomes present a low number of genomic rearrangements, associated with the absence or low number of transposable elements (either ISs, mobile elements or group II introns) (Figure 6). Our data support the paradigm that a major difference between *Wolbachia* from filarial nematodes and those from arthropods is a reduced genome containing fewer (or even zero) transposable elements, prophage-related genes or repeat-motif proteins (as ankyrin domains) (24). Further, our results highlight the distinction between supergroups C and J *Wolbachia* versus the supergroup D and F *Wolbachia*. In addition to having larger genomes, more transposable elements were identified in these genomes: in supergroups D and F, *w*Lbra, *w*Wb and *w*Mhie contain more mobile elements; and *w*Lsig, *w*Lbra and *w*Mhie more ISs (Figure 6). Traditionally, studies of genome reduction in the cases of symbiotic bacteria indicate an expansion of mobile genetic elements in the initial stages of bacterial adaptation to a host-dependent lifestyle and an absence of mobile genetic elements in long-term obligate symbiosis associations (91). This suggests that the different associations of *Wolbachia*-filarial nematodes represent different stages of host-dependent adaptation. Initially, it had been suggested that *Wolbachia* symbionts coevolved with their filarial nematodes (4). Supergroup F *Wolbachia* were thought to be the only example of horizontal transfer among the filarial nematodes (34, 46). A recent revision of the co-phylogenetic patterns of *Wolbachia* in filariae based on multi-locus phylogenies suggests that only supergroup C *Wolbachia* exhibit strong co-speciation with their hosts (36). Indeed, our global-fit analyses are not compatible with a global pattern of coevolution, but rather support the hypothesis of two independent acquisitions of supergroup D and J (Figure 4). These results highlight a differential evolution of *Wolbachia* symbiosis among the various filarial nematodes, likely having evolved from different acquisitions and subject to different selective pressures.

Another important aspect of *Wolbachia* diversity is the association between some *Wolbachia* and the WO bacteriophage (92–95). Indeed, prophage regions have been identified in numerous *Wolbachia* genomes and the fact that these insertions have not been eliminated by selective pressure support the hypothesis that they could provide factors of importance to *Wolbachia* (93, 96). In the case of *Wolbachia* from arthropods, these insertions can constitute a large proportion of the *Wolbachia* genome. For example, it was recently shown that 25.4% of the *w*Fol genome comprises five phage WO regions (85). Our analyses indicate that the large-sized genomes, such as *w*Stri or *w*VulC, have large regions of WO prophage (Figure 5, Table S9). No intact region or only vestiges of prophage regions had been observed in previously studied *Wolbachia* genomes infecting filarial nematodes (23, 24). Our data support this absence of prophages in the newly studied genomes in this report. However, we detected some genes annotated as phage-like in the cases of *w*Bm, *w*Bp and *w*Wb (supergroup D) and *w*Mhie (supergroup F), unlike other representatives of supergroup D (*w*Lsig or *w*Lbra), and the genomes belonging to supergroups C and J *Wolbachia* (Table S10). Interestingly, while the bedbug symbiont *w*Cle (supergroup F) has fewer phage genes than other *Wolbachia* from arthropods, numerous phage elements have been found in *Wolbachia* sequences integrated into the nuclear genome of a strongyloidean nematode (*Dictyocaulus viviparus*) (83), which were allocated to supergroup F, suggesting significant variation in the role of phage WO within this clade. So far, *w*Wb, *w*Bm and *w*Bp have the largest *Wolbachia* genomes of filarial nematodes and while these phage-like insertions represent a negligible proportion of the entire genome, they nevertheless suggest that *w*Wb, *w*Bm and *w*Bp were in contact with bacteriophages which successfully inserted DNA in their respective genomes. At the same time, our study shows that numerous genes involved in HR and the cell cycle pathway (Figure 7) are absent in the *Wolbachia* from filarial nematodes other than *w*Bm, *w*Bp and *w*Wb, and thus insertion of DNA might not be possible for the bacteriophages due to the nature of these genomes themselves.

Supergroup F is particularly interesting as it represents the only clade composed of both *Wolbachia* symbionts of arthropods and filarial nematodes, suggesting horizontal transfer between the two phyla (84). Previous studies suggested it is more likely that the infection by supergroup F *Wolbachia* derived from multiple independent host switch events in the Filarioidea because they infected species that are not closely related (34, 36). In addition, recent phylogenomic studies suggest that supergroup F is a derived clade in the evolutionary history of *Wolbachia* (3, 17, 97). The *w*Mhie genome belonging to the supergroup F is closely related to the bedbug symbiont *w*Cle; however, the characteristics of the genome (small size, few transposase elements, few phage genes, absence of prophage region) are more similar to those observed in representatives of supergroup D.

Previous genomics studies of *Wolbachia* from filarial nematodes have hypothesized mechanisms which could underpin the obligate mutualism (23, 24). Our data indicate that both haem and nucleotide (pyrimidine and purine) metabolism are particularly conserved among all analyzed *Wolbachia* genomes, even the smallest ones, and thus support suggestions of potential provisioning of these resources by *Wolbachia* (Figure 7). The hypothesis of mutualism based on nutritional provisioning has been revised after the detection of the incomplete riboflavin (vitamin B2) pathway in *w*Oo (24). Notably, the genomes of supergroup D show an incomplete folate metabolism pathway (vitamin B9), which is complete for the small genomes of both supergroup C and J. The riboflavin pathway (vitamin B2) appears incomplete in supergroup C but almost complete for supergroup J. Another interesting vitamin B pathway is the biotin operon (vitamin B7). It was previously suggested that the evolution of this operon is not congruent with proposed *Wolbachia* evolutionary history (89). Our data show the operon is present in *w*Nfla (supergroup A), *w*stri, *w*Lug, *w*VulC (supergroup B), *w*Cle (supergroup F) and *w*CfeJ (not belonging to a described supergroup), and that it might have been acquired horizontally as a nutritional requirement. For *w*Cle, *w*stri and *w*Nfla, biotin supplementation by *Wolbachia* increases insect host fitness (17, 18). Interestingly, our study shows that incomplete metabolic pathways are not a function of *Wolbachia* genome size. Overall, the pathway analysis presented in the current study suggests that no single metabolic process governs the entire spectrum of *Wolbachia*-filarial nematode associations.

It is highly likely that such provisioning mechanisms might differ according to the particular host-symbiont association, although in cases where the *Wolbachia* host is itself a parasite (such as filarial nematodes), the potential metabolic interactions with mammalian and arthropod hosts of the filariae are highly complex. Further genomic analyses will highlight and unravel these divergent symbiotic mechanisms.

## Supporting information

two supplemental files and eleven supplementary table

## Author statements

### Author Contributions Statement

E.L., C.M., B.L.M., A.C.D., J.M.F. and B.E.S. conceived and designed the experiments. E.L., T.C., B.E.S. performed the experiments. E.L. analyzed the data. R.G., I.C., J.M.C-C., K.J., N.V-L. and C.M. provided study materials. E.L, B.E.S, J.M.F, B.L.M. and K.J. wrote the main manuscript text. All authors reviewed the manuscript.

### Ethical approval

Most of the samples were collected as described in a previous study (53) and all procedures were conducted in compliance with the rules and regulations of the respective national ethical bodies. Regarding the newly studied material: the *D.* (*D*.) *immitis* specimen was provided by the NIAID/NIH Filariasis Research Reagent Resource Center (MTA University of Wisconsin Oshkosh; www.filariasiscenter.org), and the *L. sigmodontis* specimen was provided by the National Museum of Natural History (MNHN, Paris), where the experimental procedures were carried out in strict accordance with the EU Directive 2010/63/UE and the relevant national legislation (ethical statement n°13845).

### Consent for publication

Not applicable, no consent form required

## Acknowledgments

We thank Andy Gardner, Clotilde Carlow, Tom Evans, Rich Roberts, Jim Ellard and Don Comb from New England Biolabs for their support. We also thank Laurie Mazzola, Danielle Fuchs and Kristen Augulewicz from the NEB Sequencing Core.

## Figure Legends

Figure 7. Summary of the metabolic pathways detected in different *Wolbachia* genomes using KASS. The *Wolbachia* supergroups (A-L) are indicated by different color: orange for supergroup A, dark blue for B, light green for C, light blue for D, pink for E, purple for F, yellow for J, khaki green for L, and grey when the strain is not described belonging to a supergroup.

## Abbreviations

CI: cytoplasmic incompatibility
MLST: multi-locus sequence typing
ng: nanogram
bp: DNA nucleotide base pair
CDS: Coding region of genes sequence
CCS: circular consensus sequences
IS(s): insertion sequence element(s)
PCR: polymerase chain reaction
dDDH: digital DNA-DNA hybridization
ANI: Average Nucleotide Identity

