## Supplementary material for "Diminutive, degraded but dissimilar: *Wolbachia* genomes from filarial nematodes do not conform to a single paradigm": two supplemental files and eleven supplementary table: Suppl_Table_S1_S11.pdf

**Table S1. List of Wolbachia genomes used in the study with their strain na**
The produced genome are indicated in blue. The genomes included in compara

| ***Wolbachia*** | **strain** |
|---|---|
| *Wolbachia* from *Aedes albopictus* | wAlbB |
| *Wolbachia from Atemnus politus* | wApol K5 |
| *Wolbachia* from *Drosophila simulans wAu* | wAu |
| *Wolbachia* from *Plutella australiana wAus* | wAus |
| ***Wolbachia* from *Brugia malayi*** | **wBm** |
| ***Wolbachia* from *Brugia pahangi*** | **wBp** |
| *Wolbachia* from *Hypolimnas bolina* | wBol1b |
| *Wolbachia* from *Bemisia tabaci* | wBtab |
| ***Wolbachia* from *Carposina sasakii*** | **wCauA** |
| ***Wolbachia* from *Ctenocephalides felis*** | **wCfeJ** |
| ***Wolbachia* from *Ctenocephalides felis*** | **wCfeT** |
| ***Wolbachia* from *Cimex lectularius*** | **wCle** |
| *Wolbachia* from *Cylisticus convexus* | wcon |
| ***Wolbachia* from *Cruorifilaria tuberocauda*** | **wCtub** |
| *Wolbachia* from *Dactylopius coccus* | wDacB |
| ***Wolbachia* from *Dipetalonema caudispina*** | **wDcau** |
| ***Wolbachia* from *Dirofilaria (Dirofilaria) immitis*** | **wDimm** |
| ***Wolbachia* from *Folsomia candida*** | **wFol Berlin** |
| *Wolbachia* from *Glossina morsitans morsitans* | wGmm |
| *Wolbachia* from *Drosophila simulans* | wHa |
| *Wolbachia* from *Drosophila incompta* | wIncCu |
| ***Wolbachia* from *Litomosoides brasiliensis*** | **wLbra** |
| *Wolbachia* from *Leptopilina clavipes* | wLcla |
| ***Wolbachia* from *Litomosoides sigmondontis*** | **wLsig** |
| ***Wolbachia* from *Nilaparvata lugens*** | **wLug** |
| *Wolbachia* from *Drosophila mauritiana* | wMau |
| *Wolbachia* from *Chrysomya megacephala* | wMeg |
| ***Wolbachia* from *Drosophila melanogaster*** | **wMel** |
| ***Wolbachia* from *Madathamugadia hiepei*** | **wMhie** |
| *Wolbachia* from *Nomada ferruginata* | wNfe |
| ***Wolbachia* from *Nomada flava*** | **wNfla** |
| *Wolbachia* from *Nomada leucophthalma* | wNleu |
| *Wolbachia* from *Drosophila simulans* | wNo |
| *Wolbachia* from *Nomada panzeri* | wNpa |
| ***Wolbachia* from *Onchocerca ochengi*** | **wOo** |
| ***Wolbachia* from *Onchocerca volvulus*** | **wOv Cameroo** |
| ***Wolbachia* from *Culex quinquefasciatus*** | **wPip** |

| | |
|---|---|
| ***Wolbachia* from *Pratylenchus penetrans*** | **wPpe** |
| *Wolbachia* from *Drosophila simulans wRi* | wRi |
| *Wolbachia* from *Drosphila santomea* | wsan |
| ***Wolbachia* from *Laodelphax striatella*** | **wstri** |
| *Wolbachia* from *Drosophila teissieri* | wTei |
| ***Wolbachia* from *Trichogramma pretiosum*** | **wTpre** |
| *Wolbachia* from *Muscidifurax uniraptor* | wUni |
| *Wolbachia* from *Nasonia vitripennis* | wVitA |
| *Wolbachia* from *Nasonia vitripennis* | wVitB |
| ***Wolbachia* from *Armadillidium vulgare*** | **wVulC** |
| ***Wolbachia* from *Wuchereria bancrofti*** | **wWb** |
| *Wolbachia* from *Drosophila yakuba* | wYak |

**ıme, NCBI accession number and detail of the molecular analyses.**
tive analyses are indicated in bold.

| complete genome | phylogenies 365 locus | 160 locus |
|---|---|---|
| CP031221 | yes | yes |
| WQMQ00000000 | no | yes |
| LK055284 | yes | yes |
| MRWX00000000 | no | yes |
| **NC_006833** | **yes** | **yes** |
| **CP050521** | **yes** | **yes** |
| CAOH00000000 | no | yes |
| CP016430 | yes | yes |
| **CP041215** | **yes** | **yes** |
| **CP051157** | **yes** | **yes** |
| **CP051156** | **yes** | **yes** |
| **AP013028** | **yes** | **yes** |
| QPIP00000000 | no | yes |
| **CP046579** | **yes** | **yes** |
| LSYY00000000 | no | yes |
| **CP046580** | **yes** | **yes** |
| **CP046578** | **yes** | **yes** |
| **CP015510** | **yes** | **yes** |
| AWUH00000000 | no | yes |
| NC_021089 | yes | yes |
| CP011148 | yes | yes |
| **WQMO00000000** | **no** | **yes** |
| QJHA00000000 | no | yes |
| **CP046577** | **yes** | **yes** |
| **MUIY01000000** | **no** | **yes** |
| CP034334 | yes | yes |
| CP021120 | yes | yes |
| **NC_002978** | **yes** | **yes** |
| **WQMP00000000** | **no** | **yes** |
| LYUY00000000 | no | yes |
| **LYUW00000000** | **no** | **yes** |
| LYUV00000000 | no | yes |
| NC_021084 | yes | yes |
| LYUX00000000 | no | yes |
| **NC_018267** | **yes** | **yes** |
| **HG810405** | **yes** | **yes** |
| **NC_010981** | **yes** | **yes** |

| | | |
|---|---|---|
| **MJMG01000000** | **no** | **yes** |
| NC_012416 | yes | yes |
| VCEH00000000 | no | yes |
| **MUIX00000000** | **no** | **yes** |
| VCEG01000000 | no | yes |
| **CM003641** | **yes** | **yes** |
| MUJL00000000 | no | yes |
| MUJM00000000 | no | yes |
| AERW00000000 | no | yes |
| **ALWU00000000** | **no** | **yes** |
| **NJBR00000000** | **no** | **yes** |
| VCEF00000000 | no | yes |

**Table S2. Summary of number of genes assigned to 160 different KEGG pathways fro**

| KO | KEGG pathways | wPpe | wFol | wCfeT | wPip | wLug | wstri |
|---|---|---|---|---|---|---|---|
| K00010 | Glycolysis / Gluconeogenesis | 11 | 11 | 11 | 11 | 11 | 11 |
| K00020 | Citrate cycle (TCA cycle) | 16 | 17 | 16 | 17 | 15 | 17 |
| K00030 | Pentose phosphate pathway | 6 | 6 | 6 | 6 | 6 | 6 |
| K00040 | Pentose and glucuronate interconversions | 1 | 2 | 1 | 1 | 2 | 2 |
| K00051 | Fructose and mannose metabolism | 4 | 5 | 5 | 5 | 5 | 5 |
| K00053 | Ascorbate and aldarate metabolism | - | 1 | - | - | 1 | 1 |
| K00520 | Amino sugar and nucleotide sugar metabolism | 5 | 7 | 6 | 6 | 7 | 7 |
| K00620 | Pyruvate metabolism | 8 | 8 | 8 | 8 | 8 | 8 |
| K00630 | Glyoxylate and dicarboxylate metabolism | 8 | 9 | 9 | 9 | 9 | 9 |
| K00640 | Propanoate metabolism | 5 | 6 | 6 | 6 | 6 | 6 |
| K00650 | Butanoate metabolism | 4 | 4 | 3 | 4 | 3 | 4 |
| K00660 | C5-Branched dibasic acid metabolism | 2 | 2 | 2 | 2 | 2 | 2 |
| K00562 | Inositol phosphate metabolism | 2 | 2 | 2 | 2 | 2 | 2 |
| **K00190** | **Oxidative phosphorylation** | **35** | **37** | **35** | **37** | 36 | 36 |
| K00195 | Photosynthesis | 7 | 8 | 8 | 8 | 8 | 8 |
| K00710 | Carbon fixation in photosynthetic organism | 10 | 10 | 10 | 10 | 10 | 10 |
| K00720 | Carbon fixation pathways in prokaryotes | 11 | 12 | 11 | 12 | 11 | 12 |
| K00680 | Methane metabolism | 6 | 6 | 6 | 6 | 6 | 6 |
| K00910 | Nitrogen metabolism | 2 | 2 | 2 | 2 | 2 | 2 |
| **K00061** | **Fatty acid biosynthesis** | **6** | **8** | **8** | **8** | 8 | 8 |
| K00561 | Glycerolipid metabolism | 3 | 4 | 4 | 4 | 4 | 4 |
| **K00564** | **Glycerophospholipid metabolism** | **9** | **11** | **9** | **10** | 10 | 10 |
| **K00230** | **Purine metabolism** | **22** | **21** | **21** | **22** | 21 | 21 |
| **K00240** | **Pyrimidine metabolism** | **16** | **17** | **17** | **17** | 17 | 17 |
| K00250 | Alanine, aspartate and glutamate metabolism | 11 | 12 | 11 | 12 | 11 | 11 |
| K00260 | Glycine, serine and threonine metabolism | 7 | 8 | 7 | 7 | 8 | 8 |
| K00270 | Cysteine and methionine metabolism | 7 | 8 | 8 | 8 | 8 | 8 |
| K00280 | Valine, leucine and isoleucine degradation | 2 | 3 | 3 | 3 | 3 | 3 |
| K00300 | Lysine biosynthesis | 10 | 9 | 10 | 10 | 10 | 10 |
| K00310 | Lysine degradation | 2 | 2 | 2 | 2 | 1 | 2 |
| K00220 | Arginine biosynthesis | 6 | 6 | 6 | 6 | 6 | 6 |
| K00330 | Arginine and proline metabolism | 2 | 2 | 2 | 2 | 2 | 2 |
| K00350 | Tyrosine metabolism | 1 | 1 | 1 | 1 | 1 | 1 |
| K00360 | Phenylalanine metabolism | 1 | 1 | 1 | 1 | 1 | 1 |
| K00380 | Tryptophan metabolism | 2 | 2 | 2 | 2 | 1 | 2 |
| K00400 | Phenylalanine, tyrosine and tryptophan biosynthe | 1 | 1 | 1 | 1 | 1 | 1 |
| K00410 | beta-Alanine metabolism | 1 | 1 | 2 | 1 | 1 | 1 |
| K00430 | Taurine and hypotaurine metabolism | 1 | 1 | 1 | 1 | 1 | 1 |

| | | | | | | | |
|---|---|---|---|---|---|---|---|
| K00450 | Selenocompound metabolism | 3 | 3 | 3 | 3 | 3 | 3 |
| K00460 | Cyanoamino acid metabolism | 1 | 1 | 1 | 1 | 1 | 1 |
| K00471 | D-Glutamine and D-glutamate metabolism | 3 | 3 | 3 | 3 | 3 | 3 |
| K00473 | D-Alanine metabolism | 1 | 1 | 1 | 1 | 1 | 1 |
| K00480 | Glutathione metabolism | 2 | 5 | 5 | 5 | 5 | 5 |
| K00550 | Peptidoglycan biosynthesis | 10 | 11 | 13 | 11 | 11 | 11 |
| **K00730** | **Thiamine metabolism** | 2 | 3 | 9 | 4 | 4 | 4 |
| **K00740** | **Riboflavin metabolism** | **1** | **6** | **6** | **6** | 6 | 6 |
| **K00750** | **Vitamin B6 metabolism** | **2** | **2** | - | **2** | 2 | 2 |
| K00760 | Nicotinate and nicotinamide metabolism | 2 | 2 | 2 | 2 | 2 | 2 |
| K00770 | Pantothenate and CoA biosynthesis | 3 | 2 | **5** | 3 | 3 | 3 |
| **K00780** | **Biotin metabolism** | **4** | **5** | 9 | **5** | 10 | 10 |
| K00785 | Lipoic acid metabolism | 2 | 2 | 2 | 2 | 2 | 2 |
| **K00790** | **Folate biosynthesis** | **2** | **2** | **2** | **4** | 5 | 5 |
| K00670 | One carbon pool by folate | 7 | 6 | 7 | 7 | 7 | 7 |
| **K00860** | **Porphyrin and chlorophyll metabolism** | **11** | **11** | **11** | **11** | 11 | 11 |
| K00130 | Ubiquinone and other terpenoid-quinone biosynt | 7 | 7 | 7 | 7 | 7 | 7 |
| K00900 | Terpenoid backbone biosynthesis | 8 | 8 | 8 | 8 | 8 | 8 |
| K00908 | Zeatin biosynthesis | 1 | 1 | 1 | 1 | 1 | 1 |
| K01051 | Biosynthesis of ansamycins | 1 | 1 | 1 | 1 | 1 | 1 |
| K00940 | Phenylpropanoid biosynthesis | 1 | 1 | 1 | 1 | 1 | 1 |
| K00950 | Isoquinoline alkaloid biosynthesis | 1 | 1 | 1 | 1 | 1 | 1 |
| K00960 | Tropane, piperidine and pyridine alkaloid biosynth | 1 | 1 | 1 | 1 | 1 | 1 |
| K00261 | Monobactam biosynthesis | 4 | 4 | 4 | 4 | 4 | 4 |
| K00521 | Streptomycin biosynthesis | 1 | 1 | 1 | 1 | 2 | 2 |
| K00401 | Novobiocin biosynthesis | 1 | 1 | 1 | 1 | 1 | 1 |
| K00333 | Prodigiosin biosynthesis | 3 | 3 | 3 | 3 | 3 | 3 |
| K00998 | Biosynthesis of secondary metabolites - other ant | 1 | 1 | 1 | 1 | 1 | 1 |
| K00362 | Benzoate degradation | 1 | 1 | 1 | - | 1 | 1 |
| K00627 | Aminobenzoate degradation | 1 | 1 | 1 | 1 | 1 | 1 |
| K00624 | Polycyclic aromatic hydrocarbon degradation | 1 | 1 | 1 | 1 | 1 | 1 |
| K00980 | Metabolism of xenobiotics by cytochrome P450 | 1 | 1 | 1 | 1 | 1 | 1 |
| K00982 | Drug metabolism - cytochrome P450 | 1 | 1 | 1 | 1 | 1 | 1 |
| K00983 | Drug metabolism - other enzymes | 4 | 5 | 5 | 5 | 5 | 5 |
| K03020 | RNA polymerase | 3 | 3 | 4 | 3 | 3 | 3 |
| K03010 | Ribosome | 51 | 53 | 51 | 53 | 53 | 53 |
| K00970 | Aminoacyl-tRNA biosynthesis | 22 | 24 | 24 | 24 | 24 | 24 |
| K03008 | Ribosome biogenesis in eukaryotes | - | 1 | 1 | 1 | 1 | 1 |
| K03060 | Protein export | 15 | 15 | 14 | 15 | 14 | 14 |
| K04141 | Protein processing in endoplasmic reticulum | 2 | 1 | 2 | 2 | 2 | 2 |
| K04122 | Sulfur relay system | 2 | 2 | 3 | 2 | 2 | 2 |
| K03018 | RNA degradation | 8 | 9 | 10 | 8 | 9 | 9 |

| | | | | | | | |
|---|---|---|---|---|---|---|---|
| K03030 | DNA replication | 13 | 13 | 13 | 13 | 13 | 13 |
| K03410 | Base excision repair | 8 | 7 | 8 | 8 | 8 | 7 |
| K03420 | Nucleotide excision repair | 3 | 6 | 6 | 6 | 6 | 6 |
| K03430 | Mismatch repair | 12 | 12 | 12 | 12 | 12 | 12 |
| **K03440** | **Homologous recombination** | **17** | **17** | **18** | **17** | 18 | 18 |
| **K02010** | **ABC transporters** | **20** | **20** | **20** | **20** | 20 | 20 |
| **K03070** | **Bacterial secretion system** | **22** | **23** | **22** | **23** | 22 | 22 |
| K02020 | Two-component system | 10 | 12 | 14 | 12 | 12 | 12 |
| K04013 | MAPK signaling pathway - fly | 1 | 1 | 1 | 1 | 1 | 1 |
| K04016 | MAPK signaling pathway - plant | 1 | 1 | 1 | 1 | 1 | 1 |
| K04066 | HIF-1 signaling pathway | 5 | 5 | 5 | 5 | 5 | 5 |
| K04068 | FoxO signaling pathway | 1 | 1 | 1 | 1 | 1 | 1 |
| K04070 | Phosphatidylinositol signaling system | 2 | 3 | 3 | 3 | 3 | 3 |
| K04151 | PI3K-Akt signaling pathway | 1 | 1 | 1 | 1 | 1 | 1 |
| K04152 | AMPK signaling pathway | 1 | 1 | 1 | 1 | 1 | 1 |
| K04146 | Peroxisome | 3 | 3 | 3 | 3 | 3 | 3 |
| **K04112** | **Cell cycle - Caulobacter** | **11** | **14** | 16 | **13** | 13 | 13 |
| K04210 | Apoptosis | - | 1 | 1 | 1 | 1 | 1 |
| K04214 | Apoptosis - fly | 2 | 2 | 2 | 2 | 2 | 2 |
| K04215 | Apoptosis - multiple species | - | 1 | 1 | 1 | 1 | 1 |
| K04217 | Necroptosis | 2 | 2 | 2 | 2 | 2 | 2 |
| K04115 | p53 signaling pathway | - | 1 | 1 | 1 | 1 | 1 |
| K02024 | Quorum sensing | 9 | 11 | 11 | 11 | 10 | 10 |
| K02026 | Biofilm formation - Escherichia coli | 1 | 1 | 1 | 1 | 1 | 1 |
| K04621 | NOD-like receptor signaling pathway | 2 | 2 | 2 | 2 | 1 | 1 |
| K04612 | Antigen processing and presentation | 1 | 1 | 1 | 1 | 2 | 2 |
| K04659 | Th17 cell differentiation | 1 | 1 | 1 | 1 | 1 | 1 |
| K04657 | IL-17 signaling pathway | 1 | 1 | 1 | 1 | 1 | 1 |
| K04922 | Glucagon signaling pathway | 2 | 2 | 2 | 2 | 1 | 1 |
| K03320 | PPAR signaling pathway | 1 | 1 | 1 | 1 | 2 | 2 |
| K04915 | Estrogen signaling pathway | 1 | 1 | 1 | 1 | 1 | 1 |
| K04914 | Progesterone-mediated oocyte maturation | 1 | 1 | 1 | 1 | 1 | 1 |
| K04260 | Cardiac muscle contraction | 3 | 3 | 3 | 3 | 1 | 1 |
| K04724 | Glutamatergic synapse | 1 | 1 | 1 | 1 | 3 | 3 |
| K04727 | GABAergic synapse | 1 | 1 | 1 | 1 | 1 | 1 |
| K04211 | Longevity regulating pathway | 1 | 1 | 1 | 1 | 1 | 1 |
| K04212 | Longevity regulating pathway - worm | 3 | 5 | 5 | 5 | 1 | 1 |
| K04213 | Longevity regulating pathway - multiple species | 2 | 1 | 2 | 2 | 5 | 5 |
| K04714 | Thermogenesis | 6 | 6 | 6 | 6 | 2 | 2 |
| K04626 | Plant-pathogen interaction | 2 | 2 | 3 | 2 | 6 | 6 |
| K05200 | Pathways in cancer | 2 | 4 | 4 | 4 | 3 | 3 |
| K05205 | Proteoglycans in cancer | - | 1 | 1 | 1 | 4 | 4 |

| | | | | | | | |
|---|---|---|---|---|---|---|---|
| K05204 | Chemical carcinogenesis | - | 1 | 1 | 1 | 1 | 1 |
| K05230 | Central carbon metabolism in cancer | 2 | 2 | 2 | 2 | 1 | 1 |
| K05231 | Choline metabolism in cancer | - | 1 | 1 | 1 | 2 | 2 |
| K05210 | Colorectal cancer | - | 1 | 1 | 1 | 1 | 1 |
| K05225 | Hepatocellular carcinoma | - | 1 | 1 | 1 | 1 | 1 |
| K05211 | Renal cell carcinoma | 1 | 1 | 1 | 1 | 1 | 1 |
| K05215 | Prostate cancer | 1 | 1 | 1 | 1 | 1 | 1 |
| K05222 | Small cell lung cancer | - | 1 | 1 | 1 | 1 | 1 |
| K05010 | Alzheimer disease | 4 | 5 | 5 | 5 | 5 | 5 |
| K05012 | Parkinson disease | 3 | 4 | 4 | 4 | 4 | 4 |
| K05014 | Amyotrophic lateral sclerosis (ALS) | - | 1 | 1 | 1 | 1 | 1 |
| K05016 | Huntington disease | 4 | 5 | 5 | 5 | 5 | 5 |
| K05418 | Fluid shear stress and atherosclerosis | 2 | 3 | 3 | 3 | 3 | 3 |
| K05416 | Viral myocarditis | - | 1 | 1 | 1 | 1 | 1 |
| K04940 | Type I diabetes mellitus | - | 1 | 1 | 1 | 1 | 1 |
| K04932 | Non-alcoholic fatty liver disease (NAFLD) | 3 | 4 | 4 | 4 | 4 | 4 |
| K04931 | Insulin resistance | 1 | 1 | 1 | 1 | 1 | 1 |
| K04934 | Cushing syndrome | 1 | 1 | 1 | 1 | 1 | 1 |
| K05120 | Epithelial cell signaling in Helicobacterpylori infect | 1 | 1 | 1 | 1 | 1 | 1 |
| K05130 | Pathogenic Escherichia coli infection | 1 | 2 | 2 | 2 | 2 | 2 |
| K05133 | Pertussis | 1 | 1 | 1 | 1 | 1 | 1 |
| K05134 | Legionellosis | 2 | 4 | 4 | 4 | 4 | 4 |
| K05152 | Tuberculosis | 2 | 4 | 4 | 4 | 4 | 4 |
| K05170 | Human immunodeficiency virus 1 infection | - | 1 | 1 | 1 | 1 | 1 |
| K05162 | Measles | - | 1 | 1 | 1 | 1 | 1 |
| K05164 | Influenza A | - | 1 | 1 | 1 | 1 | 1 |
| K05161 | Hepatitis B | - | 1 | 1 | 1 | 1 | 1 |
| K05160 | Hepatitis C | - | 1 | 1 | 1 | 1 | 1 |
| K05168 | Herpes simplex virus 1 infection | - | 1 | 1 | 1 | 1 | 1 |
| K05163 | Human cytomegalovirus infection | - | 1 | 1 | 1 | 1 | 1 |
| K05167 | Kaposi sarcoma-associated herpesvirus infection | - | 1 | 1 | 1 | 1 | 1 |
| K05169 | Epstein-Barr virus infection | - | 1 | 1 | 1 | 1 | 1 |
| K05145 | Toxoplasmosis | - | 1 | 1 | 1 | 1 | 1 |
| K01501 | beta-Lactam resistance | 1 | 3 | 3 | 2 | 2 | 2 |
| K01502 | Vancomycin resistance | 4 | 4 | 4 | 4 | 4 | 4 |
| K01503 | Cationic antimicrobial peptide (CAMP) resistance | 2 | 2 | 2 | 2 | 2 | 2 |
| K01524 | Platinum drug resistance | - | 2 | 2 | 2 | 2 | 2 |
| K01523 | Antifolate resistance | 2 | 2 | 2 | 3 | 3 | 3 |

**hways from Wolbachia genomes using KASS.**

| Wolbachia | | | | | | | | | | | | | |
|---|---|---|---|---|---|---|---|---|---|---|---|---|---|
| **wTpre** | **wVulC** | **wMel** | **wCau** | **wNfla** | **wBm** | **wBp** | **wWb** | **wLs** | **wLbra** | **wCle** | **wMhie** | **wCfeJ** | **wOv** |
| 11 | 11 | 11 | 11 | 11 | 11 | 11 | 10 | 11 | 11 | 11 | 10 | 11 | 11 |
| 17 | 17 | 16 | 17 | 16 | 17 | 17 | 16 | 17 | 17 | 17 | 17 | 17 | 17 |
| 6 | 6 | 6 | 6 | 6 | 6 | 6 | 6 | 6 | 6 | 6 | 4 | 6 | 6 |
| 1 | 1 | 2 | 2 | 2 | 1 | 1 | 1 | 1 | 1 | 2 | - | 1 | 1 |
| 4 | 5 | 4 | 4 | 4 | 4 | 4 | 5 | 4 | 4 | 4 | 4 | 5 | 4 |
| - | - | 1 | 1 | 1 | - | - | - | - | - | 1 | - | - | - |
| 5 | 6 | 6 | 6 | 6 | 5 | 4 | 6 | 5 | 5 | 6 | 6 | 6 | 3 |
| 7 | 8 | 7 | 8 | 8 | 7 | 8 | 7 | 8 | 8 | 7 | 8 | 8 | 7 |
| 9 | 9 | 9 | 8 | 9 | 9 | 9 | 9 | 9 | 9 | 9 | 9 | 9 | 9 |
| 6 | 6 | 6 | 6 | 6 | 6 | 6 | 6 | 6 | 6 | 6 | 6 | 6 | 6 |
| 4 | 4 | 3 | 4 | 3 | 4 | 4 | 4 | 4 | 4 | 4 | 4 | 4 | 4 |
| 2 | 2 | 2 | 2 | 2 | 2 | 2 | 2 | 2 | 2 | 2 | 2 | 2 | 2 |
| 2 | 2 | 2 | 2 | 2 | 2 | 2 | 2 | 2 | 2 | 2 | 2 | 2 | 2 |
| **37** | 36 | **38** | **39** | 37 | **37** | **36** | **36** | **37** | **37** | **36** | **36** | **38** | **36** |
| 8 | 8 | 8 | 8 | 8 | 8 | 8 | 8 | 8 | 8 | 8 | 8 | 8 | 8 |
| 9 | 10 | 9 | 10 | 10 | 9 | 10 | 10 | 10 | 10 | 9 | 8 | 10 | 9 |
| 12 | 12 | 11 | 12 | 11 | 12 | 12 | 12 | 12 | 12 | 12 | 12 | 12 | 12 |
| 6 | 6 | 6 | 6 | 6 | 6 | 6 | 6 | 6 | 6 | 6 | 5 | 6 | 6 |
| 2 | 2 | 2 | 2 | 2 | 2 | 2 | 2 | 2 | 2 | 2 | 2 | 2 | 2 |
| **8** | 8 | **8** | **8** | 8 | **8** | **8** | **7** | **8** | **8** | **8** | **8** | **8** | **8** |
| 3 | 4 | 4 | 4 | 4 | 3 | 3 | 3 | 3 | 3 | 3 | 3 | 3 | 3 |
| **9** | 11 | **10** | **10** | 10 | **8** | **9** | **9** | **8** | **7** | **9** | **9** | **10** | **8** |
| **22** | 22 | **22** | **22** | 21 | **22** | **22** | **22** | **21** | **20** | **21** | **20** | **21** | **20** |
| **17** | 17 | **17** | **17** | 17 | **17** | **17** | **17** | **17** | **17** | **17** | **16** | **17** | **17** |
| 12 | 12 | 12 | 12 | 11 | 11 | 11 | 12 | 11 | 11 | 11 | 11 | 11 | 10 |
| 7 | 7 | 8 | 8 | 8 | 7 | 7 | 7 | 7 | 7 | 7 | 7 | 7 | 7 |
| 7 | 8 | 8 | 8 | 8 | 8 | 8 | 8 | 8 | 8 | 7 | 7 | 8 | 7 |
| 3 | 3 | 3 | 3 | 3 | 3 | 3 | 3 | 3 | 3 | 3 | 3 | 3 | 3 |
| 9 | 10 | 10 | 10 | 10 | 10 | 10 | 10 | 10 | 10 | 10 | 10 | 10 | 9 |
| 2 | 2 | 2 | 2 | 2 | 2 | 2 | 2 | 2 | 2 | 2 | 2 | 2 | 2 |
| 6 | 6 | 6 | 6 | 6 | 6 | 6 | 6 | 6 | 6 | 6 | 6 | 6 | 5 |
| 2 | 2 | 2 | 2 | 2 | 2 | 2 | 2 | 2 | 2 | 2 | 2 | 2 | 2 |
| 1 | 1 | 1 | 1 | 1 | 1 | 1 | 1 | 1 | 1 | 1 | 1 | 1 | 1 |
| 1 | 1 | 1 | 1 | 1 | 1 | 1 | 1 | 1 | 1 | 1 | 1 | 1 | 1 |
| 2 | 2 | 2 | 2 | 2 | 2 | 2 | 2 | 2 | 2 | 2 | 2 | 2 | 2 |
| 1 | 1 | 1 | 1 | 1 | 1 | 1 | 1 | 1 | 1 | 1 | 1 | 1 | 1 |
| 1 | 1 | 1 | 1 | 1 | 1 | 1 | 1 | 1 | 1 | 1 | 1 | 1 | 1 |
| 1 | 1 | 1 | 1 | 1 | 1 | 1 | 1 | 1 | 1 | 1 | 1 | 1 | 1 |

| | | | | | | | | | | | | | |
|---|---|---|---|---|---|---|---|---|---|---|---|---|---|
| 3 | 3 | 3 | 3 | 3 | 3 | 3 | 3 | 3 | 3 | 3 | 3 | 3 | 3 |
| 1 | 1 | 1 | 1 | 1 | 1 | 1 | 1 | 1 | 1 | 1 | 1 | 1 | 1 |
| 3 | 3 | 3 | 3 | 3 | 3 | 3 | 3 | 3 | 3 | 3 | 3 | 3 | 3 |
| 1 | 1 | 1 | 1 | 1 | 1 | 1 | - | 1 | 1 | 1 | 1 | 1 | 1 |
| 5 | 5 | 5 | 5 | 5 | 5 | 5 | 5 | 5 | 5 | 5 | 5 | 5 | 4 |
| 11 | 12 | 12 | 12 | 12 | 11 | 12 | 10 | 11 | 12 | 12 | 12 | 13 | 11 |
| 2 | 3 | 4 | 4 | 4 | 2 | 2 | 2 | 2 | 2 | 4 | 2 | 4 | 2 |
| 6 | 6 | 6 | 6 | 6 | 6 | 6 | 6 | 6 | 5 | 6 | 5 | 5 | 1 |
| 2 | 2 | 2 | 2 | 2 | - | - | 2 | - | 1 | 2 | 2 | 2 | 1 |
| 2 | 2 | 2 | 2 | 2 | 2 | 2 | 2 | 2 | 2 | 2 | 2 | 2 | 2 |
| 3 | 3 | 3 | 3 | 3 | 3 | 3 | 3 | 3 | 3 | 3 | 3 | 3 | 1 |
| 5 | 10 | 5 | 5 | 10 | 5 | 5 | 4 | 5 | 5 | 10 | 5 | 5 | 5 |
| 2 | 2 | 2 | 2 | 2 | 2 | 2 | 2 | 2 | 2 | 2 | 2 | 2 | 2 |
| 5 | 5 | 3 | 5 | 5 | 1 | 1 | 1 | 1 | - | 4 | 4 | 5 | 5 |
| 7 | 7 | 7 | 7 | 7 | 6 | 7 | 6 | 6 | 5 | 7 | 7 | 5 | 7 |
| 11 | 11 | 11 | 11 | 11 | 10 | 11 | 9 | 11 | 11 | 11 | 11 | 11 | 10 |
| 7 | 7 | 7 | 7 | 7 | 7 | 7 | 7 | 7 | 6 | 7 | 7 | 7 | 7 |
| 8 | 8 | 8 | 8 | 8 | 8 | 8 | 8 | 8 | 8 | 8 | 8 | 8 | 4 |
| 1 | 1 | 1 | 1 | 1 | 1 | 1 | 1 | 1 | 1 | 1 | 1 | 1 | 1 |
| 1 | 1 | 1 | 1 | 1 | 1 | 1 | 1 | 1 | 1 | 1 | 1 | 1 | 1 |
| 1 | 1 | 1 | 1 | 1 | 1 | 1 | 1 | 1 | 1 | 1 | 1 | 1 | 1 |
| 1 | 1 | 1 | 1 | 1 | 1 | 1 | 1 | 1 | 1 | 1 | 1 | 1 | 1 |
| 1 | 1 | 1 | 1 | 1 | 1 | 1 | 1 | 1 | 1 | 1 | 1 | 1 | 1 |
| 4 | 4 | 4 | 4 | 4 | 4 | 4 | 4 | 4 | 4 | 4 | 4 | 4 | 4 |
| 1 | 1 | 2 | 2 | 2 | 1 | 1 | 1 | 1 | 1 | 1 | 1 | 1 | 1 |
| 1 | 1 | 1 | 1 | 1 | 1 | 1 | 1 | 1 | 1 | 1 | 1 | 1 | 1 |
| 3 | 3 | 3 | 3 | 3 | 3 | 3 | 3 | 3 | 3 | 3 | 3 | 3 | 3 |
| 1 | 1 | 1 | 1 | 1 | 1 | 1 | 1 | 1 | 1 | 1 | 1 | 1 | 1 |
| - | 1 | 1 | - | 1 | 1 | 1 | 1 | - | - | 1 | 1 | 1 | 1 |
| 1 | 1 | 1 | 1 | 1 | 1 | 1 | 1 | 1 | 1 | 1 | 1 | 1 | 1 |
| - | 1 | 1 | - | 1 | 1 | 1 | 1 | - | - | 1 | 1 | 1 | 1 |
| 1 | 1 | 1 | 1 | 1 | 1 | 1 | 1 | 1 | 1 | 1 | 1 | 1 | 1 |
| 1 | 1 | 1 | 1 | 1 | 1 | 1 | 1 | 1 | 1 | 1 | 1 | 1 | 1 |
| 5 | 5 | 5 | 5 | 5 | 5 | 5 | 5 | 5 | 5 | 5 | 5 | 5 | 5 |
| 3 | 3 | 3 | 3 | 3 | 3 | 3 | 3 | 3 | 3 | 3 | 3 | 3 | 3 |
| 52 | 52 | 52 | 53 | 52 | 52 | 53 | 49 | 51 | 52 | 51 | 48 | 51 | 52 |
| 24 | 24 | 24 | 24 | 24 | 24 | 24 | 22 | 24 | 23 | 24 | 24 | 24 | 24 |
| 1 | 1 | 1 | 1 | 1 | 1 | 1 | 1 | 1 | 1 | 1 | 1 | 1 | 1 |
| 15 | 15 | 15 | 15 | 14 | 15 | 15 | 15 | 15 | 15 | 15 | 15 | 14 | 15 |
| 2 | 1 | 1 | 2 | 1 | 1 | 1 | 1 | 1 | 1 | 1 | 1 | 2 | 1 |
| 2 | 2 | 2 | 2 | 2 | 2 | 2 | 2 | 2 | 2 | 2 | 2 | 2 | 2 |
| 9 | 9 | 9 | 9 | 9 | 9 | 9 | 8 | 9 | 9 | 9 | 9 | 10 | 9 |

| | | | | | | | | | | | | | |
|---|---|---|---|---|---|---|---|---|---|---|---|---|---|
| 13 | 13 | 13 | 13 | 13 | 13 | 13 | 12 | 13 | 13 | 13 | 13 | 13 | 13 |
| 8 | 8 | 8 | 8 | 8 | 8 | 8 | 8 | 8 | 8 | 8 | 8 | 8 | 8 |
| 6 | 6 | 6 | 6 | 6 | 6 | 6 | 5 | 6 | 6 | 6 | 6 | 6 | 6 |
| 12 | 12 | 12 | 12 | 12 | 12 | 12 | 12 | 12 | 12 | 12 | 12 | 12 | 12 |
| 18 | 18 | 18 | 18 | 18 | 18 | 18 | 18 | 13 | 11 | 18 | 18 | 18 | 10 |
| 20 | 20 | 19 | 19 | 19 | 20 | 20 | 20 | 17 | 18 | 20 | 20 | 20 | 15 |
| 23 | 23 | 23 | 23 | 22 | 22 | 22 | 22 | 23 | 23 | 23 | 23 | 22 | 22 |
| 12 | 12 | 14 | 14 | 14 | 12 | 14 | 10 | 12 | 12 | 10 | 12 | 16 | 10 |
| 1 | 1 | 1 | 1 | 1 | 1 | 1 | 1 | 1 | 1 | 1 | 1 | 1 | 1 |
| 1 | 1 | 1 | 1 | 1 | 1 | 1 | 1 | 1 | 1 | 1 | 1 | 1 | 1 |
| 5 | 5 | 5 | 5 | 5 | 5 | 5 | 5 | 5 | 5 | 5 | 5 | 5 | 5 |
| 1 | 1 | 1 | 1 | 1 | 1 | 1 | 1 | 1 | 1 | 1 | 1 | 1 | 1 |
| 2 | 3 | 3 | 3 | 3 | 2 | 2 | 2 | 2 | 2 | 2 | 2 | 2 | 2 |
| 1 | 1 | 1 | 1 | 1 | 1 | 1 | 1 | 1 | 1 | 1 | 1 | 1 | 1 |
| 1 | 1 | 1 | 1 | 1 | 1 | 1 | 1 | 1 | 1 | 1 | 1 | 1 | 1 |
| 3 | 3 | 3 | 3 | 3 | 3 | 3 | 3 | 3 | 3 | 3 | 3 | 3 | 3 |
| 13 | 13 | 14 | 14 | 14 | 13 | 15 | 13 | 12 | 13 | 12 | 13 | 16 | 10 |
| 1 | 1 | 1 | 1 | 1 | 1 | 1 | 1 | 1 | 1 | 1 | 1 | 1 | 1 |
| 2 | 2 | 2 | 2 | 2 | 2 | 2 | 2 | 2 | 2 | 2 | 2 | 2 | 2 |
| 1 | 1 | 1 | 1 | 1 | 1 | 1 | 1 | 1 | 1 | 1 | 1 | 1 | 1 |
| 2 | 2 | 2 | 2 | 2 | 2 | 2 | 2 | 2 | 2 | 2 | 2 | 2 | 2 |
| 1 | 1 | 1 | 1 | 1 | 1 | 1 | 1 | 1 | 1 | 1 | 1 | 1 | 1 |
| 11 | 11 | 11 | 11 | 10 | 11 | 11 | 11 | 11 | 10 | 11 | 10 | 10 | 10 |
| 1 | 1 | 1 | 1 | 1 | 1 | 1 | 1 | 1 | 1 | - | 1 | 1 | 1 |
| 2 | 1 | 2 | 2 | 1 | 2 | 2 | 2 | 2 | 2 | 2 | 2 | 2 | 2 |
| 1 | 2 | 1 | 1 | 2 | 1 | 1 | 1 | 1 | 1 | 1 | 1 | 1 | 1 |
| 1 | 1 | 1 | 1 | 1 | 1 | 1 | 1 | 1 | 1 | 1 | 1 | 1 | 1 |
| 1 | 1 | 1 | 1 | 1 | 1 | 1 | 1 | 1 | 1 | 2 | 1 | 1 | 1 |
| 2 | 1 | 2 | 2 | 1 | 2 | 2 | 2 | 2 | 2 | 2 | 2 | 2 | 2 |
| - | 2 | - | 1 | 2 | - | 1 | 1 | 1 | 1 | - | 1 | 1 | - |
| 1 | 1 | 1 | 1 | 1 | 1 | 1 | 1 | 1 | 1 | 1 | 1 | 1 | 1 |
| 1 | 1 | 1 | 1 | 1 | 1 | 1 | 1 | 1 | 1 | 1 | 1 | 1 | - |
| 3 | 1 | 3 | 3 | 1 | 3 | 3 | 2 | 3 | 3 | 3 | 3 | 3 | 3 |
| 1 | 3 | 1 | 1 | 3 | 1 | 1 | 1 | 1 | 1 | 1 | 1 | 1 | 1 |
| 1 | 1 | 1 | 1 | 1 | 1 | 1 | 1 | 1 | 1 | 1 | 1 | 1 | 1 |
| 1 | 1 | 1 | 1 | 1 | 1 | 1 | 1 | 1 | 1 | 1 | 1 | 1 | 1 |
| 5 | 1 | 5 | 5 | 1 | 5 | 5 | 5 | 5 | 5 | 5 | 5 | 5 | 5 |
| 2 | 5 | 2 | 2 | 5 | 2 | 2 | 2 | 2 | 2 | 2 | 2 | 2 | 2 |
| 6 | 2 | 6 | 6 | 2 | 6 | 6 | 5 | 6 | 6 | 6 | 6 | 6 | 6 |
| 2 | 6 | 2 | 2 | 6 | 2 | 3 | 2 | 3 | 2 | 2 | 2 | 3 | 2 |
| 4 | 3 | 4 | 4 | 3 | 4 | 4 | 4 | 4 | 4 | 4 | 4 | 4 | 4 |
| 1 | 4 | 1 | 1 | 4 | 1 | 1 | 1 | 1 | 1 | 1 | 1 | 1 | 1 |

| | | | | | | | | | | | | | |
|---|---|---|---|---|---|---|---|---|---|---|---|---|---|
| 1 | 1 | 1 | 1 | 1 | 1 | 1 | 1 | 1 | 1 | 1 | 1 | 1 | 1 |
| 2 | 1 | 2 | 2 | 1 | 2 | 2 | 2 | 2 | 2 | 2 | 2 | 2 | 2 |
| - | 2 | 1 | 1 | 2 | - | - | - | - | - | - | - | - | - |
| 1 | 1 | 1 | 1 | 1 | 1 | 1 | 1 | 1 | 1 | 1 | 1 | 1 | 1 |
| 1 | 1 | 1 | 1 | 1 | 1 | 1 | 1 | 1 | 1 | 1 | 1 | 1 | 1 |
| 1 | 1 | 1 | 1 | 1 | 1 | 1 | 1 | 1 | 1 | 1 | 1 | 1 | 1 |
| 1 | 1 | 1 | 1 | 1 | 1 | 1 | 1 | 1 | 1 | 1 | 1 | 1 | 1 |
| 1 | 1 | 1 | 1 | 1 | 1 | 1 | 1 | 1 | 1 | 1 | 1 | 1 | 1 |
| 5 | 5 | 5 | 5 | 5 | 5 | 5 | 4 | 5 | 5 | 5 | 5 | 5 | 5 |
| 4 | 4 | 4 | 4 | 4 | 4 | 4 | 3 | 4 | 4 | 4 | 4 | 4 | 4 |
| 1 | 1 | 1 | 1 | 1 | 1 | 1 | 1 | 1 | 1 | 1 | 1 | 1 | 1 |
| 5 | 5 | 5 | 5 | 5 | 5 | 5 | 4 | 5 | 5 | 5 | 5 | 5 | 5 |
| 3 | 3 | 3 | 3 | 3 | 3 | 3 | 3 | 3 | 3 | 3 | 3 | 3 | 3 |
| 1 | 1 | 1 | 1 | 1 | 1 | 1 | 1 | 1 | 1 | 1 | 1 | 1 | 1 |
| 1 | 1 | 1 | 1 | 1 | 1 | 1 | 1 | 1 | 1 | 1 | 1 | 1 | 1 |
| 4 | 4 | 4 | 4 | 4 | 4 | 4 | 3 | 4 | 4 | 4 | 4 | 4 | 4 |
| 1 | 1 | 1 | 1 | 1 | 1 | 1 | 1 | 1 | 1 | 1 | 1 | 1 | - |
| 1 | 1 | 1 | 1 | 1 | 1 | 1 | 1 | 1 | 1 | 1 | 1 | 1 | 1 |
| 1 | 1 | 1 | 1 | 1 | 1 | 1 | 1 | 1 | 1 | 1 | 1 | 1 | 1 |
| 2 | 2 | 2 | 2 | 2 | 2 | 2 | 2 | 2 | 2 | 2 | 2 | 2 | 2 |
| 1 | 1 | 1 | 1 | 1 | 1 | 1 | 1 | 1 | 1 | 1 | 1 | 1 | 1 |
| 4 | 4 | 4 | 4 | 4 | 4 | 4 | 4 | 4 | 4 | 4 | 4 | 4 | 4 |
| 4 | 4 | 4 | 4 | 4 | 4 | 4 | 4 | 4 | 4 | 4 | 4 | 4 | 4 |
| 1 | 1 | 1 | 1 | 1 | 1 | 1 | 1 | 1 | 1 | 1 | 1 | 1 | 1 |
| 1 | 1 | 1 | 1 | 1 | 1 | 1 | 1 | 1 | 1 | 1 | 1 | 1 | 1 |
| 1 | 1 | 1 | 1 | 1 | 1 | 1 | 1 | 1 | 1 | 1 | 1 | 1 | 1 |
| 1 | 1 | 1 | 1 | 1 | 1 | 1 | 1 | 1 | 1 | 1 | 1 | 1 | 1 |
| 1 | 1 | 1 | 1 | 1 | 1 | 1 | 1 | 1 | 1 | 1 | 1 | 1 | 1 |
| 1 | 1 | 1 | 1 | 1 | 1 | 1 | 1 | 1 | 1 | 1 | 1 | 1 | 1 |
| 1 | 1 | 1 | 1 | 1 | 1 | 1 | 1 | 1 | 1 | 1 | 1 | 1 | 1 |
| 1 | 1 | 1 | 1 | 1 | 1 | 1 | 1 | 1 | 1 | 1 | 1 | 1 | 1 |
| 1 | 1 | 1 | 1 | 1 | 1 | 1 | 1 | 1 | 1 | 1 | 1 | 1 | 1 |
| 1 | 1 | 1 | 1 | 1 | 1 | 1 | 1 | 1 | 1 | 1 | 1 | 1 | 1 |
| 2 | 3 | 2 | 3 | 3 | 2 | 2 | 2 | 2 | 3 | 3 | 3 | 3 | 2 |
| 4 | 4 | 4 | 4 | 4 | 4 | 4 | 3 | 4 | 4 | 4 | 4 | 4 | 4 |
| 2 | 2 | 2 | 2 | 2 | 2 | 2 | 1 | 2 | 2 | 2 | 2 | 2 | 2 |
| 2 | 2 | 2 | 2 | 2 | 2 | 2 | 2 | 2 | 2 | 2 | 2 | 2 | 2 |
| 3 | 3 | 3 | 3 | 3 | 2 | 2 | 2 | 2 | 1 | 3 | 3 | 3 | 3 |

| wOo | wDi | wDcau | wCtub |
|---|---|---|---|
| 11 | 11 | 11 | 11 |
| 17 | 17 | 17 | 17 |
| 6 | 6 | 6 | 6 |
| 1 | 1 | 1 | 1 |
| 4 | 4 | 4 | 4 |
| - | - | - | - |
| 4 | 5 | 5 | 5 |
| 8 | 8 | 8 | 8 |
| 9 | 9 | 9 | 9 |
| 6 | 6 | 6 | 6 |
| 4 | 4 | 4 | 4 |
| 2 | 2 | 2 | 2 |
| 2 | 2 | 2 | 2 |
| **36** | **36** | **36** | **36** |
| 8 | 8 | 8 | 8 |
| 10 | 10 | 10 | 10 |
| 12 | 12 | 12 | 12 |
| 6 | 6 | 6 | 6 |
| 2 | 2 | 2 | 2 |
| **8** | **8** | **8** | **8** |
| 3 | 3 | 3 | 3 |
| **9** | **10** | **9** | **9** |
| **21** | **21** | **21** | **21** |
| **17** | **17** | **17** | **17** |
| 10 | 10 | 10 | 10 |
| 7 | 7 | 7 | 7 |
| 8 | 8 | 8 | 8 |
| 3 | 3 | 3 | 3 |
| 10 | 10 | 10 | 10 |
| 2 | 2 | 2 | 2 |
| 6 | 6 | 6 | 6 |
| 2 | 1 | 1 | 1 |
| 1 | 1 | 1 | 1 |
| 1 | 1 | 1 | 1 |
| 2 | 2 | 1 | 2 |
| 1 | 1 | 1 | 1 |
| 1 | 1 | 1 | 1 |
| 1 | 1 | 1 | 1 |

Difference of number genes:

<-5 -5 -4 -3 -2 -1 0 1 2 3 4 5

| | | | |
|---|---|---|---|
| 3 | 3 | 3 | 3 |
| 1 | 1 | 1 | 1 |
| 3 | 3 | 3 | 3 |
| 1 | 1 | 1 | 1 |
| 5 | 5 | 4 | 4 |
| 11 | 11 | 11 | 11 |
| 2 | 2 | 2 | 2 |
| **4** | **5** | **5** | **5** |
| **2** | **2** | **2** | **2** |
| 2 | 2 | 2 | 2 |
| 3 | 3 | 3 | 3 |
| **5** | **5** | **5** | **5** |
| 2 | 2 | 2 | 2 |
| **5** | **5** | **4** | **4** |
| 7 | 7 | 7 | 7 |
| **10** | **10** | **10** | **10** |
| 7 | 7 | 7 | 7 |
| 5 | 7 | 8 | 8 |
| 1 | 1 | 1 | 1 |
| 1 | 1 | 1 | 1 |
| 1 | 1 | 1 | 1 |
| 1 | 1 | 1 | 1 |
| 1 | 1 | 1 | 1 |
| 4 | 4 | 4 | 4 |
| 1 | 1 | 1 | 1 |
| 1 | 1 | 1 | 1 |
| 3 | 3 | 3 | 3 |
| 1 | 1 | 1 | 1 |
| 1 | 1 | 1 | 1 |
| 1 | 1 | 1 | 1 |
| 1 | 1 | 1 | 1 |
| 1 | 1 | 1 | - |
| 1 | 1 | 1 | - |
| 5 | 5 | 4 | 4 |
| 3 | 3 | 3 | 3 |
| 50 | 52 | 52 | 52 |
| 24 | 24 | 24 | 24 |
| 1 | 1 | 1 | 1 |
| 15 | 15 | 15 | 15 |
| 1 | 1 | 1 | 1 |
| 2 | 2 | 2 | 2 |
| 9 | 9 | 9 | 9 |

| | | | |
|---|---|---|---|
| 13 | 13 | 13 | 13 |
| 8 | 8 | 7 | 7 |
| 6 | 6 | 5 | 5 |
| 12 | 12 | 11 | 12 |
| **11** | **11** | **8** | **17** |
| **15** | **17** | **12** | **12** |
| **22** | **22** | **22** | **22** |
| 12 | 11 | 11 | 11 |
| 1 | 1 | 1 | 1 |
| 1 | 1 | 1 | 1 |
| 5 | 5 | 5 | 5 |
| 1 | 1 | 1 | 1 |
| 2 | 2 | 2 | 2 |
| 1 | 1 | 1 | 1 |
| 1 | 1 | 1 | 1 |
| 3 | 3 | 3 | 3 |
| **12** | **11** | **8** | **9** |
| 1 | 1 | 1 | 1 |
| 2 | 2 | 2 | 2 |
| 1 | 1 | 1 | 1 |
| 2 | 2 | 2 | 2 |
| 1 | 1 | 1 | 1 |
| 10 | 11 | 10 | 10 |
| 1 | 1 | 1 | 1 |
| 2 | 2 | 2 | 2 |
| 1 | 1 | 1 | 1 |
| 1 | 1 | 1 | 1 |
| 1 | 1 | 1 | 1 |
| 2 | 2 | 2 | 2 |
| 1 | 1 | 1 | 1 |
| 1 | 1 | 1 | 1 |
| 1 | 1 | 1 | 1 |
| 3 | 3 | 3 | 3 |
| 1 | 1 | 1 | 1 |
| 1 | 1 | 1 | 1 |
| 1 | 1 | 1 | 1 |
| 5 | 5 | 4 | 4 |
| 2 | 2 | 2 | 2 |
| 6 | 6 | 6 | 6 |
| 2 | 3 | 2 | 2 |
| 4 | 4 | 3 | 3 |
| 1 | 1 | 1 | 1 |

| | | | |
|---|---|---|---|
| 1 | 1 | - | - |
| 2 | 2 | 2 | 2 |
| - | - | - | - |
| 1 | 1 | 1 | 1 |
| 1 | 1 | - | - |
| 1 | 1 | 1 | 1 |
| 1 | 1 | 1 | 1 |
| 1 | 1 | 1 | 1 |
| 5 | 5 | 5 | 5 |
| 4 | 4 | 4 | 4 |
| 1 | 1 | 1 | 1 |
| 5 | 5 | 5 | 5 |
| 3 | 3 | 2 | 2 |
| 1 | 1 | 1 | 1 |
| 1 | 1 | 1 | 1 |
| 4 | 4 | 4 | 4 |
| - | 1 | 1 | 1 |
| 1 | 1 | 1 | 1 |
| 1 | 1 | 1 | 1 |
| 2 | 2 | 2 | 2 |
| 1 | 1 | 2 | 1 |
| 4 | 4 | 4 | 4 |
| 4 | 4 | 4 | 4 |
| 1 | 1 | 1 | 1 |
| 1 | 1 | 1 | 1 |
| 1 | 1 | 1 | 1 |
| 1 | 1 | 1 | 1 |
| 1 | 1 | 1 | 1 |
| 1 | 1 | 1 | 1 |
| 1 | 1 | 1 | 1 |
| 1 | 1 | 1 | 1 |
| 1 | 1 | 1 | 1 |
| 1 | 1 | 1 | 1 |
| 2 | 2 | 2 | 2 |
| 4 | 4 | 4 | 4 |
| 2 | 2 | 2 | 2 |
| 2 | 2 | 1 | 1 |
| 3 | 3 | 3 | 3 |

**Table S3. Summary of number of genes assigned to 165 different KEGG pathwa**

| | K0 | Wolbachia wBm | wBp |
|---|---|---|---|
| **K00730 Thiamine metabolism** | | | |
| iscS, NFS1 | **K04487** | WP_011256223.1 | WP_011256223.1 |
| | | WP_041571528.1 | WP_167896281.1 |
| tenA | **K03707** | | |
| adk, AK | **K00939** | WP_011256520.1 | WP_167895918.1 |
| total | | 3 | 3 |
| **K00740 Riboflavin metabolism** | | | |
| ribA, RIB1 | **K01497** | WP_011256477.1 | WP_167895986.1 |
| ribD | **K11752** | WP_011256228.1 | WP_167895712.1 |
| ribB, RIB3 | **K02858** | WP_011256511.1 | WP_167895926.1 |
| ribH, RIB4 | **K00794** | WP_011256390.1 | WP_011256390.1 |
| ribE, RIB5 | **K00793** | WP_011256285.1 | WP_167896219.1 |
| ribF | **K11753** | WP_011256614.1 | WP_011256614.1 |
| total | | 6 | 6 |
| **K00780 Biotin metabolism** | | | |
| bioC | **K02169** | | |
| fabF, OXSM, CEM1 | **K09458** | WP_011256936.1 | WP_011256936.1 |
| fabG, OAR1 | **K00059** | WP_011256509.1 | WP_011256509.1 |
| fabZ | **K02372** | WP_011256254.1 | WP_167896200.1 |
| fabI | **K00208** | WP_011256668.1 | WP_167895823.1 |
| bioF | **K00652** | | |
| bioA | **K00833** | | |
| bioD | **K01935** | | |
| bioB | **K01012** | | |
| birA | **K03524** | WP_011256818.1 | WP_167896086.1 |
| total | | 5 | 5 |
| **K00061 Fatty acid biosynthesis** | | | |
| fabD | **K00645** | WP_011256600.1 | WP_167895869.1 |
| fabH | **K00648** | WP_011256812.1 | WP_167896082.1 |
| fabF, OXSM, CEM1 | **K09458** | WP_011256936.1 | WP_011256936.1 |
| fabG, OAR1 | **K00059** | WP_011256509.1 | WP_011256509.1 |
| fabZ | **K02372** | WP_011256254.1 | WP_167896200.1 |
| fabI | **K00208** | WP_011256668.1 | WP_167895823.1 |
| fabK | **K02371** | WP_011256950.1 | WP_167896189.1 |
| total | | 7 | 7 |
| **K00790 Folate biosynthesis** | | | |
| ribA, RIB1 | **K01497** | WP_011256477.1 | WP_167895986.1 |
| folB | **K01633** | | |

| | | | |
|---|---|---|---|
| folKP | **K13941** | | |
| folC | **K11754** | | |
| DHFR, folA | **K00287** | | |
| total | | 1 | 1 |
| **K03070 Bacterial secretion system** | | | |
| tolC | **K12340** | WP_011256773.1 | WP_167895745.1 |
| gspD | **K02453** | | |
| secD | **K03072** | WP_041571521.1 | WP_167896268.1 |
| secF | **K03074** | WP_011256387.1 | WP_167895973.1 |
| secE | **K03073** | WP_041571587.1 | WP_041571587.1 |
| SecG | **K03075** | WP_041571540.1 | WP_167895943.1 |
| SecY | **K03076** | WP_011256521.1 | WP_011256521.1 |
| YajC | **K03210** | WP_011256423.1 | WP_167896027.1 |
| yidC, spoIIIJ, OXA1, ccfA | **K03217** | WP_011256389.1 | WP_167895972.1 |
| SecA | **K03070** | WP_011256465.1 | WP_167895993.1 |
| ftsY | **K03110** | WP_041571510.1 | WP_041571510.1 |
| SecB | **K03071** | WP_011256739.1 | WP_011256739.1 |
| ffh | **K03106** | WP_011256251.1 | WP_167895727.1 |
| TatA | **K03116** | WP_006013840.1 | WP_006013840.1 |
| TatC | **K03118** | WP_011256496.1 | WP_011256496.1 |
| virB3, lvhB3 | **K03198** | WP_011256995.1 | WP_011256995.1 |
| virB9, lvhB9 | **K03204** | WP_011256789.1 | WP_041571550.1 |
| | | WP_041571550.1 | WP_167895737.1 |
| virB6, lvhB6 | **K03201** | WP_011256991.1 | WP_167896235.1 |
| | | WP_041571522.1 | WP_167896236.1 |
| | | WP_050707683.1 | WP_167896237.1 |
| | | | WP_167896238.1 |
| virB8 | **K03203** | WP_011256478.1 | WP_167895985.1 |
| | | WP_011256839.1 | WP_167896107.1 |
| virB10 | **K03195** | WP_011256480.1 | WP_167895984.1 |
| virB4 | **K03199** | WP_011256947.1 | WP_167896186.1 |
| | | WP_011256994.1 | WP_167896239.1 |
| virB11 | **K03196** | WP_011256481.1 | WP_167895983.1 |
| virD4 | **K03205** | WP_011256482.1 | WP_167895982.1 |
| total | | 27 | 28 |
| **K00190 Oxidative phosphorylation** | | | |
| **NADH Dehydrogenase** | | | |
| nuoA | **K00330** | WP_011256440.1 | WP_167896015.1 |
| nuoB | **K00331** | WP_011256441.1 | WP_167896014.1 |
| nuoC | **K00332** | WP_011256442.1 | WP_167896013.1 |
| nuoD | **K00333** | WP_011256326.1 | WP_011256326.1 |
| nuoE | **K00334** | WP_011256792.1 | WP_167895735.1 |

| | | | |
|---|---|---|---|
| nuoF | **K00335** | WP_041571573.1 | WP_167896295.1 |
| nuoG | **K00336** | WP_011256574.1 | WP_167895884.1 |
| nuoH | **K00337** | WP_011256573.1 | WP_011256573.1 |
| nuoI | **K00338** | WP_011256669.1 | WP_167895822.1 |
| nuoJ | **K00339** | WP_041571486.1 | WP_167896091.1 |
| nuoK | **K00340** | WP_011256822.1 | WP_167896090.1 |
| nuoL | **K00341** | WP_011256821.1 | WP_167896089.1 |
| nuoM | **K00342** | WP_011256820.1 | WP_167896088.1 |
| nuoN | **K00343** | WP_011256819.1 | WP_167896087.1 |
| **succinate dehydrogenase** | | | |
| sdhC, frdC | **K00241** | WP_011256684.1 | WP_167895807.1 |
| sdhD, frdD | **K00242** | WP_011256683.1 | WP_167895808.1 |
| sdhA, frdA | **K00239** | WP_011256646.1 | WP_167895839.1 |
| sdhB, frdB | **K00240** | WP_011256798.1 | WP_011256798.1 |
| **F-type ATPase, prokaryotes and chloroplasts** | | | |
| ATPF1A, atpA | **K02111** | WP_011256513.1 | WP_167895925.1 |
| ATPF1B, atpD | **K02112** | WP_041571591.1 | WP_041571591.1 |
| ATPF1G, atpG | **K02115** | WP_011256902.1 | WP_167896148.1 |
| ATPF1D, atpH | **K02113** | WP_011256514.1 | WP_167895924.1 |
| ATPF1E, atpC | **K02114** | WP_011256885.1 | WP_167896136.1 |
| ATPF0A, atpB | **K02108** | WP_011256658.1 | WP_167895832.1 |
| ATPF0B, atpF | **K02109** | WP_011256655.1 | WP_011256655.1 |
| | | WP_011256656.1 | WP_167895833.1 |
| ATPF0C, atpE | **K02110** | WP_011256657.1 | WP_011256657.1 |
| **cytochrome c reductase** | | | |
| UQCRFS1, RIP1, petA | **K00411** | WP_011256604.1 | WP_167895876.1 |
| CYTB, petB | **K00412** | WP_011256972.1 | WP_167896250.1 |
| CYC1, CYT1, petC | **K00413** | WP_041571516.1 | WP_167896337.1 |
| **Cytochrome c oxidase** | | | |
| COX10, ctaB, cyoE | **K02257** | WP_011256505.1 | WP_167895929.1 |
| COX11, ctaG | **K02258** | WP_011256689.1 | WP_167895802.1 |
| COX15, ctaA | **K02259** | WP_050707669.1 | WP_167895834.1 |
| coxC, ctaE | **K02276** | WP_011256895.1 | WP_167896140.1 |
| coxA, ctaD | **K02274** | WP_011256506.1 | WP_167895928.1 |
| coxB, ctaC | **K02275** | WP_011256507.1 | WP_011256507.1 |
| **Cytochrome bd ubiquinol oxidase** | | | |
| cydA | **K00425** | | |
| cydB | **K00426** | | |

| | | | |
|---|---|---|---|
| total | | 36 | 36 |
| **K04112 Cell cycle - Caulobacter** | | | |
| murG | **K02563** | WP_011256755.1 | WP_167895754.1 |
| ftsW, spoVE | **K03588** | | |
| ftsZ | **K03531** | WP_011256800.1 | WP_167896066.1 |
| ftsQ | **K03589** | WP_011256769.1 | WP_167895747.1 |
| ftsA | **K03590** | WP_011256314.1 | WP_167896204.1 |
| PerP | **K06985** | WP_041571541.1 | WP_041571541.1 |
| pleC | **K07716** | WP_011256329.1 | WP_167896060.1 |
| rseP | **K11749** | WP_011256257.1 | WP_011256257.1 |
| pleD | **K02488** | WP_011256385.1 | WP_167895954.1 |
| clpX, CLPX | **K03544** | WP_011256750.1 | WP_011256750.1 |
| clpP, CLPP | **K01358** | WP_011256751.1 | WP_011256751.1 |
| dnaA | **K02313** | WP_011256475.1 | WP_167895988.1 |
| dnaB | **K02314** | WP_011256498.1 | WP_167895959.1 |
| lon | **K01338** | WP_011256749.1 | WP_167895757.1 |
| total | | 13 | 13 |
| **K03440 Homologous recombination** | | | |
| recJ | **K07462** | WP_011256325.1 | WP_167896336.1 |
| ssb | **K03111** | WP_011256412.1 | WP_167896037.1 |
| recA | **K03553** | WP_011256625.1 | WP_011256625.1 |
| recF | **K03629** | WP_011256327.1 | WP_167896062.1 |
| recO | **K03584** | WP_011256487.1 | WP_167895978.1 |
| recR | **K06187** | WP_011256943.1 | WP_167896183.1 |
| polA | **K02335** | WP_011256922.1 | WP_167896163.1 |
| dnaE | **K02337** | WP_011256697.1 | WP_167895797.1 |
| ruvA | **K03550** | WP_011256450.1 | WP_167896006.1 |
| ruvB | **K03551** | WP_011256451.1 | WP_167896005.1 |
| ruvC | **K01159** | WP_011256894.1 | WP_167896139.1 |
| recG | **K03655** | WP_011256833.1 | WP_167896101.1 |
| total | | 12 | 12 |
| **K02010 ABC transporters** | | | |
| **iron(III) transport system** | | | |
| AfuA | **K02012** | WP_011256596.1 | WP_167895868.1 |
| AfuB | **K02011** | WP_011256241.1 | WP_167895723.1 |
| AfuC | **K02010** | WP_011256875.1 | WP_011256875.1 |
| **phosphate transport system** | | | |
| PstS | **K02040** | WP_011256492.1 | WP_011256492.1 |
| PstC | **K02037** | WP_041571592.1 | WP_167896331.1 |
| PstA | **K02038** | WP_011256432.1 | WP_167896021.1 |

| | | | |
|---|---|---|---|
| PstB | **K02036** | WP_011256379.1 | WP_167895948.1 |
| **lipoprotein-releasing system** | | | |
| lolC_E | **K09808** | WP_041571469.1 | WP_167895810.1 |
| LolD | **K09810** | WP_011256225.1 | WP_167895711.1 |
| **heme exporter** | | | |
| CcmC | **K02195** | WP_011256832.1 | WP_167896100.1 |
| CcmB | **K02194** | WP_011256776.1 | WP_167895744.1 |
| CcmA | **K02193** | WP_011256216.1 | WP_167895704.1 |
| **phospholipid transport system** | | | |
| MlaC | **K07323** | WP_041571568.1 | WP_167895845.1 |
| MlaD | **K02067** | WP_011256220.1 | WP_167895708.1 |
| MlaE | **K02066** | WP_011256815.1 | WP_167896083.1 |
| MlaF | **K02065** | WP_011256816.1 | WP_167896084.1 |
| **zinc transport system** | | | |
| ZnuA | **K09815** | WP_041571504.1 | WP_041571504.1 |
| ZnuB | **K09816** | WP_011256376.1 | WP_011256376.1 |
| ZnuC | **K09817** | WP_011256912.1 | WP_167896156.1 |
| **biotin transport system** | | | |
| BioY | **K03523** | WP_011256714.1 | WP_167895782.1 |
| total | | 20 | 20 |
| **K00564 Glycerophospholipid metabolism** | | | |
| araM, egsA | **K00096** | | |
| gpsA | **K00057** | WP_011256795.1 | WP_011256795.1 |
| plsY | **K08591** | WP_011256621.1 | WP_167895855.1 |
| plsC | **K00655** | WP_041571507.1 | WP_167896171.1 |
| dgkA, DGK | **K00901** | | |
| pld | **K17717** | | WP_167896304.1 |
| E2.7.7.41, CDS1, CDS2, cdsA | **K00981** | WP_011257000.1 | WP_167896277.1 |
| CHO1, pssA | **K17103** | WP_011256623.1 | WP_011256623.1 |
| psd, PISD | **K01613** | WP_041571460.1 | WP_167895853.1 |
| pgsA, PGS1 | **K00995** | WP_011256871.1 | |
| pgpA | **K01095** | WP_041571481.1 | WP_167896321.1 |
| total | | 8 | 8 |
| **K00860 Porphyrin and chlorophyll metabolism** | | | |
| ALAS | **K00643** | WP_011256334.1 | WP_011256334.1 |
| EARS, gltX | **K01885** | WP_011256643.1 | WP_167895837.1 |
| | | WP_011256651.1 | WP_167895841.1 |
| hemB, ALAD | **K01698** | WP_041571452.1 | WP_167896305.1 |
| hemC, HMBS | **K01749** | WP_011256974.1 | WP_167896255.1 |
| hemD, UROS | **K01719** | WP_011256925.1 | WP_011256925.1 |
| hemE, UROD | **K01599** | WP_041571524.1 | WP_167895693.1 |

| | | | |
|---|---|---|---|
| CPOX, hemF | **K00228** | WP_011256906.1 | WP_167896151.1 |
| hemJ | **K08973** | WP_041571401.1 | WP_167895936.1 |
| hemH, FECH | **K01772** | WP_011256916.1 | WP_011256916.1 |
| bfr | **K03594** | | WP_167896320.1 |
| COX10, ctaB, cyoE | **K02257** | WP_011256505.1 | WP_167895929.1 |
| COX15, ctaA | **K02259** | WP_050707669.1 | WP_167895834.1 |
| total | | 12 | 13 |
| **K00230 Purine metabolism** | | | |
| PRPS, prsA | **K00948** | WP_011256305.1 | WP_011256305.1 |
| purF, PPAT | **K00764** | WP_011256454.1 | WP_167896003.1 |
| purD | **K01945** | WP_011256663.1 | WP_167895827.1 |
| purN | **K11175** | WP_041571458.1 | WP_167896299.1 |
| purQ | **K23265** | WP_011256433.1 | WP_167896020.1 |
| purSL | **K23270** | WP_011256470.1 | WP_167895992.1 |
| purM | **K01933** | WP_041571547.1 | WP_167896025.1 |
| purE | **K01588** | WP_011256595.1 | WP_011256595.1 |
| purC | **K01923** | WP_011256428.1 | WP_011256428.1 |
| purB, ADSL | **K01756** | WP_011256701.1 | WP_167895794.1 |
| purH | **K00602** | WP_041571563.1 | WP_167896301.1 |
| ndk, NME | **K00940** | WP_011256914.1 | WP_167896157.1 |
| purA, ADSS | **K01939** | WP_011256472.1 | WP_167895990.1 |
| gmk | **K00942** | WP_011256778.1 | WP_011256778.1 |
| dgt | **K01129** | WP_011256227.1 | WP_011256227.1 |
| surE | **K03787** | WP_011256747.1 | WP_011256747.1 |
| nrdA, nrdE | **K00525** | WP_011256892.1 | WP_011256892.1 |
| nrdB, nrdF | **K00526** | WP_011256466.1 | WP_011256466.1 |
| guaA, GMPS | **K01951** | WP_011256641.1 | WP_167895843.1 |
| purK | **K01589** | WP_011256243.1 | WP_167895724.1 |
| IMPDH, guaB | **K00088** | WP_011256725.1 | WP_167895777.1 |
| total | | 21 | 21 |
| **K00240 Pyrimidine metabolism** | | | |
| carB, CPA2 | **K01955** | WP_011256710.1 | WP_167895786.1 |
| carA, CPA1 | **K01956** | WP_011256852.1 | WP_167896117.1 |
| pyrB, PYR2 | **K00609** | WP_011256583.1 | WP_011256583.1 |
| URA4, pyrC | **K01465** | WP_011256644.1 | WP_011256644.1 |
| DHODH, pyrD | **K00254** | WP_011256300.1 | WP_011256300.1 |
| pyrE | **K00762** | WP_011256987.1 | WP_011256987.1 |
| pyrF | **K01591** | WP_011256984.1 | WP_167896267.1 |
| surE | **K03787** | WP_011256747.1 | WP_011256747.1 |
| pyrH | **K09903** | WP_011257003.1 | WP_167896279.1 |
| ndk, NME | **K00940** | WP_011256914.1 | WP_167896157.1 |
| dcd | **K01494** | WP_011256491.1 | WP_167895976.1 |

| | | | |
|---|---|---|---|
| pyrG, CTPS | **K01937** | WP_011256370.1 | WP_167895942.1 |
| ndk, NME | **K00940** | WP_011256914.1 | WP_167896157.1 |
| E1.17.4.1A, nrdA, nrdE | **K00525** | WP_011256892.1 | WP_011256892.1 |
| E1.17.4.1B, nrdB, nrdF | **K00526** | WP_011256466.1 | WP_011256466.1 |
| dcd | **K01494** | WP_011256491.1 | WP_167895976.1 |
| dut, DUT | **K01520** | WP_041571555.1 | WP_167896308.1 |
| tmk, DTYMK | **K00943** | WP_011256349.1 | WP_167896044.1 |
| thyX, thy1 | **K03465** | WP_011256938.1 | WP_167896177.1 |
| total | | 19 | 19 |
| **K00750 Pyridoxine metabolism** | | | |
| pdxH, PNPO | **K00275** | | |
| pdxJ | **K03474** | | |
| total | | 0 | 0 |

**ays from Wolbachia genomes using KASS.**

| wCauA | wCfeJ | wCfeT | wCle | wCtub |
|---|---|---|---|---|
| QDH18146.1 | wCfeJ_00074 | wCfeT_00368 | WP_041045648.1 | wCtub_t1_00121 |
| QDH18693.1 | wCfeJ_00245 | wCfeT_01262 | WP_041045861.1 | wCtub_t1_00161 |
| QDH18495.1 | wCfeJ_00019 | wCfeT_00958 | WP_041045607.1 | |
| | wCfeJ_00020 | wCfeT_00959 | | |
| QDH18726.1 | wCfeJ_01020 | wCfeT_00162 | WP_041045236.1 | wCtub_t1_00434 |
| 4 | 5 | 5 | 4 | 3 |
| QDH18591.1 | | wCfeT_00027 | WP_041046499.1 | |
| QDH18687.1 | wCfeJ_00068 | wCfeT_00693 | WP_041045850.1 | wCtub_t1_00155 |
| QDH18735.1 | wCfeJ_00914 | wCfeT_00171 | WP_041045526.1 | wCtub_t1_00299 |
| QDH18637.1 | wCfeJ_00664 | wCfeT_00014 | WP_041045945.1 | wCtub_t1_00207 |
| QDH18442.1 | wCfeJ_00607 | wCfeT_00072 | WP_041046143.1 | wCtub_t1_00472 |
| QDH18875.1 | wCfeJ_00117 | wCfeT_00090 | WP_041046413.1 | wCtub_t1_00641 |
| 6 | 5 | 6 | 6 | 5 |
| | | | WP_041046000.1 | |
| QDH18260.1 | wCfeJ_00563 | wCfeT_00486 | WP_041046165.1 | wCtub_t1_00663 |
| QDH18738.1 | wCfeJ_00911 | wCfeT_01005 | WP_041045521.1 | wCtub_t1_00301 |
| QDH19233.1 | wCfeJ_00191 | wCfeT_00885 | WP_041046228.1 | wCtub_t1_00569 |
| QDH18549.1 | wCfeJ_00446 | wCfeT_01287 | WP_041046022.1 | wCtub_t1_00491 |
| | | wCfeT_01265 | WP_052463260.1 | |
| | | wCfeT_01269 | WP_041045996.1 | |
| | | wCfeT_01268 | WP_041045998.1 | |
| | | wCfeT_01264 | WP_041046003.1 | |
| QDH18166.1 | wCfeJ_00395 | wCfeT_00808 | WP_041045174.1 | wCtub_t1_00145 |
| 5 | 5 | 9 | 10 | 5 |
| QDH19181.1 | wCfeJ_00103 | wCfeT_00563 | WP_041046382.1 | wCtub_t1_00606 |
| QDH18153.1 | wCfeJ_00525 | wCfeT_01176 | WP_041045411.1 | wCtub_t1_00150 |
| QDH18260.1 | wCfeJ_00563 | wCfeT_00486 | WP_041046165.1 | wCtub_t1_00663 |
| QDH18738.1 | wCfeJ_00911 | wCfeT_01005 | WP_041045521.1 | wCtub_t1_00301 |
| QDH19233.1 | wCfeJ_00191 | wCfeT_00885 | WP_041046228.1 | wCtub_t1_00569 |
| QDH18549.1 | wCfeJ_00446 | wCfeT_01287 | WP_041046022.1 | wCtub_t1_00491 |
| QDH18242.1 | wCfeJ_00136 | wCfeT_00737 | WP_041045454.1 | wCtub_t1_00517 |
| 7 | 7 | 7 | 7 | 7 |
| QDH18591.1 | | wCfeT_00027 | WP_041046499.1 | |
| QDH19194.1 | wCfeJ_00748 | | WP_041046394.1 | wCtub_t1_00386 |

| | | | | |
|---|---|---|---|---|
| QDH19193.1 | wCfeJ_00747 | | WP_041046396.1 | wCtub_t1_00385 |
| QDH19257.1 | wCfeJ_00338 | wCfeT_01555 | | wCtub_t1_00180 |
| QDH19192.1 | wCfeJ_00746 | | WP_041046398.1 | wCtub_t1_00384 |
| 5 | 4 | 2 | 4 | 4 |
| QDH18501.1 | wCfeJ_00712 | wCfeT_01435 | WP_041045017.1 | wCtub_t1_00050 |
| QDH19118.1 | wCfeJ_00402 | wCfeT_00700 | WP_041046248.1 | |
| QDH18548.1 | wCfeJ_00752 | wCfeT_00908 | WP_041044998.1 | wCtub_t1_00363 |
| QDH18640.1 | wCfeJ_00661 | wCfeT_00833 | WP_041045951.1 | wCtub_t1_00210 |
| QDH18579.1 | | | WP_041046716.1 | wCtub_t1_00091 |
| QDH19088.1 | wCfeJ_00877 | wCfeT_01419 | WP_041045280.1 | wCtub_t1_00277 |
| QDH18725.1 | wCfeJ_01021 | wCfeT_00161 | WP_041046552.1 | wCtub_t1_00435 |
| QDH19026.1 | wCfeJ_00399 | wCfeT_00921 | WP_041045884.1 | wCtub_t1_00588 |
| QDH18638.1 | wCfeJ_00663 | wCfeT_00015 | WP_041045947.1 | wCtub_t1_00208 |
| QDH19019.1 | wCfeJ_00621 | wCfeT_00190 | WP_041046459.1 | wCtub_t1_00511 |
| QDH19153.1 | wCfeJ_00532 | wCfeT_00432 | WP_041045901.1 | wCtub_t1_00528 |
| QDH18533.1 | wCfeJ_00695 | wCfeT_01189 | WP_041045575.1 | wCtub_t1_00244 |
| QDH19237.1 | wCfeJ_00187 | wCfeT_01104 | WP_041045807.1 | wCtub_t1_00566 |
| QDH18480.1 | wCfeJ_00646 | wCfeT_00851 | WP_006013840.1 | wCtub_t1_00026 |
| QDH19075.1 | wCfeJ_00903 | wCfeT_00608 | WP_041045428.1 | wCtub_t1_00309 |
| QDH19171.1 | wCfeJ_00239 | wCfeT_01125 | WP_041046206.1 | wCtub_t1_00729 |
| QDH18223.1 | wCfeJ_00360 | wCfeT_00025 | WP_041044918.1 | wCtub_t1_00198 |
| QDH18589.1 | wCfeJ_00675 | wCfeT_00193 | WP_041045008.1 | wCtub_t1_00477 |
| QDH19166.1 | wCfeJ_00234 | wCfeT_01127 | WP_041046210.1 | wCtub_t1_00723 |
| QDH19167.1 | wCfeJ_00235 | wCfeT_01128 | WP_041046212.1 | wCtub_t1_00725 |
| QDH19168.1 | wCfeJ_00236 | wCfeT_01129 | WP_041046214.1 | wCtub_t1_00726 |
| QDH19169.1 | wCfeJ_00237 | wCfeT_01130 | WP_041046216.1 | wCtub_t1_00727 |
| QDH18590.1 | wCfeJ_00271 | wCfeT_00026 | WP_041044916.1 | wCtub_t1_00197 |
| QDH19127.1 | wCfeJ_00676 | wCfeT_00505 | WP_041046014.1 | wCtub_t1_00585 |
| QDH18588.1 | wCfeJ_00674 | wCfeT_00024 | WP_041044920.1 | wCtub_t1_00199 |
| QDH18244.1 | wCfeJ_00238 | wCfeT_01072 | WP_041045460.1 | wCtub_t1_00519 |
| QDH19170.1 | wCfeJ_00588 | wCfeT_01126 | WP_041046208.1 | wCtub_t1_00728 |
| QDH18587.1 | wCfeJ_00673 | wCfeT_00023 | WP_041044922.1 | wCtub_t1_00200 |
| QDH18586.1 | wCfeJ_00672 | wCfeT_00022 | WP_041044924.1 | wCtub_t1_00201 |
| 29 | 28 | 28 | 29 | 28 |
| QDH18203.1 | wCfeJ_00545 | wCfeT_00065 | WP_041045630.1 | wCtub_t1_00171 |
| QDH18204.1 | wCfeJ_00544 | wCfeT_00066 | WP_041045628.1 | wCtub_t1_00172 |
| QDH18205.1 | wCfeJ_00543 | wCfeT_00067 | WP_041045626.1 | wCtub_t1_00173 |
| QDH18667.1 | wCfeJ_00359 | wCfeT_00049 | WP_041045597.1 | wCtub_t1_00476 |
| QDH18894.1 | wCfeJ_00047 | wCfeT_00479 | WP_041046177.1 | wCtub_t1_00657 |

| | | | | |
|---|---|---|---|---|
| QDH18160.1 | wCfeJ_00220 | wCfeT_01300 | WP_041045789.1 | wCtub_t1_00494 |
| QDH18472.1 | wCfeJ_00218 | wCfeT_00037 | WP_041046491.1 | wCtub_t1_00052 |
| QDH18473.1 | wCfeJ_00217 | wCfeT_01298 | WP_041046489.1 | wCtub_t1_00053 |
| QDH18156.1 | wCfeJ_00445 | wCfeT_01038 | WP_041046024.1 | wCtub_t1_00490 |
| QDH18171.1 | wCfeJ_00390 | wCfeT_00803 | WP_041045164.1 | wCtub_t1_00140 |
| QDH18170.1 | wCfeJ_00391 | wCfeT_00804 | WP_041045166.1 | wCtub_t1_00141 |
| QDH18169.1 | wCfeJ_00392 | wCfeT_00805 | WP_041045168.1 | wCtub_t1_00142 |
| QDH18168.1 | wCfeJ_00393 | wCfeT_00806 | WP_041045170.1 | wCtub_t1_00143 |
| QDH18167.1 | wCfeJ_00394 | wCfeT_00807 | WP_041045172.1 | wCtub_t1_00144 |
| QDH18285.1 | wCfeJ_00005 | wCfeT_01066 | WP_041045380.1 | wCtub_t1_00421 |
| QDH18286.1 | wCfeJ_00004 | | WP_041045378.1 | wCtub_t1_00420 |
| QDH19062.1 | wCfeJ_00305 | wCfeT_00717 | WP_041045103.1 | wCtub_t1_00673 |
| QDH18901.1 | wCfeJ_00054 | wCfeT_01423 | WP_041044980.1 | wCtub_t1_00693 |
| QDH18733.1 | wCfeJ_00916 | wCfeT_00169 | WP_041045530.1 | wCtub_t1_00297 |
| QDH19309.1 | wCfeJ_00635 | wCfeT_00847 | WP_041046666.1 | wCtub_t1_00016 |
| QDH18293.1 | wCfeJ_00313 | wCfeT_00777 | WP_041045916.1 | wCtub_t1_00102 |
| QDH18732.1 | wCfeJ_00917 | wCfeT_00168 | WP_041045532.1 | wCtub_t1_00296 |
| QDH18567.1 | wCfeJ_00636 | wCfeT_00848 | WP_041045689.1 | wCtub_t1_00017 |
| QDH19052.1 | wCfeJ_00335 | wCfeT_00110<br>wCfeT_00121 | WP_041045659.1 | wCtub_t1_00078 |
| QDH19054.1 | wCfeJ_00332 | wCfeT_00112 | WP_041045653.1 | wCtub_t1_00080 |
| QDH19055.1 | wCfeJ_00333 | wCfeT_00113<br>wCfeT_00123<br>wCfeT_00124 | WP_041045655.1 | wCtub_t1_00081 |
| QDH19053.1 | wCfeJ_00334 | wCfeT_00111<br>wCfeT_00122 | WP_041045657.1 | wCtub_t1_00079 |
| QDH18266.1 | wCfeJ_00252 | wCfeT_00086 | WP_041045911.1 | wCtub_t1_00548 |
| QDH19241.1 | wCfeJ_00742 | wCfeT_00818 | WP_041045067.1 | wCtub_t1_00377 |
| QDH19373.1 | wCfeJ_00743 | wCfeT_00819 | WP_041046525.1 | wCtub_t1_00378 |
| QDH18752.1 | wCfeJ_00907 | wCfeT_01387 | WP_041045518.1 | wCtub_t1_00306 |
| QDH18229.1 | wCfeJ_00011 | wCfeT_00954 | WP_041045326.1 | wCtub_t1_00428 |
| QDH19057.1 | wCfeJ_00330 | wCfeT_00131 | WP_052463207.1 | wCtub_t1_00038 |
| QDH18493.1 | wCfeJ_00018 | wCfeT_01045 | WP_041045605.1 | wCtub_t1_00269 |
| QDH18751.1 | wCfeJ_00908 | wCfeT_01388 | WP_041045519.1 | wCtub_t1_00305 |
| QDH18750.1 | wCfeJ_00909 | wCfeT_01389 | WP_041045520.1 | wCtub_t1_00303 |
| QDH18890.1 | wCfeJ_00801 | | | |
| QDH19341.1 | wCfeJ_00802 | | | |

| | | | | |
|---|---|---|---|---|
| 38 | 38 | 39 | 36 | 36 |
| QDH18761.1 | wCfeJ_00814 | wCfeT_01099 | WP_041046749.1 | wCtub_t1_00226 |
| QDH18834.1 | wCfeJ_00982 | wCfeT_00316 | WP_041045840.1 | |
| QDH18905.1 | wCfeJ_00057 | wCfeT_00103 | WP_041044986.1 | |
| QDH18539.1 | wCfeJ_00823 | wCfeT_00082 | WP_041045083.1 | wCtub_t1_00084 |
| QDH18466.1 | wCfeJ_00034 | wCfeT_00369 | WP_041045268.1 | |
| QDH18639.1 | wCfeJ_00662 | wCfeT_00016 | WP_041045949.1 | |
| QDH18624.1 | wCfeJ_00416 | wCfeT_01230 | | |
| QDH19230.1 | wCfeJ_00194 | wCfeT_00888 | WP_041046729.1 | wCtub_t1_00572 |
| QDH18650.1 | wCfeJ_00787 | wCfeT_00533 | | wCtub_t1_00224 |
| QDH18765.1 | wCfeJ_00821 | wCfeT_00639 | WP_041046352.1 | wCtub_t1_00232 |
| QDH18764.1 | wCfeJ_00820 | wCfeT_00640 | WP_041046354.1 | wCtub_t1_00231 |
| QDH18593.1 | wCfeJ_00679 | wCfeT_00995 | WP_041044912.1 | wCtub_t1_00393 |
| QDH18777.1 | wCfeJ_00061 | wCfeT_01400 | WP_041045127.1 | wCtub_t1_00325 |
| QDH18766.1 | wCfeJ_00822 | wCfeT_00638 | WP_041046350.1 | wCtub_t1_00233 |
| 14 | 14 | 14 | 12 | 9 |
| QDH18808.1 | wCfeJ_00079 | wCfeT_00788 | WP_041046660.1 | wCtub_t1_00735 |
| QDH18863.1 | wCfeJ_00460 | wCfeT_00697 | WP_041046254.1 | wCtub_t1_00118 |
| QDH19258.1 | wCfeJ_00244 | wCfeT_01155 | WP_041045712.1 | wCtub_t1_00578 |
| QDH18623.1 | wCfeJ_00418 | wCfeT_00070 | WP_041046102.1 | wCtub_t1_00699 |
| QDH18651.1 | wCfeJ_01046 | wCfeT_00917 | WP_041045191.1 | wCtub_t1_00085 |
| QDH18250.1 | wCfeJ_00597 | wCfeT_00292 | WP_041046430.1 | wCtub_t1_00522 |
| QDH18141.1 | wCfeJ_01064 | wCfeT_00005 | WP_041045260.1 | wCtub_t1_00350 |
| QDH19338.1 | wCfeJ_00843 | wCfeT_01549 | WP_041046477.1 | wCtub_t1_00255 |
| QDH18214.1 | wCfeJ_00111 | wCfeT_00862 | WP_041046324.1 | wCtub_t1_00647 |
| QDH18215.1 | wCfeJ_00112 | wCfeT_00861 | WP_041046322.1 | wCtub_t1_00646 |
| QDH18492.1 | wCfeJ_00017 | wCfeT_01046 | WP_041045603.1 | wCtub_t1_00270 |
| QDH19134.1 | wCfeJ_00406 | wCfeT_00248 | WP_041044937.1 | |
| 12 | 12 | 12 | 12 | 11 |
| QDH19159.1 | wCfeJ_00355 | wCfeT_00407 | WP_041046376.1 | |
| QDH18193.1 | wCfeJ_00165 | wCfeT_00820 | WP_041046008.1 | |
| QDH18479.1 | wCfeJ_00647 | wCfeT_00852 | WP_041045636.1 | |
| QDH18822.1 | wCfeJ_01014 | wCfeT_00705 | WP_041045248.1 | wCtub_t1_00429 |
| QDH19308.1 | wCfeJ_00634 | wCfeT_01134 | WP_041046667.1 | wCtub_t1_00015 |
| QDH18125.1 | wCfeJ_00369 | wCfeT_01086 | WP_041045776.1 | wCtub_t1_00617 |

| | | | | |
|---|---|---|---|---|
| QDH18815.1 | wCfeJ_00991 | wCfeT_01147 | WP_041045036.1 | wCtub_t1_00284 |
| QDH18289.1 | wCfeJ_01055 | wCfeT_00045 | WP_041046260.1 | wCtub_t1_00418 |
| QDH18690.1 | wCfeJ_00071 | wCfeT_01020 | WP_041045856.1 | wCtub_t1_00158 |
| QDH18789.1 | wCfeJ_00199 | wCfeT_01309 | WP_041046697.1 | wCtub_t1_00138 |
| QDH19224.1 | wCfeJ_00715 | wCfeT_01438 | WP_041045010.1 | wCtub_t1_00068 |
| QDH19038.1 | wCfeJ_00515 | wCfeT_00680 | WP_041045869.1 | wCtub_t1_00483 |
| | wCfeJ_00339 | wCfeT_01554 | WP_041046505.1 | |
| QDH18222.1 | wCfeJ_00382 | wCfeT_01058 | WP_041046042.1 | |
| QDH18151.1 | wCfeJ_00311 | wCfeT_01531 | WP_041045181.1 | |
| QDH18150.1 | wCfeJ_00312 | wCfeT_01530 | WP_041045178.1 | |
| QDH19211.1 | wCfeJ_00740 | wCfeT_01243 | WP_041045061.1 | wCtub_t1_00359 |
| QDH18770.1 | wCfeJ_00996 | wCfeT_00221 | WP_041045133.1 | wCtub_t1_00281 |
| QDH19212.1 | wCfeJ_00739 | wCfeT_01242 | WP_041045063.1 | wCtub_t1_00360 |
| QDH18270.1 | wCfeJ_00689 | wCfeT_00521 | WP_041045585.1 | |
| 19 | 20 | 20 | 20 | 12 |
| QDH18853.1 | wCfeJ_00827 | | WP_041045073.1 | wCtub_t1_00039 |
| QDH18897.1 | wCfeJ_00050 | wCfeT_00354 | WP_041044970.1 | wCtub_t1_00697 |
| QDH19376.1 | wCfeJ_00284 | wCfeT_00364 | WP_041045704.1 | wCtub_t1_00649 |
| QDH18184.1 | wCfeJ_00701 | wCfeT_00914 | WP_041044962.1 | wCtub_t1_00237 |
| QDH18178.1 | | wCfeT_01190 | | |
| | wCfeJ_00211 | | | |
| QDH19032.1 | wCfeJ_00794 | wCfeT_00418 | WP_041046114.1 | wCtub_t1_00407 |
| QDH19260.1 | wCfeJ_00287 | wCfeT_00402 | WP_041045708.1 | wCtub_t1_00651 |
| QDH19259.1 | wCfeJ_00288 | wCfeT_00401 | WP_041045710.1 | wCtub_t1_00652 |
| QDH19107.1 | | | WP_041045757.1 | wCtub_t1_00373 |
| QDH18898.1 | wCfeJ_00051 | wCfeT_00355 | WP_041044972.1 | wCtub_t1_00696 |
| 10 | 9 | 8 | 9 | 9 |
| QDH18628.1 | wCfeJ_00410 | wCfeT_00205 | WP_041046044.1 | wCtub_t1_00705 |
| QDH18860.1 | wCfeJ_00327 | wCfeT_00683 | WP_041045649.1 | wCtub_t1_00036 |
| QDH19060.1 | wCfeJ_00216 | wCfeT_01538 | WP_041046680.1 | wCtub_t1_00677 |
| QDH18474.1 | wCfeJ_00485 | wCfeT_01033 | WP_041045162.1 | wCtub_t1_00120 |
| QDH19022.1 | wCfeJ_01100 | wCfeT_00420 | WP_041045183.1 | wCtub_t1_00364 |
| QDH18143.1 | wCfeJ_00805 | wCfeT_00775 | WP_041046238.1 | wCtub_t1_00345 |
| QDH19272.1 | wCfeJ_00259 | wCfeT_01286 | WP_041046675.1 | wCtub_t1_00625 |

| | | | | |
|---|---|---|---|---|
| QDH18278.1 | wCfeJ_00971 | wCfeT_01007 | WP_041044932.1 | wCtub_t1_00318 |
| QDH19044.1 | wCfeJ_00988 | wCfeT_00460 | WP_041045043.1 | wCtub_t1_00287 |
| QDH18253.1 | wCfeJ_01070 | wCfeT_00350 | WP_041045057.1 | wCtub_t1_00356 |
| QDH18305.1 | wCfeJ_00209 | wCfeT_01200 | WP_041046194.1 | |
| QDH18752.1 | wCfeJ_00907 | wCfeT_01387 | WP_041045518.1 | wCtub_t1_00306 |
| QDH19057.1 | wCfeJ_00330 | wCfeT_00131 | WP_052463207.1 | wCtub_t1_00038 |
| 13 | 13 | 13 | 13 | 12 |
| QDH18522.1 | wCfeJ_00273 | wCfeT_01312 | WP_041046242.1 | wCtub_t1_00592 |
| QDH18218.1 | wCfeJ_00353 | wCfeT_00896 | WP_041046402.1 | wCtub_t1_00192 |
| QDH18525.1 | wCfeJ_00075 | wCfeT_00234 | WP_041046425.1 | wCtub_t1_00633 |
| QDH18871.1 | wCfeJ_00123 | wCfeT_00937 | WP_041046407.1 | wCtub_t1_00637 |
| QDH18126.1 | wCfeJ_00368 | wCfeT_01085 | WP_041045778.1 | wCtub_t1_00616 |
| QDH19199.1 | wCfeJ_00152 | wCfeT_00684 | WP_041045388.1 | wCtub_t1_00194 |
| QDH18121.1 | wCfeJ_00373 | wCfeT_01290 | WP_041045768.1 | wCtub_t1_00621 |
| QDH18610.1 | wCfeJ_00420 | wCfeT_00675 | WP_041046091.1 | wCtub_t1_00104 |
| QDH18122.1 | wCfeJ_00372 | wCfeT_01293 | WP_041045770.1 | wCtub_t1_00620 |
| QDH18854.1 | | | | |
| QDH19179.1 | wCfeJ_00810 | wCfeT_01319 | WP_041046184.1 | wCtub_t1_00732 |
| QDH18251.1 | wCfeJ_01072 | wCfeT_00549 | WP_041045059.1 | wCtub_t1_00358 |
| QDH19332.1 | wCfeJ_00682 | wCfeT_00391 | WP_041046767.1 | wCtub_t1_00395 |
| QDH19064.1 | wCfeJ_00479 | wCfeT_01440 | WP_041046741.1 | wCtub_t1_00066 |
| QDH18688.1 | wCfeJ_00069 | wCfeT_00692 | WP_041045852.1 | wCtub_t1_00156 |
| QDH19238.1 | wCfeJ_00751 | wCfeT_00476 | WP_041045599.1 | wCtub_t1_00388 |
| QDH18562.1 | wCfeJ_00629 | wCfeT_01080 | WP_041045695.1 | wCtub_t1_00008 |
| QDH18654.1 | wCfeJ_00611 | wCfeT_00924 | WP_041046461.1 | wCtub_t1_00510 |
| QDH18561.1 | wCfeJ_00127 | wCfeT_00467 | WP_041046429.1 | wCtub_t1_00634 |
| QDH18190.1 | wCfeJ_00163 | wCfeT_01117 | WP_041046004.1 | wCtub_t1_00559 |
| QDH18546.1 | wCfeJ_00354 | wCfeT_00907 | WP_041044994.1 | wCtub_t1_00070 |
| 21 | 20 | 20 | 20 | 20 |
| QDH18619.1 | wCfeJ_00918 | wCfeT_00084 | WP_041045384.1 | wCtub_t1_00414 |
| QDH18703.1 | wCfeJ_00495 | wCfeT_00137 | WP_041046065.1 | wCtub_t1_00459 |
| QDH19158.1 | wCfeJ_00357 | wCfeT_01008 | WP_041045966.1 | wCtub_t1_00601 |
| QDH18643.1 | wCfeJ_00215 | wCfeT_00308 | WP_041045721.1 | wCtub_t1_00676 |
| QDH18298.1 | wCfeJ_00131 | wCfeT_00570 | WP_041045394.1 | wCtub_t1_00668 |
| QDH18645.1 | wCfeJ_00755 | wCfeT_01151 | WP_041045002.1 | wCtub_t1_00362 |
| QDH19082.1 | wCfeJ_00925 | wCfeT_00403 | WP_041046286.1 | wCtub_t1_00320 |
| QDH19238.1 | wCfeJ_00751 | wCfeT_00476 | WP_041045599.1 | wCtub_t1_00388 |
| QDH19029.1 | wCfeJ_00797 | wCfeT_00415 | WP_041046121.1 | wCtub_t1_00410 |
| QDH18251.1 | wCfeJ_01072 | wCfeT_00549 | WP_041045059.1 | wCtub_t1_00358 |
| QDH18823.1 | wCfeJ_01009 | wCfeT_00973 | WP_041045442.1 | wCtub_t1_00327 |

| | | | | |
|---|---|---|---|---|
| QDH19089.1 | wCfeJ_00876 | wCfeT_01418 | WP_041045278.1 | wCtub_t1_00276 |
| QDH18251.1 | wCfeJ_01072 | wCfeT_00549 | WP_041045059.1 | wCtub_t1_00358 |
| QDH18562.1 | wCfeJ_00629 | wCfeT_01080 | WP_041045695.1 | wCtub_t1_00008 |
| QDH18654.1 | wCfeJ_00611 | wCfeT_00924 | WP_041046461.1 | wCtub_t1_00510 |
| QDH18823.1 | wCfeJ_01009 | wCfeT_00973 | WP_041045442.1 | wCtub_t1_00327 |
| QDH18634.1 | wCfeJ_00176 | wCfeT_00635 | WP_041046540.1 | wCtub_t1_00688 |
| QDH19288.1 | wCfeJ_00248 | wCfeT_01441 | WP_041046727.1 | wCtub_t1_00552 |
| QDH18263.1 | wCfeJ_00309 | wCfeT_01145 | WP_041045735.1 | wCtub_t1_00545 |
| 19 | 19 | 19 | 19 | 19 |

| | | | | |
|---|---|---|---|---|
| QDH19279.1 | wCfeJ_00522 | | WP_041045403.1 | wCtub_t1_00480 |
| QDH18504.1 | wCfeJ_00708 | | WP_041045024.1 | wCtub_t1_00047 |
| 2 | 2 | 0 | 2 | 2 |

| wDcau | wDi | wFol | wLbra | wLsig |
|---|---|---|---|---|
| wDcau_t1_00195 | wDi2_02130 | WP_110409687.1 | LOGDKPFN_00194 | wLs20_03220 |
| wDcau_t1_00229 | wDi2_02520 | WP_110410487.1 | LOGDKPFN_00887 | wLs20_06930 |
| | | WP_110409931.1 | | |
| | | WP_110409932.1 | | |
| wDcau_t1_00482 | wDi2_05540 | WP_110409453.1 | LOGDKPFN_00587 | wLs20_07400 |
| 3 | 3 | 5 | 3 | 3 |
| | wDi2_02950 | WP_110409421.1 | | wLs20_00530 |
| wDcau_t1_00223 | wDi2_02470 | WP_110409590.1 | LOGDKPFN_00881 | wLs20_06990 |
| wDcau_t1_00418 | wDi2_04020 | WP_110409444.1 | LOGDKPFN_00578 | wLs20_07310 |
| wDcau_t1_00276 | wDi2_03070 | WP_110410142.1 | LOGDKPFN_00425 | wLs20_00650 |
| wDcau_t1_00153 | WP_110409930.1 | LOGDKPFN_00983 | wLs20_07900 | WP_010962384.1 |
| wDcau_t1_00020 | wDi2_07790 | WP_110410546.1 | LOGDKPFN_01000 | wLs20_03330 |
| 5 | 6 | 6 | 5 | 6 |
| wDcau_t1_00033 | wDi2_07550 | WP_110409604.1 | LOGDKPFN_00107 | wLs20_03840 |
| wDcau_t1_00415 | wDi2_04050 | WP_110409528.1 | LOGDKPFN_00575 | wLs20_07280 |
| wDcau_t1_00061 | wDi2_00570 | WP_110409511.1 | LOGDKPFN_00921 | wLs20_08750 |
| wDcau_t1_00143 | wDi2_01460 | WP_110410193.1 | LOGDKPFN_00909 | wLs20_03390 |
| wDcau_t1_00214 | wDi2_02370 | WP_110409976.1 | LOGDKPFN_00078 | wLs20_04030 |
| 5 | 5 | 5 | 5 | 5 |
| wDcau_t1_00665 | wDi2_00220 | WP_112477189.1 | LOGDKPFN_00857 | wLs20_08430 |
| wDcau_t1_00219 | wDi2_02420 | WP_110409948.1 | LOGDKPFN_00071 | wLs20_04110 |
| wDcau_t1_00033 | wDi2_07550 | WP_110409604.1 | LOGDKPFN_00107 | wLs20_03840 |
| wDcau_t1_00415 | wDi2_04050 | WP_110409528.1 | LOGDKPFN_00575 | wLs20_07280 |
| wDcau_t1_00061 | wDi2_00570 | WP_110409511.1 | LOGDKPFN_00921 | wLs20_08750 |
| wDcau_t1_00143 | wDi2_01460 | WP_110410193.1 | LOGDKPFN_00909 | wLs20_03390 |
| wDcau_t1_00115 | wDi2_01180 | WP_110409562.1 | LOGDKPFN_00705 | wLs20_04850 |
| 7 | 7 | 7 | 7 | 7 |
| | wDi2_02950 | WP_110409421.1 | | wLs20_00530 |
| wDcau_t1_00294 | wDi2_04980 | | | |

| | | | | |
|---|---|---|---|---|
| wDcau_t1_00295 | wDi2_04970 | | | |
| wDcau_t1_00249 | wDi2_02730 | WP_110410545.1 | | |
| wDcau_t1_00296 | wDi2_04960 | | | |
| 4 | 5 | 2 | | 1 |
| wDcau_t1_00526 | wDi2_05990 | WP_110409432.1 | LOGDKPFN_00628 | wLs20_08250 |
| | | WP_110410356.1 | LOGDKPFN_00973 | wLs20_02520 |
| wDcau_t1_00312 | wDi2_04740 | WP_110409614.1 | LOGDKPFN_00464 | wLs20_01280 |
| wDcau_t1_00278 | wDi2_03090 | WP_110409898.1 | LOGDKPFN_00422 | wLs20_00630 |
| wDcau_t1_00180 | wDi2_01950 | WP_110409549.1 | LOGDKPFN_00234 | wLs20_02950 |
| wDcau_t1_00348 | wDi2_03790 | WP_110410302.1 | LOGDKPFN_00164 | wLs20_01850 |
| wDcau_t1_00483 | wDi2_05550 | WP_110409454.1 | LOGDKPFN_00588 | wLs20_07410 |
| wDcau_t1_00651 | wDi2_00390 | WP_110409595.1 | LOGDKPFN_00101 | wLs20_03890 |
| wDcau_t1_00277 | wDi2_03080 | WP_110410143.1 | LOGDKPFN_00424 | wLs20_00640 |
| wDcau_t1_00121 | wDi2_01300 | WP_110409731.1 | LOGDKPFN_00616 | wLs20_07710 |
| wDcau_t1_00104 | wDi2_01050 | WP_110409517.1 | LOGDKPFN_00693 | wLs20_06050 |
| wDcau_t1_00380 | wDi2_03450 | WP_110409254.1 | LOGDKPFN_00351 | wLs20_02070 |
| wDcau_t1_00064 | wDi2_00600 | WP_110409582.1 | LOGDKPFN_00831 | wLs20_06840 |
| wDcau_t1_00567 | wDi2_06470 | WP_110410593.1 | LOGDKPFN_00543 | wLs20_00320 |
| wDcau_t1_00408 | wDi2_04130 | WP_110410299.1 | LOGDKPFN_00495 | wLs20_01370 |
| wDcau_t1_00601 | wDi2_06860 | WP_110409799.1 | LOGDKPFN_00117 | wLs20_03710 |
| wDcau_t1_00149 | wDi2_01610 | WP_110409834.1 | LOGDKPFN_00018 | wLs20_00550 |
| wDcau_t1_00267 | wDi2_02970 | WP_110410531.1 | LOGDKPFN_00415 | wLs20_04540 |
| wDcau_t1_00603 | wDi2_06880 | WP_110409796.1 | LOGDKPFN_00119 | wLs20_03660 |
| wDcau_t1_00604 | wDi2_06890 | WP_110409797.1 | LOGDKPFN_00120 | wLs20_03670 |
| wDcau_t1_00605 | wDi2_06900 | WP_110410558.1 | LOGDKPFN_00121 | wLs20_03680 |
| wDcau_t1_00606 | wDi2_06910 | WP_112477239.1 | LOGDKPFN_00122 | wLs20_03690 |
| wDcau_t1_00047 | wDi2_00420 | WP_110409420.1 | LOGDKPFN_00247 | wLs20_00540 |
| wDcau_t1_00266 | wDi2_02960 | WP_110410455.1 | LOGDKPFN_00414 | wLs20_02830 |
| wDcau_t1_00268 | wDi2_02980 | WP_110409419.1 | LOGDKPFN_00416 | wLs20_00560 |
| wDcau_t1_00113 | wDi2_01160 | WP_110409723.1 | LOGDKPFN_00118 | wLs20_03700 |
| wDcau_t1_00602 | wDi2_06870 | WP_110409798.1 | LOGDKPFN_00942 | wLs20_04810 |
| wDcau_t1_00269 | wDi2_02990 | WP_110409418.1 | LOGDKPFN_00417 | wLs20_00570 |
| wDcau_t1_00270 | wDi2_03000 | WP_110409417.1 | LOGDKPFN_00418 | wLs20_00580 |
| 28 | 28 | 29 | 29 | 29 |
| wDcau_t1_00238 | wDi2_02640 | WP_110410184.1 | LOGDKPFN_00338 | wLs20_06200 |
| wDcau_t1_00239 | wDi2_02650 | WP_110410183.1 | LOGDKPFN_00337 | wLs20_06210 |
| wDcau_t1_00240 | wDi2_02660 | WP_110410182.1 | LOGDKPFN_00336 | wLs20_06220 |
| wDcau_t1_00148 | wDi2_01620 | WP_110409234.1 | LOGDKPFN_00778 | wLs20_03560 |
| wDcau_t1_00039 | wDi2_07620 | WP_110410158.1 | LOGDKPFN_00021 | wLs20_04510 |

| | | | | |
|---|---|---|---|---|
| wDcau_t1_00145 | wDi2_01430 | WP_110410174.1 | LOGDKPFN_00938 | wLs20_04750 |
| wDcau_t1_00541 | wDi2_06180 | WP_110409810.1 | LOGDKPFN_01005 | wLs20_05220 |
| wDcau_t1_00540 | wDi2_06170 | WP_110409808.1 | LOGDKPFN_01006 | wLs20_05210 |
| wDcau_t1_00142 | wDi2_01470 | WP_110409755.1 | LOGDKPFN_00908 | wLs20_03380 |
| wDcau_t1_00209 | wDi2_02320 | WP_110409970.1 | LOGDKPFN_00083 | wLs20_03980 |
| wDcau_t1_00210 | wDi2_02330 | WP_110409971.1 | LOGDKPFN_00082 | wLs20_03990 |
| wDcau_t1_00211 | wDi2_02340 | WP_110409972.1 | LOGDKPFN_00081 | wLs20_04000 |
| wDcau_t1_00212 | wDi2_02350 | WP_110409973.1 | LOGDKPFN_00080 | wLs20_04010 |
| wDcau_t1_00213 | wDi2_02360 | WP_110409974.1 | LOGDKPFN_00079 | wLs20_04020 |
| wDcau_t1_00469 | wDi2_05430 | WP_110409772.1 | LOGDKPFN_00742 | wLs20_00880 |
| wDcau_t1_00468 | wDi2_05420 | WP_110409773.1 | LOGDKPFN_00743 | wLs20_00890 |
| wDcau_t1_00650 | wDi2_07410 | WP_110409873.1 | LOGDKPFN_00041 | wLs20_04370 |
| wDcau_t1_00635 | wDi2_07230 | WP_110409860.1 | LOGDKPFN_00863 | wLs20_08880 |
| wDcau_t1_00420 | wDi2_04000 | WP_110409446.1 | LOGDKPFN_00580 | wLs20_07330 |
| wDcau_t1_00576 | wDi2_06560 | WP_110410534.1 | LOGDKPFN_00527 | wLs20_00200 |
| wDcau_t1_00170 | wDi2_01840 | WP_110410512.1 | LOGDKPFN_00404 | wLs20_00430 |
| wDcau_t1_00421 | wDi2_03990 | WP_110409447.1 | LOGDKPFN_00581 | wLs20_07340 |
| wDcau_t1_00575 | wDi2_06550 | WP_110409480.1 | LOGDKPFN_00528 | wLs20_00210 |
| wDcau_t1_00552 | wDi2_06310 | WP_110410255.1 | LOGDKPFN_00056 | wLs20_04240 |
| wDcau_t1_00554 | wDi2_06330 | WP_110410252.1 | LOGDKPFN_00053 | wLs20_04260 |
| wDcau_t1_00555 | wDi2_06340 | WP_110410253.1 | LOGDKPFN_00054 | wLs20_04270 |
| wDcau_t1_00553 | wDi2_06320 | WP_110410254.1 | LOGDKPFN_00055 | wLs20_04250 |
| wDcau_t1_00081 | wDi2_00830 | WP_110409374.1 | LOGDKPFN_00935 | wLs20_05780 |
| wDcau_t1_00299 | wDi2_04920 | WP_110409821.1 | LOGDKPFN_00379 | wLs20_02280 |
| wDcau_t1_00298 | wDi2_04930 | WP_110410562.1 | LOGDKPFN_00380 | wLs20_02290 |
| wDcau_t1_00411 | wDi2_04100 | WP_110409537.1 | LOGDKPFN_00472 | wLs20_01330 |
| wDcau_t1_00475 | wDi2_05490 | WP_110409692.1 | LOGDKPFN_00735 | wLs20_00820 |
| wDcau_t1_00556 | wDi2_06350 | WP_110410251.1 | LOGDKPFN_00050 | wLs20_04290 |
| wDcau_t1_00402 | wDi2_03700 | WP_110409895.1 | LOGDKPFN_00516 | wLs20_00090 |
| wDcau_t1_00412 | wDi2_04090 | WP_110409538.1 | LOGDKPFN_00471 | wLs20_01320 |
| wDcau_t1_00413 | wDi2_04080 | WP_110409539.1 | LOGDKPFN_00470 | wLs20_01310 |

| | | | | |
|---|---|---|---|---|
| 36 | 36 | 36 | 36 | 36 |
| wDcau_t1_00365 | wDi2_03270 | WP_110410532.1 | LOGDKPFN_00655 | wLs20_08000 |
| | | WP_110409641.1 | LOGDKPFN_00440 | |
| | wDi2_07250 | WP_110409902.1 | LOGDKPFN_00860 | wLs20_08910 |
| | wDi2_02000 | WP_110409427.1 | LOGDKPFN_00633 | wLs20_08200 |
| | wDi2_06230 | WP_110410210.1 | LOGDKPFN_00216 | wLs20_03090 |
| | | WP_110410144.1 | | |
| | | WP_110409598.1 | LOGDKPFN_00368 | wLs20_02180 |
| wDcau_t1_00058 | wDi2_00540 | WP_110409508.1 | LOGDKPFN_00924 | wLs20_08720 |
| wDcau_t1_00290 | wDi2_03240 | WP_110409226.1 | LOGDKPFN_00131 | wLs20_01590 |
| wDcau_t1_00371 | wDi2_03330 | WP_110409702.1 | LOGDKPFN_00661 | wLs20_08060 |
| wDcau_t1_00370 | wDi2_03320 | WP_110409703.1 | LOGDKPFN_00660 | wLs20_08050 |
| wDcau_t1_00360 | wDi2_05070 | WP_110409593.1 | LOGDKPFN_00411 | wLs20_00500 |
| wDcau_t1_00452 | wDi2_04350 | WP_110409984.1 | LOGDKPFN_00557 | wLs20_07150 |
| wDcau_t1_00372 | wDi2_03340 | WP_110409701.1 | LOGDKPFN_00662 | wLs20_08070 |
| 8 | 11 | 14 | 13 | 12 |
| | wDi2_06790 | WP_110409532.1 | LOGDKPFN_00777 | wLs20_03550 |
| wDcau_t1_00192 | wDi2_02100 | WP_110409575.1 | LOGDKPFN_00029 | wLs20_04470 |
| | | WP_110410451.1 | | wLs20_05870 |
| | | WP_110409675.1 | | |
| | | WP_110409777.1 | | |
| | | WP_110409781.1 | | |
| wDcau_t1_00324 | wDi2_04600 | WP_110410263.1 | LOGDKPFN_01016 | wLs20_08650 |
| wDcau_t1_00389 | wDi2_03550 | WP_110409962.1 | LOGDKPFN_00137 | wLs20_01600 |
| | | WP_110409770.1 | | |
| | | WP_110409769.1 | | |
| | | WP_110409896.1 | | wLs20_00110 |
| | wDi2_06820 | WP_112477178.1 | LOGDKPFN_00255 | wLs20_02750 |
| 3 | 5 | 12 | 5 | 7 |
| | wDi2_00250 | WP_110409389.1 | | |
| | wDi2_00730 | WP_110409663.1 | LOGDKPFN_00207 | wLs20_03140 |
| | | | LOGDKPFN_00208 | |
| | wDi2_06460 | WP_110410468.1 | LOGDKPFN_00541 | wLs20_00310 |
| wDcau_t1_00477 | wDi2_05500 | WP_110410334.1 | LOGDKPFN_00570 | wLs20_07230 |
| wDcau_t1_00577 | wDi2_06570 | WP_110409481.1 | LOGDKPFN_00526 | wLs20_00190 |
| wDcau_t1_00678 | wDi2_00090 | WP_110409647.1 | LOGDKPFN_00682 | wLs20_05950 |

| | | | | |
|---|---|---|---|---|
| wDcau_t1_00341 | wDi2_03860 | WP_110409954.1 | LOGDKPFN_00490 | wLs20_01420 |
| wDcau_t1_00465 | wDi2_05400 | WP_110410353.1 | LOGDKPFN_00154 | wLs20_01770 |
| wDcau_t1_00226 | wDi2_02500 | WP_110410541.1 | LOGDKPFN_00884 | wLs20_06960 |
| wDcau_t1_00207 | wDi2_02300 | WP_110409416.1 | LOGDKPFN_00259 | wLs20_02720 |
| wDcau_t1_00528 | wDi2_06010 | WP_110409364.1 | LOGDKPFN_00625 | wLs20_08290 |
| wDcau_t1_00136 | wDi2_01530 | WP_110409788.1 | LOGDKPFN_00375 | wLs20_02230 |
| | wDi2_02740 | WP_110409677.1 | LOGDKPFN_00945 | wLs20_05670 |
| | | | | wLs20_05690 |
| | | WP_110409250.1 | LOGDKPFN_00370 | |
| | | WP_110410529.1 | LOGDKPFN_00074 | wLs20_04080 |
| | | WP_110409375.1 | | |
| wDcau_t1_00316 | wDi2_04700 | WP_110409413.1 | LOGDKPFN_00726 | wLs20_09030 |
| wDcau_t1_00344 | wDi2_03830 | WP_110409829.1 | LOGDKPFN_00487 | wLs20_01500 |
| wDcau_t1_00315 | wDi2_04710 | WP_110409412.1 | LOGDKPFN_00725 | wLs20_09040 |
| | wDi2_03510 | WP_110409280.1 | LOGDKPFN_00811 | wLs20_05350 |
| 12 | 16 | 20 | 18 | 17 |
| wDcau_t1_00514 | wDi2_05880 | WP_110409407.1 | | |
| wDcau_t1_00631 | wDi2_07190 | WP_112477233.1 | LOGDKPFN_00866 | wLs20_08850 |
| wDcau_t1_00024 | wDi2_07710 | WP_110410530.1 | LOGDKPFN_00676 | wLs20_05930 |
| wDcau_t1_00374 | wDi2_03390 | WP_110410521.1 | LOGDKPFN_00005 | wLs20_04660 |
| | | WP_110409833.1 | | |
| | | WP_110410581.1 | | |
| wDcau_t1_00439 | wDi2_05280 | WP_110409887.1 | LOGDKPFN_00453 | wLs20_01160 |
| wDcau_t1_00026 | wDi2_07690 | WP_110410449.1 | LOGDKPFN_00672 | wLs20_05890 |
| wDcau_t1_00027 | wDi2_07680 | WP_110410450.1 | LOGDKPFN_00671 | wLs20_05880 |
| wDcau_t1_00303 | wDi2_04870 | WP_110409856.1 | LOGDKPFN_00994 | wLs20_01040 |
| wDcau_t1_00632 | wDi2_07200 | WP_110409863.1 | | |
| 9 | 9 | 11 | 7 | 7 |
| wDcau_t1_00624 | wDi2_07120 | WP_110409513.1 | LOGDKPFN_00363 | wLs20_02130 |
| wDcau_t1_00558 | wDi2_06370 | WP_110409937.1 | LOGDKPFN_00047 | wLs20_04320 |
| wDcau_t1_00647 | wDi2_07370 | WP_110410484.1 | LOGDKPFN_00934 | wLs20_05760 |
| wDcau_t1_00194 | wDi2_02120 | WP_110409806.1 | LOGDKPFN_00891 | wLs20_05180 |
| wDcau_t1_00311 | wDi2_04750 | WP_110409616.1 | LOGDKPFN_00158 | wLs20_01790 |
| wDcau_t1_00327 | wDi2_04550 | WP_110409698.1 | LOGDKPFN_00814 | wLs20_05320 |
| wDcau_t1_00002 | wDi2_00010 | WP_110409669.1 | LOGDKPFN_00317 | wLs20_06420 |

| | | | | |
|---|---|---|---|---|
| wDcau_t1_00428 | wDi2_04250 | WP_110409498.1 | LOGDKPFN_00549 | wLs20_00370 |
| wDcau_t1_00338 | wDi2_03890 | WP_110410277.1 | LOGDKPFN_00167 | wLs20_01870 |
| wDcau_t1_00319 | wDi2_04660 | WP_110409942.1 | LOGDKPFN_00729 | wLs20_09000 |
| | | WP_110409743.1 | LOGDKPFN_00820 | wLs20_06760 |
| wDcau_t1_00411 | wDi2_04100 | WP_110409537.1 | LOGDKPFN_00472 | wLs20_01330 |
| wDcau_t1_00556 | wDi2_06350 | WP_110410251.1 | LOGDKPFN_00050 | wLs20_04290 |
| 12 | 12 | 13 | 13 | 13 |

| | | | | |
|---|---|---|---|---|
| wDcau_t1_00654 | wDi2_00350 | WP_110409377.1 | LOGDKPFN_00903 | wLs20_05100 |
| wDcau_t1_00260 | wDi2_02870 | WP_110410305.1 | LOGDKPFN_00332 | wLs20_06260 |
| wDcau_t1_00010 | wDi2_07870 | WP_110410561.1 | LOGDKPFN_00062 | wLs20_04190 |
| wDcau_t1_00016 | wDi2_07830 | WP_110409673.1 | LOGDKPFN_00876 | wLs20_07080 |
| wDcau_t1_00677 | wDi2_00100 | WP_110409648.1 | LOGDKPFN_00681 | wLs20_05940 |
| wDcau_t1_00263 | wDi2_02900 | WP_110410539.1 | LOGDKPFN_00503 | wLs20_00020 |
| wDcau_t1_00682 | wDi2_00050 | WP_110410191.1 | LOGDKPFN_00687 | wLs20_06000 |
| wDcau_t1_00168 | wDi2_01820 | WP_110409920.1 | LOGDKPFN_00399 | wLs20_02430 |
| wDcau_t1_00681 | wDi2_00060 | WP_110409643.1 | LOGDKPFN_00686 | wLs20_05990 |
| | | WP_110409965.1 | | |
| wDcau_t1_00598 | wDi2_06830 | WP_110409952.1 | | wLs20_03290 |
| wDcau_t1_00317 | wDi2_04680 | WP_110409940.1 | LOGDKPFN_00727 | wLs20_09020 |
| wDcau_t1_00357 | wDi2_05100 | WP_110409230.1 | LOGDKPFN_00407 | wLs20_00450 |
| wDcau_t1_00530 | wDi2_06030 | WP_110409362.1 | LOGDKPFN_00623 | wLs20_08310 |
| wDcau_t1_00224 | wDi2_02480 | WP_110409589.1 | LOGDKPFN_00882 | wLs20_06980 |
| wDcau_t1_00291 | wDi2_05010 | WP_110410589.1 | LOGDKPFN_01018 | wLs20_07940 |
| wDcau_t1_00584 | wDi2_06630 | WP_110409368.1 | LOGDKPFN_00519 | wLs20_00130 |
| wDcau_t1_00122 | wDi2_01310 | WP_110410517.1 | LOGDKPFN_00554 | wLs20_07130 |
| wDcau_t1_00011 | wDi2_07860 | WP_110409734.1 | LOGDKPFN_00930 | wLs20_05730 |
| wDcau_t1_00091 | wDi2_00710 | | LOGDKPFN_00211 | wLs20_03120 |
| wDcau_t1_00543 | wDi2_06200 | WP_110410241.1 | LOGDKPFN_00718 | wLs20_04940 |
| 20 | 20 | 20 | 19 | 20 |

| | | | | |
|---|---|---|---|---|
| wDcau_t1_00446 | wDi2_05350 | WP_110409721.1 | LOGDKPFN_00753 | wLs20_00980 |
| wDcau_t1_00507 | wDi2_05780 | WP_110409476.1 | LOGDKPFN_00229 | wLs20_03000 |
| wDcau_t1_00661 | wDi2_00260 | WP_110410150.1 | LOGDKPFN_00852 | wLs20_08480 |
| wDcau_t1_00648 | wDi2_07390 | WP_110410493.1 | LOGDKPFN_00039 | wLs20_04390 |
| wDcau_t1_00028 | wDi2_07500 | WP_110409871.1 | LOGDKPFN_00894 | wLs20_05170 |
| wDcau_t1_00313 | wDi2_04730 | WP_110409640.1 | LOGDKPFN_00462 | wLs20_01290 |
| wDcau_t1_00329 | wDi2_04260 | WP_110410318.1 | LOGDKPFN_00752 | wLs20_00970 |
| wDcau_t1_00291 | wDi2_05010 | WP_110410589.1 | LOGDKPFN_01018 | wLs20_07940 |
| wDcau_t1_00442 | wDi2_05310 | WP_110409892.1 | LOGDKPFN_00457 | wLs20_01190 |
| wDcau_t1_00317 | wDi2_04680 | WP_110409940.1 | LOGDKPFN_00727 | wLs20_09020 |
| wDcau_t1_00332 | wDi2_04310 | WP_110409218.1 | LOGDKPFN_00567 | wLs20_07220 |

| | | | | |
|---|---|---|---|---|
| wDcau_t1_00349 | wDi2_03780 | WP_110410303.1 | LOGDKPFN_00163 | wLs20_01840 |
| wDcau_t1_00317 | wDi2_04680 | WP_110409940.1 | LOGDKPFN_00727 | wLs20_09020 |
| wDcau_t1_00584 | wDi2_06630 | WP_110409368.1 | LOGDKPFN_00519 | wLs20_00130 |
| wDcau_t1_00122 | wDi2_01310 | WP_110410517.1 | LOGDKPFN_00554 | wLs20_07130 |
| wDcau_t1_00332 | wDi2_04310 | WP_110409218.1 | LOGDKPFN_00567 | wLs20_07220 |
| wDcau_t1_00639 | wDi2_07280 | WP_110409768.1 | LOGDKPFN_00976 | wLs20_07860 |
| wDcau_t1_00085 | wDi2_00790 | WP_110409817.1 | LOGDKPFN_00816 | wLs20_06720 |
| wDcau_t1_00078 | wDi2_00890 | WP_110409558.1 | LOGDKPFN_00109 | wLs20_03820 |
| 19 | 19 | 19 | 19 | 19 |

| | | | | |
|---|---|---|---|---|
| wDcau_t1_00132 | wDi2_01590 | WP_110410165.1 | LOGDKPFN_00956 | |
| wDcau_t1_00522 | wDi2_05950 | WP_110409428.1 | | |
| 2 | 2 | 2 | 1 | 0 |

| wLug | wMel | wMhie | wNfla | wOo |
|---|---|---|---|---|
| wLug_00451 | WP_010081988.1 | KIHBGAEE_00515 | LKKDFFFJ_00530 | wOo_06060 |
| wLug_00581 | WP_010962973.1 | KIHBGAEE_00569 | LKKDFFFJ_01116 | wOo_06420 |
| wLug_00292 | WP_010962390.1 | | LKKDFFFJ_01228 | |
| wLug_00293 | | | | |
| wLug_00237 | WP_010962746.1 | KIHBGAEE_00346 | LKKDFFFJ_00183 | wOo_07920 |
| 5 | 4 | 3 | 4 | 3 |
| wLug_00148 | WP_010082649.1 | | LKKDFFFJ_00254 | wOo_05610 |
| wLug_00575 | WP_010962779.1 | KIHBGAEE_00521 | LKKDFFFJ_01110 | |
| wLug_00228 | WP_010962739.1 | KIHBGAEE_00681 | LKKDFFFJ_00175 | wOo_06530 |
| wLug_01192 | WP_006279647.1 | KIHBGAEE_00541 | LKKDFFFJ_00418 | wOo_05500 |
| wLug_00306 | KIHBGAEE_00823 | | LKKDFFFJ_00376 | |
| wLug_00777 | WP_010082493.1 | KIHBGAEE_00977 | LKKDFFFJ_00080 | wOo_03530 |
| 6 | 6 | 4 | 6 | 4 |
| wLug_01483 | | | LKKDFFFJ_00584 | |
| wLug_01404 | WP_010963089.1 | KIHBGAEE_00613 | LKKDFFFJ_00346 | wOo_03770 |
| wLug_00225 | WP_010962736.1 | KIHBGAEE_00684 | LKKDFFFJ_00172 | wOo_06500 |
| wLug_00404 | WP_010082082.1 | KIHBGAEE_00100 | LKKDFFFJ_00462 | wOo_02740 |
| wLug_01351 | WP_006280349.1 | KIHBGAEE_00857 | LKKDFFFJ_01204 | wOo_01730 |
| wLug_01485 | | | LKKDFFFJ_00582 | |
| wLug_01481 | | | LKKDFFFJ_00586 | |
| wLug_01482 | | | LKKDFFFJ_00585 | |
| wLug_01486 | | | LKKDFFFJ_00581 | |
| wLug_00471 | WP_010082374.1 | KIHBGAEE_00256 | LKKDFFFJ_00556 | wOo_06210 |
| 10 | 5 | 5 | 10 | 5 |
| wLug_00738 | WP_010962894.1 | KIHBGAEE_00246 | LKKDFFFJ_00750 | wOo_03130 |
| wLug_00413 | WP_010962965.1 | KIHBGAEE_00784 | LKKDFFFJ_00539 | wOo_06160 |
| wLug_01404 | WP_010963089.1 | KIHBGAEE_00613 | LKKDFFFJ_00346 | wOo_03770 |
| wLug_00225 | WP_010962736.1 | KIHBGAEE_00684 | LKKDFFFJ_00172 | wOo_06500 |
| wLug_00404 | WP_010082082.1 | KIHBGAEE_00100 | LKKDFFFJ_00462 | wOo_02740 |
| wLug_01351 | WP_006280349.1 | KIHBGAEE_00857 | LKKDFFFJ_01204 | wOo_01730 |
| wLug_01276 | WP_010082023.1 | KIHBGAEE_00948 | LKKDFFFJ_00325 | wOo_02040 |
| 7 | 7 | 7 | 7 | 7 |
| wLug_00148 | WP_010082649.1 | | LKKDFFFJ_00254 | wOo_05610 |
| wLug_01317 | | KIHBGAEE_00239 | LKKDFFFJ_00086 | wOo_05290 |

| | | | | |
|---|---|---|---|---|
| wLug_00511 | | KIHBGAEE_00238 | LKKDFFFJ_00087 | wOo_05280 |
| wLug_01316 | WP_010963010.1 | KIHBGAEE_00169 | LKKDFFFJ_01246 | wOo_05830 |
| wLug_01315 | WP_010962904.1 | KIHBGAEE_00237 | LKKDFFFJ_00959 | wOo_05270 |
| 5 | 3 | 4 | 5 | 5 |

| | | | | |
|---|---|---|---|---|
| wLug_01251 | WP_010082597.1 | KIHBGAEE_00027 | LKKDFFFJ_01165 | wOo_00090 |
| wLug_00880 | WP_022626270.1 | KIHBGAEE_00080 | LKKDFFFJ_01072 | |
| wLug_01337 | WP_010962357.1 | KIHBGAEE_00723 | LKKDFFFJ_01205 | wOo_05050 |
| wLug_01195 | WP_010082185.1 | KIHBGAEE_00996 | LKKDFFFJ_00421 | wOo_05480 |
| | WP_006279543.1 | KIHBGAEE_00215 | | wOo_01230 |
| wLug_00169 | WP_010962615.1 | KIHBGAEE_00563 | LKKDFFFJ_00681 | wOo_06770 |
| wLug_00238 | WP_010962747.1 | KIHBGAEE_00345 | LKKDFFFJ_00184 | wOo_07940 |
| wLug_00163 | WP_010081840.1 | KIHBGAEE_00145 | LKKDFFFJ_00398 | wOo_02930 |
| wLug_01193 | WP_010962455.1 | KIHBGAEE_00540 | LKKDFFFJ_00419 | wOo_05490 |
| wLug_00864 | WP_022626296.1 | KIHBGAEE_00827 | LKKDFFFJ_01073 | wOo_01950 |
| wLug_00135 | WP_010962907.1 | KIHBGAEE_00816 | LKKDFFFJ_00813 | wOo_02240 |
| wLug_00346 | WP_006279761.1 | KIHBGAEE_00558 | LKKDFFFJ_00825 | wOo_07150 |
| wLug_00408 | WP_010963030.1 | KIHBGAEE_00671 | LKKDFFFJ_00465 | wOo_02710 |
| wLug_00946 | WP_006013840.1 | KIHBGAEE_00689 | LKKDFFFJ_00273 | wOo_00560 |
| wLug_01040 | WP_010962605.1 | KIHBGAEE_00513 | LKKDFFFJ_00613 | wOo_04430 |
| wLug_01246 | WP_010962887.1 | KIHBGAEE_00002 | LKKDFFFJ_00740 | wOo_01000 |
| wLug_00146 | WP_010082330.1 | KIHBGAEE_00032 | LKKDFFFJ_00252 | wOo_01580 |
| wLug_00440 | WP_010962300.1 | KIHBGAEE_00108 | LKKDFFFJ_00668 | wOo_05590 |
| wLug_01241 | WP_010962882.1 | KIHBGAEE_00004 | LKKDFFFJ_00735 | wOo_01020 |
| wLug_01242 | WP_010962883.1 | KIHBGAEE_00005 | LKKDFFFJ_00736 | wOo_01030 |
| wLug_01243 | WP_010962884.1 | KIHBGAEE_00006 | LKKDFFFJ_00737 | wOo_01040 |
| wLug_01244 | WP_010962885.1 | KIHBGAEE_00007 | LKKDFFFJ_00738 | wOo_01050 |
| wLug_00147 | WP_010962299.1 | KIHBGAEE_00107 | LKKDFFFJ_00109 | wOo_02890 |
| wLug_01015 | WP_010962854.1 | KIHBGAEE_00740 | LKKDFFFJ_00253 | wOo_05600 |
| wLug_00145 | WP_010962301.1 | KIHBGAEE_00109 | LKKDFFFJ_00251 | wOo_05580 |
| wLug_01245 | WP_010962886.1 | KIHBGAEE_00003 | LKKDFFFJ_00328 | wOo_01010 |
| wLug_01279 | WP_022626146.1 | KIHBGAEE_00223 | LKKDFFFJ_00739 | wOo_02070 |
| wLug_00144 | WP_010962302.1 | KIHBGAEE_00110 | LKKDFFFJ_00250 | wOo_05570 |
| wLug_00143 | WP_010962303.1 | KIHBGAEE_00111 | LKKDFFFJ_00249 | wOo_05560 |
| 28 | 29 | 29 | 28 | 28 |

| | | | | |
|---|---|---|---|---|
| wLug_00419 | WP_006279576.1 | KIHBGAEE_00311 | LKKDFFFJ_00783 | wOo_05960 |
| wLug_00420 | WP_006279574.1 | KIHBGAEE_00310 | LKKDFFFJ_00784 | wOo_05950 |
| wLug_00421 | WP_006279573.1 | KIHBGAEE_00309 | LKKDFFFJ_00785 | wOo_05940 |
| wLug_01274 | WP_010962676.1 | KIHBGAEE_00204 | LKKDFFFJ_01067 | wOo_01570 |
| wLug_00717 | WP_010962799.1 | | LKKDFFFJ_01179 | wOo_03710 |

| | | | | |
|---|---|---|---|---|
| wLug_00463 | WP_010962959.1 | KIHBGAEE_00506 | LKKDFFFJ_00548 | wOo_01750 |
| wLug_00340 | WP_010962403.1 | KIHBGAEE_00467 | LKKDFFFJ_00264 | wOo_00260 |
| wLug_00339 | WP_010962402.1 | KIHBGAEE_00466 | LKKDFFFJ_00265 | wOo_00250 |
| wLug_00460 | WP_022626393.1 | KIHBGAEE_00858 | LKKDFFFJ_00543 | wOo_01720 |
| wLug_00477 | WP_010962952.1 | KIHBGAEE_00261 | LKKDFFFJ_00561 | wOo_06260 |
| wLug_00476 | WP_010962953.1 | KIHBGAEE_00260 | LKKDFFFJ_00560 | wOo_06250 |
| wLug_00475 | WP_022626390.1 | KIHBGAEE_00259 | LKKDFFFJ_00559 | wOo_06240 |
| wLug_00474 | WP_010962955.1 | KIHBGAEE_00258 | LKKDFFFJ_00558 | wOo_06230 |
| wLug_00473 | WP_010962956.1 | KIHBGAEE_00257 | LKKDFFFJ_00557 | wOo_06220 |
| wLug_00370 | WP_010963112.1 | KIHBGAEE_00355 | LKKDFFFJ_01176 | wOo_07800 |
| | | KIHBGAEE_00356 | | wOo_07790 |
| wLug_00728 | WP_010962592.1 | KIHBGAEE_00712 | LKKDFFFJ_00430 | wOo_03910 |
| wLug_00705 | WP_010082472.1 | KIHBGAEE_00474 | LKKDFFFJ_01187 | wOo_04120 |
| wLug_00230 | WP_010962741.1 | KIHBGAEE_00679 | LKKDFFFJ_00177 | wOo_06550 |
| wLug_00269/00278 | WP_010962432.1 | KIHBGAEE_00118 | LKKDFFFJ_00690 | wOo_00670 |
| wLug_00362 | WP_010081973.1 | KIHBGAEE_00691 | LKKDFFFJ_00776 | wOo_01350 |
| wLug_00231 | WP_010962742.1 | KIHBGAEE_00929 | LKKDFFFJ_00178 | wOo_06560 |
| wLug_00268/00277 | WP_006279902.1 | KIHBGAEE_00117 | LKKDFFFJ_00689 | wOo_00660 |
| wLug_01265 | WP_010962583.1 | KIHBGAEE_00263 | LKKDFFFJ_00514 | wOo_00400 |
| wLug_01262 | WP_010962584.1 | KIHBGAEE_00265 | LKKDFFFJ_00516 | wOo_00420 |
| wLug_01263 | WP_010962585.1 | KIHBGAEE_00266 | LKKDFFFJ_00517 | wOo_00430 |
| wLug_01264 | WP_006014985.1 | KIHBGAEE_00264 | LKKDFFFJ_00515 | wOo_00410 |
| wLug_00604 | WP_010963095.1 | KIHBGAEE_00998 | LKKDFFFJ_00351 | wOo_02510 |
| wLug_01429 | WP_006280242.1 | KIHBGAEE_00233 | LKKDFFFJ_01264 | wOo_05230 |
| wLug_01428 | WP_010963022.1 | KIHBGAEE_00234 | LKKDFFFJ_01263 | wOo_05240 |
| wLug_00939 | WP_010962500.1 | KIHBGAEE_00424 | LKKDFFFJ_00915 | wOo_04400 |
| wLug_00583 | WP_010082343.1 | KIHBGAEE_00053 | LKKDFFFJ_00577 | wOo_07860 |
| wLug_01260 | WP_022626285.1 | KIHBGAEE_00268 | LKKDFFFJ_00519 | wOo_00440 |
| wLug_00291 | WP_006279354.1 | KIHBGAEE_01014 | LKKDFFFJ_01230 | wOo_06840 |
| wLug_00938 | WP_010962499.1 | KIHBGAEE_00423 | LKKDFFFJ_00916 | wOo_04390 |
| wLug_00937 | WP_010962498.1 | KIHBGAEE_00422 | LKKDFFFJ_00917 | wOo_04380 |
| | WP_010962801.1 | | LKKDFFFJ_00721 | |
| | WP_010962802.1 | | LKKDFFFJ_00720 | |

| | | | | |
|---|---|---|---|---|
| 35 | 37 | 35 | 37 | 36 |
| wLug_00535 | WP_007549072.1 | KIHBGAEE_00154 | LKKDFFFJ_00488 | wOo_07340 |
| | WP_010962558.1 | KIHBGAEE_00981 | LKKDFFFJ_01035 | |
| wLug_00702 | WP_010962790.1 | | LKKDFFFJ_01191 | wOo_04100 |
| wLug_00805 | WP_010962363.1 | KIHBGAEE_00092 | LKKDFFFJ_00972 | wOo_01160 |
| wLug_01470 | WP_010962406.1 | KIHBGAEE_00778 | LKKDFFFJ_01293 | wOo_00320 |
| wLug_01194 | WP_006279648.1 | KIHBGAEE_00995 | LKKDFFFJ_00420 | |
| wLug_01380 | WP_010963156.1 | KIHBGAEE_00156 | LKKDFFFJ_00877 | wOo_04190 |
| | | | | wOo_04200 |
| wLug_00401 | WP_010963034.1 | KIHBGAEE_00097 | LKKDFFFJ_00459 | wOo_02770 |
| wLug_00824 | WP_010962441.1 | KIHBGAEE_00208 | LKKDFFFJ_00231 | wOo_05330 |
| wLug_01206 | WP_010962512.1 | KIHBGAEE_00148 | LKKDFFFJ_00482 | wOo_07280 |
| wLug_01207 | WP_010962513.1 | KIHBGAEE_00149 | LKKDFFFJ_00483 | wOo_07290 |
| wLug_01196 | WP_010962298.1 | KIHBGAEE_00104 | LKKDFFFJ_00257 | wOo_07410 |
| wLug_00809 | WP_010962532.1 | KIHBGAEE_00062 | LKKDFFFJ_00605 | wOo_04630 |
| wLug_01205 | WP_010962511.1 | KIHBGAEE_00147 | LKKDFFFJ_00481 | wOo_07270 |
| 13 | 14 | 13 | 14 | 13 |
| wLug_01272 | WP_010962506.1 | KIHBGAEE_00733 | LKKDFFFJ_01218 | wOo_00900 |
| wLug_00590 | WP_010082448.1 | KIHBGAEE_00076 | LKKDFFFJ_00064 | wOo_06450 |
| wLug_00496 | WP_010963008.1 | KIHBGAEE_00401 | LKKDFFFJ_01243 | |
| wLug_00532 | WP_010963158.1 | KIHBGAEE_00158 | LKKDFFFJ_00879 | |
| wLug_00825 | WP_010962439.1 | KIHBGAEE_01000 | LKKDFFFJ_00230 | |
| wLug_01391 | WP_010963080.1 | KIHBGAEE_00814 | LKKDFFFJ_00335 | |
| wLug_00314 | WP_010962979.1 | KIHBGAEE_00532 | LKKDFFFJ_00523 | wOo_04910 |
| wLug_01301 | WP_010962830.1 | KIHBGAEE_00639 | LKKDFFFJ_00988 | wOo_07050 |
| wLug_00483 | WP_010082382.1 | KIHBGAEE_00609 | LKKDFFFJ_00660 | wOo_03590 |
| wLug_00484 | WP_010963048.1 | KIHBGAEE_00608 | LKKDFFFJ_00661 | |
| wLug_00290 | WP_010962392.1 | KIHBGAEE_01013 | LKKDFFFJ_01231 | |
| wLug_00871 | WP_010962860.1 | KIHBGAEE_00828 | LKKDFFFJ_00117 | |
| 12 | 12 | 12 | 12 | 5 |
| wLug_00734 | WP_010082338.1 | KIHBGAEE_00973 | LKKDFFFJ_00806 | wOo_03090 |
| wLug_01445 | WP_010963057.1 | KIHBGAEE_01024 | LKKDFFFJ_00310 | wOo_02620 |
| wLug_00947 | WP_006280146.1 | KIHBGAEE_00690 | LKKDFFFJ_00272 | wOo_00550 |
| wLug_00157 | WP_010962546.1 | KIHBGAEE_00716 | LKKDFFFJ_01127 | wOo_07870 |
| wLug_00270/00279 | WP_038228098.1 | KIHBGAEE_00119 | LKKDFFFJ_00691 | wOo_00680 |
| wLug_00551 | WP_010962988.1 | KIHBGAEE_00523 | LKKDFFFJ_00644 | wOo_03250 |

| | | | | |
|---|---|---|---|---|
| wLug_00970 | WP_010962543.1 | KIHBGAEE_00017 | LKKDFFFJ_00592 | wOo_06690 |
| wLug_00365 | WP_010963118.1 | KIHBGAEE_00069 | LKKDFFFJ_01213 | wOo_07770 |
| wLug_00578 | WP_010962776.1 | KIHBGAEE_00518 | LKKDFFFJ_01113 | wOo_06090 |
| wLug_00882 | WP_010962525.1 | KIHBGAEE_00478 | LKKDFFFJ_01289 | wOo_06290 |
| wLug_01067 | WP_010963039.1 | KIHBGAEE_00030 | LKKDFFFJ_00453 | wOo_00110 |
| wLug_00028 | WP_007548816.1 | KIHBGAEE_00657 | LKKDFFFJ_00449 | wOo_01660 |
| wLug_00510 | | KIHBGAEE_00170 | | |
| wLug_00447 | WP_010082331.1 | KIHBGAEE_00159 | LKKDFFFJ_00667 | |
| wLug_00410 | WP_010962968.1 | KIHBGAEE_00252 | LKKDFFFJ_00536 | |
| wLug_00409 | WP_006279859.1 | KIHBGAEE_00253 | LKKDFFFJ_00535 | |
| wLug_01030 | WP_010962934.1 | KIHBGAEE_00230 | LKKDFFFJ_00803 | wOo_05010 |
| wLug_01052 | WP_010962536.1 | KIHBGAEE_00067 | LKKDFFFJ_00597 | wOo_06730 |
| wLug_01031 | WP_022626380.1 | KIHBGAEE_00231 | LKKDFFFJ_00804 | wOo_05020 |
| wLug_00609 | WP_010963099.1 | KIHBGAEE_00792 | LKKDFFFJ_00356 | |
| 20 | 19 | 20 | 19 | 15 |
| wLug_01306 | WP_010962834.1 | KIHBGAEE_00799 | LKKDFFFJ_00993 | wOo_08290 |
| wLug_00709 | WP_010962796.1 | KIHBGAEE_00470 | LKKDFFFJ_01183 | wOo_04160 |
| wLug_00324 | WP_010963004.1 | KIHBGAEE_00406 | LKKDFFFJ_00923 | wOo_03620 |
| wLug_00631 | WP_010963063.1 | KIHBGAEE_00907 | LKKDFFFJ_00301 | wOo_07210 |
| wLug_00522 | WP_007549043.1 | | LKKDFFFJ_00321 | |
| wLug_00201 | WP_010082212.1 | KIHBGAEE_00369 | LKKDFFFJ_00390 | wOo_07610 |
| wLug_00320 | WP_010963007.1 | KIHBGAEE_00403 | LKKDFFFJ_00926 | wOo_03640 |
| wLug_00319 | WP_007548750.1 | KIHBGAEE_00402 | LKKDFFFJ_00927 | wOo_03650 |
| wLug_00216 | WP_010962629.1 | KIHBGAEE_00497 | LKKDFFFJ_01054 | wOo_05190 |
| wLug_00708 | WP_010962795.1 | KIHBGAEE_00471 | LKKDFFFJ_01184 | wOo_04150 |
| 10 | 10 | 9 | 10 | 9 |
| wLug_01386 | WP_010963153.1 | KIHBGAEE_00810 | LKKDFFFJ_00872 | wOo_04240 |
| wLug_01049 | WP_010962590.1 | KIHBGAEE_00271 | LKKDFFFJ_00061 | wOo_00460 |
| wLug_01257 | WP_010962828.1 | KIHBGAEE_00499 | LKKDFFFJ_00522 | wOo_03950 |
| wLug_00338 | WP_010962401.1 | KIHBGAEE_00934 | LKKDFFFJ_00266 | wOo_06430 |
| wLug_01340 | WP_010962666.1 | KIHBGAEE_00701 | LKKDFFFJ_00402 | wOo_05060 |
| wLug_00311 | WP_010962976.1 | KIHBGAEE_00921 | LKKDFFFJ_00526 | wOo_04870 |
| wLug_00559 | WP_010082013.1 | KIHBGAEE_00457 | LKKDFFFJ_00653 | wOo_03330 |

| | | | | |
|---|---|---|---|---|
| wLug_00621 | WP_010963106.1 | KIHBGAEE_00277 | LKKDFFFJ_01015 | wOo_04530 |
| wLug_00935 | WP_010962574.1 | KIHBGAEE_00014 | LKKDFFFJ_00443 | wOo_06670 |
| wLug_01412 | WP_010082099.1 | KIHBGAEE_00227 | LKKDFFFJ_00338 | wOo_04970 |
| wLug_01083 | WP_010081932.1 | KIHBGAEE_00410 | LKKDFFFJ_01106 | |
| wLug_00939 | WP_010962500.1 | KIHBGAEE_00424 | LKKDFFFJ_00915 | wOo_04400 |
| wLug_01260 | WP_022626285.1 | KIHBGAEE_00268 | LKKDFFFJ_00519 | wOo_00440 |
| 13 | 13 | 13 | 13 | 12 |
| wLug_01099 | WP_010962322.1 | KIHBGAEE_00084 | LKKDFFFJ_00861 | wOo_02980 |
| wLug_00639 | WP_010082377.1 | KIHBGAEE_00890 | LKKDFFFJ_00663 | wOo_05680 |
| wLug_01228 | WP_010962317.1 | KIHBGAEE_00879 | LKKDFFFJ_00467 | wOo_03440 |
| wLug_00772 | WP_010082490.1 | KIHBGAEE_00759 | LKKDFFFJ_00075 | wOo_03490 |
| wLug_00550 | WP_006279487.1 | KIHBGAEE_00522 | LKKDFFFJ_00643 | wOo_03240 |
| wLug_01091 | WP_010962921.1 | KIHBGAEE_00537 | LKKDFFFJ_00818 | wOo_05650 |
| wLug_00555 | WP_010962991.1 | KIHBGAEE_00527 | LKKDFFFJ_00649 | wOo_03290 |
| wLug_00383 | WP_010963167.1 | KIHBGAEE_00644 | LKKDFFFJ_00895 | wOo_01370 |
| wLug_00554 | WP_010081959.1 | KIHBGAEE_00526 | LKKDFFFJ_00648 | wOo_03280 |
| | WP_010962833.1 | | | |
| wLug_00750 | WP_010962893.1 | KIHBGAEE_00860 | LKKDFFFJ_00747 | wOo_00960 |
| wLug_01414 | WP_010082097.1 | KIHBGAEE_00229 | LKKDFFFJ_00336 | wOo_05000 |
| wLug_00999 | WP_010962523.1 | KIHBGAEE_00894 | LKKDFFFJ_01284 | wOo_07430 |
| wLug_00730 | WP_022626283.1 | KIHBGAEE_00383 | LKKDFFFJ_00428 | wOo_00130 |
| wLug_00576 | WP_010962778.1 | KIHBGAEE_00520 | LKKDFFFJ_01111 | wOo_06110 |
| wLug_01437 | WP_010963027.1 | KIHBGAEE_00202 | LKKDFFFJ_01269 | wOo_05320 |
| wLug_00288 | WP_006280151.1 | KIHBGAEE_00124 | LKKDFFFJ_00697 | wOo_00740 |
| wLug_00828 | WP_006280119.1 | KIHBGAEE_00132 | LKKDFFFJ_00837 | wOo_01930 |
| wLug_00879 | WP_010962427.1 | KIHBGAEE_01016 | LKKDFFFJ_01206 | wOo_03460 |
| wLug_01443 | WP_010963059.1 | | LKKDFFFJ_00308 | wOo_02640 |
| wLug_00798 | WP_022626203.1 | KIHBGAEE_00938 | LKKDFFFJ_00979 | wOo_00290 |
| 20 | 21 | 19 | 20 | 20 |
| wLug_00493 | WP_010963164.1 | KIHBGAEE_00352 | LKKDFFFJ_00883 | wOo_07690 |
| wLug_00262 | WP_010962760.1 | KIHBGAEE_00189 | LKKDFFFJ_00130 | wOo_08180 |
| wLug_00733 | WP_006280416.1 | KIHBGAEE_00640 | LKKDFFFJ_00807 | wOo_03080 |
| wLug_00813 | WP_022626228.1 | KIHBGAEE_00659 | LKKDFFFJ_00240 | wOo_03930 |
| wLug_00351 | WP_010082053.1 | KIHBGAEE_00832 | LKKDFFFJ_00768 | wOo_03820 |
| wLug_01343 | WP_010962448.1 | KIHBGAEE_00725 | LKKDFFFJ_00238 | wOo_05040 |
| wLug_00975 | WP_010082622.1 | | LKKDFFFJ_00606 | wOo_04540 |
| wLug_01437 | WP_010963027.1 | KIHBGAEE_00202 | LKKDFFFJ_01269 | wOo_05320 |
| wLug_00159 | WP_010962659.1 | KIHBGAEE_00366 | LKKDFFFJ_00394 | wOo_07650 |
| wLug_01414 | WP_010082097.1 | KIHBGAEE_00229 | LKKDFFFJ_00336 | wOo_05000 |
| wLug_00154 | WP_006280485.1 | KIHBGAEE_01018 | LKKDFFFJ_01124 | wOo_04600 |

| | | | | |
|---|---|---|---|---|
| wLug_00170 | WP_010962616.1 | KIHBGAEE_00564 | LKKDFFFJ_00680 | wOo_06780 |
| wLug_01414 | WP_010082097.1 | KIHBGAEE_00229 | LKKDFFFJ_00336 | wOo_05000 |
| wLug_00288 | WP_006280151.1 | KIHBGAEE_00124 | LKKDFFFJ_00697 | wOo_00740 |
| wLug_00828 | WP_006280119.1 | KIHBGAEE_00132 | LKKDFFFJ_00837 | wOo_01930 |
| wLug_00154 | WP_006280485.1 | KIHBGAEE_01018 | LKKDFFFJ_01124 | wOo_04600 |
| wLug_01189 | WP_006279458.1 | KIHBGAEE_00976 | LKKDFFFJ_00415 | wOo_04050 |
| wLug_00528 | WP_038227975.1 | KIHBGAEE_00414 | LKKDFFFJ_01101 | wOo_02550 |
| wLug_01402 | WP_010082033.1 | KIHBGAEE_00664 | LKKDFFFJ_00348 | wOo_02430 |
| 19 | 19 | 18 | 19 | 19 |
| wLug_00547 | WP_022626142.1 | KIHBGAEE_00288 | LKKDFFFJ_01149 | wOo_01610 |
| wLug_01255 | WP_006279877.1 | KIHBGAEE_00023 | LKKDFFFJ_01169 | wOo_00050 |
| 2 | 2 | 2 | 2 | 2 |

| wOv | wPip | wPpe | wStri |
|---|---|---|---|
| WP_025264087.1 | WP_007302229.1 | WP_070064703.1 | AOCBPJOI_00882 |
| WP_025264112.1 | WP_007302400.1 | WP_070065168.1 | AOCBPJOI_00975 |
| | WP_007302512.1 | | AOCBPJOI_01135 |
| | WP_007302513.1 | | AOCBPJOI_01136 |
| WP_025264197.1 | WP_007302552.1 | WP_070065334.1 | AOCBPJOI_01178 |
| 3 | 5 | 3 | 5 |
| WP_025264063.1 | WP_007302608.1 | | AOCBPJOI_01253 |
| | WP_012481923.1 | | AOCBPJOI_00876 |
| WP_007302560.1 | WP_070065343.1 | WP_070065343.1 | AOCBPJOI_01187 |
| | WP_007302797.1 | | AOCBPJOI_00061 |
| | WP_007302503.1 | | AOCBPJOI_01124 |
| | WP_007301939.1 | | AOCBPJOI_00457 |
| 2 | 6 | 1 | 6 |
| | | | AOCBPJOI_00230 |
| WP_025263950.1 | WP_007302174.1 | | AOCBPJOI_00719 |
| WP_025264118.1 | WP_007302563.1 | WP_070064622.1 | AOCBPJOI_01190 |
| WP_014868871.1 | WP_006014455.1 | WP_070064951.1 | AOCBPJOI_01023 |
| WP_014868810.1 | WP_007302843.1 | WP_070065243.1 | AOCBPJOI_01535 |
| | | | AOCBPJOI_00232 |
| | | | AOCBPJOI_00228 |
| | | | AOCBPJOI_00229 |
| | | | AOCBPJOI_00233 |
| WP_025264096.1 | WP_007302384.1 | WP_070064755.1 | AOCBPJOI_00949 |
| 5 | 5 | 4 | 10 |
| WP_025264388.1 | WP_007302089.1 | WP_070064750.1 | AOCBPJOI_00493 |
| WP_025264093.1 | WP_012481973.1 | WP_070065180.1 | AOCBPJOI_01014 |
| WP_025263950.1 | WP_007302174.1 | | AOCBPJOI_00719 |
| WP_025264118.1 | WP_007302563.1 | WP_070064622.1 | AOCBPJOI_01190 |
| WP_014868871.1 | WP_006014455.1 | WP_070064951.1 | AOCBPJOI_01023 |
| WP_014868810.1 | WP_007302843.1 | WP_070065243.1 | AOCBPJOI_01535 |
| WP_025264322.1 | WP_007302056.1 | | AOCBPJOI_00601 |
| 7 | 7 | 5 | 7 |
| WP_025264063.1 | WP_007302608.1 | | AOCBPJOI_01253 |
| WP_025264044.1 | WP_007302114.1 | | AOCBPJOI_00655 |

| | | | |
|---|---|---|---|
| WP_025264043.1 | | | AOCBPJOI_00654 |
| WP_025264075.1 | WP_012481955.1 | WP_070064581.1 | AOCBPJOI_00816 |
| WP_025264042.1 | WP_007302111.1 | | AOCBPJOI_00653 |
| 5 | 4 | 1 | 5 |
| WP_025264220.1 | WP_012481865.1 | WP_070065063.1 | AOCBPJOI_00576 |
| | WP_044475426.1 | WP_070064591.1 | AOCBPJOI_00388 |
| WP_025264030.1 | WP_012481842.1 | WP_081326513.1 | AOCBPJOI_00675 |
| WP_014869084.1 | WP_007302794.1 | WP_070064926.1 | AOCBPJOI_00065 |
| WP_025264279.1 | WP_012481857.1 | WP_081326526.1 | |
| WP_025264135.1 | WP_007302593.1 | WP_070064867.1 | AOCBPJOI_01226 |
| WP_025264198.1 | WP_007302551.1 | WP_070065333.1 | AOCBPJOI_01177 |
| WP_014868887.1 | WP_006012489.1 | WP_070064804.1 | AOCBPJOI_01234 |
| WP_025264056.1 | WP_007302796.1 | WP_070065122.1 | AOCBPJOI_00062 |
| WP_025264315.1 | WP_012481813.1 | WP_070065308.1 | AOCBPJOI_00367 |
| WP_014868835.1 | WP_007302619.1 | WP_070064642.1 | AOCBPJOI_01264 |
| WP_025264149.1 | WP_007302467.1 | WP_081326532.1 | AOCBPJOI_01085 |
| WP_025264366.1 | WP_007302411.1 | WP_070064671.1 | AOCBPJOI_01019 |
| WP_014868718.1 | WP_006013840.1 | WP_070065314.1 | AOCBPJOI_01527 |
| WP_014869000.1 | WP_007302699.1 | WP_070064595.1 | AOCBPJOI_01442 |
| WP_014868750.1 | WP_006015054.1 | WP_070064713.1 | AOCBPJOI_00571 |
| WP_014868799.1 | WP_007302273.1 | WP_070064991.1 | AOCBPJOI_00983 |
| WP_052307037.1 | WP_007302610.1 | WP_070065046.1 | AOCBPJOI_01255 |
| WP_025264268.1 | WP_007302028.1 | WP_070064715.1 | AOCBPJOI_00566 |
| WP_025264269.1 | WP_007302029.1 | WP_070064717.1 | AOCBPJOI_00567 |
| WP_025264271.1 | WP_012481863.1 | WP_081326504.1 | AOCBPJOI_00568 |
| | WP_012481864.1 | WP_081326505.1 | AOCBPJOI_00569 |
| WP_025264062.1 | WP_007302609.1 | WP_070064992.1 | AOCBPJOI_01254 |
| WP_025264375.1 | WP_007302720.1 | WP_070065082.1 | AOCBPJOI_01467 |
| WP_025264061.1 | WP_007302611.1 | | AOCBPJOI_01256 |
| WP_025264267.1 | WP_007302030.1 | WP_070064714.1 | AOCBPJOI_00570 |
| WP_025264323.1 | WP_007302059.1 | | AOCBPJOI_00604 |
| WP_025264060.1 | WP_007302612.1 | WP_070064990.1 | AOCBPJOI_01257 |
| WP_025264059.1 | WP_007302613.1 | WP_070064791.1 | AOCBPJOI_01258 |
| 27 | 29 | 27 | 28 |
| WP_014869121.1 | WP_012481929.1 | WP_070064575.1 | AOCBPJOI_01005 |
| WP_014869120.1 | WP_007302257.1 | WP_070064574.1 | AOCBPJOI_01004 |
| WP_014869119.1 | WP_007302258.1 | WP_070064573.1 | AOCBPJOI_01003 |
| WP_025264298.1 | WP_007302054.1 | WP_070065280.1 | AOCBPJOI_00599 |
| WP_025263945.1 | WP_007302017.1 | WP_070065307.1 | AOCBPJOI_00514 |

| | | | |
|---|---|---|---|
| WP_025264306.1 | WP_007302390.1 | WP_070065164.1 | AOCBPJOI_00963 |
| WP_025264230.1 | WP_007302472.1 | WP_070064587.1 | AOCBPJOI_01091 |
| WP_025264229.1 | WP_007302473.1 | WP_070064586.1 | AOCBPJOI_01093 |
| WP_014868809.1 | WP_012481971.1 | WP_070065244.1 | AOCBPJOI_00966 |
| WP_025264101.1 | WP_012481965.1 | WP_070065103.1 | AOCBPJOI_00943 |
| WP_025264100.1 | WP_006012916.1 | WP_070065104.1 | AOCBPJOI_00944 |
| WP_025264099.1 | WP_012481966.1 | WP_070065105.1 | AOCBPJOI_00945 |
| WP_025264098.1 | WP_012481967.1 | WP_070065106.1 | AOCBPJOI_00946 |
| WP_025264097.1 | WP_012481968.1 | WP_070065107.1 | AOCBPJOI_00947 |
| WP_025264187.1 | WP_012481985.1 | WP_070065130.1 | AOCBPJOI_01063 |
| WP_025264186.1 | WP_012481986.1 | WP_070065129.1 | AOCBPJOI_01064 |
| WP_025263958.1 | WP_012481881.1 | WP_070064874.1 | AOCBPJOI_00503 |
| WP_025263973.1 | WP_007302009.1 | WP_070065230.1 | AOCBPJOI_00522 |
| WP_025264120.1 | WP_007302558.1 | | AOCBPJOI_01185 |
| WP_029784320.1 | WP_019078549.1 | WP_081326572.1 | AOCBPJOI_01147 |
| WP_025264286.1 | WP_007302454.1 | WP_070064900.1 | AOCBPJOI_01071 |
| WP_025264121.1 | WP_007302557.1 | WP_070065341.1 | AOCBPJOI_01184 |
| WP_014868725.1 | WP_012482002.1 | WP_070065353.1 | AOCBPJOI_01148 |
| WP_014868703.1 | WP_007302047.1 | WP_070064846.1 | AOCBPJOI_00590 |
| WP_014868706.1 | WP_007302045.1 | WP_070064667.1 | AOCBPJOI_00587 |
| WP_025264238.1 | WP_007302046.1 | WP_081326500.1 | AOCBPJOI_00588 |
| WP_014868704.1 | WP_006014985.1 | WP_070064666.1 | AOCBPJOI_00589 |
| WP_025264353.1 | WP_006014178.1 | WP_070064727.1 | AOCBPJOI_00901 |
| WP_025264038.1 | WP_007302150.1 | WP_070065368.1 | AOCBPJOI_00744 |
| WP_025264039.1 | WP_007302151.1 | WP_081326574.1 | AOCBPJOI_00743 |
| WP_014868997.1 | WP_007302763.1 | WP_070065172.1 | AOCBPJOI_01519 |
| WP_025264192.1 | WP_010407017.1 | WP_081326502.1 | AOCBPJOI_00884 |
| WP_044104096.1 | WP_012481867.1 | WP_070064579.1 | AOCBPJOI_00585 |
| WP_025264140.1 | WP_007302514.1 | WP_070065161.1 | AOCBPJOI_01137 |
| WP_025263989.1 | WP_012481731.1 | WP_070065171.1 | AOCBPJOI_01518 |
| WP_014868995.1 | WP_007302761.1 | WP_070065210.1 | AOCBPJOI_01517 |

| | | | |
|---|---|---|---|
| 36 | 36 | 35 | 36 |
| WP_025264157.1 | WP_007302295.1 | WP_070064664.1 | AOCBPJOI_00839 |
| WP_025263971.1 | WP_012481860.1 | | AOCBPJOI_00525 |
| WP_025264276.1 | WP_007302888.1 | WP_081326543.1 | AOCBPJOI_00413 |
| WP_025264234.1 | WP_007302316.1 | WP_070065058.1 | AOCBPJOI_00788 |
| | WP_007302795.1 | WP_081326545.1 | AOCBPJOI_00063 |
| | WP_010403276.1 | WP_070064863.1 | AOCBPJOI_00699 |
| WP_025264370.1 | WP_007302416.1 | WP_070064948.1 | AOCBPJOI_01026 |
| | WP_007302914.1 | WP_070065194.1 | AOCBPJOI_00435 |
| WP_014869236.1 | WP_007302000.1 | WP_070065016.1 | AOCBPJOI_00531 |
| WP_014869237.1 | WP_007301999.1 | WP_070065015.1 | AOCBPJOI_00532 |
| WP_025264162.1 | WP_012481713.1 | | AOCBPJOI_00066 |
| WP_025264003.1 | WP_007302722.1 | WP_070065072.1 | AOCBPJOI_00453 |
| WP_025264155.1 | WP_007302001.1 | WP_081326530.1 | AOCBPJOI_00530 |
| 10 | 13 | 11 | 13 |
| WP_025264262.1 | WP_007302053.1 | WP_070065272.1 | AOCBPJOI_00597 |
| WP_025264114.1 | WP_006014313.1 | WP_070065313.1 | AOCBPJOI_00891 |
| | WP_007302323.1 | WP_070065226.1 | AOCBPJOI_00802 |
| | WP_007302298.1 | WP_070065277.1 | AOCBPJOI_00836 |
| | WP_007302913.1 | WP_070065047.1 | AOCBPJOI_00434 |
| | WP_012481907.1 | WP_070065239.1 | AOCBPJOI_00710 |
| WP_025264018.1 | WP_007302475.1 | WP_070065099.1 | AOCBPJOI_01116 |
| WP_025264145.1 | WP_012481876.1 | WP_070065008.1 | AOCBPJOI_00632 |
| | WP_007302375.1 | WP_070064943.1 | AOCBPJOI_00938 |
| | WP_007302374.1 | | AOCBPJOI_00937 |
| | WP_012481999.1 | WP_070065162.1 | AOCBPJOI_01138 |
| | | WP_070065345.1 | AOCBPJOI_00380 |
| 4 | 11 | 11 | 12 |
| WP_025264384.1 | WP_007302093.1 | WP_070064782.1 | AOCBPJOI_00497 |
| WP_025264361.1 | WP_007302135.1 | WP_070064682.1 | AOCBPJOI_00763 |
| WP_014868717.1 | WP_007302769.1 | WP_081326558.1 | AOCBPJOI_01528 |
| WP_025264193.1 | WP_012482020.1 | WP_070064753.1 | AOCBPJOI_01240 |
| WP_014868727.1 | WP_012482001.1 | WP_070065351.1 | AOCBPJOI_01146 |
| WP_025263916.1 | WP_007302279.1 | WP_070064618.1 | AOCBPJOI_00854 |

| | | | |
|---|---|---|---|
| WP_025264129.1 | WP_012481738.1 | WP_070064616.1 | AOCBPJOI_00046 |
| WP_025264184.1 | WP_007302450.1 | WP_070065102.1 | AOCBPJOI_01067 |
| WP_025264089.1 | WP_007302231.1 | WP_070065030.1 | AOCBPJOI_00879 |
| WP_025264103.1 | WP_007301928.1 | WP_070064705.1 | AOCBPJOI_00390 |
| WP_025264222.1 | WP_007302672.1 | WP_081326557.1 | AOCBPJOI_01411 |
| WP_014868803.1 | WP_007302587.1 | WP_070065371.1 | AOCBPJOI_01221 |
| | WP_019236836.1 | WP_070064582.1 | AOCBPJOI_00815 |
| | WP_007302404.1 | WP_070064774.1 | AOCBPJOI_00979 |
| | WP_012481974.1 | WP_070064751.1 | AOCBPJOI_01017 |
| | WP_012481975.1 | WP_070064752.1 | AOCBPJOI_01018 |
| WP_025264026.1 | WP_007302707.1 | WP_070064643.1 | AOCBPJOI_01453 |
| WP_025264131.1 | WP_007302687.1 | WP_070064597.1 | AOCBPJOI_01428 |
| WP_025264027.1 | WP_007302706.1 | WP_070064644.1 | AOCBPJOI_01452 |
| | WP_007302344.1 | WP_070065036.1 | AOCBPJOI_00906 |
| 15 | 20 | 20 | 20 |
| | WP_007302109.1 | WP_070064937.1 | AOCBPJOI_00640 |
| WP_025263977.1 | WP_007302013.1 | WP_070064625.1 | AOCBPJOI_00518 |
| WP_014868937.1 | WP_007302484.1 | WP_070064740.1 | AOCBPJOI_01107 |
| WP_025264153.1 | WP_012481964.1 | WP_070064552.1 | AOCBPJOI_00931 |
| | WP_007302306.1 | | AOCBPJOI_00826 |
| WP_025264174.1 | WP_007302837.1 | WP_070064932.1 | AOCBPJOI_00325 |
| WP_014868939.1 | WP_007302481.1 | WP_070064851.1 | AOCBPJOI_01110 |
| WP_025263942.1 | WP_007302480.1 | WP_070064737.1 | AOCBPJOI_01111 |
| WP_014869058.1 | WP_006012406.1 | WP_070064728.1 | AOCBPJOI_01202 |
| WP_025263976.1 | WP_012481861.1 | WP_081326496.1 | AOCBPJOI_00519 |
| 8 | 10 | 9 | 10 |
| WP_025263980.1 | WP_019078653.1 | WP_070065211.1 | AOCBPJOI_00706 |
| WP_025263961.1 | WP_007302040.1 | WP_070064545.1 | AOCBPJOI_00582 |
| WP_025264241.1 | WP_007302690.1 | WP_070065196.1 | AOCBPJOI_01432 |
| WP_014869164.1 | WP_007302474.1 | WP_070064982.1 | AOCBPJOI_01094 |
| WP_014869049.1 | WP_007301916.1 | WP_070064660.1 | AOCBPJOI_00677 |
| WP_025264015.1 | WP_007302498.1 | WP_070065369.1 | AOCBPJOI_01119 |
| WP_025263922.1 | WP_007302249.1 | WP_081326497.1 | AOCBPJOI_00862 |

| | | | |
|---|---|---|---|
| WP_025263996.1 | WP_007302354.1 | WP_070064763.1 | AOCBPJOI_00917 |
| WP_014869184.1 | WP_012481732.1 | WP_070064778.1 | AOCBPJOI_01515 |
| WP_025264022.1 | WP_007302167.1 | WP_070065309.1 | AOCBPJOI_00727 |
| | WP_007302660.1 | WP_070065124.1 | AOCBPJOI_01394 |
| WP_014868997.1 | WP_007302763.1 | WP_070065172.1 | AOCBPJOI_01519 |
| WP_044104096.1 | WP_012481867.1 | WP_070064579.1 | AOCBPJOI_00585 |
| 12 | 13 | 13 | 13 |

| | | | |
|---|---|---|---|
| WP_025264378.1 | WP_007302646.1 | WP_070065204.1 | AOCBPJOI_01742 |
| WP_025264066.1 | WP_007302311.1 | WP_070064738.1 | AOCBPJOI_00796 |
| WP_025263931.1 | WP_007301981.1 | WP_070064735.1 | AOCBPJOI_00553 |
| WP_014868928.1 | WP_007301943.1 | WP_070064652.1 | AOCBPJOI_00462 |
| WP_025263915.1 | WP_007302280.1 | WP_070064617.1 | AOCBPJOI_00853 |
| WP_025264064.1 | WP_007302652.1 | WP_070065056.1 | AOCBPJOI_01749 |
| WP_025263919.1 | WP_007302275.1 | WP_070065241.1 | AOCBPJOI_00858 |
| WP_025264288.1 | WP_007302433.1 | WP_070064541.1 | AOCBPJOI_01047 |
| | WP_007302276.1 | WP_070064620.1 | AOCBPJOI_00857 |
| | WP_007302075.1 | WP_070065011.1 | |
| WP_025264266.1 | WP_007301976.1 | WP_070064683.1 | AOCBPJOI_00480 |
| WP_014869043.1 | WP_007302165.1 | WP_070065312.1 | AOCBPJOI_00729 |
| WP_025264164.1 | WP_007302822.1 | WP_070064736.1 | AOCBPJOI_01549 |
| WP_025264223.1 | WP_007302097.1 | WP_070064629.1 | AOCBPJOI_00501 |
| WP_014869135.1 | WP_007302233.1 | WP_070065029.1 | AOCBPJOI_00877 |
| WP_014869070.1 | WP_007302143.1 | WP_070065354.1 | AOCBPJOI_00754 |
| WP_025264253.1 | WP_007302519.1 | WP_070065300.1 | AOCBPJOI_01140 |
| WP_014868819.1 | WP_007302810.1 | WP_070064921.1 | AOCBPJOI_00431 |
| WP_029784303.1 | WP_007301932.1 | WP_070065298.1 | AOCBPJOI_00387 |
| WP_014868861.1 | WP_029237555.1 | WP_070064543.1 | AOCBPJOI_00761 |
| WP_025264231.1 | WP_012481851.1 | WP_070064799.1 | AOCBPJOI_00404 |
| 19 | 21 | 21 | 20 |

| | | | |
|---|---|---|---|
| WP_025264179.1 | WP_007302307.1 | WP_070065304.1 | AOCBPJOI_00801 |
| WP_025264206.1 | WP_007302530.1 | | AOCBPJOI_01153 |
| WP_025264383.1 | WP_007302094.1 | WP_070064538.1 | AOCBPJOI_00498 |
| WP_025263960.1 | WP_007302727.1 | WP_070065159.1 | AOCBPJOI_00447 |
| WP_025263953.1 | WP_007302462.1 | WP_070064854.1 | AOCBPJOI_01080 |
| WP_025264029.1 | WP_007301912.1 | WP_070065080.1 | AOCBPJOI_00680 |
| WP_025263997.1 | WP_012481735.1 | WP_081326559.1 | AOCBPJOI_01499 |
| WP_014869070.1 | WP_007302143.1 | WP_070065354.1 | AOCBPJOI_00754 |
| WP_025264176.1 | WP_010407219.1 | WP_070064929.1 | AOCBPJOI_01238 |
| WP_014869043.1 | WP_007302165.1 | WP_070065312.1 | AOCBPJOI_00729 |
| WP_014869015.1 | WP_007302605.1 | WP_070065153.1 | AOCBPJOI_01243 |

| | | | |
|---|---|---|---|
| WP_014869194.1 | WP_007302592.1 | WP_070064887.1 | AOCBPJOI_01225 |
| WP_014869043.1 | WP_007302165.1 | WP_070065312.1 | AOCBPJOI_00729 |
| WP_025264253.1 | WP_007302519.1 | WP_070065300.1 | AOCBPJOI_01140 |
| WP_014868819.1 | WP_007302810.1 | WP_070064921.1 | AOCBPJOI_00431 |
| WP_014869015.1 | WP_007302605.1 | WP_070065153.1 | AOCBPJOI_01243 |
| WP_044104110.1 | WP_007302799.1 | WP_070065206.1 | AOCBPJOI_00058 |
| WP_025264356.1 | WP_007302302.1 | WP_070064942.1 | AOCBPJOI_00832 |
| WP_025264349.1 | WP_007302176.1 | WP_070064593.1 | AOCBPJOI_00717 |
| 19 | 19 | 18 | 19 |
| WP_014868800.1 | WP_006014087.1 | WP_070065260.1 | AOCBPJOI_00851 |
| | WP_007302038.1 | WP_070065135.1 | AOCBPJOI_00580 |
| 1 | 2 | 2 | 2 |

| wTpre | wVulC | wWb |
|---|---|---|
| WP_068652018.1 | KLT22481.1 | WP_088414480.1 |
| WP_068652059.1 | KLT22335.1 | WP_088415327.1 |
| | KLT22624.1 | |
| | KLT22625.1 | |
| WP_068651644.1 | KLT23222.1 | WP_088414083.1 |
| 3 | 5 | 3 |
| WP_068652574.1 | KLT22475.1 | WP_088415180.1 |
| WP_068652028.1 | KLT22653.1 | WP_088415325.1 |
| WP_068651629.1 | KLT23213.1 | WP_088414101.1 |
| WP_068651225.1 | KLT22974.1 | WP_088415468.1 |
| WP_068651717.1 | KLT23169.1 | WP_088415045.1 |
| WP_068651113.1 | KLT21729.1 | WP_088414322.1 |
| 6 | 6 | 6 |
| | KLT21798.1 | |
| WP_068652321.1 | KLT22033.1 | |
| WP_068651623.1 | KLT23210.1 | WP_088414107.1 |
| WP_068651904.1 | KLT22206.1 | WP_088414233.1 |
| WP_068651336.1 | KLT22992.1 | WP_088414739.1 |
| | KLT21796.1 | |
| | KLT21800.1 | |
| | KLT21799.1 | |
| | KLT21795.1 | |
| WP_068651942.1 | KLT22316.1 | WP_088415346.1 |
| 5 | 10 | 4 |
| WP_068651076.1 | KLT22440.1 | WP_088414257.1 |
| WP_068651888.1 | KLT21979.1 | WP_088415336.1 |
| WP_068652321.1 | KLT22033.1 | |
| WP_068651623.1 | KLT23210.1 | WP_088414107.1 |
| WP_068651904.1 | KLT22206.1 | WP_088414233.1 |
| WP_068651336.1 | KLT22992.1 | WP_088414739.1 |
| WP_068652375.1 | KLT22569.1 | WP_088414883.1 |
| 7 | 7 | 6 |
| WP_068652574.1 | KLT22653.1 | WP_088415180.1 |
| WP_068652070.1 | KLT22021.1 | |

| | | |
|---|---|---|
| WP_068652067.1 | KLT22020.1 | |
| WP_068652177.1 | KLT21699.1 | |
| WP_068652065.1 | KLT22019.1 | |
| 5 | 5 | 1 |

| | | |
|---|---|---|
| WP_068652676.1 | KLT22114.1 | WP_088414005.1 |
| WP_068651130.1 | KLT22978.1 | |
| WP_068651145.1 | KLT22990.1 | WP_088414312.1 |
| WP_068651216.1 | KLT22970.1 | WP_088415464.1 |
| WP_068652482.1 | KLT23377.1 | WP_088415412.1 |
| WP_068652570.1 | KLT22895.1 | WP_088415305.1 |
| WP_068651646.1 | KLT23223.1 | WP_088414081.1 |
| WP_068651320.1 | KLT21938.1 | WP_088415517.1 |
| WP_068651220.1 | KLT22973.1 | WP_088415466.1 |
| WP_068651578.1 | KLT22169.1 | WP_088414298.1 |
| WP_068651517.1 | KLT23183.1 | WP_088414879.1 |
| WP_068651781.1 | KLT22281.1 | WP_088414841.1 |
| WP_068651896.1 | KLT22202.1 | WP_088414476.1 |
| WP_006013840.1 | KLT23260.1 | WP_006013840.1 |
| WP_068651390.1 | KLT22877.1 | WP_088414135.1 |
| WP_068652414.1 | KLT22118.1 | WP_011256995.1 |
| WP_068652081.1 | KLT22651.1 | WP_088414033.1 |
| WP_068652572.1 | KLT22340.1 | WP_088415194.1 |
| WP_068652418.1 | KLT22120.1 | WP_088414779.1 |
| WP_068652421.1 | KLT22121.1 | WP_088414781.1 |
| WP_068652423.1 | KLT22122.1 | WP_088414783.1 |
| WP_068652678.1 | KLT22123.1 | |
| WP_068651308.1 | KLT22652.1 | WP_088415178.1 |
| WP_068651427.1 | KLT22664.1 | WP_088415388.1 |
| WP_068651310.1 | KLT22650.1 | WP_088415176.1 |
| WP_068652368.1 | KLT22566.1 | WP_088414785.1 |
| WP_068652416.1 | KLT22119.1 | WP_088414887.1 |
| WP_068651312.1 | KLT22649.1 | WP_088415174.1 |
| WP_068651314.1 | KLT22648.1 | WP_088415172.1 |
| 29 | 29 | 27 |

| | | |
|---|---|---|
| WP_068651876.1 | KLT21973.1 | WP_088414392.1 |
| WP_068651874.1 | KLT21972.1 | WP_088414394.1 |
| WP_068651872.1 | KLT21971.1 | WP_088414396.1 |
| WP_068652379.1 | KLT22571.1 | WP_088414805.1 |
| WP_068652438.1 | KLT22734.1 | WP_088414037.1 |

| | | |
|---|---|---|
| WP_068651928.1 | KLT22322.1 | WP_088414617.1 |
| WP_068652608.1 | KLT22015.1 | WP_088414211.1 |
| WP_068651773.1 | KLT22014.1 | WP_088414209.1 |
| WP_068652629.1 | KLT22326.1 | WP_088414741.1 |
| WP_068651951.1 | KLT22311.1 | WP_088415356.1 |
| WP_068651949.1 | KLT22312.1 | WP_088415354.1 |
| WP_068651947.1 | KLT22313.1 | WP_088415352.1 |
| WP_068651945.1 | KLT22314.1 | WP_088415350.1 |
| WP_068651944.1 | KLT22315.1 | WP_088415348.1 |
| WP_068651812.1 | KLT22260.1 | WP_088415000.1 |
| WP_068651810.1 | KLT22261.1 | WP_088414998.1 |
| WP_068652546.1 | KLT22451.1 | WP_088415258.1 |
| WP_068652446.1 | KLT23194.1 | WP_088415154.1 |
| WP_068651633.1 | KLT23215.1 | WP_088414097.1 |
| WP_068651694.1 | KLT22582.1 | WP_088414577.1 |
| WP_068651800.1 | KLT22267.1 | WP_088414563.1 |
| WP_068651635.1 | KLT23216.1 | WP_088414095.1 |
| WP_068651692.1 | KLT22581.1 | WP_088414531.1 |
| WP_068652394.1 | KLT21774.1 | WP_088415450.1 |
| WP_068652397.1 | KLT21771.1 | WP_088415444.1 |
| WP_068652398.1 | KLT21772.1 | WP_088415446.1 |
| WP_068652396.1 | KLT21773.1 | WP_088415448.1 |
| WP_068652187.1 | KLT21674.1 | WP_088415224.1 |
| WP_068652285.1 | KLT22059.1 | |
| WP_068652287.1 | KLT22058.1 | WP_088414970.1 |
| WP_068651283.1 | KLT23266.1 | WP_088414113.1 |
| WP_068652016.1 | KLT22483.1 | WP_088415011.1 |
| WP_068652399.1 | KLT21769.1 | WP_088415442.1 |
| WP_068652606.1 | KLT22623.1 | WP_088414542.1 |
| WP_068651281.1 | KLT23267.1 | WP_088414111.1 |
| WP_068651279.1 | KLT23268.1 | WP_088414109.1 |

| | | |
|---|---|---|
| 36 | 36 | 35 |
| WP_068652109.1 | KLT22560.1 | WP_088413973.1 |
| WP_068652440.1 | KLT23198.1 | WP_088415152.1 |
| WP_068651163.1 | KLT23388.1 | WP_088413995.1 |
| WP_068652140.1 | KLT23319.1 | WP_088415434.1 |
| WP_068651218.1 | KLT22972.1 | WP_088415478.1 |
| WP_068652238.1 | KLT23426.1 | WP_088414629.1 |
| WP_068651909.1 | KLT22209.1 | WP_088414238.1 |
| WP_068651584.1 | KLT22180.1 | WP_088415303.1 |
| WP_068652468.1 | KLT23204.1 | WP_088413963.1 |
| WP_068652682.1 | KLT23205.1 | WP_088413965.1 |
| WP_068651214.1 | KLT22969.1 | WP_088415184.1 |
| WP_068651429.1 | KLT21624.1 | WP_088414131.1 |
| WP_068652466.1 | KLT23203.1 | WP_088413961.1 |
| 13 | 13 | 13 |
| WP_068652380.1 | KLT21655.1 | WP_088414803.1 |
| WP_068652009.1 | KLT22487.1 | WP_088415491.1 |
| WP_068652151.1 | KLT21684.1 | WP_088415100.1 |
| WP_068652113.1 | KLT23021.1 | WP_088414633.1 |
| WP_068651582.1 | KLT22179.1 | WP_088414753.1 |
| WP_068651981.1 | KLT21947.1 | WP_088414482.1 |
| WP_068651731.1 | KLT22073.1 | WP_088414189.1 |
| WP_068652338.1 | KLT21703.1 | WP_088414496.1 |
| WP_068651959.1 | KLT22299.1 | WP_088414412.1 |
| WP_068651961.1 | KLT22298.1 | WP_088414414.1 |
| WP_068651707.1 | KLT22622.1 | WP_088414540.1 |
| WP_068651133.1 | KLT22656.1 | WP_088415374.1 |
| 12 | 12 | 12 |
| WP_068651071.1 | KLT22444.1 | WP_088414251.1 |
| WP_068652267.1 | KLT23129.1 | WP_088414456.1 |
| WP_068651293.1 | KLT23259.1 | WP_088414699.1 |
| WP_068651328.1 | KLT23413.1 | WP_088414900.1 |
| WP_068651696.1 | KLT22583.1 | WP_088414579.1 |
| WP_068652095.1 | KLT21636.1 | WP_088414376.1 |

| | | |
|---|---|---|
| WP_068651249.1 | KLT21956.1 | WP_088415291.1 |
| WP_068651806.1 | KLT22264.1 | WP_088414992.1 |
| WP_068652022.1 | KLT22478.1 | WP_088415319.1 |
| WP_068652562.1 | KLT22804.1 | WP_088415372.1 |
| WP_068651380.1 | KLT23307.1 | WP_088414007.1 |
| WP_068651124.1 | KLT22185.1 | WP_088414358.1 |
| WP_068652648.1 | KLT21698.1 | WP_088414604.1 |
| WP_068652051.1 | KLT22339.1 | WP_088415315.1 |
| WP_068651892.1 | KLT21982.1 | WP_088415340.1 |
| WP_068651894.1 | KLT21983.1 | WP_088415342.1 |
| WP_068651408.1 | KLT23347.1 | WP_088414171.1 |
| WP_068651358.1 | KLT23066.1 | WP_088415284.1 |
| WP_068651406.1 | KLT23348.1 | WP_088414169.1 |
| WP_068652195.1 | KLT21678.1 | WP_088414687.1 |
| 20 | 20 | 20 |
| WP_068652330.1 | KLT21708.1 | WP_088413989.1 |
| WP_068652454.1 | KLT23190.1 | WP_088415162.1 |
| WP_068651749.1 | KLT22083.1 | WP_088415108.1 |
| WP_068652226.1 | KLT22290.1 | WP_088414715.1 |
| | KLT21600.1 | |
| | KLT22761.1 | |
| WP_068651348.1 | KLT22999.1 | WP_088414512.1 |
| WP_068651741.1 | KLT22080.1 | WP_088415104.1 |
| WP_068651739.1 | KLT22079.1 | WP_088415102.1 |
| WP_068651615.1 | KLT22903.1 | WP_088414703.1 |
| WP_068652452.1 | KLT23191.1 | WP_088415160.1 |
| 9 | 11 | 9 |
| WP_068651971.1 | KLT21952.1 | WP_088414619.1 |
| WP_068651437.1 | KLT23069.1 | WP_088415252.1 |
| WP_068652401.1 | KLT21766.1 | |
| WP_068651772.1 | KLT22013.1 | WP_088414205.1 |
| WP_068651147.1 | KLT22988.1 | |
| WP_068651725.1 | KLT22591.1 | WP_088414725.1 |
| WP_068652047.1 | KLT21781.1 | WP_088414912.1 |

| | | |
|---|---|---|
| WP_068652209.1 | KLT23331.1 | WP_088414568.1 |
| WP_068651275.1 | KLT23270.1 | WP_088414199.1 |
| WP_068652308.1 | KLT22042.1 | WP_088414177.1 |
| WP_068651546.1 | KLT23053.1 | |
| WP_068651283.1 | KLT23266.1 | WP_088414113.1 |
| WP_068652399.1 | KLT21769.1 | WP_088415442.1 |
| 13 | 13 | 10 |
| WP_068651566.1 | KLT22855.1 | WP_088414771.1 |
| WP_068652134.1 | KLT22724.1 | WP_088414418.1 |
| WP_068652494.1 | KLT23367.1 | WP_088415460.1 |
| WP_068651109.1 | KLT21725.1 | WP_088415116.1 |
| WP_068652097.1 | KLT21637.1 | WP_088414378.1 |
| WP_068651555.1 | KLT22861.1 | WP_088414952.1 |
| WP_068652087.1 | KLT21632.1 | WP_088414368.1 |
| WP_068651834.1 | KLT22223.1 | WP_088415206.1 |
| WP_068652089.1 | KLT21633.1 | WP_088414370.1 |
| WP_068652332.1 | KLT21707.1 | WP_088414504.1 |
| WP_068651086.1 | KLT23091.1 | WP_088414360.1 |
| WP_068652306.1 | KLT22044.1 | WP_088414173.1 |
| WP_068651480.1 | KLT21964.1 | WP_088415190.1 |
| WP_068652550.1 | KLT22449.1 | WP_088414011.1 |
| WP_068652026.1 | KLT22476.1 | WP_088415323.1 |
| WP_068652278.1 | KLT23120.1 | WP_088413957.1 |
| WP_068651703.1 | KLT22587.1 | WP_088414537.1 |
| WP_068651462.1 | KLT22176.1 | WP_011256466.1 |
| WP_068651129.1 | KLT23036.1 | WP_088415248.1 |
| WP_068652650.1 | KLT23127.1 | WP_088414460.1 |
| WP_068652508.1 | KLT23398.1 | WP_088414675.1 |
| 21 | 21 | 21 |
| WP_068652125.1 | KLT23284.1 | WP_088414947.1 |
| WP_068651684.1 | KLT23245.1 | WP_088415242.1 |
| WP_068651070.1 | KLT22445.1 | WP_088414342.1 |
| WP_068651447.1 | KLT21621.1 | WP_088415254.1 |
| WP_068651786.1 | KLT22275.1 | WP_088414763.1 |
| WP_068651153.1 | KLT22985.1 | WP_088414306.1 |
| WP_068651246.1 | KLT22923.1 | WP_088415120.1 |
| WP_068652278.1 | KLT23120.1 | WP_088413957.1 |
| WP_068651324.1 | KLT21941.1 | WP_088414518.1 |
| WP_068652306.1 | KLT22044.1 | WP_088414173.1 |
| WP_068651304.1 | KLT22825.1 | WP_088414749.1 |

| | | |
|---|---|---|
| WP_068651299.1 | KLT22894.1 | WP_088415278.1 |
| WP_068652306.1 | KLT22044.1 | WP_088414173.1 |
| WP_068651703.1 | KLT22587.1 | WP_088414537.1 |
| WP_068651462.1 | KLT22176.1 | WP_011256466.1 |
| WP_068651304.1 | KLT22825.1 | WP_088414749.1 |
| WP_068651231.1 | KLT22100.1 | WP_088414591.1 |
| WP_068652121.1 | KLT23017.1 | WP_088414817.1 |
| WP_068652325.1 | KLT22031.1 | WP_088414799.1 |
| 19 | 19 | 19 |
| WP_068652098.1 | KLT21606.1 | WP_088414027.1 |
| WP_068652403.1 | KLT22110.1 | WP_088413997.1 |
| 2 | 2 | 2 |

**Table 4. Summary of BUSCOs analyses of Wolbachia and filarial nematodes complete ge**
Dataset used for the analysis was a proteobacteria dataset containing 1520 species representir

| organism | species/strain | NCBI accession | C | | S | | D | | F | |
|---|---|---|---|---|---|---|---|---|---|---|
| **organism** | **species/strain** | **NCBI accession** | **n** | **%** | **n** | **%** | **n** | **%** | **n** | **%** |
| | wLsig | **CP046577** | 168 | 76.0 | 167 | 75.6 | 1 | 0.5 | 8 | 3.6 |
| | wDimm | **CP046578** | 168 | 76.0 | 168 | 76.0 | 0 | 0.0 | 8 | 3.6 |
| | wCtub | **CP046579** | 171 | 77.4 | 171 | 77.4 | 0 | 0.0 | 6 | 2.7 |
| | wDcau | **CP046580** | 162 | 73.4 | 161 | 72.9 | 1 | 0.5 | 6 | 2.7 |
| | wLbra | **WQMO00000000** | 169 | 76.5 | 168 | 76.0 | 1 | 0.5 | 5 | 2.3 |
| | wMhie | **WQMP00000000** | 167 | 75.6 | 166 | 75.1 | 1 | 0.5 | 5 | 2.3 |
| | wBm | NC_006833 | 175 | 79.2 | 175 | 79.2 | 0 | 0.0 | 4 | 1.8 |
| | wBp | CP050521 | 178 | 81.3 | 178 | 81.3 | 0 | 0.0 | 7 | 3.2 |
| | wCau | CP041215 | 180 | 81.4 | 180 | 81.4 | 0 | 0.0 | 3 | 1.4 |
| | wCfeJ | CP051157 | 185 | 84.5 | 184 | 84.0 | 1 | 0.5 | 3 | 1.4 |
| | wCfeT | CP051156 | 184 | 84.1 | 183 | 83.6 | 1 | 0.5 | 4 | 1.8 |
| ***Wolbachia*** | wCle | AP013028 | 177 | 80.1 | 177 | 80.1 | 0 | 0.0 | 9 | 4.1 |
| | wFol | CP015510 | 179 | 81.0 | 179 | 81.0 | 0 | 0.0 | 4 | 1.8 |
| | wMel | NC_002978 | 180 | 81.4 | 180 | 81.4 | 0 | 0.0 | 2 | 0.9 |
| | wOo | NC_018267 | 165 | 74.7 | 165 | 74.7 | 0 | 0.0 | 6 | 2.7 |
| | wOv | HG810405 | 166 | 75.1 | 166 | 75.1 | 0 | 0.0 | 4 | 1.8 |
| | wPel | NC_010981 | 178 | 80.5 | 178 | 80.5 | 0 | 0.0 | 5 | 2.3 |
| | wPpe | MJMG01000000 | 161 | 72.9 | 160 | 72.4 | 1 | 0.5 | 12 | 5.4 |
| | wTpre | CM003641 | 180 | 81.4 | 180 | 81.4 | 0 | 0.0 | 4 | 1.8 |
| | wNfla | LYUW00000000 | 179 | 81.0 | 177 | 80.1 | 2 | 0.9 | 4 | 1.8 |
| | wLug | MUIY01000000 | 179 | 81.0 | 177 | 80.1 | 2 | 0.9 | 4 | 1.8 |
| | wstri | MUIX00000000 | 179 | 81.0 | 179 | 81.0 | 0 | 0.0 | 4 | 1.8 |
| | wVulC | ALWU00000000 | 180 | 81.5 | 179 | 81.0 | 1 | 0.5 | 4 | 1.8 |
| | B. malayi | AAQA00000000 | 860 | 87.5 | 842 | 86 | 18 | 1.8 | 57 | 5.8 |
| | B. pahangi | JRWH00000000 | 827 | 84.2 | 817 | 83 | 10 | 1.0 | 76 | 7.7 |
| | C. tuberocauda | **JABVXU000000000** | 939 | 95.6 | 935 | 95 | 4 | 0.4 | 31 | 3.2 |
| | D. immitis | **JABVXT000000000** | 932 | 94.9 | 926 | 94 | 6 | 0.6 | 30 | 3.1 |
| | D. caudispina | **JABVXV000000000** | 959 | 97.7 | 956 | 97.4 | 3 | 0.3 | 18 | 1.8 |
| ***Nematode*** | L. brasiliensis | **JABVXR000000000** | 917 | 93.4 | 911 | 92.8 | 6 | 0.6 | 46 | 4.7 |
| | L. sigmodontis | **JABVXW000000000** | 924 | 94 | 919 | 94 | 5 | 1 | 40 | 4 |
| | M. hiepei | **JABVXS000000000** | 849 | 86 | 789 | 80 | 60 | 6 | 73 | 7 |
| | O. ochengi | CAWC010000000 | 841 | 85.6 | 836 | 85 | 5 | 0.5 | 93 | 9.5 |
| | O. volvulus | CBVM000000000 | 958 | 97.6 | 955 | 97 | 3 | 0.3 | 17 | 1.7 |
| | W. bancrofti | LAQH01000000 | 917 | 93.4 | 852 | 87 | 65 | 6.6 | 38 | 3.9 |

**mplete genomes and draft genomes**

epresenting 221 BUSCOs for Wolbachia and a nematodes dataset containing 8 species representing 982 BUSCOs

| M | | Total |
|---|---|---|
| n | % | |
| 45 | 20 | 221 |
| 45 | 20 | 221 |
| 44 | 20 | 221 |
| 53 | 24 | 221 |
| 47 | 21.2 | 221 |
| 49 | 22.1 | 221 |
| 41 | 19.0 | 221 |
| 34 | 15.5 | 219 |
| 38 | 17.2 | 221 |
| 31 | 14.1 | 219 |
| 31 | 14.1 | 219 |
| 35 | 15.8 | 221 |
| 38 | 17.2 | 221 |
| 39 | 17.7 | 221 |
| 50 | 22.6 | 221 |
| 51 | 23.1 | 221 |
| 38 | 17.2 | 221 |
| 48 | 21.7 | 221 |
| 37 | 16.8 | 221 |
| 38 | 17.2 | 221 |
| 38 | 17.2 | 221 |
| 38 | 17.2 | 221 |
| 37 | 16.7 | 221 |
| 65 | 6.7 | 982 |
| 79 | 8.0 | 982 |
| 12 | 1.2 | 982 |
| 20 | 2.0 | 982 |
| 5 | 0.5 | 982 |
| 19 | 1.9 | 982 |
| 18 | 2 | 982 |
| 60 | 6 | 982 |
| 48 | 4.9 | 982 |
| 7 | 0.7 | 982 |
| 27 | 2.7 | 982 |

esenting 982 BUSCOs for the nematodes. Abbreviation: "C": Complete BUSCOs; "S": Complete and single-copy BU

Complete and single-copy BUSCOs; "D": Complete and duplicated BUSCOs (D); "F": Fragmented BUSCOs; "M": Mis agmented BUSCOs; "M": Missing BUSCOs.

**Table S5. nblast similai**

| gene | accession nb |
|---|---|
| 16S | AJ548802 |
| | FR827938 |
| | KU255234 |
| | KU255233 |
| | KU255232 |
| gltA | AJ609648 |
| | FJ390335 |
| groEL | AJ609658 |
| | FJ390369 |
| | FR827918 |
| | KU255204 |
| | KU255203 |
| ftsZ | FR827924 |
| dnaA | FJ390365 |
| | KU255295 |
| | KU255296 |
| | KU255297 |
| coxA | FJ390250 |
| | KU255265 |
| | KU255264 |
| | KU255263 |
| gatB | KU255375 |
| | KU255374 |
| fbpA | KU255325 |
| | KU255324 |
| nuoD | FJ390289 |
| atpD | FJ390360 |
| aspC | FJ390330 |
| NADH | FJ390209 |
| pdhB | FJ390153 |
| hyp | FJ390302 |
| hyp | FJ390237 |
| hyp | FJ390176 |

rity of sequences annoted as belonging to *Wolbachia* from Dipetalonema gracile previously sequ

| description |
| --- |
| Wolbachia pipientis partial 16S rRNA gene, specific host Dipetalonema gracile |
| Wolbachia endosymbiont of Dipetalonema gracile partial 16S rRNA gene (MIB:ZPL:01175) |
| Wolbachia endosymbiont of Dipetalonema gracile isolate 63YT MNHN 16S |
| Wolbachia endosymbiont of Dipetalonema gracile isolate 215YU ind4 MNHN 16S |
| Wolbachia endosymbiont of Dipetalonema gracile isolate 124CV MNHN 16S |
| Wolbachia pipientis (from Dipetalonema gracile) partial gltA gene for citrate synthase |
| Wolbachia endosymbiont of Dipetalonema gracile citrate synthase (gltA) gene |
| Wolbachia pipientis (from Dipetalonema gracile) partial gene for groEL protein |
| Wolbachia endosymbiont of Dipetalonema gracile chaperonin GroEL (groEL) gene |
| Wolbachia endosymbiont of Dipetalonema gracile partial groEL gene |
| Wolbachia endosymbiont of Dipetalonema gracile isolate 215YU ind4 MNHN groEL |
| Wolbachia endosymbiont of Dipetalonema gracile isolate 124CV MNHN groEL |
| Wolbachia endosymbiont of Dipetalonema gracile partial ftsZ gene for cell division protein |
| Wolbachia endosymbiont of Dipetalonema gracile chromosomal replication initiation protein (dnaA) gen |
| Wolbachia endosymbiont of Dipetalonema gracile isolate 124CV MNHN dnaA gene |
| Wolbachia endosymbiont of Dipetalonema gracile isolate 215YU ind4 MNHN dnaA gene |
| Wolbachia endosymbiont of Dipetalonema gracile isolate 63YT MNHN dnaA gene |
| Wolbachia endosymbiont of Dipetalonema gracile cytochrome c oxidase subunit I (coxA) gene |
| Wolbachia endosymbiont of Dipetalonema gracile isolate 63YT MNHN coxA gene |
| Wolbachia endosymbiont of Dipetalonema gracile isolate 215YU ind4 MNHN coxA gene |
| Wolbachia endosymbiont of Dipetalonema gracile isolate 124CV MNHN coxA gene |
| Wolbachia endosymbiont of Dipetalonema gracile isolate 215YU ind4 gatB gene |
| Wolbachia endosymbiont of Dipetalonema gracile isolate 124CV MNHN gatB |
| Wolbachia endosymbiont of Dipetalonema gracile isolate 215YU ind4 MNHN fbpA gene |
| Wolbachia endosymbiont of Dipetalonema gracile isolate 124CV MNHN gatB fbpA gene |
| Wolbachia endosymbiont of Dipetalonema gracile NADH dehydrogenase subunit D (nuoD) gene |
| Wolbachia endosymbiont of Dipetalonema gracile ATP synthase F1 beta subunit (atpD) gene |
| Wolbachia endosymbiont of Dipetalonema gracile aspartate aminotransferase (aspC) gene |
| Wolbachia endosymbiont of Dipetalonema gracile NADH dehydrogenase I subunit F gene |
| Wolbachia endosymbiont of Dipetalonema gracile pyruvate dehydrogenase beta subunit (pdhB) gene |
| Wolbachia endosymbiont of Dipetalonema gracile hypothetical protein gene |
| Wolbachia endosymbiont of Dipetalonema gracile hypothetical protein gene |
| Wolbachia endosymbiont of Dipetalonema gracile hypothetical protein gene |

| % identity with wDcau |
|---|
| 99.71% |
| 99.71% |
| 99.89% |
| 99.18% |
| 99.47% |
| 99.06% |
| 98.95% |
| 97.81% |
| 97.81% |
| 97.81% |
| 98.60% |
| 98.34% |
| 72.37%* |
| 83.42% |
| 96.60% |
| 95.36% |
| 98.98% |
| 98.18% |
| 99.98% |
| 98.39% |
| 98.32% |
| 98.39% |
| 98.02% |
| 96.88% |
| 97.40% |
| 97.88% |
| 89.91% |
| 81.37% |
| 98.92% |
| 85.71% |
| 98.28% |
| 82.96% |
| 98.85% |

| nblast with NCBI database description |
|---|
| Wolbachia endosymbiont of Dipetalonema gracile partial 16S rRNA gene (MIB:ZPL:01175) |
| Wolbachia pipientis partial 16S rRNA gene, specific host Dipetalonema gracile |
| Wolbachia endosymbiont of Dipetalonema gracile isolate 215YU ind4 MNHN 16S |
| Wolbachia endosymbiont of Dipetalonema gracile partial 16S rRNA gene (MIB:ZPL:01175) |
| Wolbachia endosymbiont of Dipetalonema gracile partial 16S rRNA gene (MIB:ZPL:01175) |
| Wolbachia endosymbiont of Dipetalonema gracile citrate synthase (gltA) gene |
| Wolbachia pipientis (from Dipetalonema gracile) partial gltA gene for citrate synthase |
| Wolbachia pipientis (from Dipetalonema gracile) partial gene for groEL protein |
| Wolbachia pipientis (from Dipetalonema gracile) partial gene for groEL protein |
| Wolbachia endosymbiont of Dipetalonema gracile chaperonin GroEL gene |
| Wolbachia endosymbiont of Dipetalonema gracile chaperonin GroEL gene |
| Wolbachia endosymbiont of Dipetalonema gracile chaperonin GroEL gene |
| Wolbachia endosymbiont of Onchocerca skrjabini partial ftsZ gene for cell division protein |
| Wolbachia pipientis strain wMel_ZH26 chromosome, complete genome |
| Wolbachia endosymbiont of Dipetalonema gracile isolate 215YU ind4 MNHN dnaA gene |
| Wolbachia endosymbiont of Dipetalonema gracile isolate 124CV MNHN dnaA gene |
| Wolbachia endosymbiont of Dipetalonema robini isolate 217YU ind7 MNHN dnaA gene |
| Wolbachia endosymbiont of Dipetalonema gracile isolate 124CV MNHN cytochrome c oxidase sub |
| Wolbachia endosymbiont of Dipetalonema gracile isolate 215YU ind4 MNHN coxA gene |
| Wolbachia endosymbiont of Dipetalonema gracile isolate 124CV MNHN coxA gene |
| Wolbachia endosymbiont of Dipetalonema gracile isolate 215YU ind4 MNHN coxA gene |
| Wolbachia endosymbiont of Dipetalonema gracile isolate 124CV MNHN gatB |
| Wolbachia endosymbiont of Dipetalonema gracile isolate 215YU ind4 gatB gene |
| Wolbachia endosymbiont of Dipetalonema gracile isolate 124CV MNHN fbpA gene |
| Wolbachia endosymbiont of Dipetalonema gracile isolate 215YU ind4 MNHN fbpA gene |
| Wolbachia endosymbiont of Onchocerca volvulus str. Cameroon W_O_volvulus_Cameroon_v3 |
| Wolbachia pipientis strain wMel_ZH26 chromosome, complete genome |
| Wolbachia pipientis strain wAlbB-HN2016 chromosome, complete genome |
| Wolbachia endosymbiont of Drosophila simulans wHa, complete genome |
| Wolbachia pipientis strain wTei pyruvate dehydrogenase beta subunit (pdhB) gene |
| Wolbachia endosymbiont of Cordylochernes scorpioides hypothetical protein gene |
| Wolbachia pipientis strain wMel_ZH26 chromosome, complete genome |
| Wolbachia endosymbiont of Onchocerca ochengi complete genome, strain wOo |

| accession | % identity | reference | |
|---|---|---|---|
| FR827938 | 99.78% | Casiraghi et al. 2004 | |
| AJ548802 | 99.78% | Ferri and al. 2011 | |
| KU255233 | 99.89% | Lefoulon and al. 2016 | |
| FR827938 | 99.08% | Lefoulon and al. 2016 | |
| FR827938 | 99.79% | Lefoulon and al. 2016 | |
| FJ390335 | 100.00% | Casiraghi et al. 2005 | |
| AJ609648 | 100.00% | Bordenstein and al. 2009 | |
| FJ390369 | 100.00% | Casiraghi et al. 2005 | |
| AJ609658 | 100.00% | Bordenstein and al. 2009 | |
| FJ390369 | 100.00% | Ferri and al. 2011 | |
| FJ390369 | 99.11% | Lefoulon and al. 2016 | |
| FJ390369 | 99.62% | Lefoulon and al. 2016 | |
| FR827925 | 96.53% | Ferri and al. 2011 | * 54% coverage only |
| CP042445 | 100.00% | Bordenstein and al. 2009 | |
| KU255296 | 98.98% | Lefoulon and al. 2016 | |
| KU255295 | 98.98% | Lefoulon and al. 2016 | |
| KU255301 | 97.28% | Lefoulon and al. 2016 | |
| KU255263 | 99.22% | Bordenstein and al. 2009 | |
| KU255264 | 99.47% | Lefoulon and al. 2016 | |
| KU255263 | 99.28% | Lefoulon and al. 2016 | |
| KU255264 | 98.36% | Lefoulon and al. 2016 | |
| KU255374 | 98.56% | Lefoulon and al. 2016 | |
| KU255375 | 98.56% | Lefoulon and al. 2016 | |
| KU255324 | 99.48% | Lefoulon and al. 2016 | |
| KU255325 | 99.48% | Lefoulon and al. 2016 | |
| HG810405 | 88.67 | Bordenstein and al. 2009 | |
| CP042445 | 99.83% | Bordenstein and al. 2009 | |
| CP041924 | 85.92% | Bordenstein and al. 2009 | |
| CP003884 | 85.18% | Bordenstein and al. 2009 | |
| DQ235372 | 99.70% | Bordenstein and al. 2009 | |
| FJ390303 | 83.88% | Bordenstein and al. 2009 | |
| CP042445 | 99.81% | Bordenstein and al. 2009 | |
| HE660029 | 84.78% | Bordenstein and al. 2009 | |

**Table S6. List of potential insertion sequences element (ISs) identify using ISSAGA.**

| | IS family | ISL3 | IS3 | | | |
|---|---|---|---|---|---|---|
| | sous-group | | IS3 | IS51 | IS150 | IS4 |
| wBm | complete | 0 | 0 | 0 | 0 | 0 |
| | partial | 0 | 0 | 0 | 0 | 0 |
| | unknown | 0 | 0 | 0 | 0 | 0 |
| | total | 0 | 0 | 0 | 0 | 0 |
| wBp | complete | 0 | 0 | 0 | 0 | 0 |
| | partial | 0 | 0 | 0 | 0 | 0 |
| | unknown | 0 | 0 | 0 | 0 | 0 |
| | total | 0 | 0 | 0 | 0 | 0 |
| wCau | complete | 1 | 2 | 0 | 0 | 0 |
| | partial | 0 | 0 | 0 | 2 | 6 |
| | unknown | 4 | 10 | 0 | 0 | 0 |
| | total | 5 | 12 | 0 | 2 | 6 |
| wCfeJ | complete | 0 | 0 | 0 | 0 | 0 |
| | partial | 0 | 0 | 0 | 0 | 1 |
| | unknown | 0 | 0 | 0 | 0 | 0 |
| | total | 0 | 0 | 0 | 0 | 1 |
| wCfeT | complete | 0 | 0 | 0 | 0 | 0 |
| | partial | 0 | 0 | 0 | 0 | 0 |
| | unknown | 0 | 0 | 0 | 0 | 0 |
| | total | 0 | 0 | 0 | 0 | 0 |
| wCle | complete | 0 | 0 | 0 | 0 | 0 |
| | partial | 2 | 0 | 0 | 0 | 0 |
| | unknown | 0 | 0 | 0 | 0 | 1 |
| | total | 2 | 0 | 0 | 0 | 1 |
| wCtub | complete | 0 | 0 | 0 | 0 | 0 |
| | partial | 0 | 0 | 0 | 0 | 0 |
| | unknown | 0 | 0 | 0 | 0 | 0 |
| | total | 0 | 0 | 0 | 0 | 0 |
| wDcau | complete | 0 | 0 | 0 | 0 | 0 |
| | partial | 0 | 0 | 0 | 0 | 0 |
| | unknown | 0 | 0 | 0 | 0 | 0 |
| | total | 0 | 0 | 0 | 0 | 0 |
| wDimm | complete | 0 | 0 | 0 | 0 | 0 |
| | partial | 0 | 0 | 0 | 0 | 0 |
| | unknown | 0 | 0 | 0 | 0 | 0 |
| | total | 0 | 0 | 0 | 0 | 0 |
| wFol | complete | 0 | 0 | 0 | 0 | 0 |
| | partial | 0 | 1 | 0 | 0 | 0 |

| | | | | | | |
|---|---|---|---|---|---|---|
| [illegible] | unknown | 1 | 0 | 0 | 0 | 16 |
| | total | 1 | 1 | 0 | 0 | 16 |
| wLsig | complete | 0 | 0 | 0 | 0 | 0 |
| | partial | 0 | 0 | 0 | 0 | 0 |
| | unknown | 0 | 0 | 0 | 0 | 0 |
| | total | 0 | 0 | 0 | 0 | 0 |
| wLbra | complete | 0 | 0 | 0 | 0 | 0 |
| | partial | 0 | 0 | 0 | 0 | 0 |
| | unknown | 0 | 0 | 0 | 0 | 0 |
| | total | 0 | 0 | 0 | 0 | 0 |
| wLug | complete | 1 | 3 | 0 | 0 | 0 |
| | partial | 0 | 20 | 0 | 0 | 0 |
| | unknown | 1 | 0 | 0 | 0 | 13 |
| | total | 2 | 23 | 0 | 0 | 13 |
| wMel | complete | 0 | 3 | 0 | 0 | 0 |
| | partial | 0 | 0 | 1 | 0 | 8 |
| | unknown | 1 | 6 | 0 | 0 | 0 |
| | total | 1 | 9 | 1 | 0 | 8 |
| wMhie | complete | 1 | 0 | 0 | 0 | 0 |
| | partial | 0 | 0 | 0 | 0 | 0 |
| | unknown | 0 | 0 | 0 | 0 | 0 |
| | total | 1 | 0 | 0 | 0 | 0 |
| wNfla | complete | 0 | 2 | 0 | 0 | 0 |
| | partial | 0 | 0 | 1 | 0 | 4 |
| | unknown | 1 | 1 | 0 | 0 | 0 |
| | total | 1 | 3 | 1 | 0 | 4 |
| wOo | complete | 0 | 0 | 0 | 0 | 0 |
| | partial | 0 | 0 | 0 | 0 | 0 |
| | unknown | 0 | 0 | 0 | 0 | 0 |
| | total | 0 | 0 | 0 | 0 | 0 |
| wOv | complete | 0 | 0 | 0 | 0 | 0 |
| | partial | 0 | 0 | 0 | 0 | 0 |
| | unknown | 0 | 0 | 0 | 0 | 0 |
| | total | 0 | 0 | 0 | 0 | 0 |
| wPip | complete | 0 | 0 | 0 | 0 | 0 |
| | partial | 1 | 0 | 0 | 0 | 0 |
| | unknown | 2 | 0 | 0 | 0 | 0 |
| | total | 3 | 0 | 0 | 0 | 0 |
| wPpe | complete | 0 | 0 | 0 | 0 | 0 |
| | partial | 0 | 0 | 0 | 0 | 0 |
| | unknown | 0 | 0 | 0 | 0 | 0 |
| | total | 0 | 0 | 0 | 0 | 0 |

| | | | | | | |
|---|---|---|---|---|---|---|
| wstri | complete | 2 | 21 | 0 | 0 | 0 |
| | partial | 0 | 0 | 0 | 0 | 0 |
| | unknown | 1 | 0 | 0 | 0 | 0 |
| | total | 3 | 21 | 0 | 0 | 0 |
| wTpre | complete | 0 | 0 | 0 | 0 | 0 |
| | partial | 0 | 0 | 0 | 0 | 0 |
| | unknown | 0 | 0 | 0 | 0 | 0 |
| | total | 0 | 0 | 0 | 0 | 0 |
| wVulC | complete | 0 | 0 | 0 | 17 | 9 |
| | partial | 2 | 0 | 0 | 8 | 6 |
| | unknown | 1 | 3 | 0 | 0 | 1 |
| | total | 3 | 3 | 0 | 25 | 16 |
| wWb | complete | 0 | 0 | 0 | 0 | 0 |
| | partial | 1 | 0 | 0 | 0 | 0 |
| | unknown | 0 | 0 | 0 | 0 | 0 |
| | total | 1 | 0 | 0 | 0 | 0 |

| potential insertion sequences element (ISs) | | | | | | | | |
|---|---|---|---|---|---|---|---|---|
| IS4 | IS5 | | | | | IS6 | IS200_IS605 | IS256 |
| IS50 | IS231 | IS427 | IS1031 | ISL2 | IS903 | | IS200 | |
| 0 | 0 | 0 | 0 | 0 | 0 | 0 | 0 | 0 |
| 0 | 0 | 0 | 1 | 0 | 0 | 0 | 0 | 0 |
| 0 | 0 | 0 | 0 | 0 | 0 | 0 | 0 | 0 |
| 0 | 0 | 0 | 1 | 0 | 0 | 0 | 0 | 0 |
| 0 | 0 | 0 | 0 | 0 | 0 | 0 | 0 | 0 |
| 0 | 0 | 0 | 1 | 0 | 0 | 0 | 0 | 0 |
| 0 | 0 | 0 | 0 | 0 | 0 | 0 | 0 | 0 |
| 0 | 0 | 0 | 1 | 0 | 0 | 0 | 0 | 0 |
| 0 | 3 | 0 | 54 | 0 | 0 | 0 | 0 | 0 |
| 5 | 1 | 0 | 58 | 0 | 0 | 0 | 0 | 0 |
| 0 | 0 | 0 | 0 | 0 | 16 | 0 | 0 | 0 |
| 5 | 4 | 0 | 112 | 0 | 16 | 0 | 0 | 0 |
| 0 | 0 | 0 | 1 | 0 | 0 | 0 | 0 | 0 |
| 0 | 0 | 1 | 3 | 0 | 0 | 0 | 0 | 0 |
| 0 | 0 | 0 | 0 | 0 | 0 | 0 | 0 | 0 |
| 0 | 0 | 1 | 4 | 0 | 0 | 0 | 0 | 0 |
| 0 | 0 | 1 | 0 | 0 | 0 | 0 | 0 | 0 |
| 0 | 0 | 0 | 32 | 0 | 1 | 0 | 0 | 0 |
| 0 | 0 | 0 | 0 | 0 | 55 | 0 | 0 | 0 |
| 0 | 0 | 1 | 32 | 0 | 56 | 0 | 0 | 0 |
| 0 | 0 | 0 | 39 | 0 | 0 | 0 | 0 | 0 |
| 1 | 0 | 0 | 181 | 0 | 0 | 0 | 0 | 0 |
| 1 | 0 | 0 | 1 | 0 | 0 | 0 | 0 | 0 |
| 2 | 0 | 0 | 221 | 0 | 0 | 0 | 0 | 0 |
| 0 | 0 | 0 | 0 | 0 | 0 | 0 | 0 | 0 |
| 0 | 0 | 0 | 0 | 0 | 0 | 0 | 0 | 0 |
| 0 | 0 | 0 | 0 | 0 | 0 | 0 | 0 | 0 |
| 0 | 0 | 0 | 0 | 0 | 0 | 0 | 0 | 0 |
| 0 | 0 | 0 | 0 | 0 | 0 | 0 | 0 | 0 |
| 0 | 0 | 0 | 0 | 0 | 0 | 0 | 0 | 0 |
| 0 | 0 | 0 | 0 | 0 | 0 | 0 | 0 | 0 |
| 0 | 0 | 0 | 0 | 0 | 0 | 0 | 0 | 0 |
| 0 | 0 | 0 | 0 | 0 | 0 | 0 | 0 | 0 |
| 0 | 0 | 0 | 0 | 0 | 0 | 0 | 0 | 0 |
| 0 | 0 | 0 | 0 | 0 | 0 | 0 | 0 | 0 |
| 0 | 0 | 0 | 0 | 0 | 0 | 0 | 0 | 0 |
| 1 | 0 | 6 | 0 | 0 | 0 | 0 | 0 | 2 |
| 0 | 0 | 0 | 0 | 0 | 0 | 0 | 0 | 0 |

| | | | | | | | | |
|---|---|---|---|---|---|---|---|---|
| 20 | 0 | 0 | 0 | 0 | 13 | 0 | 0 | 0 |
| 21 | 0 | 6 | 0 | 0 | 13 | 0 | 0 | 2 |
| 0 | 0 | 0 | 0 | 0 | 0 | 0 | 0 | 0 |
| 0 | 0 | 0 | 0 | 0 | 0 | 0 | 0 | 0 |
| 0 | 0 | 0 | 0 | 0 | 0 | 0 | 0 | 0 |
| 0 | 0 | 0 | 0 | 0 | 0 | 0 | 0 | 0 |
| 0 | 0 | 0 | 0 | 0 | 0 | 0 | 0 | 0 |
| 0 | 0 | 0 | 0 | 0 | 0 | 0 | 0 | 0 |
| 0 | 0 | 0 | 0 | 0 | 0 | 0 | 0 | 0 |
| 0 | 0 | 0 | 0 | 0 | 0 | 0 | 0 | 0 |
| 0 | 4 | 0 | 4 | 0 | 0 | 3 | 1 | 0 |
| 2 | 18 | 1 | 14 | 1 | 0 | 0 | 4 | 1 |
| 0 | 0 | 0 | 0 | 0 | 1 | 0 | 0 | 0 |
| 2 | 22 | 1 | 18 | 1 | 1 | 3 | 5 | 1 |
| 0 | 3 | 0 | 13 | 0 | 0 | 0 | 0 | 1 |
| 5 | 0 | 0 | 16 | 0 | 0 | 0 | 0 | 1 |
| 0 | 0 | 0 | 0 | 0 | 16 | 0 | 0 | 0 |
| 5 | 3 | 0 | 29 | 0 | 16 | 0 | 0 | 2 |
| 0 | 0 | 0 | 0 | 0 | 0 | 0 | 0 | 0 |
| 0 | 0 | 1 | 1 | 0 | 0 | 0 | 0 | 0 |
| 0 | 0 | 1 | 0 | 0 | 0 | 0 | 0 | 0 |
| 0 | 0 | 2 | 1 | 0 | 0 | 0 | 0 | 0 |
| 0 | 0 | 0 | 0 | 2 | 0 | 0 | 0 | 0 |
| 4 | 3 | 0 | 0 | 23 | 0 | 0 | 0 | 3 |
| 0 | 0 | 0 | 0 | 0 | 3 | 0 | 0 | 0 |
| 4 | 3 | 0 | 0 | 25 | 3 | 0 | 0 | 3 |
| 0 | 0 | 0 | 0 | 0 | 0 | 0 | 0 | 0 |
| 0 | 1 | 0 | 0 | 0 | 0 | 0 | 0 | 0 |
| 0 | 0 | 0 | 0 | 0 | 0 | 0 | 0 | 0 |
| 0 | 1 | 0 | 0 | 0 | 0 | 0 | 0 | 0 |
| 0 | 0 | 0 | 0 | 0 | 0 | 0 | 0 | 0 |
| 0 | 0 | 0 | 0 | 0 | 0 | 0 | 0 | 0 |
| 0 | 0 | 0 | 0 | 0 | 0 | 0 | 0 | 0 |
| 0 | 0 | 0 | 0 | 0 | 0 | 0 | 0 | 0 |
| 0 | 0 | 0 | 1 | 0 | 0 | 0 | 0 | 4 |
| 0 | 0 | 0 | 3 | 0 | 1 | 9 | 3 | 36 |
| 0 | 0 | 0 | 0 | 0 | 26 | 0 | 0 | 0 |
| 1 | 0 | 0 | 4 | 0 | 27 | 9 | 3 | 40 |
| 0 | 0 | 0 | 0 | 0 | 0 | 0 | 0 | 0 |
| 0 | 0 | 0 | 0 | 6 | 0 | 0 | 0 | 0 |
| 0 | 0 | 0 | 0 | 1 | 0 | 0 | 0 | 0 |
| 0 | 0 | 0 | 0 | 7 | 0 | 0 | 0 | 0 |

| | | | | | | | | |
|---|---|---|---|---|---|---|---|---|
| 0 | 0 | 0 | 5 | 0 | 0 | 1 | 11 | 0 |
| 0 | 1 | 0 | 4 | 1 | 0 | 0 | 1 | 3 |
| 0 | 0 | 0 | 0 | 0 | 2 | 0 | 0 | 0 |
| 0 | 1 | 0 | 9 | 1 | 2 | 1 | 12 | 3 |
| 0 | 0 | 0 | 0 | 0 | 0 | 0 | 0 | 0 |
| 0 | 19 | 0 | 2 | 1 | 0 | 0 | 0 | 0 |
| 0 | 0 | 0 | 0 | 0 | 0 | 0 | 0 | 0 |
| 0 | 19 | 0 | 2 | 1 | 0 | 0 | 0 | 0 |
| 0 | 0 | 0 | 0 | 0 | 0 | 0 | 0 | 0 |
| 29 | 3 | 2 | 2 | 7 | 0 | 0 | 0 | 31 |
| 0 | 0 | 0 | 0 | 0 | 5 | 0 | 0 | 0 |
| 29 | 3 | 2 | 2 | 7 | 5 | 0 | 0 | 31 |
| 0 | 0 | 0 | 0 | 0 | 0 | 0 | 0 | 0 |
| 0 | 0 | 0 | 1 | 0 | 0 | 0 | 0 | 2 |
| 0 | 0 | 0 | 0 | 0 | 0 | 0 | 0 | 0 |
| 0 | 0 | 0 | 1 | 0 | 0 | 0 | 0 | 2 |

| IS481 | IS66 | IS982 | IS630 | IS110 | | ISNCY | total |
|---|---|---|---|---|---|---|---|
| | ISBst12 | | | | IS1111 | | |
| 0 | 0 | 0 | 0 | 0 | 0 | 0 | 0 |
| 0 | 0 | 0 | 2 | 0 | 0 | 0 | 3 |
| 0 | 0 | 0 | 0 | 0 | 0 | 0 | 0 |
| 0 | 0 | 0 | 2 | 0 | 0 | 0 | 3 |
| 0 | 0 | 0 | 0 | 0 | 0 | 0 | 0 |
| 0 | 0 | 0 | 0 | 0 | 0 | 0 | 1 |
| 0 | 0 | 0 | 0 | 0 | 0 | 0 | 0 |
| 0 | 0 | 0 | 0 | 0 | 0 | 0 | 1 |
| 1 | 0 | 0 | 3 | 2 | 2 | 0 | 68 |
| 2 | 5 | 1 | 11 | 9 | 1 | 0 | 101 |
| 0 | 0 | 0 | 1 | 4 | 1 | 0 | 36 |
| 3 | 5 | 1 | 15 | 15 | 4 | 0 | 205 |
| 0 | 0 | 2 | 0 | 2 | 0 | 0 | 5 |
| 9 | 0 | 1 | 0 | 0 | 0 | 0 | 15 |
| 0 | 0 | 0 | 0 | 0 | 0 | 0 | 0 |
| 9 | 0 | 3 | 0 | 2 | 0 | 0 | 20 |
| 0 | 104 | 1 | 0 | 0 | 8 | 0 | 114 |
| 0 | 87 | 0 | 0 | 0 | 0 | 0 | 120 |
| 0 | 0 | 0 | 0 | 0 | 0 | 0 | 55 |
| 0 | 191 | 1 | 0 | 0 | 8 | 0 | 289 |
| 0 | 0 | 0 | 0 | 0 | 0 | 0 | 39 |
| 0 | 0 | 0 | 1 | 4 | 0 | 0 | 189 |
| 0 | 0 | 0 | 0 | 0 | 0 | 0 | 3 |
| 0 | 0 | 0 | 1 | 4 | 0 | 0 | 231 |
| 0 | 0 | 0 | 0 | 0 | 0 | 0 | 0 |
| 0 | 0 | 0 | 0 | 0 | 0 | 0 | 0 |
| 0 | 0 | 0 | 0 | 0 | 0 | 0 | 0 |
| 0 | 0 | 0 | 0 | 0 | 0 | 0 | 0 |
| 0 | 0 | 0 | 0 | 0 | 0 | 0 | 0 |
| 0 | 0 | 0 | 0 | 0 | 0 | 0 | 0 |
| 0 | 0 | 0 | 0 | 0 | 0 | 0 | 0 |
| 0 | 0 | 0 | 0 | 0 | 0 | 0 | 0 |
| 0 | 0 | 0 | 0 | 0 | 0 | 0 | 0 |
| 0 | 0 | 0 | 0 | 0 | 0 | 0 | 0 |
| 0 | 0 | 0 | 0 | 0 | 0 | 0 | 0 |
| 0 | 0 | 0 | 0 | 0 | 0 | 0 | 0 |
| 0 | 0 | 0 | 3 | 15 | 0 | 0 | 27 |
| 0 | 0 | 3 | 0 | 12 | 0 | 0 | 16 |

| | | | | | | | |
|---|---|---|---|---|---|---|---|
| 20 | 0 | 0 | 0 | 5 | 0 | 0 | 75 |
| 20 | 0 | 3 | 3 | 32 | 0 | 0 | 118 |
| 0 | 0 | 0 | 0 | 0 | 0 | 0 | 0 |
| 0 | 0 | 0 | 15 | 0 | 0 | 0 | 15 |
| 0 | 0 | 0 | 2 | 0 | 0 | 0 | 2 |
| 0 | 0 | 0 | 17 | 0 | 0 | 0 | 17 |
| 0 | 0 | 0 | 1 | 0 | 0 | 0 | 1 |
| 0 | 0 | 0 | 32 | 0 | 0 | 0 | 32 |
| 0 | 0 | 0 | 6 | 0 | 0 | 0 | 6 |
| 0 | 0 | 0 | 39 | 0 | 0 | 0 | 39 |
| 0 | 2 | 0 | 1 | 3 | 0 | 0 | 22 |
| 8 | 0 | 5 | 7 | 4 | 0 | 0 | 85 |
| 0 | 0 | 0 | 1 | 0 | 0 | 0 | 16 |
| 8 | 2 | 5 | 9 | 7 | 0 | 0 | 123 |
| 1 | 0 | 0 | 3 | 2 | 0 | 0 | 26 |
| 0 | 3 | 0 | 2 | 8 | 1 | 0 | 45 |
| 0 | 0 | 0 | 1 | 3 | 0 | 0 | 27 |
| 1 | 3 | 0 | 6 | 13 | 1 | 0 | 98 |
| 0 | 0 | 0 | 0 | 0 | 0 | 0 | 1 |
| 2 | 1 | 8 | 0 | 0 | 0 | 0 | 13 |
| 0 | 0 | 0 | 0 | 0 | 0 | 0 | 1 |
| 2 | 1 | 8 | 0 | 0 | 0 | 0 | 15 |
| 1 | 0 | 0 | 1 | 2 | 0 | 0 | 8 |
| 2 | 3 | 2 | 5 | 6 | 1 | 0 | 57 |
| 0 | 0 | 0 | 0 | 2 | 1 | 0 | 8 |
| 3 | 3 | 2 | 6 | 10 | 2 | 0 | 73 |
| 0 | 0 | 0 | 0 | 0 | 0 | 0 | 0 |
| 0 | 0 | 0 | 0 | 0 | 0 | 0 | 1 |
| 0 | 0 | 0 | 0 | 0 | 0 | 0 | 0 |
| 0 | 0 | 0 | 0 | 0 | 0 | 0 | 1 |
| 0 | 0 | 0 | 0 | 0 | 0 | 0 | 0 |
| 0 | 0 | 0 | 0 | 0 | 0 | 0 | 0 |
| 0 | 0 | 0 | 0 | 0 | 0 | 0 | 0 |
| 0 | 0 | 0 | 0 | 0 | 0 | 0 | 0 |
| 0 | 1 | 44 | 10 | 3 | 0 | 0 | 63 |
| 0 | 0 | 8 | 4 | 0 | 0 | 0 | 65 |
| 0 | 0 | 0 | 0 | 0 | 0 | 1 | 29 |
| 0 | 0 | 52 | 14 | 3 | 0 | 1 | 157 |
| 0 | 0 | 0 | 0 | 0 | 0 | 0 | 0 |
| 0 | 0 | 0 | 0 | 0 | 0 | 0 | 6 |
| 0 | 0 | 0 | 0 | 0 | 0 | 0 | 1 |
| 0 | 0 | 0 | 0 | 0 | 0 | 0 | 7 |

| | | | | | | | |
|---|---|---|---|---|---|---|---|
| 0 | 0 | 6 | 3 | 14 | 0 | 0 | 63 |
| 0 | 0 | 0 | 1 | 6 | 0 | 0 | 17 |
| 0 | 0 | 0 | 0 | 0 | 0 | 0 | 3 |
| 0 | 0 | 6 | 4 | 20 | 0 | 0 | 83 |
| 0 | 0 | 0 | 0 | 0 | 0 | 0 | 0 |
| 2 | 0 | 0 | 1 | 1 | 0 | 0 | 26 |
| 0 | 0 | 0 | 0 | 0 | 0 | 0 | 0 |
| 2 | 0 | 0 | 1 | 1 | 0 | 0 | 26 |
| 0 | 2 | 0 | 5 | 9 | 0 | 0 | 42 |
| 11 | 4 | 0 | 10 | 46 | 6 | 0 | 167 |
| 0 | 0 | 0 | 0 | 0 | 0 | 0 | 10 |
| 11 | 6 | 0 | 15 | 55 | 6 | 0 | 219 |
| 0 | 0 | 0 | 0 | 0 | 0 | 0 | 0 |
| 0 | 0 | 0 | 0 | 0 | 0 | 0 | 4 |
| 0 | 0 | 0 | 0 | 0 | 0 | 0 | 0 |
| 0 | 0 | 0 | 0 | 0 | 0 | 0 | 4 |

**Table S7. List of Group II intron-associated genes detectd by RAST pipeline.**

| genome | Feature ID | Type | Contig | Start | Stop |
|---|---|---|---|---|---|
| wBm | - | - | - | - | - |
| wBp | - | - | - | - | - |
| wCauA | fig\|2591635.5.peg.438 | CDS | CP041215.1 | 409378 | 409653 |
| | fig\|2591635.5.peg.1150 | CDS | CP041215.1 | 1085440 | 1086753 |
| wCtub | - | - | - | - | - |
| wCfeJ | fig\|212123.3.peg.24 | CDS | CP051157.1 | 20515 | 19613 |
| | fig\|212123.3.peg.25 | CDS | CP051157.1 | 21483 | 20473 |
| | fig\|212123.3.peg.915 | CDS | CP051157.1 | 943435 | 945081 |
| wCfeT | fig\|212123.4.peg.109 | CDS | CP051156.1 | 104955 | 106331 |
| | fig\|212123.4.peg.120 | CDS | CP051156.1 | 112631 | 114007 |
| | fig\|212123.4.peg.131 | CDS | CP051156.1 | 120307 | 121683 |
| | fig\|212123.4.peg.904 | CDS | CP051156.1 | 913030 | 911393 |
| | fig\|212123.4.peg.1096 | CDS | CP051156.1 | 1111462 | 1112943 |
| | fig\|212123.4.peg.1236 | CDS | CP051156.1 | 1238861 | 1236990 |
| | fig\|212123.4.peg.1413 | CDS | CP051156.1 | 1396973 | 1395642 |
| | fig\|212123.4.peg.1457 | CDS | CP051156.1 | 1436914 | 1438551 |
| wCle | - | - | - | - | - |
| wDimm | - | - | - | - | - |
| wDcau | - | - | - | - | - |
| wFol | fig\|169402.10.peg.411 | CDS | NZ_CP015510.2 | 425791 | 426789 |
| | fig\|169402.10.peg.1022 | CDS | NZ_CP015510.2 | 1083651 | 1082887 |
| | fig\|169402.10.peg.1146 | CDS | NZ_CP015510.2 | 1223669 | 1223989 |
| | fig\|169402.10.peg.1149 | CDS | NZ_CP015510.2 | 1226526 | 1227566 |
| | fig\|169402.10.peg.1169 | CDS | NZ_CP015510.2 | 1246407 | 1246850 |
| wMel | fig\|163164.1.peg.621 | CDS | NC_002978 | 669520 | 671067 |
| | fig\|163164.1.peg.887 | CDS | NC_002978 | 951605 | 953152 |
| | fig\|163164.1.peg.1020 | CDS | NC_002978 | 1090833 | 1089286 |
| | fig\|163164.1.peg.1198 | CDS | NC_002978 | 48840 | 48064 |
| | fig\|163164.1.peg.1218 | CDS | NC_002978 | 245008 | 243932 |
| | fig\|163164.1.peg.1241 | CDS | NC_002978 | 378869 | 377574 |
| | fig\|163164.1.peg.1246 | CDS | NC_002978 | 487718 | 486642 |
| | fig\|163164.1.peg.1248 | CDS | NC_002978 | 509575 | 508823 |
| | fig\|163164.1.peg.1251 | CDS | NC_002978 | 523124 | 524419 |
| | fig\|163164.1.peg.1260 | CDS | NC_002978 | 583296 | 584372 |
| wMhie | - | - | - | - | - |
| wLbra | - | - | - | - | - |
| wLsig | - | - | - | - | - |
| wLug | fig\|1335053.6.peg.47 | CDS | MUIY01000001.1 | 45754 | 47004 |
| | fig\|1335053.6.peg.55 | CDS | MUIY01000001.1 | 52785 | 54035 |

| | | | | | |
|---|---|---|---|---|---|
| | fig\|1335053.6.peg.94 | CDS | MUIY01000001.1 | 85363 | 84092 |
| | fig\|1335053.6.peg.598 | CDS | MUIY01000001.1 | 559490 | 560740 |
| | fig\|1335053.6.peg.651 | CDS | MUIY01000001.1 | 604803 | 606050 |
| | fig\|1335053.6.peg.909 | CDS | MUIY01000001.1 | 842776 | 843441 |
| | fig\|1335053.6.peg.914 | CDS | MUIY01000001.1 | 849817 | 848855 |
| | fig\|1335053.6.peg.994 | CDS | MUIY01000001.1 | 932421 | 931174 |
| | fig\|1335053.6.peg.1161 | CDS | MUIY01000001.1 | 1091876 | 1093126 |
| | fig\|1335053.6.peg.1219 | CDS | MUIY01000001.1 | 1144340 | 1143090 |
| | fig\|1335053.6.peg.1258 | CDS | MUIY01000002.1 | 18649 | 17402 |
| | fig\|1335053.6.peg.1442 | CDS | MUIY01000002.1 | 214193 | 215443 |
| | fig\|1335053.6.peg.1451 | CDS | MUIY01000002.1 | 224755 | 223667 |
| | fig\|1335053.6.peg.1532 | CDS | MUIY01000002.1 | 306864 | 305617 |
| wNfla | fig\|1854759.6.peg.501 | CDS | NZ_LYUW010000 | 14040 | 13885 |
| | fig\|1854759.6.peg.1363 | CDS | NZ_LYUW010000 | 327 | 37 |
| | fig\|1854759.6.peg.1364 | CDS | NZ_LYUW010000 | 1204 | 308 |
| | fig\|1854759.6.peg.1441 | CDS | NZ_LYUW010001 | 535 | 1785 |
| | fig\|1854759.6.peg.1507 | CDS | NZ_LYUW010001 | 243 | 1 |
| | fig\|1854759.6.peg.1509 | CDS | NZ_LYUW010001 | 369 | 1 |
| wOv | - | - | - | - | - |
| wOo | - | - | - | - | - |
| wPip | fig\|955.1.peg.354 | CDS | culex88a09.p1k | 1374 | 2717 |
| | fig\|955.1.peg.1056 | CDS | culex35f07.p1k | 1768 | 1013 |
| | fig\|955.1.peg.1057 | CDS | culex35f07.p1k | 3045 | 1801 |
| | fig\|955.1.peg.1059 | CDS | culex35f07.p1k | 4408 | 3557 |
| | fig\|955.1.peg.1062 | CDS | culex35f07.p1k | 7184 | 5841 |
| | fig\|955.1.peg.1375 | CDS | culex143c03.p1k | 18716 | 17415 |
| wPpe | - | - | - | - | - |
| wstri | fig\|368602.8.peg.17 | CDS | MUIX01000001.1 | 14980 | 13754 |
| | fig\|368602.8.peg.28 | CDS | MUIX01000001.1 | 22475 | 23326 |
| | fig\|368602.8.peg.30 | CDS | MUIX01000001.1 | 23840 | 25084 |
| | fig\|368602.8.peg.31 | CDS | MUIX01000001.1 | 25138 | 25872 |
| | fig\|368602.8.peg.37 | CDS | MUIX01000001.1 | 28602 | 29453 |
| | fig\|368602.8.peg.39 | CDS | MUIX01000001.1 | 29967 | 31211 |
| | fig\|368602.8.peg.40 | CDS | MUIX01000001.1 | 31265 | 32230 |
| | fig\|368602.8.peg.43 | CDS | MUIX01000001.1 | 38813 | 40057 |
| | fig\|368602.8.peg.44 | CDS | MUIX01000001.1 | 40111 | 40845 |
| | fig\|368602.8.peg.81 | CDS | MUIX01000001.1 | 75109 | 73883 |
| | fig\|368602.8.peg.108 | CDS | MUIX01000001.1 | 98449 | 97136 |
| | fig\|368602.8.peg.132 | CDS | MUIX01000001.1 | 118288 | 119757 |
| | fig\|368602.8.peg.190 | CDS | MUIX01000001.1 | 177581 | 176355 |
| | fig\|368602.8.peg.208 | CDS | MUIX01000001.1 | 190567 | 191880 |
| | fig\|368602.8.peg.219 | CDS | MUIX01000001.1 | 210525 | 210046 |

| | | | | | |
|---|---|---|---|---|---|
| | fig\|368602.8.peg.279 | CDS | MUIX01000001.1 | 258901 | 257630 |
| | fig\|368602.8.peg.342 | CDS | MUIX01000001.1 | 318580 | 317309 |
| | fig\|368602.8.peg.353 | CDS | MUIX01000001.1 | 329350 | 330594 |
| | fig\|368602.8.peg.354 | CDS | MUIX01000001.1 | 330648 | 331382 |
| | fig\|368602.8.peg.394 | CDS | MUIX01000001.1 | 373387 | 372161 |
| | fig\|368602.8.peg.644 | CDS | MUIX01000001.1 | 642493 | 643974 |
| | fig\|368602.8.peg.656 | CDS | MUIX01000001.1 | 657736 | 656510 |
| | fig\|368602.8.peg.660 | CDS | MUIX01000001.1 | 662316 | 661003 |
| | fig\|368602.8.peg.701 | CDS | MUIX01000001.1 | 696321 | 695050 |
| | fig\|368602.8.peg.767 | CDS | MUIX01000001.1 | 781667 | 780198 |
| | fig\|368602.8.peg.977 | CDS | MUIX01000001.1 | 970601 | 969375 |
| | fig\|368602.8.peg.1080 | CDS | MUIX01000001.1 | 1060427 | 1059156 |
| | fig\|368602.8.peg.1261 | CDS | MUIX01000001.1 | 1239295 | 1238561 |
| | fig\|368602.8.peg.1262 | CDS | MUIX01000001.1 | 1240593 | 1239349 |
| | fig\|368602.8.peg.1264 | CDS | MUIX01000001.1 | 1241958 | 1241107 |
| | fig\|368602.8.peg.1312 | CDS | MUIX01000001.1 | 1287747 | 1288988 |
| | fig\|368602.8.peg.1343 | CDS | MUIX01000001.1 | 1314385 | 1315854 |
| | fig\|368602.8.peg.1376 | CDS | MUIX01000001.1 | 1340599 | 1342146 |
| | fig\|368602.8.peg.1443 | CDS | MUIX01000001.1 | 1413840 | 1415321 |
| | fig\|368602.8.peg.1506 | CDS | MUIX01000001.1 | 1474450 | 1473209 |
| | fig\|368602.8.peg.1514 | CDS | MUIX01000001.1 | 1481600 | 1480119 |
| | fig\|368602.8.peg.1524 | CDS | MUIX01000001.1 | 1488474 | 1489700 |
| | fig\|368602.8.peg.1580 | CDS | MUIX01000001.1 | 1549498 | 1550769 |
| | fig\|368602.8.peg.1587 | CDS | MUIX01000001.1 | 1561281 | 1562594 |
| | fig\|368602.8.peg.1591 | CDS | MUIX01000001.1 | 1568851 | 1567370 |
| | fig\|368602.8.peg.1599 | CDS | MUIX01000001.1 | 1576075 | 1577346 |
| | fig\|368602.8.peg.1604 | CDS | MUIX01000001.1 | 1582692 | 1581451 |
| | fig\|368602.8.peg.1610 | CDS | MUIX01000001.1 | 1586192 | 1584879 |
| | fig\|368602.8.peg.1618 | CDS | MUIX01000001.1 | 1592503 | 1591262 |
| | fig\|368602.8.peg.1624 | CDS | MUIX01000001.1 | 1596003 | 1594690 |
| | fig\|368602.8.peg.1654 | CDS | MUIX01000002.1 | 16889 | 15576 |
| | fig\|368602.8.peg.1657 | CDS | MUIX01000002.1 | 19278 | 20759 |
| | fig\|368602.8.peg.1659 | CDS | MUIX01000002.1 | 21319 | 21798 |
| | fig\|368602.8.peg.1708 | CDS | MUIX01000002.1 | 63777 | 62464 |
| | fig\|368602.8.peg.1715 | CDS | MUIX01000002.1 | 70074 | 71345 |
| | fig\|368602.8.peg.1762 | CDS | MUIX01000002.1 | 109901 | 108630 |
| | fig\|368602.8.peg.1798 | CDS | MUIX01000002.1 | 152978 | 154162 |
| | fig\|368602.8.peg.1799 | CDS | MUIX01000002.1 | 154216 | 154950 |
| | fig\|368602.8.peg.1805 | CDS | MUIX01000002.1 | 160079 | 161305 |
| wTpre | fig\|125593.4.peg.526 | CDS | NZ_CM003641.1 | 421271 | 420969 |
| | fig\|125593.4.peg.527 | CDS | NZ_CM003641.1 | 421627 | 421268 |
| | fig\|125593.4.peg.528 | CDS | NZ_CM003641.1 | 421909 | 421640 |

| | | | | | |
|---|---|---|---|---|---|
| wVulC | fig\|1220511.6.peg.141 | CDS | ALWU01000001.1 | 131094 | 130579 |
| | fig\|1220511.6.peg.214 | CDS | ALWU01000001.1 | 194427 | 193747 |
| | fig\|1220511.6.peg.216 | CDS | ALWU01000001.1 | 194982 | 195479 |
| | fig\|1220511.6.peg.344 | CDS | ALWU01000001.1 | 300284 | 299736 |
| | fig\|1220511.6.peg.345 | CDS | ALWU01000001.1 | 301515 | 300343 |
| | fig\|1220511.6.peg.347 | CDS | ALWU01000001.1 | 302964 | 302620 |
| | fig\|1220511.6.peg.352 | CDS | ALWU01000001.1 | 304451 | 304873 |
| | fig\|1220511.6.peg.363 | CDS | ALWU01000001.1 | 311311 | 311826 |
| | fig\|1220511.6.peg.379 | CDS | ALWU01000001.1 | 323123 | 322341 |
| | fig\|1220511.6.peg.699 | CDS | ALWU01000002.1 | 14691 | 13576 |
| | fig\|1220511.6.peg.700 | CDS | ALWU01000002.1 | 15415 | 14744 |
| | fig\|1220511.6.peg.847 | CDS | ALWU01000005.1 | 30121 | 30366 |
| | fig\|1220511.6.peg.857 | CDS | ALWU01000005.1 | 36848 | 37093 |
| | fig\|1220511.6.peg.946 | CDS | ALWU01000005.1 | 113463 | 112963 |
| | fig\|1220511.6.peg.1040 | CDS | ALWU01000005.1 | 190520 | 192301 |
| | fig\|1220511.6.peg.1079 | CDS | ALWU01000006.1 | 3901 | 5682 |
| | fig\|1220511.6.peg.1108 | CDS | ALWU01000006.1 | 28951 | 29604 |
| | fig\|1220511.6.peg.1113 | CDS | ALWU01000006.1 | 32799 | 33422 |
| | fig\|1220511.6.peg.1327 | CDS | ALWU01000008.1 | 175500 | 174985 |
| | fig\|1220511.6.peg.1530 | CDS | ALWU01000010.1 | 148506 | 148321 |
| | fig\|1220511.6.peg.1602 | CDS | ALWU01000010.1 | 208558 | 208139 |
| | fig\|1220511.6.peg.1603 | CDS | ALWU01000010.1 | 208853 | 208548 |
| | fig\|1220511.6.peg.1610 | CDS | ALWU01000010.1 | 212470 | 212775 |
| | fig\|1220511.6.peg.1611 | CDS | ALWU01000010.1 | 212765 | 213184 |
| | fig\|1220511.6.peg.1690 | CDS | ALWU01000010.1 | 276574 | 277398 |
| | fig\|1220511.6.peg.1691 | CDS | ALWU01000010.1 | 277373 | 277630 |
| | fig\|1220511.6.peg.1692 | CDS | ALWU01000010.1 | 277620 | 278039 |
| | fig\|1220511.6.peg.1799 | CDS | ALWU01000010.1 | 377187 | 376435 |
| | fig\|1220511.6.peg.1801 | CDS | ALWU01000010.1 | 377742 | 378239 |
| | fig\|1220511.6.peg.1941 | CDS | ALWU01000010.1 | 507287 | 507802 |
| wWb | - | - | - | - | - |

| Strand | Length (bp) | Function |
|---|---|---|
| - | - | - |
| - | - | - |
| + | 276 | Retron-type RNA-directed DNA polymerase (EC 2.7.7.49) |
| + | 1314 | Retron-type RNA-directed DNA polymerase (EC 2.7.7.49) |
| - | - | - |
| - | 903 | Retron-type RNA-directed DNA polymerase (EC 2.7.7.49) |
| - | 1011 | Retron-type RNA-directed DNA polymerase (EC 2.7.7.49) |
| + | 1647 | Retron-type RNA-directed DNA polymerase (EC 2.7.7.49) |
| + | 1377 | Retron-type RNA-directed DNA polymerase (EC 2.7.7.49) |
| + | 1377 | Retron-type RNA-directed DNA polymerase (EC 2.7.7.49) |
| + | 1377 | Retron-type RNA-directed DNA polymerase (EC 2.7.7.49) |
| - | 1638 | Retron-type RNA-directed DNA polymerase (EC 2.7.7.49) |
| + | 1482 | Retron-type RNA-directed DNA polymerase (EC 2.7.7.49) |
| - | 1872 | Retron-type RNA-directed DNA polymerase (EC 2.7.7.49) |
| - | 1332 | Retron-type RNA-directed DNA polymerase (EC 2.7.7.49) |
| + | 1638 | Retron-type RNA-directed DNA polymerase (EC 2.7.7.49) |
| - | - | - |
| - | - | - |
| - | - | - |
| + | 999 | Retron-type RNA-directed DNA polymerase (EC 2.7.7.49) |
| - | 765 | Retron-type RNA-directed DNA polymerase (EC 2.7.7.49) |
| + | 321 | Retron-type RNA-directed DNA polymerase (EC 2.7.7.49) |
| + | 1041 | Retron-type RNA-directed DNA polymerase (EC 2.7.7.49) |
| + | 444 | Retron-type RNA-directed DNA polymerase (EC 2.7.7.49) |
| + | 1548 | Retron-type RNA-directed DNA polymerase (EC 2.7.7.49) |
| + | 1548 | Retron-type RNA-directed DNA polymerase (EC 2.7.7.49) |
| - | 1548 | Retron-type RNA-directed DNA polymerase (EC 2.7.7.49) |
| - | 777 | Retron-type RNA-directed DNA polymerase (EC 2.7.7.49) |
| - | 1077 | Retron-type RNA-directed DNA polymerase (EC 2.7.7.49) |
| - | 1296 | Retron-type RNA-directed DNA polymerase (EC 2.7.7.49) |
| - | 1077 | Retron-type RNA-directed DNA polymerase (EC 2.7.7.49) |
| - | 753 | Retron-type RNA-directed DNA polymerase (EC 2.7.7.49) |
| + | 1296 | Retron-type RNA-directed DNA polymerase (EC 2.7.7.49) |
| + | 1077 | Retron-type RNA-directed DNA polymerase (EC 2.7.7.49) |
| - | - | - |
| - | - | - |
| - | - | - |
| + | 1251 | Retron-type RNA-directed DNA polymerase (EC 2.7.7.49) |
| + | 1251 | Retron-type RNA-directed DNA polymerase (EC 2.7.7.49) |

| | | | |
|---|---|---|---|
| - | | 1272 | Retron-type RNA-directed DNA polymerase (EC 2.7.7.49) |
| + | | 1251 | Retron-type RNA-directed DNA polymerase (EC 2.7.7.49) |
| + | | 1248 | Retron-type RNA-directed DNA polymerase (EC 2.7.7.49) |
| + | | 666 | Retron-type RNA-directed DNA polymerase (EC 2.7.7.49) |
| - | | 963 | Retron-type RNA-directed DNA polymerase (EC 2.7.7.49) |
| - | | 1248 | Retron-type RNA-directed DNA polymerase (EC 2.7.7.49) |
| + | | 1251 | Retron-type RNA-directed DNA polymerase (EC 2.7.7.49) |
| - | | 1251 | Retron-type RNA-directed DNA polymerase (EC 2.7.7.49) |
| - | | 1248 | Retron-type RNA-directed DNA polymerase (EC 2.7.7.49) |
| + | | 1251 | Retron-type RNA-directed DNA polymerase (EC 2.7.7.49) |
| - | | 1089 | Retron-type RNA-directed DNA polymerase (EC 2.7.7.49) |
| - | | 1248 | Retron-type RNA-directed DNA polymerase (EC 2.7.7.49) |
| - | | 156 | Retron-type RNA-directed DNA polymerase (EC 2.7.7.49) |
| - | | 291 | Retron-type RNA-directed DNA polymerase (EC 2.7.7.49) |
| - | | 897 | Retron-type RNA-directed DNA polymerase (EC 2.7.7.49) |
| + | | 1251 | Retron-type RNA-directed DNA polymerase (EC 2.7.7.49) |
| - | | 243 | Retron-type RNA-directed DNA polymerase (EC 2.7.7.49) |
| - | | 369 | Retron-type RNA-directed DNA polymerase (EC 2.7.7.49) |
| - | - | | - |
| - | - | | - |
| + | | 1344 | Retron-type RNA-directed DNA polymerase (EC 2.7.7.49) |
| - | | 756 | Retron-type RNA-directed DNA polymerase (EC 2.7.7.49) |
| - | | 1245 | Retron-type RNA-directed DNA polymerase (EC 2.7.7.49) |
| - | | 852 | Retron-type RNA-directed DNA polymerase (EC 2.7.7.49) |
| - | | 1344 | Retron-type RNA-directed DNA polymerase (EC 2.7.7.49) |
| - | | 1302 | Retron-type RNA-directed DNA polymerase (EC 2.7.7.49) |
| - | - | | - |
| - | | 1227 | Retron-type RNA-directed DNA polymerase (EC 2.7.7.49) |
| + | | 852 | Retron-type RNA-directed DNA polymerase (EC 2.7.7.49) |
| + | | 1245 | Retron-type RNA-directed DNA polymerase (EC 2.7.7.49) |
| + | | 735 | Retron-type RNA-directed DNA polymerase (EC 2.7.7.49) |
| + | | 852 | Retron-type RNA-directed DNA polymerase (EC 2.7.7.49) |
| + | | 1245 | Retron-type RNA-directed DNA polymerase (EC 2.7.7.49) |
| + | | 966 | Retron-type RNA-directed DNA polymerase (EC 2.7.7.49) |
| + | | 1245 | Retron-type RNA-directed DNA polymerase (EC 2.7.7.49) |
| + | | 735 | Retron-type RNA-directed DNA polymerase (EC 2.7.7.49) |
| - | | 1227 | Retron-type RNA-directed DNA polymerase (EC 2.7.7.49) |
| - | | 1314 | Retron-type RNA-directed DNA polymerase (EC 2.7.7.49) |
| + | | 1470 | Retron-type RNA-directed DNA polymerase (EC 2.7.7.49) |
| - | | 1227 | Retron-type RNA-directed DNA polymerase (EC 2.7.7.49) |
| + | | 1314 | Retron-type RNA-directed DNA polymerase (EC 2.7.7.49) |
| - | | 480 | Retron-type RNA-directed DNA polymerase (EC 2.7.7.49) |

| | | |
|---|---|---|
| - | 1272 | Retron-type RNA-directed DNA polymerase (EC 2.7.7.49) |
| - | 1272 | Retron-type RNA-directed DNA polymerase (EC 2.7.7.49) |
| + | 1245 | Retron-type RNA-directed DNA polymerase (EC 2.7.7.49) |
| + | 735 | Retron-type RNA-directed DNA polymerase (EC 2.7.7.49) |
| - | 1227 | Retron-type RNA-directed DNA polymerase (EC 2.7.7.49) |
| + | 1482 | Retron-type RNA-directed DNA polymerase (EC 2.7.7.49) |
| - | 1227 | Retron-type RNA-directed DNA polymerase (EC 2.7.7.49) |
| - | 1314 | Retron-type RNA-directed DNA polymerase (EC 2.7.7.49) |
| - | 1272 | Retron-type RNA-directed DNA polymerase (EC 2.7.7.49) |
| - | 1470 | Retron-type RNA-directed DNA polymerase (EC 2.7.7.49) |
| - | 1227 | Retron-type RNA-directed DNA polymerase (EC 2.7.7.49) |
| - | 1272 | Retron-type RNA-directed DNA polymerase (EC 2.7.7.49) |
| - | 735 | Retron-type RNA-directed DNA polymerase (EC 2.7.7.49) |
| - | 1245 | Retron-type RNA-directed DNA polymerase (EC 2.7.7.49) |
| - | 852 | Retron-type RNA-directed DNA polymerase (EC 2.7.7.49) |
| + | 1242 | Retron-type RNA-directed DNA polymerase (EC 2.7.7.49) |
| + | 1470 | Retron-type RNA-directed DNA polymerase (EC 2.7.7.49) |
| + | 1548 | Retron-type RNA-directed DNA polymerase (EC 2.7.7.49) |
| + | 1482 | Retron-type RNA-directed DNA polymerase (EC 2.7.7.49) |
| - | 1242 | Retron-type RNA-directed DNA polymerase (EC 2.7.7.49) |
| - | 1482 | Retron-type RNA-directed DNA polymerase (EC 2.7.7.49) |
| + | 1227 | Retron-type RNA-directed DNA polymerase (EC 2.7.7.49) |
| + | 1272 | Retron-type RNA-directed DNA polymerase (EC 2.7.7.49) |
| + | 1314 | Retron-type RNA-directed DNA polymerase (EC 2.7.7.49) |
| - | 1482 | Retron-type RNA-directed DNA polymerase (EC 2.7.7.49) |
| + | 1272 | Retron-type RNA-directed DNA polymerase (EC 2.7.7.49) |
| - | 1242 | Retron-type RNA-directed DNA polymerase (EC 2.7.7.49) |
| - | 1314 | Retron-type RNA-directed DNA polymerase (EC 2.7.7.49) |
| - | 1242 | Retron-type RNA-directed DNA polymerase (EC 2.7.7.49) |
| - | 1314 | Retron-type RNA-directed DNA polymerase (EC 2.7.7.49) |
| - | 1314 | Retron-type RNA-directed DNA polymerase (EC 2.7.7.49) |
| + | 1482 | Retron-type RNA-directed DNA polymerase (EC 2.7.7.49) |
| + | 480 | Retron-type RNA-directed DNA polymerase (EC 2.7.7.49) |
| - | 1314 | Retron-type RNA-directed DNA polymerase (EC 2.7.7.49) |
| + | 1272 | Retron-type RNA-directed DNA polymerase (EC 2.7.7.49) |
| - | 1272 | Retron-type RNA-directed DNA polymerase (EC 2.7.7.49) |
| + | 1185 | Retron-type RNA-directed DNA polymerase (EC 2.7.7.49) |
| + | 735 | Retron-type RNA-directed DNA polymerase (EC 2.7.7.49) |
| + | 1227 | Retron-type RNA-directed DNA polymerase (EC 2.7.7.49) |
| - | 303 | Retron-type RNA-directed DNA polymerase (EC 2.7.7.49) |
| - | 360 | Retron-type RNA-directed DNA polymerase (EC 2.7.7.49) |
| - | 270 | Retron-type RNA-directed DNA polymerase (EC 2.7.7.49) |

| | | |
|---|---|---|
| - | 516 | Retron-type RNA-directed DNA polymerase (EC 2.7.7.49) |
| - | 681 | Retron-type RNA-directed DNA polymerase (EC 2.7.7.49) |
| + | 498 | Retron-type RNA-directed DNA polymerase (EC 2.7.7.49) |
| - | 549 | Retron-type RNA-directed DNA polymerase (EC 2.7.7.49) |
| - | 1173 | Retron-type RNA-directed DNA polymerase (EC 2.7.7.49) |
| - | 345 | Retron-type RNA-directed DNA polymerase (EC 2.7.7.49) |
| + | 423 | Retron-type RNA-directed DNA polymerase (EC 2.7.7.49) |
| + | 516 | Retron-type RNA-directed DNA polymerase (EC 2.7.7.49) |
| - | 783 | Retron-type RNA-directed DNA polymerase (EC 2.7.7.49) |
| - | 1116 | Retron-type RNA-directed DNA polymerase (EC 2.7.7.49) |
| - | 672 | Retron-type RNA-directed DNA polymerase (EC 2.7.7.49) |
| + | 246 | Retron-type RNA-directed DNA polymerase (EC 2.7.7.49) |
| + | 246 | Retron-type RNA-directed DNA polymerase (EC 2.7.7.49) |
| - | 501 | Retron-type RNA-directed DNA polymerase (EC 2.7.7.49) |
| + | 1782 | Retron-type RNA-directed DNA polymerase (EC 2.7.7.49) |
| + | 1782 | Retron-type RNA-directed DNA polymerase (EC 2.7.7.49) |
| + | 654 | Retron-type RNA-directed DNA polymerase (EC 2.7.7.49) |
| + | 624 | Retron-type RNA-directed DNA polymerase (EC 2.7.7.49) |
| - | 516 | Retron-type RNA-directed DNA polymerase (EC 2.7.7.49) |
| - | 186 | Retron-type RNA-directed DNA polymerase (EC 2.7.7.49) |
| - | 420 | Retron-type RNA-directed DNA polymerase (EC 2.7.7.49) |
| - | 306 | Retron-type RNA-directed DNA polymerase (EC 2.7.7.49) |
| + | 306 | Retron-type RNA-directed DNA polymerase (EC 2.7.7.49) |
| + | 420 | Retron-type RNA-directed DNA polymerase (EC 2.7.7.49) |
| + | 825 | Retron-type RNA-directed DNA polymerase (EC 2.7.7.49) |
| + | 258 | Retron-type RNA-directed DNA polymerase (EC 2.7.7.49) |
| + | 420 | Retron-type RNA-directed DNA polymerase (EC 2.7.7.49) |
| - | 753 | Retron-type RNA-directed DNA polymerase (EC 2.7.7.49) |
| + | 498 | Retron-type RNA-directed DNA polymerase (EC 2.7.7.49) |
| + | 516 | Retron-type RNA-directed DNA polymerase (EC 2.7.7.49) |
| - | - | - |

| Subsystem |
| --- |
| - |
| - |
| Group II intron-associated genes |
| Group II intron-associated genes |
| - |
| Group II intron-associated genes |
| Group II intron-associated genes |
| Group II intron-associated genes |
| Group II intron-associated genes |
| Group II intron-associated genes |
| Group II intron-associated genes |
| Group II intron-associated genes |
| Group II intron-associated genes |
| Group II intron-associated genes |
| Group II intron-associated genes |
| Group II intron-associated genes |
| - |
| - |
| - |
| Group II intron-associated genes |
| Group II intron-associated genes |
| Group II intron-associated genes |
| Group II intron-associated genes |
| Group II intron-associated genes |
| Group II intron-associated genes |
| Group II intron-associated genes |
| Group II intron-associated genes |
| Group II intron-associated genes |
| Group II intron-associated genes |
| Group II intron-associated genes |
| Group II intron-associated genes |
| Group II intron-associated genes |
| Group II intron-associated genes |
| Group II intron-associated genes |
| - |
| - |
| - |
| Group II intron-associated genes |
| Group II intron-associated genes |

| |
|---|
| Group II intron-associated genes |
| Group II intron-associated genes |
| Group II intron-associated genes |
| Group II intron-associated genes |
| Group II intron-associated genes |
| Group II intron-associated genes |
| Group II intron-associated genes |
| Group II intron-associated genes |
| Group II intron-associated genes |
| Group II intron-associated genes |
| Group II intron-associated genes |
| Group II intron-associated genes |
| Group II intron-associated genes |
| Group II intron-associated genes |
| Group II intron-associated genes |
| Group II intron-associated genes |
| Group II intron-associated genes |
| Group II intron-associated genes |
| - |
| - |
| Group II intron-associated genes |
| Group II intron-associated genes |
| Group II intron-associated genes |
| Group II intron-associated genes |
| Group II intron-associated genes |
| Group II intron-associated genes |
| - |
| Group II intron-associated genes |
| Group II intron-associated genes |
| Group II intron-associated genes |
| Group II intron-associated genes |
| Group II intron-associated genes |
| Group II intron-associated genes |
| Group II intron-associated genes |
| Group II intron-associated genes |
| Group II intron-associated genes |
| Group II intron-associated genes |
| Group II intron-associated genes |
| Group II intron-associated genes |
| Group II intron-associated genes |
| Group II intron-associated genes |
| Group II intron-associated genes |

| |
|---|
| Group II intron-associated genes |
| Group II intron-associated genes |
| Group II intron-associated genes |
| Group II intron-associated genes |
| Group II intron-associated genes |
| Group II intron-associated genes |
| Group II intron-associated genes |
| Group II intron-associated genes |
| Group II intron-associated genes |
| Group II intron-associated genes |
| Group II intron-associated genes |
| Group II intron-associated genes |
| Group II intron-associated genes |
| Group II intron-associated genes |
| Group II intron-associated genes |
| Group II intron-associated genes |
| Group II intron-associated genes |
| Group II intron-associated genes |
| Group II intron-associated genes |
| Group II intron-associated genes |
| Group II intron-associated genes |
| Group II intron-associated genes |
| Group II intron-associated genes |
| Group II intron-associated genes |
| Group II intron-associated genes |
| Group II intron-associated genes |
| Group II intron-associated genes |
| Group II intron-associated genes |
| Group II intron-associated genes |
| Group II intron-associated genes |
| Group II intron-associated genes |
| Group II intron-associated genes |
| Group II intron-associated genes |
| Group II intron-associated genes |
| Group II intron-associated genes |
| Group II intron-associated genes |
| Group II intron-associated genes |
| Group II intron-associated genes |
| Group II intron-associated genes |
| Group II intron-associated genes |
| Group II intron-associated genes |
| Group II intron-associated genes |

| |
|---|
| Group II intron-associated genes |
| Group II intron-associated genes |
| Group II intron-associated genes |
| Group II intron-associated genes |
| Group II intron-associated genes |
| Group II intron-associated genes |
| Group II intron-associated genes |
| Group II intron-associated genes |
| Group II intron-associated genes |
| Group II intron-associated genes |
| Group II intron-associated genes |
| Group II intron-associated genes |
| Group II intron-associated genes |
| Group II intron-associated genes |
| Group II intron-associated genes |
| Group II intron-associated genes |
| Group II intron-associated genes |
| Group II intron-associated genes |
| Group II intron-associated genes |
| Group II intron-associated genes |
| Group II intron-associated genes |
| Group II intron-associated genes |
| Group II intron-associated genes |
| Group II intron-associated genes |
| Group II intron-associated genes |
| Group II intron-associated genes |
| Group II intron-associated genes |
| Group II intron-associated genes |
| Group II intron-associated genes |
| Group II intron-associated genes |
| - |

**Table S8. List of mobile element genes detected using RAST pipeline.**

| genome | Feature ID | Type | Contig | Start | Stop |
|---|---|---|---|---|---|
| wBm | fig\|292805.13.peg.74 | CDS | wBm_reseq_reve | 45903 | 45631 |
| | fig\|292805.13.peg.515 | CDS | wBm_reseq_reve | 356167 | 355979 |
| | fig\|292805.13.peg.760 | CDS | wBm_reseq_reve | 522826 | 522984 |
| wBp | fig\|96495.5.peg.194 | CDS | NZ_CP050521.1 | 152947 | 153219 |
| | fig\|96495.5.peg.254 | CDS | NZ_CP050521.1 | 200393 | 200121 |
| | fig\|96495.5.peg.1182 | CDS | NZ_CP050521.1 | 969737 | 969579 |
| | fig\|96495.5.peg.1215 | CDS | NZ_CP050521.1 | 991912 | 992184 |
| | fig\|96495.5.peg.1244 | CDS | NZ_CP050521.1 | 1028753 | 1028592 |
| | fig\|96495.5.peg.1285 | CDS | NZ_CP050521.1 | 1054315 | 1054587 |
| | fig\|96495.5.peg.1302 | CDS | NZ_CP050521.1 | 1068612 | 1068884 |
| wCtub | - | - | - | - | - |
| wCauA | fig\|2591635.5.peg.23 | CDS | CP041215.1 | 24095 | 24379 |
| | fig\|2591635.5.peg.25 | CDS | CP041215.1 | 25017 | 25145 |
| | fig\|2591635.5.peg.34 | CDS | CP041215.1 | 35460 | 35825 |
| | fig\|2591635.5.peg.53 | CDS | CP041215.1 | 51796 | 52161 |
| | fig\|2591635.5.peg.58 | CDS | CP041215.1 | 54789 | 55154 |
| | fig\|2591635.5.peg.70 | CDS | CP041215.1 | 65690 | 65502 |
| | fig\|2591635.5.peg.74 | CDS | CP041215.1 | 66495 | 66292 |
| | fig\|2591635.5.peg.76 | CDS | CP041215.1 | 67349 | 66984 |
| | fig\|2591635.5.peg.139 | CDS | CP041215.1 | 122153 | 122482 |
| | fig\|2591635.5.peg.140 | CDS | CP041215.1 | 122466 | 123101 |
| | fig\|2591635.5.peg.150 | CDS | CP041215.1 | 131020 | 131289 |
| | fig\|2591635.5.peg.156 | CDS | CP041215.1 | 136492 | 136857 |
| | fig\|2591635.5.peg.206 | CDS | CP041215.1 | 190832 | 190197 |
| | fig\|2591635.5.peg.207 | CDS | CP041215.1 | 191145 | 190816 |
| | fig\|2591635.5.peg.218 | CDS | CP041215.1 | 201739 | 201374 |
| | fig\|2591635.5.peg.232 | CDS | CP041215.1 | 216153 | 215788 |
| | fig\|2591635.5.peg.259 | CDS | CP041215.1 | 241662 | 241297 |
| | fig\|2591635.5.peg.262 | CDS | CP041215.1 | 243248 | 242916 |
| | fig\|2591635.5.peg.276 | CDS | CP041215.1 | 254122 | 253757 |
| | fig\|2591635.5.peg.299 | CDS | CP041215.1 | 270346 | 270651 |
| | fig\|2591635.5.peg.311 | CDS | CP041215.1 | 287679 | 287314 |
| | fig\|2591635.5.peg.331 | CDS | CP041215.1 | 305256 | 305621 |
| | fig\|2591635.5.peg.369 | CDS | CP041215.1 | 339797 | 340111 |
| | fig\|2591635.5.peg.370 | CDS | CP041215.1 | 340181 | 341134 |
| | fig\|2591635.5.peg.393 | CDS | CP041215.1 | 374395 | 374769 |
| | fig\|2591635.5.peg.394 | CDS | CP041215.1 | 374821 | 375489 |
| | fig\|2591635.5.peg.395 | CDS | CP041215.1 | 375537 | 375665 |
| | fig\|2591635.5.peg.397 | CDS | CP041215.1 | 376428 | 376063 |

| | | | | |
|---|---|---|---|---|
| fig\|2591635.5.peg.410 | CDS | CP041215.1 | 385857 | 385492 |
| fig\|2591635.5.peg.411 | CDS | CP041215.1 | 386001 | 385900 |
| fig\|2591635.5.peg.412 | CDS | CP041215.1 | 386293 | 386081 |
| fig\|2591635.5.peg.413 | CDS | CP041215.1 | 387359 | 386691 |
| fig\|2591635.5.peg.431 | CDS | CP041215.1 | 406509 | 406144 |
| fig\|2591635.5.peg.434 | CDS | CP041215.1 | 408304 | 408140 |
| fig\|2591635.5.peg.437 | CDS | CP041215.1 | 409183 | 408881 |
| fig\|2591635.5.peg.443 | CDS | CP041215.1 | 412418 | 412053 |
| fig\|2591635.5.peg.446 | CDS | CP041215.1 | 413999 | 413634 |
| fig\|2591635.5.peg.448 | CDS | CP041215.1 | 415537 | 415902 |
| fig\|2591635.5.peg.479 | CDS | CP041215.1 | 443377 | 443742 |
| fig\|2591635.5.peg.483 | CDS | CP041215.1 | 445690 | 445325 |
| fig\|2591635.5.peg.504 | CDS | CP041215.1 | 460412 | 460714 |
| fig\|2591635.5.peg.507 | CDS | CP041215.1 | 461291 | 461455 |
| fig\|2591635.5.peg.516 | CDS | CP041215.1 | 470726 | 471154 |
| fig\|2591635.5.peg.517 | CDS | CP041215.1 | 471637 | 471449 |
| fig\|2591635.5.peg.518 | CDS | CP041215.1 | 471943 | 472890 |
| fig\|2591635.5.peg.528 | CDS | CP041215.1 | 482617 | 482982 |
| fig\|2591635.5.peg.535 | CDS | CP041215.1 | 488566 | 488931 |
| fig\|2591635.5.peg.537 | CDS | CP041215.1 | 489465 | 489749 |
| fig\|2591635.5.peg.570 | CDS | CP041215.1 | 519654 | 519289 |
| fig\|2591635.5.peg.573 | CDS | CP041215.1 | 521550 | 521843 |
| fig\|2591635.5.peg.576 | CDS | CP041215.1 | 522420 | 522584 |
| fig\|2591635.5.peg.606 | CDS | CP041215.1 | 549419 | 549784 |
| fig\|2591635.5.peg.608 | CDS | CP041215.1 | 550413 | 550192 |
| fig\|2591635.5.peg.646 | CDS | CP041215.1 | 598063 | 597698 |
| fig\|2591635.5.peg.661 | CDS | CP041215.1 | 612161 | 611901 |
| fig\|2591635.5.peg.681 | CDS | CP041215.1 | 637982 | 638347 |
| fig\|2591635.5.peg.704 | CDS | CP041215.1 | 661183 | 660818 |
| fig\|2591635.5.peg.720 | CDS | CP041215.1 | 679773 | 679408 |
| fig\|2591635.5.peg.726 | CDS | CP041215.1 | 684487 | 683333 |
| fig\|2591635.5.peg.731 | CDS | CP041215.1 | 688898 | 688614 |
| fig\|2591635.5.peg.742 | CDS | CP041215.1 | 699152 | 699517 |
| fig\|2591635.5.peg.758 | CDS | CP041215.1 | 723892 | 724257 |
| fig\|2591635.5.peg.760 | CDS | CP041215.1 | 724793 | 724665 |
| fig\|2591635.5.peg.773 | CDS | CP041215.1 | 733962 | 734327 |
| fig\|2591635.5.peg.775 | CDS | CP041215.1 | 735459 | 734944 |
| fig\|2591635.5.peg.776 | CDS | CP041215.1 | 735700 | 735449 |
| fig\|2591635.5.peg.824 | CDS | CP041215.1 | 772655 | 772939 |
| fig\|2591635.5.peg.826 | CDS | CP041215.1 | 773590 | 773955 |
| fig\|2591635.5.peg.833 | CDS | CP041215.1 | 781559 | 781711 |
| fig\|2591635.5.peg.839 | CDS | CP041215.1 | 785191 | 785475 |

| | | | | |
|---|---|---|---|---|
| fig\|2591635.5.peg.847 | CDS | CP041215.1 | 795122 | 794715 |
| fig\|2591635.5.peg.848 | CDS | CP041215.1 | 795342 | 795142 |
| fig\|2591635.5.peg.849 | CDS | CP041215.1 | 795655 | 795326 |
| fig\|2591635.5.peg.852 | CDS | CP041215.1 | 798078 | 798443 |
| fig\|2591635.5.peg.871 | CDS | CP041215.1 | 815731 | 815366 |
| fig\|2591635.5.peg.879 | CDS | CP041215.1 | 823006 | 822641 |
| fig\|2591635.5.peg.892 | CDS | CP041215.1 | 837236 | 836952 |
| fig\|2591635.5.peg.900 | CDS | CP041215.1 | 843638 | 843934 |
| fig\|2591635.5.peg.901 | CDS | CP041215.1 | 844127 | 844492 |
| fig\|2591635.5.peg.908 | CDS | CP041215.1 | 849586 | 849221 |
| fig\|2591635.5.peg.910 | CDS | CP041215.1 | 851497 | 851862 |
| fig\|2591635.5.peg.914 | CDS | CP041215.1 | 855002 | 855271 |
| fig\|2591635.5.peg.921 | CDS | CP041215.1 | 863125 | 863490 |
| fig\|2591635.5.peg.923 | CDS | CP041215.1 | 863979 | 864344 |
| fig\|2591635.5.peg.931 | CDS | CP041215.1 | 869247 | 869549 |
| fig\|2591635.5.peg.934 | CDS | CP041215.1 | 870126 | 870290 |
| fig\|2591635.5.peg.962 | CDS | CP041215.1 | 900311 | 900676 |
| fig\|2591635.5.peg.964 | CDS | CP041215.1 | 902658 | 901315 |
| fig\|2591635.5.peg.978 | CDS | CP041215.1 | 914510 | 914875 |
| fig\|2591635.5.peg.1019 | CDS | CP041215.1 | 958304 | 958597 |
| fig\|2591635.5.peg.1021 | CDS | CP041215.1 | 958979 | 959230 |
| fig\|2591635.5.peg.1088 | CDS | CP041215.1 | 1023095 | 1022730 |
| fig\|2591635.5.peg.1104 | CDS | CP041215.1 | 1048366 | 1048124 |
| fig\|2591635.5.peg.1106 | CDS | CP041215.1 | 1048684 | 1049049 |
| fig\|2591635.5.peg.1110 | CDS | CP041215.1 | 1051302 | 1050937 |
| fig\|2591635.5.peg.1127 | CDS | CP041215.1 | 1064148 | 1063915 |
| fig\|2591635.5.peg.1128 | CDS | CP041215.1 | 1064248 | 1064613 |
| fig\|2591635.5.peg.1167 | CDS | CP041215.1 | 1098415 | 1098720 |
| fig\|2591635.5.peg.1181 | CDS | CP041215.1 | 1117546 | 1117418 |
| fig\|2591635.5.peg.1183 | CDS | CP041215.1 | 1118468 | 1118184 |
| fig\|2591635.5.peg.1211 | CDS | CP041215.1 | 1142602 | 1142474 |
| fig\|2591635.5.peg.1213 | CDS | CP041215.1 | 1143524 | 1143240 |
| fig\|2591635.5.peg.1250 | CDS | CP041215.1 | 1191920 | 1191555 |
| fig\|2591635.5.peg.1279 | CDS | CP041215.1 | 1216896 | 1217261 |
| fig\|2591635.5.peg.1285 | CDS | CP041215.1 | 1224736 | 1224101 |
| fig\|2591635.5.peg.1286 | CDS | CP041215.1 | 1225049 | 1224720 |
| fig\|2591635.5.peg.1303 | CDS | CP041215.1 | 1237452 | 1237087 |
| fig\|2591635.5.peg.1315 | CDS | CP041215.1 | 1250757 | 1251041 |
| fig\|2591635.5.peg.1318 | CDS | CP041215.1 | 1252076 | 1253230 |
| fig\|2591635.5.peg.1335 | CDS | CP041215.1 | 1267744 | 1267109 |
| fig\|2591635.5.peg.1336 | CDS | CP041215.1 | 1268027 | 1267728 |
| fig\|2591635.5.peg.1351 | CDS | CP041215.1 | 1287126 | 1286830 |

| | | | | | |
|---|---|---|---|---|---|
| | fig\|2591635.5.peg.1352 | CDS | CP041215.1 | 1287236 | 1287409 |
| | fig\|2591635.5.peg.1354 | CDS | CP041215.1 | 1287746 | 1288057 |
| | fig\|2591635.5.peg.1356 | CDS | CP041215.1 | 1288532 | 1288714 |
| | fig\|2591635.5.peg.1357 | CDS | CP041215.1 | 1288806 | 1289000 |
| | fig\|2591635.5.peg.1359 | CDS | CP041215.1 | 1289435 | 1289800 |
| | fig\|2591635.5.peg.1372 | CDS | CP041215.1 | 1301081 | 1301509 |
| | fig\|2591635.5.peg.1373 | CDS | CP041215.1 | 1301992 | 1301804 |
| | fig\|2591635.5.peg.1374 | CDS | CP041215.1 | 1302351 | 1302716 |
| | fig\|2591635.5.peg.1379 | CDS | CP041215.1 | 1304667 | 1304260 |
| | fig\|2591635.5.peg.1380 | CDS | CP041215.1 | 1304887 | 1304687 |
| | fig\|2591635.5.peg.1381 | CDS | CP041215.1 | 1305200 | 1304871 |
| | fig\|2591635.5.peg.1408 | CDS | CP041215.1 | 1338817 | 1338452 |
| | fig\|2591635.5.peg.1412 | CDS | CP041215.1 | 1340776 | 1340348 |
| | fig\|2591635.5.peg.1420 | CDS | CP041215.1 | 1346824 | 1346630 |
| | fig\|2591635.5.peg.1421 | CDS | CP041215.1 | 1347098 | 1346916 |
| | fig\|2591635.5.peg.1423 | CDS | CP041215.1 | 1347851 | 1348216 |
| | fig\|2591635.5.peg.1433 | CDS | CP041215.1 | 1353969 | 1354175 |
| | fig\|2591635.5.peg.1434 | CDS | CP041215.1 | 1354293 | 1354676 |
| | fig\|2591635.5.peg.1453 | CDS | CP041215.1 | 1378271 | 1377924 |
| | fig\|2591635.5.peg.1455 | CDS | CP041215.1 | 1380538 | 1380350 |
| | fig\|2591635.5.peg.1458 | CDS | CP041215.1 | 1381879 | 1382178 |
| | fig\|2591635.5.peg.1459 | CDS | CP041215.1 | 1382162 | 1382797 |
| | fig\|2591635.5.peg.1479 | CDS | CP041215.1 | 1400752 | 1400387 |
| | fig\|2591635.5.peg.1496 | CDS | CP041215.1 | 1418544 | 1417909 |
| | fig\|2591635.5.peg.1497 | CDS | CP041215.1 | 1418857 | 1418528 |
| | fig\|2591635.5.peg.1519 | CDS | CP041215.1 | 1440382 | 1440017 |
| wCfeJ | fig\|212123.3.peg.15 | CDS | CP051157.1 | 13901 | 13032 |
| | fig\|212123.3.peg.93 | CDS | CP051157.1 | 89982 | 90935 |
| | fig\|212123.3.peg.269 | CDS | CP051157.1 | 275016 | 275261 |
| | fig\|212123.3.peg.530 | CDS | CP051157.1 | 539929 | 540117 |
| | fig\|212123.3.peg.547 | CDS | CP051157.1 | 562196 | 562561 |
| | fig\|212123.3.peg.701 | CDS | CP051157.1 | 716149 | 716298 |
| | fig\|212123.3.peg.905 | CDS | CP051157.1 | 934347 | 934646 |
| | fig\|212123.3.peg.906 | CDS | CP051157.1 | 934840 | 935007 |
| | fig\|212123.3.peg.909 | CDS | CP051157.1 | 936536 | 937183 |
| | fig\|212123.3.peg.910 | CDS | CP051157.1 | 937180 | 937440 |
| | fig\|212123.3.peg.1152 | CDS | CP051157.1 | 1173814 | 1174230 |
| wCfeT | fig\|212123.4.peg.1 | CDS | CP051156.1 | 531 | 139 |
| | fig\|212123.4.peg.3 | CDS | CP051156.1 | 4905 | 3544 |
| | fig\|212123.4.peg.8 | CDS | CP051156.1 | 10203 | 9376 |
| | fig\|212123.4.peg.9 | CDS | CP051156.1 | 10557 | 10345 |
| | fig\|212123.4.peg.30 | CDS | CP051156.1 | 29990 | 29034 |

| | | | | |
|---|---|---|---|---|
| fig\|212123.4.peg.43 | CDS | CP051156.1 | 41140 | 41370 |
| fig\|212123.4.peg.48 | CDS | CP051156.1 | 47068 | 45728 |
| fig\|212123.4.peg.75 | CDS | CP051156.1 | 72746 | 71406 |
| fig\|212123.4.peg.79 | CDS | CP051156.1 | 75727 | 76098 |
| fig\|212123.4.peg.80 | CDS | CP051156.1 | 76240 | 77067 |
| fig\|212123.4.peg.93 | CDS | CP051156.1 | 91154 | 90327 |
| fig\|212123.4.peg.94 | CDS | CP051156.1 | 91667 | 91296 |
| fig\|212123.4.peg.108 | CDS | CP051156.1 | 104468 | 103599 |
| fig\|212123.4.peg.119 | CDS | CP051156.1 | 112144 | 111275 |
| fig\|212123.4.peg.130 | CDS | CP051156.1 | 119820 | 118951 |
| fig\|212123.4.peg.134 | CDS | CP051156.1 | 123480 | 124091 |
| fig\|212123.4.peg.135 | CDS | CP051156.1 | 125053 | 124097 |
| fig\|212123.4.peg.136 | CDS | CP051156.1 | 125690 | 125319 |
| fig\|212123.4.peg.178 | CDS | CP051156.1 | 154323 | 155684 |
| fig\|212123.4.peg.183 | CDS | CP051156.1 | 159234 | 160037 |
| fig\|212123.4.peg.186 | CDS | CP051156.1 | 160878 | 161048 |
| fig\|212123.4.peg.207 | CDS | CP051156.1 | 187656 | 188996 |
| fig\|212123.4.peg.209 | CDS | CP051156.1 | 191115 | 190159 |
| fig\|212123.4.peg.213 | CDS | CP051156.1 | 194651 | 195991 |
| fig\|212123.4.peg.215 | CDS | CP051156.1 | 198936 | 197596 |
| fig\|212123.4.peg.224 | CDS | CP051156.1 | 211293 | 209953 |
| fig\|212123.4.peg.225 | CDS | CP051156.1 | 212788 | 211427 |
| fig\|212123.4.peg.226 | CDS | CP051156.1 | 214417 | 213056 |
| fig\|212123.4.peg.235 | CDS | CP051156.1 | 227384 | 226428 |
| fig\|212123.4.peg.237 | CDS | CP051156.1 | 230591 | 231931 |
| fig\|212123.4.peg.239 | CDS | CP051156.1 | 233235 | 232309 |
| fig\|212123.4.peg.243 | CDS | CP051156.1 | 238649 | 237693 |
| fig\|212123.4.peg.245 | CDS | CP051156.1 | 242441 | 241101 |
| fig\|212123.4.peg.264 | CDS | CP051156.1 | 263363 | 264319 |
| fig\|212123.4.peg.289 | CDS | CP051156.1 | 286594 | 285638 |
| fig\|212123.4.peg.291 | CDS | CP051156.1 | 287718 | 288089 |
| fig\|212123.4.peg.292 | CDS | CP051156.1 | 288231 | 289058 |
| fig\|212123.4.peg.296 | CDS | CP051156.1 | 291867 | 293207 |
| fig\|212123.4.peg.299 | CDS | CP051156.1 | 295758 | 297098 |
| fig\|212123.4.peg.321 | CDS | CP051156.1 | 321273 | 320317 |
| fig\|212123.4.peg.324 | CDS | CP051156.1 | 324435 | 323479 |
| fig\|212123.4.peg.327 | CDS | CP051156.1 | 327731 | 326391 |
| fig\|212123.4.peg.337 | CDS | CP051156.1 | 337975 | 337817 |
| fig\|212123.4.peg.338 | CDS | CP051156.1 | 338067 | 338438 |
| fig\|212123.4.peg.341 | CDS | CP051156.1 | 339510 | 340265 |
| fig\|212123.4.peg.373 | CDS | CP051156.1 | 370736 | 369375 |
| fig\|212123.4.peg.377 | CDS | CP051156.1 | 375660 | 376616 |

| | | | | |
|---|---|---|---|---|
| fig\|212123.4.peg.379 | CDS | CP051156.1 | 377056 | 378396 |
| fig\|212123.4.peg.381 | CDS | CP051156.1 | 378909 | 379865 |
| fig\|212123.4.peg.383 | CDS | CP051156.1 | 382478 | 381423 |
| fig\|212123.4.peg.384 | CDS | CP051156.1 | 383869 | 382529 |
| fig\|212123.4.peg.400 | CDS | CP051156.1 | 400810 | 400007 |
| fig\|212123.4.peg.401 | CDS | CP051156.1 | 401323 | 400952 |
| fig\|212123.4.peg.420 | CDS | CP051156.1 | 417498 | 418838 |
| fig\|212123.4.peg.423 | CDS | CP051156.1 | 420909 | 420709 |
| fig\|212123.4.peg.430 | CDS | CP051156.1 | 430156 | 428795 |
| fig\|212123.4.peg.435 | CDS | CP051156.1 | 435523 | 434609 |
| fig\|212123.4.peg.438 | CDS | CP051156.1 | 436589 | 437929 |
| fig\|212123.4.peg.442 | CDS | CP051156.1 | 441077 | 439737 |
| fig\|212123.4.peg.443 | CDS | CP051156.1 | 441285 | 442241 |
| fig\|212123.4.peg.456 | CDS | CP051156.1 | 455453 | 456409 |
| fig\|212123.4.peg.461 | CDS | CP051156.1 | 463678 | 462338 |
| fig\|212123.4.peg.464 | CDS | CP051156.1 | 467059 | 465743 |
| fig\|212123.4.peg.479 | CDS | CP051156.1 | 480399 | 479572 |
| fig\|212123.4.peg.480 | CDS | CP051156.1 | 480912 | 480541 |
| fig\|212123.4.peg.483 | CDS | CP051156.1 | 485154 | 484198 |
| fig\|212123.4.peg.487 | CDS | CP051156.1 | 490628 | 489672 |
| fig\|212123.4.peg.495 | CDS | CP051156.1 | 494934 | 496274 |
| fig\|212123.4.peg.506 | CDS | CP051156.1 | 505358 | 504684 |
| fig\|212123.4.peg.507 | CDS | CP051156.1 | 505871 | 505500 |
| fig\|212123.4.peg.530 | CDS | CP051156.1 | 533698 | 532742 |
| fig\|212123.4.peg.537 | CDS | CP051156.1 | 539018 | 538191 |
| fig\|212123.4.peg.538 | CDS | CP051156.1 | 539531 | 539160 |
| fig\|212123.4.peg.543 | CDS | CP051156.1 | 542537 | 543877 |
| fig\|212123.4.peg.548 | CDS | CP051156.1 | 548794 | 547607 |
| fig\|212123.4.peg.551 | CDS | CP051156.1 | 550679 | 549723 |
| fig\|212123.4.peg.556 | CDS | CP051156.1 | 556041 | 556997 |
| fig\|212123.4.peg.557 | CDS | CP051156.1 | 558424 | 557084 |
| fig\|212123.4.peg.571 | CDS | CP051156.1 | 572896 | 572165 |
| fig\|212123.4.peg.572 | CDS | CP051156.1 | 573409 | 573038 |
| fig\|212123.4.peg.580 | CDS | CP051156.1 | 581602 | 582942 |
| fig\|212123.4.peg.592 | CDS | CP051156.1 | 595051 | 595422 |
| fig\|212123.4.peg.593 | CDS | CP051156.1 | 595564 | 596391 |
| fig\|212123.4.peg.594 | CDS | CP051156.1 | 596551 | 596378 |
| fig\|212123.4.peg.599 | CDS | CP051156.1 | 599055 | 599426 |
| fig\|212123.4.peg.600 | CDS | CP051156.1 | 599568 | 600395 |
| fig\|212123.4.peg.604 | CDS | CP051156.1 | 603636 | 602680 |
| fig\|212123.4.peg.630 | CDS | CP051156.1 | 628914 | 630254 |
| fig\|212123.4.peg.631 | CDS | CP051156.1 | 631507 | 630251 |

| | | | | |
|---|---|---|---|---|
| fig\|212123.4.peg.637 | CDS | CP051156.1 | 639236 | 638052 |
| fig\|212123.4.peg.644 | CDS | CP051156.1 | 646539 | 645583 |
| fig\|212123.4.peg.647 | CDS | CP051156.1 | 647873 | 648460 |
| fig\|212123.4.peg.687 | CDS | CP051156.1 | 688045 | 687089 |
| fig\|212123.4.peg.690 | CDS | CP051156.1 | 692253 | 690913 |
| fig\|212123.4.peg.697 | CDS | CP051156.1 | 700240 | 698879 |
| fig\|212123.4.peg.698 | CDS | CP051156.1 | 701545 | 700589 |
| fig\|212123.4.peg.700 | CDS | CP051156.1 | 702474 | 703814 |
| fig\|212123.4.peg.706 | CDS | CP051156.1 | 708547 | 709503 |
| fig\|212123.4.peg.709 | CDS | CP051156.1 | 711654 | 712610 |
| fig\|212123.4.peg.714 | CDS | CP051156.1 | 716733 | 715393 |
| fig\|212123.4.peg.737 | CDS | CP051156.1 | 743664 | 744023 |
| fig\|212123.4.peg.738 | CDS | CP051156.1 | 744165 | 744992 |
| fig\|212123.4.peg.751 | CDS | CP051156.1 | 762080 | 762451 |
| fig\|212123.4.peg.752 | CDS | CP051156.1 | 762593 | 763420 |
| fig\|212123.4.peg.758 | CDS | CP051156.1 | 767626 | 768966 |
| fig\|212123.4.peg.762 | CDS | CP051156.1 | 773298 | 773576 |
| fig\|212123.4.peg.763 | CDS | CP051156.1 | 774538 | 773582 |
| fig\|212123.4.peg.787 | CDS | CP051156.1 | 795485 | 796123 |
| fig\|212123.4.peg.793 | CDS | CP051156.1 | 801938 | 800598 |
| fig\|212123.4.peg.799 | CDS | CP051156.1 | 809694 | 808738 |
| fig\|212123.4.peg.800 | CDS | CP051156.1 | 809735 | 810556 |
| fig\|212123.4.peg.804 | CDS | CP051156.1 | 813372 | 813743 |
| fig\|212123.4.peg.806 | CDS | CP051156.1 | 814126 | 814713 |
| fig\|212123.4.peg.825 | CDS | CP051156.1 | 831373 | 832329 |
| fig\|212123.4.peg.827 | CDS | CP051156.1 | 832769 | 834109 |
| fig\|212123.4.peg.835 | CDS | CP051156.1 | 840507 | 841463 |
| fig\|212123.4.peg.837 | CDS | CP051156.1 | 841717 | 842304 |
| fig\|212123.4.peg.838 | CDS | CP051156.1 | 843225 | 842284 |
| fig\|212123.4.peg.839 | CDS | CP051156.1 | 843402 | 843632 |
| fig\|212123.4.peg.842 | CDS | CP051156.1 | 847093 | 845753 |
| fig\|212123.4.peg.848 | CDS | CP051156.1 | 853107 | 851767 |
| fig\|212123.4.peg.856 | CDS | CP051156.1 | 860154 | 859198 |
| fig\|212123.4.peg.861 | CDS | CP051156.1 | 866387 | 867343 |
| fig\|212123.4.peg.870 | CDS | CP051156.1 | 879404 | 878091 |
| fig\|212123.4.peg.879 | CDS | CP051156.1 | 887615 | 886659 |
| fig\|212123.4.peg.880 | CDS | CP051156.1 | 888082 | 889422 |
| fig\|212123.4.peg.890 | CDS | CP051156.1 | 898806 | 899177 |
| fig\|212123.4.peg.898 | CDS | CP051156.1 | 906081 | 905125 |
| fig\|212123.4.peg.906 | CDS | CP051156.1 | 915870 | 914530 |
| fig\|212123.4.peg.908 | CDS | CP051156.1 | 917266 | 916310 |
| fig\|212123.4.peg.922 | CDS | CP051156.1 | 930706 | 929366 |

| | | | | |
|---|---|---|---|---|
| fig\|212123.4.peg.925 | CDS | CP051156.1 | 932288 | 933628 |
| fig\|212123.4.peg.940 | CDS | CP051156.1 | 945097 | 944510 |
| fig\|212123.4.peg.947 | CDS | CP051156.1 | 950143 | 949187 |
| fig\|212123.4.peg.964 | CDS | CP051156.1 | 967780 | 968736 |
| fig\|212123.4.peg.968 | CDS | CP051156.1 | 970958 | 972298 |
| fig\|212123.4.peg.990 | CDS | CP051156.1 | 993516 | 992560 |
| fig\|212123.4.peg.997 | CDS | CP051156.1 | 999100 | 1000440 |
| fig\|212123.4.peg.998 | CDS | CP051156.1 | 1000539 | 1001879 |
| fig\|212123.4.peg.1004 | CDS | CP051156.1 | 1007115 | 1006159 |
| fig\|212123.4.peg.1052 | CDS | CP051156.1 | 1059547 | 1058591 |
| fig\|212123.4.peg.1056 | CDS | CP051156.1 | 1062311 | 1062682 |
| fig\|212123.4.peg.1057 | CDS | CP051156.1 | 1062824 | 1063651 |
| fig\|212123.4.peg.1077 | CDS | CP051156.1 | 1083495 | 1082539 |
| fig\|212123.4.peg.1080 | CDS | CP051156.1 | 1087812 | 1088042 |
| fig\|212123.4.peg.1082 | CDS | CP051156.1 | 1089376 | 1088420 |
| fig\|212123.4.peg.1102 | CDS | CP051156.1 | 1119077 | 1117737 |
| fig\|212123.4.peg.1126 | CDS | CP051156.1 | 1141921 | 1142658 |
| fig\|212123.4.peg.1127 | CDS | CP051156.1 | 1142625 | 1142876 |
| fig\|212123.4.peg.1135 | CDS | CP051156.1 | 1151598 | 1150642 |
| fig\|212123.4.peg.1140 | CDS | CP051156.1 | 1155674 | 1157014 |
| fig\|212123.4.peg.1146 | CDS | CP051156.1 | 1160219 | 1161256 |
| fig\|212123.4.peg.1147 | CDS | CP051156.1 | 1161329 | 1161559 |
| fig\|212123.4.peg.1166 | CDS | CP051156.1 | 1180793 | 1180563 |
| fig\|212123.4.peg.1192 | CDS | CP051156.1 | 1201427 | 1201657 |
| fig\|212123.4.peg.1233 | CDS | CP051156.1 | 1234458 | 1233118 |
| fig\|212123.4.peg.1237 | CDS | CP051156.1 | 1239445 | 1240785 |
| fig\|212123.4.peg.1248 | CDS | CP051156.1 | 1249535 | 1250875 |
| fig\|212123.4.peg.1254 | CDS | CP051156.1 | 1257621 | 1256281 |
| fig\|212123.4.peg.1271 | CDS | CP051156.1 | 1276151 | 1275195 |
| fig\|212123.4.peg.1272 | CDS | CP051156.1 | 1276172 | 1276477 |
| fig\|212123.4.peg.1274 | CDS | CP051156.1 | 1276808 | 1277446 |
| fig\|212123.4.peg.1280 | CDS | CP051156.1 | 1284273 | 1285613 |
| fig\|212123.4.peg.1284 | CDS | CP051156.1 | 1289018 | 1288146 |
| fig\|212123.4.peg.1289 | CDS | CP051156.1 | 1295154 | 1294198 |
| fig\|212123.4.peg.1293 | CDS | CP051156.1 | 1296985 | 1296347 |
| fig\|212123.4.peg.1294 | CDS | CP051156.1 | 1297687 | 1296989 |
| fig\|212123.4.peg.1302 | CDS | CP051156.1 | 1304851 | 1304213 |
| fig\|212123.4.peg.1307 | CDS | CP051156.1 | 1309594 | 1308767 |
| fig\|212123.4.peg.1308 | CDS | CP051156.1 | 1310041 | 1309736 |
| fig\|212123.4.peg.1309 | CDS | CP051156.1 | 1310062 | 1311018 |
| fig\|212123.4.peg.1315 | CDS | CP051156.1 | 1315810 | 1316181 |
| fig\|212123.4.peg.1316 | CDS | CP051156.1 | 1316323 | 1317150 |

| | | | | | |
|---|---|---|---|---|---|
| | fig\|212123.4.peg.1327 | CDS | CP051156.1 | 1326222 | 1325395 |
| | fig\|212123.4.peg.1328 | CDS | CP051156.1 | 1326576 | 1326364 |
| | fig\|212123.4.peg.1334 | CDS | CP051156.1 | 1331433 | 1330795 |
| | fig\|212123.4.peg.1336 | CDS | CP051156.1 | 1332102 | 1331764 |
| | fig\|212123.4.peg.1344 | CDS | CP051156.1 | 1341193 | 1340366 |
| | fig\|212123.4.peg.1345 | CDS | CP051156.1 | 1341706 | 1341335 |
| | fig\|212123.4.peg.1353 | CDS | CP051156.1 | 1347441 | 1346614 |
| | fig\|212123.4.peg.1354 | CDS | CP051156.1 | 1347934 | 1347590 |
| | fig\|212123.4.peg.1357 | CDS | CP051156.1 | 1348703 | 1350043 |
| | fig\|212123.4.peg.1366 | CDS | CP051156.1 | 1358646 | 1357819 |
| | fig\|212123.4.peg.1367 | CDS | CP051156.1 | 1359159 | 1358788 |
| | fig\|212123.4.peg.1397 | CDS | CP051156.1 | 1383513 | 1382686 |
| | fig\|212123.4.peg.1398 | CDS | CP051156.1 | 1384026 | 1383655 |
| | fig\|212123.4.peg.1404 | CDS | CP051156.1 | 1389956 | 1389207 |
| | fig\|212123.4.peg.1461 | CDS | CP051156.1 | 1441907 | 1443268 |
| | fig\|212123.4.peg.1462 | CDS | CP051156.1 | 1443346 | 1444692 |
| | fig\|212123.4.peg.1469 | CDS | CP051156.1 | 1449907 | 1450863 |
| | fig\|212123.4.peg.1474 | CDS | CP051156.1 | 1455126 | 1455497 |
| | fig\|212123.4.peg.1475 | CDS | CP051156.1 | 1455639 | 1456466 |
| | fig\|212123.4.peg.1489 | CDS | CP051156.1 | 1473809 | 1474765 |
| | fig\|212123.4.peg.1495 | CDS | CP051156.1 | 1481119 | 1479779 |
| | fig\|212123.4.peg.1502 | CDS | CP051156.1 | 1489851 | 1488895 |
| | fig\|212123.4.peg.1508 | CDS | CP051156.1 | 1494125 | 1494721 |
| | fig\|212123.4.peg.1509 | CDS | CP051156.1 | 1495538 | 1494708 |
| wCle | fig\|246273.9.peg.13 | CDS | wCle_NZ_AP0130 | 13005 | 13319 |
| | fig\|246273.9.peg.14 | CDS | wCle_NZ_AP0130 | 13438 | 13794 |
| | fig\|246273.9.peg.21 | CDS | wCle_NZ_AP0130 | 26655 | 26299 |
| | fig\|246273.9.peg.22 | CDS | wCle_NZ_AP0130 | 27088 | 26774 |
| | fig\|246273.9.peg.50 | CDS | wCle_NZ_AP0130 | 51176 | 50754 |
| | fig\|246273.9.peg.51 | CDS | wCle_NZ_AP0130 | 51307 | 51621 |
| | fig\|246273.9.peg.52 | CDS | wCle_NZ_AP0130 | 51633 | 52097 |
| | fig\|246273.9.peg.53 | CDS | wCle_NZ_AP0130 | 52366 | 52094 |
| | fig\|246273.9.peg.76 | CDS | wCle_NZ_AP0130 | 78119 | 77655 |
| | fig\|246273.9.peg.77 | CDS | wCle_NZ_AP0130 | 78445 | 78131 |
| | fig\|246273.9.peg.109 | CDS | wCle_NZ_AP0130 | 105865 | 106329 |
| | fig\|246273.9.peg.131 | CDS | wCle_NZ_AP0130 | 127772 | 128086 |
| | fig\|246273.9.peg.132 | CDS | wCle_NZ_AP0130 | 128205 | 128612 |
| | fig\|246273.9.peg.145 | CDS | wCle_NZ_AP0130 | 137718 | 138074 |
| | fig\|246273.9.peg.147 | CDS | wCle_NZ_AP0130 | 139024 | 139338 |
| | fig\|246273.9.peg.148 | CDS | wCle_NZ_AP0130 | 139457 | 139813 |
| | fig\|246273.9.peg.156 | CDS | wCle_NZ_AP0130 | 149781 | 150095 |
| | fig\|246273.9.peg.157 | CDS | wCle_NZ_AP0130 | 150214 | 150570 |

| | | | | |
|---|---|---|---|---|
| fig\|246273.9.peg.166 | CDS | wCle_NZ_AP0130 | 157008 | 157364 |
| fig\|246273.9.peg.185 | CDS | wCle_NZ_AP0130 | 178293 | 177829 |
| fig\|246273.9.peg.186 | CDS | wCle_NZ_AP0130 | 178619 | 178305 |
| fig\|246273.9.peg.194 | CDS | wCle_NZ_AP0130 | 186666 | 186980 |
| fig\|246273.9.peg.195 | CDS | wCle_NZ_AP0130 | 186992 | 187456 |
| fig\|246273.9.peg.199 | CDS | wCle_NZ_AP0130 | 189861 | 189397 |
| fig\|246273.9.peg.200 | CDS | wCle_NZ_AP0130 | 190187 | 189873 |
| fig\|246273.9.peg.204 | CDS | wCle_NZ_AP0130 | 192383 | 192697 |
| fig\|246273.9.peg.205 | CDS | wCle_NZ_AP0130 | 192816 | 193172 |
| fig\|246273.9.peg.219 | CDS | wCle_NZ_AP0130 | 205679 | 205218 |
| fig\|246273.9.peg.220 | CDS | wCle_NZ_AP0130 | 206005 | 205691 |
| fig\|246273.9.peg.262 | CDS | wCle_NZ_AP0130 | 233485 | 233799 |
| fig\|246273.9.peg.263 | CDS | wCle_NZ_AP0130 | 233811 | 234173 |
| fig\|246273.9.peg.267 | CDS | wCle_NZ_AP0130 | 236273 | 235917 |
| fig\|246273.9.peg.268 | CDS | wCle_NZ_AP0130 | 236706 | 236392 |
| fig\|246273.9.peg.272 | CDS | wCle_NZ_AP0130 | 237804 | 238118 |
| fig\|246273.9.peg.273 | CDS | wCle_NZ_AP0130 | 238237 | 238593 |
| fig\|246273.9.peg.274 | CDS | wCle_NZ_AP0130 | 239251 | 238895 |
| fig\|246273.9.peg.275 | CDS | wCle_NZ_AP0130 | 239684 | 239370 |
| fig\|246273.9.peg.293 | CDS | wCle_NZ_AP0130 | 258275 | 257919 |
| fig\|246273.9.peg.294 | CDS | wCle_NZ_AP0130 | 258708 | 258394 |
| fig\|246273.9.peg.301 | CDS | wCle_NZ_AP0130 | 265341 | 264985 |
| fig\|246273.9.peg.302 | CDS | wCle_NZ_AP0130 | 265774 | 265460 |
| fig\|246273.9.peg.319 | CDS | wCle_NZ_AP0130 | 282111 | 281755 |
| fig\|246273.9.peg.320 | CDS | wCle_NZ_AP0130 | 282544 | 282335 |
| fig\|246273.9.peg.323 | CDS | wCle_NZ_AP0130 | 285184 | 284828 |
| fig\|246273.9.peg.324 | CDS | wCle_NZ_AP0130 | 285619 | 285305 |
| fig\|246273.9.peg.331 | CDS | wCle_NZ_AP0130 | 292811 | 292347 |
| fig\|246273.9.peg.332 | CDS | wCle_NZ_AP0130 | 292942 | 293256 |
| fig\|246273.9.peg.333 | CDS | wCle_NZ_AP0130 | 293375 | 293731 |
| fig\|246273.9.peg.334 | CDS | wCle_NZ_AP0130 | 294000 | 293728 |
| fig\|246273.9.peg.343 | CDS | wCle_NZ_AP0130 | 300242 | 299886 |
| fig\|246273.9.peg.344 | CDS | wCle_NZ_AP0130 | 300675 | 300361 |
| fig\|246273.9.peg.349 | CDS | wCle_NZ_AP0130 | 306943 | 307152 |
| fig\|246273.9.peg.350 | CDS | wCle_NZ_AP0130 | 307269 | 307733 |
| fig\|246273.9.peg.378 | CDS | wCle_NZ_AP0130 | 338132 | 337776 |
| fig\|246273.9.peg.379 | CDS | wCle_NZ_AP0130 | 338565 | 338251 |
| fig\|246273.9.peg.389 | CDS | wCle_NZ_AP0130 | 350581 | 350895 |
| fig\|246273.9.peg.390 | CDS | wCle_NZ_AP0130 | 351016 | 351372 |
| fig\|246273.9.peg.394 | CDS | wCle_NZ_AP0130 | 355568 | 355212 |
| fig\|246273.9.peg.395 | CDS | wCle_NZ_AP0130 | 356001 | 355687 |
| fig\|246273.9.peg.397 | CDS | wCle_NZ_AP0130 | 356563 | 356877 |

| | | | | |
|---|---|---|---|---|
| fig\|246273.9.peg.398 | CDS | wCle_NZ_AP0130 | 356996 | 357352 |
| fig\|246273.9.peg.400 | CDS | wCle_NZ_AP0130 | 360282 | 359926 |
| fig\|246273.9.peg.401 | CDS | wCle_NZ_AP0130 | 360715 | 360401 |
| fig\|246273.9.peg.409 | CDS | wCle_NZ_AP0130 | 370459 | 369995 |
| fig\|246273.9.peg.410 | CDS | wCle_NZ_AP0130 | 370785 | 370576 |
| fig\|246273.9.peg.430 | CDS | wCle_NZ_AP0130 | 395841 | 396113 |
| fig\|246273.9.peg.431 | CDS | wCle_NZ_AP0130 | 396253 | 396110 |
| fig\|246273.9.peg.432 | CDS | wCle_NZ_AP0130 | 396573 | 396211 |
| fig\|246273.9.peg.433 | CDS | wCle_NZ_AP0130 | 396899 | 396585 |
| fig\|246273.9.peg.434 | CDS | wCle_NZ_AP0130 | 397137 | 397493 |
| fig\|246273.9.peg.435 | CDS | wCle_NZ_AP0130 | 397933 | 398304 |
| fig\|246273.9.peg.436 | CDS | wCle_NZ_AP0130 | 398320 | 398802 |
| fig\|246273.9.peg.444 | CDS | wCle_NZ_AP0130 | 406327 | 405845 |
| fig\|246273.9.peg.445 | CDS | wCle_NZ_AP0130 | 406714 | 406343 |
| fig\|246273.9.peg.448 | CDS | wCle_NZ_AP0130 | 407857 | 408171 |
| fig\|246273.9.peg.449 | CDS | wCle_NZ_AP0130 | 408290 | 408646 |
| fig\|246273.9.peg.459 | CDS | wCle_NZ_AP0130 | 416874 | 417188 |
| fig\|246273.9.peg.460 | CDS | wCle_NZ_AP0130 | 417307 | 417663 |
| fig\|246273.9.peg.471 | CDS | wCle_NZ_AP0130 | 430526 | 430882 |
| fig\|246273.9.peg.480 | CDS | wCle_NZ_AP0130 | 439007 | 439183 |
| fig\|246273.9.peg.482 | CDS | wCle_NZ_AP0130 | 440419 | 439955 |
| fig\|246273.9.peg.483 | CDS | wCle_NZ_AP0130 | 440745 | 440431 |
| fig\|246273.9.peg.486 | CDS | wCle_NZ_AP0130 | 443059 | 443373 |
| fig\|246273.9.peg.487 | CDS | wCle_NZ_AP0130 | 443385 | 443849 |
| fig\|246273.9.peg.490 | CDS | wCle_NZ_AP0130 | 445312 | 445626 |
| fig\|246273.9.peg.491 | CDS | wCle_NZ_AP0130 | 445638 | 446012 |
| fig\|246273.9.peg.492 | CDS | wCle_NZ_AP0130 | 446473 | 446009 |
| fig\|246273.9.peg.493 | CDS | wCle_NZ_AP0130 | 446799 | 446485 |
| fig\|246273.9.peg.505 | CDS | wCle_NZ_AP0130 | 452208 | 452522 |
| fig\|246273.9.peg.506 | CDS | wCle_NZ_AP0130 | 452643 | 452999 |
| fig\|246273.9.peg.543 | CDS | wCle_NZ_AP0130 | 488634 | 488170 |
| fig\|246273.9.peg.544 | CDS | wCle_NZ_AP0130 | 488960 | 488646 |
| fig\|246273.9.peg.546 | CDS | wCle_NZ_AP0130 | 489839 | 489696 |
| fig\|246273.9.peg.547 | CDS | wCle_NZ_AP0130 | 490159 | 489797 |
| fig\|246273.9.peg.548 | CDS | wCle_NZ_AP0130 | 490485 | 490171 |
| fig\|246273.9.peg.553 | CDS | wCle_NZ_AP0130 | 494223 | 493570 |
| fig\|246273.9.peg.554 | CDS | wCle_NZ_AP0130 | 494584 | 494255 |
| fig\|246273.9.peg.555 | CDS | wCle_NZ_AP0130 | 494822 | 495178 |
| fig\|246273.9.peg.576 | CDS | wCle_NZ_AP0130 | 510286 | 510600 |
| fig\|246273.9.peg.577 | CDS | wCle_NZ_AP0130 | 510719 | 511075 |
| fig\|246273.9.peg.592 | CDS | wCle_NZ_AP0130 | 526798 | 527112 |
| fig\|246273.9.peg.593 | CDS | wCle_NZ_AP0130 | 527231 | 527587 |

| | | | | |
|---|---|---|---|---|
| fig\|246273.9.peg.615 | CDS | wCle_NZ_AP0130 | 545423 | 545067 |
| fig\|246273.9.peg.616 | CDS | wCle_NZ_AP0130 | 545856 | 545542 |
| fig\|246273.9.peg.625 | CDS | wCle_NZ_AP0130 | 554222 | 553866 |
| fig\|246273.9.peg.626 | CDS | wCle_NZ_AP0130 | 554655 | 554341 |
| fig\|246273.9.peg.635 | CDS | wCle_NZ_AP0130 | 561106 | 561420 |
| fig\|246273.9.peg.636 | CDS | wCle_NZ_AP0130 | 561432 | 561896 |
| fig\|246273.9.peg.650 | CDS | wCle_NZ_AP0130 | 574601 | 574915 |
| fig\|246273.9.peg.651 | CDS | wCle_NZ_AP0130 | 575034 | 575390 |
| fig\|246273.9.peg.652 | CDS | wCle_NZ_AP0130 | 575659 | 575387 |
| fig\|246273.9.peg.661 | CDS | wCle_NZ_AP0130 | 583494 | 583030 |
| fig\|246273.9.peg.662 | CDS | wCle_NZ_AP0130 | 583820 | 583506 |
| fig\|246273.9.peg.664 | CDS | wCle_NZ_AP0130 | 584419 | 584733 |
| fig\|246273.9.peg.665 | CDS | wCle_NZ_AP0130 | 584852 | 585208 |
| fig\|246273.9.peg.699 | CDS | wCle_NZ_AP0130 | 622093 | 622407 |
| fig\|246273.9.peg.700 | CDS | wCle_NZ_AP0130 | 622419 | 622883 |
| fig\|246273.9.peg.707 | CDS | wCle_NZ_AP0130 | 627489 | 627133 |
| fig\|246273.9.peg.708 | CDS | wCle_NZ_AP0130 | 627922 | 627608 |
| fig\|246273.9.peg.717 | CDS | wCle_NZ_AP0130 | 637381 | 637695 |
| fig\|246273.9.peg.718 | CDS | wCle_NZ_AP0130 | 637814 | 638170 |
| fig\|246273.9.peg.746 | CDS | wCle_NZ_AP0130 | 662977 | 662513 |
| fig\|246273.9.peg.747 | CDS | wCle_NZ_AP0130 | 663303 | 662989 |
| fig\|246273.9.peg.765 | CDS | wCle_NZ_AP0130 | 680675 | 680319 |
| fig\|246273.9.peg.766 | CDS | wCle_NZ_AP0130 | 681110 | 680796 |
| fig\|246273.9.peg.774 | CDS | wCle_NZ_AP0130 | 684882 | 685196 |
| fig\|246273.9.peg.775 | CDS | wCle_NZ_AP0130 | 685208 | 685672 |
| fig\|246273.9.peg.777 | CDS | wCle_NZ_AP0130 | 687491 | 687135 |
| fig\|246273.9.peg.778 | CDS | wCle_NZ_AP0130 | 687926 | 687612 |
| fig\|246273.9.peg.783 | CDS | wCle_NZ_AP0130 | 692034 | 692348 |
| fig\|246273.9.peg.784 | CDS | wCle_NZ_AP0130 | 692467 | 692823 |
| fig\|246273.9.peg.791 | CDS | wCle_NZ_AP0130 | 695944 | 696258 |
| fig\|246273.9.peg.792 | CDS | wCle_NZ_AP0130 | 696270 | 696734 |
| fig\|246273.9.peg.815 | CDS | wCle_NZ_AP0130 | 718850 | 718386 |
| fig\|246273.9.peg.816 | CDS | wCle_NZ_AP0130 | 719176 | 718862 |
| fig\|246273.9.peg.833 | CDS | wCle_NZ_AP0130 | 738398 | 738670 |
| fig\|246273.9.peg.834 | CDS | wCle_NZ_AP0130 | 739023 | 738667 |
| fig\|246273.9.peg.835 | CDS | wCle_NZ_AP0130 | 739456 | 739142 |
| fig\|246273.9.peg.859 | CDS | wCle_NZ_AP0130 | 768349 | 768098 |
| fig\|246273.9.peg.860 | CDS | wCle_NZ_AP0130 | 768782 | 768468 |
| fig\|246273.9.peg.868 | CDS | wCle_NZ_AP0130 | 775257 | 774793 |
| fig\|246273.9.peg.869 | CDS | wCle_NZ_AP0130 | 775583 | 775269 |
| fig\|246273.9.peg.872 | CDS | wCle_NZ_AP0130 | 779276 | 778920 |
| fig\|246273.9.peg.873 | CDS | wCle_NZ_AP0130 | 779709 | 779395 |

| | | | | |
|---|---|---|---|---|
| fig\|246273.9.peg.890 | CDS | wCle_NZ_AP0130 | 794064 | 794378 |
| fig\|246273.9.peg.891 | CDS | wCle_NZ_AP0130 | 794390 | 794752 |
| fig\|246273.9.peg.892 | CDS | wCle_NZ_AP0130 | 794710 | 794853 |
| fig\|246273.9.peg.895 | CDS | wCle_NZ_AP0130 | 797258 | 796794 |
| fig\|246273.9.peg.896 | CDS | wCle_NZ_AP0130 | 797584 | 797270 |
| fig\|246273.9.peg.900 | CDS | wCle_NZ_AP0130 | 801633 | 801169 |
| fig\|246273.9.peg.901 | CDS | wCle_NZ_AP0130 | 801959 | 801645 |
| fig\|246273.9.peg.906 | CDS | wCle_NZ_AP0130 | 804943 | 805257 |
| fig\|246273.9.peg.907 | CDS | wCle_NZ_AP0130 | 805269 | 805733 |
| fig\|246273.9.peg.913 | CDS | wCle_NZ_AP0130 | 810737 | 810273 |
| fig\|246273.9.peg.914 | CDS | wCle_NZ_AP0130 | 811063 | 810749 |
| fig\|246273.9.peg.925 | CDS | wCle_NZ_AP0130 | 818691 | 819005 |
| fig\|246273.9.peg.926 | CDS | wCle_NZ_AP0130 | 819017 | 819481 |
| fig\|246273.9.peg.929 | CDS | wCle_NZ_AP0130 | 822089 | 822403 |
| fig\|246273.9.peg.930 | CDS | wCle_NZ_AP0130 | 822522 | 822878 |
| fig\|246273.9.peg.934 | CDS | wCle_NZ_AP0130 | 825930 | 825616 |
| fig\|246273.9.peg.935 | CDS | wCle_NZ_AP0130 | 826363 | 826049 |
| fig\|246273.9.peg.939 | CDS | wCle_NZ_AP0130 | 831099 | 831368 |
| fig\|246273.9.peg.946 | CDS | wCle_NZ_AP0130 | 835948 | 835592 |
| fig\|246273.9.peg.947 | CDS | wCle_NZ_AP0130 | 836381 | 836067 |
| fig\|246273.9.peg.961 | CDS | wCle_NZ_AP0130 | 848608 | 848252 |
| fig\|246273.9.peg.962 | CDS | wCle_NZ_AP0130 | 849041 | 848727 |
| fig\|246273.9.peg.966 | CDS | wCle_NZ_AP0130 | 853742 | 853386 |
| fig\|246273.9.peg.967 | CDS | wCle_NZ_AP0130 | 854175 | 853861 |
| fig\|246273.9.peg.975 | CDS | wCle_NZ_AP0130 | 857947 | 858261 |
| fig\|246273.9.peg.976 | CDS | wCle_NZ_AP0130 | 858380 | 858736 |
| fig\|246273.9.peg.981 | CDS | wCle_NZ_AP0130 | 862932 | 863246 |
| fig\|246273.9.peg.982 | CDS | wCle_NZ_AP0130 | 863258 | 863620 |
| fig\|246273.9.peg.983 | CDS | wCle_NZ_AP0130 | 863578 | 863721 |
| fig\|246273.9.peg.1012 | CDS | wCle_NZ_AP0130 | 893235 | 893549 |
| fig\|246273.9.peg.1013 | CDS | wCle_NZ_AP0130 | 893561 | 893923 |
| fig\|246273.9.peg.1014 | CDS | wCle_NZ_AP0130 | 893881 | 894024 |
| fig\|246273.9.peg.1019 | CDS | wCle_NZ_AP0130 | 895780 | 896094 |
| fig\|246273.9.peg.1020 | CDS | wCle_NZ_AP0130 | 896215 | 896571 |
| fig\|246273.9.peg.1043 | CDS | wCle_NZ_AP0130 | 920622 | 920266 |
| fig\|246273.9.peg.1044 | CDS | wCle_NZ_AP0130 | 921055 | 920741 |
| fig\|246273.9.peg.1051 | CDS | wCle_NZ_AP0130 | 925594 | 925908 |
| fig\|246273.9.peg.1052 | CDS | wCle_NZ_AP0130 | 926027 | 926341 |
| fig\|246273.9.peg.1053 | CDS | wCle_NZ_AP0130 | 926651 | 926379 |
| fig\|246273.9.peg.1075 | CDS | wCle_NZ_AP0130 | 943312 | 943626 |
| fig\|246273.9.peg.1076 | CDS | wCle_NZ_AP0130 | 943745 | 944101 |
| fig\|246273.9.peg.1084 | CDS | wCle_NZ_AP0130 | 950402 | 950716 |

| | | | | |
|---|---|---|---|---|
| fig\|246273.9.peg.1085 | CDS | wCle_NZ_AP0130 | 950728 | 951192 |
| fig\|246273.9.peg.1090 | CDS | wCle_NZ_AP0130 | 954264 | 953800 |
| fig\|246273.9.peg.1091 | CDS | wCle_NZ_AP0130 | 954590 | 954276 |
| fig\|246273.9.peg.1110 | CDS | wCle_NZ_AP0130 | 970103 | 969639 |
| fig\|246273.9.peg.1111 | CDS | wCle_NZ_AP0130 | 970429 | 970115 |
| fig\|246273.9.peg.1112 | CDS | wCle_NZ_AP0130 | 971044 | 970580 |
| fig\|246273.9.peg.1113 | CDS | wCle_NZ_AP0130 | 971370 | 971056 |
| fig\|246273.9.peg.1119 | CDS | wCle_NZ_AP0130 | 974882 | 975196 |
| fig\|246273.9.peg.1120 | CDS | wCle_NZ_AP0130 | 975208 | 975570 |
| fig\|246273.9.peg.1164 | CDS | wCle_NZ_AP0130 | 1027754 | 1028068 |
| fig\|246273.9.peg.1165 | CDS | wCle_NZ_AP0130 | 1028080 | 1028544 |
| fig\|246273.9.peg.1174 | CDS | wCle_NZ_AP0130 | 1036077 | 1036391 |
| fig\|246273.9.peg.1175 | CDS | wCle_NZ_AP0130 | 1036510 | 1036890 |
| fig\|246273.9.peg.1182 | CDS | wCle_NZ_AP0130 | 1042475 | 1042119 |
| fig\|246273.9.peg.1183 | CDS | wCle_NZ_AP0130 | 1042908 | 1042594 |
| fig\|246273.9.peg.1185 | CDS | wCle_NZ_AP0130 | 1043508 | 1043822 |
| fig\|246273.9.peg.1186 | CDS | wCle_NZ_AP0130 | 1043941 | 1044297 |
| fig\|246273.9.peg.1215 | CDS | wCle_NZ_AP0130 | 1074844 | 1075158 |
| fig\|246273.9.peg.1216 | CDS | wCle_NZ_AP0130 | 1075277 | 1075633 |
| fig\|246273.9.peg.1219 | CDS | wCle_NZ_AP0130 | 1078868 | 1079182 |
| fig\|246273.9.peg.1220 | CDS | wCle_NZ_AP0130 | 1079194 | 1079658 |
| fig\|246273.9.peg.1237 | CDS | wCle_NZ_AP0130 | 1094020 | 1093664 |
| fig\|246273.9.peg.1238 | CDS | wCle_NZ_AP0130 | 1094453 | 1094139 |
| fig\|246273.9.peg.1243 | CDS | wCle_NZ_AP0130 | 1098512 | 1098156 |
| fig\|246273.9.peg.1244 | CDS | wCle_NZ_AP0130 | 1098945 | 1098631 |
| fig\|246273.9.peg.1248 | CDS | wCle_NZ_AP0130 | 1101807 | 1102121 |
| fig\|246273.9.peg.1249 | CDS | wCle_NZ_AP0130 | 1102133 | 1102597 |
| fig\|246273.9.peg.1287 | CDS | wCle_NZ_AP0130 | 1138396 | 1138668 |
| fig\|246273.9.peg.1288 | CDS | wCle_NZ_AP0130 | 1139129 | 1138665 |
| fig\|246273.9.peg.1289 | CDS | wCle_NZ_AP0130 | 1139455 | 1139141 |
| fig\|246273.9.peg.1290 | CDS | wCle_NZ_AP0130 | 1139693 | 1140049 |
| fig\|246273.9.peg.1299 | CDS | wCle_NZ_AP0130 | 1148985 | 1148521 |
| fig\|246273.9.peg.1300 | CDS | wCle_NZ_AP0130 | 1149311 | 1148997 |
| fig\|246273.9.peg.1308 | CDS | wCle_NZ_AP0130 | 1155935 | 1155579 |
| fig\|246273.9.peg.1309 | CDS | wCle_NZ_AP0130 | 1156368 | 1156054 |
| fig\|246273.9.peg.1326 | CDS | wCle_NZ_AP0130 | 1169551 | 1169159 |
| fig\|246273.9.peg.1327 | CDS | wCle_NZ_AP0130 | 1169984 | 1169670 |
| fig\|246273.9.peg.1343 | CDS | wCle_NZ_AP0130 | 1187338 | 1187652 |
| fig\|246273.9.peg.1344 | CDS | wCle_NZ_AP0130 | 1187664 | 1188128 |
| fig\|246273.9.peg.1364 | CDS | wCle_NZ_AP0130 | 1207471 | 1207007 |
| fig\|246273.9.peg.1365 | CDS | wCle_NZ_AP0130 | 1207797 | 1207483 |
| fig\|246273.9.peg.1367 | CDS | wCle_NZ_AP0130 | 1208957 | 1209271 |

| | | | | | |
|---|---|---|---|---|---|
| | fig\|246273.9.peg.1368 | CDS | wCle_NZ_AP0130 | 1209390 | 1209746 |
| | fig\|246273.9.peg.1381 | CDS | wCle_NZ_AP0130 | 1223391 | 1223705 |
| | fig\|246273.9.peg.1382 | CDS | wCle_NZ_AP0130 | 1223824 | 1224276 |
| wDimm | - | - | - | - | - |
| wDcau | - | - | - | - | - |
| wFol | fig\|169402.10.peg.67 | CDS | wFol_NZ_CP0155 | 85045 | 85617 |
| | fig\|169402.10.peg.83 | CDS | wFol_NZ_CP0155 | 102814 | 102065 |
| | fig\|169402.10.peg.123 | CDS | wFol_NZ_CP0155 | 151236 | 150664 |
| | fig\|169402.10.peg.189 | CDS | wFol_NZ_CP0155 | 199332 | 198874 |
| | fig\|169402.10.peg.203 | CDS | wFol_NZ_CP0155 | 208998 | 209969 |
| | fig\|169402.10.peg.205 | CDS | wFol_NZ_CP0155 | 211839 | 211531 |
| | fig\|169402.10.peg.207 | CDS | wFol_NZ_CP0155 | 213865 | 213344 |
| | fig\|169402.10.peg.221 | CDS | wFol_NZ_CP0155 | 231074 | 230103 |
| | fig\|169402.10.peg.225 | CDS | wFol_NZ_CP0155 | 234594 | 235565 |
| | fig\|169402.10.peg.227 | CDS | wFol_NZ_CP0155 | 237341 | 238291 |
| | fig\|169402.10.peg.246 | CDS | wFol_NZ_CP0155 | 260870 | 261841 |
| | fig\|169402.10.peg.247 | CDS | wFol_NZ_CP0155 | 261958 | 262917 |
| | fig\|169402.10.peg.258 | CDS | wFol_NZ_CP0155 | 276312 | 275353 |
| | fig\|169402.10.peg.287 | CDS | wFol_NZ_CP0155 | 306562 | 307311 |
| | fig\|169402.10.peg.318 | CDS | wFol_NZ_CP0155 | 340321 | 340629 |
| | fig\|169402.10.peg.320 | CDS | wFol_NZ_CP0155 | 343163 | 342192 |
| | fig\|169402.10.peg.338 | CDS | wFol_NZ_CP0155 | 363464 | 363333 |
| | fig\|169402.10.peg.339 | CDS | wFol_NZ_CP0155 | 365540 | 363516 |
| | fig\|169402.10.peg.341 | CDS | wFol_NZ_CP0155 | 366410 | 366063 |
| | fig\|169402.10.peg.345 | CDS | wFol_NZ_CP0155 | 369085 | 369939 |
| | fig\|169402.10.peg.347 | CDS | wFol_NZ_CP0155 | 370478 | 372502 |
| | fig\|169402.10.peg.348 | CDS | wFol_NZ_CP0155 | 372554 | 372685 |
| | fig\|169402.10.peg.380 | CDS | wFol_NZ_CP0155 | 398067 | 396043 |
| | fig\|169402.10.peg.382 | CDS | wFol_NZ_CP0155 | 399460 | 398606 |
| | fig\|169402.10.peg.408 | CDS | wFol_NZ_CP0155 | 422342 | 423196 |
| | fig\|169402.10.peg.410 | CDS | wFol_NZ_CP0155 | 423735 | 425759 |
| | fig\|169402.10.peg.463 | CDS | wFol_NZ_CP0155 | 513040 | 513333 |
| | fig\|169402.10.peg.469 | CDS | wFol_NZ_CP0155 | 517685 | 518656 |
| | fig\|169402.10.peg.470 | CDS | wFol_NZ_CP0155 | 519689 | 518730 |
| | fig\|169402.10.peg.508 | CDS | wFol_NZ_CP0155 | 562897 | 563868 |
| | fig\|169402.10.peg.578 | CDS | wFol_NZ_CP0155 | 638367 | 638236 |
| | fig\|169402.10.peg.579 | CDS | wFol_NZ_CP0155 | 640443 | 638419 |
| | fig\|169402.10.peg.581 | CDS | wFol_NZ_CP0155 | 641313 | 640966 |
| | fig\|169402.10.peg.587 | CDS | wFol_NZ_CP0155 | 644907 | 645761 |
| | fig\|169402.10.peg.589 | CDS | wFol_NZ_CP0155 | 646300 | 648324 |
| | fig\|169402.10.peg.590 | CDS | wFol_NZ_CP0155 | 648376 | 648507 |
| | fig\|169402.10.peg.652 | CDS | wFol_NZ_CP0155 | 697686 | 697555 |

| | | | | |
|---|---|---|---|---|
| fig\|169402.10.peg.653 | CDS | wFol_NZ_CP0155 | 699762 | 697738 |
| fig\|169402.10.peg.693 | CDS | wFol_NZ_CP0155 | 735037 | 736257 |
| fig\|169402.10.peg.701 | CDS | wFol_NZ_CP0155 | 756671 | 757525 |
| fig\|169402.10.peg.703 | CDS | wFol_NZ_CP0155 | 758064 | 760088 |
| fig\|169402.10.peg.704 | CDS | wFol_NZ_CP0155 | 760140 | 760271 |
| fig\|169402.10.peg.735 | CDS | wFol_NZ_CP0155 | 790591 | 791163 |
| fig\|169402.10.peg.753 | CDS | wFol_NZ_CP0155 | 805472 | 805897 |
| fig\|169402.10.peg.754 | CDS | wFol_NZ_CP0155 | 805930 | 806430 |
| fig\|169402.10.peg.755 | CDS | wFol_NZ_CP0155 | 807266 | 806475 |
| fig\|169402.10.peg.773 | CDS | wFol_NZ_CP0155 | 825224 | 825973 |
| fig\|169402.10.peg.789 | CDS | wFol_NZ_CP0155 | 844113 | 843142 |
| fig\|169402.10.peg.792 | CDS | wFol_NZ_CP0155 | 844561 | 844857 |
| fig\|169402.10.peg.840 | CDS | wFol_NZ_CP0155 | 894075 | 893782 |
| fig\|169402.10.peg.842 | CDS | wFol_NZ_CP0155 | 894761 | 895564 |
| fig\|169402.10.peg.877 | CDS | wFol_NZ_CP0155 | 939075 | 938104 |
| fig\|169402.10.peg.897 | CDS | wFol_NZ_CP0155 | 965463 | 964504 |
| fig\|169402.10.peg.924 | CDS | wFol_NZ_CP0155 | 990543 | 989572 |
| fig\|169402.10.peg.927 | CDS | wFol_NZ_CP0155 | 992683 | 992390 |
| fig\|169402.10.peg.976 | CDS | wFol_NZ_CP0155 | 1039236 | 1040423 |
| fig\|169402.10.peg.1021 | CDS | wFol_NZ_CP0155 | 1081816 | 1082934 |
| fig\|169402.10.peg.1023 | CDS | wFol_NZ_CP0155 | 1085665 | 1083641 |
| fig\|169402.10.peg.1025 | CDS | wFol_NZ_CP0155 | 1087058 | 1086204 |
| fig\|169402.10.peg.1028 | CDS | wFol_NZ_CP0155 | 1089357 | 1089136 |
| fig\|169402.10.peg.1043 | CDS | wFol_NZ_CP0155 | 1116273 | 1117289 |
| fig\|169402.10.peg.1072 | CDS | wFol_NZ_CP0155 | 1142833 | 1142084 |
| fig\|169402.10.peg.1106 | CDS | wFol_NZ_CP0155 | 1171445 | 1170429 |
| fig\|169402.10.peg.1138 | CDS | wFol_NZ_CP0155 | 1206584 | 1207093 |
| fig\|169402.10.peg.1148 | CDS | wFol_NZ_CP0155 | 1224512 | 1226536 |
| fig\|169402.10.peg.1168 | CDS | wFol_NZ_CP0155 | 1246264 | 1245755 |
| fig\|169402.10.peg.1183 | CDS | wFol_NZ_CP0155 | 1258327 | 1259547 |
| fig\|169402.10.peg.1186 | CDS | wFol_NZ_CP0155 | 1262251 | 1263000 |
| fig\|169402.10.peg.1227 | CDS | wFol_NZ_CP0155 | 1322890 | 1321919 |
| fig\|169402.10.peg.1261 | CDS | wFol_NZ_CP0155 | 1353973 | 1354944 |
| fig\|169402.10.peg.1289 | CDS | wFol_NZ_CP0155 | 1380980 | 1380231 |
| fig\|169402.10.peg.1342 | CDS | wFol_NZ_CP0155 | 1428324 | 1429283 |
| fig\|169402.10.peg.1429 | CDS | wFol_NZ_CP0155 | 1535699 | 1536007 |
| fig\|169402.10.peg.1431 | CDS | wFol_NZ_CP0155 | 1538541 | 1537570 |
| fig\|169402.10.peg.1457 | CDS | wFol_NZ_CP0155 | 1569518 | 1568547 |
| fig\|169402.10.peg.1512 | CDS | wFol_NZ_CP0155 | 1630319 | 1631290 |
| fig\|169402.10.peg.1517 | CDS | wFol_NZ_CP0155 | 1640441 | 1639470 |
| fig\|169402.10.peg.1539 | CDS | wFol_NZ_CP0155 | 1668023 | 1667565 |
| fig\|169402.10.peg.1594 | CDS | wFol_NZ_CP0155 | 1725463 | 1724492 |

| | | | | | |
|---|---|---|---|---|---|
| | fig\|169402.10.peg.1595 | CDS | wFol_NZ_CP0155 | 1725754 | 1726326 |
| | fig\|169402.10.peg.1604 | CDS | wFol_NZ_CP0155 | 1735518 | 1736477 |
| | fig\|169402.10.peg.1634 | CDS | wFol_NZ_CP0155 | 1763802 | 1764773 |
| | fig\|169402.10.peg.1651 | CDS | wFol_NZ_CP0155 | 1782463 | 1781492 |
| | fig\|169402.10.peg.1652 | CDS | wFol_NZ_CP0155 | 1782754 | 1783326 |
| | fig\|169402.10.peg.1669 | CDS | wFol_NZ_CP0155 | 1797182 | 1798153 |
| wLbra | fig\|1812117.6.peg.13 | CDS | N762_781_tig19 | 7146 | 6934 |
| | fig\|1812117.6.peg.40 | CDS | N762_781_tig19 | 16351 | 16229 |
| | fig\|1812117.6.peg.61 | CDS | N851_N842_tig2( | 6679 | 6876 |
| | fig\|1812117.6.peg.107 | CDS | NODE_153_lengtl | 6539 | 6718 |
| | fig\|1812117.6.peg.124 | CDS | NODE_153_lengtl | 17139 | 16960 |
| | fig\|1812117.6.peg.360 | CDS | NODE_231_lengtl | 68282 | 68533 |
| | fig\|1812117.6.peg.406 | CDS | NODE_243_lengtl | 8405 | 8283 |
| | fig\|1812117.6.peg.474 | CDS | NODE_243_lengtl | 51712 | 51852 |
| | fig\|1812117.6.peg.521 | CDS | NODE_243_lengtl | 74785 | 74895 |
| | fig\|1812117.6.peg.531 | CDS | NODE_243_lengtl | 79556 | 79762 |
| | fig\|1812117.6.peg.564 | CDS | NODE_316_lengtl | 15888 | 16136 |
| | fig\|1812117.6.peg.619 | CDS | NODE_316_lengtl | 51369 | 51491 |
| | fig\|1812117.6.peg.662 | CDS | NODE_339_lengtl | 18017 | 17781 |
| | fig\|1812117.6.peg.701 | CDS | NODE_339_lengtl | 46665 | 46432 |
| | fig\|1812117.6.peg.720 | CDS | NODE_360_lengtl | 236 | 3 |
| | fig\|1812117.6.peg.796 | CDS | NODE_363_lengtl | 2794 | 2510 |
| | fig\|1812117.6.peg.840 | CDS | NODE_363_lengtl | 30258 | 30103 |
| | fig\|1812117.6.peg.951 | CDS | NODE_368_lengtl | 51526 | 51738 |
| | fig\|1812117.6.peg.1082 | CDS | NODE_444_lengtl | 24157 | 23897 |
| | fig\|1812117.6.peg.1324 | CDS | NODE_602_lengtl | 211 | 2 |
| | fig\|1812117.6.peg.1342 | CDS | NODE_602_lengtl | 8601 | 8416 |
| | fig\|1812117.6.peg.1430 | CDS | NODE_686_lengtl | 13983 | 13867 |
| | fig\|1812117.6.peg.1442 | CDS | NODE_735_lengtl | 3278 | 3099 |
| | fig\|1812117.6.peg.1569 | CDS | NODE_814_lengtl | 8355 | 8510 |
| | fig\|1812117.6.peg.1614 | CDS | NODE_939_lengtl | 3496 | 3660 |
| wLsig | fig\|80850.8.peg.147 | CDS | canuwLsigCt1p4_ | 93557 | 93673 |
| | fig\|80850.8.peg.415 | CDS | canuwLsigCt1p4_ | 257572 | 257733 |
| | fig\|80850.8.peg.535 | CDS | canuwLsigCt1p4_ | 337457 | 337570 |
| | fig\|80850.8.peg.587 | CDS | canuwLsigCt1p4_ | 379343 | 379552 |
| | fig\|80850.8.peg.876 | CDS | canuwLsigCt1p4_ | 615885 | 616058 |
| | fig\|80850.8.peg.881 | CDS | canuwLsigCt1p4_ | 617049 | 616882 |
| | fig\|80850.8.peg.1246 | CDS | canuwLsigCt1p4_ | 890552 | 890719 |
| | fig\|80850.8.peg.1299 | CDS | canuwLsigCt1p4_ | 926592 | 926401 |
| | fig\|80850.8.peg.1303 | CDS | canuwLsigCt1p4_ | 927234 | 927425 |
| | fig\|80850.8.peg.1410 | CDS | canuwLsigCt1p4_ | 997075 | 997251 |
| wLug | fig\|1335053.6.peg.34 | CDS | MUIY01000001.1 | 35358 | 35507 |

| | | | | |
|---|---|---|---|---|
| fig\|1335053.6.peg.36 | CDS | MUIY01000001.1 | 35987 | 36202 |
| fig\|1335053.6.peg.37 | CDS | MUIY01000001.1 | 36171 | 36362 |
| fig\|1335053.6.peg.60 | CDS | MUIY01000001.1 | 57573 | 56227 |
| fig\|1335053.6.peg.78 | CDS | MUIY01000001.1 | 72180 | 71839 |
| fig\|1335053.6.peg.79 | CDS | MUIY01000001.1 | 72671 | 72291 |
| fig\|1335053.6.peg.180 | CDS | MUIY01000001.1 | 179694 | 180512 |
| fig\|1335053.6.peg.181 | CDS | MUIY01000001.1 | 180630 | 180974 |
| fig\|1335053.6.peg.184 | CDS | MUIY01000001.1 | 182588 | 183406 |
| fig\|1335053.6.peg.185 | CDS | MUIY01000001.1 | 183524 | 183868 |
| fig\|1335053.6.peg.188 | CDS | MUIY01000001.1 | 184722 | 186068 |
| fig\|1335053.6.peg.204 | CDS | MUIY01000001.1 | 202204 | 202049 |
| fig\|1335053.6.peg.207 | CDS | MUIY01000001.1 | 203006 | 202893 |
| fig\|1335053.6.peg.214 | CDS | MUIY01000001.1 | 210047 | 209754 |
| fig\|1335053.6.peg.215 | CDS | MUIY01000001.1 | 210181 | 210062 |
| fig\|1335053.6.peg.216 | CDS | MUIY01000001.1 | 210571 | 210308 |
| fig\|1335053.6.peg.379 | CDS | MUIY01000001.1 | 364741 | 365178 |
| fig\|1335053.6.peg.440 | CDS | MUIY01000001.1 | 422985 | 423170 |
| fig\|1335053.6.peg.441 | CDS | MUIY01000001.1 | 423541 | 423852 |
| fig\|1335053.6.peg.449 | CDS | MUIY01000001.1 | 428972 | 429235 |
| fig\|1335053.6.peg.462 | CDS | MUIY01000001.1 | 439800 | 440018 |
| fig\|1335053.6.peg.463 | CDS | MUIY01000001.1 | 440165 | 440350 |
| fig\|1335053.6.peg.501 | CDS | MUIY01000001.1 | 476720 | 476595 |
| fig\|1335053.6.peg.502 | CDS | MUIY01000001.1 | 477223 | 476999 |
| fig\|1335053.6.peg.503 | CDS | MUIY01000001.1 | 477437 | 477267 |
| fig\|1335053.6.peg.524 | CDS | MUIY01000001.1 | 495640 | 495467 |
| fig\|1335053.6.peg.525 | CDS | MUIY01000001.1 | 496032 | 496145 |
| fig\|1335053.6.peg.580 | CDS | MUIY01000001.1 | 544729 | 544181 |
| fig\|1335053.6.peg.609 | CDS | MUIY01000001.1 | 569720 | 569388 |
| fig\|1335053.6.peg.611 | CDS | MUIY01000001.1 | 570811 | 570527 |
| fig\|1335053.6.peg.642 | CDS | MUIY01000001.1 | 597920 | 597624 |
| fig\|1335053.6.peg.643 | CDS | MUIY01000001.1 | 598416 | 598084 |
| fig\|1335053.6.peg.645 | CDS | MUIY01000001.1 | 599082 | 599378 |
| fig\|1335053.6.peg.657 | CDS | MUIY01000001.1 | 610183 | 609458 |
| fig\|1335053.6.peg.658 | CDS | MUIY01000001.1 | 610780 | 610574 |
| fig\|1335053.6.peg.659 | CDS | MUIY01000001.1 | 611016 | 610825 |
| fig\|1335053.6.peg.660 | CDS | MUIY01000001.1 | 611200 | 610985 |
| fig\|1335053.6.peg.662 | CDS | MUIY01000001.1 | 611829 | 611680 |
| fig\|1335053.6.peg.665 | CDS | MUIY01000001.1 | 612568 | 612759 |
| fig\|1335053.6.peg.666 | CDS | MUIY01000001.1 | 613140 | 612796 |
| fig\|1335053.6.peg.667 | CDS | MUIY01000001.1 | 614076 | 613258 |
| fig\|1335053.6.peg.668 | CDS | MUIY01000001.1 | 614637 | 614467 |
| fig\|1335053.6.peg.669 | CDS | MUIY01000001.1 | 614909 | 614718 |

| | | | | |
|---|---|---|---|---|
| fig\|1335053.6.peg.670 | CDS | MUIY01000001.1 | 615093 | 614878 |
| fig\|1335053.6.peg.675 | CDS | MUIY01000001.1 | 624453 | 624157 |
| fig\|1335053.6.peg.681 | CDS | MUIY01000001.1 | 629063 | 629185 |
| fig\|1335053.6.peg.682 | CDS | MUIY01000001.1 | 629205 | 629378 |
| fig\|1335053.6.peg.683 | CDS | MUIY01000001.1 | 629344 | 629529 |
| fig\|1335053.6.peg.685 | CDS | MUIY01000001.1 | 629905 | 630528 |
| fig\|1335053.6.peg.690 | CDS | MUIY01000001.1 | 636330 | 636866 |
| fig\|1335053.6.peg.695 | CDS | MUIY01000001.1 | 640321 | 639785 |
| fig\|1335053.6.peg.724 | CDS | MUIY01000001.1 | 666965 | 668257 |
| fig\|1335053.6.peg.726 | CDS | MUIY01000001.1 | 670348 | 670611 |
| fig\|1335053.6.peg.727 | CDS | MUIY01000001.1 | 670627 | 670752 |
| fig\|1335053.6.peg.728 | CDS | MUIY01000001.1 | 670842 | 671138 |
| fig\|1335053.6.peg.795 | CDS | MUIY01000001.1 | 734441 | 734611 |
| fig\|1335053.6.peg.796 | CDS | MUIY01000001.1 | 734655 | 734768 |
| fig\|1335053.6.peg.797 | CDS | MUIY01000001.1 | 734965 | 735138 |
| fig\|1335053.6.peg.798 | CDS | MUIY01000001.1 | 735150 | 735275 |
| fig\|1335053.6.peg.799 | CDS | MUIY01000001.1 | 735952 | 735608 |
| fig\|1335053.6.peg.800 | CDS | MUIY01000001.1 | 736888 | 736070 |
| fig\|1335053.6.peg.809 | CDS | MUIY01000001.1 | 744727 | 744431 |
| fig\|1335053.6.peg.810 | CDS | MUIY01000001.1 | 745221 | 744817 |
| fig\|1335053.6.peg.821 | CDS | MUIY01000001.1 | 757302 | 757177 |
| fig\|1335053.6.peg.822 | CDS | MUIY01000001.1 | 757512 | 757384 |
| fig\|1335053.6.peg.823 | CDS | MUIY01000001.1 | 757663 | 757514 |
| fig\|1335053.6.peg.835 | CDS | MUIY01000001.1 | 770803 | 771183 |
| fig\|1335053.6.peg.836 | CDS | MUIY01000001.1 | 771279 | 771635 |
| fig\|1335053.6.peg.865 | CDS | MUIY01000001.1 | 799879 | 800028 |
| fig\|1335053.6.peg.867 | CDS | MUIY01000001.1 | 800508 | 800723 |
| fig\|1335053.6.peg.868 | CDS | MUIY01000001.1 | 800692 | 800883 |
| fig\|1335053.6.peg.903 | CDS | MUIY01000001.1 | 835989 | 835645 |
| fig\|1335053.6.peg.904 | CDS | MUIY01000001.1 | 836925 | 836107 |
| fig\|1335053.6.peg.913 | CDS | MUIY01000001.1 | 848819 | 848571 |
| fig\|1335053.6.peg.917 | CDS | MUIY01000001.1 | 850193 | 850342 |
| fig\|1335053.6.peg.919 | CDS | MUIY01000001.1 | 850822 | 851037 |
| fig\|1335053.6.peg.920 | CDS | MUIY01000001.1 | 851006 | 851197 |
| fig\|1335053.6.peg.931 | CDS | MUIY01000001.1 | 864132 | 863947 |
| fig\|1335053.6.peg.932 | CDS | MUIY01000001.1 | 864497 | 864279 |
| fig\|1335053.6.peg.935 | CDS | MUIY01000001.1 | 866408 | 866064 |
| fig\|1335053.6.peg.936 | CDS | MUIY01000001.1 | 867344 | 866526 |
| fig\|1335053.6.peg.942 | CDS | MUIY01000001.1 | 871548 | 871766 |
| fig\|1335053.6.peg.955 | CDS | MUIY01000001.1 | 897293 | 898585 |
| fig\|1335053.6.peg.971 | CDS | MUIY01000001.1 | 915253 | 915411 |
| fig\|1335053.6.peg.972 | CDS | MUIY01000001.1 | 915414 | 915572 |

| | | | | |
|---|---|---|---|---|
| fig\|1335053.6.peg.978 | CDS | MUIY01000001.1 | 922717 | 922526 |
| fig\|1335053.6.peg.979 | CDS | MUIY01000001.1 | 922901 | 922686 |
| fig\|1335053.6.peg.981 | CDS | MUIY01000001.1 | 923530 | 923381 |
| fig\|1335053.6.peg.986 | CDS | MUIY01000001.1 | 926800 | 926985 |
| fig\|1335053.6.peg.987 | CDS | MUIY01000001.1 | 927356 | 927667 |
| fig\|1335053.6.peg.988 | CDS | MUIY01000001.1 | 927960 | 927745 |
| fig\|1335053.6.peg.989 | CDS | MUIY01000001.1 | 928085 | 927972 |
| fig\|1335053.6.peg.990 | CDS | MUIY01000001.1 | 928370 | 928116 |
| fig\|1335053.6.peg.1012 | CDS | MUIY01000001.1 | 954244 | 952952 |
| fig\|1335053.6.peg.1015 | CDS | MUIY01000001.1 | 959719 | 959549 |
| fig\|1335053.6.peg.1016 | CDS | MUIY01000001.1 | 959991 | 959800 |
| fig\|1335053.6.peg.1017 | CDS | MUIY01000001.1 | 960175 | 959960 |
| fig\|1335053.6.peg.1019 | CDS | MUIY01000001.1 | 960804 | 960655 |
| fig\|1335053.6.peg.1043 | CDS | MUIY01000001.1 | 982363 | 982250 |
| fig\|1335053.6.peg.1179 | CDS | MUIY01000001.1 | 1108978 | 1108847 |
| fig\|1335053.6.peg.1180 | CDS | MUIY01000001.1 | 1110037 | 1109369 |
| fig\|1335053.6.peg.1190 | CDS | MUIY01000001.1 | 1117346 | 1117552 |
| fig\|1335053.6.peg.1191 | CDS | MUIY01000001.1 | 1117659 | 1118207 |
| fig\|1335053.6.peg.1198 | CDS | MUIY01000001.1 | 1125869 | 1125696 |
| fig\|1335053.6.peg.1203 | CDS | MUIY01000001.1 | 1130274 | 1130606 |
| fig\|1335053.6.peg.1204 | CDS | MUIY01000001.1 | 1130769 | 1131065 |
| fig\|1335053.6.peg.1213 | CDS | MUIY01000001.1 | 1139222 | 1139554 |
| fig\|1335053.6.peg.1214 | CDS | MUIY01000001.1 | 1139717 | 1140013 |
| fig\|1335053.6.peg.1216 | CDS | MUIY01000001.1 | 1141833 | 1141522 |
| fig\|1335053.6.peg.1217 | CDS | MUIY01000001.1 | 1142389 | 1142204 |
| fig\|1335053.6.peg.1387 | CDS | MUIY01000002.1 | 162663 | 162352 |
| fig\|1335053.6.peg.1388 | CDS | MUIY01000002.1 | 163219 | 163034 |
| fig\|1335053.6.peg.1397 | CDS | MUIY01000002.1 | 170549 | 170734 |
| fig\|1335053.6.peg.1398 | CDS | MUIY01000002.1 | 171105 | 171416 |
| fig\|1335053.6.peg.1402 | CDS | MUIY01000002.1 | 174522 | 174211 |
| fig\|1335053.6.peg.1403 | CDS | MUIY01000002.1 | 175078 | 174893 |
| fig\|1335053.6.peg.1408 | CDS | MUIY01000002.1 | 177309 | 176965 |
| fig\|1335053.6.peg.1409 | CDS | MUIY01000002.1 | 178245 | 177427 |
| fig\|1335053.6.peg.1411 | CDS | MUIY01000002.1 | 179940 | 181232 |
| fig\|1335053.6.peg.1420 | CDS | MUIY01000002.1 | 193344 | 193192 |
| fig\|1335053.6.peg.1421 | CDS | MUIY01000002.1 | 193940 | 193515 |
| fig\|1335053.6.peg.1423 | CDS | MUIY01000002.1 | 194175 | 194324 |
| fig\|1335053.6.peg.1425 | CDS | MUIY01000002.1 | 194804 | 195019 |
| fig\|1335053.6.peg.1426 | CDS | MUIY01000002.1 | 194988 | 195179 |
| fig\|1335053.6.peg.1427 | CDS | MUIY01000002.1 | 196606 | 195260 |
| fig\|1335053.6.peg.1449 | CDS | MUIY01000002.1 | 222511 | 221585 |
| fig\|1335053.6.peg.1453 | CDS | MUIY01000002.1 | 225676 | 225275 |

| | | | | | |
|---|---|---|---|---|---|
| | fig\|1335053.6.peg.1508 | CDS | MUIY01000002.1 | 282380 | 281856 |
| | fig\|1335053.6.peg.1509 | CDS | MUIY01000002.1 | 282867 | 282736 |
| | fig\|1335053.6.peg.1534 | CDS | MUIY01000002.1 | 307626 | 307384 |
| | fig\|1335053.6.peg.1539 | CDS | MUIY01000002.1 | 309540 | 310076 |
| | fig\|1335053.6.peg.1614 | CDS | MUIY01000002.1 | 371439 | 371029 |
| | fig\|1335053.6.peg.1619 | CDS | MUIY01000002.1 | 381193 | 380975 |
| wMel | fig\|163164.1.peg.40 | CDS | wMel_NC_00297ε | 46860 | 46174 |
| | fig\|163164.1.peg.41 | CDS | wMel_NC_00297ε | 47198 | 46857 |
| | fig\|163164.1.peg.42 | CDS | wMel_NC_00297ε | 47689 | 47309 |
| | fig\|163164.1.peg.100 | CDS | wMel_NC_00297ε | 107885 | 109213 |
| | fig\|163164.1.peg.122 | CDS | wMel_NC_00297ε | 126290 | 126670 |
| | fig\|163164.1.peg.123 | CDS | wMel_NC_00297ε | 126781 | 127122 |
| | fig\|163164.1.peg.161 | CDS | wMel_NC_00297ε | 165751 | 165245 |
| | fig\|163164.1.peg.191 | CDS | wMel_NC_00297ε | 197767 | 197426 |
| | fig\|163164.1.peg.192 | CDS | wMel_NC_00297ε | 198258 | 197878 |
| | fig\|163164.1.peg.225 | CDS | wMel_NC_00297ε | 239076 | 237748 |
| | fig\|163164.1.peg.226 | CDS | wMel_NC_00297ε | 240427 | 239195 |
| | fig\|163164.1.peg.291 | CDS | wMel_NC_00297ε | 309070 | 308729 |
| | fig\|163164.1.peg.292 | CDS | wMel_NC_00297ε | 309561 | 309181 |
| | fig\|163164.1.peg.407 | CDS | wMel_NC_00297ε | 437813 | 438193 |
| | fig\|163164.1.peg.408 | CDS | wMel_NC_00297ε | 438304 | 438645 |
| | fig\|163164.1.peg.460 | CDS | wMel_NC_00297ε | 507960 | 507580 |
| | fig\|163164.1.peg.461 | CDS | wMel_NC_00297ε | 508294 | 507953 |
| | fig\|163164.1.peg.462 | CDS | wMel_NC_00297ε | 508785 | 508405 |
| | fig\|163164.1.peg.485 | CDS | wMel_NC_00297ε | 532315 | 532695 |
| | fig\|163164.1.peg.486 | CDS | wMel_NC_00297ε | 532806 | 533147 |
| | fig\|163164.1.peg.499 | CDS | wMel_NC_00297ε | 547952 | 547731 |
| | fig\|163164.1.peg.500 | CDS | wMel_NC_00297ε | 548107 | 549435 |
| | fig\|163164.1.peg.523 | CDS | wMel_NC_00297ε | 568583 | 568242 |
| | fig\|163164.1.peg.524 | CDS | wMel_NC_00297ε | 569074 | 568694 |
| | fig\|163164.1.peg.575 | CDS | wMel_NC_00297ε | 634537 | 634058 |
| | fig\|163164.1.peg.576 | CDS | wMel_NC_00297ε | 634725 | 635105 |
| | fig\|163164.1.peg.577 | CDS | wMel_NC_00297ε | 635216 | 635557 |
| | fig\|163164.1.peg.788 | CDS | wMel_NC_00297ε | 848079 | 847783 |
| | fig\|163164.1.peg.814 | CDS | wMel_NC_00297ε | 872362 | 872748 |
| | fig\|163164.1.peg.815 | CDS | wMel_NC_00297ε | 874056 | 873715 |
| | fig\|163164.1.peg.816 | CDS | wMel_NC_00297ε | 874547 | 874167 |
| | fig\|163164.1.peg.824 | CDS | wMel_NC_00297ε | 886563 | 886222 |
| | fig\|163164.1.peg.825 | CDS | wMel_NC_00297ε | 887054 | 886674 |
| | fig\|163164.1.peg.834 | CDS | wMel_NC_00297ε | 896358 | 896606 |
| | fig\|163164.1.peg.835 | CDS | wMel_NC_00297ε | 896627 | 897007 |
| | fig\|163164.1.peg.836 | CDS | wMel_NC_00297ε | 897118 | 897459 |

| | | | | | |
|---|---|---|---|---|---|
| | fig\|163164.1.peg.843 | CDS | wMel_NC_00297 | 908139 | 908546 |
| | fig\|163164.1.peg.1102 | CDS | wMel_NC_00297 | 1172316 | 1172696 |
| | fig\|163164.1.peg.1103 | CDS | wMel_NC_00297 | 1172807 | 1173148 |
| | fig\|163164.1.peg.1214 | CDS | wMel_NC_00297 | 226892 | 227221 |
| | fig\|163164.1.peg.1215 | CDS | wMel_NC_00297 | 227205 | 227405 |
| | fig\|163164.1.peg.1216 | CDS | wMel_NC_00297 | 227425 | 227817 |
| | fig\|163164.1.peg.1229 | CDS | wMel_NC_00297 | 310033 | 309626 |
| | fig\|163164.1.peg.1230 | CDS | wMel_NC_00297 | 310253 | 310053 |
| | fig\|163164.1.peg.1231 | CDS | wMel_NC_00297 | 310566 | 310237 |
| | fig\|163164.1.peg.1270 | CDS | wMel_NC_00297 | 723000 | 722692 |
| | fig\|163164.1.peg.1271 | CDS | wMel_NC_00297 | 723618 | 723325 |
| | fig\|163164.1.peg.1284 | CDS | wMel_NC_00297 | 868830 | 869237 |
| | fig\|163164.1.peg.1285 | CDS | wMel_NC_00297 | 873202 | 872774 |
| | fig\|163164.1.peg.1287 | CDS | wMel_NC_00297 | 874805 | 875197 |
| | fig\|163164.1.peg.1288 | CDS | wMel_NC_00297 | 897773 | 898165 |
| | fig\|163164.1.peg.1301 | CDS | wMel_NC_00297 | 1068928 | 1068521 |
| | fig\|163164.1.peg.1302 | CDS | wMel_NC_00297 | 1069148 | 1068948 |
| | fig\|163164.1.peg.1303 | CDS | wMel_NC_00297 | 1069461 | 1069132 |
| wMhie | NODE_13561_length_1237_co | CDS | | 49 | 213 |
| | NODE_1527_length_14466_co | CDS | | 98 | 262 |
| | NODE_1788_length_12468_co | CDS | | 12467 | 12207 |
| | NODE_2037_length_10977_co | CDS | | 7126 | 6770 |
| | NODE_2047_length_10911_co | CDS | | 1 | 318 |
| | NODE_2794_length_7599_cov_ | CDS | | 624 | 22 |
| | NODE_3238_length_6354_cov_ | CDS | | 439 | 17 |
| | NODE_3292_length_6258_cov_ | CDS | | 6256 | 6005 |
| | NODE_3460_length_5886_cov_ | CDS | | 5884 | 5648 |
| | NODE_3707_length_5372_cov_ | CDS | | 254 | 3 |
| | NODE_4051_length_4797_cov_ | CDS | | 4691 | 4795 |
| | NODE_4620_length_3999_cov_ | CDS | | 1 | 162 |
| | NODE_4627_length_3995_cov_ | CDS | | 3820 | 3993 |
| | NODE_4649_length_3964_cov_ | CDS | | 3 | 110 |
| | NODE_5236_length_3419_cov_ | CDS | | 251 | 60 |
| | NODE_5396_length_3316_cov_ | CDS | | 2 | 304 |
| | NODE_6358_length_2708_cov_ | CDS | | 2706 | 2566 |
| | NODE_8063_length_2068_cov_ | CDS | | 224 | 565 |
| wNfla | fig\|1854759.6.peg.2 | CDS | NZ_LYUW010000 | 720 | 989 |
| | fig\|1854759.6.peg.64 | CDS | NZ_LYUW010000 | 56688 | 56948 |
| | fig\|1854759.6.peg.65 | CDS | NZ_LYUW010000 | 56938 | 57060 |
| | fig\|1854759.6.peg.113 | CDS | NZ_LYUW010000 | 52251 | 52469 |
| | fig\|1854759.6.peg.155 | CDS | NZ_LYUW010000 | 38862 | 38674 |
| | fig\|1854759.6.peg.158 | CDS | NZ_LYUW010000 | 39669 | 39466 |

| | | | | |
|---|---|---|---|---|
| fig\|1854759.6.peg.205 | CDS | NZ_LYUW010000 | 303 | 1 |
| fig\|1854759.6.peg.239 | CDS | NZ_LYUW010000 | 65 | 229 |
| fig\|1854759.6.peg.412 | CDS | NZ_LYUW010000 | 221 | 3 |
| fig\|1854759.6.peg.414 | CDS | NZ_LYUW010000 | 1214 | 1354 |
| fig\|1854759.6.peg.415 | CDS | NZ_LYUW010000 | 1869 | 2138 |
| fig\|1854759.6.peg.581 | CDS | NZ_LYUW010000 | 1 | 231 |
| fig\|1854759.6.peg.582 | CDS | NZ_LYUW010000 | 1107 | 646 |
| fig\|1854759.6.peg.583 | CDS | NZ_LYUW010000 | 1324 | 1142 |
| fig\|1854759.6.peg.584 | CDS | NZ_LYUW010000 | 1665 | 1321 |
| fig\|1854759.6.peg.603 | CDS | NZ_LYUW010000 | 17926 | 18144 |
| fig\|1854759.6.peg.627 | CDS | NZ_LYUW010000 | 435 | 301 |
| fig\|1854759.6.peg.708 | CDS | NZ_LYUW010000 | 159 | 1 |
| fig\|1854759.6.peg.740 | CDS | NZ_LYUW010000 | 14906 | 15169 |
| fig\|1854759.6.peg.765 | CDS | NZ_LYUW010000 | 13803 | 14135 |
| fig\|1854759.6.peg.801 | CDS | NZ_LYUW010000 | 221 | 3 |
| fig\|1854759.6.peg.812 | CDS | NZ_LYUW010000 | 8812 | 7787 |
| fig\|1854759.6.peg.871 | CDS | NZ_LYUW010000 | 221 | 3 |
| fig\|1854759.6.peg.900 | CDS | NZ_LYUW010000 | 11442 | 11140 |
| fig\|1854759.6.peg.901 | CDS | NZ_LYUW010000 | 11671 | 11414 |
| fig\|1854759.6.peg.902 | CDS | NZ_LYUW010000 | 11927 | 11658 |
| fig\|1854759.6.peg.903 | CDS | NZ_LYUW010000 | 12201 | 11932 |
| fig\|1854759.6.peg.916 | CDS | NZ_LYUW010000 | 194 | 3 |
| fig\|1854759.6.peg.960 | CDS | NZ_LYUW010000 | 234 | 112 |
| fig\|1854759.6.peg.961 | CDS | NZ_LYUW010000 | 484 | 224 |
| fig\|1854759.6.peg.1006 | CDS | NZ_LYUW010000 | 10873 | 11127 |
| fig\|1854759.6.peg.1007 | CDS | NZ_LYUW010000 | 11151 | 11234 |
| fig\|1854759.6.peg.1060 | CDS | NZ_LYUW010000 | 221 | 3 |
| fig\|1854759.6.peg.1091 | CDS | NZ_LYUW010000 | 1030 | 779 |
| fig\|1854759.6.peg.1092 | CDS | NZ_LYUW010000 | 1179 | 1000 |
| fig\|1854759.6.peg.1093 | CDS | NZ_LYUW010000 | 1705 | 1412 |
| fig\|1854759.6.peg.1114 | CDS | NZ_LYUW010000 | 234 | 112 |
| fig\|1854759.6.peg.1115 | CDS | NZ_LYUW010000 | 484 | 224 |
| fig\|1854759.6.peg.1227 | CDS | NZ_LYUW010000 | 6489 | 6572 |
| fig\|1854759.6.peg.1248 | CDS | NZ_LYUW010000 | 6113 | 6331 |
| fig\|1854759.6.peg.1266 | CDS | NZ_LYUW010000 | 221 | 3 |
| fig\|1854759.6.peg.1283 | CDS | NZ_LYUW010000 | 369 | 1 |
| fig\|1854759.6.peg.1297 | CDS | NZ_LYUW010000 | 5182 | 5352 |
| fig\|1854759.6.peg.1333 | CDS | NZ_LYUW010000 | 433 | 95 |
| fig\|1854759.6.peg.1335 | CDS | NZ_LYUW010000 | 1061 | 1357 |
| fig\|1854759.6.peg.1349 | CDS | NZ_LYUW010000 | 2677 | 2273 |
| fig\|1854759.6.peg.1378 | CDS | NZ_LYUW010000 | 3157 | 3375 |
| fig\|1854759.6.peg.1400 | CDS | NZ_LYUW010000 | 2815 | 2495 |

| | | | | | |
|---|---|---|---|---|---|
| | fig\|1854759.6.peg.1411 | CDS | NZ_LYUW010001 | 2649 | 2867 |
| | fig\|1854759.6.peg.1428 | CDS | NZ_LYUW010001 | 221 | 3 |
| | fig\|1854759.6.peg.1448 | CDS | NZ_LYUW010001 | 277 | 954 |
| | fig\|1854759.6.peg.1449 | CDS | NZ_LYUW010001 | 1138 | 1614 |
| | fig\|1854759.6.peg.1463 | CDS | NZ_LYUW010001 | 248 | 1045 |
| | fig\|1854759.6.peg.1468 | CDS | NZ_LYUW010001 | 1134 | 1352 |
| | fig\|1854759.6.peg.1476 | CDS | NZ_LYUW010001 | 221 | 3 |
| | fig\|1854759.6.peg.1479 | CDS | NZ_LYUW010001 | 81 | 461 |
| | fig\|1854759.6.peg.1483 | CDS | NZ_LYUW010001 | 234 | 112 |
| | fig\|1854759.6.peg.1484 | CDS | NZ_LYUW010001 | 484 | 224 |
| | fig\|1854759.6.peg.1486 | CDS | NZ_LYUW010001 | 173 | 466 |
| | fig\|1854759.6.peg.1487 | CDS | NZ_LYUW010001 | 758 | 522 |
| | fig\|1854759.6.peg.1488 | CDS | NZ_LYUW010001 | 74 | 358 |
| | fig\|1854759.6.peg.1494 | CDS | NZ_LYUW010001 | 551 | 24 |
| | fig\|1854759.6.peg.1496 | CDS | NZ_LYUW010001 | 400 | 621 |
| | fig\|1854759.6.peg.1498 | CDS | NZ_LYUW010001 | 230 | 78 |
| | fig\|1854759.6.peg.1499 | CDS | NZ_LYUW010001 | 221 | 3 |
| | fig\|1854759.6.peg.1500 | CDS | NZ_LYUW010001 | 511 | 329 |
| | fig\|1854759.6.peg.1505 | CDS | NZ_LYUW010001 | 81 | 422 |
| | fig\|1854759.6.peg.1506 | CDS | NZ_LYUW010001 | 435 | 208 |
| | fig\|1854759.6.peg.1512 | CDS | NZ_LYUW010001 | 3 | 185 |
| | fig\|1854759.6.peg.1515 | CDS | NZ_LYUW010001 | 296 | 3 |
| | fig\|1854759.6.peg.1516 | CDS | NZ_LYUW010001 | 1 | 249 |
| | fig\|1854759.6.peg.1517 | CDS | NZ_LYUW010001 | 32 | 235 |
| | fig\|1854759.6.peg.1520 | CDS | NZ_LYUW010001 | 202 | 2 |
| wOo | - | - | - | - | - |
| wOv | - | - | - | - | - |
| wPpe | fig\|6666666.425803.peg.109 | CDS | wPpe_NZ_MJMG | 98290 | 98141 |
| | fig\|6666666.425803.peg.431 | CDS | NZ_MJMG01000 | 22176 | 21970 |
| | fig\|6666666.425803.peg.938 | CDS | NZ_MJMG01000 | 53277 | 53366 |
| | fig\|6666666.425803.peg.959 | CDS | NZ_MJMG01000 | 13531 | 13698 |
| | fig\|6666666.425803.peg.961 | CDS | NZ_MJMG01000 | 13850 | 14083 |
| wPip | fig\|955.1.peg.1 | CDS | wPip_culex15b06 | 967 | 494 |
| | fig\|955.1.peg.4 | CDS | wPip_culex15b06 | 2316 | 2738 |
| | fig\|955.1.peg.57 | CDS | culex182c05.q1k | 733 | 128 |
| | fig\|955.1.peg.58 | CDS | culex182c05.q1k | 1290 | 883 |
| | fig\|955.1.peg.59 | CDS | culex182c05.q1k | 1478 | 1290 |
| | fig\|955.1.peg.62 | CDS | culex169h11.p1k | 269 | 649 |
| | fig\|955.1.peg.75 | CDS | culex57g05.p1k | 1944 | 1072 |
| | fig\|955.1.peg.87 | CDS | culex57g05.p1k | 12448 | 12630 |
| | fig\|955.1.peg.113 | CDS | culex8d01.q1ka | 23914 | 24786 |
| | fig\|955.1.peg.186 | CDS | culex8d01.q1ka | 112015 | 112683 |

| | | | | |
|---|---|---|---|---|
| fig\|955.1.peg.187 | CDS | culex8d01.q1ka | 112655 | 113245 |
| fig\|955.1.peg.230 | CDS | culex136g12.p1k | 19996 | 19124 |
| fig\|955.1.peg.240 | CDS | culex136g12.p1k | 26263 | 26745 |
| fig\|955.1.peg.252 | CDS | culex164g03.p1k | 80 | 538 |
| fig\|955.1.peg.255 | CDS | culex164g03.p1k | 2734 | 1862 |
| fig\|955.1.peg.261 | CDS | culex164g03.p1k | 8856 | 9074 |
| fig\|955.1.peg.266 | CDS | culex56c01.p1k | 3284 | 3607 |
| fig\|955.1.peg.276 | CDS | culex56c01.p1k | 13242 | 12370 |
| fig\|955.1.peg.278 | CDS | culex56c01.p1k | 13726 | 14202 |
| fig\|955.1.peg.279 | CDS | culex56c01.p1k | 14186 | 14350 |
| fig\|955.1.peg.280 | CDS | culex56c01.p1k | 14310 | 14666 |
| fig\|955.1.peg.284 | CDS | culex56c01.p1k | 16776 | 16420 |
| fig\|955.1.peg.285 | CDS | culex56c01.p1k | 16900 | 16736 |
| fig\|955.1.peg.286 | CDS | culex56c01.p1k | 17360 | 16884 |
| fig\|955.1.peg.288 | CDS | culex56c01.p1k | 19617 | 19874 |
| fig\|955.1.peg.289 | CDS | culex56c01.p1k | 20056 | 20196 |
| fig\|955.1.peg.290 | CDS | culex56c01.p1k | 20263 | 20502 |
| fig\|955.1.peg.294 | CDS | culex56c01.p1k | 25446 | 24574 |
| fig\|955.1.peg.331 | CDS | culex61a11.q1k | 9530 | 9123 |
| fig\|955.1.peg.332 | CDS | culex61a11.q1k | 10499 | 9627 |
| fig\|955.1.peg.345 | CDS | culex61a11.q1k | 26704 | 26348 |
| fig\|955.1.peg.346 | CDS | culex61a11.q1k | 26828 | 26664 |
| fig\|955.1.peg.347 | CDS | culex61a11.q1k | 27288 | 26812 |
| fig\|955.1.peg.360 | CDS | culex12a01.q1k | 3952 | 3080 |
| fig\|955.1.peg.391 | CDS | culex6c12.q1k | 127 | 723 |
| fig\|955.1.peg.402 | CDS | culex6c12.q1k | 11993 | 11586 |
| fig\|955.1.peg.405 | CDS | culex35f11.p1k | 96 | 968 |
| fig\|955.1.peg.409 | CDS | culex35f11.p1k | 3601 | 3272 |
| fig\|955.1.peg.412 | CDS | culex35f11.p1k | 5690 | 5100 |
| fig\|955.1.peg.413 | CDS | culex35f11.p1k | 6330 | 5662 |
| fig\|955.1.peg.426 | CDS | culex35f11.p1k | 21351 | 20113 |
| fig\|955.1.peg.429 | CDS | culex35f11.p1k | 22690 | 21818 |
| fig\|955.1.peg.469 | CDS | culex171h04.q1k | 6 | 716 |
| fig\|955.1.peg.500 | CDS | culex98d12.p1k | 19840 | 20712 |
| fig\|955.1.peg.501 | CDS | culex98d12.p1k | 21014 | 20790 |
| fig\|955.1.peg.503 | CDS | culex98d12.p1k | 21935 | 21720 |
| fig\|955.1.peg.505 | CDS | culex98d12.p1k | 22606 | 22400 |
| fig\|955.1.peg.566 | CDS | culex102f01.p1k | 49864 | 48992 |
| fig\|955.1.peg.576 | CDS | culex107d08.p1k | 11768 | 10788 |
| fig\|955.1.peg.577 | CDS | culex107d08.p1k | 12026 | 11778 |
| fig\|955.1.peg.597 | CDS | culex107d08.p1k | 29347 | 30219 |
| fig\|955.1.peg.638 | CDS | culex184h04.q1k | 37738 | 37938 |

| | | | | |
|---|---|---|---|---|
| fig\|955.1.peg.641 | CDS | culex22a03.p1k | 217 | 807 |
| fig\|955.1.peg.642 | CDS | culex22a03.p1k | 1186 | 857 |
| fig\|955.1.peg.644 | CDS | culex48c09.p1k | 458 | 198 |
| fig\|955.1.peg.645 | CDS | culex48c09.p1k | 1302 | 520 |
| fig\|955.1.peg.652 | CDS | culex95h05.q1k | 13131 | 12259 |
| fig\|955.1.peg.656 | CDS | culex95h05.q1k | 16191 | 15319 |
| fig\|955.1.peg.659 | CDS | culex95h05.q1k | 18585 | 18833 |
| fig\|955.1.peg.660 | CDS | culex95h05.q1k | 18843 | 19253 |
| fig\|955.1.peg.661 | CDS | culex95h05.q1k | 19225 | 19815 |
| fig\|955.1.peg.662 | CDS | culex95h05.q1k | 20164 | 19835 |
| fig\|955.1.peg.674 | CDS | culex46e07.p1k | 3743 | 4843 |
| fig\|955.1.peg.675 | CDS | culex46e07.p1k | 4849 | 4980 |
| fig\|955.1.peg.685 | CDS | culex46e07.p1k | 16478 | 16678 |
| fig\|955.1.peg.686 | CDS | culex46e07.p1k | 16755 | 17627 |
| fig\|955.1.peg.723 | CDS | culex111d03.p1k | 738 | 37 |
| fig\|955.1.peg.728 | CDS | culex111d03.p1k | 4889 | 5761 |
| fig\|955.1.peg.731 | CDS | culex111d03.p1k | 9470 | 9048 |
| fig\|955.1.peg.734 | CDS | culex111d03.p1k | 12195 | 12971 |
| fig\|955.1.peg.735 | CDS | culex111d03.p1k | 12995 | 13243 |
| fig\|955.1.peg.736 | CDS | culex111d03.p1k | 13253 | 13663 |
| fig\|955.1.peg.737 | CDS | culex111d03.p1k | 13635 | 14045 |
| fig\|955.1.peg.738 | CDS | culex111d03.p1k | 14094 | 14225 |
| fig\|955.1.peg.752 | CDS | culex111d03.p1k | 34110 | 34532 |
| fig\|955.1.peg.756 | CDS | culex111d03.p1k | 36581 | 35709 |
| fig\|955.1.peg.759 | CDS | culex111d03.p1k | 38450 | 39322 |
| fig\|955.1.peg.774 | CDS | culex111d03.p1k | 55089 | 55565 |
| fig\|955.1.peg.775 | CDS | culex111d03.p1k | 55549 | 55713 |
| fig\|955.1.peg.776 | CDS | culex111d03.p1k | 55673 | 56029 |
| fig\|955.1.peg.786 | CDS | culex111d03.p1k | 65158 | 64802 |
| fig\|955.1.peg.791 | CDS | culex60b10.p1k | 312 | 1 |
| fig\|955.1.peg.798 | CDS | culex60b10.p1k | 2259 | 2450 |
| fig\|955.1.peg.806 | CDS | culex60b10.p1k | 11112 | 10540 |
| fig\|955.1.peg.809 | CDS | culex161e12.p1k | 2192 | 3064 |
| fig\|955.1.peg.846 | CDS | culex40h12.p1k | 321 | 73 |
| fig\|955.1.peg.851 | CDS | culex40h12.p1k | 3347 | 3820 |
| fig\|955.1.peg.852 | CDS | culex40h12.p1k | 4609 | 3737 |
| fig\|955.1.peg.857 | CDS | culex40h12.p1k | 7390 | 6800 |
| fig\|955.1.peg.858 | CDS | culex40h12.p1k | 8030 | 7362 |
| fig\|955.1.peg.862 | CDS | culex40h12.p1k | 9215 | 10147 |
| fig\|955.1.peg.884 | CDS | culex74f02.p1k | 18546 | 17674 |
| fig\|955.1.peg.916 | CDS | culex154c10.q1k | 672 | 1307 |
| fig\|955.1.peg.917 | CDS | culex154c10.q1k | 1273 | 1908 |

| | | | | |
|---|---|---|---|---|
| fig\|955.1.peg.922 | CDS | culex154c10.q1k | 4895 | 4566 |
| fig\|955.1.peg.933 | CDS | culex154c10.q1k | 11862 | 12482 |
| fig\|955.1.peg.951 | CDS | culex6a11.q1k | 19335 | 19583 |
| fig\|955.1.peg.952 | CDS | culex6a11.q1k | 19593 | 19904 |
| fig\|955.1.peg.953 | CDS | culex183b05.p1k | 647 | 291 |
| fig\|955.1.peg.954 | CDS | culex183b05.p1k | 771 | 607 |
| fig\|955.1.peg.955 | CDS | culex183b05.p1k | 1231 | 755 |
| fig\|955.1.peg.977 | CDS | culex181f03.p1k | 6851 | 7723 |
| fig\|955.1.peg.979 | CDS | culex181f03.p1k | 8734 | 9210 |
| fig\|955.1.peg.980 | CDS | culex181f03.p1k | 9194 | 9358 |
| fig\|955.1.peg.981 | CDS | culex181f03.p1k | 9318 | 9674 |
| fig\|955.1.peg.998 | CDS | culex147c02.q1k | 3641 | 4513 |
| fig\|955.1.peg.999 | CDS | culex147c02.q1k | 4610 | 5017 |
| fig\|955.1.peg.1002 | CDS | culex180a01.p1k | 140 | 3 |
| fig\|955.1.peg.1003 | CDS | culex180a01.p1k | 780 | 112 |
| fig\|955.1.peg.1005 | CDS | culex180a01.p1k | 1849 | 2721 |
| fig\|955.1.peg.1032 | CDS | culex180a01.p1k | 27157 | 27633 |
| fig\|955.1.peg.1033 | CDS | culex180a01.p1k | 27617 | 27745 |
| fig\|955.1.peg.1036 | CDS | culex150d03.p1k | 4866 | 4444 |
| fig\|955.1.peg.1043 | CDS | culex150d03.p1k | 11328 | 11678 |
| fig\|955.1.peg.1045 | CDS | culex150d03.p1k | 11967 | 12557 |
| fig\|955.1.peg.1051 | CDS | culex150d03.p1k | 20112 | 20588 |
| fig\|955.1.peg.1052 | CDS | culex150d03.p1k | 20572 | 20736 |
| fig\|955.1.peg.1053 | CDS | culex150d03.p1k | 20696 | 21052 |
| fig\|955.1.peg.1084 | CDS | culex35f07.p1k | 28920 | 29792 |
| fig\|955.1.peg.1111 | CDS | culex126f06.p1k | 358 | 1230 |
| fig\|955.1.peg.1112 | CDS | culex126f06.p1k | 1573 | 1409 |
| fig\|955.1.peg.1113 | CDS | culex126f06.p1k | 2033 | 1557 |
| fig\|955.1.peg.1116 | CDS | culex126f06.p1k | 4283 | 4152 |
| fig\|955.1.peg.1117 | CDS | culex126f06.p1k | 4420 | 4289 |
| fig\|955.1.peg.1135 | CDS | culex79g04.p1k | 13719 | 14195 |
| fig\|955.1.peg.1136 | CDS | culex79g04.p1k | 14179 | 14343 |
| fig\|955.1.peg.1137 | CDS | culex79g04.p1k | 14303 | 14659 |
| fig\|955.1.peg.1167 | CDS | culex79g04.p1k | 47332 | 46976 |
| fig\|955.1.peg.1168 | CDS | culex79g04.p1k | 47474 | 47292 |
| fig\|955.1.peg.1169 | CDS | culex79g04.p1k | 48362 | 47490 |
| fig\|955.1.peg.1170 | CDS | culex79g04.p1k | 48902 | 48432 |
| fig\|955.1.peg.1176 | CDS | culex79g04.p1k | 56389 | 56865 |
| fig\|955.1.peg.1177 | CDS | culex79g04.p1k | 56849 | 57013 |
| fig\|955.1.peg.1178 | CDS | culex79g04.p1k | 56973 | 57329 |
| fig\|955.1.peg.1225 | CDS | culex176d05.q1k | 399 | 1637 |
| fig\|955.1.peg.1240 | CDS | culex176d05.q1k | 15732 | 14494 |

| | | | | | |
|---|---|---|---|---|---|
| | fig\|955.1.peg.1253 | CDS | culex176d05.q1k | 28740 | 27868 |
| | fig\|955.1.peg.1261 | CDS | culex176d05.q1k | 35974 | 35102 |
| | fig\|955.1.peg.1262 | CDS | culex176d05.q1k | 36149 | 36505 |
| | fig\|955.1.peg.1269 | CDS | culex70a11.p1k | 314 | 904 |
| | fig\|955.1.peg.1279 | CDS | culex70a11.p1k | 10273 | 9791 |
| | fig\|955.1.peg.1285 | CDS | culex70a11.p1k | 15897 | 15532 |
| | fig\|955.1.peg.1324 | CDS | culex180d12.p1k | 8580 | 8912 |
| | fig\|955.1.peg.1329 | CDS | culex180d12.p1k | 15056 | 15928 |
| | fig\|955.1.peg.1334 | CDS | culex180d12.p1k | 19616 | 20488 |
| | fig\|955.1.peg.1358 | CDS | culex143c03.p1k | 883 | 5 |
| | fig\|955.1.peg.1367 | CDS | culex143c03.p1k | 11951 | 12823 |
| | fig\|955.1.peg.1368 | CDS | culex143c03.p1k | 13004 | 13672 |
| | fig\|955.1.peg.1369 | CDS | culex143c03.p1k | 13644 | 14234 |
| | fig\|955.1.peg.1402 | CDS | culex143c03.p1k | 42759 | 43241 |
| | fig\|955.1.peg.1412 | CDS | culex143c03.p1k | 53318 | 52953 |
| | fig\|955.1.peg.1414 | CDS | culex125c07.q1k | 866 | 3 |
| | fig\|955.1.peg.1451 | CDS | culex125c07.q1k | 29492 | 28620 |
| | fig\|955.1.peg.1465 | CDS | culex37f09.p1k | 89 | 355 |
| | fig\|955.1.peg.1466 | CDS | culex37f09.p1k | 339 | 503 |
| | fig\|955.1.peg.1467 | CDS | culex37f09.p1k | 463 | 819 |
| | fig\|955.1.peg.1474 | CDS | culex37f09.p1k | 5160 | 4678 |
| | fig\|955.1.peg.1484 | CDS | culex37f09.p1k | 15988 | 15581 |
| | fig\|955.1.peg.1485 | CDS | culex37f09.p1k | 16516 | 16133 |
| | fig\|955.1.peg.1486 | CDS | culex37f09.p1k | 17413 | 16682 |
| | fig\|955.1.peg.1487 | CDS | culex160e12.q1k | 772 | 182 |
| | fig\|955.1.peg.1507 | CDS | culex160e12.q1k | 25453 | 26325 |
| | fig\|955.1.peg.1508 | CDS | culex160e12.q1k | 26848 | 26372 |
| | fig\|955.1.peg.1509 | CDS | culex160e12.q1k | 27939 | 27067 |
| | fig\|955.1.peg.1511 | CDS | culex160e12.q1k | 28406 | 28735 |
| | fig\|955.1.peg.1514 | CDS | culex160e12.q1k | 30140 | 29544 |
| | fig\|955.1.peg.1526 | CDS | culex16c12.p1k | 14148 | 15020 |
| | fig\|955.1.peg.1547 | CDS | culex16c12.p1k | 33201 | 33064 |
| | fig\|955.1.peg.1549 | CDS | culex102c04.p1k | 885 | 94 |
| | fig\|955.1.peg.1560 | CDS | culex102c04.p1k | 12390 | 13490 |
| | fig\|955.1.peg.1561 | CDS | culex102c04.p1k | 13496 | 13627 |
| | fig\|955.1.peg.1570 | CDS | culex102c04.p1k | 21652 | 22524 |
| | fig\|955.1.peg.1584 | CDS | wPip_culex15b06 | 29 | 313 |
| wstri | fig\|368602.8.peg.23 | CDS | MUIX01000001.1 | 17370 | 18716 |
| | fig\|368602.8.peg.96 | CDS | MUIX01000001.1 | 88853 | 89002 |
| | fig\|368602.8.peg.103 | CDS | MUIX01000001.1 | 95367 | 95218 |
| | fig\|368602.8.peg.196 | CDS | MUIX01000001.1 | 179971 | 181317 |
| | fig\|368602.8.peg.227 | CDS | MUIX01000001.1 | 216498 | 216647 |

| | | | | |
|---|---|---|---|---|
| fig\|368602.8.peg.235 | CDS | MUIX01000001.1 | 223177 | 223326 |
| fig\|368602.8.peg.252 | CDS | MUIX01000001.1 | 242563 | 241616 |
| fig\|368602.8.peg.262 | CDS | MUIX01000001.1 | 247657 | 247508 |
| fig\|368602.8.peg.275 | CDS | MUIX01000001.1 | 255418 | 255567 |
| fig\|368602.8.peg.287 | CDS | MUIX01000001.1 | 264423 | 265121 |
| fig\|368602.8.peg.295 | CDS | MUIX01000001.1 | 280878 | 279931 |
| fig\|368602.8.peg.317 | CDS | MUIX01000001.1 | 296748 | 295402 |
| fig\|368602.8.peg.319 | CDS | MUIX01000001.1 | 298277 | 298498 |
| fig\|368602.8.peg.321 | CDS | MUIX01000001.1 | 299137 | 299253 |
| fig\|368602.8.peg.344 | CDS | MUIX01000001.1 | 319167 | 320039 |
| fig\|368602.8.peg.366 | CDS | MUIX01000001.1 | 344772 | 344921 |
| fig\|368602.8.peg.380 | CDS | MUIX01000001.1 | 362060 | 361650 |
| fig\|368602.8.peg.381 | CDS | MUIX01000001.1 | 362483 | 362103 |
| fig\|368602.8.peg.391 | CDS | MUIX01000001.1 | 370613 | 371110 |
| fig\|368602.8.peg.414 | CDS | MUIX01000001.1 | 392882 | 393829 |
| fig\|368602.8.peg.440 | CDS | MUIX01000001.1 | 417169 | 417318 |
| fig\|368602.8.peg.443 | CDS | MUIX01000001.1 | 419148 | 418999 |
| fig\|368602.8.peg.457 | CDS | MUIX01000001.1 | 433674 | 433264 |
| fig\|368602.8.peg.458 | CDS | MUIX01000001.1 | 434097 | 433717 |
| fig\|368602.8.peg.554 | CDS | MUIX01000001.1 | 533338 | 533033 |
| fig\|368602.8.peg.576 | CDS | MUIX01000001.1 | 560179 | 560655 |
| fig\|368602.8.peg.577 | CDS | MUIX01000001.1 | 560621 | 560740 |
| fig\|368602.8.peg.578 | CDS | MUIX01000001.1 | 560755 | 561102 |
| fig\|368602.8.peg.628 | CDS | MUIX01000001.1 | 626680 | 626829 |
| fig\|368602.8.peg.669 | CDS | MUIX01000001.1 | 667114 | 666965 |
| fig\|368602.8.peg.689 | CDS | MUIX01000001.1 | 683754 | 683617 |
| fig\|368602.8.peg.703 | CDS | MUIX01000001.1 | 696903 | 697778 |
| fig\|368602.8.peg.706 | CDS | MUIX01000001.1 | 707063 | 706191 |
| fig\|368602.8.peg.715 | CDS | MUIX01000001.1 | 719300 | 718803 |
| fig\|368602.8.peg.786 | CDS | MUIX01000001.1 | 798428 | 797919 |
| fig\|368602.8.peg.787 | CDS | MUIX01000001.1 | 798913 | 798800 |
| fig\|368602.8.peg.811 | CDS | MUIX01000001.1 | 823415 | 822069 |
| fig\|368602.8.peg.840 | CDS | MUIX01000001.1 | 851013 | 850840 |
| fig\|368602.8.peg.842 | CDS | MUIX01000001.1 | 851928 | 852041 |
| fig\|368602.8.peg.845 | CDS | MUIX01000001.1 | 852731 | 852886 |
| fig\|368602.8.peg.922 | CDS | MUIX01000001.1 | 921951 | 921784 |
| fig\|368602.8.peg.923 | CDS | MUIX01000001.1 | 922204 | 922079 |
| fig\|368602.8.peg.981 | CDS | MUIX01000001.1 | 971913 | 972062 |
| fig\|368602.8.peg.992 | CDS | MUIX01000001.1 | 982194 | 982343 |
| fig\|368602.8.peg.1005 | CDS | MUIX01000001.1 | 992272 | 991400 |
| fig\|368602.8.peg.1008 | CDS | MUIX01000001.1 | 995596 | 994250 |
| fig\|368602.8.peg.1010 | CDS | MUIX01000001.1 | 995884 | 996339 |

| | | | | | |
|---|---|---|---|---|---|
| | fig\|368602.8.peg.1011 | CDS | MUIX01000001.1 | 996348 | 996761 |
| | fig\|368602.8.peg.1012 | CDS | MUIX01000001.1 | 996866 | 996997 |
| | fig\|368602.8.peg.1054 | CDS | MUIX01000001.1 | 1035134 | 1034187 |
| | fig\|368602.8.peg.1056 | CDS | MUIX01000001.1 | 1037348 | 1036002 |
| | fig\|368602.8.peg.1060 | CDS | MUIX01000001.1 | 1040617 | 1040468 |
| | fig\|368602.8.peg.1082 | CDS | MUIX01000001.1 | 1061014 | 1061886 |
| | fig\|368602.8.peg.1161 | CDS | MUIX01000001.1 | 1159348 | 1159473 |
| | fig\|368602.8.peg.1212 | CDS | MUIX01000001.1 | 1197114 | 1196965 |
| | fig\|368602.8.peg.1245 | CDS | MUIX01000001.1 | 1223093 | 1223242 |
| | fig\|368602.8.peg.1247 | CDS | MUIX01000001.1 | 1224153 | 1225499 |
| | fig\|368602.8.peg.1292 | CDS | MUIX01000001.1 | 1273206 | 1273057 |
| | fig\|368602.8.peg.1370 | CDS | MUIX01000001.1 | 1337921 | 1337769 |
| | fig\|368602.8.peg.1388 | CDS | MUIX01000001.1 | 1367815 | 1368243 |
| | fig\|368602.8.peg.1404 | CDS | MUIX01000001.1 | 1378346 | 1378197 |
| | fig\|368602.8.peg.1410 | CDS | MUIX01000001.1 | 1382200 | 1382349 |
| | fig\|368602.8.peg.1432 | CDS | MUIX01000001.1 | 1402164 | 1402322 |
| | fig\|368602.8.peg.1492 | CDS | MUIX01000001.1 | 1461761 | 1461582 |
| | fig\|368602.8.peg.1500 | CDS | MUIX01000001.1 | 1465595 | 1466185 |
| | fig\|368602.8.peg.1509 | CDS | MUIX01000001.1 | 1477470 | 1476523 |
| | fig\|368602.8.peg.1511 | CDS | MUIX01000001.1 | 1478191 | 1478511 |
| | fig\|368602.8.peg.1512 | CDS | MUIX01000001.1 | 1478514 | 1478810 |
| | fig\|368602.8.peg.1563 | CDS | MUIX01000001.1 | 1532161 | 1532012 |
| | fig\|368602.8.peg.1578 | CDS | MUIX01000001.1 | 1548911 | 1548039 |
| | fig\|368602.8.peg.1581 | CDS | MUIX01000001.1 | 1551001 | 1550858 |
| | fig\|368602.8.peg.1650 | CDS | MUIX01000002.1 | 13813 | 13664 |
| | fig\|368602.8.peg.1662 | CDS | MUIX01000002.1 | 24502 | 25089 |
| | fig\|368602.8.peg.1676 | CDS | MUIX01000002.1 | 41983 | 41036 |
| | fig\|368602.8.peg.1684 | CDS | MUIX01000002.1 | 46876 | 45929 |
| wTpre | fig\|125593.4.peg.19 | CDS | NZ_CM003641.1 | 19000 | 19152 |
| | fig\|125593.4.peg.20 | CDS | NZ_CM003641.1 | 19459 | 19599 |
| | fig\|125593.4.peg.83 | CDS | NZ_CM003641.1 | 63818 | 63934 |
| | fig\|125593.4.peg.84 | CDS | NZ_CM003641.1 | 64010 | 64162 |
| | fig\|125593.4.peg.103 | CDS | NZ_CM003641.1 | 81741 | 81454 |
| | fig\|125593.4.peg.116 | CDS | NZ_CM003641.1 | 92046 | 92351 |
| | fig\|125593.4.peg.133 | CDS | NZ_CM003641.1 | 102732 | 102869 |
| | fig\|125593.4.peg.143 | CDS | NZ_CM003641.1 | 110438 | 110322 |
| | fig\|125593.4.peg.305 | CDS | NZ_CM003641.1 | 246336 | 246452 |
| | fig\|125593.4.peg.327 | CDS | NZ_CM003641.1 | 258863 | 259054 |
| | fig\|125593.4.peg.353 | CDS | NZ_CM003641.1 | 280927 | 281193 |
| | fig\|125593.4.peg.354 | CDS | NZ_CM003641.1 | 281316 | 281621 |
| | fig\|125593.4.peg.404 | CDS | NZ_CM003641.1 | 315522 | 315394 |
| | fig\|125593.4.peg.405 | CDS | NZ_CM003641.1 | 315730 | 315599 |

| | | | | | |
|---|---|---|---|---|---|
| | fig\|125593.4.peg.456 | CDS | NZ_CM003641.1 | 360148 | 359957 |
| | fig\|125593.4.peg.532 | CDS | NZ_CM003641.1 | 425795 | 426100 |
| | fig\|125593.4.peg.543 | CDS | NZ_CM003641.1 | 439570 | 440172 |
| | fig\|125593.4.peg.564 | CDS | NZ_CM003641.1 | 455497 | 455679 |
| | fig\|125593.4.peg.565 | CDS | NZ_CM003641.1 | 455683 | 455811 |
| | fig\|125593.4.peg.566 | CDS | NZ_CM003641.1 | 455838 | 456071 |
| | fig\|125593.4.peg.567 | CDS | NZ_CM003641.1 | 456225 | 456827 |
| | fig\|125593.4.peg.768 | CDS | NZ_CM003641.1 | 625607 | 625491 |
| | fig\|125593.4.peg.847 | CDS | NZ_CM003641.1 | 688739 | 688137 |
| | fig\|125593.4.peg.848 | CDS | NZ_CM003641.1 | 689126 | 688893 |
| | fig\|125593.4.peg.849 | CDS | NZ_CM003641.1 | 689281 | 689153 |
| | fig\|125593.4.peg.850 | CDS | NZ_CM003641.1 | 689467 | 689285 |
| | fig\|125593.4.peg.933 | CDS | NZ_CM003641.1 | 754707 | 754105 |
| | fig\|125593.4.peg.934 | CDS | NZ_CM003641.1 | 755094 | 754861 |
| | fig\|125593.4.peg.935 | CDS | NZ_CM003641.1 | 755249 | 755121 |
| | fig\|125593.4.peg.936 | CDS | NZ_CM003641.1 | 755435 | 755253 |
| | fig\|125593.4.peg.937 | CDS | NZ_CM003641.1 | 755848 | 755654 |
| | fig\|125593.4.peg.938 | CDS | NZ_CM003641.1 | 755963 | 755838 |
| | fig\|125593.4.peg.939 | CDS | NZ_CM003641.1 | 756131 | 755988 |
| | fig\|125593.4.peg.964 | CDS | NZ_CM003641.1 | 772019 | 772135 |
| | fig\|125593.4.peg.987 | CDS | NZ_CM003641.1 | 789463 | 788861 |
| | fig\|125593.4.peg.988 | CDS | NZ_CM003641.1 | 789850 | 789617 |
| | fig\|125593.4.peg.989 | CDS | NZ_CM003641.1 | 790005 | 789877 |
| | fig\|125593.4.peg.990 | CDS | NZ_CM003641.1 | 790191 | 790009 |
| | fig\|125593.4.peg.1095 | CDS | NZ_CM003641.1 | 874360 | 874542 |
| | fig\|125593.4.peg.1096 | CDS | NZ_CM003641.1 | 874546 | 874674 |
| | fig\|125593.4.peg.1097 | CDS | NZ_CM003641.1 | 874701 | 874934 |
| | fig\|125593.4.peg.1098 | CDS | NZ_CM003641.1 | 875088 | 875690 |
| | fig\|125593.4.peg.1104 | CDS | NZ_CM003641.1 | 882093 | 882209 |
| | fig\|125593.4.peg.1176 | CDS | NZ_CM003641.1 | 938835 | 938692 |
| | fig\|125593.4.peg.1192 | CDS | NZ_CM003641.1 | 952419 | 952276 |
| | fig\|125593.4.peg.1193 | CDS | NZ_CM003641.1 | 952656 | 952498 |
| | fig\|125593.4.peg.1194 | CDS | NZ_CM003641.1 | 952864 | 952733 |
| | fig\|125593.4.peg.1298 | CDS | NZ_CM003641.1 | 1057695 | 1057877 |
| | fig\|125593.4.peg.1299 | CDS | NZ_CM003641.1 | 1057881 | 1058009 |
| | fig\|125593.4.peg.1300 | CDS | NZ_CM003641.1 | 1058036 | 1058269 |
| | fig\|125593.4.peg.1301 | CDS | NZ_CM003641.1 | 1058423 | 1059025 |
| | fig\|125593.4.peg.1360 | CDS | NZ_CM003641.1 | 1110505 | 1110347 |
| wVulC | fig\|1220511.6.peg.1 | CDS | ALWU01000001.1 | 486 | 229 |
| | fig\|1220511.6.peg.2 | CDS | ALWU01000001.1 | 719 | 555 |
| | fig\|1220511.6.peg.9 | CDS | ALWU01000001.1 | 6218 | 6790 |
| | fig\|1220511.6.peg.10 | CDS | ALWU01000001.1 | 7059 | 7457 |

| | | | | |
|---|---|---|---|---|
| fig\|1220511.6.peg.11 | CDS | ALWU01000001.1 | 7489 | 7695 |
| fig\|1220511.6.peg.12 | CDS | ALWU01000001.1 | 7703 | 8392 |
| fig\|1220511.6.peg.17 | CDS | ALWU01000001.1 | 11853 | 13052 |
| fig\|1220511.6.peg.20 | CDS | ALWU01000001.1 | 15013 | 14717 |
| fig\|1220511.6.peg.103 | CDS | ALWU01000001.1 | 101350 | 101787 |
| fig\|1220511.6.peg.104 | CDS | ALWU01000001.1 | 101835 | 102008 |
| fig\|1220511.6.peg.121 | CDS | ALWU01000001.1 | 119941 | 119159 |
| fig\|1220511.6.peg.122 | CDS | ALWU01000001.1 | 120201 | 120010 |
| fig\|1220511.6.peg.125 | CDS | ALWU01000001.1 | 122714 | 122956 |
| fig\|1220511.6.peg.126 | CDS | ALWU01000001.1 | 123164 | 123298 |
| fig\|1220511.6.peg.127 | CDS | ALWU01000001.1 | 123592 | 123723 |
| fig\|1220511.6.peg.130 | CDS | ALWU01000001.1 | 125723 | 125331 |
| fig\|1220511.6.peg.131 | CDS | ALWU01000001.1 | 125959 | 125828 |
| fig\|1220511.6.peg.132 | CDS | ALWU01000001.1 | 126387 | 126253 |
| fig\|1220511.6.peg.133 | CDS | ALWU01000001.1 | 126711 | 126595 |
| fig\|1220511.6.peg.143 | CDS | ALWU01000001.1 | 131924 | 132277 |
| fig\|1220511.6.peg.144 | CDS | ALWU01000001.1 | 132473 | 132592 |
| fig\|1220511.6.peg.148 | CDS | ALWU01000001.1 | 133518 | 133796 |
| fig\|1220511.6.peg.155 | CDS | ALWU01000001.1 | 140437 | 140751 |
| fig\|1220511.6.peg.156 | CDS | ALWU01000001.1 | 140755 | 140922 |
| fig\|1220511.6.peg.209 | CDS | ALWU01000001.1 | 192037 | 191348 |
| fig\|1220511.6.peg.211 | CDS | ALWU01000001.1 | 192719 | 192588 |
| fig\|1220511.6.peg.212 | CDS | ALWU01000001.1 | 193147 | 193013 |
| fig\|1220511.6.peg.213 | CDS | ALWU01000001.1 | 193597 | 193355 |
| fig\|1220511.6.peg.217 | CDS | ALWU01000001.1 | 195487 | 195747 |
| fig\|1220511.6.peg.218 | CDS | ALWU01000001.1 | 196019 | 196246 |
| fig\|1220511.6.peg.219 | CDS | ALWU01000001.1 | 196338 | 196508 |
| fig\|1220511.6.peg.233 | CDS | ALWU01000001.1 | 207218 | 207084 |
| fig\|1220511.6.peg.234 | CDS | ALWU01000001.1 | 207668 | 207426 |
| fig\|1220511.6.peg.238 | CDS | ALWU01000001.1 | 210025 | 209894 |
| fig\|1220511.6.peg.239 | CDS | ALWU01000001.1 | 210453 | 210319 |
| fig\|1220511.6.peg.240 | CDS | ALWU01000001.1 | 210903 | 210661 |
| fig\|1220511.6.peg.266 | CDS | ALWU01000001.1 | 233597 | 233767 |
| fig\|1220511.6.peg.267 | CDS | ALWU01000001.1 | 234108 | 234236 |
| fig\|1220511.6.peg.285 | CDS | ALWU01000001.1 | 254207 | 254013 |
| fig\|1220511.6.peg.293 | CDS | ALWU01000001.1 | 256508 | 256822 |
| fig\|1220511.6.peg.294 | CDS | ALWU01000001.1 | 256779 | 257138 |
| fig\|1220511.6.peg.305 | CDS | ALWU01000001.1 | 266087 | 266440 |
| fig\|1220511.6.peg.319 | CDS | ALWU01000001.1 | 279147 | 279266 |
| fig\|1220511.6.peg.320 | CDS | ALWU01000001.1 | 279325 | 279447 |
| fig\|1220511.6.peg.321 | CDS | ALWU01000001.1 | 280487 | 279855 |
| fig\|1220511.6.peg.322 | CDS | ALWU01000001.1 | 281443 | 281036 |

| | | | | |
|---|---|---|---|---|
| fig\|1220511.6.peg.323 | CDS | ALWU01000001.1 | 282118 | 281828 |
| fig\|1220511.6.peg.325 | CDS | ALWU01000001.1 | 283228 | 283019 |
| fig\|1220511.6.peg.330 | CDS | ALWU01000001.1 | 287291 | 287196 |
| fig\|1220511.6.peg.350 | CDS | ALWU01000001.1 | 303646 | 304131 |
| fig\|1220511.6.peg.351 | CDS | ALWU01000001.1 | 304283 | 304438 |
| fig\|1220511.6.peg.356 | CDS | ALWU01000001.1 | 307442 | 308194 |
| fig\|1220511.6.peg.358 | CDS | ALWU01000001.1 | 309528 | 309839 |
| fig\|1220511.6.peg.359 | CDS | ALWU01000001.1 | 310346 | 309861 |
| fig\|1220511.6.peg.374 | CDS | ALWU01000001.1 | 320649 | 320807 |
| fig\|1220511.6.peg.375 | CDS | ALWU01000001.1 | 320768 | 320932 |
| fig\|1220511.6.peg.376 | CDS | ALWU01000001.1 | 321314 | 321550 |
| fig\|1220511.6.peg.378 | CDS | ALWU01000001.1 | 321911 | 322090 |
| fig\|1220511.6.peg.381 | CDS | ALWU01000001.1 | 324347 | 324514 |
| fig\|1220511.6.peg.400 | CDS | ALWU01000001.1 | 347078 | 346962 |
| fig\|1220511.6.peg.401 | CDS | ALWU01000001.1 | 347456 | 347331 |
| fig\|1220511.6.peg.402 | CDS | ALWU01000001.1 | 347609 | 347490 |
| fig\|1220511.6.peg.489 | CDS | ALWU01000001.1 | 418396 | 418241 |
| fig\|1220511.6.peg.490 | CDS | ALWU01000001.1 | 418520 | 418362 |
| fig\|1220511.6.peg.491 | CDS | ALWU01000001.1 | 418812 | 418621 |
| fig\|1220511.6.peg.492 | CDS | ALWU01000001.1 | 419048 | 418893 |
| fig\|1220511.6.peg.493 | CDS | ALWU01000001.1 | 419421 | 419302 |
| fig\|1220511.6.peg.513 | CDS | ALWU01000001.1 | 433083 | 432952 |
| fig\|1220511.6.peg.514 | CDS | ALWU01000001.1 | 433511 | 433377 |
| fig\|1220511.6.peg.515 | CDS | ALWU01000001.1 | 433982 | 433719 |
| fig\|1220511.6.peg.523 | CDS | ALWU01000001.1 | 441991 | 442206 |
| fig\|1220511.6.peg.524 | CDS | ALWU01000001.1 | 442250 | 442564 |
| fig\|1220511.6.peg.525 | CDS | ALWU01000001.1 | 442569 | 442751 |
| fig\|1220511.6.peg.528 | CDS | ALWU01000001.1 | 445242 | 445114 |
| fig\|1220511.6.peg.530 | CDS | ALWU01000001.1 | 445814 | 445677 |
| fig\|1220511.6.peg.542 | CDS | ALWU01000001.1 | 455260 | 455123 |
| fig\|1220511.6.peg.543 | CDS | ALWU01000001.1 | 455685 | 455530 |
| fig\|1220511.6.peg.544 | CDS | ALWU01000001.1 | 455849 | 455673 |
| fig\|1220511.6.peg.555 | CDS | ALWU01000001.1 | 467015 | 466848 |
| fig\|1220511.6.peg.556 | CDS | ALWU01000001.1 | 467491 | 467357 |
| fig\|1220511.6.peg.557 | CDS | ALWU01000001.1 | 468345 | 467593 |
| fig\|1220511.6.peg.564 | CDS | ALWU01000001.1 | 476088 | 475399 |
| fig\|1220511.6.peg.565 | CDS | ALWU01000001.1 | 476290 | 476096 |
| fig\|1220511.6.peg.570 | CDS | ALWU01000001.1 | 482031 | 481900 |
| fig\|1220511.6.peg.571 | CDS | ALWU01000001.1 | 482459 | 482325 |
| fig\|1220511.6.peg.572 | CDS | ALWU01000001.1 | 482909 | 482667 |
| fig\|1220511.6.peg.600 | CDS | ALWU01000001.1 | 513153 | 512971 |
| fig\|1220511.6.peg.601 | CDS | ALWU01000001.1 | 513472 | 513158 |

| | | | | |
|---|---|---|---|---|
| fig\|1220511.6.peg.602 | CDS | ALWU01000001.1 | 513731 | 513516 |
| fig\|1220511.6.peg.608 | CDS | ALWU01000001.1 | 518428 | 518201 |
| fig\|1220511.6.peg.609 | CDS | ALWU01000001.1 | 518960 | 518700 |
| fig\|1220511.6.peg.610 | CDS | ALWU01000001.1 | 519153 | 518968 |
| fig\|1220511.6.peg.625 | CDS | ALWU01000001.1 | 536523 | 536729 |
| fig\|1220511.6.peg.626 | CDS | ALWU01000001.1 | 536737 | 537426 |
| fig\|1220511.6.peg.628 | CDS | ALWU01000001.1 | 539346 | 539149 |
| fig\|1220511.6.peg.629 | CDS | ALWU01000001.1 | 539530 | 539417 |
| fig\|1220511.6.peg.640 | CDS | ALWU01000001.1 | 551851 | 552099 |
| fig\|1220511.6.peg.641 | CDS | ALWU01000001.1 | 552254 | 552457 |
| fig\|1220511.6.peg.652 | CDS | ALWU01000001.1 | 562995 | 562864 |
| fig\|1220511.6.peg.653 | CDS | ALWU01000001.1 | 563423 | 563289 |
| fig\|1220511.6.peg.654 | CDS | ALWU01000001.1 | 563873 | 563631 |
| fig\|1220511.6.peg.666 | CDS | ALWU01000001.1 | 572408 | 572268 |
| fig\|1220511.6.peg.671 | CDS | ALWU01000001.1 | 576644 | 576159 |
| fig\|1220511.6.peg.675 | CDS | ALWU01000001.1 | 578067 | 578180 |
| fig\|1220511.6.peg.676 | CDS | ALWU01000001.1 | 578370 | 578242 |
| fig\|1220511.6.peg.687 | CDS | ALWU01000002.1 | 3779 | 3384 |
| fig\|1220511.6.peg.733 | CDS | ALWU01000003.1 | 899 | 117 |
| fig\|1220511.6.peg.734 | CDS | ALWU01000003.1 | 1159 | 968 |
| fig\|1220511.6.peg.738 | CDS | ALWU01000003.1 | 4170 | 3943 |
| fig\|1220511.6.peg.739 | CDS | ALWU01000003.1 | 4597 | 4442 |
| fig\|1220511.6.peg.740 | CDS | ALWU01000003.1 | 4761 | 4585 |
| fig\|1220511.6.peg.742 | CDS | ALWU01000003.1 | 5252 | 5025 |
| fig\|1220511.6.peg.743 | CDS | ALWU01000003.1 | 5784 | 5524 |
| fig\|1220511.6.peg.744 | CDS | ALWU01000003.1 | 6031 | 5792 |
| fig\|1220511.6.peg.752 | CDS | ALWU01000003.1 | 13094 | 13342 |
| fig\|1220511.6.peg.753 | CDS | ALWU01000003.1 | 13497 | 13700 |
| fig\|1220511.6.peg.758 | CDS | ALWU01000003.1 | 18497 | 17298 |
| fig\|1220511.6.peg.765 | CDS | ALWU01000003.1 | 25105 | 25221 |
| fig\|1220511.6.peg.766 | CDS | ALWU01000003.1 | 25429 | 25563 |
| fig\|1220511.6.peg.767 | CDS | ALWU01000003.1 | 25857 | 25988 |
| fig\|1220511.6.peg.768 | CDS | ALWU01000003.1 | 26394 | 26059 |
| fig\|1220511.6.peg.773 | CDS | ALWU01000003.1 | 28454 | 29158 |
| fig\|1220511.6.peg.817 | CDS | ALWU01000005.1 | 1003 | 17 |
| fig\|1220511.6.peg.819 | CDS | ALWU01000005.1 | 1437 | 1589 |
| fig\|1220511.6.peg.820 | CDS | ALWU01000005.1 | 1543 | 1719 |
| fig\|1220511.6.peg.822 | CDS | ALWU01000005.1 | 2709 | 2915 |
| fig\|1220511.6.peg.823 | CDS | ALWU01000005.1 | 2923 | 3612 |
| fig\|1220511.6.peg.853 | CDS | ALWU01000005.1 | 35459 | 34677 |
| fig\|1220511.6.peg.854 | CDS | ALWU01000005.1 | 35719 | 35528 |
| fig\|1220511.6.peg.888 | CDS | ALWU01000005.1 | 62018 | 61236 |

| | | | | |
|---|---|---|---|---|
| fig\|1220511.6.peg.889 | CDS | ALWU01000005.1 | 62278 | 62087 |
| fig\|1220511.6.peg.890 | CDS | ALWU01000005.1 | 63679 | 62495 |
| fig\|1220511.6.peg.899 | CDS | ALWU01000005.1 | 74512 | 73760 |
| fig\|1220511.6.peg.901 | CDS | ALWU01000005.1 | 75096 | 75230 |
| fig\|1220511.6.peg.902 | CDS | ALWU01000005.1 | 75524 | 75655 |
| fig\|1220511.6.peg.903 | CDS | ALWU01000005.1 | 76284 | 75850 |
| fig\|1220511.6.peg.944 | CDS | ALWU01000005.1 | 111706 | 111897 |
| fig\|1220511.6.peg.945 | CDS | ALWU01000005.1 | 111966 | 112748 |
| fig\|1220511.6.peg.948 | CDS | ALWU01000005.1 | 114600 | 113848 |
| fig\|1220511.6.peg.955 | CDS | ALWU01000005.1 | 120253 | 119471 |
| fig\|1220511.6.peg.956 | CDS | ALWU01000005.1 | 120513 | 120322 |
| fig\|1220511.6.peg.967 | CDS | ALWU01000005.1 | 132323 | 132192 |
| fig\|1220511.6.peg.968 | CDS | ALWU01000005.1 | 132751 | 132617 |
| fig\|1220511.6.peg.969 | CDS | ALWU01000005.1 | 133075 | 132959 |
| fig\|1220511.6.peg.971 | CDS | ALWU01000005.1 | 133903 | 133739 |
| fig\|1220511.6.peg.972 | CDS | ALWU01000005.1 | 134204 | 133908 |
| fig\|1220511.6.peg.983 | CDS | ALWU01000005.1 | 146558 | 146677 |
| fig\|1220511.6.peg.990 | CDS | ALWU01000005.1 | 152037 | 151783 |
| fig\|1220511.6.peg.997 | CDS | ALWU01000005.1 | 157573 | 157836 |
| fig\|1220511.6.peg.998 | CDS | ALWU01000005.1 | 158044 | 158178 |
| fig\|1220511.6.peg.999 | CDS | ALWU01000005.1 | 158472 | 158603 |
| fig\|1220511.6.peg.1002 | CDS | ALWU01000005.1 | 159420 | 160052 |
| fig\|1220511.6.peg.1015 | CDS | ALWU01000005.1 | 175722 | 174523 |
| fig\|1220511.6.peg.1016 | CDS | ALWU01000005.1 | 176263 | 176072 |
| fig\|1220511.6.peg.1017 | CDS | ALWU01000005.1 | 176970 | 176311 |
| fig\|1220511.6.peg.1033 | CDS | ALWU01000005.1 | 186831 | 187583 |
| fig\|1220511.6.peg.1035 | CDS | ALWU01000005.1 | 189109 | 188660 |
| fig\|1220511.6.peg.1037 | CDS | ALWU01000005.1 | 189422 | 189574 |
| fig\|1220511.6.peg.1044 | CDS | ALWU01000005.1 | 193794 | 193516 |
| fig\|1220511.6.peg.1045 | CDS | ALWU01000005.1 | 194091 | 194297 |
| fig\|1220511.6.peg.1046 | CDS | ALWU01000005.1 | 194305 | 194994 |
| fig\|1220511.6.peg.1090 | CDS | ALWU01000006.1 | 13400 | 13876 |
| fig\|1220511.6.peg.1093 | CDS | ALWU01000006.1 | 15257 | 15448 |
| fig\|1220511.6.peg.1094 | CDS | ALWU01000006.1 | 15517 | 16299 |
| fig\|1220511.6.peg.1102 | CDS | ALWU01000006.1 | 26817 | 26906 |
| fig\|1220511.6.peg.1103 | CDS | ALWU01000006.1 | 26916 | 27140 |
| fig\|1220511.6.peg.1104 | CDS | ALWU01000006.1 | 27172 | 27333 |
| fig\|1220511.6.peg.1105 | CDS | ALWU01000006.1 | 27405 | 27632 |
| fig\|1220511.6.peg.1106 | CDS | ALWU01000006.1 | 27625 | 27930 |
| fig\|1220511.6.peg.1109 | CDS | ALWU01000006.1 | 30985 | 30290 |
| fig\|1220511.6.peg.1117 | CDS | ALWU01000006.1 | 34414 | 35013 |
| fig\|1220511.6.peg.1143 | CDS | ALWU01000008.1 | 8373 | 7591 |

| | | | | |
|---|---|---|---|---|
| fig\|1220511.6.peg.1144 | CDS | ALWU01000008.1 | 8633 | 8442 |
| fig\|1220511.6.peg.1177 | CDS | ALWU01000008.1 | 38367 | 38095 |
| fig\|1220511.6.peg.1213 | CDS | ALWU01000008.1 | 76751 | 76461 |
| fig\|1220511.6.peg.1215 | CDS | ALWU01000008.1 | 77819 | 78367 |
| fig\|1220511.6.peg.1216 | CDS | ALWU01000008.1 | 78507 | 78749 |
| fig\|1220511.6.peg.1217 | CDS | ALWU01000008.1 | 78957 | 79091 |
| fig\|1220511.6.peg.1227 | CDS | ALWU01000008.1 | 86923 | 87129 |
| fig\|1220511.6.peg.1228 | CDS | ALWU01000008.1 | 87137 | 87826 |
| fig\|1220511.6.peg.1229 | CDS | ALWU01000008.1 | 88296 | 88120 |
| fig\|1220511.6.peg.1258 | CDS | ALWU01000008.1 | 121359 | 120919 |
| fig\|1220511.6.peg.1259 | CDS | ALWU01000008.1 | 121573 | 121367 |
| fig\|1220511.6.peg.1278 | CDS | ALWU01000008.1 | 137147 | 137338 |
| fig\|1220511.6.peg.1279 | CDS | ALWU01000008.1 | 137407 | 137895 |
| fig\|1220511.6.peg.1280 | CDS | ALWU01000008.1 | 137853 | 138188 |
| fig\|1220511.6.peg.1316 | CDS | ALWU01000008.1 | 169494 | 169913 |
| fig\|1220511.6.peg.1318 | CDS | ALWU01000008.1 | 170793 | 170659 |
| fig\|1220511.6.peg.1319 | CDS | ALWU01000008.1 | 171117 | 171001 |
| fig\|1220511.6.peg.1329 | CDS | ALWU01000008.1 | 176330 | 176683 |
| fig\|1220511.6.peg.1330 | CDS | ALWU01000008.1 | 176879 | 176998 |
| fig\|1220511.6.peg.1334 | CDS | ALWU01000008.1 | 177924 | 178202 |
| fig\|1220511.6.peg.1359 | CDS | ALWU01000010.1 | 2644 | 3129 |
| fig\|1220511.6.peg.1364 | CDS | ALWU01000010.1 | 7529 | 7329 |
| fig\|1220511.6.peg.1365 | CDS | ALWU01000010.1 | 7795 | 8004 |
| fig\|1220511.6.peg.1367 | CDS | ALWU01000010.1 | 8905 | 9195 |
| fig\|1220511.6.peg.1369 | CDS | ALWU01000010.1 | 9558 | 9788 |
| fig\|1220511.6.peg.1370 | CDS | ALWU01000010.1 | 9785 | 10522 |
| fig\|1220511.6.peg.1375 | CDS | ALWU01000010.1 | 16761 | 16952 |
| fig\|1220511.6.peg.1376 | CDS | ALWU01000010.1 | 17021 | 17803 |
| fig\|1220511.6.peg.1408 | CDS | ALWU01000010.1 | 50302 | 50051 |
| fig\|1220511.6.peg.1409 | CDS | ALWU01000010.1 | 51024 | 50620 |
| fig\|1220511.6.peg.1410 | CDS | ALWU01000010.1 | 51239 | 51054 |
| fig\|1220511.6.peg.1422 | CDS | ALWU01000010.1 | 59385 | 58696 |
| fig\|1220511.6.peg.1423 | CDS | ALWU01000010.1 | 59599 | 59393 |
| fig\|1220511.6.peg.1424 | CDS | ALWU01000010.1 | 60029 | 59631 |
| fig\|1220511.6.peg.1425 | CDS | ALWU01000010.1 | 60870 | 60298 |
| fig\|1220511.6.peg.1427 | CDS | ALWU01000010.1 | 61511 | 61675 |
| fig\|1220511.6.peg.1429 | CDS | ALWU01000010.1 | 61960 | 62106 |
| fig\|1220511.6.peg.1432 | CDS | ALWU01000010.1 | 64820 | 65329 |
| fig\|1220511.6.peg.1464 | CDS | ALWU01000010.1 | 96463 | 96615 |
| fig\|1220511.6.peg.1496 | CDS | ALWU01000010.1 | 124352 | 124543 |
| fig\|1220511.6.peg.1497 | CDS | ALWU01000010.1 | 124612 | 125394 |
| fig\|1220511.6.peg.1498 | CDS | ALWU01000010.1 | 125681 | 125517 |

| | | | | |
|---|---|---|---|---|
| fig\|1220511.6.peg.1519 | CDS | ALWU01000010.1 | 139281 | 139487 |
| fig\|1220511.6.peg.1520 | CDS | ALWU01000010.1 | 139495 | 140184 |
| fig\|1220511.6.peg.1532 | CDS | ALWU01000010.1 | 148970 | 149722 |
| fig\|1220511.6.peg.1533 | CDS | ALWU01000010.1 | 149966 | 150085 |
| fig\|1220511.6.peg.1545 | CDS | ALWU01000010.1 | 159626 | 159480 |
| fig\|1220511.6.peg.1547 | CDS | ALWU01000010.1 | 160075 | 159911 |
| fig\|1220511.6.peg.1550 | CDS | ALWU01000010.1 | 161910 | 162047 |
| fig\|1220511.6.peg.1576 | CDS | ALWU01000010.1 | 188072 | 188263 |
| fig\|1220511.6.peg.1577 | CDS | ALWU01000010.1 | 188332 | 189114 |
| fig\|1220511.6.peg.1591 | CDS | ALWU01000010.1 | 198550 | 198663 |
| fig\|1220511.6.peg.1592 | CDS | ALWU01000010.1 | 198734 | 198931 |
| fig\|1220511.6.peg.1597 | CDS | ALWU01000010.1 | 204452 | 204234 |
| fig\|1220511.6.peg.1598 | CDS | ALWU01000010.1 | 204786 | 204487 |
| fig\|1220511.6.peg.1606 | CDS | ALWU01000010.1 | 210161 | 211360 |
| fig\|1220511.6.peg.1621 | CDS | ALWU01000010.1 | 222489 | 222013 |
| fig\|1220511.6.peg.1635 | CDS | ALWU01000010.1 | 231331 | 230378 |
| fig\|1220511.6.peg.1636 | CDS | ALWU01000010.1 | 231643 | 231461 |
| fig\|1220511.6.peg.1697 | CDS | ALWU01000010.1 | 292023 | 290779 |
| fig\|1220511.6.peg.1711 | CDS | ALWU01000010.1 | 305934 | 305152 |
| fig\|1220511.6.peg.1712 | CDS | ALWU01000010.1 | 306194 | 306003 |
| fig\|1220511.6.peg.1727 | CDS | ALWU01000010.1 | 315542 | 316051 |
| fig\|1220511.6.peg.1737 | CDS | ALWU01000010.1 | 323950 | 323738 |
| fig\|1220511.6.peg.1738 | CDS | ALWU01000010.1 | 324441 | 325640 |
| fig\|1220511.6.peg.1741 | CDS | ALWU01000010.1 | 326276 | 326464 |
| fig\|1220511.6.peg.1761 | CDS | ALWU01000010.1 | 350833 | 350714 |
| fig\|1220511.6.peg.1781 | CDS | ALWU01000010.1 | 364925 | 364806 |
| fig\|1220511.6.peg.1782 | CDS | ALWU01000010.1 | 365057 | 364935 |
| fig\|1220511.6.peg.1783 | CDS | ALWU01000010.1 | 365483 | 365079 |
| fig\|1220511.6.peg.1784 | CDS | ALWU01000010.1 | 365660 | 365755 |
| fig\|1220511.6.peg.1788 | CDS | ALWU01000010.1 | 369184 | 369321 |
| fig\|1220511.6.peg.1791 | CDS | ALWU01000010.1 | 370529 | 370681 |
| fig\|1220511.6.peg.1793 | CDS | ALWU01000010.1 | 371696 | 371986 |
| fig\|1220511.6.peg.1794 | CDS | ALWU01000010.1 | 372932 | 372798 |
| fig\|1220511.6.peg.1795 | CDS | ALWU01000010.1 | 373382 | 373140 |
| fig\|1220511.6.peg.1802 | CDS | ALWU01000010.1 | 378247 | 378507 |
| fig\|1220511.6.peg.1803 | CDS | ALWU01000010.1 | 378779 | 379006 |
| fig\|1220511.6.peg.1804 | CDS | ALWU01000010.1 | 379098 | 379268 |
| fig\|1220511.6.peg.1874 | CDS | ALWU01000010.1 | 442603 | 442469 |
| fig\|1220511.6.peg.1911 | CDS | ALWU01000010.1 | 478879 | 478097 |
| fig\|1220511.6.peg.1912 | CDS | ALWU01000010.1 | 479139 | 478948 |
| fig\|1220511.6.peg.1931 | CDS | ALWU01000010.1 | 502449 | 502096 |
| fig\|1220511.6.peg.1935 | CDS | ALWU01000010.1 | 505265 | 505119 |

| | | | | | |
|---|---|---|---|---|---|
| | fig\|1220511.6.peg.1936 | CDS | ALWU01000010.1 | 505539 | 505399 |
| wWb | fig\|96496.9.peg.245 | CDS | NZ_NJBR0200001 | 3174 | 2374 |
| | fig\|96496.9.peg.655 | CDS | NZ_NJBR0200004 | 78 | 320 |
| | fig\|96496.9.peg.656 | CDS | NZ_NJBR0200004 | 997 | 1215 |
| | fig\|96496.9.peg.681 | CDS | NZ_NJBR0200005 | 3735 | 4562 |
| | fig\|96496.9.peg.832 | CDS | NZ_NJBR0200006 | 17486 | 17124 |
| | fig\|96496.9.peg.835 | CDS | NZ_NJBR0200007 | 1091 | 693 |
| | fig\|96496.9.peg.895 | CDS | NZ_NJBR0200007 | 20475 | 21638 |
| | fig\|96496.9.peg.1056 | CDS | NZ_NJBR0200009 | 2858 | 2995 |

| Strand | Length (bp) | Function |
|---|---|---|
| - | 273 | Mobile element protein |
| - | 189 | Mobile element protein |
| + | 159 | Mobile element protein |
| + | 273 | Mobile element protein |
| - | 273 | Mobile element protein |
| - | 159 | Mobile element protein |
| + | 273 | Mobile element protein |
| - | 162 | Mobile element protein |
| + | 273 | Mobile element protein |
| + | 273 | Mobile element protein |
| - | - | - |
| + | 285 | Mobile element protein |
| + | 129 | Mobile element protein |
| + | 366 | Mobile element protein |
| + | 366 | Mobile element protein |
| + | 366 | Mobile element protein |
| - | 189 | Mobile element protein |
| - | 204 | Mobile element protein |
| - | 366 | Mobile element protein |
| + | 330 | Mobile element protein |
| + | 636 | Mobile element protein |
| + | 270 | Mobile element protein |
| + | 366 | Mobile element protein |
| - | 636 | Mobile element protein |
| - | 330 | Mobile element protein |
| - | 366 | Mobile element protein |
| - | 366 | Mobile element protein |
| - | 366 | Mobile element protein |
| - | 333 | Mobile element protein |
| - | 366 | Mobile element protein |
| + | 306 | Mobile element protein |
| - | 366 | Mobile element protein |
| + | 366 | Mobile element protein |
| + | 315 | Mobile element protein |
| + | 954 | Mobile element protein |
| + | 375 | Mobile element protein |
| + | 669 | Mobile element protein |
| + | 129 | Mobile element protein |
| - | 366 | Mobile element protein |

| | | |
|---|---|---|
| - | 366 | Mobile element protein |
| - | 102 | Mobile element protein |
| - | 213 | Mobile element protein |
| - | 669 | Mobile element protein |
| - | 366 | Mobile element protein |
| - | 165 | Mobile element protein |
| - | 303 | Mobile element protein |
| - | 366 | Mobile element protein |
| - | 366 | Mobile element protein |
| + | 366 | Mobile element protein |
| + | 366 | Mobile element protein |
| - | 366 | Mobile element protein |
| + | 303 | Mobile element protein |
| + | 165 | Mobile element protein |
| + | 429 | Mobile element protein |
| - | 189 | Mobile element protein |
| + | 948 | Mobile element protein |
| + | 366 | Mobile element protein |
| + | 366 | Mobile element protein |
| + | 285 | Mobile element protein |
| - | 366 | Mobile element protein |
| + | 294 | Mobile element protein |
| + | 165 | Mobile element protein |
| + | 366 | Mobile element protein |
| - | 222 | Mobile element protein |
| - | 366 | Mobile element protein |
| - | 261 | Mobile element protein |
| + | 366 | Mobile element protein |
| - | 366 | Mobile element protein |
| - | 366 | Mobile element protein |
| - | 1155 | Mobile element protein |
| - | 285 | Mobile element protein |
| + | 366 | Mobile element protein |
| + | 366 | Mobile element protein |
| - | 129 | Mobile element protein |
| + | 366 | Mobile element protein |
| - | 516 | Mobile element protein |
| - | 252 | Mobile element protein |
| + | 285 | Mobile element protein |
| + | 366 | Mobile element protein |
| + | 153 | Mobile element protein |
| + | 285 | Mobile element protein |

| | | |
|---|---|---|
| - | 408 | Mobile element protein |
| - | 201 | Mobile element protein |
| - | 330 | Mobile element protein |
| + | 366 | Mobile element protein |
| - | 366 | Mobile element protein |
| - | 366 | Mobile element protein |
| - | 285 | Mobile element protein |
| + | 297 | Mobile element protein |
| + | 366 | Mobile element protein |
| - | 366 | Mobile element protein |
| + | 366 | Mobile element protein |
| + | 270 | Mobile element protein |
| + | 366 | Mobile element protein |
| + | 366 | Mobile element protein |
| + | 303 | Mobile element protein |
| + | 165 | Mobile element protein |
| + | 366 | Mobile element protein |
| - | 1344 | Mobile element protein |
| + | 366 | Mobile element protein |
| + | 294 | Mobile element protein |
| + | 252 | Mobile element protein |
| - | 366 | Mobile element protein |
| - | 243 | Mobile element protein |
| + | 366 | Mobile element protein |
| - | 366 | Mobile element protein |
| - | 234 | Mobile element protein |
| + | 366 | Mobile element protein |
| + | 306 | Mobile element protein |
| - | 129 | Mobile element protein |
| - | 285 | Mobile element protein |
| - | 129 | Mobile element protein |
| - | 285 | Mobile element protein |
| - | 366 | Mobile element protein |
| + | 366 | Mobile element protein |
| - | 636 | Mobile element protein |
| - | 330 | Mobile element protein |
| - | 366 | Mobile element protein |
| + | 285 | Mobile element protein |
| + | 1155 | Mobile element protein |
| - | 636 | Mobile element protein |
| - | 300 | Mobile element protein |
| - | 297 | Mobile element protein |

| | | |
|---|---|---|
| + | 174 | Mobile element protein |
| + | 312 | Mobile element protein |
| + | 183 | Mobile element protein |
| + | 195 | Mobile element protein |
| + | 366 | Mobile element protein |
| + | 429 | Mobile element protein |
| - | 189 | Mobile element protein |
| + | 366 | Mobile element protein |
| - | 408 | Mobile element protein |
| - | 201 | Mobile element protein |
| - | 330 | Mobile element protein |
| - | 366 | Mobile element protein |
| - | 429 | Mobile element protein |
| - | 195 | Mobile element protein |
| - | 183 | Mobile element protein |
| + | 366 | Mobile element protein |
| + | 207 | Mobile element protein |
| + | 384 | Mobile element protein |
| - | 348 | Mobile element protein |
| - | 189 | Mobile element protein |
| + | 300 | Mobile element protein |
| + | 636 | Mobile element protein |
| - | 366 | Mobile element protein |
| - | 636 | Mobile element protein |
| - | 330 | Mobile element protein |
| - | 366 | Mobile element protein |
| - | 870 | Mobile element protein |
| + | 954 | Mobile element protein |
| + | 246 | Mobile element protein |
| + | 189 | Mobile element protein |
| + | 366 | Mobile element protein |
| + | 150 | Mobile element protein |
| + | 300 | Mobile element protein |
| + | 168 | Mobile element protein |
| + | 648 | Mobile element protein |
| + | 261 | Mobile element protein |
| + | 417 | Mobile element protein |
| - | 393 | Mobile element protein |
| - | 1362 | Mobile element protein |
| - | 828 | Mobile element protein |
| - | 213 | Mobile element protein |
| - | 957 | Mobile element protein |

| | | |
|---|---|---|
| + | 231 | Mobile element protein |
| - | 1341 | Mobile element protein |
| - | 1341 | Mobile element protein |
| + | 372 | Mobile element protein |
| + | 828 | Mobile element protein |
| - | 828 | Mobile element protein |
| - | 372 | Mobile element protein |
| - | 870 | Mobile element protein |
| - | 870 | Mobile element protein |
| - | 870 | Mobile element protein |
| + | 612 | Mobile element protein |
| - | 957 | Mobile element protein |
| - | 372 | Mobile element protein |
| + | 1362 | Mobile element protein |
| + | 804 | Mobile element protein |
| + | 171 | Mobile element protein |
| + | 1341 | Mobile element protein |
| - | 957 | Mobile element protein |
| + | 1341 | Mobile element protein |
| - | 1341 | Mobile element protein |
| - | 1341 | Mobile element protein |
| - | 1362 | Mobile element protein |
| - | 1362 | Mobile element protein |
| - | 957 | Mobile element protein |
| + | 1341 | Mobile element protein |
| - | 927 | Mobile element protein |
| - | 957 | Mobile element protein |
| - | 1341 | Mobile element protein |
| + | 957 | Mobile element protein |
| - | 957 | Mobile element protein |
| + | 372 | Mobile element protein |
| + | 828 | Mobile element protein |
| + | 1341 | Mobile element protein |
| + | 1341 | Mobile element protein |
| - | 957 | Mobile element protein |
| - | 957 | Mobile element protein |
| - | 1341 | Mobile element protein |
| - | 159 | Mobile element protein |
| + | 372 | Mobile element protein |
| + | 756 | Mobile element protein |
| - | 1362 | Mobile element protein |
| + | 957 | Mobile element protein |

| | | |
|---|---|---|
| + | 1341 | Mobile element protein |
| + | 957 | Mobile element protein |
| - | 1056 | Mobile element protein |
| - | 1341 | Mobile element protein |
| - | 804 | Mobile element protein |
| - | 372 | Mobile element protein |
| + | 1341 | Mobile element protein |
| - | 201 | Mobile element protein |
| - | 1362 | Mobile element protein |
| - | 915 | Mobile element protein |
| + | 1341 | Mobile element protein |
| - | 1341 | Mobile element protein |
| + | 957 | Mobile element protein |
| + | 957 | Mobile element protein |
| - | 1341 | Mobile element protein |
| - | 1317 | Mobile element protein |
| - | 828 | Mobile element protein |
| - | 372 | Mobile element protein |
| - | 957 | Mobile element protein |
| - | 957 | Mobile element protein |
| + | 1341 | Mobile element protein |
| - | 675 | Mobile element protein |
| - | 372 | Mobile element protein |
| - | 957 | Mobile element protein |
| - | 828 | Mobile element protein |
| - | 372 | Mobile element protein |
| + | 1341 | Mobile element protein |
| - | 1188 | Mobile element protein |
| - | 957 | Mobile element protein |
| + | 957 | Mobile element protein |
| - | 1341 | Mobile element protein |
| - | 732 | Mobile element protein |
| - | 372 | Mobile element protein |
| + | 1341 | Mobile element protein |
| + | 372 | Mobile element protein |
| + | 828 | Mobile element protein |
| - | 174 | Mobile element protein |
| + | 372 | Mobile element protein |
| + | 828 | Mobile element protein |
| - | 957 | Mobile element protein |
| + | 1341 | Mobile element protein |
| - | 1257 | Mobile element protein |

| | | |
|---|---|---|
| - | 1185 | Mobile element protein |
| - | 957 | Mobile element protein |
| + | 588 | Mobile element protein |
| - | 957 | Mobile element protein |
| - | 1341 | Mobile element protein |
| - | 1362 | Mobile element protein |
| - | 957 | Mobile element protein |
| + | 1341 | Mobile element protein |
| + | 957 | Mobile element protein |
| + | 957 | Mobile element protein |
| - | 1341 | Mobile element protein |
| + | 360 | Mobile element protein |
| + | 828 | Mobile element protein |
| + | 372 | Mobile element protein |
| + | 828 | Mobile element protein |
| + | 1341 | Mobile element protein |
| + | 279 | Mobile element protein |
| - | 957 | Mobile element protein |
| + | 639 | Mobile element protein |
| - | 1341 | Mobile element protein |
| - | 957 | Mobile element protein |
| + | 822 | Mobile element protein |
| + | 372 | Mobile element protein |
| + | 588 | Mobile element protein |
| + | 957 | Mobile element protein |
| + | 1341 | Mobile element protein |
| + | 957 | Mobile element protein |
| + | 588 | Mobile element protein |
| - | 942 | Mobile element protein |
| + | 231 | Mobile element protein |
| - | 1341 | Mobile element protein |
| - | 1341 | Mobile element protein |
| - | 957 | Mobile element protein |
| + | 957 | Mobile element protein |
| - | 1314 | Mobile element protein |
| - | 957 | Mobile element protein |
| + | 1341 | Mobile element protein |
| + | 372 | Mobile element protein |
| - | 957 | Mobile element protein |
| - | 1341 | Mobile element protein |
| - | 957 | Mobile element protein |
| - | 1341 | Mobile element protein |

| | | |
|---|---|---|
| + | 1341 | Mobile element protein |
| - | 588 | Mobile element protein |
| - | 957 | Mobile element protein |
| + | 957 | Mobile element protein |
| + | 1341 | Mobile element protein |
| - | 957 | Mobile element protein |
| + | 1341 | Mobile element protein |
| + | 1341 | Mobile element protein |
| - | 957 | Mobile element protein |
| - | 957 | Mobile element protein |
| + | 372 | Mobile element protein |
| + | 828 | Mobile element protein |
| - | 957 | Mobile element protein |
| + | 231 | Mobile element protein |
| - | 957 | Mobile element protein |
| - | 1341 | Mobile element protein |
| + | 738 | Mobile element protein |
| + | 252 | Mobile element protein |
| - | 957 | Mobile element protein |
| + | 1341 | Mobile element protein |
| + | 1038 | Mobile element protein |
| + | 231 | Mobile element protein |
| - | 231 | Mobile element protein |
| + | 231 | Mobile element protein |
| - | 1341 | Mobile element protein |
| + | 1341 | Mobile element protein |
| + | 1341 | Mobile element protein |
| - | 1341 | Mobile element protein |
| - | 957 | Mobile element protein |
| + | 306 | Mobile element protein |
| + | 639 | Mobile element protein |
| + | 1341 | Mobile element protein |
| - | 873 | Mobile element protein |
| - | 957 | Mobile element protein |
| - | 639 | Mobile element protein |
| - | 699 | Mobile element protein |
| - | 639 | Mobile element protein |
| - | 828 | Mobile element protein |
| - | 306 | Mobile element protein |
| + | 957 | Mobile element protein |
| + | 372 | Mobile element protein |
| + | 828 | Mobile element protein |

| | | |
|---|---|---|
| - | 828 | Mobile element protein |
| - | 213 | Mobile element protein |
| - | 639 | Mobile element protein |
| - | 339 | Mobile element protein |
| - | 828 | Mobile element protein |
| - | 372 | Mobile element protein |
| - | 828 | Mobile element protein |
| - | 345 | Mobile element protein |
| + | 1341 | Mobile element protein |
| - | 828 | Mobile element protein |
| - | 372 | Mobile element protein |
| - | 828 | Mobile element protein |
| - | 372 | Mobile element protein |
| - | 750 | Mobile element protein |
| + | 1362 | Mobile element protein |
| + | 1347 | Mobile element protein |
| + | 957 | Mobile element protein |
| + | 372 | Mobile element protein |
| + | 828 | Mobile element protein |
| + | 957 | Mobile element protein |
| - | 1341 | Mobile element protein |
| - | 957 | Mobile element protein |
| + | 597 | Mobile element protein |
| - | 831 | Mobile element protein |
| + | 315 | Mobile element protein |
| + | 357 | Mobile element protein |
| - | 357 | Mobile element protein |
| - | 315 | Mobile element protein |
| - | 423 | Mobile element protein |
| + | 315 | Mobile element protein |
| + | 465 | Mobile element protein |
| - | 273 | Mobile element protein |
| - | 465 | Mobile element protein |
| - | 315 | Mobile element protein |
| + | 465 | Mobile element protein |
| + | 315 | Mobile element protein |
| + | 408 | Mobile element protein |
| + | 357 | Mobile element protein |
| + | 315 | Mobile element protein |
| + | 357 | Mobile element protein |
| + | 315 | Mobile element protein |
| + | 357 | Mobile element protein |

| | | |
|---|---|---|
| + | 357 | Mobile element protein |
| - | 465 | Mobile element protein |
| - | 315 | Mobile element protein |
| + | 315 | Mobile element protein |
| + | 465 | Mobile element protein |
| - | 465 | Mobile element protein |
| - | 315 | Mobile element protein |
| + | 315 | Mobile element protein |
| + | 357 | Mobile element protein |
| - | 462 | Mobile element protein |
| - | 315 | Mobile element protein |
| + | 315 | Mobile element protein |
| + | 363 | Mobile element protein |
| - | 357 | Mobile element protein |
| - | 315 | Mobile element protein |
| + | 315 | Mobile element protein |
| + | 357 | Mobile element protein |
| - | 357 | Mobile element protein |
| - | 315 | Mobile element protein |
| - | 357 | Mobile element protein |
| - | 315 | Mobile element protein |
| - | 357 | Mobile element protein |
| - | 315 | Mobile element protein |
| - | 357 | Mobile element protein |
| - | 210 | Mobile element protein |
| - | 357 | Mobile element protein |
| - | 315 | Mobile element protein |
| - | 465 | Mobile element protein |
| + | 315 | Mobile element protein |
| + | 357 | Mobile element protein |
| - | 273 | Mobile element protein |
| - | 357 | Mobile element protein |
| - | 315 | Mobile element protein |
| + | 210 | Mobile element protein |
| + | 465 | Mobile element protein |
| - | 357 | Mobile element protein |
| - | 315 | Mobile element protein |
| + | 315 | Mobile element protein |
| + | 357 | Mobile element protein |
| - | 357 | Mobile element protein |
| - | 315 | Mobile element protein |
| + | 315 | Mobile element protein |

| | | |
|---|---|---|
| + | 357 | Mobile element protein |
| - | 357 | Mobile element protein |
| - | 315 | Mobile element protein |
| - | 465 | Mobile element protein |
| - | 210 | Mobile element protein |
| + | 273 | Mobile element protein |
| - | 144 | Mobile element protein |
| - | 363 | Mobile element protein |
| - | 315 | Mobile element protein |
| + | 357 | Mobile element protein |
| + | 372 | Mobile element protein |
| + | 483 | Mobile element protein |
| - | 483 | Mobile element protein |
| - | 372 | Mobile element protein |
| + | 315 | Mobile element protein |
| + | 357 | Mobile element protein |
| + | 315 | Mobile element protein |
| + | 357 | Mobile element protein |
| + | 357 | Mobile element protein |
| + | 177 | Mobile element protein |
| - | 465 | Mobile element protein |
| - | 315 | Mobile element protein |
| + | 315 | Mobile element protein |
| + | 465 | Mobile element protein |
| + | 315 | Mobile element protein |
| + | 375 | Mobile element protein |
| - | 465 | Mobile element protein |
| - | 315 | Mobile element protein |
| + | 315 | Mobile element protein |
| + | 357 | Mobile element protein |
| - | 465 | Mobile element protein |
| - | 315 | Mobile element protein |
| - | 144 | Mobile element protein |
| - | 363 | Mobile element protein |
| - | 315 | Mobile element protein |
| - | 654 | Mobile element protein |
| - | 330 | Mobile element protein |
| + | 357 | Mobile element protein |
| + | 315 | Mobile element protein |
| + | 357 | Mobile element protein |
| + | 315 | Mobile element protein |
| + | 357 | Mobile element protein |

| | | |
|---|---|---|
| - | 357 | Mobile element protein |
| - | 315 | Mobile element protein |
| - | 357 | Mobile element protein |
| - | 315 | Mobile element protein |
| + | 315 | Mobile element protein |
| + | 465 | Mobile element protein |
| + | 315 | Mobile element protein |
| + | 357 | Mobile element protein |
| - | 273 | Mobile element protein |
| - | 465 | Mobile element protein |
| - | 315 | Mobile element protein |
| + | 315 | Mobile element protein |
| + | 357 | Mobile element protein |
| + | 315 | Mobile element protein |
| + | 465 | Mobile element protein |
| - | 357 | Mobile element protein |
| - | 315 | Mobile element protein |
| + | 315 | Mobile element protein |
| + | 357 | Mobile element protein |
| - | 465 | Mobile element protein |
| - | 315 | Mobile element protein |
| - | 357 | Mobile element protein |
| - | 315 | Mobile element protein |
| + | 315 | Mobile element protein |
| + | 465 | Mobile element protein |
| - | 357 | Mobile element protein |
| - | 315 | Mobile element protein |
| + | 315 | Mobile element protein |
| + | 357 | Mobile element protein |
| + | 315 | Mobile element protein |
| + | 465 | Mobile element protein |
| - | 465 | Mobile element protein |
| - | 315 | Mobile element protein |
| + | 273 | Mobile element protein |
| - | 357 | Mobile element protein |
| - | 315 | Mobile element protein |
| - | 252 | Mobile element protein |
| - | 315 | Mobile element protein |
| - | 465 | Mobile element protein |
| - | 315 | Mobile element protein |
| - | 357 | Mobile element protein |
| - | 315 | Mobile element protein |

| | | |
|---|---|---|
| + | 315 | Mobile element protein |
| + | 363 | Mobile element protein |
| + | 144 | Mobile element protein |
| - | 465 | Mobile element protein |
| - | 315 | Mobile element protein |
| - | 465 | Mobile element protein |
| - | 315 | Mobile element protein |
| + | 315 | Mobile element protein |
| + | 465 | Mobile element protein |
| - | 465 | Mobile element protein |
| - | 315 | Mobile element protein |
| + | 315 | Mobile element protein |
| + | 465 | Mobile element protein |
| + | 315 | Mobile element protein |
| + | 357 | Mobile element protein |
| - | 315 | Mobile element protein |
| - | 315 | Mobile element protein |
| + | 270 | Mobile element protein |
| - | 357 | Mobile element protein |
| - | 315 | Mobile element protein |
| - | 357 | Mobile element protein |
| - | 315 | Mobile element protein |
| - | 357 | Mobile element protein |
| - | 315 | Mobile element protein |
| + | 315 | Mobile element protein |
| + | 357 | Mobile element protein |
| + | 315 | Mobile element protein |
| + | 363 | Mobile element protein |
| + | 144 | Mobile element protein |
| + | 315 | Mobile element protein |
| + | 363 | Mobile element protein |
| + | 144 | Mobile element protein |
| + | 315 | Mobile element protein |
| + | 357 | Mobile element protein |
| - | 357 | Mobile element protein |
| - | 315 | Mobile element protein |
| + | 315 | Mobile element protein |
| + | 315 | Mobile element protein |
| - | 273 | Mobile element protein |
| + | 315 | Mobile element protein |
| + | 357 | Mobile element protein |
| + | 315 | Mobile element protein |

| | | |
|---|---|---|
| + | 465 | Mobile element protein |
| - | 465 | Mobile element protein |
| - | 315 | Mobile element protein |
| - | 465 | Mobile element protein |
| - | 315 | Mobile element protein |
| - | 465 | Mobile element protein |
| - | 315 | Mobile element protein |
| + | 315 | Mobile element protein |
| + | 363 | Mobile element protein |
| + | 315 | Mobile element protein |
| + | 465 | Mobile element protein |
| + | 315 | Mobile element protein |
| + | 381 | Mobile element protein |
| - | 357 | Mobile element protein |
| - | 315 | Mobile element protein |
| + | 315 | Mobile element protein |
| + | 357 | Mobile element protein |
| + | 315 | Mobile element protein |
| + | 357 | Mobile element protein |
| + | 315 | Mobile element protein |
| + | 465 | Mobile element protein |
| - | 357 | Mobile element protein |
| - | 315 | Mobile element protein |
| - | 357 | Mobile element protein |
| - | 315 | Mobile element protein |
| + | 315 | Mobile element protein |
| + | 465 | Mobile element protein |
| + | 273 | Mobile element protein |
| - | 465 | Mobile element protein |
| - | 315 | Mobile element protein |
| + | 357 | Mobile element protein |
| - | 465 | Mobile element protein |
| - | 315 | Mobile element protein |
| - | 357 | Mobile element protein |
| - | 315 | Mobile element protein |
| - | 393 | Mobile element protein |
| - | 315 | Mobile element protein |
| + | 315 | Mobile element protein |
| + | 465 | Mobile element protein |
| - | 465 | Mobile element protein |
| - | 315 | Mobile element protein |
| + | 315 | Mobile element protein |

| | | | |
|---|---|---|---|
| + | | 357 | Mobile element protein |
| + | | 315 | Mobile element protein |
| + | | 453 | Mobile element protein |
| - | - | | - |
| - | - | | - |
| + | | 573 | Mobile element protein |
| - | | 750 | Mobile element protein |
| - | | 573 | Mobile element protein |
| - | | 459 | Mobile element protein |
| + | | 972 | Mobile element protein |
| - | | 309 | Mobile element protein |
| - | | 522 | Mobile element protein |
| - | | 972 | Mobile element protein |
| + | | 972 | Mobile element protein |
| + | | 951 | Mobile element protein |
| + | | 972 | Mobile element protein |
| + | | 960 | Mobile element protein |
| - | | 960 | Mobile element protein |
| + | | 750 | Mobile element protein |
| + | | 309 | Mobile element protein |
| - | | 972 | Mobile element protein |
| - | | 132 | Mobile element protein |
| - | | 2025 | Mobile element protein |
| - | | 348 | Mobile element protein |
| + | | 855 | Mobile element protein |
| + | | 2025 | Mobile element protein |
| + | | 132 | Mobile element protein |
| - | | 2025 | Mobile element protein |
| - | | 855 | Mobile element protein |
| + | | 855 | Mobile element protein |
| + | | 2025 | Mobile element protein |
| + | | 294 | Mobile element protein |
| + | | 972 | Mobile element protein |
| - | | 960 | Mobile element protein |
| + | | 972 | Mobile element protein |
| - | | 132 | Mobile element protein |
| - | | 2025 | Mobile element protein |
| - | | 348 | Mobile element protein |
| + | | 855 | Mobile element protein |
| + | | 2025 | Mobile element protein |
| + | | 132 | Mobile element protein |
| - | | 132 | Mobile element protein |

| | | |
|---|---|---|
| - | 2025 | Mobile element protein |
| + | 1221 | Mobile element protein |
| + | 855 | Mobile element protein |
| + | 2025 | Mobile element protein |
| + | 132 | Mobile element protein |
| + | 573 | Mobile element protein |
| + | 426 | Mobile element protein |
| + | 501 | Mobile element protein |
| - | 792 | Mobile element protein |
| + | 750 | Mobile element protein |
| - | 972 | Mobile element protein |
| + | 297 | Mobile element protein |
| - | 294 | Mobile element protein |
| + | 804 | Mobile element protein |
| - | 972 | Mobile element protein |
| - | 960 | Mobile element protein |
| - | 972 | Mobile element protein |
| - | 294 | Mobile element protein |
| + | 1188 | Mobile element protein |
| + | 1119 | Mobile element protein |
| - | 2025 | Mobile element protein |
| - | 855 | Mobile element protein |
| - | 222 | Mobile element protein |
| + | 1017 | Mobile element protein |
| - | 750 | Mobile element protein |
| - | 1017 | Mobile element protein |
| + | 510 | Mobile element protein |
| + | 2025 | Mobile element protein |
| - | 510 | Mobile element protein |
| + | 1221 | Mobile element protein |
| + | 750 | Mobile element protein |
| - | 972 | Mobile element protein |
| + | 972 | Mobile element protein |
| - | 750 | Mobile element protein |
| + | 960 | Mobile element protein |
| + | 309 | Mobile element protein |
| - | 972 | Mobile element protein |
| - | 972 | Mobile element protein |
| + | 972 | Mobile element protein |
| - | 972 | Mobile element protein |
| - | 459 | Mobile element protein |
| - | 972 | Mobile element protein |

| | | |
|---|---|---|
| + | 573 | Mobile element protein |
| + | 960 | Mobile element protein |
| + | 972 | Mobile element protein |
| - | 972 | Mobile element protein |
| + | 573 | Mobile element protein |
| + | 972 | Mobile element protein |
| - | 213 | Mobile element protein |
| - | 123 | Mobile element protein |
| + | 198 | Mobile element protein |
| + | 180 | Mobile element protein |
| - | 180 | Mobile element protein |
| + | 252 | Mobile element protein |
| - | 123 | Mobile element protein |
| + | 141 | Mobile element protein |
| + | 111 | Mobile element protein |
| + | 207 | Mobile element protein |
| + | 249 | Mobile element protein |
| + | 123 | Mobile element protein |
| - | 237 | Mobile element protein |
| - | 234 | Mobile element protein |
| - | 234 | Mobile element protein |
| - | 285 | Mobile element protein |
| - | 156 | Mobile element protein |
| + | 213 | Mobile element protein |
| - | 261 | Mobile element protein |
| - | 210 | Mobile element protein |
| - | 186 | Mobile element protein |
| - | 117 | Mobile element protein |
| - | 180 | Mobile element protein |
| + | 156 | Mobile element protein |
| + | 165 | Mobile element protein |
| + | 117 | Mobile element protein |
| + | 162 | Mobile element protein |
| + | 114 | Mobile element protein |
| + | 210 | Mobile element protein |
| + | 174 | Mobile element protein |
| - | 168 | Mobile element protein |
| + | 168 | Mobile element protein |
| - | 192 | Mobile element protein |
| + | 192 | Mobile element protein |
| + | 177 | Mobile element protein |
| + | 150 | Mobile element protein |

| | | |
|---|---|---|
| + | 216 | Mobile element protein |
| + | 192 | Mobile element protein |
| - | 1347 | Mobile element protein |
| - | 342 | Mobile element protein |
| - | 381 | Mobile element protein |
| + | 819 | Mobile element protein |
| + | 345 | Mobile element protein |
| + | 819 | Mobile element protein |
| + | 345 | Mobile element protein |
| + | 1347 | Mobile element protein |
| - | 156 | Mobile element protein |
| - | 114 | Mobile element protein |
| - | 294 | Mobile element protein |
| - | 120 | Mobile element protein |
| - | 264 | Mobile element protein |
| + | 438 | Mobile element protein |
| + | 186 | Mobile element protein |
| + | 312 | Mobile element protein |
| + | 264 | Mobile element protein |
| + | 219 | Mobile element protein |
| + | 186 | Mobile element protein |
| - | 126 | Mobile element protein |
| - | 225 | Mobile element protein |
| - | 171 | Mobile element protein |
| - | 174 | Mobile element protein |
| + | 114 | Mobile element protein |
| - | 549 | Mobile element protein |
| - | 333 | Mobile element protein |
| - | 285 | Mobile element protein |
| - | 297 | Mobile element protein |
| - | 333 | Mobile element protein |
| + | 297 | Mobile element protein |
| - | 726 | Mobile element protein |
| - | 207 | Mobile element protein |
| - | 192 | Mobile element protein |
| - | 216 | Mobile element protein |
| - | 150 | Mobile element protein |
| + | 192 | Mobile element protein |
| - | 345 | Mobile element protein |
| - | 819 | Mobile element protein |
| - | 171 | Mobile element protein |
| - | 192 | Mobile element protein |

| | | |
|---|---|---|
| - | 216 | Mobile element protein |
| - | 297 | Mobile element protein |
| + | 123 | Mobile element protein |
| + | 174 | Mobile element protein |
| + | 186 | Mobile element protein |
| + | 624 | Mobile element protein |
| + | 537 | Mobile element protein |
| - | 537 | Mobile element protein |
| + | 1293 | Mobile element protein |
| + | 264 | Mobile element protein |
| + | 126 | Mobile element protein |
| + | 297 | Mobile element protein |
| + | 171 | Mobile element protein |
| + | 114 | Mobile element protein |
| + | 174 | Mobile element protein |
| + | 126 | Mobile element protein |
| - | 345 | Mobile element protein |
| - | 819 | Mobile element protein |
| - | 297 | Mobile element protein |
| - | 405 | Mobile element protein |
| - | 126 | Mobile element protein |
| - | 129 | Mobile element protein |
| - | 150 | Mobile element protein |
| + | 381 | Mobile element protein |
| + | 357 | Mobile element protein |
| + | 150 | Mobile element protein |
| + | 216 | Mobile element protein |
| + | 192 | Mobile element protein |
| - | 345 | Mobile element protein |
| - | 819 | Mobile element protein |
| - | 249 | Mobile element protein |
| + | 150 | Mobile element protein |
| + | 216 | Mobile element protein |
| + | 192 | Mobile element protein |
| - | 186 | Mobile element protein |
| - | 219 | Mobile element protein |
| - | 345 | Mobile element protein |
| - | 819 | Mobile element protein |
| + | 219 | Mobile element protein |
| + | 1293 | Mobile element protein |
| + | 159 | Mobile element protein |
| + | 159 | Mobile element protein |

| | | |
|---|---|---|
| - | 192 | Mobile element protein |
| - | 216 | Mobile element protein |
| - | 150 | Mobile element protein |
| + | 186 | Mobile element protein |
| + | 312 | Mobile element protein |
| - | 216 | Mobile element protein |
| - | 114 | Mobile element protein |
| - | 255 | Mobile element protein |
| - | 1293 | Mobile element protein |
| - | 171 | Mobile element protein |
| - | 192 | Mobile element protein |
| - | 216 | Mobile element protein |
| - | 150 | Mobile element protein |
| - | 114 | Mobile element protein |
| - | 132 | Mobile element protein |
| - | 669 | Mobile element protein |
| + | 207 | Mobile element protein |
| + | 549 | Mobile element protein |
| - | 174 | Mobile element protein |
| + | 333 | Mobile element protein |
| + | 297 | Mobile element protein |
| + | 333 | Mobile element protein |
| + | 297 | Mobile element protein |
| - | 312 | Mobile element protein |
| - | 186 | Mobile element protein |
| - | 312 | Mobile element protein |
| - | 186 | Mobile element protein |
| + | 186 | Mobile element protein |
| + | 312 | Mobile element protein |
| - | 312 | Mobile element protein |
| - | 186 | Mobile element protein |
| - | 345 | Mobile element protein |
| - | 819 | Mobile element protein |
| + | 1293 | Mobile element protein |
| - | 153 | Mobile element protein |
| - | 426 | Mobile element protein |
| + | 150 | Mobile element protein |
| + | 216 | Mobile element protein |
| + | 192 | Mobile element protein |
| - | 1347 | Mobile element protein |
| - | 927 | Mobile element protein |
| - | 402 | Mobile element protein |

| | | |
|---|---|---|
| - | 525 | Mobile element protein |
| - | 132 | Mobile element protein |
| - | 243 | Mobile element protein |
| + | 537 | Mobile element protein |
| - | 411 | Mobile element protein |
| - | 219 | Mobile element protein |
| - | 687 | Mobile element protein |
| - | 342 | Mobile element protein |
| - | 381 | Mobile element protein |
| + | 1329 | Mobile element protein |
| + | 381 | Mobile element protein |
| + | 342 | Mobile element protein |
| - | 507 | Mobile element protein |
| - | 342 | Mobile element protein |
| - | 381 | Mobile element protein |
| - | 1329 | Mobile element protein |
| - | 1233 | Mobile element protein |
| - | 342 | Mobile element protein |
| - | 381 | Mobile element protein |
| + | 381 | Mobile element protein |
| + | 342 | Mobile element protein |
| - | 381 | Mobile element protein |
| - | 342 | Mobile element protein |
| - | 381 | Mobile element protein |
| + | 381 | Mobile element protein |
| + | 342 | Mobile element protein |
| - | 222 | Mobile element protein |
| + | 1329 | Mobile element protein |
| - | 342 | Mobile element protein |
| - | 381 | Mobile element protein |
| - | 480 | Mobile element protein |
| + | 381 | Mobile element protein |
| + | 342 | Mobile element protein |
| - | 297 | Mobile element protein |
| + | 387 | Mobile element protein |
| - | 342 | Mobile element protein |
| - | 381 | Mobile element protein |
| - | 342 | Mobile element protein |
| - | 381 | Mobile element protein |
| + | 249 | Mobile element protein |
| + | 381 | Mobile element protein |
| + | 342 | Mobile element protein |

| | | |
|---|---|---|
| + | 408 | Mobile element protein |
| + | 381 | Mobile element protein |
| + | 342 | Mobile element protein |
| + | 330 | Mobile element protein |
| + | 201 | Mobile element protein |
| + | 393 | Mobile element protein |
| - | 408 | Mobile element protein |
| - | 201 | Mobile element protein |
| - | 330 | Mobile element protein |
| - | 309 | Mobile element protein |
| - | 294 | Mobile element protein |
| + | 408 | Mobile element protein |
| - | 429 | Mobile element protein |
| + | 393 | Mobile element protein |
| + | 393 | Mobile element protein |
| - | 408 | Mobile element protein |
| - | 201 | Mobile element protein |
| - | 330 | Mobile element protein |
| + | 165 | Mobile element protein |
| + | 165 | Mobile element protein |
| - | 261 | Mobile element protein |
| - | 357 | Mobile element protein |
| + | 318 | Mobile element protein |
| - | 603 | Mobile element protein |
| - | 423 | Mobile element protein |
| - | 252 | Mobile element protein |
| - | 237 | Mobile element protein |
| - | 252 | Mobile element protein |
| + | 105 | Mobile element protein |
| + | 162 | Mobile element protein |
| + | 174 | Mobile element protein |
| + | 108 | Mobile element protein |
| - | 192 | Mobile element protein |
| + | 303 | Mobile element protein |
| - | 141 | Mobile element protein |
| + | 342 | Mobile element protein |
| + | 270 | Mobile element protein |
| + | 261 | Mobile element protein |
| + | 123 | Mobile element protein |
| + | 219 | Mobile element protein |
| - | 189 | Mobile element protein |
| - | 204 | Mobile element protein |

| | | |
|---|---|---|
| - | 303 | Mobile element protein |
| + | 165 | Mobile element protein |
| - | 219 | Mobile element protein |
| + | 141 | Mobile element protein |
| + | 270 | Mobile element protein |
| + | 231 | Mobile element protein |
| - | 462 | Mobile element protein |
| - | 183 | Mobile element protein |
| - | 345 | Mobile element protein |
| + | 219 | Mobile element protein |
| - | 135 | Mobile element protein |
| - | 159 | Mobile element protein |
| + | 264 | Mobile element protein |
| + | 333 | Mobile element protein |
| - | 219 | Mobile element protein |
| - | 1026 | Mobile element protein |
| - | 219 | Mobile element protein |
| - | 303 | Mobile element protein |
| - | 258 | Mobile element protein |
| - | 270 | Mobile element protein |
| - | 270 | Mobile element protein |
| - | 192 | Mobile element protein |
| - | 123 | Mobile element protein |
| - | 261 | Mobile element protein |
| + | 255 | Mobile element protein |
| + | 84 | Mobile element protein |
| - | 219 | Mobile element protein |
| - | 252 | Mobile element protein |
| - | 180 | Mobile element protein |
| - | 294 | Mobile element protein |
| - | 123 | Mobile element protein |
| - | 261 | Mobile element protein |
| + | 84 | Mobile element protein |
| + | 219 | Mobile element protein |
| - | 219 | Mobile element protein |
| - | 369 | Mobile element protein |
| + | 171 | Mobile element protein |
| - | 339 | Mobile element protein |
| + | 297 | Mobile element protein |
| - | 405 | Mobile element protein |
| + | 219 | Mobile element protein |
| - | 321 | Mobile element protein |

| | | | |
|---|---|---|---|
| + | | 219 | Mobile element protein |
| - | | 219 | Mobile element protein |
| + | | 678 | Mobile element protein |
| + | | 477 | Mobile element protein |
| + | | 798 | Mobile element protein |
| + | | 219 | Mobile element protein |
| - | | 219 | Mobile element protein |
| + | | 381 | Mobile element protein |
| - | | 123 | Mobile element protein |
| - | | 261 | Mobile element protein |
| + | | 294 | Mobile element protein |
| - | | 237 | Mobile element protein |
| + | | 285 | Mobile element protein |
| - | | 528 | Mobile element protein |
| + | | 222 | Mobile element protein |
| - | | 153 | Mobile element protein |
| - | | 219 | Mobile element protein |
| - | | 183 | Mobile element protein |
| + | | 342 | Mobile element protein |
| - | | 228 | Mobile element protein |
| + | | 183 | Mobile element protein |
| - | | 294 | Mobile element protein |
| + | | 249 | Mobile element protein |
| + | | 204 | Mobile element protein |
| - | | 201 | Mobile element protein |
| - | - | | - |
| - | - | | - |
| - | | 150 | Mobile element protein |
| - | | 207 | Mobile element protein |
| + | | 90 | Mobile element protein |
| + | | 168 | Mobile element protein |
| + | | 234 | Mobile element protein |
| - | | 474 | Mobile element protein |
| + | | 423 | Mobile element protein |
| - | | 606 | Mobile element protein |
| - | | 408 | Mobile element protein |
| - | | 189 | Mobile element protein |
| + | | 381 | Mobile element protein |
| - | | 873 | Mobile element protein |
| + | | 183 | Mobile element protein |
| + | | 873 | Mobile element protein |
| + | | 669 | Mobile element protein |

| | | |
|---|---|---|
| + | 591 | Mobile element protein |
| - | 873 | Mobile element protein |
| + | 483 | Mobile element protein |
| + | 459 | Mobile element protein |
| - | 873 | Mobile element protein |
| + | 219 | Mobile element protein |
| + | 324 | Mobile element protein |
| - | 873 | Mobile element protein |
| + | 477 | Mobile element protein |
| + | 165 | Mobile element protein |
| + | 357 | Mobile element protein |
| - | 357 | Mobile element protein |
| - | 165 | Mobile element protein |
| - | 477 | Mobile element protein |
| + | 258 | Mobile element protein |
| + | 141 | Mobile element protein |
| + | 240 | Mobile element protein |
| - | 873 | Mobile element protein |
| - | 408 | Mobile element protein |
| - | 873 | Mobile element protein |
| - | 357 | Mobile element protein |
| - | 165 | Mobile element protein |
| - | 477 | Mobile element protein |
| - | 873 | Mobile element protein |
| + | 597 | Mobile element protein |
| - | 408 | Mobile element protein |
| + | 873 | Mobile element protein |
| - | 330 | Mobile element protein |
| - | 591 | Mobile element protein |
| - | 669 | Mobile element protein |
| - | 1239 | Mobile element protein |
| - | 873 | Mobile element protein |
| + | 711 | Mobile element protein |
| + | 873 | Mobile element protein |
| - | 225 | Mobile element protein |
| - | 216 | Mobile element protein |
| - | 207 | Mobile element protein |
| - | 873 | Mobile element protein |
| - | 981 | Mobile element protein |
| - | 249 | Mobile element protein |
| + | 873 | Mobile element protein |
| + | 201 | Mobile element protein |

| | | |
|---|---|---|
| + | 591 | Mobile element protein |
| - | 330 | Mobile element protein |
| - | 261 | Mobile element protein |
| - | 783 | Mobile element protein |
| - | 873 | Mobile element protein |
| - | 873 | Mobile element protein |
| + | 249 | Mobile element protein |
| + | 411 | Mobile element protein |
| + | 591 | Mobile element protein |
| - | 330 | Mobile element protein |
| + | 1101 | Mobile element protein |
| + | 132 | Mobile element protein |
| + | 201 | Mobile element protein |
| + | 873 | Mobile element protein |
| - | 702 | Mobile element protein |
| + | 873 | Mobile element protein |
| - | 423 | Mobile element protein |
| + | 777 | Mobile element protein |
| + | 249 | Mobile element protein |
| + | 411 | Mobile element protein |
| + | 411 | Mobile element protein |
| + | 132 | Mobile element protein |
| + | 423 | Mobile element protein |
| - | 873 | Mobile element protein |
| + | 873 | Mobile element protein |
| + | 477 | Mobile element protein |
| + | 165 | Mobile element protein |
| + | 357 | Mobile element protein |
| - | 357 | Mobile element protein |
| - | 312 | Mobile element protein |
| + | 192 | Mobile element protein |
| - | 573 | Mobile element protein |
| + | 873 | Mobile element protein |
| - | 249 | Mobile element protein |
| + | 474 | Mobile element protein |
| - | 873 | Mobile element protein |
| - | 591 | Mobile element protein |
| - | 669 | Mobile element protein |
| + | 933 | Mobile element protein |
| - | 873 | Mobile element protein |
| + | 636 | Mobile element protein |
| + | 636 | Mobile element protein |

| | | |
|---|---|---|
| - | 330 | Mobile element protein |
| + | 621 | Mobile element protein |
| + | 249 | Mobile element protein |
| + | 312 | Mobile element protein |
| - | 357 | Mobile element protein |
| - | 165 | Mobile element protein |
| - | 477 | Mobile element protein |
| + | 873 | Mobile element protein |
| + | 477 | Mobile element protein |
| + | 165 | Mobile element protein |
| + | 357 | Mobile element protein |
| + | 873 | Mobile element protein |
| + | 408 | Mobile element protein |
| - | 138 | Mobile element protein |
| - | 669 | Mobile element protein |
| + | 873 | Mobile element protein |
| + | 477 | Mobile element protein |
| + | 129 | Mobile element protein |
| - | 423 | Mobile element protein |
| + | 351 | Mobile element protein |
| + | 591 | Mobile element protein |
| + | 477 | Mobile element protein |
| + | 165 | Mobile element protein |
| + | 357 | Mobile element protein |
| + | 873 | Mobile element protein |
| + | 873 | Mobile element protein |
| - | 165 | Mobile element protein |
| - | 477 | Mobile element protein |
| - | 132 | Mobile element protein |
| - | 132 | Mobile element protein |
| + | 477 | Mobile element protein |
| + | 165 | Mobile element protein |
| + | 357 | Mobile element protein |
| - | 357 | Mobile element protein |
| - | 183 | Mobile element protein |
| - | 873 | Mobile element protein |
| - | 471 | Mobile element protein |
| + | 477 | Mobile element protein |
| + | 165 | Mobile element protein |
| + | 357 | Mobile element protein |
| + | 1239 | Mobile element protein |
| - | 1239 | Mobile element protein |

| | | |
|---|---|---|
| - | 873 | Mobile element protein |
| - | 873 | Mobile element protein |
| + | 357 | Mobile element protein |
| + | 591 | Mobile element protein |
| - | 483 | Mobile element protein |
| - | 366 | Mobile element protein |
| + | 333 | Mobile element protein |
| + | 873 | Mobile element protein |
| + | 873 | Mobile element protein |
| - | 879 | Mobile element protein |
| + | 873 | Mobile element protein |
| + | 669 | Mobile element protein |
| + | 591 | Mobile element protein |
| + | 483 | Mobile element protein |
| - | 366 | Mobile element protein |
| - | 864 | Mobile element protein |
| - | 873 | Mobile element protein |
| + | 267 | Mobile element protein |
| + | 165 | Mobile element protein |
| + | 357 | Mobile element protein |
| - | 483 | Mobile element protein |
| - | 408 | Mobile element protein |
| - | 384 | Mobile element protein |
| - | 732 | Mobile element protein |
| - | 591 | Mobile element protein |
| + | 873 | Mobile element protein |
| - | 477 | Mobile element protein |
| - | 873 | Mobile element protein |
| + | 330 | Mobile element protein |
| - | 597 | Mobile element protein |
| + | 873 | Mobile element protein |
| - | 138 | Mobile element protein |
| - | 792 | Mobile element protein |
| + | 1101 | Mobile element protein |
| + | 132 | Mobile element protein |
| + | 873 | Mobile element protein |
| + | 285 | Mobile element protein |
| + | 1347 | Mobile element protein |
| + | 150 | Mobile element protein |
| - | 150 | Mobile element protein |
| + | 1347 | Mobile element protein |
| + | 150 | Mobile element protein |

| | | |
|---|---|---|
| + | 150 | Mobile element protein |
| - | 948 | Mobile element protein |
| - | 150 | Mobile element protein |
| + | 150 | Mobile element protein |
| + | 699 | Mobile element protein |
| - | 948 | Mobile element protein |
| - | 1347 | Mobile element protein |
| + | 222 | Mobile element protein |
| + | 117 | Mobile element protein |
| + | 873 | Mobile element protein |
| + | 150 | Mobile element protein |
| - | 411 | Mobile element protein |
| - | 381 | Mobile element protein |
| + | 498 | Mobile element protein |
| + | 948 | Mobile element protein |
| + | 150 | Mobile element protein |
| - | 150 | Mobile element protein |
| - | 411 | Mobile element protein |
| - | 381 | Mobile element protein |
| - | 306 | Mobile element protein |
| + | 477 | Mobile element protein |
| + | 120 | Mobile element protein |
| + | 348 | Mobile element protein |
| + | 150 | Mobile element protein |
| - | 150 | Mobile element protein |
| - | 138 | Mobile element protein |
| + | 876 | Mobile element protein |
| - | 873 | Mobile element protein |
| - | 498 | Mobile element protein |
| - | 510 | Mobile element protein |
| - | 114 | Mobile element protein |
| - | 1347 | Mobile element protein |
| - | 174 | Mobile element protein |
| + | 114 | Mobile element protein |
| + | 156 | Mobile element protein |
| - | 168 | Mobile element protein |
| - | 126 | Mobile element protein |
| + | 150 | Mobile element protein |
| + | 150 | Mobile element protein |
| - | 873 | Mobile element protein |
| - | 1347 | Mobile element protein |
| + | 456 | Mobile element protein |

| | | |
|---|---|---|
| + | 414 | Mobile element protein |
| + | 132 | Mobile element protein |
| - | 948 | Mobile element protein |
| - | 1347 | Mobile element protein |
| - | 150 | Mobile element protein |
| + | 873 | Mobile element protein |
| + | 126 | Mobile element protein |
| - | 150 | Mobile element protein |
| + | 150 | Mobile element protein |
| + | 1347 | Mobile element protein |
| - | 150 | Mobile element protein |
| - | 153 | Mobile element protein |
| + | 429 | Mobile element protein |
| - | 150 | Mobile element protein |
| + | 150 | Mobile element protein |
| + | 159 | Mobile element protein |
| - | 180 | Mobile element protein |
| + | 591 | Mobile element protein |
| - | 948 | Mobile element protein |
| + | 321 | Mobile element protein |
| + | 297 | Mobile element protein |
| - | 150 | Mobile element protein |
| - | 873 | Mobile element protein |
| - | 144 | Mobile element protein |
| - | 150 | Mobile element protein |
| + | 588 | Mobile element protein |
| - | 948 | Mobile element protein |
| - | 948 | Mobile element protein |
| + | 153 | Mobile element protein |
| + | 141 | Mobile element protein |
| + | 117 | Mobile element protein |
| + | 153 | Mobile element protein |
| - | 288 | Mobile element protein |
| + | 306 | Mobile element protein |
| + | 138 | Mobile element protein |
| - | 117 | Mobile element protein |
| + | 117 | Mobile element protein |
| + | 192 | Mobile element protein |
| + | 267 | Mobile element protein |
| + | 306 | Mobile element protein |
| - | 129 | Mobile element protein |
| - | 132 | Mobile element protein |

| | | |
|---|---|---|
| - | 192 | Mobile element protein |
| + | 306 | Mobile element protein |
| + | 603 | Mobile element protein |
| + | 183 | Mobile element protein |
| + | 129 | Mobile element protein |
| + | 234 | Mobile element protein |
| + | 603 | Mobile element protein |
| - | 117 | Mobile element protein |
| - | 603 | Mobile element protein |
| - | 234 | Mobile element protein |
| - | 129 | Mobile element protein |
| - | 183 | Mobile element protein |
| - | 603 | Mobile element protein |
| - | 234 | Mobile element protein |
| - | 129 | Mobile element protein |
| - | 183 | Mobile element protein |
| - | 195 | Mobile element protein |
| - | 126 | Mobile element protein |
| - | 144 | Mobile element protein |
| + | 117 | Mobile element protein |
| - | 603 | Mobile element protein |
| - | 234 | Mobile element protein |
| - | 129 | Mobile element protein |
| - | 183 | Mobile element protein |
| + | 183 | Mobile element protein |
| + | 129 | Mobile element protein |
| + | 234 | Mobile element protein |
| + | 603 | Mobile element protein |
| + | 117 | Mobile element protein |
| - | 144 | Mobile element protein |
| - | 144 | Mobile element protein |
| - | 159 | Mobile element protein |
| - | 132 | Mobile element protein |
| + | 183 | Mobile element protein |
| + | 129 | Mobile element protein |
| + | 234 | Mobile element protein |
| + | 603 | Mobile element protein |
| - | 159 | Mobile element protein |
| - | 258 | Mobile element protein |
| - | 165 | Mobile element protein |
| + | 573 | Mobile element protein |
| + | 399 | Mobile element protein |

| | | |
|---|---|---|
| + | 207 | Mobile element protein |
| + | 690 | Mobile element protein |
| + | 1200 | Mobile element protein |
| - | 297 | Mobile element protein |
| + | 438 | Mobile element protein |
| + | 174 | Mobile element protein |
| - | 783 | Mobile element protein |
| - | 192 | Mobile element protein |
| + | 243 | Mobile element protein |
| + | 135 | Mobile element protein |
| + | 132 | Mobile element protein |
| - | 393 | Mobile element protein |
| - | 132 | Mobile element protein |
| - | 135 | Mobile element protein |
| - | 117 | Mobile element protein |
| + | 354 | Mobile element protein |
| + | 120 | Mobile element protein |
| + | 279 | Mobile element protein |
| + | 315 | Mobile element protein |
| + | 168 | Mobile element protein |
| - | 690 | Mobile element protein |
| - | 132 | Mobile element protein |
| - | 135 | Mobile element protein |
| - | 243 | Mobile element protein |
| + | 261 | Mobile element protein |
| + | 228 | Mobile element protein |
| + | 171 | Mobile element protein |
| - | 135 | Mobile element protein |
| - | 243 | Mobile element protein |
| - | 132 | Mobile element protein |
| - | 135 | Mobile element protein |
| - | 243 | Mobile element protein |
| + | 171 | Mobile element protein |
| + | 129 | Mobile element protein |
| - | 195 | Mobile element protein |
| + | 315 | Mobile element protein |
| + | 360 | Mobile element protein |
| + | 354 | Mobile element protein |
| + | 120 | Mobile element protein |
| + | 123 | Mobile element protein |
| - | 633 | Mobile element protein |
| - | 408 | Mobile element protein |

| | | |
|---|---|---|
| - | 291 | Mobile element protein |
| - | 210 | Mobile element protein |
| - | 96 | Mobile element protein |
| + | 486 | Mobile element protein |
| + | 156 | Mobile element protein |
| + | 753 | Mobile element protein |
| + | 312 | Mobile element protein |
| - | 486 | Mobile element protein |
| + | 159 | Mobile element protein |
| + | 165 | Mobile element protein |
| + | 237 | Mobile element protein |
| + | 180 | Mobile element protein |
| + | 168 | Mobile element protein |
| - | 117 | Mobile element protein |
| - | 126 | Mobile element protein |
| - | 120 | Mobile element protein |
| - | 156 | Mobile element protein |
| - | 159 | Mobile element protein |
| - | 192 | Mobile element protein |
| - | 156 | Mobile element protein |
| - | 120 | Mobile element protein |
| - | 132 | Mobile element protein |
| - | 135 | Mobile element protein |
| - | 264 | Mobile element protein |
| + | 216 | Mobile element protein |
| + | 315 | Mobile element protein |
| + | 183 | Mobile element protein |
| - | 129 | Mobile element protein |
| - | 138 | Mobile element protein |
| - | 138 | Mobile element protein |
| - | 156 | Mobile element protein |
| - | 177 | Mobile element protein |
| - | 168 | Mobile element protein |
| - | 135 | Mobile element protein |
| - | 753 | Mobile element protein |
| - | 690 | Mobile element protein |
| - | 195 | Mobile element protein |
| - | 132 | Mobile element protein |
| - | 135 | Mobile element protein |
| - | 243 | Mobile element protein |
| - | 183 | Mobile element protein |
| - | 315 | Mobile element protein |

| | | |
|---|---|---|
| - | 216 | Mobile element protein |
| - | 228 | Mobile element protein |
| - | 261 | Mobile element protein |
| - | 186 | Mobile element protein |
| + | 207 | Mobile element protein |
| + | 690 | Mobile element protein |
| - | 198 | Mobile element protein |
| - | 114 | Mobile element protein |
| + | 249 | Mobile element protein |
| + | 204 | Mobile element protein |
| - | 132 | Mobile element protein |
| - | 135 | Mobile element protein |
| - | 243 | Mobile element protein |
| - | 141 | Mobile element protein |
| - | 486 | Mobile element protein |
| + | 114 | Mobile element protein |
| - | 129 | Mobile element protein |
| - | 396 | Mobile element protein |
| - | 783 | Mobile element protein |
| - | 192 | Mobile element protein |
| - | 228 | Mobile element protein |
| - | 156 | Mobile element protein |
| - | 177 | Mobile element protein |
| - | 228 | Mobile element protein |
| - | 261 | Mobile element protein |
| - | 240 | Mobile element protein |
| + | 249 | Mobile element protein |
| + | 204 | Mobile element protein |
| - | 1200 | Mobile element protein |
| + | 117 | Mobile element protein |
| + | 135 | Mobile element protein |
| + | 132 | Mobile element protein |
| - | 336 | Mobile element protein |
| + | 705 | Mobile element protein |
| - | 987 | Mobile element protein |
| + | 153 | Mobile element protein |
| + | 177 | Mobile element protein |
| + | 207 | Mobile element protein |
| + | 690 | Mobile element protein |
| - | 783 | Mobile element protein |
| - | 192 | Mobile element protein |
| - | 783 | Mobile element protein |

| | | |
|---|---|---|
| - | 192 | Mobile element protein |
| - | 1185 | Mobile element protein |
| - | 753 | Mobile element protein |
| + | 135 | Mobile element protein |
| + | 132 | Mobile element protein |
| - | 435 | Mobile element protein |
| + | 192 | Mobile element protein |
| + | 783 | Mobile element protein |
| - | 753 | Mobile element protein |
| - | 783 | Mobile element protein |
| - | 192 | Mobile element protein |
| - | 132 | Mobile element protein |
| - | 135 | Mobile element protein |
| - | 117 | Mobile element protein |
| - | 165 | Mobile element protein |
| - | 297 | Mobile element protein |
| + | 120 | Mobile element protein |
| - | 255 | Mobile element protein |
| + | 264 | Mobile element protein |
| + | 135 | Mobile element protein |
| + | 132 | Mobile element protein |
| + | 633 | Mobile element protein |
| - | 1200 | Mobile element protein |
| - | 192 | Mobile element protein |
| - | 660 | Mobile element protein |
| + | 753 | Mobile element protein |
| - | 450 | Mobile element protein |
| + | 153 | Mobile element protein |
| - | 279 | Mobile element protein |
| + | 207 | Mobile element protein |
| + | 690 | Mobile element protein |
| + | 477 | Mobile element protein |
| + | 192 | Mobile element protein |
| + | 783 | Mobile element protein |
| + | 90 | Mobile element protein |
| + | 225 | Mobile element protein |
| + | 162 | Mobile element protein |
| + | 228 | Mobile element protein |
| + | 306 | Mobile element protein |
| - | 696 | Mobile element protein |
| + | 600 | Mobile element protein |
| - | 783 | Mobile element protein |

| | | |
|---|---|---|
| - | 192 | Mobile element protein |
| - | 273 | Mobile element protein |
| - | 291 | Mobile element protein |
| + | 549 | Mobile element protein |
| + | 243 | Mobile element protein |
| + | 135 | Mobile element protein |
| + | 207 | Mobile element protein |
| + | 690 | Mobile element protein |
| - | 177 | Mobile element protein |
| - | 441 | Mobile element protein |
| - | 207 | Mobile element protein |
| + | 192 | Mobile element protein |
| + | 489 | Mobile element protein |
| + | 336 | Mobile element protein |
| + | 420 | Mobile element protein |
| - | 135 | Mobile element protein |
| - | 117 | Mobile element protein |
| + | 354 | Mobile element protein |
| + | 120 | Mobile element protein |
| + | 279 | Mobile element protein |
| + | 486 | Mobile element protein |
| - | 201 | Mobile element protein |
| + | 210 | Mobile element protein |
| + | 291 | Mobile element protein |
| + | 231 | Mobile element protein |
| + | 738 | Mobile element protein |
| + | 192 | Mobile element protein |
| + | 783 | Mobile element protein |
| - | 252 | Mobile element protein |
| - | 405 | Mobile element protein |
| - | 186 | Mobile element protein |
| - | 690 | Mobile element protein |
| - | 207 | Mobile element protein |
| - | 399 | Mobile element protein |
| - | 573 | Mobile element protein |
| + | 165 | Mobile element protein |
| + | 147 | Mobile element protein |
| + | 510 | Mobile element protein |
| + | 153 | Mobile element protein |
| + | 192 | Mobile element protein |
| + | 783 | Mobile element protein |
| - | 165 | Mobile element protein |

| | | |
|---|---|---|
| + | 207 | Mobile element protein |
| + | 690 | Mobile element protein |
| + | 753 | Mobile element protein |
| + | 120 | Mobile element protein |
| - | 147 | Mobile element protein |
| - | 165 | Mobile element protein |
| + | 138 | Mobile element protein |
| + | 192 | Mobile element protein |
| + | 783 | Mobile element protein |
| + | 114 | Mobile element protein |
| + | 198 | Mobile element protein |
| - | 219 | Mobile element protein |
| - | 300 | Mobile element protein |
| + | 1200 | Mobile element protein |
| - | 477 | Mobile element protein |
| - | 954 | Mobile element protein |
| - | 183 | Mobile element protein |
| - | 1245 | Mobile element protein |
| - | 783 | Mobile element protein |
| - | 192 | Mobile element protein |
| + | 510 | Mobile element protein |
| - | 213 | Mobile element protein |
| + | 1200 | Mobile element protein |
| + | 189 | Mobile element protein |
| - | 120 | Mobile element protein |
| - | 120 | Mobile element protein |
| - | 123 | Mobile element protein |
| - | 405 | Mobile element protein |
| + | 96 | Mobile element protein |
| + | 138 | Mobile element protein |
| + | 153 | Mobile element protein |
| + | 291 | Mobile element protein |
| - | 135 | Mobile element protein |
| - | 243 | Mobile element protein |
| + | 261 | Mobile element protein |
| + | 228 | Mobile element protein |
| + | 171 | Mobile element protein |
| - | 135 | Mobile element protein |
| - | 783 | Mobile element protein |
| - | 192 | Mobile element protein |
| - | 354 | Mobile element protein |
| - | 147 | Mobile element protein |

| | | |
|---|---|---|
| - | 141 | Mobile element protein |
| - | 801 | Mobile element protein |
| + | 243 | Mobile element protein |
| + | 219 | Mobile element protein |
| + | 822 | Mobile element protein |
| - | 360 | Mobile element protein |
| - | 399 | Mobile element protein |
| + | 1152 | Mobile element protein |
| + | 138 | Mobile element protein |

| Subsystem |
| --- |
| - none - |
| - none - |
| - none - |
| - none - |
| - none - |
| - none - |
| - none - |
| - none - |
| - none - |
| - none - |
| - |
| - none - |
| - none - |
| - none - |
| - none - |
| - none - |
| - none - |
| - none - |
| - none - |
| - none - |
| - none - |
| - none - |
| - none - |
| - none - |
| - none - |
| - none - |
| - none - |
| - none - |
| - none - |
| - none - |
| - none - |
| - none - |
| - none - |
| - none - |
| - none - |
| - none - |
| - none - |
| - none - |
| - none - |

- none -
- none -
- none -
- none -
- none -
- none -
- none -
- none -
- none -
- none -
- none -
- none -
- none -
- none -
- none -
- none -
- none -
- none -
- none -
- none -
- none -
- none -
- none -
- none -
- none -
- none -
- none -
- none -
- none -
- none -
- none -
- none -
- none -
- none -
- none -
- none -
- none -
- none -
- none -
- none -
- none -
- none -

- none -
- none -
- none -
- none -
- none -
- none -
- none -
- none -
- none -
- none -
- none -
- none -
- none -
- none -
- none -
- none -
- none -
- none -
- none -
- none -
- none -
- none -
- none -
- none -
- none -
- none -
- none -
- none -
- none -
- none -
- none -
- none -
- none -
- none -
- none -
- none -
- none -
- none -
- none -
- none -
- none -
- none -

- none -
- none -
- none -
- none -
- none -
- none -
- none -
- none -
- none -
- none -
- none -
- none -
- none -
- none -
- none -
- none -
- none -
- none -
- none -
- none -
- none -
- none -
- none -
- none -
- none -
- none -
- none -
- none -
- none -
- none -
- none -
- none -
- none -
- none -
- none -
- none -
- none -
- none -
- none -
- none -
- none -
- none -

- none -
- none -
- none -
- none -
- none -
- none -
- none -
- none -
- none -
- none -
- none -
- none -
- none -
- none -
- none -
- none -
- none -
- none -
- none -
- none -
- none -
- none -
- none -
- none -
- none -
- none -
- none -
- none -
- none -
- none -
- none -
- none -
- none -
- none -
- none -
- none -
- none -
- none -
- none -
- none -
- none -
- none -

- none -
- none -
- none -
- none -
- none -
- none -
- none -
- none -
- none -
- none -
- none -
- none -
- none -
- none -
- none -
- none -
- none -
- none -
- none -
- none -
- none -
- none -
- none -
- none -
- none -
- none -
- none -
- none -
- none -
- none -
- none -
- none -
- none -
- none -
- none -
- none -
- none -
- none -
- none -
- none -
- none -
- none -

- none -
- none -
- none -
- none -
- none -
- none -
- none -
- none -
- none -
- none -
- none -
- none -
- none -
- none -
- none -
- none -
- none -
- none -
- none -
- none -
- none -
- none -
- none -
- none -
- none -
- none -
- none -
- none -
- none -
- none -
- none -
- none -
- none -
- none -
- none -
- none -
- none -
- none -
- none -
- none -
- none -
- none -

- none -
- none -
- none -
- none -
- none -
- none -
- none -
- none -
- none -
- none -
- none -
- none -
- none -
- none -
- none -
- none -
- none -
- none -
- none -
- none -
- none -
- none -
- none -
- none -
- none -
- none -
- none -
- none -
- none -
- none -
- none -
- none -
- none -
- none -
- none -
- none -
- none -
- none -
- none -
- none -
- none -
- none -

- none -
- none -
- none -
- none -
- none -
- none -
- none -
- none -
- none -
- none -
- none -
- none -
- none -
- none -
- none -
- none -
- none -
- none -
- none -
- none -
- none -
- none -
- none -
- none -
- none -
- none -
- none -
- none -
- none -
- none -
- none -
- none -
- none -
- none -
- none -
- none -
- none -
- none -
- none -
- none -
- none -
- none -

- none -
- none -
- none -
- none -
- none -
- none -
- none -
- none -
- none -
- none -
- none -
- none -
- none -
- none -
- none -
- none -
- none -
- none -
- none -
- none -
- none -
- none -
- none -
- none -
- none -
- none -
- none -
- none -
- none -
- none -
- none -
- none -
- none -
- none -
- none -
- none -
- none -
- none -
- none -
- none -
- none -
- none -

- none -
- none -
- none -
- none -
- none -
- none -
- none -
- none -
- none -
- none -
- none -
- none -
- none -
- none -
- none -
- none -
- none -
- none -
- none -
- none -
- none -
- none -
- none -
- none -
- none -
- none -
- none -
- none -
- none -
- none -
- none -
- none -
- none -
- none -
- none -
- none -
- none -
- none -
- none -
- none -
- none -
- none -

- none -
- none -
- none -
- none -
- none -
- none -
- none -
- none -
- none -
- none -
- none -
- none -
- none -
- none -
- none -
- none -
- none -
- none -
- none -
- none -
- none -
- none -
- none -
- none -
- none -
- none -
- none -
- none -
- none -
- none -
- none -
- none -
- none -
- none -
- none -
- none -
- none -
- none -
- none -
- none -
- none -
- none -

- none -
- none -
- none -
- none -
- none -
- none -
- none -
- none -
- none -
- none -
- none -
- none -
- none -
- none -
- none -
- none -
- none -
- none -
- none -
- none -
- none -
- none -
- none -
- none -
- none -
- none -
- none -
- none -
- none -
- none -
- none -
- none -
- none -
- none -
- none -
- none -
- none -
- none -
- none -
- none -
- none -
- none -

- none -
- none -
- none -
- none -
- none -
- none -
- none -
- none -
- none -
- none -
- none -
- none -
- none -
- none -
- none -
- none -
- none -
- none -
- none -
- none -
- none -
- none -
- none -
- none -
- none -
- none -
- none -
- none -
- none -
- none -
- none -
- none -
- none -
- none -
- none -
- none -
- none -
- none -
- none -
- none -
- none -
- none -

- none -
- none -
- none -
-
-
- none -
- none -
- none -
- none -
- none -
- none -
- none -
- none -
- none -
- none -
- none -
- none -
- none -
- none -
- none -
- none -
- none -
- none -
- none -
- none -
- none -
- none -
- none -
- none -
- none -
- none -
- none -
- none -
- none -
- none -
- none -
- none -
- none -
- none -
- none -
- none -
- none -

- none -
- none -
- none -
- none -
- none -
- none -
- none -
- none -
- none -
- none -
- none -
- none -
- none -
- none -
- none -
- none -
- none -
- none -
- none -
- none -
- none -
- none -
- none -
- none -
- none -
- none -
- none -
- none -
- none -
- none -
- none -
- none -
- none -
- none -
- none -
- none -
- none -
- none -
- none -
- none -
- none -
- none -

- none -
- none -
- none -
- none -
- none -
- none -
- none -
- none -
- none -
- none -
- none -
- none -
- none -
- none -
- none -
- none -
- none -
- none -
- none -
- none -
- none -
- none -
- none -
- none -
- none -
- none -
- none -
- none -
- none -
- none -
- none -
- none -
- none -
- none -
- none -
- none -
- none -
- none -
- none -
- none -
- none -
- none -

- none -
- none -
- none -
- none -
- none -
- none -
- none -
- none -
- none -
- none -
- none -
- none -
- none -
- none -
- none -
- none -
- none -
- none -
- none -
- none -
- none -
- none -
- none -
- none -
- none -
- none -
- none -
- none -
- none -
- none -
- none -
- none -
- none -
- none -
- none -
- none -
- none -
- none -
- none -
- none -
- none -
- none -

- none -
- none -
- none -
- none -
- none -
- none -
- none -
- none -
- none -
- none -
- none -
- none -
- none -
- none -
- none -
- none -
- none -
- none -
- none -
- none -
- none -
- none -
- none -
- none -
- none -
- none -
- none -
- none -
- none -
- none -
- none -
- none -
- none -
- none -
- none -
- none -
- none -
- none -
- none -
- none -
- none -
- none -

- none -
- none -
- none -
- none -
- none -
- none -
- none -
- none -
- none -
- none -
- none -
- none -
- none -
- none -
- none -
- none -
- none -
- none -
- none -
- none -
- none -
- none -
- none -
- none -
- none -
- none -
- none -
- none -
- none -
- none -
- none -
- none -
- none -
- none -
- none -
- none -
- none -
- none -
- none -
- none -
- none -
- none -

- none -
- none -
- none -
- none -
- none -
- none -
- none -
- none -
- none -
- none -
- none -
- none -
- none -
- none -
- none -
- none -
- none -
- none -
- none -
- none -
- none -
- none -
- none -
- none -
- none -
- none -
- none -
- none -
- none -
- none -
- none -
- none -
- none -
- none -
- none -
- none -
- none -
- none -
- none -
- none -
- none -
- none -

- none -
- none -
- none -
- none -
- none -
- none -
- none -
- none -
- none -
- none -
- none -
- none -
- none -
- none -
- none -
- none -
- none -
- none -
- none -
- none -
- none -
- none -
- none -
- none -
- none -
- none -
- none -
- none -
- none -
- none -
- none -
- none -
- none -
- none -
- none -
- none -
- none -
- none -
- none -
- none -
- none -
- none -

- none -
- none -
- none -
- none -
- none -
- none -
- none -
- none -
- none -
- none -
- none -
- none -
- none -
- none -
- none -
- none -
- none -
- none -
- none -
- none -
- none -
- none -
- none -
- none -
- none -
- none -
- none -
- none -
- none -
- none -
- none -
- none -
- none -
- none -
- none -
- none -
- none -
- none -
- none -
- none -
- none -
- none -

- none -
- none -
- none -
- none -
- none -
- none -
- none -
- none -
- none -
- none -
- none -
- none -
- none -
- none -
- none -
- none -
- none -
- none -
- none -
- none -
- none -
- none -
- none -
- none -
- none -
-
-
- none -
- none -
- none -
- none -
- none -
- none -
- none -
- none -
- none -
- none -
- none -
- none -
- none -
- none -
- none -

- none -
- none -
- none -
- none -
- none -
- none -
- none -
- none -
- none -
- none -
- none -
- none -
- none -
- none -
- none -
- none -
- none -
- none -
- none -
- none -
- none -
- none -
- none -
- none -
- none -
- none -
- none -
- none -
- none -
- none -
- none -
- none -
- none -
- none -
- none -
- none -
- none -
- none -
- none -
- none -
- none -
- none -

- none -
- none -
- none -
- none -
- none -
- none -
- none -
- none -
- none -
- none -
- none -
- none -
- none -
- none -
- none -
- none -
- none -
- none -
- none -
- none -
- none -
- none -
- none -
- none -
- none -
- none -
- none -
- none -
- none -
- none -
- none -
- none -
- none -
- none -
- none -
- none -
- none -
- none -
- none -
- none -
- none -
- none -

- none -
- none -
- none -
- none -
- none -
- none -
- none -
- none -
- none -
- none -
- none -
- none -
- none -
- none -
- none -
- none -
- none -
- none -
- none -
- none -
- none -
- none -
- none -
- none -
- none -
- none -
- none -
- none -
- none -
- none -
- none -
- none -
- none -
- none -
- none -
- none -
- none -
- none -
- none -
- none -
- none -
- none -

- none -
- none -
- none -
- none -
- none -
- none -
- none -
- none -
- none -
- none -
- none -
- none -
- none -
- none -
- none -
- none -
- none -
- none -
- none -
- none -
- none -
- none -
- none -
- none -
- none -
- none -
- none -
- none -
- none -
- none -
- none -
- none -
- none -
- none -
- none -
- none -
- none -
- none -
- none -
- none -
- none -
- none -

- none -
- none -
- none -
- none -
- none -
- none -
- none -
- none -
- none -
- none -
- none -
- none -
- none -
- none -
- none -
- none -
- none -
- none -
- none -
- none -
- none -
- none -
- none -
- none -
- none -
- none -
- none -
- none -
- none -
- none -
- none -
- none -
- none -
- none -
- none -
- none -
- none -
- none -
- none -
- none -
- none -
- none -

- none -
- none -
- none -
- none -
- none -
- none -
- none -
- none -
- none -
- none -
- none -
- none -
- none -
- none -
- none -
- none -
- none -
- none -
- none -
- none -
- none -
- none -
- none -
- none -
- none -
- none -
- none -
- none -
- none -
- none -
- none -
- none -
- none -
- none -
- none -
- none -
- none -
- none -
- none -
- none -
- none -
- none -

- none -
- none -
- none -
- none -
- none -
- none -
- none -
- none -
- none -
- none -
- none -
- none -
- none -
- none -
- none -
- none -
- none -
- none -
- none -
- none -
- none -
- none -
- none -
- none -
- none -
- none -
- none -
- none -
- none -
- none -
- none -
- none -
- none -
- none -
- none -
- none -
- none -
- none -
- none -
- none -
- none -
- none -

- none -
- none -
- none -
- none -
- none -
- none -
- none -
- none -
- none -
- none -
- none -
- none -
- none -
- none -
- none -
- none -
- none -
- none -
- none -
- none -
- none -
- none -
- none -
- none -
- none -
- none -
- none -
- none -
- none -
- none -
- none -
- none -
- none -
- none -
- none -
- none -
- none -
- none -
- none -
- none -
- none -
- none -

- none -
- none -
- none -
- none -
- none -
- none -
- none -
- none -
- none -
- none -
- none -
- none -
- none -
- none -
- none -
- none -
- none -
- none -
- none -
- none -
- none -
- none -
- none -
- none -
- none -
- none -
- none -
- none -
- none -
- none -
- none -
- none -
- none -
- none -
- none -
- none -
- none -
- none -
- none -
- none -
- none -
- none -

- none -
- none -
- none -
- none -
- none -
- none -
- none -
- none -
- none -
- none -
- none -
- none -
- none -
- none -
- none -
- none -
- none -
- none -
- none -
- none -
- none -
- none -
- none -
- none -
- none -
- none -
- none -
- none -
- none -
- none -
- none -
- none -
- none -
- none -
- none -
- none -
- none -
- none -
- none -
- none -
- none -
- none -

- none -
- none -
- none -
- none -
- none -
- none -
- none -
- none -
- none -
- none -
- none -
- none -
- none -
- none -
- none -
- none -
- none -
- none -
- none -
- none -
- none -
- none -
- none -
- none -
- none -
- none -
- none -
- none -
- none -
- none -
- none -
- none -
- none -
- none -
- none -
- none -
- none -
- none -
- none -
- none -
- none -
- none -

- none -
- none -
- none -
- none -
- none -
- none -
- none -
- none -
- none -
- none -
- none -
- none -
- none -
- none -
- none -
- none -
- none -
- none -
- none -
- none -
- none -
- none -
- none -
- none -
- none -
- none -
- none -
- none -
- none -
- none -
- none -
- none -
- none -
- none -
- none -
- none -
- none -
- none -
- none -
- none -
- none -
- none -

- none -
- none -
- none -
- none -
- none -
- none -
- none -
- none -
- none -
- none -
- none -
- none -
- none -
- none -
- none -
- none -
- none -
- none -
- none -
- none -
- none -
- none -
- none -
- none -
- none -
- none -
- none -
- none -
- none -
- none -
- none -
- none -
- none -
- none -
- none -
- none -
- none -
- none -
- none -
- none -
- none -
- none -

| |
|---|
| - none - |
| - none - |
| - none - |
| - none - |
| - none - |
| - none - |
| - none - |
| - none - |
| - none - |

**Table S9. List of potential prophage region detected using PHASTER.** The color indicated

| genome | Region | Region Length | Completeness | Score | # Total Proteins |
|---|---|---|---|---|---|
| wBm | - | - | - | - | - |
| wBp | - | - | - | - | - |
| wCtub | - | - | - | - | - |
| wCauA | 1 | 61Kb | intact | 150 | 51 |
| | 2 | 44.5Kb | intact | 150 | 50 |
| | 3 | 10.9Kb | questionable | 70 | 9 |
| | 4 | 15Kb | intact | 100 | 18 |
| | 5 | 17.1Kb | questionable | 90 | 22 |
| wCfeJ | - | - | - | - | - |
| wCfeT | 1 | 40.5Kb | intact | 150 | 46 |
| wCle | 1 | 7.6Kb | intact | 140 | 14 |
| | 2 | 6.7Kb | questionable | 80 | 11 |
| wDimm | - | - | - | - | - |
| wDcau | - | - | - | - | - |
| wFol | 1 | 36.7Kb | intact | 140 | 31 |
| | 2 | 54.5Kb | intact | 150 | 54 |
| | 3 | 21.4Kb | questionable | 90 | 19 |
| | 4 | 9.3Kb | questionable | 70 | 7 |
| | 5 | 31.7Kb | intact | 120 | 25 |
| | 6 | 44.1Kb | intact | 150 | 49 |
| | 7 | 59.7Kb | intact | 150 | 56 |
| wLbra* | - | - | - | - | - |
| wLsig | - | - | - | - | - |
| wLug* | wLug scaffold1 | 28.3Kb | incomplete | 40 | 14 |
| | wLug scaffold1 | 8.8Kb | incomplete | 20 | 8 |
| | wLug scaffold1 | 26.6Kb | questionable | 80 | 18 |
| | wLug scaffold1 | 15.9Kb | incomplete | 40 | 12 |
| | wLug scaffold2 | 8.9Kb | incomplete | 60 | 12 |
| | wLug scaffold2 | 41.1Kb | intact | 150 | 51 |
| wMel | 1 | 37.2Kb | intact | 110 | 23 |
| | 2 | 9.9Kb | questionable | 70 | 14 |
| | 3 | 10.2Kb | incomplete | 50 | 10 |
| | 4 | 16.1Kb | questionable | 80 | 18 |
| wMhie* | - | - | - | - | - |
| wNfla* | 1 | 22.2Kb | intact | 140 | 30 |
| | 2 | 36.4Kb | intact | 100 | 23 |
| | 3 | 15.7Kb | incomplete | 30 | 17 |
| | 4 | 12.3Kb | incomplete | 50 | 17 |
| | 5 | 11.5Kb | incomplete | 50 | 12 |

| | | | | | |
|---|---|---|---|---|---|
| | 6 | 6.6Kb | incomplete | 60 | 10 |
| wOo | - | - | - | - | - |
| wOv | - | - | - | - | - |
| wPpe* | 1 | 11.9Kb | incomplete | 50 | 11 |
| wPip | 1 | 21Kb | questionable | 80 | 24 |
| | 2 | 23.9Kb | questionable | 80 | 24 |
| | 3 | 38Kb | questionable | 90 | 22 |
| | 4 | 40.1Kb | questionable | 80 | 21 |
| | 5 | 22.3Kb | incomplete | 60 | 24 |
| | 6 | 42.7Kb | intact | 150 | 47 |
| wstri* | wstri scaffold1 | 7.5Kb | incomplete | 50 | 13 |
| | wstri scaffold1 | 14.8Kb | incomplete | 50 | 23 |
| | wstri scaffold1 | 61.3Kb | intact | 150 | 72 |
| | wstri scaffold1 | 17.6Kb | incomplete | 60 | 15 |
| | wstri scaffold1 | 18.1Kb | questionable | 70 | 29 |
| | wstri scaffold2 | 59.3Kb | intact | 150 | 59 |
| | wstri scaffold2 | 10.7Kb | incomplete | 20 | 16 |
| wTpre | 1 | 8Kb | incomplete | 30 | 9 |
| wVulC* | wVulC_contig01 | 9.4Kb | questionable | 90 | 18 |
| | wVulC_contig02 | 17.3Kb | intact | 110 | 22 |
| | wVulC_contig04 | 15.6Kb | incomplete | 40 | 11 |
| | wVulC_contig05 | 23.2Kb | questionable | 90 | 24 |
| | wVulC_contig05 | 6.3Kb | incomplete | 40 | 12 |
| | wVulC_contig06 | 6.6Kb | incomplete | 50 | 9 |
| | wVulC_contig08 | 23.3Kb | incomplete | 40 | 22 |
| | wVulC_contig09 | 9.1Kb | incomplete | 60 | 13 |
| | wVulC_contig10 | 34.4Kb | intact | 110 | 31 |
| | wVulC_contig10 | 45.1Kb | intact | 150 | 58 |
| wWb* | - | - | - | - | - |

the completeness of the prophage region: red for incomplet, blue for questionable and green for int

| Region Position | Most Common Phage | GC % |
|---|---|---|
| - | - | - |
| - | - | - |
| - | - | - |
| 245484-306575 info_outline | PHAGE_Escher_vB_EcoM_ep3_NC_025430(10) | 33.85% |
| 305256-349803 info_outline | PHAGE_Escher_vB_EcoM_ECO1230_10_NC_027995(12) | 35.62% |
| 732730-743690 info_outline | PHAGE_Paenib_Tripp_NC_028930(2) | 36.32% |
| 999801-1014803 info_outline | PHAGE_Escher_vB_EcoM_ep3_NC_025430(5) | 37.24% |
| 1077288-1094462 info_outline | PHAGE_Escher_vB_EcoM_ep3_NC_025430(5) | 36.39% |
| - | - | - |
| 1388834-1429333 info_outline | PHAGE_Escher_vB_EcoM_ECO1230_10_NC_027995(9) | 36.41% |
| 487551-495178 info_outline | PHAGE_Paenib_Tripp_NC_028930(2) | 37.01% |
| 857941-864678 info_outline | PHAGE_Paenib_Tripp_NC_028930(2) | 35.53% |
| - | - | - |
| - | - | - |
| 368051-404817 info_outline | PHAGE_Escher_vB_EcoM_ECO1230_10_NC_027995(6) | 34.83% |
| 648544-703067 info_outline | PHAGE_Escher_vB_EcoM_ep3_NC_025430(11) | 35.28% |
| 709069-730486 info_outline | PHAGE_Escher_vB_EcoM_ep3_NC_025430(5) | 35.30% |
| 805472-814846 info_outline | PHAGE_Caulob_swift_NC_019411(2) | 35.64% |
| 1056324-1088090 info_outline | PHAGE_Escher_vB_EcoM_ECO1230_10_NC_027995(6) | 34.66% |
| 1115459-1159579 info_outline | PHAGE_Escher_vB_EcoM_ECO1230_10_NC_027995(11) | 35.73% |
| 1200316-1260099 info_outline | PHAGE_Escher_vB_EcoM_ECO1230_10_NC_027995(10) | 34.55% |
| - | - | - |
| - | - | - |
| 40679-69013 info_outline | PHAGE_Vibrio_VP58.5_NC_027981(4) | 35.46% |
| 437781-446632 info_outline | PHAGE_Shigel_SfIV_NC_022749(2) | 36.60% |
| 840736-867413 info_outline | PHAGE_Lactob_phiPYB5_NC_027982(1) | 31.43% |
| 1068306-1084226 info_outline | PHAGE_Escher_vB_EcoM_ep3_NC_025430(5) | 35.03% |
| 302917-311874 info_outline | PHAGE_Escher_RCS47_NC_042128(2) | 36.81% |
| 319077-360264 info_outline | PHAGE_Escher_vB_EcoM_ep3_NC_025430(9) | 35.44% |
| 236286-273546 info_outline | PHAGE_Escher_vB_EcoM_ep3_NC_025430(4) | 36.34% |
| 548107-558081 info_outline | PHAGE_Pseudo_PPpW_3_NC_023006(6) | 34.10% |
| 560258-570525 info_outline | PHAGE_Paenib_Tripp_NC_028930(2) | 36.38% |
| 626320-642425 info_outline | PHAGE_Escher_vB_EcoM_ep3_NC_025430(5) | 35.04% |
| - | - | - |
| 25289-47574 info_outline | PHAGE_Escher_vB_EcoM_ep3_NC_025430(9) | 37.02% |
| 134523-171016 info_outline | PHAGE_Escher_vB_EcoM_ep3_NC_025430(6) | 35.76% |
| 192739-208472 info_outline | PHAGE_Vibrio_VP58.5_NC_027981(4) | 35.34% |
| 301484-313844 info_outline | PHAGE_Pseudo_PPpW_3_NC_023006(3) | 33.58% |
| 934607-946146 info_outline | PHAGE_Escher_vB_EcoM_ep3_NC_025430(4) | 34.55% |

| | | |
|---|---|---|
| 985436-992091 info_outline | PHAGE_Escher_vB_EcoM_ep3_NC_025430(3) | 34.83% |
| - | - | - |
| - | - | - |
| 12404-24395 info_outline | PHAGE_Agroba_Atu_ph07_NC_042013(2) | 32.81% |
| 247653-268692 info_outline | PHAGE_Vibrio_vB_VpaM_MAR | 35.29% |
| 317593-341555 info_outline | PHAGE_Vibrio_vB_VpaM_MAR | 35.33% |
| 337503-375565 info_outline | PHAGE_Vibrio_vB_VpaM_MAR | 34.77% |
| 424334-464512 info_outline | PHAGE_Vibrio_vB_VpaM_MAR | 34.17% |
| 465011-487365 info_outline | PHAGE_Yersin_413C | 35.14% |
| 1373737-1416457 info_outline | PHAGE_Vibrio_vB_VpaM_MAR | 35.06% |
| 82896-90494 info_outline | PHAGE_Vibrio_vB_VpaM_MAR_NC_019722(2) | 36.10% |
| 94364-109251 info_outline | PHAGE_Escher_vB_EcoM_ECOO78_NC_041926(5) | 35.73% |
| 233933-295249 info_outline | PHAGE_Escher_vB_EcoM_ep3_NC_025430(8) | 33.75% |
| 1288808-1306494 info_outline | PHAGE_Escher_vB_EcoM_ep3_NC_025430(5) | 29.58% |
| 1581451-1599644 info_outline | PHAGE_Escher_vB_EcoM_ECOO78_NC_041926(5) | 36.45% |
| 5544-64915 info_outline | PHAGE_Escher_vB_EcoM_ep3_NC_025430(10) | 34.96% |
| 97729-108512 info_outline | PHAGE_Escher_vB_EcoM_ECOO78_NC_041926(5) | 34.46% |
| 660724-668752 info_outline | PHAGE_Cellul_phi17:1_NC_021795(1) | 36.48% |
| 119159-128605 info_outline | PHAGE_Bacill_PfEFR_5_NC_031055(1) | 34.02% |
| 8417-25727 info_outline | PHAGE_Escher_vB_EcoM_ep3_NC_025430(6) | 34.58% |
| 21780-37451 info_outline | PHAGE_Vibrio_vB_VpaM_MAR_NC_019722(3) | 35.99% |
| 85659-108896 info_outline | PHAGE_Escher_vB_EcoM_ep3_NC_025430(4) | 36.34% |
| 188660-194994 info_outline | PHAGE_Prochl_P_HM2_NC_015284(1) | 35.41% |
| 2027-8648 info_outline | PHAGE_Entero_Arya_NC_031048(5) | 35.46% |
| 121973-145317 info_outline | PHAGE_Bacill_G_NC_023719(2) | 34.44% |
| 26-9201 info_outline | PHAGE_Entero_Arya_NC_031048(5) | 37.02% |
| 189796-224236 info_outline | PHAGE_Shigel_SfIV_NC_022749(2) | 34.28% |
| 227250-272417 info_outline | PHAGE_Escher_vB_EcoM_ep3_NC_025430(9) | 35.37% |
| - | - | - |

tact. The draft genome are indicated by a star.

**Table S10. List of phage-like genes detectd by RAST pipeline.**

| genome | Feature ID | Type | Contig | Start |
|---|---|---|---|---|
| wBm | fig\|292805.13.peg.289 | CDS | wBm_reseq_reversed | 194277 |
| | fig\|292805.13.peg.290 | CDS | wBm_reseq_reversed | 194440 |
| | fig\|292805.13.peg.292 | CDS | wBm_reseq_reversed | 195156 |
| | fig\|292805.13.peg.392 | CDS | wBm_reseq_reversed | 264389 |
| | fig\|292805.13.peg.393 | CDS | wBm_reseq_reversed | 264678 |
| | fig\|292805.13.peg.551 | CDS | wBm_reseq_reversed | 376109 |
| | fig\|292805.13.peg.829 | CDS | wBm_reseq_reversed | 557961 |
| | fig\|292805.13.peg.996 | CDS | wBm_reseq_reversed | 673208 |
| | fig\|292805.13.peg.998 | CDS | wBm_reseq_reversed | 673924 |
| | fig\|292805.13.peg.999 | CDS | wBm_reseq_reversed | 674087 |
| | fig\|292805.13.peg.1217 | CDS | wBm_reseq_reversed | 813513 |
| | fig\|292805.13.peg.1276 | CDS | wBm_reseq_reversed | 854793 |
| | fig\|292805.13.peg.1277 | CDS | wBm_reseq_reversed | 854956 |
| | fig\|292805.13.peg.1279 | CDS | wBm_reseq_reversed | 855672 |
| wBp | fig\|96495.5.peg.513 | CDS | NZ_CP050521.1 | 419977 |
| | fig\|96495.5.peg.514 | CDS | NZ_CP050521.1 | 420080 |
| | fig\|96495.5.peg.648 | CDS | NZ_CP050521.1 | 532215 |
| | fig\|96495.5.peg.649 | CDS | NZ_CP050521.1 | 532318 |
| | fig\|96495.5.peg.944 | CDS | NZ_CP050521.1 | 776704 |
| | fig\|96495.5.peg.945 | CDS | NZ_CP050521.1 | 776807 |
| | fig\|96495.5.peg.1033 | CDS | NZ_CP050521.1 | 846814 |
| | fig\|96495.5.peg.1277 | CDS | NZ_CP050521.1 | 1048705 |
| wCauA | fig\|2591635.5.peg.10 | CDS | CP041215.1 | 8020 |
| | fig\|2591635.5.peg.11 | CDS | CP041215.1 | 8386 |
| | fig\|2591635.5.peg.15 | CDS | CP041215.1 | 12227 |
| | fig\|2591635.5.peg.106 | CDS | CP041215.1 | 97850 |
| | fig\|2591635.5.peg.278 | CDS | CP041215.1 | 254553 |
| | fig\|2591635.5.peg.279 | CDS | CP041215.1 | 255968 |
| | fig\|2591635.5.peg.280 | CDS | CP041215.1 | 257077 |
| | fig\|2591635.5.peg.281 | CDS | CP041215.1 | 257471 |
| | fig\|2591635.5.peg.283 | CDS | CP041215.1 | 258779 |
| | fig\|2591635.5.peg.284 | CDS | CP041215.1 | 259297 |
| | fig\|2591635.5.peg.285 | CDS | CP041215.1 | 259757 |
| | fig\|2591635.5.peg.287 | CDS | CP041215.1 | 260503 |
| | fig\|2591635.5.peg.288 | CDS | CP041215.1 | 260841 |
| | fig\|2591635.5.peg.289 | CDS | CP041215.1 | 261920 |
| | fig\|2591635.5.peg.292 | CDS | CP041215.1 | 264470 |
| | fig\|2591635.5.peg.293 | CDS | CP041215.1 | 264903 |
| | fig\|2591635.5.peg.321 | CDS | CP041215.1 | 295196 |

| | | | |
|---|---|---|---|
| fig\|2591635.5.peg.322 | CDS | CP041215.1 | 295406 |
| fig\|2591635.5.peg.323 | CDS | CP041215.1 | 295756 |
| fig\|2591635.5.peg.324 | CDS | CP041215.1 | 298044 |
| fig\|2591635.5.peg.325 | CDS | CP041215.1 | 298414 |
| fig\|2591635.5.peg.326 | CDS | CP041215.1 | 298832 |
| fig\|2591635.5.peg.327 | CDS | CP041215.1 | 299881 |
| fig\|2591635.5.peg.338 | CDS | CP041215.1 | 310107 |
| fig\|2591635.5.peg.339 | CDS | CP041215.1 | 310507 |
| fig\|2591635.5.peg.340 | CDS | CP041215.1 | 312691 |
| fig\|2591635.5.peg.343 | CDS | CP041215.1 | 313572 |
| fig\|2591635.5.peg.344 | CDS | CP041215.1 | 314745 |
| fig\|2591635.5.peg.358 | CDS | CP041215.1 | 327588 |
| fig\|2591635.5.peg.361 | CDS | CP041215.1 | 331475 |
| fig\|2591635.5.peg.363 | CDS | CP041215.1 | 333458 |
| fig\|2591635.5.peg.365 | CDS | CP041215.1 | 335513 |
| fig\|2591635.5.peg.366 | CDS | CP041215.1 | 336919 |
| fig\|2591635.5.peg.367 | CDS | CP041215.1 | 338021 |
| fig\|2591635.5.peg.368 | CDS | CP041215.1 | 338422 |
| fig\|2591635.5.peg.371 | CDS | CP041215.1 | 341227 |
| fig\|2591635.5.peg.372 | CDS | CP041215.1 | 341446 |
| fig\|2591635.5.peg.373 | CDS | CP041215.1 | 341940 |
| fig\|2591635.5.peg.374 | CDS | CP041215.1 | 342400 |
| fig\|2591635.5.peg.377 | CDS | CP041215.1 | 344222 |
| fig\|2591635.5.peg.378 | CDS | CP041215.1 | 344551 |
| fig\|2591635.5.peg.379 | CDS | CP041215.1 | 345336 |
| fig\|2591635.5.peg.382 | CDS | CP041215.1 | 347885 |
| fig\|2591635.5.peg.383 | CDS | CP041215.1 | 348273 |
| fig\|2591635.5.peg.944 | CDS | CP041215.1 | 879470 |
| fig\|2591635.5.peg.1058 | CDS | CP041215.1 | 997233 |
| fig\|2591635.5.peg.1059 | CDS | CP041215.1 | 997707 |
| fig\|2591635.5.peg.1061 | CDS | CP041215.1 | 999801 |
| fig\|2591635.5.peg.1064 | CDS | CP041215.1 | 1002917 |
| fig\|2591635.5.peg.1065 | CDS | CP041215.1 | 1004335 |
| fig\|2591635.5.peg.1066 | CDS | CP041215.1 | 1005440 |
| fig\|2591635.5.peg.1067 | CDS | CP041215.1 | 1005837 |
| fig\|2591635.5.peg.1069 | CDS | CP041215.1 | 1006926 |
| fig\|2591635.5.peg.1070 | CDS | CP041215.1 | 1007404 |
| fig\|2591635.5.peg.1071 | CDS | CP041215.1 | 1007864 |
| fig\|2591635.5.peg.1073 | CDS | CP041215.1 | 1008588 |
| fig\|2591635.5.peg.1074 | CDS | CP041215.1 | 1008933 |
| fig\|2591635.5.peg.1075 | CDS | CP041215.1 | 1009737 |
| fig\|2591635.5.peg.1082 | CDS | CP041215.1 | 1017033 |

| | | | | |
|---|---|---|---|---|
| | fig\|2591635.5.peg.1084 | CDS | CP041215.1 | 1019792 |
| | fig\|2591635.5.peg.1118 | CDS | CP041215.1 | 1057192 |
| | fig\|2591635.5.peg.1137 | CDS | CP041215.1 | 1073915 |
| | fig\|2591635.5.peg.1139 | CDS | CP041215.1 | 1075755 |
| | fig\|2591635.5.peg.1141 | CDS | CP041215.1 | 1077738 |
| | fig\|2591635.5.peg.1145 | CDS | CP041215.1 | 1080548 |
| | fig\|2591635.5.peg.1146 | CDS | CP041215.1 | 1081957 |
| | fig\|2591635.5.peg.1147 | CDS | CP041215.1 | 1083047 |
| | fig\|2591635.5.peg.1148 | CDS | CP041215.1 | 1083507 |
| | fig\|2591635.5.peg.1151 | CDS | CP041215.1 | 1087067 |
| | fig\|2591635.5.peg.1152 | CDS | CP041215.1 | 1087366 |
| | fig\|2591635.5.peg.1153 | CDS | CP041215.1 | 1087826 |
| | fig\|2591635.5.peg.1155 | CDS | CP041215.1 | 1088572 |
| | fig\|2591635.5.peg.1156 | CDS | CP041215.1 | 1088910 |
| | fig\|2591635.5.peg.1157 | CDS | CP041215.1 | 1089989 |
| | fig\|2591635.5.peg.1160 | CDS | CP041215.1 | 1092539 |
| | fig\|2591635.5.peg.1161 | CDS | CP041215.1 | 1092972 |
| | fig\|2591635.5.peg.1259 | CDS | CP041215.1 | 1199236 |
| | fig\|2591635.5.peg.1342 | CDS | CP041215.1 | 1274393 |
| | fig\|2591635.5.peg.1343 | CDS | CP041215.1 | 1274670 |
| wCtub | - | - | - | - |
| wCfeJ | fig\|212123.3.peg.171 | CDS | CP051157.1 | 159434 |
| | fig\|212123.3.peg.172 | CDS | CP051157.1 | 160027 |
| | fig\|212123.3.peg.293 | CDS | CP051157.1 | 301523 |
| | fig\|212123.3.peg.498 | CDS | CP051157.1 | 511005 |
| | fig\|212123.3.peg.780 | CDS | CP051157.1 | 788877 |
| | fig\|212123.3.peg.792 | CDS | CP051157.1 | 801831 |
| | fig\|212123.3.peg.825 | CDS | CP051157.1 | 837780 |
| | fig\|212123.3.peg.914 | CDS | CP051157.1 | 942443 |
| | fig\|212123.3.peg.916 | CDS | CP051157.1 | 945050 |
| | fig\|212123.3.peg.986 | CDS | CP051157.1 | 1015314 |
| | fig\|212123.3.peg.1022 | CDS | CP051157.1 | 1057430 |
| | fig\|212123.3.peg.1146 | CDS | CP051157.1 | 1171611 |
| wCfeT | fig\|212123.4.peg.375 | CDS | CP051156.1 | 371618 |
| | fig\|212123.4.peg.386 | CDS | CP051156.1 | 385298 |
| | fig\|212123.4.peg.387 | CDS | CP051156.1 | 387529 |
| | fig\|212123.4.peg.399 | CDS | CP051156.1 | 398722 |
| | fig\|212123.4.peg.705 | CDS | CP051156.1 | 707133 |
| | fig\|212123.4.peg.1240 | CDS | CP051156.1 | 1243041 |
| | fig\|212123.4.peg.1241 | CDS | CP051156.1 | 1244232 |
| | fig\|212123.4.peg.1298 | CDS | CP051156.1 | 1300553 |
| | fig\|212123.4.peg.1407 | CDS | CP051156.1 | 1391897 |

| | | | | |
|---|---|---|---|---|
| | fig\|212123.4.peg.1408 | CDS | CP051156.1 | 1392229 |
| | fig\|212123.4.peg.1410 | CDS | CP051156.1 | 1393058 |
| | fig\|212123.4.peg.1411 | CDS | CP051156.1 | 1393521 |
| | fig\|212123.4.peg.1421 | CDS | CP051156.1 | 1405308 |
| | fig\|212123.4.peg.1423 | CDS | CP051156.1 | 1407318 |
| | fig\|212123.4.peg.1425 | CDS | CP051156.1 | 1409394 |
| | fig\|212123.4.peg.1427 | CDS | CP051156.1 | 1411608 |
| | fig\|212123.4.peg.1428 | CDS | CP051156.1 | 1413020 |
| | fig\|212123.4.peg.1429 | CDS | CP051156.1 | 1414218 |
| | fig\|212123.4.peg.1430 | CDS | CP051156.1 | 1414595 |
| | fig\|212123.4.peg.1432 | CDS | CP051156.1 | 1416003 |
| | fig\|212123.4.peg.1440 | CDS | CP051156.1 | 1423315 |
| | fig\|212123.4.peg.1441 | CDS | CP051156.1 | 1424564 |
| | fig\|212123.4.peg.1442 | CDS | CP051156.1 | 1425111 |
| | fig\|212123.4.peg.1443 | CDS | CP051156.1 | 1425478 |
| | fig\|212123.4.peg.1444 | CDS | CP051156.1 | 1427823 |
| | fig\|212123.4.peg.1445 | CDS | CP051156.1 | 1428176 |
| | fig\|212123.4.peg.1446 | CDS | CP051156.1 | 1428392 |
| wCle | fig\|246273.9.peg.151 | CDS | wCle_NZ_AP013028 | 144250 |
| | fig\|246273.9.peg.169 | CDS | wCle_NZ_AP013028 | 158182 |
| | fig\|246273.9.peg.299 | CDS | wCle_NZ_AP013028 | 262640 |
| | fig\|246273.9.peg.306 | CDS | wCle_NZ_AP013028 | 268534 |
| | fig\|246273.9.peg.307 | CDS | wCle_NZ_AP013028 | 269365 |
| | fig\|246273.9.peg.313 | CDS | wCle_NZ_AP013028 | 274012 |
| | fig\|246273.9.peg.532 | CDS | wCle_NZ_AP013028 | 481043 |
| | fig\|246273.9.peg.698 | CDS | wCle_NZ_AP013028 | 620688 |
| | fig\|246273.9.peg.703 | CDS | wCle_NZ_AP013028 | 625421 |
| | fig\|246273.9.peg.759 | CDS | wCle_NZ_AP013028 | 676109 |
| | fig\|246273.9.peg.920 | CDS | wCle_NZ_AP013028 | 816276 |
| | fig\|246273.9.peg.921 | CDS | wCle_NZ_AP013028 | 816422 |
| | fig\|246273.9.peg.922 | CDS | wCle_NZ_AP013028 | 817421 |
| | fig\|246273.9.peg.948 | CDS | wCle_NZ_AP013028 | 836503 |
| | fig\|246273.9.peg.1081 | CDS | wCle_NZ_AP013028 | 948026 |
| | fig\|246273.9.peg.1231 | CDS | wCle_NZ_AP013028 | 1089572 |
| | fig\|246273.9.peg.1245 | CDS | wCle_NZ_AP013028 | 1099171 |
| | fig\|246273.9.peg.1247 | CDS | wCle_NZ_AP013028 | 1101100 |
| | fig\|246273.9.peg.1372 | CDS | wCle_NZ_AP013028 | 1214854 |
| | fig\|246273.9.peg.1373 | CDS | wCle_NZ_AP013028 | 1215789 |
| wDimm | - | - | - | - |
| wDcau | - | - | - | - |
| wFol | fig\|169402.10.peg.230 | CDS | wFol_NZ_CP015510 | 240820 |
| | fig\|169402.10.peg.300 | CDS | wFol_NZ_CP015510 | 318830 |

| | | | |
|---|---|---|---|
| fig\|169402.10.peg.365 | CDS | wFol_NZ_CP015510 | 385331 |
| fig\|169402.10.peg.372 | CDS | wFol_NZ_CP015510 | 390734 |
| fig\|169402.10.peg.373 | CDS | wFol_NZ_CP015510 | 391928 |
| fig\|169402.10.peg.403 | CDS | wFol_NZ_CP015510 | 419737 |
| fig\|169402.10.peg.404 | CDS | wFol_NZ_CP015510 | 420173 |
| fig\|169402.10.peg.604 | CDS | wFol_NZ_CP015510 | 659568 |
| fig\|169402.10.peg.605 | CDS | wFol_NZ_CP015510 | 659921 |
| fig\|169402.10.peg.606 | CDS | wFol_NZ_CP015510 | 662251 |
| fig\|169402.10.peg.610 | CDS | wFol_NZ_CP015510 | 663740 |
| fig\|169402.10.peg.611 | CDS | wFol_NZ_CP015510 | 664934 |
| fig\|169402.10.peg.632 | CDS | wFol_NZ_CP015510 | 682428 |
| fig\|169402.10.peg.636 | CDS | wFol_NZ_CP015510 | 686563 |
| fig\|169402.10.peg.640 | CDS | wFol_NZ_CP015510 | 689334 |
| fig\|169402.10.peg.643 | CDS | wFol_NZ_CP015510 | 692331 |
| fig\|169402.10.peg.645 | CDS | wFol_NZ_CP015510 | 693723 |
| fig\|169402.10.peg.649 | CDS | wFol_NZ_CP015510 | 695417 |
| fig\|169402.10.peg.650 | CDS | wFol_NZ_CP015510 | 695755 |
| fig\|169402.10.peg.668 | CDS | wFol_NZ_CP015510 | 712200 |
| fig\|169402.10.peg.672 | CDS | wFol_NZ_CP015510 | 716312 |
| fig\|169402.10.peg.674 | CDS | wFol_NZ_CP015510 | 718546 |
| fig\|169402.10.peg.677 | CDS | wFol_NZ_CP015510 | 721560 |
| fig\|169402.10.peg.679 | CDS | wFol_NZ_CP015510 | 722992 |
| fig\|169402.10.peg.683 | CDS | wFol_NZ_CP015510 | 724685 |
| fig\|169402.10.peg.689 | CDS | wFol_NZ_CP015510 | 728834 |
| fig\|169402.10.peg.730 | CDS | wFol_NZ_CP015510 | 786766 |
| fig\|169402.10.peg.757 | CDS | wFol_NZ_CP015510 | 807757 |
| fig\|169402.10.peg.1007 | CDS | wFol_NZ_CP015510 | 1068113 |
| fig\|169402.10.peg.1008 | CDS | wFol_NZ_CP015510 | 1068466 |
| fig\|169402.10.peg.1009 | CDS | wFol_NZ_CP015510 | 1070796 |
| fig\|169402.10.peg.1012 | CDS | wFol_NZ_CP015510 | 1074008 |
| fig\|169402.10.peg.1013 | CDS | wFol_NZ_CP015510 | 1075221 |
| fig\|169402.10.peg.1045 | CDS | wFol_NZ_CP015510 | 1119089 |
| fig\|169402.10.peg.1046 | CDS | wFol_NZ_CP015510 | 1119427 |
| fig\|169402.10.peg.1050 | CDS | wFol_NZ_CP015510 | 1121124 |
| fig\|169402.10.peg.1052 | CDS | wFol_NZ_CP015510 | 1122530 |
| fig\|169402.10.peg.1055 | CDS | wFol_NZ_CP015510 | 1125436 |
| fig\|169402.10.peg.1058 | CDS | wFol_NZ_CP015510 | 1128534 |
| fig\|169402.10.peg.1062 | CDS | wFol_NZ_CP015510 | 1132659 |
| fig\|169402.10.peg.1084 | CDS | wFol_NZ_CP015510 | 1153660 |
| fig\|169402.10.peg.1085 | CDS | wFol_NZ_CP015510 | 1154888 |
| fig\|169402.10.peg.1088 | CDS | wFol_NZ_CP015510 | 1155802 |
| fig\|169402.10.peg.1089 | CDS | wFol_NZ_CP015510 | 1158075 |

| | | | | |
|---|---|---|---|---|
| | fig\|169402.10.peg.1090 | CDS | wFol_NZ_CP015510 | 1158428 |
| | fig\|169402.10.peg.1132 | CDS | wFol_NZ_CP015510 | 1201478 |
| | fig\|169402.10.peg.1133 | CDS | wFol_NZ_CP015510 | 1201822 |
| | fig\|169402.10.peg.1136 | CDS | wFol_NZ_CP015510 | 1205158 |
| | fig\|169402.10.peg.1137 | CDS | wFol_NZ_CP015510 | 1206384 |
| | fig\|169402.10.peg.1144 | CDS | wFol_NZ_CP015510 | 1222334 |
| | fig\|169402.10.peg.1150 | CDS | wFol_NZ_CP015510 | 1227862 |
| | fig\|169402.10.peg.1173 | CDS | wFol_NZ_CP015510 | 1247975 |
| | fig\|169402.10.peg.1179 | CDS | wFol_NZ_CP015510 | 1252124 |
| | fig\|169402.10.peg.1198 | CDS | wFol_NZ_CP015510 | 1291238 |
| | fig\|169402.10.peg.1417 | CDS | wFol_NZ_CP015510 | 1523771 |
| wMel | fig\|163164.1.peg.234 | CDS | wMel_NC_002978 | 248608 |
| | fig\|163164.1.peg.236 | CDS | wMel_NC_002978 | 250515 |
| | fig\|163164.1.peg.242 | CDS | wMel_NC_002978 | 258344 |
| | fig\|163164.1.peg.244 | CDS | wMel_NC_002978 | 260904 |
| | fig\|163164.1.peg.246 | CDS | wMel_NC_002978 | 261229 |
| | fig\|163164.1.peg.249 | CDS | wMel_NC_002978 | 262919 |
| | fig\|163164.1.peg.250 | CDS | wMel_NC_002978 | 263248 |
| | fig\|163164.1.peg.252 | CDS | wMel_NC_002978 | 265391 |
| | fig\|163164.1.peg.253 | CDS | wMel_NC_002978 | 266255 |
| | fig\|163164.1.peg.254 | CDS | wMel_NC_002978 | 266878 |
| | fig\|163164.1.peg.255 | CDS | wMel_NC_002978 | 267309 |
| | fig\|163164.1.peg.258 | CDS | wMel_NC_002978 | 271048 |
| | fig\|163164.1.peg.259 | CDS | wMel_NC_002978 | 273255 |
| | fig\|163164.1.peg.339 | CDS | wMel_NC_002978 | 361426 |
| | fig\|163164.1.peg.401 | CDS | wMel_NC_002978 | 431961 |
| | fig\|163164.1.peg.409 | CDS | wMel_NC_002978 | 440219 |
| | fig\|163164.1.peg.504 | CDS | wMel_NC_002978 | 552598 |
| | fig\|163164.1.peg.505 | CDS | wMel_NC_002978 | 552808 |
| | fig\|163164.1.peg.506 | CDS | wMel_NC_002978 | 553155 |
| | fig\|163164.1.peg.507 | CDS | wMel_NC_002978 | 555194 |
| | fig\|163164.1.peg.511 | CDS | wMel_NC_002978 | 556883 |
| | fig\|163164.1.peg.528 | CDS | wMel_NC_002978 | 572259 |
| | fig\|163164.1.peg.530 | CDS | wMel_NC_002978 | 573982 |
| | fig\|163164.1.peg.531 | CDS | wMel_NC_002978 | 575460 |
| | fig\|163164.1.peg.535 | CDS | wMel_NC_002978 | 579610 |
| | fig\|163164.1.peg.537 | CDS | wMel_NC_002978 | 581154 |
| | fig\|163164.1.peg.563 | CDS | wMel_NC_002978 | 623343 |
| | fig\|163164.1.peg.564 | CDS | wMel_NC_002978 | 628119 |
| | fig\|163164.1.peg.565 | CDS | wMel_NC_002978 | 628553 |
| | fig\|163164.1.peg.566 | CDS | wMel_NC_002978 | 629024 |
| | fig\|163164.1.peg.567 | CDS | wMel_NC_002978 | 629676 |

| | | | | |
|---|---|---|---|---|
| | fig\|163164.1.peg.569 | CDS | wMel_NC_002978 | 631903 |
| | fig\|163164.1.peg.570 | CDS | wMel_NC_002978 | 632232 |
| | fig\|163164.1.peg.572 | CDS | wMel_NC_002978 | 632970 |
| | fig\|163164.1.peg.574 | CDS | wMel_NC_002978 | 633879 |
| | fig\|163164.1.peg.673 | CDS | wMel_NC_002978 | 726462 |
| | fig\|163164.1.peg.903 | CDS | wMel_NC_002978 | 976186 |
| | fig\|163164.1.peg.907 | CDS | wMel_NC_002978 | 980557 |
| | fig\|163164.1.peg.1014 | CDS | wMel_NC_002978 | 1082174 |
| | fig\|163164.1.peg.1029 | CDS | wMel_NC_002978 | 1098113 |
| | fig\|163164.1.peg.1219 | CDS | wMel_NC_002978 | 253852 |
| | fig\|163164.1.peg.1220 | CDS | wMel_NC_002978 | 254208 |
| | fig\|163164.1.peg.1258 | CDS | wMel_NC_002978 | 578196 |
| | fig\|163164.1.peg.1259 | CDS | wMel_NC_002978 | 579419 |
| wMhie | fig\|1241303.4.peg.37 | CDS | NODE_11105_length_1 | 168 |
| | fig\|1241303.4.peg.102 | CDS | NODE_13107_length_1 | 72 |
| | fig\|1241303.4.peg.143 | CDS | NODE_15112_length_1 | 229 |
| | fig\|1241303.4.peg.729 | CDS | NODE_3460_length_58 | 1958 |
| | fig\|1241303.4.peg.730 | CDS | NODE_3460_length_58 | 2751 |
| | fig\|1241303.4.peg.731 | CDS | NODE_3460_length_58 | 3302 |
| | fig\|1241303.4.peg.732 | CDS | NODE_3460_length_58 | 3867 |
| | fig\|1241303.4.peg.1031 | CDS | NODE_5322_length_33 | 3288 |
| | fig\|1241303.4.peg.1079 | CDS | NODE_5846_length_29 | 2739 |
| wLbra | - | - | - | - |
| wLsig | - | - | - | - |
| wLug | fig\|1335053.6.peg.2 | CDS | MUIY01000001.1 | 960 |
| | fig\|1335053.6.peg.3 | CDS | MUIY01000001.1 | 1520 |
| | fig\|1335053.6.peg.22 | CDS | MUIY01000001.1 | 26889 |
| | fig\|1335053.6.peg.25 | CDS | MUIY01000001.1 | 29282 |
| | fig\|1335053.6.peg.43 | CDS | MUIY01000001.1 | 43093 |
| | fig\|1335053.6.peg.46 | CDS | MUIY01000001.1 | 45621 |
| | fig\|1335053.6.peg.48 | CDS | MUIY01000001.1 | 47238 |
| | fig\|1335053.6.peg.50 | CDS | MUIY01000001.1 | 48805 |
| | fig\|1335053.6.peg.51 | CDS | MUIY01000001.1 | 49452 |
| | fig\|1335053.6.peg.52 | CDS | MUIY01000001.1 | 49859 |
| | fig\|1335053.6.peg.53 | CDS | MUIY01000001.1 | 50979 |
| | fig\|1335053.6.peg.54 | CDS | MUIY01000001.1 | 52235 |
| | fig\|1335053.6.peg.56 | CDS | MUIY01000001.1 | 54263 |
| | fig\|1335053.6.peg.59 | CDS | MUIY01000001.1 | 56065 |
| | fig\|1335053.6.peg.61 | CDS | MUIY01000001.1 | 58972 |
| | fig\|1335053.6.peg.64 | CDS | MUIY01000001.1 | 60923 |
| | fig\|1335053.6.peg.66 | CDS | MUIY01000001.1 | 62731 |
| | fig\|1335053.6.peg.83 | CDS | MUIY01000001.1 | 76016 |

| | | | |
|---|---|---|---|
| fig\|1335053.6.peg.85 | CDS | MUIY01000001.1 | 76383 |
| fig\|1335053.6.peg.86 | CDS | MUIY01000001.1 | 78647 |
| fig\|1335053.6.peg.87 | CDS | MUIY01000001.1 | 78991 |
| fig\|1335053.6.peg.88 | CDS | MUIY01000001.1 | 79201 |
| fig\|1335053.6.peg.127 | CDS | MUIY01000001.1 | 127284 |
| fig\|1335053.6.peg.395 | CDS | MUIY01000001.1 | 377706 |
| fig\|1335053.6.peg.397 | CDS | MUIY01000001.1 | 380184 |
| fig\|1335053.6.peg.400 | CDS | MUIY01000001.1 | 385605 |
| fig\|1335053.6.peg.460 | CDS | MUIY01000001.1 | 439214 |
| fig\|1335053.6.peg.655 | CDS | MUIY01000001.1 | 608051 |
| fig\|1335053.6.peg.656 | CDS | MUIY01000001.1 | 609049 |
| fig\|1335053.6.peg.677 | CDS | MUIY01000001.1 | 626036 |
| fig\|1335053.6.peg.678 | CDS | MUIY01000001.1 | 627493 |
| fig\|1335053.6.peg.922 | CDS | MUIY01000001.1 | 852099 |
| fig\|1335053.6.peg.1001 | CDS | MUIY01000001.1 | 939456 |
| fig\|1335053.6.peg.1022 | CDS | MUIY01000001.1 | 961963 |
| fig\|1335053.6.peg.1137 | CDS | MUIY01000001.1 | 1077559 |
| fig\|1335053.6.peg.1138 | CDS | MUIY01000001.1 | 1078344 |
| fig\|1335053.6.peg.1139 | CDS | MUIY01000001.1 | 1078682 |
| fig\|1335053.6.peg.1141 | CDS | MUIY01000001.1 | 1079411 |
| fig\|1335053.6.peg.1142 | CDS | MUIY01000001.1 | 1079871 |
| fig\|1335053.6.peg.1143 | CDS | MUIY01000001.1 | 1080371 |
| fig\|1335053.6.peg.1145 | CDS | MUIY01000001.1 | 1081237 |
| fig\|1335053.6.peg.1146 | CDS | MUIY01000001.1 | 1081658 |
| fig\|1335053.6.peg.1149 | CDS | MUIY01000001.1 | 1084198 |
| fig\|1335053.6.peg.1150 | CDS | MUIY01000001.1 | 1085259 |
| fig\|1335053.6.peg.1151 | CDS | MUIY01000001.1 | 1085597 |
| fig\|1335053.6.peg.1153 | CDS | MUIY01000001.1 | 1086314 |
| fig\|1335053.6.peg.1154 | CDS | MUIY01000001.1 | 1086774 |
| fig\|1335053.6.peg.1155 | CDS | MUIY01000001.1 | 1087274 |
| fig\|1335053.6.peg.1157 | CDS | MUIY01000001.1 | 1088582 |
| fig\|1335053.6.peg.1158 | CDS | MUIY01000001.1 | 1089029 |
| fig\|1335053.6.peg.1159 | CDS | MUIY01000001.1 | 1090079 |
| fig\|1335053.6.peg.1160 | CDS | MUIY01000001.1 | 1091326 |
| fig\|1335053.6.peg.1162 | CDS | MUIY01000001.1 | 1095487 |
| fig\|1335053.6.peg.1164 | CDS | MUIY01000001.1 | 1097451 |
| fig\|1335053.6.peg.1166 | CDS | MUIY01000001.1 | 1098055 |
| fig\|1335053.6.peg.1196 | CDS | MUIY01000001.1 | 1124794 |
| fig\|1335053.6.peg.1200 | CDS | MUIY01000001.1 | 1127909 |
| fig\|1335053.6.peg.1342 | CDS | MUIY01000002.1 | 123586 |
| fig\|1335053.6.peg.1551 | CDS | MUIY01000002.1 | 321363 |
| fig\|1335053.6.peg.1554 | CDS | MUIY01000002.1 | 323903 |

| | | | | |
|---|---|---|---|---|
| | fig\|1335053.6.peg.1555 | CDS | MUIY01000002.1 | 324964 |
| | fig\|1335053.6.peg.1556 | CDS | MUIY01000002.1 | 325302 |
| | fig\|1335053.6.peg.1559 | CDS | MUIY01000002.1 | 327092 |
| | fig\|1335053.6.peg.1560 | CDS | MUIY01000002.1 | 327552 |
| | fig\|1335053.6.peg.1561 | CDS | MUIY01000002.1 | 328052 |
| | fig\|1335053.6.peg.1563 | CDS | MUIY01000002.1 | 329458 |
| | fig\|1335053.6.peg.1564 | CDS | MUIY01000002.1 | 329905 |
| | fig\|1335053.6.peg.1565 | CDS | MUIY01000002.1 | 330955 |
| | fig\|1335053.6.peg.1566 | CDS | MUIY01000002.1 | 332193 |
| | fig\|1335053.6.peg.1567 | CDS | MUIY01000002.1 | 333904 |
| | fig\|1335053.6.peg.1568 | CDS | MUIY01000002.1 | 335186 |
| | fig\|1335053.6.peg.1571 | CDS | MUIY01000002.1 | 337137 |
| | fig\|1335053.6.peg.1573 | CDS | MUIY01000002.1 | 338945 |
| | fig\|1335053.6.peg.1593 | CDS | MUIY01000002.1 | 354335 |
| | fig\|1335053.6.peg.1594 | CDS | MUIY01000002.1 | 355496 |
| | fig\|1335053.6.peg.1595 | CDS | MUIY01000002.1 | 356085 |
| | fig\|1335053.6.peg.1597 | CDS | MUIY01000002.1 | 356452 |
| | fig\|1335053.6.peg.1598 | CDS | MUIY01000002.1 | 358760 |
| | fig\|1335053.6.peg.1599 | CDS | MUIY01000002.1 | 359104 |
| | fig\|1335053.6.peg.1600 | CDS | MUIY01000002.1 | 359314 |
| wNfla | fig\|1854759.6.peg.24 | CDS | NZ_LYUW01000001.1 | 26459 |
| | fig\|1854759.6.peg.25 | CDS | NZ_LYUW01000001.1 | 26850 |
| | fig\|1854759.6.peg.26 | CDS | NZ_LYUW01000001.1 | 29025 |
| | fig\|1854759.6.peg.28 | CDS | NZ_LYUW01000001.1 | 29907 |
| | fig\|1854759.6.peg.29 | CDS | NZ_LYUW01000001.1 | 31109 |
| | fig\|1854759.6.peg.37 | CDS | NZ_LYUW01000001.1 | 38416 |
| | fig\|1854759.6.peg.38 | CDS | NZ_LYUW01000001.1 | 39220 |
| | fig\|1854759.6.peg.39 | CDS | NZ_LYUW01000001.1 | 39565 |
| | fig\|1854759.6.peg.41 | CDS | NZ_LYUW01000001.1 | 40289 |
| | fig\|1854759.6.peg.42 | CDS | NZ_LYUW01000001.1 | 40749 |
| | fig\|1854759.6.peg.43 | CDS | NZ_LYUW01000001.1 | 41227 |
| | fig\|1854759.6.peg.45 | CDS | NZ_LYUW01000001.1 | 42628 |
| | fig\|1854759.6.peg.46 | CDS | NZ_LYUW01000001.1 | 43026 |
| | fig\|1854759.6.peg.47 | CDS | NZ_LYUW01000001.1 | 44124 |
| | fig\|1854759.6.peg.48 | CDS | NZ_LYUW01000001.1 | 45539 |
| | fig\|1854759.6.peg.50 | CDS | NZ_LYUW01000001.1 | 47574 |
| | fig\|1854759.6.peg.52 | CDS | NZ_LYUW01000001.1 | 49544 |
| | fig\|1854759.6.peg.54 | CDS | NZ_LYUW01000001.1 | 50171 |
| | fig\|1854759.6.peg.411 | CDS | NZ_LYUW01000011.1 | 22014 |
| | fig\|1854759.6.peg.419 | CDS | NZ_LYUW01000012.1 | 9233 |
| | fig\|1854759.6.peg.420 | CDS | NZ_LYUW01000012.1 | 9661 |
| | fig\|1854759.6.peg.421 | CDS | NZ_LYUW01000012.1 | 10105 |

| | | | |
|---|---|---|---|
| fig\|1854759.6.peg.422 | CDS | NZ_LYUW01000012.1 | 10236 |
| fig\|1854759.6.peg.425 | CDS | NZ_LYUW01000012.1 | 13144 |
| fig\|1854759.6.peg.426 | CDS | NZ_LYUW01000012.1 | 13944 |
| fig\|1854759.6.peg.427 | CDS | NZ_LYUW01000012.1 | 14180 |
| fig\|1854759.6.peg.430 | CDS | NZ_LYUW01000012.1 | 16183 |
| fig\|1854759.6.peg.431 | CDS | NZ_LYUW01000012.1 | 17347 |
| fig\|1854759.6.peg.436 | CDS | NZ_LYUW01000012.1 | 18831 |
| fig\|1854759.6.peg.437 | CDS | NZ_LYUW01000012.1 | 21407 |
| fig\|1854759.6.peg.473 | CDS | NZ_LYUW01000014.1 | 64 |
| fig\|1854759.6.peg.474 | CDS | NZ_LYUW01000014.1 | 970 |
| fig\|1854759.6.peg.475 | CDS | NZ_LYUW01000014.1 | 2079 |
| fig\|1854759.6.peg.476 | CDS | NZ_LYUW01000014.1 | 2488 |
| fig\|1854759.6.peg.478 | CDS | NZ_LYUW01000014.1 | 3800 |
| fig\|1854759.6.peg.494 | CDS | NZ_LYUW01000015.1 | 3321 |
| fig\|1854759.6.peg.556 | CDS | NZ_LYUW01000018.1 | 18709 |
| fig\|1854759.6.peg.557 | CDS | NZ_LYUW01000018.1 | 18919 |
| fig\|1854759.6.peg.558 | CDS | NZ_LYUW01000018.1 | 19263 |
| fig\|1854759.6.peg.729 | CDS | NZ_LYUW01000028.1 | 2729 |
| fig\|1854759.6.peg.730 | CDS | NZ_LYUW01000028.1 | 3506 |
| fig\|1854759.6.peg.788 | CDS | NZ_LYUW01000032.1 | 2447 |
| fig\|1854759.6.peg.818 | CDS | NZ_LYUW01000033.1 | 13262 |
| fig\|1854759.6.peg.819 | CDS | NZ_LYUW01000033.1 | 13473 |
| fig\|1854759.6.peg.820 | CDS | NZ_LYUW01000033.1 | 13616 |
| fig\|1854759.6.peg.1051 | CDS | NZ_LYUW01000050.1 | 5312 |
| fig\|1854759.6.peg.1054 | CDS | NZ_LYUW01000050.1 | 7870 |
| fig\|1854759.6.peg.1055 | CDS | NZ_LYUW01000050.1 | 8960 |
| fig\|1854759.6.peg.1056 | CDS | NZ_LYUW01000050.1 | 9298 |
| fig\|1854759.6.peg.1058 | CDS | NZ_LYUW01000050.1 | 10020 |
| fig\|1854759.6.peg.1059 | CDS | NZ_LYUW01000050.1 | 10534 |
| fig\|1854759.6.peg.1106 | CDS | NZ_LYUW01000055.1 | 3379 |
| fig\|1854759.6.peg.1107 | CDS | NZ_LYUW01000055.1 | 4605 |
| fig\|1854759.6.peg.1110 | CDS | NZ_LYUW01000055.1 | 5881 |
| fig\|1854759.6.peg.1112 | CDS | NZ_LYUW01000055.1 | 6239 |
| fig\|1854759.6.peg.1113 | CDS | NZ_LYUW01000055.1 | 8539 |
| fig\|1854759.6.peg.1128 | CDS | NZ_LYUW01000057.1 | 6823 |
| fig\|1854759.6.peg.1164 | CDS | NZ_LYUW01000061.1 | 3348 |
| fig\|1854759.6.peg.1165 | CDS | NZ_LYUW01000061.1 | 4155 |
| fig\|1854759.6.peg.1166 | CDS | NZ_LYUW01000061.1 | 4571 |
| fig\|1854759.6.peg.1167 | CDS | NZ_LYUW01000061.1 | 5166 |
| fig\|1854759.6.peg.1169 | CDS | NZ_LYUW01000061.1 | 5533 |
| fig\|1854759.6.peg.1257 | CDS | NZ_LYUW01000073.1 | 22 |
| fig\|1854759.6.peg.1258 | CDS | NZ_LYUW01000073.1 | 165 |

| | | | | |
|---|---|---|---|---|
| | fig\|1854759.6.peg.1259 | CDS | NZ_LYUW01000073.1 | 375 |
| | fig\|1854759.6.peg.1446 | CDS | NZ_LYUW01000114.1 | 39 |
| | fig\|1854759.6.peg.1456 | CDS | NZ_LYUW01000117.1 | 1101 |
| | fig\|1854759.6.peg.1474 | CDS | NZ_LYUW01000126.1 | 1082 |
| wOo | - | - | - | - |
| wOv | - | - | - | - |
| wPip | fig\|955.1.peg.49 | CDS | culex94e01.p1k | 11967 |
| | fig\|955.1.peg.359 | CDS | culex12a01.q1k | 2928 |
| | fig\|955.1.peg.449 | CDS | culex17f04.q1k | 4008 |
| | fig\|955.1.peg.450 | CDS | culex17f04.q1k | 4361 |
| | fig\|955.1.peg.451 | CDS | culex17f04.q1k | 6625 |
| | fig\|955.1.peg.456 | CDS | culex147f12.p1k | 1227 |
| | fig\|955.1.peg.459 | CDS | culex147f12.p1k | 2938 |
| | fig\|955.1.peg.463 | CDS | culex69g03.p1k | 1544 |
| | fig\|955.1.peg.475 | CDS | culex140f11.p1k | 691 |
| | fig\|955.1.peg.476 | CDS | culex140f11.p1k | 927 |
| | fig\|955.1.peg.480 | CDS | culex141a08.q1k | 1867 |
| | fig\|955.1.peg.511 | CDS | culex102f01.p1k | 185 |
| | fig\|955.1.peg.512 | CDS | culex102f01.p1k | 763 |
| | fig\|955.1.peg.527 | CDS | culex102f01.p1k | 17434 |
| | fig\|955.1.peg.528 | CDS | culex102f01.p1k | 18628 |
| | fig\|955.1.peg.533 | CDS | culex102f01.p1k | 20112 |
| | fig\|955.1.peg.534 | CDS | culex102f01.p1k | 22403 |
| | fig\|955.1.peg.535 | CDS | culex102f01.p1k | 22957 |
| | fig\|955.1.peg.547 | CDS | culex102f01.p1k | 33590 |
| | fig\|955.1.peg.548 | CDS | culex102f01.p1k | 33928 |
| | fig\|955.1.peg.552 | CDS | culex102f01.p1k | 35606 |
| | fig\|955.1.peg.555 | CDS | culex102f01.p1k | 37412 |
| | fig\|955.1.peg.557 | CDS | culex102f01.p1k | 40018 |
| | fig\|955.1.peg.559 | CDS | culex102f01.p1k | 42077 |
| | fig\|955.1.peg.563 | CDS | culex102f01.p1k | 45831 |
| | fig\|955.1.peg.567 | CDS | culex102f01.p1k | 50379 |
| | fig\|955.1.peg.842 | CDS | culex131h09.q1k | 1367 |
| | fig\|955.1.peg.843 | CDS | culex50h05.p1k | 6 |
| | fig\|955.1.peg.853 | CDS | culex40h12.p1k | 4714 |
| | fig\|955.1.peg.870 | CDS | culex40h12.p1k | 33536 |
| | fig\|955.1.peg.909 | CDS | culex18h04.p1k | 28 |
| | fig\|955.1.peg.910 | CDS | culex18h04.p1k | 570 |
| | fig\|955.1.peg.911 | CDS | culex18h04.p1k | 923 |
| | fig\|955.1.peg.1182 | CDS | culex79g04.p1k | 60509 |
| | fig\|955.1.peg.1317 | CDS | culex180d12.p1k | 4625 |
| | fig\|955.1.peg.1320 | CDS | culex180d12.p1k | 6408 |

| | | | | |
|---|---|---|---|---|
| | fig\|955.1.peg.1338 | CDS | culex180d12.p1k | 26226 |
| | fig\|955.1.peg.1372 | CDS | culex143c03.p1k | 16074 |
| | fig\|955.1.peg.1377 | CDS | culex143c03.p1k | 21327 |
| | fig\|955.1.peg.1378 | CDS | culex143c03.p1k | 21665 |
| | fig\|955.1.peg.1382 | CDS | culex143c03.p1k | 23331 |
| | fig\|955.1.peg.1386 | CDS | culex143c03.p1k | 25488 |
| | fig\|955.1.peg.1391 | CDS | culex143c03.p1k | 31843 |
| | fig\|955.1.peg.1395 | CDS | culex143c03.p1k | 38991 |
| | fig\|955.1.peg.1529 | CDS | culex16c12.p1k | 16431 |
| wPpe | - | - | - | - |
| wstri | fig\|368602.8.peg.1 | CDS | MUIX01000001.1 | 2 |
| | fig\|368602.8.peg.5 | CDS | MUIX01000001.1 | 6352 |
| | fig\|368602.8.peg.8 | CDS | MUIX01000001.1 | 8932 |
| | fig\|368602.8.peg.9 | CDS | MUIX01000001.1 | 10010 |
| | fig\|368602.8.peg.10 | CDS | MUIX01000001.1 | 10348 |
| | fig\|368602.8.peg.12 | CDS | MUIX01000001.1 | 10794 |
| | fig\|368602.8.peg.13 | CDS | MUIX01000001.1 | 11507 |
| | fig\|368602.8.peg.14 | CDS | MUIX01000001.1 | 12016 |
| | fig\|368602.8.peg.19 | CDS | MUIX01000001.1 | 15546 |
| | fig\|368602.8.peg.83 | CDS | MUIX01000001.1 | 76352 |
| | fig\|368602.8.peg.85 | CDS | MUIX01000001.1 | 81034 |
| | fig\|368602.8.peg.87 | CDS | MUIX01000001.1 | 82896 |
| | fig\|368602.8.peg.90 | CDS | MUIX01000001.1 | 85646 |
| | fig\|368602.8.peg.91 | CDS | MUIX01000001.1 | 87052 |
| | fig\|368602.8.peg.92 | CDS | MUIX01000001.1 | 87607 |
| | fig\|368602.8.peg.93 | CDS | MUIX01000001.1 | 88170 |
| | fig\|368602.8.peg.94 | CDS | MUIX01000001.1 | 88504 |
| | fig\|368602.8.peg.99 | CDS | MUIX01000001.1 | 91161 |
| | fig\|368602.8.peg.110 | CDS | MUIX01000001.1 | 99602 |
| | fig\|368602.8.peg.111 | CDS | MUIX01000001.1 | 99812 |
| | fig\|368602.8.peg.112 | CDS | MUIX01000001.1 | 100156 |
| | fig\|368602.8.peg.113 | CDS | MUIX01000001.1 | 102464 |
| | fig\|368602.8.peg.114 | CDS | MUIX01000001.1 | 102831 |
| | fig\|368602.8.peg.115 | CDS | MUIX01000001.1 | 103420 |
| | fig\|368602.8.peg.116 | CDS | MUIX01000001.1 | 103750 |
| | fig\|368602.8.peg.117 | CDS | MUIX01000001.1 | 104012 |
| | fig\|368602.8.peg.119 | CDS | MUIX01000001.1 | 106062 |
| | fig\|368602.8.peg.121 | CDS | MUIX01000001.1 | 108001 |
| | fig\|368602.8.peg.123 | CDS | MUIX01000001.1 | 109830 |
| | fig\|368602.8.peg.149 | CDS | MUIX01000001.1 | 133841 |
| | fig\|368602.8.peg.150 | CDS | MUIX01000001.1 | 135032 |
| | fig\|368602.8.peg.151 | CDS | MUIX01000001.1 | 135621 |

| | | | |
|---|---|---|---|
| fig\|368602.8.peg.153 | CDS | MUIX01000001.1 | 135988 |
| fig\|368602.8.peg.154 | CDS | MUIX01000001.1 | 138408 |
| fig\|368602.8.peg.155 | CDS | MUIX01000001.1 | 138752 |
| fig\|368602.8.peg.156 | CDS | MUIX01000001.1 | 138963 |
| fig\|368602.8.peg.178 | CDS | MUIX01000001.1 | 168987 |
| fig\|368602.8.peg.181 | CDS | MUIX01000001.1 | 171533 |
| fig\|368602.8.peg.182 | CDS | MUIX01000001.1 | 172611 |
| fig\|368602.8.peg.183 | CDS | MUIX01000001.1 | 172949 |
| fig\|368602.8.peg.185 | CDS | MUIX01000001.1 | 173395 |
| fig\|368602.8.peg.186 | CDS | MUIX01000001.1 | 174108 |
| fig\|368602.8.peg.187 | CDS | MUIX01000001.1 | 174617 |
| fig\|368602.8.peg.192 | CDS | MUIX01000001.1 | 178147 |
| fig\|368602.8.peg.197 | CDS | MUIX01000001.1 | 182296 |
| fig\|368602.8.peg.201 | CDS | MUIX01000001.1 | 185034 |
| fig\|368602.8.peg.203 | CDS | MUIX01000001.1 | 186998 |
| fig\|368602.8.peg.205 | CDS | MUIX01000001.1 | 188785 |
| fig\|368602.8.peg.206 | CDS | MUIX01000001.1 | 190153 |
| fig\|368602.8.peg.209 | CDS | MUIX01000001.1 | 194485 |
| fig\|368602.8.peg.212 | CDS | MUIX01000001.1 | 196557 |
| fig\|368602.8.peg.223 | CDS | MUIX01000001.1 | 214568 |
| fig\|368602.8.peg.224 | CDS | MUIX01000001.1 | 215761 |
| fig\|368602.8.peg.246 | CDS | MUIX01000001.1 | 237828 |
| fig\|368602.8.peg.249 | CDS | MUIX01000001.1 | 240360 |
| fig\|368602.8.peg.250 | CDS | MUIX01000001.1 | 241151 |
| fig\|368602.8.peg.251 | CDS | MUIX01000001.1 | 241387 |
| fig\|368602.8.peg.253 | CDS | MUIX01000001.1 | 242890 |
| fig\|368602.8.peg.256 | CDS | MUIX01000001.1 | 244691 |
| fig\|368602.8.peg.257 | CDS | MUIX01000001.1 | 245154 |
| fig\|368602.8.peg.258 | CDS | MUIX01000001.1 | 245672 |
| fig\|368602.8.peg.260 | CDS | MUIX01000001.1 | 246578 |
| fig\|368602.8.peg.265 | CDS | MUIX01000001.1 | 250380 |
| fig\|368602.8.peg.266 | CDS | MUIX01000001.1 | 250747 |
| fig\|368602.8.peg.269 | CDS | MUIX01000001.1 | 251870 |
| fig\|368602.8.peg.270 | CDS | MUIX01000001.1 | 253072 |
| fig\|368602.8.peg.291 | CDS | MUIX01000001.1 | 275913 |
| fig\|368602.8.peg.294 | CDS | MUIX01000001.1 | 278019 |
| fig\|368602.8.peg.296 | CDS | MUIX01000001.1 | 281205 |
| fig\|368602.8.peg.298 | CDS | MUIX01000001.1 | 281923 |
| fig\|368602.8.peg.299 | CDS | MUIX01000001.1 | 282383 |
| fig\|368602.8.peg.300 | CDS | MUIX01000001.1 | 282871 |
| fig\|368602.8.peg.301 | CDS | MUIX01000001.1 | 283091 |
| fig\|368602.8.peg.302 | CDS | MUIX01000001.1 | 284268 |

| | | | |
|---|---|---|---|
| fig\|368602.8.peg.303 | CDS | MUIX01000001.1 | 284662 |
| fig\|368602.8.peg.304 | CDS | MUIX01000001.1 | 285783 |
| fig\|368602.8.peg.305 | CDS | MUIX01000001.1 | 286582 |
| fig\|368602.8.peg.306 | CDS | MUIX01000001.1 | 287441 |
| fig\|368602.8.peg.307 | CDS | MUIX01000001.1 | 287835 |
| fig\|368602.8.peg.308 | CDS | MUIX01000001.1 | 288956 |
| fig\|368602.8.peg.309 | CDS | MUIX01000001.1 | 290799 |
| fig\|368602.8.peg.310 | CDS | MUIX01000001.1 | 291193 |
| fig\|368602.8.peg.311 | CDS | MUIX01000001.1 | 292314 |
| fig\|368602.8.peg.312 | CDS | MUIX01000001.1 | 293719 |
| fig\|368602.8.peg.316 | CDS | MUIX01000001.1 | 295249 |
| fig\|368602.8.peg.643 | CDS | MUIX01000001.1 | 641854 |
| fig\|368602.8.peg.645 | CDS | MUIX01000001.1 | 644368 |
| fig\|368602.8.peg.990 | CDS | MUIX01000001.1 | 981690 |
| fig\|368602.8.peg.1055 | CDS | MUIX01000001.1 | 1035843 |
| fig\|368602.8.peg.1057 | CDS | MUIX01000001.1 | 1038207 |
| fig\|368602.8.peg.1063 | CDS | MUIX01000001.1 | 1046164 |
| fig\|368602.8.peg.1317 | CDS | MUIX01000001.1 | 1293083 |
| fig\|368602.8.peg.1318 | CDS | MUIX01000001.1 | 1293226 |
| fig\|368602.8.peg.1319 | CDS | MUIX01000001.1 | 1293869 |
| fig\|368602.8.peg.1320 | CDS | MUIX01000001.1 | 1294207 |
| fig\|368602.8.peg.1322 | CDS | MUIX01000001.1 | 1294936 |
| fig\|368602.8.peg.1323 | CDS | MUIX01000001.1 | 1295396 |
| fig\|368602.8.peg.1324 | CDS | MUIX01000001.1 | 1295905 |
| fig\|368602.8.peg.1326 | CDS | MUIX01000001.1 | 1297213 |
| fig\|368602.8.peg.1327 | CDS | MUIX01000001.1 | 1297660 |
| fig\|368602.8.peg.1328 | CDS | MUIX01000001.1 | 1298113 |
| fig\|368602.8.peg.1331 | CDS | MUIX01000001.1 | 1302009 |
| fig\|368602.8.peg.1332 | CDS | MUIX01000001.1 | 1304089 |
| fig\|368602.8.peg.1334 | CDS | MUIX01000001.1 | 1305918 |
| fig\|368602.8.peg.1355 | CDS | MUIX01000001.1 | 1325332 |
| fig\|368602.8.peg.1356 | CDS | MUIX01000001.1 | 1326529 |
| fig\|368602.8.peg.1357 | CDS | MUIX01000001.1 | 1327118 |
| fig\|368602.8.peg.1359 | CDS | MUIX01000001.1 | 1327485 |
| fig\|368602.8.peg.1360 | CDS | MUIX01000001.1 | 1329793 |
| fig\|368602.8.peg.1361 | CDS | MUIX01000001.1 | 1330137 |
| fig\|368602.8.peg.1362 | CDS | MUIX01000001.1 | 1330347 |
| fig\|368602.8.peg.1386 | CDS | MUIX01000001.1 | 1367344 |
| fig\|368602.8.peg.1390 | CDS | MUIX01000001.1 | 1370517 |
| fig\|368602.8.peg.1393 | CDS | MUIX01000001.1 | 1373097 |
| fig\|368602.8.peg.1394 | CDS | MUIX01000001.1 | 1374175 |
| fig\|368602.8.peg.1395 | CDS | MUIX01000001.1 | 1374513 |

| | | | |
|---|---|---|---|
| fig\|368602.8.peg.1397 | CDS | MUIX01000001.1 | 1374959 |
| fig\|368602.8.peg.1398 | CDS | MUIX01000001.1 | 1375672 |
| fig\|368602.8.peg.1399 | CDS | MUIX01000001.1 | 1376181 |
| fig\|368602.8.peg.1525 | CDS | MUIX01000001.1 | 1490714 |
| fig\|368602.8.peg.1612 | CDS | MUIX01000001.1 | 1587343 |
| fig\|368602.8.peg.1613 | CDS | MUIX01000001.1 | 1587553 |
| fig\|368602.8.peg.1614 | CDS | MUIX01000001.1 | 1587897 |
| fig\|368602.8.peg.1615 | CDS | MUIX01000001.1 | 1590205 |
| fig\|368602.8.peg.1616 | CDS | MUIX01000001.1 | 1590572 |
| fig\|368602.8.peg.1617 | CDS | MUIX01000001.1 | 1591161 |
| fig\|368602.8.peg.1626 | CDS | MUIX01000001.1 | 1597154 |
| fig\|368602.8.peg.1627 | CDS | MUIX01000001.1 | 1597364 |
| fig\|368602.8.peg.1628 | CDS | MUIX01000001.1 | 1597708 |
| fig\|368602.8.peg.1629 | CDS | MUIX01000001.1 | 1599644 |
| fig\|368602.8.peg.1630 | CDS | MUIX01000001.1 | 1600254 |
| fig\|368602.8.peg.1632 | CDS | MUIX01000002.1 | 326 |
| fig\|368602.8.peg.1633 | CDS | MUIX01000002.1 | 661 |
| fig\|368602.8.peg.1634 | CDS | MUIX01000002.1 | 3099 |
| fig\|368602.8.peg.1635 | CDS | MUIX01000002.1 | 3334 |
| fig\|368602.8.peg.1639 | CDS | MUIX01000002.1 | 6494 |
| fig\|368602.8.peg.1640 | CDS | MUIX01000002.1 | 6705 |
| fig\|368602.8.peg.1641 | CDS | MUIX01000002.1 | 7040 |
| fig\|368602.8.peg.1642 | CDS | MUIX01000002.1 | 9478 |
| fig\|368602.8.peg.1643 | CDS | MUIX01000002.1 | 9845 |
| fig\|368602.8.peg.1644 | CDS | MUIX01000002.1 | 10398 |
| fig\|368602.8.peg.1645 | CDS | MUIX01000002.1 | 10630 |
| fig\|368602.8.peg.1669 | CDS | MUIX01000002.1 | 36941 |
| fig\|368602.8.peg.1670 | CDS | MUIX01000002.1 | 37240 |
| fig\|368602.8.peg.1673 | CDS | MUIX01000002.1 | 39766 |
| fig\|368602.8.peg.1674 | CDS | MUIX01000002.1 | 40566 |
| fig\|368602.8.peg.1675 | CDS | MUIX01000002.1 | 40664 |
| fig\|368602.8.peg.1677 | CDS | MUIX01000002.1 | 42310 |
| fig\|368602.8.peg.1680 | CDS | MUIX01000002.1 | 44111 |
| fig\|368602.8.peg.1681 | CDS | MUIX01000002.1 | 44574 |
| fig\|368602.8.peg.1682 | CDS | MUIX01000002.1 | 45092 |
| fig\|368602.8.peg.1685 | CDS | MUIX01000002.1 | 47203 |
| fig\|368602.8.peg.1687 | CDS | MUIX01000002.1 | 47903 |
| fig\|368602.8.peg.1688 | CDS | MUIX01000002.1 | 48366 |
| fig\|368602.8.peg.1689 | CDS | MUIX01000002.1 | 48539 |
| fig\|368602.8.peg.1690 | CDS | MUIX01000002.1 | 48823 |
| fig\|368602.8.peg.1692 | CDS | MUIX01000002.1 | 49795 |
| fig\|368602.8.peg.1693 | CDS | MUIX01000002.1 | 50153 |

| | | | | |
|---|---|---|---|---|
| | fig\|368602.8.peg.1694 | CDS | MUIX01000002.1 | 51579 |
| | fig\|368602.8.peg.1696 | CDS | MUIX01000002.1 | 53629 |
| | fig\|368602.8.peg.1699 | CDS | MUIX01000002.1 | 55614 |
| | fig\|368602.8.peg.1701 | CDS | MUIX01000002.1 | 58285 |
| | fig\|368602.8.peg.1703 | CDS | MUIX01000002.1 | 60114 |
| | fig\|368602.8.peg.1730 | CDS | MUIX01000002.1 | 83815 |
| | fig\|368602.8.peg.1731 | CDS | MUIX01000002.1 | 85006 |
| | fig\|368602.8.peg.1733 | CDS | MUIX01000002.1 | 86639 |
| | fig\|368602.8.peg.1735 | CDS | MUIX01000002.1 | 87006 |
| | fig\|368602.8.peg.1738 | CDS | MUIX01000002.1 | 90748 |
| | fig\|368602.8.peg.1739 | CDS | MUIX01000002.1 | 91939 |
| | fig\|368602.8.peg.1741 | CDS | MUIX01000002.1 | 93572 |
| | fig\|368602.8.peg.1743 | CDS | MUIX01000002.1 | 93939 |
| | fig\|368602.8.peg.1747 | CDS | MUIX01000002.1 | 97729 |
| | fig\|368602.8.peg.1748 | CDS | MUIX01000002.1 | 98920 |
| | fig\|368602.8.peg.1750 | CDS | MUIX01000002.1 | 100553 |
| | fig\|368602.8.peg.1752 | CDS | MUIX01000002.1 | 100920 |
| | fig\|368602.8.peg.1753 | CDS | MUIX01000002.1 | 102367 |
| | fig\|368602.8.peg.1754 | CDS | MUIX01000002.1 | 103193 |
| | fig\|368602.8.peg.1755 | CDS | MUIX01000002.1 | 103537 |
| | fig\|368602.8.peg.1756 | CDS | MUIX01000002.1 | 103747 |
| | fig\|368602.8.peg.1792 | CDS | MUIX01000002.1 | 147856 |
| | fig\|368602.8.peg.1795 | CDS | MUIX01000002.1 | 150759 |
| | fig\|368602.8.peg.1813 | CDS | MUIX01000002.1 | 171756 |
| | fig\|368602.8.peg.1814 | CDS | MUIX01000002.1 | 172947 |
| | fig\|368602.8.peg.1816 | CDS | MUIX01000002.1 | 174580 |
| | fig\|368602.8.peg.1818 | CDS | MUIX01000002.1 | 174947 |
| | fig\|368602.8.peg.1819 | CDS | MUIX01000002.1 | 177385 |
| | fig\|368602.8.peg.1820 | CDS | MUIX01000002.1 | 177720 |
| | fig\|368602.8.peg.1821 | CDS | MUIX01000002.1 | 177931 |
| wTpre | fig\|125593.4.peg.73 | CDS | NZ_CM003641.1 | 55803 |
| | fig\|125593.4.peg.74 | CDS | NZ_CM003641.1 | 56306 |
| | fig\|125593.4.peg.177 | CDS | NZ_CM003641.1 | 136658 |
| | fig\|125593.4.peg.178 | CDS | NZ_CM003641.1 | 137078 |
| | fig\|125593.4.peg.206 | CDS | NZ_CM003641.1 | 158015 |
| | fig\|125593.4.peg.217 | CDS | NZ_CM003641.1 | 164222 |
| | fig\|125593.4.peg.218 | CDS | NZ_CM003641.1 | 164840 |
| | fig\|125593.4.peg.734 | CDS | NZ_CM003641.1 | 602681 |
| | fig\|125593.4.peg.735 | CDS | NZ_CM003641.1 | 603110 |
| | fig\|125593.4.peg.736 | CDS | NZ_CM003641.1 | 603382 |
| | fig\|125593.4.peg.737 | CDS | NZ_CM003641.1 | 604108 |
| | fig\|125593.4.peg.739 | CDS | NZ_CM003641.1 | 604411 |

| | | | | |
|---|---|---|---|---|
| | fig\|125593.4.peg.740 | CDS | NZ_CM003641.1 | 604538 |
| | fig\|125593.4.peg.814 | CDS | NZ_CM003641.1 | 662026 |
| | fig\|125593.4.peg.815 | CDS | NZ_CM003641.1 | 662762 |
| | fig\|125593.4.peg.953 | CDS | NZ_CM003641.1 | 765303 |
| | fig\|125593.4.peg.954 | CDS | NZ_CM003641.1 | 765641 |
| | fig\|125593.4.peg.1339 | CDS | NZ_CM003641.1 | 1096320 |
| | fig\|125593.4.peg.1363 | CDS | NZ_CM003641.1 | 1115627 |
| | fig\|125593.4.peg.1364 | CDS | NZ_CM003641.1 | 1115961 |
| | fig\|125593.4.peg.1365 | CDS | NZ_CM003641.1 | 1116525 |
| wVulC | fig\|1220511.6.peg.28 | CDS | ALWU01000001.1 | 24120 |
| | fig\|1220511.6.peg.118 | CDS | ALWU01000001.1 | 117882 |
| | fig\|1220511.6.peg.137 | CDS | ALWU01000001.1 | 129054 |
| | fig\|1220511.6.peg.367 | CDS | ALWU01000001.1 | 313351 |
| | fig\|1220511.6.peg.703 | CDS | ALWU01000002.1 | 17665 |
| | fig\|1220511.6.peg.704 | CDS | ALWU01000002.1 | 17875 |
| | fig\|1220511.6.peg.705 | CDS | ALWU01000002.1 | 18225 |
| | fig\|1220511.6.peg.706 | CDS | ALWU01000002.1 | 20495 |
| | fig\|1220511.6.peg.707 | CDS | ALWU01000002.1 | 20865 |
| | fig\|1220511.6.peg.708 | CDS | ALWU01000002.1 | 21418 |
| | fig\|1220511.6.peg.709 | CDS | ALWU01000002.1 | 21827 |
| | fig\|1220511.6.peg.710 | CDS | ALWU01000002.1 | 22614 |
| | fig\|1220511.6.peg.727 | CDS | ALWU01000002.1 | 38746 |
| | fig\|1220511.6.peg.730 | CDS | ALWU01000002.1 | 42735 |
| | fig\|1220511.6.peg.774 | CDS | ALWU01000004.1 | 1192 |
| | fig\|1220511.6.peg.775 | CDS | ALWU01000004.1 | 1559 |
| | fig\|1220511.6.peg.776 | CDS | ALWU01000004.1 | 2140 |
| | fig\|1220511.6.peg.777 | CDS | ALWU01000004.1 | 3333 |
| | fig\|1220511.6.peg.796 | CDS | ALWU01000004.1 | 21208 |
| | fig\|1220511.6.peg.799 | CDS | ALWU01000004.1 | 25150 |
| | fig\|1220511.6.peg.802 | CDS | ALWU01000004.1 | 27108 |
| | fig\|1220511.6.peg.806 | CDS | ALWU01000004.1 | 30196 |
| | fig\|1220511.6.peg.807 | CDS | ALWU01000004.1 | 30319 |
| | fig\|1220511.6.peg.808 | CDS | ALWU01000004.1 | 31593 |
| | fig\|1220511.6.peg.809 | CDS | ALWU01000004.1 | 32692 |
| | fig\|1220511.6.peg.810 | CDS | ALWU01000004.1 | 33088 |
| | fig\|1220511.6.peg.815 | CDS | ALWU01000004.1 | 36957 |
| | fig\|1220511.6.peg.864 | CDS | ALWU01000005.1 | 42324 |
| | fig\|1220511.6.peg.911 | CDS | ALWU01000005.1 | 87955 |
| | fig\|1220511.6.peg.914 | CDS | ALWU01000005.1 | 90517 |
| | fig\|1220511.6.peg.916 | CDS | ALWU01000005.1 | 91586 |
| | fig\|1220511.6.peg.917 | CDS | ALWU01000005.1 | 91924 |
| | fig\|1220511.6.peg.919 | CDS | ALWU01000005.1 | 92669 |

| | | | |
|---|---|---|---|
| fig\|1220511.6.peg.920 | CDS | ALWU01000005.1 | 93132 |
| fig\|1220511.6.peg.921 | CDS | ALWU01000005.1 | 93646 |
| fig\|1220511.6.peg.926 | CDS | ALWU01000005.1 | 97512 |
| fig\|1220511.6.peg.927 | CDS | ALWU01000005.1 | 97908 |
| fig\|1220511.6.peg.928 | CDS | ALWU01000005.1 | 98411 |
| fig\|1220511.6.peg.930 | CDS | ALWU01000005.1 | 100660 |
| fig\|1220511.6.peg.931 | CDS | ALWU01000005.1 | 102090 |
| fig\|1220511.6.peg.934 | CDS | ALWU01000005.1 | 104621 |
| fig\|1220511.6.peg.937 | CDS | ALWU01000005.1 | 106584 |
| fig\|1220511.6.peg.939 | CDS | ALWU01000005.1 | 108371 |
| fig\|1220511.6.peg.1038 | CDS | ALWU01000005.1 | 189620 |
| fig\|1220511.6.peg.1074 | CDS | ALWU01000006.1 | 1821 |
| fig\|1220511.6.peg.1075 | CDS | ALWU01000006.1 | 2027 |
| fig\|1220511.6.peg.1076 | CDS | ALWU01000006.1 | 2538 |
| fig\|1220511.6.peg.1077 | CDS | ALWU01000006.1 | 3001 |
| fig\|1220511.6.peg.1082 | CDS | ALWU01000006.1 | 7492 |
| fig\|1220511.6.peg.1083 | CDS | ALWU01000006.1 | 7866 |
| fig\|1220511.6.peg.1084 | CDS | ALWU01000006.1 | 8648 |
| fig\|1220511.6.peg.1087 | CDS | ALWU01000006.1 | 11219 |
| fig\|1220511.6.peg.1088 | CDS | ALWU01000006.1 | 11653 |
| fig\|1220511.6.peg.1111 | CDS | ALWU01000006.1 | 31788 |
| fig\|1220511.6.peg.1192 | CDS | ALWU01000008.1 | 54805 |
| fig\|1220511.6.peg.1300 | CDS | ALWU01000008.1 | 156193 |
| fig\|1220511.6.peg.1323 | CDS | ALWU01000008.1 | 173460 |
| fig\|1220511.6.peg.1336 | CDS | ALWU01000009.1 | 805 |
| fig\|1220511.6.peg.1337 | CDS | ALWU01000009.1 | 1143 |
| fig\|1220511.6.peg.1339 | CDS | ALWU01000009.1 | 1861 |
| fig\|1220511.6.peg.1340 | CDS | ALWU01000009.1 | 2321 |
| fig\|1220511.6.peg.1341 | CDS | ALWU01000009.1 | 2817 |
| fig\|1220511.6.peg.1342 | CDS | ALWU01000009.1 | 3037 |
| fig\|1220511.6.peg.1343 | CDS | ALWU01000009.1 | 4214 |
| fig\|1220511.6.peg.1344 | CDS | ALWU01000009.1 | 4626 |
| fig\|1220511.6.peg.1345 | CDS | ALWU01000009.1 | 5748 |
| fig\|1220511.6.peg.1346 | CDS | ALWU01000009.1 | 7151 |
| fig\|1220511.6.peg.1348 | CDS | ALWU01000009.1 | 9201 |
| fig\|1220511.6.peg.1350 | CDS | ALWU01000009.1 | 11187 |
| fig\|1220511.6.peg.1352 | CDS | ALWU01000009.1 | 12998 |
| fig\|1220511.6.peg.1440 | CDS | ALWU01000010.1 | 72470 |
| fig\|1220511.6.peg.1442 | CDS | ALWU01000010.1 | 74923 |
| fig\|1220511.6.peg.1528 | CDS | ALWU01000010.1 | 147132 |
| fig\|1220511.6.peg.1529 | CDS | ALWU01000010.1 | 148295 |
| fig\|1220511.6.peg.1607 | CDS | ALWU01000010.1 | 211437 |

| | | | | |
|---|---|---|---|---|
| | fig\|1220511.6.peg.1609 | CDS | ALWU01000010.1 | 212090 |
| | fig\|1220511.6.peg.1623 | CDS | ALWU01000010.1 | 224236 |
| | fig\|1220511.6.peg.1624 | CDS | ALWU01000010.1 | 224670 |
| | fig\|1220511.6.peg.1627 | CDS | ALWU01000010.1 | 227250 |
| | fig\|1220511.6.peg.1628 | CDS | ALWU01000010.1 | 228050 |
| | fig\|1220511.6.peg.1629 | CDS | ALWU01000010.1 | 228388 |
| | fig\|1220511.6.peg.1631 | CDS | ALWU01000010.1 | 229106 |
| | fig\|1220511.6.peg.1632 | CDS | ALWU01000010.1 | 229575 |
| | fig\|1220511.6.peg.1633 | CDS | ALWU01000010.1 | 230066 |
| | fig\|1220511.6.peg.1634 | CDS | ALWU01000010.1 | 230285 |
| | fig\|1220511.6.peg.1639 | CDS | ALWU01000010.1 | 235511 |
| | fig\|1220511.6.peg.1640 | CDS | ALWU01000010.1 | 235912 |
| | fig\|1220511.6.peg.1641 | CDS | ALWU01000010.1 | 237014 |
| | fig\|1220511.6.peg.1642 | CDS | ALWU01000010.1 | 238411 |
| | fig\|1220511.6.peg.1644 | CDS | ALWU01000010.1 | 240458 |
| | fig\|1220511.6.peg.1647 | CDS | ALWU01000010.1 | 242416 |
| | fig\|1220511.6.peg.1650 | CDS | ALWU01000010.1 | 246405 |
| | fig\|1220511.6.peg.1672 | CDS | ALWU01000010.1 | 262463 |
| | fig\|1220511.6.peg.1673 | CDS | ALWU01000010.1 | 263677 |
| | fig\|1220511.6.peg.1675 | CDS | ALWU01000010.1 | 265372 |
| | fig\|1220511.6.peg.1677 | CDS | ALWU01000010.1 | 265739 |
| | fig\|1220511.6.peg.1678 | CDS | ALWU01000010.1 | 268039 |
| | fig\|1220511.6.peg.1679 | CDS | ALWU01000010.1 | 268383 |
| | fig\|1220511.6.peg.1680 | CDS | ALWU01000010.1 | 268590 |
| | fig\|1220511.6.peg.1713 | CDS | ALWU01000010.1 | 306984 |
| | fig\|1220511.6.peg.1714 | CDS | ALWU01000010.1 | 307174 |
| | fig\|1220511.6.peg.1716 | CDS | ALWU01000010.1 | 309114 |
| | fig\|1220511.6.peg.1720 | CDS | ALWU01000010.1 | 310801 |
| | fig\|1220511.6.peg.1721 | CDS | ALWU01000010.1 | 310889 |
| | fig\|1220511.6.peg.1945 | CDS | ALWU01000010.1 | 509327 |
| wWb | fig\|96496.9.peg.476 | CDS | NZ_NJBR02000028.1 | 38534 |
| | fig\|96496.9.peg.698 | CDS | NZ_NJBR02000055.1 | 97 |
| | fig\|96496.9.peg.707 | CDS | NZ_NJBR02000055.1 | 5944 |
| | fig\|96496.9.peg.952 | CDS | NZ_NJBR02000078.1 | 729 |

| Stop | Strand | Length (bp) | Function |
| --- | --- | --- | --- |
| 194104 | - | 174 | Phage uncharacterized protein |
| 194261 | - | 180 | Phage uncharacterized protein |
| 194971 | - | 186 | some similarities to phage related proteins |
| 264264 | - | 126 | Phage major capsid protein |
| 264415 | - | 264 | Phage major capsid protein |
| 376360 | + | 252 | phage major capsid protein, HK97 family |
| 557776 | - | 186 | some similarities to phage related proteins |
| 673393 | + | 186 | some similarities to phage related proteins |
| 674103 | + | 180 | Phage uncharacterized protein |
| 674260 | + | 174 | Phage uncharacterized protein |
| 813370 | - | 144 | Phage Host Specificity Protein |
| 854620 | - | 174 | Phage uncharacterized protein |
| 854777 | - | 180 | Phage uncharacterized protein |
| 855487 | - | 186 | some similarities to phage related proteins |
| 420096 | + | 120 | Phage uncharacterized protein |
| 420247 | + | 168 | Phage uncharacterized protein |
| 532048 | - | 168 | Phage uncharacterized protein |
| 532199 | - | 120 | Phage uncharacterized protein |
| 776537 | - | 168 | Phage uncharacterized protein |
| 776688 | - | 120 | Phage uncharacterized protein |
| 846689 | - | 126 | Phage major capsid protein |
| 1048911 | + | 207 | phage major capsid protein, HK97 family |
| 8319 | + | 300 | some similarities to phage related proteins |
| 9483 | + | 1098 | some similarities to phage related proteins |
| 13420 | + | 1194 | Phage portal protein |
| 96399 | - | 1452 | some similarities to phage related proteins |
| 255971 | + | 1419 | Phage head, portal protein B |
| 256999 | + | 1032 | Phage head, head-tail preconnector protease C |
| 257445 | + | 369 | Phage head, head-DNA stabilization protein D |
| 258475 | + | 1005 | Phage head, major capsid protein E |
| 259300 | + | 522 | Phage minor tail protein Z |
| 259770 | + | 474 | Phage protein |
| 260221 | + | 465 | Phage baseplate assembly protein V |
| 260838 | + | 336 | Phage baseplate assembly protein GpW |
| 261641 | + | 801 | Phage baseplate assembly protein J |
| 263086 | + | 1167 | Phage tail formation protein I |
| 264910 | + | 441 | Phage protein |
| 266393 | + | 1491 | prophage LambdaW1, site-specific recombinas |
| 294261 | - | 936 | Phage tail formation protein D |

| | | | |
|---|---|---|---|
| 295197 | - | 210 | Phage protein |
| 295409 | - | 348 | prophage P2W3, tail protein U, putative |
| 295756 | - | 2289 | Phage tail length tape-measure protein T |
| 298160 | - | 255 | Phage tail protein E |
| 298449 | - | 384 | Phage tail tube protein FII |
| 299375 | - | 507 | Phage tail tube protein FII |
| 309907 | - | 201 | prophage P2W3, tail protein X, putative |
| 310091 | - | 417 | Phage tail protein |
| 310511 | - | 2181 | Phage tail length tape-measure protein T |
| 313072 | - | 501 | Phage tail tube protein FII |
| 313594 | - | 1152 | Phage tail sheath protein FI |
| 327929 | + | 342 | Phage protein |
| 331957 | + | 483 | Phage head, terminase subunit Nu1 |
| 335284 | + | 1827 | Phage head, terminase DNA packaging protein |
| 336919 | + | 1407 | Phage head, portal protein B |
| 337953 | + | 1035 | Phage head, head-tail preconnector protease ( |
| 338389 | + | 369 | Phage head, head-DNA stabilization protein D |
| 339426 | + | 1005 | Phage head, major capsid protein E |
| 341445 | + | 219 | putative phage related protein |
| 341943 | + | 498 | Phage tail, component Z |
| 342413 | + | 474 | Phage protein |
| 342864 | + | 465 | Phage baseplate assembly protein V |
| 344548 | + | 327 | Phage baseplate assembly protein GpW |
| 345336 | + | 786 | Phage baseplate assembly protein J |
| 346496 | + | 1161 | Phage tail formation protein I |
| 348322 | + | 438 | Phage protein |
| 348758 | + | 486 | prophage LambdaW1, site-specific recombinas |
| 882796 | + | 3327 | Phage Host Specificity Protein |
| 997703 | + | 471 | Phage protein |
| 998318 | + | 612 | Phage head, terminase subunit Nu1 |
| 1001633 | + | 1833 | Phage head, terminase DNA packaging protein |
| 1004338 | + | 1422 | Phage head, portal protein B |
| 1005372 | + | 1038 | Phage head, head-tail preconnector protease ( |
| 1005811 | + | 372 | Phage head, head-DNA stabilization protein D |
| 1006529 | + | 693 | Phage head, major capsid protein E |
| 1007423 | + | 498 | Phage minor tail protein Z |
| 1007877 | + | 474 | Phage protein |
| 1008319 | + | 456 | Phage baseplate assembly protein V |
| 1008923 | + | 336 | Phage baseplate assembly protein |
| 1009730 | + | 798 | Phage baseplate assembly protein GpJ |
| 1010954 | + | 1218 | Phage tail formation protein I |
| 1018187 | + | 1155 | Phage tail sheath protein FI |

| | | | | |
|---|---|---|---|---|
| 1020298 | + | | 507 | Phage tail tube protein FII |
| 1058643 | + | | 1452 | some similarities to phage related proteins |
| 1074391 | + | | 477 | Phage protein |
| 1076237 | + | | 483 | Phage head, terminase subunit Nu1 |
| 1079564 | + | | 1827 | Phage head, terminase DNA packaging protein |
| 1081957 | + | | 1410 | Phage head, portal protein B |
| 1082988 | + | | 1032 | Phage head, head-tail preconnector protease |
| 1083502 | + | | 456 | Phage head, head-DNA stabilization protein D |
| 1084511 | + | | 1005 | Phage head, major capsid protein E |
| 1087369 | + | | 303 | Phage minor tail protein Z |
| 1087839 | + | | 474 | Phage protein |
| 1088290 | + | | 465 | Phage baseplate assembly protein V |
| 1088907 | + | | 336 | Phage baseplate assembly protein GpW |
| 1089710 | + | | 801 | Phage baseplate assembly protein J |
| 1091155 | + | | 1167 | Phage tail formation protein I |
| 1092979 | + | | 441 | Phage protein |
| 1094462 | + | | 1491 | prophage LambdaW1, site-specific recombinas |
| 1198037 | - | | 1200 | Phage major capsid protein |
| 1274629 | + | | 237 | some similarities to phage related proteins |
| 1275803 | + | | 1134 | some similarities to phage related proteins |
| - | - | - | | - |
| 159297 | - | | 138 | Phage portal protein |
| 159575 | - | | 453 | Phage portal protein |
| 300540 | - | | 984 | Phage integrase |
| 512183 | + | | 1179 | Phage portal protein |
| 790409 | + | | 1533 | some similarities to phage related proteins |
| 800350 | - | | 1482 | some similarities to phage related proteins |
| 837514 | - | | 267 | prophage LambdaW1, site-specific recombinas |
| 942817 | + | | 375 | Phage Host Specificity Protein |
| 947941 | + | | 2892 | Phage Host Specificity Protein |
| 1014112 | - | | 1203 | Phage major capsid protein |
| 1058869 | + | | 1440 | Phage tail formation protein I |
| 1171465 | - | | 147 | prophage LambdaW1, site-specific recombinas |
| 372637 | + | | 1020 | Phage portal protein |
| 387463 | + | | 2166 | Phage Host Specificity Protein |
| 388323 | + | | 795 | Phage Host Specificity Protein |
| 399882 | + | | 1161 | Phage major capsid protein |
| 708503 | + | | 1371 | Phage tail fiber protein |
| 1244219 | + | | 1179 | some similarities to phage related proteins |
| 1244375 | + | | 144 | Phage uncharacterized protein |
| 1301638 | + | | 1086 | Phage tail fiber protein |
| 1391094 | - | | 804 | Phage baseplate assembly protein GpJ |

| | | | |
|---|---|---|---|
| 1391894 | - | 336 | Phage baseplate assembly protein GpW |
| 1392597 | - | 462 | Phage baseplate assembly protein V |
| 1393048 | - | 474 | Phage protein |
| 1405820 | + | 513 | Phage protein |
| 1407797 | + | 480 | Phage head, terminase subunit Nu1 |
| 1411223 | + | 1830 | Phage head, terminase DNA packaging protein |
| 1413023 | + | 1416 | Phage head, portal protein B |
| 1414135 | + | 1116 | Phage head, head-tail preconnector protease ( |
| 1414592 | + | 375 | Phage head, head-DNA stabilization protein D |
| 1415599 | + | 1005 | Phage head, major capsid protein E |
| 1416515 | + | 513 | prophage LambdaW5, minor tail protein Z, put |
| 1424514 | + | 1200 | Phage tail sheath protein FI |
| 1425070 | + | 507 | Phage tail tube protein FII |
| 1425368 | + | 258 | Phage tail protein E |
| 1427823 | + | 2346 | Phage tail length tape-measure protein T |
| 1428179 | + | 357 | phage-related tail formation protein U |
| 1428391 | + | 216 | Phage tail protein GpX |
| 1429333 | + | 942 | Phage tail formation protein D |
| 145662 | + | 1413 | ankyrin repeat domain protein |
| 157943 | - | 240 | ankyrin repeat domain protein |
| 264079 | + | 1440 | ankyrin repeat domain protein |
| 269130 | + | 597 | Ankyrin |
| 269934 | + | 570 | prophage LambdaW1, ankyrin repeat domain p |
| 275277 | + | 1266 | ankyrin repeat domain protein |
| 480465 | - | 579 | Ankyrin |
| 621938 | + | 1251 | ankyrin repeat domain protein |
| 625684 | + | 264 | ankyrin repeat domain protein |
| 674745 | - | 1365 | ankyrin repeat domain protein |
| 815854 | - | 423 | prophage LambdaW1, ankyrin repeat domain p |
| 816273 | - | 150 | prophage LambdaW1, ankyrin repeat domain p |
| 816657 | - | 765 | Ankyrin |
| 837048 | + | 546 | ankyrin repeat domain protein |
| 948271 | + | 246 | ankyrin repeat domain protein |
| 1088325 | - | 1248 | Ankyrin repeat domain protein |
| 1099581 | + | 411 | ankyrin domain protein ank2 |
| 1101756 | + | 657 | ankyrin repeat protein, putative |
| 1214645 | - | 210 | ankyrin repeat domain protein |
| 1214950 | - | 840 | ankyrin repeat domain protein |
| - | - | - | - |
| - | - | - | - |
| 242073 | + | 1254 | Phage major capsid protein |
| 318399 | - | 432 | prophage LambdaW1, site-specific recombinas |

| | | | |
|---|---|---|---|
| 384984 | - | 348 | prophage P2W3, tail protein U, putative |
| 390228 | - | 507 | prophage P2W3, contractile tail tube protein |
| 390753 | - | 1176 | Phage tail sheath monomer |
| 418085 | - | 1653 | Phage integrase, site-specific serine recombina |
| 419721 | - | 453 | Putative bacteriophage-related protein |
| 659362 | - | 207 | prophage P2W3, tail protein X, putative |
| 659565 | - | 357 | prophage P2W3, tail protein U, putative |
| 659921 | - | 2331 | Phage tail length tape-measure protein |
| 663234 | - | 507 | prophage P2W3, contractile tail tube protein |
| 663759 | - | 1176 | Phage tail sheath monomer |
| 682916 | + | 489 | Phage protein |
| 688389 | + | 1827 | Phage terminase, large subunit |
| 690746 | + | 1413 | Phage portal protein |
| 693332 | + | 1002 | Phage protein |
| 694235 | + | 513 | prophage LambdaW5, minor tail protein Z, put |
| 695752 | + | 336 | prophage LambdaW5, baseplate assembly pro |
| 696564 | + | 810 | prophage LambdaW1, baseplate assembly pro |
| 712688 | + | 489 | Phage protein |
| 718141 | + | 1830 | Phage terminase, large subunit |
| 719958 | + | 1413 | Phage portal protein |
| 722561 | + | 1002 | Phage protein |
| 723507 | + | 516 | prophage LambdaW5, minor tail protein Z, put |
| 725020 | + | 336 | prophage LambdaW5, baseplate assembly pro |
| 730486 | + | 1653 | Phage integrase, site-specific serine recombina |
| 787959 | + | 1194 | Phage portal protein |
| 811089 | + | 3333 | Phage Host Specificity Protein |
| 1067901 | - | 213 | prophage P2W3, tail protein X, putative |
| 1068110 | - | 357 | prophage P2W3, tail protein U, putative |
| 1068466 | - | 2331 | Phage tail length tape-measure protein |
| 1073502 | - | 507 | prophage P2W3, contractile tail tube protein |
| 1074022 | - | 1200 | Phage tail sheath monomer |
| 1118280 | - | 810 | prophage LambdaW1, baseplate assembly pro |
| 1119092 | - | 336 | prophage LambdaW5, baseplate assembly pro |
| 1120609 | - | 516 | prophage LambdaW5, minor tail protein Z, put |
| 1121526 | - | 1005 | Phage protein |
| 1124030 | - | 1407 | Phage portal protein |
| 1126714 | - | 1821 | Phage terminase, large subunit |
| 1132171 | - | 489 | Phage protein |
| 1154841 | + | 1182 | Phage tail sheath monomer |
| 1155394 | + | 507 | prophage P2W3, contractile tail tube protein |
| 1158075 | + | 2274 | Phage tail length tape-measure protein |
| 1158431 | + | 357 | prophage P2W3, tail protein U, putative |

| | | | |
|---|---|---|---|
| 1158637 | + | 210 | prophage P2W3, tail protein X, putative |
| 1201272 | - | 207 | prophage P2W3, tail protein X, putative |
| 1201475 | - | 348 | prophage P2W3, tail protein U, putative |
| 1204652 | - | 507 | prophage P2W3, contractile tail tube protein |
| 1205188 | - | 1197 | Phage tail sheath monomer |
| 1221816 | - | 519 | Phage protein |
| 1227674 | - | 189 | putative phage related protein |
| 1248310 | + | 336 | prophage LambdaW5, baseplate assembly pro |
| 1253776 | + | 1653 | Phage integrase, site-specific serine recombina |
| 1289676 | - | 1563 | some similarities to phage related proteins |
| 1520121 | - | 3651 | Phage tail fiber protein |
| 249837 | + | 1230 | prophage LambdaW1, DNA methylase |
| 252341 | + | 1827 | Phage terminase, large subunit |
| 259345 | + | 1002 | elements of external origin; phage-related fun |
| 259882 | - | 1023 | prophage LambdaW1, transposase, IS110 fami |
| 261726 | + | 498 | prophage LambdaW1, minor tail protein Z, put |
| 263245 | + | 327 | prophage LambdaW1, baseplate assembly pro |
| 264057 | + | 810 | prophage LambdaW1, baseplate assembly pro |
| 265993 | + | 603 | prophage LambdaW1, ankyrin repeat domain p |
| 266845 | + | 591 | prophage LambdaW1, ankyrin repeat domain p |
| 267306 | + | 429 | Putative bacteriophage-related protein |
| 268811 | + | 1503 | prophage LambdaW1, site-specific recombinas |
| 270374 | - | 675 | prophage LambdaW1, ankyrin repeat domain p |
| 271150 | - | 2106 | prophage LambdaW1, ankyrin repeat domain p |
| 364761 | + | 3336 | Phage Host Specificity Protein |
| 431428 | - | 534 | phage prohead protease |
| 439020 | - | 1200 | Phage major capsid protein #Fam0006 |
| 551654 | - | 945 | prophage P2W3, tail protein D, putative |
| 552599 | - | 210 | prophage P2W3, tail protein X, putative |
| 552805 | - | 351 | prophage P2W3, tail protein U, putative |
| 553152 | - | 2043 | Phage tail length tape-measure protein |
| 556386 | - | 498 | prophage P2W3, contractile tail tube protein |
| 573485 | + | 1227 | prophage LambdaW4, DNA methylase |
| 575463 | + | 1482 | prophage LambdaW4, ankyrin repeat domain p |
| 577283 | + | 1824 | Phage terminase, large subunit |
| 580671 | + | 1062 | prophage LambdaW4, minor capsid protein C, |
| 582158 | + | 1005 | elements of external origin; phage-related fun |
| 626243 | + | 2901 | prophage LambdaW5, ankyrin repeat domain p |
| 626650 | - | 1470 | prophage LambdaW5, site-specific recombinas |
| 628113 | - | 441 | Putative bacteriophage-related protein |
| 628566 | - | 459 | prophage LambdaW5, ankyrin repeat domain p |
| 629050 | - | 627 | prophage LambdaW5, ankyrin repeat domain p |

| | | | |
|---|---|---|---|
| 631112 | - | 792 | prophage LambdaW5, baseplate assembly pro |
| 631906 | - | 327 | prophage LambdaW5, baseplate assembly pro |
| 632506 | - | 465 | prophage LambdaW5, baseplate assembly pro |
| 633430 | - | 450 | prophage LambdaW5, minor tail protein Z, put |
| 727391 | + | 930 | site-specific recombinase, phage integrase fam |
| 974993 | - | 1194 | Phage portal protein |
| 979094 | - | 1464 | some similarities to phage related proteins |
| 1083061 | + | 888 | Phage-related protein |
| 1099099 | + | 987 | site-specific recombinase, phage integrase fam |
| 254226 | + | 375 | Phage portal protein # N-terminal fragment |
| 255269 | + | 1062 | Phage portal protein |
| 579431 | + | 1236 | Phage portal protein |
| 579613 | + | 195 | Phage portal protein # C-terminal fragment |
| 305 | + | 138 | Phage uncharacterized protein |
| 830 | + | 759 | Phage major capsid protein |
| 498 | + | 270 | Phage protein |
| 2785 | + | 828 | Phage Host Specificity Protein |
| 3314 | + | 564 | Phage Host Specificity Protein |
| 3865 | + | 564 | Phage Host Specificity Protein |
| 5183 | + | 1317 | Phage Host Specificity Protein |
| 2809 | - | 480 | Phage protein |
| 2473 | - | 267 | Phage protein |
| - | - | - | - |
| - | - | - | - |
| 1520 | + | 561 | Phage tail length tape-measure protein |
| 1858 | + | 339 | prophage P2W3, tail protein U, putative |
| 26458 | - | 432 | Phage protein |
| 28263 | - | 1020 | Phage tail formation protein I |
| 42662 | - | 432 | Phage protein |
| 44467 | - | 1155 | Phage tail formation protein I |
| 47077 | - | 162 | Phage head, portal protein B |
| 47456 | - | 1350 | Phage head, terminase DNA packaging protein |
| 48847 | - | 606 | Phage head, major capsid protein E |
| 49485 | - | 375 | Phage head, head-DNA stabilization protein D |
| 49927 | - | 1053 | Phage head, head-tail preconnector protease |
| 50976 | - | 1260 | Phage head, portal protein B |
| 54108 | - | 156 | Phage head, portal protein B |
| 55529 | - | 537 | Phage head, terminase DNA packaging protein |
| 57611 | - | 1362 | Phage head, terminase DNA packaging protein |
| 60450 | - | 474 | Phage head, terminase subunit Nu1 |
| 62276 | - | 456 | Phage protein |
| 76273 | + | 258 | Phage tail protein E |

| | | | |
|---|---|---|---|
| 78647 | + | 2265 | Phage tail length tape-measure protein T |
| 78994 | + | 348 | phage-related tail formation protein U |
| 79200 | + | 210 | Phage tail protein GpX |
| 80145 | + | 945 | Phage tail formation protein D |
| 123955 | - | 3330 | Phage Host Specificity Protein |
| 379028 | + | 1323 | Phage tail fiber protein |
| 382616 | + | 2433 | Phage tail fiber protein |
| 387056 | + | 1452 | some similarities to phage related proteins |
| 437781 | - | 1434 | some similarities to phage related proteins |
| 609064 | + | 1014 | Phage tail fiber protein |
| 609207 | + | 159 | Phage tail fiber protein |
| 624876 | - | 1161 | Phage tail formation protein I |
| 626036 | - | 1458 | Phage baseplate assembly protein J |
| 852242 | + | 144 | prophage LambdaW1, baseplate assembly pro |
| 938293 | - | 1164 | Phage major capsid protein |
| 961646 | - | 318 | Phage protein |
| 1076399 | - | 1161 | Phage tail formation protein I |
| 1077559 | - | 786 | Phage baseplate assembly protein J |
| 1078347 | - | 336 | Phage baseplate assembly protein GpW |
| 1078947 | - | 465 | Phage baseplate assembly protein V |
| 1079398 | - | 474 | Phage protein |
| 1079868 | - | 504 | Phage minor tail protein Z |
| 1080773 | - | 465 | Phage head, major capsid protein E |
| 1081227 | - | 432 | Phage protein |
| 1083032 | - | 1167 | Phage tail formation protein I |
| 1084477 | - | 783 | Phage baseplate assembly protein J |
| 1085262 | - | 336 | Phage baseplate assembly protein GpW |
| 1085817 | - | 498 | Phage baseplate assembly protein V |
| 1086301 | - | 474 | Phage protein |
| 1086771 | - | 504 | Phage minor tail protein Z |
| 1087578 | - | 1005 | Phage head, major capsid protein E |
| 1088619 | - | 411 | Phage head, head-DNA stabilization protein D |
| 1089066 | - | 1014 | Phage head, head-tail preconnector protease |
| 1090079 | - | 1248 | Phage head, portal protein B |
| 1093199 | - | 2289 | Phage head, terminase DNA packaging protein |
| 1096987 | - | 465 | Phage head, terminase subunit Nu1 |
| 1097585 | - | 471 | Phage protein |
| 1124651 | - | 144 | prophage LambdaW1, baseplate assembly pro |
| 1127487 | - | 423 | Phage tail fiber protein |
| 122411 | - | 1176 | Phage portal protein |
| 320932 | - | 432 | Phage protein |
| 322737 | - | 1167 | Phage tail formation protein I |

| | | | |
|---|---|---|---|
| 324182 | - | 783 | Phage baseplate assembly protein J |
| 324967 | - | 336 | Phage baseplate assembly protein GpW |
| 326628 | - | 465 | Phage baseplate assembly protein V |
| 327079 | - | 474 | Phage protein |
| 327549 | - | 504 | Phage minor tail protein Z |
| 328454 | - | 1005 | Phage head, major capsid protein E |
| 329495 | - | 411 | Phage head, head-DNA stabilization protein D |
| 329942 | - | 1014 | Phage head, head-tail preconnector protease C |
| 330955 | - | 1239 | Phage head, portal protein B |
| 333434 | - | 471 | Phage head, terminase DNA packaging protein |
| 333990 | - | 1197 | Phage head, terminase DNA packaging protein |
| 336667 | - | 471 | Phage head, terminase subunit Nu1 |
| 338490 | - | 456 | Phage protein |
| 355480 | + | 1146 | Phage tail sheath protein FI |
| 356002 | + | 507 | Phage tail tube protein FII |
| 356342 | + | 258 | Phage tail protein E |
| 358758 | + | 2307 | Phage tail length tape-measure protein T |
| 359107 | + | 348 | phage-related tail formation protein U |
| 359313 | + | 210 | Phage tail protein GpX |
| 360264 | + | 951 | Phage tail formation protein D |
| 26259 | - | 201 | prophage P2W3, tail protein X, putative |
| 26443 | - | 408 | Phage tail protein |
| 26854 | - | 2172 | Phage tail length tape-measure protein T |
| 29407 | - | 501 | Phage tail tube protein FII |
| 29922 | - | 1188 | Phage tail sheath protein FI |
| 37199 | - | 1218 | Phage tail formation protein I |
| 38423 | - | 798 | Phage tail tip, host specificity protein J |
| 39230 | - | 336 | Phage baseplate assembly protein GpW |
| 39834 | - | 456 | Phage baseplate assembly protein V |
| 40276 | - | 474 | Phage protein |
| 40730 | - | 498 | Phage minor tail protein Z |
| 41624 | - | 1005 | Phage head, major capsid protein E |
| 42655 | - | 372 | Phage head, head-DNA stabilization protein D |
| 43054 | - | 1071 | Phage head, head-tail preconnector protease C |
| 44121 | - | 1419 | Phage head, portal protein B |
| 45757 | - | 1818 | Phage head, terminase DNA packaging protein |
| 49074 | - | 471 | Phage head, terminase subunit Nu1 |
| 49677 | - | 495 | Phage protein |
| 23018 | + | 1005 | some similarities to phage related proteins |
| 7737 | - | 1497 | prophage LambdaW1, site-specific recombinas |
| 9227 | - | 435 | Phage protein |
| 9704 | - | 402 | prophage LambdaW1, ankyrin repeat domain p |

| | | | |
|---|---|---|---|
| 10102 | - | 135 | prophage LambdaW1, ankyrin repeat domain [illegible] |
| 11984 | - | 1161 | Phage tail formation protein I |
| 13144 | - | 801 | Phage tail tip, host specificity protein J |
| 13947 | - | 234 | Phage baseplate assembly protein GpW |
| 17331 | + | 1149 | Phage tail sheath protein FI |
| 17853 | + | 507 | Phage tail tube protein FII |
| 21407 | + | 2577 | Phage tail length tape-measure protein T |
| 21661 | + | 255 | prophage P2W3, tail protein U, putative |
| 957 | + | 894 | Phage head, portal protein B |
| 2001 | + | 1032 | Phage head, head-tail preconnector protease [illegible] |
| 2450 | + | 372 | Phage head, head-DNA stabilization protein D |
| 3492 | + | 1005 | Phage head, major capsid protein E |
| 4285 | + | 486 | Phage minor tail protein Z |
| 3455 | + | 135 | Phage head, portal protein B |
| 17765 | - | 945 | Phage tail formation protein D |
| 18710 | - | 210 | Phage tail protein GpX |
| 18916 | - | 348 | phage-related tail formation protein U |
| 3445 | + | 717 | Phage portal protein |
| 3922 | + | 417 | Phage portal protein |
| 3646 | + | 1200 | Phage major capsid protein |
| 12318 | - | 945 | Phage tail formation protein D |
| 13264 | - | 210 | Phage tail protein GpX |
| 13470 | - | 147 | prophage P2W3, tail protein U, putative |
| 4881 | - | 432 | Phage protein |
| 6704 | - | 1167 | Phage tail formation protein I |
| 8160 | - | 801 | Phage baseplate assembly protein J |
| 8963 | - | 336 | Phage baseplate assembly protein GpW |
| 9556 | - | 465 | Phage baseplate assembly protein V |
| 10007 | - | 528 | Phage protein |
| 4593 | + | 1215 | Phage tail sheath protein FI |
| 5111 | + | 507 | Phage tail tube protein FII |
| 6138 | + | 258 | Phage tail protein E |
| 8539 | + | 2301 | Phage tail length tape-measure protein T |
| 8796 | + | 258 | phage-related tail formation protein U |
| 3497 | - | 3327 | Phage Host Specificity Protein |
| 4052 | + | 705 | Phage tail sheath protein FI |
| 4550 | + | 396 | Phage tail sheath protein FI |
| 5077 | + | 507 | Phage tail tube protein FII |
| 5423 | + | 258 | Phage tail protein E |
| 7668 | + | 2136 | Phage tail length tape-measure protein T |
| 168 | + | 147 | phage-related tail formation protein U |
| 374 | + | 210 | Phage tail protein GpX |

| | | | | | |
|---|---|---|---|---|---|
| | 1319 | + | | 945 | Phage tail formation protein D |
| | 410 | + | | 372 | phage-related tail formation protein U |
| | 472 | - | | 630 | some similarities to phage related proteins |
| | 579 | - | | 504 | Phage protein |
| - | | - | - | | - |
| - | | - | - | | - |
| | 11536 | - | | 432 | Putative bacteriophage-related protein |
| | 1495 | - | | 1434 | Phage-related protein |
| | 3799 | - | | 210 | prophage P2W3, tail protein X, putative |
| | 4005 | - | | 357 | prophage P2W3, tail protein U, putative |
| | 4361 | - | | 2265 | Phage tail length tape-measure protein |
| | 730 | - | | 498 | prophage LambdaW5, minor tail protein Z, put |
| | 2567 | - | | 372 | PROBABLE BACTERIOPHAGE-RELATED PROTEIN |
| | 1224 | - | | 321 | PROBABLE BACTERIOPHAGE-RELATED PROTEIN |
| | 2 | - | | 690 | Phage-related baseplate assembly protein |
| | 694 | - | | 234 | Phage baseplate assembly protein W |
| | 1508 | - | | 360 | prophage P2W3, tail protein D, putative |
| | 3 | - | | 183 | prophage LambdaW4, DNA methylase |
| | 299 | - | | 465 | Putative bacteriophage-related protein |
| | 18612 | + | | 1179 | Phage tail sheath monomer |
| | 19134 | + | | 507 | Phage major tail tube protein |
| | 22403 | + | | 2292 | Phage tail length tape-measure protein |
| | 22750 | + | | 348 | prophage P2W3, tail protein U, putative |
| | 23901 | + | | 945 | Phage protein D |
| | 32799 | - | | 792 | Phage-related baseplate assembly protein |
| | 33593 | - | | 336 | Phage baseplate assembly protein |
| | 35109 | - | | 498 | prophage LambdaW5, minor tail protein Z, put |
| | 37041 | - | | 372 | PROBABLE BACTERIOPHAGE-RELATED PROTEIN |
| | 38600 | - | | 1419 | Phage portal protein |
| | 40233 | - | | 1845 | Phage terminase, large subunit |
| | 45367 | - | | 465 | Putative bacteriophage-related protein |
| | 52436 | + | | 2058 | PROBABLE BACTERIOPHAGE-RELATED PROTEIN |
| | 750 | - | | 618 | Phage-related baseplate assembly protein |
| | 1808 | + | | 1803 | PROBABLE BACTERIOPHAGE-RELATED PROTEIN |
| | 5175 | + | | 462 | prophage LambdaW4, terminase large subunit |
| | 33132 | - | | 405 | Putative bacteriophage-related protein |
| | 570 | + | | 543 | Phage-related tail protein |
| | 926 | + | | 357 | prophage P2W3, tail protein U, putative |
| | 1132 | + | | 210 | prophage P2W3, tail protein X, putative |
| | 61684 | + | | 1176 | Phage portal protein |
| | 4996 | + | | 372 | PROBABLE BACTERIOPHAGE-RELATED PROTEIN |
| | 6899 | + | | 492 | prophage LambdaW5, minor tail protein Z, put |

| | | | |
|---|---|---|---|
| 24031 | - | 2196 | Phage Host Specificity Protein |
| 15643 | - | 432 | Putative bacteriophage-related protein |
| 20542 | - | 786 | Phage-related baseplate assembly protein |
| 21330 | - | 336 | Phage baseplate assembly protein |
| 22840 | - | 492 | prophage LambdaW5, minor tail protein Z, put |
| 25117 | - | 372 | PROBABLE BACTERIOPHAGE-RELATED PROTEI |
| 31379 | - | 465 | Putative bacteriophage-related protein |
| 37201 | - | 1791 | PROBABLE BACTERIOPHAGE-RELATED PROTEI |
| 17828 | + | 1398 | Phage-related protein |
| - | - | - | - |
| 211 | + | 210 | Phage tail fiber protein |
| 5921 | - | 432 | Phage protein |
| 7766 | - | 1167 | Phage tail formation protein I |
| 9210 | - | 801 | Phage baseplate assembly protein J |
| 10013 | - | 336 | Phage baseplate assembly protein GpW |
| 10627 | - | 168 | Phage baseplate assembly protein V |
| 11034 | - | 474 | Phage protein |
| 11504 | - | 513 | Phage minor tail protein Z |
| 15767 | + | 222 | Phage major capsid protein |
| 76696 | + | 345 | Phage protein |
| 81411 | + | 378 | Phage head, terminase subunit Nu1 |
| 84722 | + | 1827 | Phage head, terminase DNA packaging protein |
| 87052 | + | 1407 | Phage head, portal protein B |
| 87480 | + | 429 | Phage head, head-tail preconnector protease |
| 88095 | + | 489 | Phage head, head-tail preconnector protease |
| 88541 | + | 372 | Phage head, head-DNA stabilization protein D |
| 88653 | + | 150 | Phage head, major capsid protein E |
| 90685 | - | 477 | Phage protein |
| 98892 | - | 711 | Phage tail formation protein D |
| 99603 | - | 210 | Phage tail protein GpX |
| 99809 | - | 348 | phage-related tail formation protein U |
| 100158 | - | 2307 | Phage tail length tape-measure protein T |
| 102574 | - | 258 | Phage tail protein E |
| 102914 | - | 507 | Phage tail tube protein FII |
| 103436 | - | 315 | Phage tail sheath protein FI |
| 103704 | - | 309 | Phage head, portal protein B |
| 104227 | - | 1836 | Phage head, terminase DNA packaging protein |
| 107531 | - | 471 | Phage head, terminase subunit Nu1 |
| 109408 | - | 423 | Phage protein |
| 135016 | + | 1176 | Phage tail sheath protein FI |
| 135538 | + | 507 | Phage tail tube protein FII |
| 135878 | + | 258 | Phage tail protein E |

| | | | |
|---|---|---|---|
| 138408 | + | 2421 | Phage tail length tape-measure protein T |
| 138755 | + | 348 | phage-related tail formation protein U |
| 138961 | + | 210 | Phage tail protein GpX |
| 139913 | + | 951 | Phage tail formation protein D |
| 168556 | - | 432 | Phage protein |
| 170367 | - | 1167 | Phage tail formation protein I |
| 171811 | - | 801 | Phage baseplate assembly protein J |
| 172614 | - | 336 | Phage baseplate assembly protein GpW |
| 173228 | - | 168 | Phage baseplate assembly protein V |
| 173635 | - | 474 | Phage protein |
| 174105 | - | 513 | Phage minor tail protein Z |
| 178368 | + | 222 | Phage major capsid protein |
| 181301 | - | 996 | Phage head, portal protein B |
| 183211 | - | 1824 | Phage head, terminase DNA packaging protein |
| 186519 | - | 480 | Phage head, terminase subunit Nu1 |
| 188330 | - | 456 | Phage protein |
| 189422 | - | 732 | Prophage antirepressor |
| 195177 | + | 693 | Phage tail formation protein I |
| 196988 | + | 432 | Phage protein |
| 214062 | - | 507 | Phage tail tube protein FII |
| 214580 | - | 1182 | Phage tail sheath protein FI |
| 237397 | - | 432 | Phage protein |
| 239200 | - | 1161 | Phage tail formation protein I |
| 240360 | - | 792 | Phage baseplate assembly protein J |
| 241154 | - | 234 | Phage baseplate assembly protein GpW |
| 242777 | - | 114 | Phage baseplate assembly protein GpW |
| 244227 | - | 465 | Phage baseplate assembly protein V |
| 244678 | - | 477 | Phage protein |
| 245151 | - | 522 | Phage minor tail protein Z |
| 245973 | - | 606 | Phage head, major capsid protein E |
| 248065 | - | 2316 | Phage tail length tape-measure protein T |
| 250490 | - | 258 | Phage tail protein E |
| 251364 | - | 507 | Phage tail tube protein FII |
| 251888 | - | 1185 | Phage tail sheath protein FI |
| 275482 | - | 432 | Phage protein |
| 277327 | - | 693 | Phage tail formation protein I |
| 281107 | - | 99 | Phage baseplate assembly protein GpW |
| 281459 | - | 465 | Phage baseplate assembly protein V |
| 281910 | - | 474 | Phage protein |
| 282380 | - | 492 | Phage minor tail protein Z |
| 282873 | - | 219 | putative phage related protein |
| 283264 | - | 1005 | Phage head, major capsid protein E |

| | | | |
|---|---|---|---|
| 284294 | - | 369 | Phage head, head-DNA stabilization protein D |
| 284740 | - | 1044 | Phage head, head-tail preconnector protease |
| 285770 | - | 813 | Phage head, portal protein B |
| 286659 | - | 783 | Phage head, major capsid protein E |
| 287467 | - | 369 | Phage head, head-DNA stabilization protein D |
| 287913 | - | 1044 | Phage head, head-tail preconnector protease |
| 288943 | - | 1857 | Phage head, major capsid protein E |
| 290825 | - | 369 | Phage head, head-DNA stabilization protein D |
| 291271 | - | 1044 | Phage head, head-tail preconnector protease |
| 292301 | - | 1419 | Phage head, portal protein B |
| 294701 | - | 549 | Phage head, terminase DNA packaging protein |
| 640940 | - | 915 | Phage portal protein |
| 644141 | - | 228 | Phage portal protein |
| 982022 | + | 333 | some similarities to phage related proteins |
| 1035340 | - | 504 | some similarities to phage related proteins |
| 1037386 | - | 822 | some similarities to phage related proteins |
| 1042691 | - | 3474 | Phage tail fiber protein |
| 1291923 | - | 1161 | Phage tail formation protein I |
| 1293083 | - | 144 | prophage LambdaW1, baseplate assembly prot |
| 1293249 | - | 621 | Phage baseplate assembly protein J |
| 1293872 | - | 336 | Phage baseplate assembly protein GpW |
| 1294472 | - | 465 | Phage baseplate assembly protein V |
| 1294923 | - | 474 | Phage protein |
| 1295393 | - | 513 | prophage LambdaW5, minor tail protein Z, put |
| 1296209 | - | 1005 | Phage head, major capsid protein E |
| 1297250 | - | 411 | Phage head, head-DNA stabilization protein D |
| 1297697 | - | 417 | Phage head, head-tail preconnector protease |
| 1302863 | + | 855 | Phage head, terminase DNA packaging protein |
| 1303619 | - | 471 | Phage head, terminase subunit Nu1 |
| 1305496 | - | 423 | Phage protein |
| 1326513 | + | 1182 | Phage tail sheath protein FI |
| 1327035 | + | 507 | Phage tail tube protein FII |
| 1327375 | + | 258 | Phage tail protein E |
| 1329791 | + | 2307 | Phage tail length tape-measure protein T |
| 1330140 | + | 348 | phage-related tail formation protein U |
| 1330346 | + | 210 | Phage tail protein GpX |
| 1331297 | + | 951 | Phage tail formation protein D |
| 1367231 | - | 114 | prophage LambdaW1, site-specific recombinas |
| 1370086 | - | 432 | Phage protein |
| 1371931 | - | 1167 | Phage tail formation protein I |
| 1373375 | - | 801 | Phage baseplate assembly protein J |
| 1374178 | - | 336 | Phage baseplate assembly protein GpW |

| | | | |
|---|---|---|---|
| 1374792 | - | 168 | Phage baseplate assembly protein V |
| 1375199 | - | 474 | Phage protein |
| 1375669 | - | 513 | Phage minor tail protein Z |
| 1489785 | - | 930 | Phage major capsid protein |
| 1586666 | - | 678 | Phage tail formation protein D |
| 1587344 | - | 210 | Phage tail protein GpX |
| 1587550 | - | 348 | phage-related tail formation protein U |
| 1587899 | - | 2307 | Phage tail length tape-measure protein T |
| 1590315 | - | 258 | Phage tail protein E |
| 1590655 | - | 507 | Phage tail tube protein FII |
| 1596477 | - | 678 | Phage tail formation protein D |
| 1597155 | - | 210 | Phage tail protein GpX |
| 1597361 | - | 348 | phage-related tail formation protein U |
| 1597710 | - | 1935 | Phage tail length tape-measure protein T |
| 1599619 | - | 636 | Phage tail length tape-measure protein T |
| 117 | - | 210 | Phage tail protein GpX |
| 323 | - | 339 | prophage P2W3, tail protein U, putative |
| 661 | - | 2439 | Phage tail length tape-measure protein T |
| 3209 | - | 126 | Phage tail protein E |
| 5544 | - | 951 | Phage tail formation protein D |
| 6496 | - | 210 | Phage tail protein GpX |
| 6702 | - | 339 | prophage P2W3, tail protein U, putative |
| 7040 | - | 2439 | Phage tail length tape-measure protein T |
| 9588 | - | 258 | Phage tail protein E |
| 9892 | - | 507 | Phage tail tube protein FII |
| 10427 | - | 204 | Phage tail sheath protein FI |
| 36810 | - | 132 | Phage protein |
| 37028 | - | 213 | Phage protein |
| 38606 | - | 1161 | Phage tail formation protein I |
| 39766 | - | 801 | Phage baseplate assembly protein J |
| 40569 | - | 96 | Phage baseplate assembly protein |
| 42197 | - | 114 | Phage baseplate assembly protein GpW |
| 43647 | - | 465 | Phage baseplate assembly protein V |
| 44098 | - | 477 | Phage protein |
| 44571 | - | 522 | Phage minor tail protein Z |
| 47090 | - | 114 | Phage baseplate assembly protein GpW |
| 47460 | - | 444 | Phage baseplate assembly protein V |
| 47890 | - | 477 | Phage protein |
| 48363 | - | 177 | prophage LambdaW5, minor tail protein Z, put |
| 48575 | - | 249 | Phage minor tail protein Z |
| 49127 | - | 669 | Phage head, major capsid protein E |
| 49785 | - | 369 | Phage head, head-tail preconnector protease ( |

| | | | |
|---|---|---|---|
| 50167 | - | 1413 | Phage head, portal protein B |
| 51794 | - | 1836 | Phage head, terminase DNA packaging protein |
| 55132 | - | 483 | Phage head, terminase subunit Nu1 |
| 57815 | - | 471 | Phage head, terminase subunit Nu1 |
| 59638 | - | 477 | Phage protein |
| 84990 | + | 1176 | Phage tail sheath protein FI |
| 85512 | + | 507 | Phage tail tube protein FII |
| 86896 | + | 258 | Phage tail protein E |
| 88790 | + | 1785 | Phage tail length tape-measure protein T |
| 91923 | + | 1176 | Phage tail sheath protein FI |
| 92445 | + | 507 | Phage tail tube protein FII |
| 93829 | + | 258 | Phage tail protein E |
| 95681 | + | 1743 | Phage tail length tape-measure protein T |
| 98904 | + | 1176 | Phage tail sheath protein FI |
| 99426 | + | 507 | Phage tail tube protein FII |
| 100810 | + | 258 | Phage tail protein E |
| 102311 | + | 1392 | Phage tail length tape-measure protein T |
| 103179 | + | 813 | Phage tail length tape-measure protein T |
| 103540 | + | 348 | phage-related tail formation protein U |
| 103746 | + | 210 | Phage tail protein GpX |
| 104691 | + | 945 | Phage tail formation protein D |
| 149484 | + | 1629 | Phage head, terminase DNA packaging protein |
| 152165 | + | 1407 | Phage head, portal protein B |
| 172931 | + | 1176 | Phage tail sheath protein FI |
| 173453 | + | 507 | Phage tail tube protein FII |
| 174837 | + | 258 | Phage tail protein E |
| 177385 | + | 2439 | Phage tail length tape-measure protein T |
| 177723 | + | 339 | prophage P2W3, tail protein U, putative |
| 177929 | + | 210 | Phage tail protein GpX |
| 178881 | + | 951 | Phage tail formation protein D |
| 56231 | + | 429 | Phage tail sheath protein FI |
| 56494 | + | 189 | Phage tail sheath protein FI |
| 136957 | + | 300 | prophage LambdaW5, ankyrin repeat domain [illegible] |
| 137233 | + | 156 | prophage LambdaW5, ankyrin repeat domain [illegible] |
| 158200 | + | 186 | uncharacterized phage protein, putative |
| 164067 | - | 156 | Phage major capsid protein |
| 164577 | - | 264 | Phage major capsid protein |
| 603088 | + | 408 | Phage tail fiber protein |
| 603352 | + | 243 | Phage tail fiber protein |
| 603510 | + | 129 | Phage tail fiber protein |
| 604263 | + | 156 | Phage tail fiber protein |
| 604569 | + | 159 | Phage tail fiber protein |

| | | | |
|---|---|---|---|
| 604849 | + | 312 | Phage tail fiber protein |
| 661595 | - | 432 | some similarities to phage related proteins |
| 662151 | - | 612 | some similarities to phage related proteins |
| 764710 | - | 594 | Phage tail fiber protein |
| 765300 | - | 342 | Phage tail fiber protein |
| 1096535 | + | 216 | Phage tail formation protein D |
| 1115439 | - | 189 | prophage LambdaW1, site-specific recombinas |
| 1115839 | - | 123 | prophage LambdaW1, site-specific recombinas |
| 1115971 | - | 555 | prophage LambdaW1, site-specific recombinas |
| 27323 | + | 3204 | Phage Host Specificity Protein |
| 116743 | - | 1140 | Phage major capsid protein |
| 128713 | - | 342 | prophage LambdaW1, site-specific recombinas |
| 313692 | + | 342 | prophage LambdaW1, site-specific recombinas |
| 16730 | - | 936 | Phage tail formation protein D |
| 17666 | - | 210 | Phage protein |
| 17878 | - | 348 | prophage P2W3, tail protein U, putative |
| 18225 | - | 2271 | Phage tail length tape-measure protein T |
| 20611 | - | 255 | Phage tail protein E |
| 20912 | - | 507 | Phage tail tube protein FII |
| 21432 | - | 396 | Phage tail sheath protein FI |
| 21865 | - | 750 | Phage tail sheath protein FI |
| 39207 | + | 462 | Phage protein |
| 43217 | + | 483 | Phage head, terminase subunit Nu1 |
| 8 | - | 1185 | Phage tail length tape-measure protein T |
| 1302 | - | 258 | Phage tail protein E |
| 1634 | - | 507 | Phage tail tube protein FII |
| 2152 | - | 1182 | Phage tail sheath protein FI |
| 21672 | + | 465 | Phage protein |
| 25632 | + | 483 | Phage head, terminase subunit Nu1 |
| 28622 | + | 1515 | Phage head, terminase DNA packaging protein |
| 30306 | + | 111 | Phage head, portal protein B |
| 31593 | + | 1275 | Phage head, portal protein B |
| 32645 | + | 1053 | Phage head, head-tail preconnector protease ( |
| 33063 | + | 372 | Phage head, head-DNA stabilization protein D |
| 34089 | + | 1002 | Phage head, major capsid protein E |
| 37451 | + | 495 | Phage minor tail protein Z |
| 42716 | + | 393 | Phage tail fiber protein |
| 87524 | - | 432 | Phage protein |
| 89351 | - | 1167 | Phage tail formation protein I |
| 90804 | - | 783 | Phage tail tip, host specificity protein J |
| 91589 | - | 336 | Phage baseplate assembly protein GpW |
| 92205 | - | 465 | Phage baseplate assembly protein V |

| | | | |
|---|---|---|---|
| 92656 | - | 477 | Phage protein |
| 93152 | - | 495 | Phage minor tail protein Z |
| 96511 | - | 1002 | Phage head, major capsid protein E |
| 97537 | - | 372 | Phage head, head-DNA stabilization protein D |
| 98106 | - | 306 | Phage head, head-tail preconnector protease C |
| 99923 | - | 738 | Phage head, head-tail preconnector protease C |
| 100657 | - | 1434 | Phage head, portal protein B |
| 102801 | - | 1821 | Phage head, terminase DNA packaging protein |
| 106105 | - | 480 | Phage head, terminase subunit Nu1 |
| 107916 | - | 456 | Phage protein |
| 190084 | + | 465 | Phage baseplate assembly protein V |
| 1991 | + | 171 | putative phage related protein |
| 2518 | + | 492 | Phage minor tail protein Z |
| 3014 | + | 477 | Phage protein |
| 3465 | + | 465 | Phage baseplate assembly protein V |
| 7863 | + | 372 | Phage baseplate assembly protein GpW |
| 8648 | + | 783 | Phage baseplate assembly protein J |
| 9808 | + | 1161 | Phage tail formation protein I |
| 11650 | + | 432 | Phage protein |
| 13128 | + | 1476 | prophage LambdaW1, site-specific recombinas |
| 31444 | - | 345 | prophage LambdaW1, site-specific recombinas |
| 53426 | - | 1380 | some similarities to phage related proteins |
| 157626 | + | 1434 | some similarities to phage related proteins |
| 173119 | - | 342 | prophage LambdaW1, site-specific recombinas |
| 26 | - | 780 | Phage baseplate assembly protein J |
| 808 | - | 336 | Phage baseplate assembly protein GpW |
| 1397 | - | 465 | Phage baseplate assembly protein V |
| 1848 | - | 474 | Phage protein |
| 2338 | - | 480 | Phage minor tail protein Z |
| 2819 | - | 219 | putative phage related protein |
| 3210 | - | 1005 | Phage head, major capsid protein E |
| 4252 | - | 375 | Phage head, head-DNA stabilization protein D |
| 4744 | - | 1005 | Phage head, head-tail preconnector protease C |
| 5745 | - | 1407 | Phage head, portal protein B |
| 7366 | - | 1836 | Phage head, terminase DNA packaging protein |
| 10702 | - | 486 | Phage head, terminase subunit Nu1 |
| 12534 | - | 465 | Phage protein |
| 73645 | + | 1176 | Phage portal protein |
| 75066 | + | 144 | Phage portal protein |
| 146815 | - | 318 | prophage LambdaW1, site-specific recombinas |
| 147099 | - | 1197 | prophage LambdaW1, site-specific recombinas |
| 211583 | + | 147 | prophage LambdaW1, site-specific recombinas |

| | | | |
|---|---|---|---|
| 212446 | + | 357 | prophage LambdaW1, site-specific recombinas |
| 222761 | - | 1476 | prophage LambdaW1, site-specific recombinas |
| 224239 | - | 432 | Phage protein |
| 226090 | - | 1161 | Phage tail formation protein I |
| 227250 | - | 801 | Phage baseplate assembly protein J |
| 228053 | - | 336 | Phage baseplate assembly protein GpW |
| 228642 | - | 465 | Phage baseplate assembly protein V |
| 229093 | - | 483 | Phage protein |
| 229590 | - | 477 | Phage tail, component Z |
| 230067 | - | 219 | putative phage related protein |
| 234510 | - | 1002 | Phage head, major capsid protein E |
| 235544 | - | 369 | Phage head, head-DNA stabilization protein D |
| 235980 | - | 1035 | Phage head, head-tail preconnector protease C |
| 237014 | - | 1398 | Phage head, portal protein B |
| 238944 | - | 1515 | Phage head, terminase DNA packaging protein |
| 241934 | - | 483 | Phage head, terminase subunit Nu1 |
| 245944 | - | 462 | Phage protein |
| 263665 | + | 1203 | Phage tail sheath protein FI |
| 264183 | + | 507 | Phage tail tube protein FII |
| 265629 | + | 258 | Phage tail protein E |
| 268039 | + | 2301 | Phage tail length tape-measure protein T |
| 268386 | + | 348 | phage-related tail formation protein U |
| 268589 | + | 207 | Phage tail protein GpX |
| 269537 | + | 948 | Phage tail formation protein D |
| 306562 | - | 423 | Phage head, terminase DNA packaging protein |
| 306944 | - | 231 | Phage head, terminase DNA packaging protein |
| 308641 | - | 474 | Phage head, terminase subunit Nu1 |
| 310421 | - | 381 | Phage protein |
| 310761 | - | 129 | Phage protein |
| 509668 | + | 342 | prophage LambdaW1, site-specific recombinas |
| 38184 | - | 351 | Phage major capsid protein |
| 315 | + | 219 | Phage uncharacterized protein |
| 6150 | + | 204 | uncharacterized phage protein, putative |
| 547 | - | 183 | Phage major capsid protein |

| Subsystem |
|---|
| - none - |
| - none - |
| - none - |
| Phage capsid proteins |
| Phage capsid proteins |
| - none - |
| - none - |
| - none - |
| - none - |
| - none - |
| - none - |
| - none - |
| - none - |
| - none - |
| - none - |
| - none - |
| - none - |
| - none - |
| - none - |
| - none - |
| Phage capsid proteins |
| - none - |
| - none - |
| - none - |
| Phage packaging machinery |
| - none - |
| - none - |
| - none - |
| - none - |
| - none - |
| - none - |
| - none - |
| - none - |
| - none - |
| - none - |
| - none - |
| - none - |
| - none - |
| - none - |

| |
|---|
| - none - |
| - none - |
| - none - |
| - none - |
| - none - |
| - none - |
| - none - |
| Phage tail proteins |
| - none - |
| - none - |
| - none - |
| - none - |
| - none - |
| - none - |
| - none - |
| - none - |
| - none - |
| - none - |
| - none - |
| - none - |
| - none - |
| - none - |
| - none - |
| - none - |
| - none - |
| - none - |
| - none - |
| - none - |
| - none - |
| - none - |
| - none - |
| - none - |
| - none - |
| - none - |
| - none - |
| - none - |
| - none - |
| - none - |
| - none - |
| - none - |

- none -
- none -
- none -
- none -
- none -
- none -
- none -
- none -
- none -
- none -
- none -
- none -
- none -
- none -
- none -
- none -
- none -
Phage capsid proteins
- none -
- none -
-
Phage packaging machinery
Phage packaging machinery
- none -
Phage packaging machinery
- none -
- none -
- none -
- none -
- none -
Phage capsid proteins
- none -
- none -
Phage packaging machinery
- none -
- none -
Phage capsid proteins
Phage tail fiber proteins
- none -
- none -
Phage tail fiber proteins
- none -

- none -
- none -
- none -
- none -
- none -
- none -
- none -
- none -
- none -
- none -
- none -
- none -
- none -
- none -
- none -
- none -
- none -
- none -
- none -
- none -
- none -
- none -
- none -
- none -
- none -
- none -
- none -
- none -
- none -
- none -
- none -
- none -
- none -
- none -
- none -
- none -
- none -
- none -
-
-
Phage capsid proteins
- none -

- none -
- none -
Phage tail proteins; <br>Phage tail proteins 2
- none -
- none -
- none -
- none -
Phage tail proteins; <br>Phage tail proteins 2
- none -
Phage tail proteins; <br>Phage tail proteins 2
- none -
Phage packaging machinery
Phage packaging machinery
- none -
- none -
- none -
- none -
- none -
Phage packaging machinery
Phage packaging machinery
- none -
- none -
- none -
- none -
Phage packaging machinery
- none -
- none -
- none -
Phage tail proteins; <br>Phage tail proteins 2
- none -
Phage tail proteins; <br>Phage tail proteins 2
- none -
- none -
- none -
- none -
Phage packaging machinery
Phage packaging machinery
- none -
Phage tail proteins; <br>Phage tail proteins 2
- none -
Phage tail proteins; <br>Phage tail proteins 2
- none -

- none -
- none -
- none -
- none -
Phage tail proteins; <br>Phage tail proteins 2
- none -
- none -
- none -
- none -
- none -
Phage tail fiber proteins
- none -
- none -
- none -
- none -
- none -
- none -
- none -
- none -
- none -
- none -
- none -
- none -
- none -
- none -
- none -
Phage capsid proteins
- none -
- none -
- none -
- none -
- none -
- none -
- none -
- none -
- none -
- none -
- none -
- none -
- none -
- none -
- none -

- none -
- none -
- none -
- none -
- none -
- none -
- none -
- none -
- none -
- none -
- none -
- none -
- none -
- none -
Phage capsid proteins
- none -
- none -
- none -
- none -
- none -
- none -
- none -
-
-
Phage tail proteins; <br>Phage tail proteins 2
- none -
- none -
- none -
- none -
- none -
- none -
- none -
- none -
- none -
- none -
- none -
- none -
- none -
- none -
- none -
- none -
- none -

| |
|---|
| - none - |
| - none - |
| - none - |
| - none - |
| - none - |
| Phage tail fiber proteins |
| Phage tail fiber proteins |
| - none - |
| - none - |
| Phage tail fiber proteins |
| Phage tail fiber proteins |
| - none - |
| - none - |
| - none - |
| Phage capsid proteins |
| - none - |
| - none - |
| - none - |
| - none - |
| - none - |
| - none - |
| - none - |
| - none - |
| - none - |
| - none - |
| - none - |
| - none - |
| - none - |
| - none - |
| - none - |
| - none - |
| - none - |
| - none - |
| - none - |
| - none - |
| - none - |
| - none - |
| Phage tail fiber proteins |
| Phage packaging machinery |
| - none - |
| - none - |

- none -
- none -
- none -
- none -
- none -
- none -
- none -
- none -
- none -
- none -
- none -
- none -
- none -
- none -
- none -
- none -
- none -
- none -
- none -
- none -
- none -
Phage tail proteins
- none -
- none -
- none -
- none -
- none -
- none -
- none -
- none -
- none -
- none -
- none -
- none -
- none -
- none -
- none -
- none -
- none -
- none -
- none -
- none -

- none -
- none -
- none -
- none -
- none -
- none -
- none -
- none -
- none -
- none -
- none -
- none -
- none -
- none -
- none -
- none -
- none -
Phage packaging machinery
Phage packaging machinery
Phage capsid proteins
- none -
- none -
- none -
- none -
- none -
- none -
- none -
- none -
- none -
- none -
- none -
- none -
- none -
- none -
- none -
- none -
- none -
- none -
- none -
- none -
- none -
- none -

- none -
- none -
- none -
- none -
-
-
- none -
- none -
- none -
- none -
- none -
- none -
- none -
- none -
- none -
- none -
- none -
- none -
- none -
- none -
- none -
- none -
- none -
- none -
- none -
- none -
- none -
- none -
- none -
- none -
- none -
- none -
- none -
- none -
- none -
- none -
- none -
- none -
- none -
- none -
- none -
- none -

- none -
- none -
- none -
- none -
- none -
- none -
- none -
- none -
- none -
-
Phage tail fiber proteins
- none -
- none -
- none -
- none -
- none -
- none -
- none -
Phage capsid proteins
- none -
- none -
- none -
- none -
- none -
- none -
- none -
- none -
- none -
- none -
- none -
- none -
- none -
- none -
- none -
- none -
- none -
- none -
- none -
- none -
- none -
- none -
- none -

| |
|---|
| - none - |
| - none - |
| - none - |
| - none - |
| - none - |
| - none - |
| - none - |
| - none - |
| - none - |
| - none - |
| - none - |
| Phage capsid proteins |
| - none - |
| - none - |
| - none - |
| - none - |
| - none - |
| - none - |
| - none - |
| - none - |
| - none - |
| - none - |
| - none - |
| - none - |
| - none - |
| - none - |
| - none - |
| - none - |
| - none - |
| - none - |
| - none - |
| - none - |
| - none - |
| - none - |
| - none - |
| - none - |
| - none - |
| - none - |
| - none - |
| - none - |
| - none - |
| - none - |

| |
|---|
| - none - |
| - none - |
| - none - |
| - none - |
| - none - |
| - none - |
| - none - |
| - none - |
| - none - |
| - none - |
| - none - |
| Phage packaging machinery |
| Phage packaging machinery |
| - none - |
| - none - |
| - none - |
| Phage tail fiber proteins |
| - none - |
| - none - |
| - none - |
| - none - |
| - none - |
| - none - |
| - none - |
| - none - |
| - none - |
| - none - |
| - none - |
| - none - |
| - none - |
| - none - |
| - none - |
| - none - |
| - none - |
| - none - |
| - none - |
| - none - |
| - none - |
| - none - |
| - none - |
| - none - |
| - none - |

| |
|---|
| - none - |
| - none - |
| - none - |
| Phage capsid proteins |
| - none - |
| - none - |
| - none - |
| - none - |
| - none - |
| - none - |
| - none - |
| - none - |
| - none - |
| - none - |
| - none - |
| - none - |
| - none - |
| - none - |
| - none - |
| - none - |
| - none - |
| - none - |
| - none - |
| - none - |
| - none - |
| - none - |
| - none - |
| - none - |
| - none - |
| - none - |
| - none - |
| - none - |
| - none - |
| - none - |
| - none - |
| - none - |
| - none - |
| - none - |
| - none - |
| - none - |
| - none - |
| - none - |

- none -
- none -
- none -
- none -
- none -
- none -
- none -
- none -
- none -
- none -
- none -
- none -
- none -
- none -
- none -
- none -
- none -
- none -
- none -
- none -
- none -
- none -
- none -
- none -
- none -
- none -
- none -
- none -
- none -
- none -
- none -
- none -
- none -
- none -
- none -
Phage capsid proteins
Phage capsid proteins
Phage tail fiber proteins
Phage tail fiber proteins
Phage tail fiber proteins
Phage tail fiber proteins
Phage tail fiber proteins

Phage tail fiber proteins
- none -
- none -
Phage tail fiber proteins
Phage tail fiber proteins
- none -
- none -
- none -
- none -
- none -
Phage capsid proteins
- none -
- none -
- none -
- none -
- none -
- none -
- none -
- none -
- none -
- none -
- none -
- none -
- none -
- none -
- none -
- none -
- none -
- none -
- none -
- none -
- none -
- none -
- none -
- none -
- none -
Phage tail fiber proteins
- none -
- none -
- none -
- none -
- none -

- none -
- none -
- none -
- none -
- none -
- none -
- none -
- none -
- none -
- none -
- none -
- none -
- none -
- none -
- none -
- none -
- none -
- none -
- none -
- none -
- none -
- none -
- none -
- none -
- none -
- none -
- none -
- none -
- none -
- none -
- none -
- none -
- none -
- none -
- none -
- none -
- none -
Phage packaging machinery
Phage packaging machinery
- none -
- none -
- none -

| |
|---|
| - none - |
| - none - |
| - none - |
| - none - |
| - none - |
| - none - |
| - none - |
| - none - |
| - none - |
| - none - |
| - none - |
| - none - |
| - none - |
| - none - |
| - none - |
| - none - |
| - none - |
| - none - |
| - none - |
| - none - |
| - none - |
| - none - |
| - none - |
| - none - |
| - none - |
| - none - |
| - none - |
| - none - |
| - none - |
| - none - |
| Phage capsid proteins |
| - none - |
| - none - |
| Phage capsid proteins |

**Table S11. List of Group II intron-associated genes detectd by RAST pipeline.**

| genome | Feature ID | Type | Contig | Start |
|---|---|---|---|---|
| wBm | fig\|292805.13.peg.114 | CDS | wBm_reseq_reversed | 73102 |
| | fig\|292805.13.peg.117 | CDS | wBm_reseq_reversed | 73863 |
| | fig\|292805.13.peg.148 | CDS | wBm_reseq_reversed | 92642 |
| | fig\|292805.13.peg.359 | CDS | wBm_reseq_reversed | 244000 |
| | fig\|292805.13.peg.366 | CDS | wBm_reseq_reversed | 246465 |
| | fig\|292805.13.peg.451 | CDS | wBm_reseq_reversed | 305656 |
| | fig\|292805.13.peg.768 | CDS | wBm_reseq_reversed | 527797 |
| | fig\|292805.13.peg.770 | CDS | wBm_reseq_reversed | 528230 |
| | fig\|292805.13.peg.876 | CDS | wBm_reseq_reversed | 591256 |
| | fig\|292805.13.peg.1178 | CDS | wBm_reseq_reversed | 789538 |
| | fig\|292805.13.peg.1195 | CDS | wBm_reseq_reversed | 801082 |
| | fig\|292805.13.peg.1351 | CDS | wBm_reseq_reversed | 914605 |
| | fig\|292805.13.peg.1370 | CDS | wBm_reseq_reversed | 931551 |
| | fig\|292805.13.peg.1466 | CDS | wBm_reseq_reversed | 994127 |
| wBp | fig\|96495.5.peg.83 | CDS | NZ_CP050521.1 | 68548 |
| | fig\|96495.5.peg.84 | CDS | NZ_CP050521.1 | 68851 |
| | fig\|96495.5.peg.134 | CDS | NZ_CP050521.1 | 106398 |
| | fig\|96495.5.peg.162 | CDS | NZ_CP050521.1 | 125187 |
| | fig\|96495.5.peg.164 | CDS | NZ_CP050521.1 | 126044 |
| | fig\|96495.5.peg.360 | CDS | NZ_CP050521.1 | 284373 |
| | fig\|96495.5.peg.449 | CDS | NZ_CP050521.1 | 360334 |
| | fig\|96495.5.peg.635 | CDS | NZ_CP050521.1 | 519936 |
| | fig\|96495.5.peg.687 | CDS | NZ_CP050521.1 | 564919 |
| | fig\|96495.5.peg.688 | CDS | NZ_CP050521.1 | 565291 |
| | fig\|96495.5.peg.830 | CDS | NZ_CP050521.1 | 681971 |
| | fig\|96495.5.peg.1006 | CDS | NZ_CP050521.1 | 826430 |
| | fig\|96495.5.peg.1012 | CDS | NZ_CP050521.1 | 828893 |
| | fig\|96495.5.peg.1079 | CDS | NZ_CP050521.1 | 888052 |
| | fig\|96495.5.peg.1173 | CDS | NZ_CP050521.1 | 964250 |
| | fig\|96495.5.peg.1174 | CDS | NZ_CP050521.1 | 964912 |
| wCauA | fig\|2591635.5.peg.201 | CDS | CP041215.1 | 184749 |
| | fig\|2591635.5.peg.290 | CDS | CP041215.1 | 263255 |
| | fig\|2591635.5.peg.320 | CDS | CP041215.1 | 294202 |
| | fig\|2591635.5.peg.336 | CDS | CP041215.1 | 308423 |
| | fig\|2591635.5.peg.428 | CDS | CP041215.1 | 402304 |
| | fig\|2591635.5.peg.477 | CDS | CP041215.1 | 442575 |
| | fig\|2591635.5.peg.484 | CDS | CP041215.1 | 446226 |
| | fig\|2591635.5.peg.526 | CDS | CP041215.1 | 481560 |
| | fig\|2591635.5.peg.588 | CDS | CP041215.1 | 532168 |

| | | | | |
|---|---|---|---|---|
| | fig\|2591635.5.peg.643 | CDS | CP041215.1 | 595336 |
| | fig\|2591635.5.peg.828 | CDS | CP041215.1 | 775309 |
| | fig\|2591635.5.peg.834 | CDS | CP041215.1 | 782160 |
| | fig\|2591635.5.peg.947 | CDS | CP041215.1 | 885203 |
| | fig\|2591635.5.peg.1002 | CDS | CP041215.1 | 943133 |
| | fig\|2591635.5.peg.1158 | CDS | CP041215.1 | 1091324 |
| | fig\|2591635.5.peg.1187 | CDS | CP041215.1 | 1120192 |
| | fig\|2591635.5.peg.1238 | CDS | CP041215.1 | 1180532 |
| | fig\|2591635.5.peg.1241 | CDS | CP041215.1 | 1184408 |
| | fig\|2591635.5.peg.1309 | CDS | CP041215.1 | 1242840 |
| wCtub | fig\|6666666.333112.peg.64 | CDS | wCtub_1_pilon_pilon | 65291 |
| | fig\|6666666.333112.peg.712 | CDS | wCtub_1_pilon_pilon | 773336 |
| wCfeJ | fig\|212123.3.peg.23 | CDS | CP051157.1 | 19644 |
| | fig\|212123.3.peg.26 | CDS | CP051157.1 | 24735 |
| | fig\|212123.3.peg.170 | CDS | CP051157.1 | 159267 |
| | fig\|212123.3.peg.255 | CDS | CP051157.1 | 252846 |
| | fig\|212123.3.peg.339 | CDS | CP051157.1 | 347939 |
| | fig\|212123.3.peg.500 | CDS | CP051157.1 | 512440 |
| | fig\|212123.3.peg.523 | CDS | CP051157.1 | 532548 |
| | fig\|212123.3.peg.824 | CDS | CP051157.1 | 836134 |
| | fig\|212123.3.peg.956 | CDS | CP051157.1 | 980620 |
| | fig\|212123.3.peg.1023 | CDS | CP051157.1 | 1059287 |
| | fig\|212123.3.peg.1030 | CDS | CP051157.1 | 1068174 |
| | fig\|212123.3.peg.1157 | CDS | CP051157.1 | 1180619 |
| wCfeT | fig\|212123.4.peg.232 | CDS | CP051156.1 | 223517 |
| | fig\|212123.4.peg.263 | CDS | CP051156.1 | 263288 |
| | fig\|212123.4.peg.325 | CDS | CP051156.1 | 324510 |
| | fig\|212123.4.peg.459 | CDS | CP051156.1 | 460975 |
| | fig\|212123.4.peg.576 | CDS | CP051156.1 | 576510 |
| | fig\|212123.4.peg.683 | CDS | CP051156.1 | 684534 |
| | fig\|212123.4.peg.753 | CDS | CP051156.1 | 763460 |
| | fig\|212123.4.peg.796 | CDS | CP051156.1 | 804877 |
| | fig\|212123.4.peg.1447 | CDS | CP051156.1 | 1429394 |
| | fig\|212123.4.peg.1453 | CDS | CP051156.1 | 1432543 |
| wCle | fig\|246273.9.peg.151 | CDS | wCle_NZ_AP013028 | 144250 |
| | fig\|246273.9.peg.169 | CDS | wCle_NZ_AP013028 | 158182 |
| | fig\|246273.9.peg.299 | CDS | wCle_NZ_AP013028 | 262640 |
| | fig\|246273.9.peg.306 | CDS | wCle_NZ_AP013028 | 268534 |
| | fig\|246273.9.peg.307 | CDS | wCle_NZ_AP013028 | 269365 |
| | fig\|246273.9.peg.313 | CDS | wCle_NZ_AP013028 | 274012 |
| | fig\|246273.9.peg.532 | CDS | wCle_NZ_AP013028 | 481043 |
| | fig\|246273.9.peg.698 | CDS | wCle_NZ_AP013028 | 620688 |

| | | | | |
|---|---|---|---|---|
| | fig\|246273.9.peg.703 | CDS | wCle_NZ_AP013028 | 625421 |
| | fig\|246273.9.peg.759 | CDS | wCle_NZ_AP013028 | 676109 |
| | fig\|246273.9.peg.920 | CDS | wCle_NZ_AP013028 | 816276 |
| | fig\|246273.9.peg.921 | CDS | wCle_NZ_AP013028 | 816422 |
| | fig\|246273.9.peg.922 | CDS | wCle_NZ_AP013028 | 817421 |
| | fig\|246273.9.peg.948 | CDS | wCle_NZ_AP013028 | 836503 |
| | fig\|246273.9.peg.1081 | CDS | wCle_NZ_AP013028 | 948026 |
| | fig\|246273.9.peg.1231 | CDS | wCle_NZ_AP013028 | 1089572 |
| | fig\|246273.9.peg.1245 | CDS | wCle_NZ_AP013028 | 1099171 |
| | fig\|246273.9.peg.1247 | CDS | wCle_NZ_AP013028 | 1101100 |
| | fig\|246273.9.peg.1372 | CDS | wCle_NZ_AP013028 | 1214854 |
| | fig\|246273.9.peg.1373 | CDS | wCle_NZ_AP013028 | 1215789 |
| wDimm | fig\|1812115.6.peg.87 | CDS | canuwDimmCt1p0_circ_ | 66360 |
| | fig\|1812115.6.peg.284 | CDS | canuwDimmCt1p0_circ_ | 236311 |
| wDcau | fig\|6666666.428248.peg.164 | CDS | wDcau | 188959 |
| wFol | fig\|169402.10.peg.217 | CDS | wFol_NZ_CP015510 | 225797 |
| | fig\|169402.10.peg.234 | CDS | wFol_NZ_CP015510 | 249370 |
| | fig\|169402.10.peg.284 | CDS | wFol_NZ_CP015510 | 302682 |
| | fig\|169402.10.peg.304 | CDS | wFol_NZ_CP015510 | 321595 |
| | fig\|169402.10.peg.305 | CDS | wFol_NZ_CP015510 | 322471 |
| | fig\|169402.10.peg.335 | CDS | wFol_NZ_CP015510 | 360679 |
| | fig\|169402.10.peg.402 | CDS | wFol_NZ_CP015510 | 416676 |
| | fig\|169402.10.peg.515 | CDS | wFol_NZ_CP015510 | 570503 |
| | fig\|169402.10.peg.554 | CDS | wFol_NZ_CP015510 | 615056 |
| | fig\|169402.10.peg.602 | CDS | wFol_NZ_CP015510 | 658368 |
| | fig\|169402.10.peg.686 | CDS | wFol_NZ_CP015510 | 727231 |
| | fig\|169402.10.peg.747 | CDS | wFol_NZ_CP015510 | 802088 |
| | fig\|169402.10.peg.1005 | CDS | wFol_NZ_CP015510 | 1066904 |
| | fig\|169402.10.peg.1041 | CDS | wFol_NZ_CP015510 | 1115293 |
| | fig\|169402.10.peg.1092 | CDS | wFol_NZ_CP015510 | 1159631 |
| | fig\|169402.10.peg.1093 | CDS | wFol_NZ_CP015510 | 1160371 |
| | fig\|169402.10.peg.1176 | CDS | wFol_NZ_CP015510 | 1250521 |
| | fig\|169402.10.peg.1195 | CDS | wFol_NZ_CP015510 | 1286321 |
| wMel | fig\|163164.1.peg.32 | CDS | wMel_NC_002978 | 37671 |
| | fig\|163164.1.peg.65 | CDS | wMel_NC_002978 | 68396 |
| | fig\|163164.1.peg.130 | CDS | wMel_NC_002978 | 133174 |
| | fig\|163164.1.peg.168 | CDS | wMel_NC_002978 | 175126 |
| | fig\|163164.1.peg.252 | CDS | wMel_NC_002978 | 265391 |
| | fig\|163164.1.peg.253 | CDS | wMel_NC_002978 | 266255 |
| | fig\|163164.1.peg.258 | CDS | wMel_NC_002978 | 271048 |
| | fig\|163164.1.peg.259 | CDS | wMel_NC_002978 | 273255 |
| | fig\|163164.1.peg.261 | CDS | wMel_NC_002978 | 275225 |

| | | | | |
|---|---|---|---|---|
| | fig\|163164.1.peg.342 | CDS | wMel_NC_002978 | 367326 |
| | fig\|163164.1.peg.392 | CDS | wMel_NC_002978 | 422022 |
| | fig\|163164.1.peg.395 | CDS | wMel_NC_002978 | 425912 |
| | fig\|163164.1.peg.446 | CDS | wMel_NC_002978 | 479223 |
| | fig\|163164.1.peg.459 | CDS | wMel_NC_002978 | 507200 |
| | fig\|163164.1.peg.489 | CDS | wMel_NC_002978 | 538083 |
| | fig\|163164.1.peg.503 | CDS | wMel_NC_002978 | 551597 |
| | fig\|163164.1.peg.530 | CDS | wMel_NC_002978 | 573982 |
| | fig\|163164.1.peg.563 | CDS | wMel_NC_002978 | 623343 |
| | fig\|163164.1.peg.566 | CDS | wMel_NC_002978 | 629024 |
| | fig\|163164.1.peg.567 | CDS | wMel_NC_002978 | 629676 |
| | fig\|163164.1.peg.675 | CDS | wMel_NC_002978 | 729022 |
| | fig\|163164.1.peg.687 | CDS | wMel_NC_002978 | 738997 |
| | fig\|163164.1.peg.1090 | CDS | wMel_NC_002978 | 1160993 |
| wMhie | fig\|1241303.4.peg.485 | CDS | NODE_2425_length_89 | 4985 |
| | fig\|1241303.4.peg.533 | CDS | NODE_2622_length_81 | 8114 |
| | fig\|1241303.4.peg.833 | CDS | NODE_4004_length_48 | 4187 |
| | fig\|1241303.4.peg.889 | CDS | NODE_4371_length_43 | 333 |
| | fig\|1241303.4.peg.1182 | CDS | NODE_7789_length_21 | 1568 |
| wLbra | fig\|1812117.6.peg.155 | CDS | NODE_153_length_129 | 37638 |
| | fig\|1812117.6.peg.1035 | CDS | NODE_444_length_408 | 224 |
| | fig\|1812117.6.peg.1038 | CDS | NODE_444_length_408 | 1246 |
| wLsig | fig\|80850.8.peg.1170 | CDS | canuwLsigCt1p4_wLsig_ | 833976 |
| wLug | fig\|1335053.6.peg.6 | CDS | MUIY01000001.1 | 3064 |
| | fig\|1335053.6.peg.89 | CDS | MUIY01000001.1 | 80198 |
| | fig\|1335053.6.peg.323 | CDS | MUIY01000001.1 | 304990 |
| | fig\|1335053.6.peg.434 | CDS | MUIY01000001.1 | 417440 |
| | fig\|1335053.6.peg.435 | CDS | MUIY01000001.1 | 418000 |
| | fig\|1335053.6.peg.447 | CDS | MUIY01000001.1 | 428259 |
| | fig\|1335053.6.peg.627 | CDS | MUIY01000001.1 | 584055 |
| | fig\|1335053.6.peg.743 | CDS | MUIY01000001.1 | 684266 |
| | fig\|1335053.6.peg.746 | CDS | MUIY01000001.1 | 687966 |
| | fig\|1335053.6.peg.768 | CDS | MUIY01000001.1 | 709727 |
| | fig\|1335053.6.peg.774 | CDS | MUIY01000001.1 | 717607 |
| | fig\|1335053.6.peg.784 | CDS | MUIY01000001.1 | 726030 |
| | fig\|1335053.6.peg.996 | CDS | MUIY01000001.1 | 934070 |
| | fig\|1335053.6.peg.1065 | CDS | MUIY01000001.1 | 1002512 |
| | fig\|1335053.6.peg.1194 | CDS | MUIY01000001.1 | 1121977 |
| | fig\|1335053.6.peg.1341 | CDS | MUIY01000002.1 | 122358 |
| | fig\|1335053.6.peg.1348 | CDS | MUIY01000002.1 | 129065 |
| | fig\|1335053.6.peg.1457 | CDS | MUIY01000002.1 | 230560 |
| | fig\|1335053.6.peg.1490 | CDS | MUIY01000002.1 | 266538 |

| | | | | |
|---|---|---|---|---|
| | fig\|1335053.6.peg.1491 | CDS | MUIY01000002.1 | 267549 |
| | fig\|1335053.6.peg.1492 | CDS | MUIY01000002.1 | 267878 |
| | fig\|1335053.6.peg.1601 | CDS | MUIY01000002.1 | 360277 |
| wNfla | fig\|1854759.6.peg.321 | CDS | NZ_LYUW01000008.1 | 18514 |
| | fig\|1854759.6.peg.371 | CDS | NZ_LYUW01000010.1 | 15180 |
| | fig\|1854759.6.peg.421 | CDS | NZ_LYUW01000012.1 | 10105 |
| | fig\|1854759.6.peg.422 | CDS | NZ_LYUW01000012.1 | 10236 |
| | fig\|1854759.6.peg.555 | CDS | NZ_LYUW01000018.1 | 17712 |
| | fig\|1854759.6.peg.580 | CDS | NZ_LYUW01000019.1 | 19030 |
| | fig\|1854759.6.peg.647 | CDS | NZ_LYUW01000023.1 | 47 |
| | fig\|1854759.6.peg.651 | CDS | NZ_LYUW01000023.1 | 3869 |
| | fig\|1854759.6.peg.817 | CDS | NZ_LYUW01000033.1 | 12267 |
| | fig\|1854759.6.peg.1038 | CDS | NZ_LYUW01000049.1 | 1789 |
| | fig\|1854759.6.peg.1098 | CDS | NZ_LYUW01000054.1 | 7183 |
| | fig\|1854759.6.peg.1125 | CDS | NZ_LYUW01000057.1 | 925 |
| | fig\|1854759.6.peg.1156 | CDS | NZ_LYUW01000060.1 | 5845 |
| | fig\|1854759.6.peg.1157 | CDS | NZ_LYUW01000060.1 | 6693 |
| | fig\|1854759.6.peg.1195 | CDS | NZ_LYUW01000064.1 | 5621 |
| | fig\|1854759.6.peg.1196 | CDS | NZ_LYUW01000064.1 | 6992 |
| | fig\|1854759.6.peg.1223 | CDS | NZ_LYUW01000068.1 | 4831 |
| | fig\|1854759.6.peg.1260 | CDS | NZ_LYUW01000073.1 | 1591 |
| | fig\|1854759.6.peg.1319 | CDS | NZ_LYUW01000081.1 | 3885 |
| | fig\|1854759.6.peg.1356 | CDS | NZ_LYUW01000089.1 | 47 |
| | fig\|1854759.6.peg.1442 | CDS | NZ_LYUW01000111.1 | 335 |
| wOo | fig\|6666666.201426.peg.17 | CDS | wOo | 14070 |
| | fig\|6666666.201426.peg.249 | CDS | wOo | 275178 |
| wOv | fig\|1410384.7.peg.552 | CDS | wOo_NZ_HG810405 | 609948 |
| | fig\|1410384.7.peg.784 | CDS | wOo_NZ_HG810405 | 871356 |
| wPip | fig\|955.1.peg.8 | CDS | wPip_culex15b06 | 8463 |
| | fig\|955.1.peg.9 | CDS | wPip_culex15b06 | 10040 |
| | fig\|955.1.peg.112 | CDS | culex8d01.q1ka | 22400 |
| | fig\|955.1.peg.144 | CDS | culex8d01.q1ka | 58928 |
| | fig\|955.1.peg.157 | CDS | culex8d01.q1ka | 75760 |
| | fig\|955.1.peg.325 | CDS | culex61a11.q1k | 3975 |
| | fig\|955.1.peg.447 | CDS | culex17f04.q1k | 2806 |
| | fig\|955.1.peg.479 | CDS | culex141a08.q1k | 1463 |
| | fig\|955.1.peg.570 | CDS | culex107d08.p1k | 5240 |
| | fig\|955.1.peg.742 | CDS | culex111d03.p1k | 20757 |
| | fig\|955.1.peg.749 | CDS | culex111d03.p1k | 32311 |
| | fig\|955.1.peg.863 | CDS | culex40h12.p1k | 18377 |
| | fig\|955.1.peg.913 | CDS | culex18h04.p1k | 2125 |
| | fig\|955.1.peg.1011 | CDS | culex180a01.p1k | 7163 |

| | | | | |
|---|---|---|---|---|
| | fig\|955.1.peg.1179 | CDS | culex79g04.p1k | 57593 |
| | fig\|955.1.peg.1183 | CDS | culex79g04.p1k | 61738 |
| | fig\|955.1.peg.1192 | CDS | culex79g04.p1k | 73793 |
| | fig\|955.1.peg.1504 | CDS | culex160e12.q1k | 22114 |
| wPpe | fig\|6666666.425803.peg.122 | CDS | wPpe_NZ_MJMG01000( | 110793 |
| | fig\|6666666.425803.peg.393 | CDS | NZ_MJMG01000002.1 | 18926 |
| | fig\|6666666.425803.peg.443 | CDS | NZ_MJMG01000004.1 | 6283 |
| | fig\|6666666.425803.peg.605 | CDS | NZ_MJMG01000006.1 | 71915 |
| | fig\|6666666.425803.peg.984 | CDS | NZ_MJMG01000011.1 | 32291 |
| wstri | fig\|368602.8.peg.7 | CDS | MUIX01000001.1 | 7598 |
| | fig\|368602.8.peg.49 | CDS | MUIX01000001.1 | 43212 |
| | fig\|368602.8.peg.157 | CDS | MUIX01000001.1 | 139951 |
| | fig\|368602.8.peg.244 | CDS | MUIX01000001.1 | 234409 |
| | fig\|368602.8.peg.248 | CDS | MUIX01000001.1 | 239031 |
| | fig\|368602.8.peg.286 | CDS | MUIX01000001.1 | 263993 |
| | fig\|368602.8.peg.293 | CDS | MUIX01000001.1 | 277159 |
| | fig\|368602.8.peg.482 | CDS | MUIX01000001.1 | 456434 |
| | fig\|368602.8.peg.492 | CDS | MUIX01000001.1 | 464826 |
| | fig\|368602.8.peg.496 | CDS | MUIX01000001.1 | 472000 |
| | fig\|368602.8.peg.517 | CDS | MUIX01000001.1 | 492284 |
| | fig\|368602.8.peg.642 | CDS | MUIX01000001.1 | 640888 |
| | fig\|368602.8.peg.733 | CDS | MUIX01000001.1 | 743574 |
| | fig\|368602.8.peg.768 | CDS | MUIX01000001.1 | 782603 |
| | fig\|368602.8.peg.937 | CDS | MUIX01000001.1 | 935303 |
| | fig\|368602.8.peg.994 | CDS | MUIX01000001.1 | 983420 |
| | fig\|368602.8.peg.1003 | CDS | MUIX01000001.1 | 990385 |
| | fig\|368602.8.peg.1131 | CDS | MUIX01000001.1 | 1121774 |
| | fig\|368602.8.peg.1228 | CDS | MUIX01000001.1 | 1209426 |
| | fig\|368602.8.peg.1290 | CDS | MUIX01000001.1 | 1270986 |
| | fig\|368602.8.peg.1363 | CDS | MUIX01000001.1 | 1331331 |
| | fig\|368602.8.peg.1392 | CDS | MUIX01000001.1 | 1371763 |
| | fig\|368602.8.peg.1473 | CDS | MUIX01000001.1 | 1442122 |
| | fig\|368602.8.peg.1520 | CDS | MUIX01000001.1 | 1486802 |
| | fig\|368602.8.peg.1608 | CDS | MUIX01000001.1 | 1584441 |
| | fig\|368602.8.peg.1622 | CDS | MUIX01000001.1 | 1594252 |
| | fig\|368602.8.peg.1638 | CDS | MUIX01000002.1 | 5506 |
| | fig\|368602.8.peg.1672 | CDS | MUIX01000002.1 | 38437 |
| | fig\|368602.8.peg.1757 | CDS | MUIX01000002.1 | 104744 |
| | fig\|368602.8.peg.1822 | CDS | MUIX01000002.1 | 178919 |
| wTpre | fig\|125593.4.peg.1 | CDS | NZ_CM003641.1 | 493 |
| | fig\|125593.4.peg.40 | CDS | NZ_CM003641.1 | 33476 |
| | fig\|125593.4.peg.42 | CDS | NZ_CM003641.1 | 34357 |

| | | | |
|---|---|---|---|
| fig\|125593.4.peg.43 | CDS | NZ_CM003641.1 | 34519 |
| fig\|125593.4.peg.44 | CDS | NZ_CM003641.1 | 34699 |
| fig\|125593.4.peg.175 | CDS | NZ_CM003641.1 | 135581 |
| fig\|125593.4.peg.177 | CDS | NZ_CM003641.1 | 136658 |
| fig\|125593.4.peg.178 | CDS | NZ_CM003641.1 | 137078 |
| fig\|125593.4.peg.224 | CDS | NZ_CM003641.1 | 169242 |
| fig\|125593.4.peg.225 | CDS | NZ_CM003641.1 | 169393 |
| fig\|125593.4.peg.226 | CDS | NZ_CM003641.1 | 169778 |
| fig\|125593.4.peg.370 | CDS | NZ_CM003641.1 | 289388 |
| fig\|125593.4.peg.371 | CDS | NZ_CM003641.1 | 289864 |
| fig\|125593.4.peg.415 | CDS | NZ_CM003641.1 | 325025 |
| fig\|125593.4.peg.459 | CDS | NZ_CM003641.1 | 362087 |
| fig\|125593.4.peg.460 | CDS | NZ_CM003641.1 | 362339 |
| fig\|125593.4.peg.461 | CDS | NZ_CM003641.1 | 362815 |
| fig\|125593.4.peg.539 | CDS | NZ_CM003641.1 | 437432 |
| fig\|125593.4.peg.549 | CDS | NZ_CM003641.1 | 445846 |
| fig\|125593.4.peg.664 | CDS | NZ_CM003641.1 | 531803 |
| fig\|125593.4.peg.666 | CDS | NZ_CM003641.1 | 532311 |
| fig\|125593.4.peg.762 | CDS | NZ_CM003641.1 | 624401 |
| fig\|125593.4.peg.763 | CDS | NZ_CM003641.1 | 624531 |
| fig\|125593.4.peg.764 | CDS | NZ_CM003641.1 | 624695 |
| fig\|125593.4.peg.765 | CDS | NZ_CM003641.1 | 624874 |
| fig\|125593.4.peg.766 | CDS | NZ_CM003641.1 | 625072 |
| fig\|125593.4.peg.767 | CDS | NZ_CM003641.1 | 625416 |
| fig\|125593.4.peg.776 | CDS | NZ_CM003641.1 | 630150 |
| fig\|125593.4.peg.777 | CDS | NZ_CM003641.1 | 630329 |
| fig\|125593.4.peg.778 | CDS | NZ_CM003641.1 | 630777 |
| fig\|125593.4.peg.883 | CDS | NZ_CM003641.1 | 711136 |
| fig\|125593.4.peg.946 | CDS | NZ_CM003641.1 | 762030 |
| fig\|125593.4.peg.1077 | CDS | NZ_CM003641.1 | 858481 |
| fig\|125593.4.peg.1078 | CDS | NZ_CM003641.1 | 858894 |
| fig\|125593.4.peg.1103 | CDS | NZ_CM003641.1 | 880600 |
| fig\|125593.4.peg.1137 | CDS | NZ_CM003641.1 | 907323 |
| fig\|125593.4.peg.1138 | CDS | NZ_CM003641.1 | 907803 |
| fig\|125593.4.peg.1139 | CDS | NZ_CM003641.1 | 907998 |
| fig\|125593.4.peg.1140 | CDS | NZ_CM003641.1 | 908599 |
| fig\|125593.4.peg.1189 | CDS | NZ_CM003641.1 | 951499 |
| fig\|125593.4.peg.1190 | CDS | NZ_CM003641.1 | 951812 |
| fig\|125593.4.peg.1251 | CDS | NZ_CM003641.1 | 1009740 |
| fig\|125593.4.peg.1335 | CDS | NZ_CM003641.1 | 1094662 |
| fig\|125593.4.peg.1340 | CDS | NZ_CM003641.1 | 1096663 |
| fig\|125593.4.peg.1341 | CDS | NZ_CM003641.1 | 1096860 |

| | | | | |
|---|---|---|---|---|
| | fig\|125593.4.peg.1342 | CDS | NZ_CM003641.1 | 1097018 |
| | fig\|125593.4.peg.1371 | CDS | NZ_CM003641.1 | 1120976 |
| | fig\|125593.4.peg.1379 | CDS | NZ_CM003641.1 | 1130329 |
| | fig\|125593.4.peg.1382 | CDS | NZ_CM003641.1 | 1133807 |
| wVulC | fig\|1220511.6.peg.182 | CDS | ALWU01000001.1 | 169776 |
| | fig\|1220511.6.peg.256 | CDS | ALWU01000001.1 | 226104 |
| | fig\|1220511.6.peg.488 | CDS | ALWU01000001.1 | 418030 |
| | fig\|1220511.6.peg.516 | CDS | ALWU01000001.1 | 435504 |
| | fig\|1220511.6.peg.562 | CDS | ALWU01000001.1 | 470805 |
| | fig\|1220511.6.peg.569 | CDS | ALWU01000001.1 | 481106 |
| | fig\|1220511.6.peg.636 | CDS | ALWU01000001.1 | 548972 |
| | fig\|1220511.6.peg.683 | CDS | ALWU01000002.1 | 1104 |
| | fig\|1220511.6.peg.702 | CDS | ALWU01000002.1 | 16677 |
| | fig\|1220511.6.peg.835 | CDS | ALWU01000005.1 | 18276 |
| | fig\|1220511.6.peg.838 | CDS | ALWU01000005.1 | 21959 |
| | fig\|1220511.6.peg.1085 | CDS | ALWU01000006.1 | 9977 |
| | fig\|1220511.6.peg.1118 | CDS | ALWU01000006.1 | 35307 |
| | fig\|1220511.6.peg.1145 | CDS | ALWU01000008.1 | 9582 |
| | fig\|1220511.6.peg.1160 | CDS | ALWU01000008.1 | 25019 |
| | fig\|1220511.6.peg.1303 | CDS | ALWU01000008.1 | 159709 |
| | fig\|1220511.6.peg.1441 | CDS | ALWU01000010.1 | 73725 |
| | fig\|1220511.6.peg.1507 | CDS | ALWU01000010.1 | 130853 |
| | fig\|1220511.6.peg.1615 | CDS | ALWU01000010.1 | 214812 |
| | fig\|1220511.6.peg.1827 | CDS | ALWU01000010.1 | 401103 |
| | fig\|1220511.6.peg.1861 | CDS | ALWU01000010.1 | 436267 |
| | fig\|1220511.6.peg.1862 | CDS | ALWU01000010.1 | 437153 |
| | fig\|1220511.6.peg.1863 | CDS | ALWU01000010.1 | 438568 |
| | fig\|1220511.6.peg.1881 | CDS | ALWU01000010.1 | 452153 |
| wWb | fig\|96496.9.peg.22 | CDS | wWb_NZ_NJBR0200000 | 23837 |
| | fig\|96496.9.peg.46 | CDS | wWb_NZ_NJBR0200000 | 42646 |
| | fig\|96496.9.peg.514 | CDS | NZ_NJBR02000035.1 | 859 |
| | fig\|96496.9.peg.517 | CDS | NZ_NJBR02000035.1 | 1422 |
| | fig\|96496.9.peg.620 | CDS | NZ_NJBR02000044.1 | 5792 |
| | fig\|96496.9.peg.621 | CDS | NZ_NJBR02000044.1 | 5997 |
| | fig\|96496.9.peg.741 | CDS | NZ_NJBR02000058.1 | 2128 |
| | fig\|96496.9.peg.904 | CDS | NZ_NJBR02000073.1 | 25943 |
| | fig\|96496.9.peg.906 | CDS | NZ_NJBR02000073.1 | 26461 |
| | fig\|96496.9.peg.914 | CDS | NZ_NJBR02000073.1 | 31586 |
| | fig\|96496.9.peg.1012 | CDS | NZ_NJBR02000084.1 | 497 |

| Stop | Strand | Length (bp | Function | Subsystem |
|---|---|---|---|---|
| 72863 | - | 240 | Ankyrin repeat domain protein | - none - |
| 73522 | - | 342 | Ankyrin repeat domain protein | - none - |
| 91302 | - | 1341 | Ankyrin repeat domain protein | - none - |
| 243626 | - | 375 | ankyrin repeat domain protein | - none - |
| 246289 | - | 177 | ankyrin repeat domain protein | - none - |
| 303566 | - | 2091 | Ankyrin 3 | - none - |
| 527916 | + | 120 | ankyrin repeat domain protein | - none - |
| 528469 | + | 240 | Ankyrin repeat domain protein | - none - |
| 590702 | - | 555 | Ankyrin 1 | - none - |
| 789783 | + | 246 | Ankyrin repeat-containing protein | - none - |
| 802404 | + | 1323 | Ankyrin repeat domain protein | - none - |
| 914147 | - | 459 | Ankyrin repeat-containing protein | - none - |
| 931234 | - | 318 | Ankyrin 1 | - none - |
| 997453 | + | 3327 | Ankyrin repeat domain protein | - none - |
| 68694 | + | 147 | ankyrin repeat domain protein | - none - |
| 68994 | + | 144 | ankyrin repeat domain protein | - none - |
| 107738 | + | 1341 | Ankyrin repeat domain protein | - none - |
| 125528 | + | 342 | Ankyrin repeat domain protein | - none - |
| 126187 | + | 144 | Ankyrin repeat domain protein | - none - |
| 281047 | - | 3327 | Ankyrin repeat domain protein | - none - |
| 360915 | + | 582 | Ankyrin repeat-containing protein | - none - |
| 521258 | + | 1323 | Ankyrin repeat domain protein | - none - |
| 564788 | - | 132 | Ankyrin repeat-containing protein | - none - |
| 564962 | - | 330 | Ankyrin repeat-containing protein | - none - |
| 682525 | + | 555 | Ankyrin 1 | - none - |
| 826194 | - | 237 | ankyrin repeat domain protein | - none - |
| 828717 | - | 177 | ankyrin repeat domain protein | - none - |
| 885962 | - | 2091 | Ankyrin 3 | - none - |
| 964104 | - | 147 | Ankyrin repeat domain protein | - none - |
| 964658 | - | 255 | ankyrin repeat domain protein | - none - |
| 183538 | - | 1212 | ankyrin repeat domain protein, putative | - none - |
| 263965 | + | 711 | FOG: Ankyrin repeat | - none - |
| 293663 | - | 540 | ankyrin repeat domain protein | - none - |
| 308959 | + | 537 | ankyrin repeat domain protein | - none - |
| 405075 | + | 2772 | ankyrin repeat domain protein | - none - |
| 441520 | - | 1056 | ankyrin repeat domain protein | - none - |
| 445711 | - | 516 | ankyrin repeat domain protein | - none - |
| 480694 | - | 867 | ankyrin repeat domain protein | - none - |
| 531350 | - | 819 | ankyrin repeat domain protein | - none - |

| | | | | |
|---|---|---|---|---|
| 596529 | + | 1194 | ankyrin repeat domain protein | - none - |
| 774353 | - | 957 | ankyrin repeat domain protein | - none - |
| 782645 | + | 486 | Ankyrin 2,3/unc44 | - none - |
| 886486 | + | 1284 | ankyrin repeat domain protein | - none - |
| 940311 | - | 2823 | ankyrin repeat domain protein | - none - |
| 1092034 | + | 711 | FOG: Ankyrin repeat | - none - |
| 1120923 | + | 732 | ankyrin repeat domain protein | - none - |
| 1177956 | - | 2577 | ankyrin repeat domain protein | - none - |
| 1183206 | - | 1203 | Ankyrin repeat domain protein | - none - |
| 1244204 | + | 1365 | ankyrin repeat domain protein | - none - |
| 66523 | + | 1233 | ankyrin repeat domain protein | - none - |
| 772614 | - | 723 | Ankyrin repeat domain protein | - none - |
| 19054 | - | 591 | ankyrin repeat domain protein | - none - |
| 21877 | - | 2859 | ankyrin repeat domain protein | - none - |
| 157360 | - | 1908 | ankyrin repeat domain protein | - none - |
| 254228 | + | 1383 | ankyrin repeat domain protein | - none - |
| 351184 | + | 3246 | Ankyrin 3 | - none - |
| 513063 | + | 624 | ankyrin repeat domain protein | - none - |
| 531325 | - | 1224 | Ankyrin repeat domain protein | - none - |
| 837399 | + | 1266 | similar to ankyrin 2,3/unc44 | - none - |
| 982188 | + | 1569 | ankyrin repeat protein, putative | - none - |
| 1060123 | + | 837 | Ankyrin | - none - |
| 1065370 | - | 2805 | ankyrin repeat domain protein | - none - |
| 1179558 | - | 1062 | ankyrin repeat domain protein | - none - |
| 221640 | - | 1878 | Ankyrin | - none - |
| 262080 | - | 1209 | ankyrin repeat domain protein | - none - |
| 325271 | + | 762 | ankyrin repeat domain protein | - none - |
| 459788 | - | 1188 | Ankyrin repeat domain protein | - none - |
| 577058 | + | 549 | ankyrin repeat domain protein | - none - |
| 684055 | - | 480 | ankyrin repeat domain protein | - none - |
| 764476 | + | 1017 | ankyrin repeat domain protein | - none - |
| 805116 | + | 240 | ankyrin repeat domain protein | - none - |
| 1429915 | + | 522 | ankyrin repeat domain protein | - none - |
| 1433331 | + | 789 | ankyrin repeat domain protein | - none - |
| 145662 | + | 1413 | ankyrin repeat domain protein | - none - |
| 157943 | - | 240 | ankyrin repeat domain protein | - none - |
| 264079 | + | 1440 | ankyrin repeat domain protein | - none - |
| 269130 | + | 597 | Ankyrin | - none - |
| 269934 | + | 570 | prophage LambdaW1, ankyrin repeat doma | - none - |
| 275277 | + | 1266 | ankyrin repeat domain protein | - none - |
| 480465 | - | 579 | Ankyrin | - none - |
| 621938 | + | 1251 | ankyrin repeat domain protein | - none - |

| | | | | |
|---|---|---|---|---|
| 625684 | + | 264 | ankyrin repeat domain protein | - none - |
| 674745 | - | 1365 | ankyrin repeat domain protein | - none - |
| 815854 | - | 423 | prophage LambdaW1, ankyrin repeat doma | - none - |
| 816273 | - | 150 | prophage LambdaW1, ankyrin repeat doma | - none - |
| 816657 | - | 765 | Ankyrin | - none - |
| 837048 | + | 546 | ankyrin repeat domain protein | - none - |
| 948271 | + | 246 | ankyrin repeat domain protein | - none - |
| 1088325 | - | 1248 | Ankyrin repeat domain protein | - none - |
| 1099581 | + | 411 | ankyrin domain protein ank2 | - none - |
| 1101756 | + | 657 | ankyrin repeat protein, putative | - none - |
| 1214645 | - | 210 | ankyrin repeat domain protein | - none - |
| 1214950 | - | 840 | ankyrin repeat domain protein | - none - |
| 63784 | - | 2577 | Ankyrin repeat domain protein | - none - |
| 237453 | + | 1143 | Ankyrin repeat domain protein | - none - |
| 187757 | - | 1203 | Ankyrin repeat domain protein | - none - |
| 227014 | + | 1218 | ankyrin repeat protein, putative | - none - |
| 247841 | - | 1530 | ankyrin repeat domain protein | - none - |
| 301432 | - | 1251 | Ankyrin repeat domain protein | - none - |
| 320990 | - | 606 | Ankyrin | - none - |
| 321800 | - | 672 | Ankyrin | - none - |
| 359891 | - | 789 | Putative ankyrin repeat protein RF_0381 | - none - |
| 417902 | + | 1227 | ankyrin repeat domain protein | - none - |
| 571501 | + | 999 | ankyrin repeat domain protein | - none - |
| 618346 | + | 3291 | ankyrin domain protein ank2 | - none - |
| 657838 | - | 531 | ankyrin repeat domain protein | - none - |
| 727890 | + | 660 | Ankyrin | - none - |
| 803296 | + | 1209 | ankyrin repeat domain protein | - none - |
| 1066380 | - | 525 | ankyrin repeat domain protein | - none - |
| 1114622 | - | 672 | Ankyrin | - none - |
| 1160209 | + | 579 | ankyrin repeat domain protein | - none - |
| 1160940 | + | 570 | ankyrin repeat domain protein | - none - |
| 1251180 | + | 660 | Ankyrin | - none - |
| 1283064 | - | 3258 | Ankyrin 3 | - none - |
| 38537 | + | 867 | ankyrin repeat domain protein | - none - |
| 70798 | + | 2403 | ankyrin repeat domain protein | - none - |
| 136038 | + | 2865 | ankyrin repeat domain protein | - none - |
| 174320 | - | 807 | ankyrin repeat domain protein | - none - |
| 265993 | + | 603 | prophage LambdaW1, ankyrin repeat doma | - none - |
| 266845 | + | 591 | prophage LambdaW1, ankyrin repeat doma | - none - |
| 270374 | - | 675 | prophage LambdaW1, ankyrin repeat doma | - none - |
| 271150 | - | 2106 | prophage LambdaW1, ankyrin repeat doma | - none - |
| 273600 | - | 1626 | ankyrin repeat domain protein | - none - |

| | | | | |
|---|---|---|---|---|
| 368954 | + | 1629 | ankyrin repeat domain protein | - none - |
| 420124 | - | 1899 | ankyrin repeat domain protein | - none - |
| 424704 | - | 1209 | ankyrin repeat domain protein | - none - |
| 480230 | + | 1008 | ankyrin repeat domain protein | - none - |
| 505791 | - | 1410 | ankyrin repeat domain protein | - none - |
| 537094 | - | 990 | ankyrin repeat domain protein | - none - |
| 551076 | - | 522 | ankyrin repeat domain protein | - none - |
| 575463 | + | 1482 | prophage LambdaW4, ankyrin repeat doma | - none - |
| 626243 | + | 2901 | prophage LambdaW5, ankyrin repeat doma | - none - |
| 628566 | - | 459 | prophage LambdaW5, ankyrin repeat doma | - none - |
| 629050 | - | 627 | prophage LambdaW5, ankyrin repeat doma | - none - |
| 727721 | - | 1302 | ankyrin repeat domain protein | - none - |
| 740421 | + | 1425 | ankyrin repeat domain protein | - none - |
| 1159749 | - | 1245 | ankyrin repeat domain protein, putative | - none - |
| 6232 | + | 1248 | Ankyrin repeat domain protein | - none - |
| 6366 | - | 1749 | ankyrin repeat domain protein | - none - |
| 4699 | + | 513 | ankyrin repeat domain protein | - none - |
| 2372 | + | 2040 | Ankyrin repeat domain protein | - none - |
| 204 | - | 1365 | ankyrin repeat domain protein | - none - |
| 40688 | + | 3051 | Ankyrin repeat domain protein | - none - |
| 358 | + | 135 | Ankyrin repeat domain protein | - none - |
| 1518 | + | 273 | Ankyrin repeat domain protein | - none - |
| 830653 | - | 3324 | Ankyrin repeat domain protein | - none - |
| 3606 | + | 543 | ankyrin repeat domain protein | - none - |
| 80731 | + | 534 | ankyrin repeat domain protein | - none - |
| 304124 | - | 867 | ankyrin repeat domain protein | - none - |
| 417153 | - | 288 | ankyrin repeat protein, putative | - none - |
| 417443 | - | 558 | ankyrin repeat protein, putative | - none - |
| 428426 | + | 168 | ankyrin repeat protein, putative | - none - |
| 582916 | - | 1140 | ankyrin repeat domain protein, putative | - none - |
| 682170 | - | 2097 | ankyrin repeat domain protein | - none - |
| 686911 | - | 1056 | Ankyrin repeat domain protein | - none - |
| 711067 | + | 1341 | ankyrin repeat domain protein | - none - |
| 716420 | - | 1188 | ankyrin repeat domain protein | - none - |
| 725113 | - | 918 | ankyrin domain protein ank2 | - none - |
| 933273 | - | 798 | ankyrin repeat domain protein | - none - |
| 1000857 | - | 1656 | ankyrin repeat domain protein | - none - |
| 1121066 | - | 912 | ankyrin repeat domain protein | - none - |
| 121471 | - | 888 | ankyrin repeat protein, putative | - none - |
| 127929 | - | 1137 | ankyrin domain protein ank2 | - none - |
| 234033 | + | 3474 | Ankyrin 3 | - none - |
| 265498 | - | 1041 | ankyrin repeat domain protein | - none - |

| | | | | |
|---|---|---|---|---|
| 266959 | - | 591 | ankyrin repeat domain protein | - none - |
| 267594 | - | 285 | ankyrin repeat domain protein | - none - |
| 360813 | + | 537 | ankyrin repeat domain protein | - none - |
| 20700 | + | 2187 | ankyrin repeat domain protein | - none - |
| 13234 | - | 1947 | ankyrin repeat domain protein | - none - |
| 9704 | - | 402 | prophage LambdaW1, ankyrin repeat doma | - none - |
| 10102 | - | 135 | prophage LambdaW1, ankyrin repeat doma | - none - |
| 17179 | - | 534 | ankyrin repeat domain protein | - none - |
| 17774 | - | 1257 | ankyrin repeat domain protein | - none - |
| 775 | + | 729 | Ankyrin repeat domain protein | - none - |
| 5746 | + | 1878 | ankyrin repeat domain protein | - none - |
| 11776 | - | 492 | ankyrin repeat domain protein | - none - |
| 2655 | + | 867 | ankyrin repeat domain protein | - none - |
| 6872 | - | 312 | ankyrin repeat domain protein | - none - |
| 29 | - | 897 | ankyrin repeat domain protein | - none - |
| 6642 | + | 798 | ankyrin repeat domain protein, putative | - none - |
| 7043 | + | 351 | ankyrin repeat domain protein, putative | - none - |
| 5034 | - | 588 | ankyrin repeat domain protein | - none - |
| 5763 | - | 1230 | ankyrin repeat domain protein | - none - |
| 3896 | - | 936 | ankyrin repeat domain protein | - none - |
| 1866 | + | 276 | ankyrin repeat domain protein | - none - |
| 4409 | + | 525 | ankyrin repeat domain protein | - none - |
| 397 | + | 351 | ankyrin repeat domain protein | - none - |
| 1699 | + | 1365 | ankyrin repeat domain protein | - none - |
| 13042 | - | 1029 | ankyrin repeat domain protein | - none - |
| 274990 | - | 189 | Ankyrin 3 | - none - |
| 608794 | - | 1155 | ankyrin repeat domain protein | - none - |
| 871168 | - | 189 | Ankyrin 3 | - none - |
| 9614 | + | 1152 | UNC-44 ankyrins | - none - |
| 11734 | + | 1695 | ankyrin repeat family protein | - none - |
| 23569 | + | 1170 | Ankyrin 2 | - none - |
| 55518 | - | 3411 | Ankyrin 3 | - none - |
| 76620 | + | 861 | Ankyrin 1 | - none - |
| 3112 | - | 864 | ankyrin repeat family protein | - none - |
| 2282 | - | 525 | Chain A, Sank E3_5: An Artificial Ankyrin Rep | - none - |
| 921 | - | 543 | Ankyrin | - none - |
| 3 | - | 5238 | UNC-44 ankyrins | - none - |
| 18430 | - | 2328 | Ankyrin 1 | - none - |
| 29954 | - | 2358 | Ankyrin 2 | - none - |
| 10131 | - | 8247 | Ankyrin 2 | - none - |
| 2649 | + | 525 | Chain A, Sank E3_5: An Artificial Ankyrin Rep | - none - |
| 9139 | + | 1977 | Ankyrin 3 | - none - |

| | | | | |
|---|---|---|---|---|
| 58432 | + | 840 | Ankyrin | - none - |
| 62283 | + | 546 | Ankyrin 2 | - none - |
| 74866 | + | 1074 | Ankyrin 1 | - none - |
| 22710 | + | 597 | Ankyrin 1 | - none - |
| 109780 | - | 1014 | Ankyrin repeat domain protein | - none - |
| 17715 | - | 1212 | ankyrin repeat domain protein | - none - |
| 5336 | - | 948 | FOG: Ankyrin repeat | - none - |
| 70542 | - | 1374 | ankyrin repeat domain protein | - none - |
| 33304 | + | 1014 | ankyrin repeat domain protein | - none - |
| 6870 | - | 729 | FOG: Ankyrin repeat | - none - |
| 44108 | + | 897 | ankyrin repeat domain protein | - none - |
| 140436 | + | 486 | ankyrin repeat domain protein | - none - |
| 235461 | + | 1053 | Putative ankyrin repeat protein RF_0381 | - none - |
| 238327 | - | 705 | FOG: Ankyrin repeat | - none - |
| 264355 | + | 363 | Ankyrin repeat domain protein | - none - |
| 276431 | - | 729 | FOG: Ankyrin repeat | - none - |
| 457345 | + | 912 | ankyrin domain protein ank2 | - none - |
| 466013 | + | 1188 | ankyrin repeat domain protein | - none - |
| 470660 | - | 1341 | ankyrin repeat domain protein | - none - |
| 493264 | + | 981 | Ankyrin repeat domain protein | - none - |
| 639896 | - | 993 | ankyrin repeat protein, putative | - none - |
| 747008 | + | 3435 | Ankyrin 3 | - none - |
| 782064 | - | 540 | ankyrin repeat domain protein | - none - |
| 934164 | - | 1140 | ankyrin repeat domain protein, putative | - none - |
| 984178 | + | 759 | ankyrin repeat protein, putative | - none - |
| 990140 | - | 246 | ankyrin repeat protein, putative | - none - |
| 1122640 | + | 867 | ankyrin repeat domain protein | - none - |
| 1209022 | - | 405 | ankyrin repeat domain protein | - none - |
| 1272044 | + | 1059 | similar to ankyrin 2,3/unc44 | - none - |
| 1331846 | + | 516 | ankyrin repeat domain protein | - none - |
| 1371035 | - | 729 | FOG: Ankyrin repeat | - none - |
| 1444113 | + | 1992 | ankyrin repeat domain protein | - none - |
| 1486284 | - | 519 | ankyrin repeat domain protein | - none - |
| 1583926 | - | 516 | ankyrin repeat domain protein | - none - |
| 1593737 | - | 516 | ankyrin repeat domain protein | - none - |
| 5015 | - | 492 | ankyrin repeat domain protein | - none - |
| 37748 | - | 690 | FOG: Ankyrin repeat | - none - |
| 105277 | + | 534 | ankyrin repeat domain protein | - none - |
| 179410 | + | 492 | ankyrin repeat domain protein | - none - |
| 2 | - | 492 | ankyrin repeat domain protein | - none - |
| 33024 | - | 453 | conserved hypothetical protein (Ankyrin rep | - none - |
| 33956 | - | 402 | ankyrin repeat domain protein | - none - |

| | | | | |
|---|---|---|---|---|
| 34406 | - | 114 | ankyrin repeat domain protein | - none - |
| 34565 | - | 135 | ankyrin repeat domain protein | - none - |
| 136018 | + | 438 | ankyrin repeat domain protein | - none - |
| 136957 | + | 300 | prophage LambdaW5, ankyrin repeat doma | - none - |
| 137233 | + | 156 | prophage LambdaW5, ankyrin repeat doma | - none - |
| 169376 | + | 135 | ankyrin repeat domain protein | - none - |
| 169656 | + | 264 | ankyrin repeat domain protein | - none - |
| 169927 | + | 150 | ankyrin repeat domain protein | - none - |
| 289798 | + | 411 | ankyrin repeat domain protein | - none - |
| 290004 | + | 141 | ankyrin repeat domain protein | - none - |
| 325177 | + | 153 | ankyrin repeat domain protein | - none - |
| 361986 | - | 102 | similar to ankyrin 2,3/unc44 | - none - |
| 362091 | - | 249 | similar to ankyrin 2,3/unc44 | - none - |
| 362393 | - | 423 | similar to ankyrin 2,3/unc44 | - none - |
| 437217 | - | 216 | ankyrin repeat domain protein | - none - |
| 446034 | + | 189 | ankyrin repeat domain protein | - none - |
| 531420 | - | 384 | ankyrin repeat domain protein | - none - |
| 532144 | - | 168 | ankyrin repeat domain protein | - none - |
| 624144 | - | 258 | ankyrin repeat domain protein | - none - |
| 624418 | - | 114 | ankyrin repeat domain protein | - none - |
| 624552 | - | 144 | ankyrin repeat domain protein | - none - |
| 624701 | - | 174 | ankyrin repeat domain protein | - none - |
| 624878 | - | 195 | ankyrin repeat domain protein | - none - |
| 625093 | - | 324 | ankyrin repeat domain protein | - none - |
| 630272 | + | 123 | ankyrin repeat protein, putative | - none - |
| 630628 | + | 300 | ankyrin repeat protein, putative | - none - |
| 630914 | + | 138 | ankyrin repeat protein, putative | - none - |
| 714609 | + | 3474 | Ankyrin 3 | - none - |
| 761896 | - | 135 | ankyrin repeat protein, putative | - none - |
| 858344 | - | 138 | ankyrin repeat domain protein, putative | - none - |
| 858694 | - | 201 | ankyrin repeat domain protein, putative | - none - |
| 881127 | + | 528 | ankyrin repeat domain protein | - none - |
| 907613 | + | 291 | ankyrin repeat domain protein | - none - |
| 907973 | + | 171 | ankyrin repeat domain protein | - none - |
| 908279 | + | 282 | Ankyrin 1 | - none - |
| 909780 | + | 1182 | ankyrin repeat domain protein | - none - |
| 951756 | + | 258 | ankyrin domain protein ank2 | - none - |
| 951994 | + | 183 | ankyrin domain protein ank2 | - none - |
| 1009895 | + | 156 | ankyrin repeat domain protein | - none - |
| 1094036 | - | 627 | FOG: Ankyrin repeat | - none - |
| 1096818 | + | 156 | ankyrin repeat domain protein | - none - |
| 1097021 | + | 162 | ankyrin repeat domain protein | - none - |

| | | | | |
|---|---|---|---|---|
| 1097194 | + | 177 | ankyrin repeat domain protein | - none - |
| 1122316 | + | 1341 | ankyrin repeat domain protein | - none - |
| 1128287 | - | 2043 | ankyrin repeat domain protein | - none - |
| 1132989 | - | 819 | Ankyrin repeat domain protein | - none - |
| 168436 | - | 1341 | ankyrin repeat domain protein | - none - |
| 225553 | - | 552 | Ankyrin repeat domain protein | - none - |
| 418224 | + | 195 | ankyrin repeat domain protein | - none - |
| 434161 | - | 1344 | ankyrin repeat domain protein | - none - |
| 470467 | - | 339 | Ankyrin repeat domain protein | - none - |
| 480210 | - | 897 | ankyrin repeat domain protein | - none - |
| 548259 | - | 714 | ankyrin repeat domain protein | - none - |
| 1871 | + | 768 | ankyrin repeat domain protein | - none - |
| 16141 | - | 537 | ankyrin repeat domain protein | - none - |
| 19259 | + | 984 | Ankyrin repeat domain protein | - none - |
| 24043 | + | 2085 | ankyrin repeat domain protein | - none - |
| 10678 | + | 702 | FOG: Ankyrin repeat | - none - |
| 36791 | + | 1485 | ankyrin repeat domain protein | - none - |
| 8854 | - | 729 | ankyrin repeat domain protein | - none - |
| 24630 | - | 390 | conserved hypothetical protein (Ankyrin rep | - none - |
| 160134 | + | 426 | ankyrin repeat domain protein | - none - |
| 74813 | + | 1089 | ankyrin repeat domain protein | - none - |
| 130251 | - | 603 | ankyrin domain protein ank2 | - none - |
| 215984 | + | 1173 | ankyrin repeat domain protein | - none - |
| 404606 | + | 3504 | Ankyrin 3 | - none - |
| 436094 | - | 174 | ankyrin repeat domain protein | - none - |
| 436341 | - | 813 | Ankyrin 1 | - none - |
| 437408 | - | 1161 | ankyrin repeat domain protein | - none - |
| 451014 | - | 1140 | ankyrin repeat domain protein | - none - |
| 22472 | - | 1347 | ankyrin repeat domain protein | - none - |
| 41306 | - | 1341 | ankyrin repeat domain protein | - none - |
| 981 | + | 123 | ankyrin repeat domain protein | - none - |
| 1568 | + | 147 | ankyrin repeat domain protein | - none - |
| 5676 | - | 117 | Ankyrin repeat-containing protein | - none - |
| 5758 | - | 240 | Ankyrin repeat-containing protein | - none - |
| 908 | - | 1221 | ankyrin repeat domain protein | - none - |
| 26191 | + | 249 | ankyrin repeat domain protein | - none - |
| 26613 | + | 153 | Ankyrin repeat domain protein | - none - |
| 32056 | + | 471 | Ankyrin 1 | - none - |
| 249 | - | 249 | ankyrin repeat domain protein | - none - |
