## Supplementary material for "Diminutive, degraded but dissimilar: *Wolbachia* genomes from filarial nematodes do not conform to a single paradigm": two supplemental files and eleven supplementary table: Supplementary_file_all.pdf

**Supplementary files 1: Species and Authorities**

**Nematodes:**

*Brugia malayi* (Brug, 1927)

*Brugia pahangi* (Buckley & Edeson, 1956)

*Cercopithifilaria japonica* (Uni, 1983)

*Cruorifilaria tuberocauda* Eberhard, Morales and Orihel, 1976

*Dirofilaria* (*Dirofilaria*) *immitis* (Leidy, 1856)

*Dipetalonema caudispina* (Molin, 1858)

*Dipetalonema gracile* (Rudolphi, 1809)

*Litomosoides brasiliensis* Lins de Almeida, 1936

*Litomosoides sigmodontis* Chandler, 1931

*Madathamugadia hiepei* Hering-Hagenbeck, Boomker, Petit, Killick-Kendrick and Bain, 2000

*Onchocerca ochengi* Bwangamoi, 1969

*Onchocerca volvulus* (Leuckart, 1893)

*Pratylenchus penetrans* (Cobb, 1917)

*Wuchereria bancrofti* (Cobbold, 1877)

**Arthropods:**

*Drosophila melanogaster* Meigen, 1830

*Folsomia candida* Willem, 1902

*Carposina sasakii* Matsumura, 1898

Ctenocephalides felis (Bouché, 1835)

*Nomada flava* Panzer, 1798

*Culex quinquefasciatus* Say, 1823

*Trichogramma pretiosum* (Riley, 1879)

*Nilaparvata lugens* (Stål, 1854)

*Laodelphax striatella* (Fallen, 1826)

*Armadillidium vulgare* (Latreille, 1804)

*Cimex lectularius* Linnaeus, 1758

**Vertebrates:**

Ateles paniscus (Linnaeus, 1758)

*Canis familiaris* Linnaeus, 1758

Carollia perspicillata (Linnaeus, 1758)

*Chondrodactylus turneri* (Gray, 1864)

*Hydrochoerus hydrochaeris* (Linnaeus, 1766)

*Meriones unguiculatus* (Milne-Edwards, 1867)

**Supplementary files 2 : Validation of the circularization of the *Wolbachia* complete genomes by PCR amplification.**

**Table. Primers and PCR condition.** In bold is indicated the primers which valided the circularization, all the other primers were design to verify previous de novo assembly. The primers which were tested but the PCR product were not sequenced are indicated in grey.

| reference | position (bp) | name of oligo | sequence (forward) | sequence (reverse) |
|---|---|---|---|---|
| **wCtub** | 626,596-627,580 | wCtub_tig1D1 | CAGCCAGCATGATAGGGTTA | GGAAGGTTCTATGTTGCTTTGGA[illegible] |
| | 768,172-768,952 | wCtub_N1201N5 | ATGCAGGAGCGATAGGGATTGAAC | ATGGGCAAGTACCACAATGCGT |
| | 423,432-424,095 | wCtub_N5N1205 | TGCATCACCATGACGGGAATCT | ACAAGGAGTGGAAGGCAGGTTA[illegible] |
| | 418,342-418,921 | wCtub_N1205N268 | GCTGAAAGCCTTGCATTTGCGT | TGCTCTGCATTCTGGAGTTACC |
| | 113,621-114,156 | wCtub_N40N1206 | TGAGGAGACGTTTTGGGAGACA | TTGGGAATGGTCACTACATGCC |
| | 108,955-109,492 | wCtub_N1206N54 | AGCTTCAAGACCATGGGATGAAGA | GCGTTAGTACCACCGAAGCCAA[illegible] |
| | 774,966-775,932 | wCtub_N54N1201 | CACGGAGATGGCCAAGTATTACTG | TTGTATGAAAGGCGGTCAAGTG[illegible] |
| | 348,986-349,631 | wCtub_N268E | AACCTTTGTCCTCGTCCTGATG | TTCTACCCAATCAGGCAAAAGG[illegible] |
| | **522,139-522915** | **wCtub_P341224** | **GTATTTACTGCAGTCGCTGGAG** | **GGGAACCTTAACACCTTTAGCC** |
| | 167,628-168,831 | wCtub_P695900 | GCACTGCCGCACTATTTATGCATC | AGGCACTCGAGCAATAGAAAGT[illegible] |
| **wDcau** | **469,087-469,863** | **wDcau_UD** | **AGTGGTCGGTGGAAACGATT** | **TCATCCGCAAAACTTGAGTC** |
| | 683,984-684,538 | wDcau_N4 | AGGAATTATATACAAGACT | CAGGTGTAGCTTCGTACCTT |
| **wDimm** | **309,052-308,471** | **wDimm_tig1UAC** | **AGTAAGCATTACACTGCTCT** | **AGCTGCATTTCTGGTTCGGT** |
| | 920,005-354 | wDimm_tig1_919000 | CGGTTATATTGCAGTCGTGA | ACTAGCCTAGAGGTAGGTGA |
| | 3,093-3761 | wDimm_tig1_3000 | CTGATGTTTAGCAGAAGCCT | CTGGCCATTCTTGCTGATCT |
| **wLsig** | **910,348-910,915** | **wLsig_tig1UAC** | **TGCTGATGACCCTAAAGTGT** | **ACCGAATACTGCCAGGGTT** |
| | 166,186-166,892 | wLsig_tig1_166000 | CCACCTTGCTCTTGTATTCG | TGCTAACCAGCATAATCCGT |
| | 301778-302366 | wLsig_tig1_301000 | GGAAAAGTCGTAGAATCAGC | GCCTATCACCATATTCACCT |
| | 906,793-907,420 | wLsig_tig1_906000 | GCAAGCTTCAGAAGTAAGGA | ACCCCAACCTTTACTGCATC |
| | 966,162-966,738 | wLsig_tig1_966000 | GCAAAGTTATTTGATTGCGA | TAGCAAGTCTACAGCACATC |
| | 926,196-926,822 | wLsig_tig1_926000 | ATTATGGATTGGTCTTGACG | AGGGAGCTAAAGTCTCAGT |
| | 911,336-911,823 | wLsig_tig1_912000 | GGTCCAGCTTCAGGAGTAA | AACTCTTCCTGTCTACCTTT |

**Amplicon sequences:**

**>wCtubN1201N5**

aTAGCCTATGAcaCaGATAGTAGAAGCATTTGGCAAGCCGCATGTAAATGTGGGAACAATAGGGCACGT
AGATCACGGGAAAACAACGTTAACGGCAGCGATAACGAAGTATTATGGGCATTTCATAGCGTATGATC
AAATAGATAAGGCACCTGAAGAGAGAAAGAGGGGAATAACGATAGCAACAGCACATGTTGAGTATG
AGACAGATAAGAGGCACTATGCACATGTTGATTGTCCTGGACACGCTGACTATGTGAAGAATATGATA
GTAGGTGCAGCACAGATGGATGCAGCGATATTAGTAGTATCTGGAGTTGACGGGCCAATGCCACAAA
CAAGAGAACATATATTACTTGCAAAGCAAGTTGGTGTTAAGTATATCGTTGTGTACATAAACAAAGCT
GATGTAGCTGATCATGATATGATTGGTTTAGTAGAAATGGAAGTTAGAGAATTGCTGAGTAAGTATGA
ATTTCCCGGTGATGATGTTCCTGTAGTGATTGGGTCTGCATTAAAAGCATTAGAAGATGAAGGTAATG
AGTATGGGAAGAAATCAATAGGAAAGTTGATGGAAAAGTTGGATGAATATGTAGCAGTTCCTCCAAG
GCCTGTAGATTTGCCGTTTCTAATGCCAATTGAAGATGTATTTTCAATACCTGGgCGTGGaACGGTAGT
GACAGGAAGAATAGAGAAgGGAGAAATAAAGACAGGCGAAGAGATCGAGATAGTAGGGTTGAAAGC
aACgcgGAAgacAaTatGTACAGGGgTAGAAATGtTC

**>wCtubN1205N268**

CTGGTAgatTAgAAcTGCTCTTTTGTGacGTAaTAGCGATTTGTGTGTGAACACTACTAAAGGCTTACGGAA
GTTTCGATGGATTTGCCTACGAAGAACATGAAAGTAATTTGCAGGAGTAGAGCAATTAACTACTTGCA
TATTATCCTCTGCACAAAGTTGTAAAAATCTTTCTATACGTGCTGAGCTATGCTCAGGTCCTTGTCCTTC
ATAACCATGAGGTAAAAGTAAAACTATACCACTAGATCTCAACCACTTTGTTTCTGCTGATGCAATGA
ACTGATCGATTATAATTTGTGCGCCATTTGCAAAATCACCAAACTGCCCTTCCCAAAGCACAAGTGAA
TATGGAGAATCAAGGCTATACCCATATTCGAAGCCCATTACAGCATACTCTGATAGGGCACTATCTAT
AACTTCAAAGTAGGCTTGTTCTTCATTTATATTATTCAGTGGAGTAAACACTTCTTCTGTTACTTGGTCA
ATAAGCTTTGAATGACGGTGAGAGAAAGTTCCTCGGCCAGAATCTTGACCTGACAAGCGCACTTCTAT
TCCTTCTGTAAGTAGTGACGCaatgc

**>wCtubP341224**

TTCTAtTGtTTGtAGTATTTTCTTCTCTACCaAAAATATTCTTTCTATGACCTTTGAATAGTACTATTACCG
CCCATCCTCCTGTCCCTGGATTACCAGAACAGGCTCCATCTGTGTATATTGTTACTTCCTTTTTGTTCAT
CTTTTTACTATGTAGATAGATAAAGTATATATTTGCTTCTACGAAAGTTCAATTTTAATGAGATTTATAT
GCTAAGGCAAACTTAAACTGTATTTTTAAGTTTTACTATAGAAACAAAAAAAAGCCCAACAACGAGCA
ACATTAATAGAAAAAGCAAAAAGTAGTTTTATATTTCTGCTGTTCTTTCATATCTAGAACCAATCGAT
AGAGCAAAAAATAATACTAGCAAACAAAGAAAATAGGTAAGAAATTGAATAAGAAAACATTCTTTTT
TGTGAATGGTAATCCTTGAACTTAATAATAGATATAGCATACCAAATAAAAACACAACCTTCAAGAAT
TGCTATACTCAGGTAAAGTAGAACATTTTTTAAAAACAGTGATGGAAGTAAGCTAATTAACACTAAAA
GTATACTATAAATTAGTATATATTTTCTTGTTCTCTCTGATCCGTAAACAACATTAAACATTGGAATTG
ATGCTTTTGCATATTCTTCAGACTTGTTTAAAGACNGAGACCAAAAGTGTGGTGGAGTCCATATAAAA
ATTATTAAAAATAGAATAAAACTCTCCCaACTAACAGCATTAGTTACGGttGCCCAACCaATCATTGGaga
AGAAAGCACcTGaTGCGC

**>wDcauUDF**

agGGaagAAgtGCTCATAGTCTTGGTTCGCCGCTAGATCCaAGACTTACTTTTAATAATTTTGTAGTAGGAA
AACCGAACGAATTAGCATTTACAGCTGCAAAACGTGTAGCAGATTCTATAGATCCAATATCAGGGAGT
AATCCTCTTTTTTTATATGGTGGAGTGGGGCTTGGTAAAACACACTTAATGCATGCTATAGCTTGGTAC
ATCATTAATTCTCTTCCAATGAAAAGAAAAGTAGTATATTTATCAGCAGAAAAATTCATGTACCAATA
TATTACAGCGTTACGAAGCAAAGATATTATGTTATTTAAGGAACAATTTAGATCAGTAGATGTGTTGA
TGGTAGATGATGTACAATTTATTAGTGGTAAGGATAGTACACAGGAAGAATTTTTTCATACTTTTAATG
CGCTAATAGATCAAAATAAGCAGTTGGTTATCTCTGCTGATAGATCTCCTAGCGATCTTGATGGAGTA
GAGGAAAGGATAAAGTCTAGGCTTGGTTGGGGATTGGTAGTGGATATTAATGAAACAACTTTTGAGTT
AAGACTTGGTATATTGCAGGCCAAAATGGAACAAATGAACATGTATGTTCCTGATGATGTCCTAAAAT
TTTTAGCAAGAAATATAAAGTCTAATATAAGAGAATTGGAgGGGGCCTTAAATAAgGtTGCTCATACTT
TATTGATTGGAAGAAGTATGACAGTAGAGTCAGCTAGTGAAACCCTAGCAGATCTTCTCaGATCAAAC
CaTaAGCCAATTACAATAGCA

**>wDimm_tig1UAC**

ACTAGTCTATATTAAATATAACAGCTACTACTTATCTTTTCTTGCATAATTTTTCCGTATTTTTATATTAT
ATATACAAACTCTTCTGAATCTAGTCATAGTATTGATACTCTTAATATTTCGCACTACTGAAAACCAGC
TTATACATTAGTCCACCAAACGAAAACCCTTGATAATTACACAAAATATTCCTCTACAAGCTGAAGCA
CACCCTCACCTCCAATTAACAGTACAAAAAAGCATATTCTATTTCATCAAAACTCAGCTATTTTTTTGT
ACCTACCTAAATTTATTCATCCACTGCTTTTTTGCAGGCTACAATTACAAGTATTACACAGTAGTTATA
CTCAATGCTCATATTTTTCTCATGTTATTTAATCGGTTACCCTACTCTTCTTCTACTACCACATCCAAAT
AATATTTCCTTTATAACATGAGCAGTACCACAGACAATTGGAGACTATCCTAACCATAAGTTGAATGA
ACCTGAAGTTTTTTTAATCTTACCTATACATTACCTTTTGTTTGGTATCAAGGCTATAAATATTCCATTA
CTAAATTTATCCACCAAAAAGAACTCAA

**>wDimm_tig13000**

CTTTATCATAGCGTTTTACTTGACTTGCTGTAGGTTTCATTATCTCTTCCACGCTGTTTAAAAGTTCTCT
ATGTAATATTGTATTTGTTACCTGGTATGTGTGGCAAATGTCTGCTTAATGTATTTTGTTGTATAATTTT
GTAGCACATTGCCTATTATGCTCATACTCCTTATCTATTTTTGCTTTAGCATTATGTTATTCTGTATGTA
AGAAGATTAGTAGTTGGATAGTTATGGAGGTTTGACTTTGTTAAATTTCAATGAATTGGGAGAATATTT
ATAAAGATGTTGCTTTTAATGACTAAAACACCTACAATCCTAAATTTTAAACAAGTTTTATGCCACGGA
GGAATTTTTTAGTATGGGTGTTGGCATCACTGTTTTATGCATACCAATATGTGTTACGTGTGATTCCAA
ATATAATTGCGCCCGAGTTAATGACGAAGTTTAATGTAAATGTAGTTGATATTGGTCAATTTAGCGGCT
TATACTATGTAGGTTATACATTAGCTCATATACCTGTTGGTCTTTTTCTTGATAGGTCTGGGCTAAAGTT
TGTTTTGCCTATATGTACTATTTTGACATTTGTTGGAACGCTGCCACTTATATGTTTTGATAGATGGTAT
CCTTCAATACTTGGTAGAATAATCGTTGGAATTGGGTCATCTG

**>wDimm_tig919000**

AAACTTCAGTAAATCATTAAATATAGTAAGACCTGGCAGATCTTTCATCAGCACTTCAGTTGCACACTT
TCTTAATACATAAAACCTTTCAAAAAGAAATAGAAAGACTCTACATGCAATCCTTTCCCTCAAAGTCCT
TTTTTTGCTCAAGTTGCTTCCTATAACAGACAAAATGTTTTCATATTAGATTTATCAAGGAACTGTTGT

CAAATATTTAATAAGCATGTGAATAACACTAAATTAATCTGTGAATTTCATGTTGATATAAGATATTAA
CAGCTCCTTAACAACTTTTTCACCAGCATATACATAGTTGTACAGCAGTGTGCAACTGAAAAAAAATTT
TCACAAAGAGAGATTACTAACAATCTAAACATTCTACTTATGCACATAAATTATAACAGAAAGTCAAT
AGTATATAAAAGTTATTAACACACTTATTCACAAATTAGGTAAAGAAAATCTTCACCT

**>wLsig_tig1UAC**

TTTTATCTTACTACACAATCCAATACTAATGACTAGAGCTCCATGATTGGAAATAGTGTGACAGCTCAC
TAAAAACAATAAAATAAAAAATATTAGCACTTGCATCTCTATTAATAAATCACTTAAAGCTACAAACA
TTAACAAATCACCTGTCGTCAAATTTTGTAAAAGTTAAATTGACTGATTTTTGATATCGCTTAATATAA
TATTTACTTCAGAAAATAATAGGTGTTCTATGTTATTGTTAGAAGCAGATCCAATATATAAACCTTTTA
ATTATCCTTGGGCGTATGATGCCTGGTTACAGCAGCAAAGAATACATTGGATACCAGAAGAAGTTCCT
CTTGCTGATGATGTGAAAGACTGGAAAGCTAAACTTTCGAATGTAGAAAAAAATTTACTAACTCAGAT
TTTCAGGTTCTTTACTCAAGCAGACATTGAAGTAAACAACTGCTATATGAGACATTACTCAAATATATT
TAAACCAACAGAAATATGCATGATGCTCGCAAGCTTTTCCAATATGGAAACCATACACATTGCAGCCT
ATTCTTATCTTTTAGATACAATTGGCATGCCAGAAAGTGAATATCAAGCATTTTTAAAATATGATGCTA
TGAGAAAGAAATATGAATATATGCTAGAATTTGAGGAGTGTAAA

**>wLsig_tig1_166000**

ATTTGTATTTTTTCTTATATCATCTAAACCTTTAGCTAATTTTGCATTGAAATTCTCTGTGAGTTTTGTAG
CTAGTGCTTTTTCATTTTTGTTAAAAGCTAGTATATAGCATTGAAATCAAAATTTTTAAGTAAGAAAAT
CATAGAGTCAGCAAAAAAAATGCTTGAAGGAAGTGTAAAATAATGCATATGTTGCAAAAAAACACTAA
ATGCTGTAATTGTAGCAAAAAAGTACTGTGTAACAGCTGCAGCAAAGTTTCATTTAAAAGAAGAAAAT
TGTTTGCTCTTCTTAAGATCGTAGAAAAACTAGATTGAATCAGAGTCGGTTTGAATGAGTTGAAGTATG
AATGCAAGAACATTCTGATATTTTCATTAAAGAAATGAGATTCAGGAAAAATTTGGCTTGAATATGTG
AATCTACAGCCATACCATAATATGTAGAGAATGAAGTTTTCGTATATTGCATCAAGATCAATTCATAAT
GAACAAGATAAAGGTAGACAGGAAGAGTTTTAAAAAACTTAATGAGACTATTGGCAAGTATCATGAA
CAAGAGTTATTCATCTTTGATGAATTATAGTTTAGCACCACACTTGAAAGTTAGACAATGGTAGTTAGA
AAAGGACGTTAAAACACAGGTTAAAGTAAAATTAGGTAGATAAAATTTTTATCTCCACAATGTGGTTA
ATTCTAGAGATGGAGAGAGCCC

**>wLsig_tig1_301000**

ACTAAATGCTGTGATTGCAGCAAAAAAAAAGCATAATATAGCAACCTTAAGAAAAATATATTTCTCTTT
CAAGAACAGCACTAACTGCGTAGATAAAGCACCTAAAACTAGGAAGAGAAGAAAAAATTGGTTACTTC
CACCCTTCAACGCCATAGAAAAATTAGATTGAAATTAGAGTAACTTCGAATAAGTTGAAGTATGGATA
CAAGAGAATCCTAATGTTACTGTAAAAAATGAGAATAAGAATCTAGGAAAAATTTATTTTGAATATCA
GTAAATCTACAGTACACCTTAGAATATGCAAAAAATGAAATTTTCGTATATTACGCCAAAACCGGTTT
ATAATGAGCAAAATAGAGGTAGACAAGAAGAATTATAAGAACTTAATAAGAGACTATTTGGTAAGTA
TTCTAAACAAGAGCTATTATTCTTTAATAAACTGCGGTTTGATATACAATAGAAAATTGGACATCGGTA
GTTTAAAAAGGGTGTTGTACCAACTCTTATTATTAATGAGTCAAATTATTATTAGTGGGTCAAATAAGT
AACATTACAGAGCTATTTAATTGCGGAGTGAGAAAGAAGAAAGGTAATAAATTTGCACAAGCACTCTT

AAGCAAGGCAATTGTATTGTAGGCTTTATTATGCAAA

**>wLsig_tig1_906000**

AAAAATACGAAAGTTGGATCGTATACAACCCAATCACAATTTCCCCGGACAAAACAGTTGCAGAAGC
AATTTCATTGATGAAAGAGCATGATTATTCTTGTATTCCTGTAGTTGAGCAATGCAAGTTAGTCGGAAT
TTTAACTAACAAAGATATAAGATTTATTGAARATCAGAACATGAGTACAAAAGTCTCTGAGGTAATGA
CAAAAGAGAAATTAGTTACAGTACGGGAGCAGGGGATAGACAGCACTTCAGCAATGAAACTCCTGCA
TGAAAACAGGATAGAGAAGCTTTTGGTCATAGATGAAAACTTCTGCTGCATAGGTCTGATCACAGCTA
AAGACATTAAAGAATACAATAGATACCCCAATTCATGCAAAGATAGTAAAGGGCGGCTCAGAGTTGC
CGCTGCAGTTGGCACTGGTAAAAAAGATGGTATAGAAAGATGTGAAGCTTTGATCAGAGAAGAGATT
GACGTGATTATTGTAGATACTGCTCACGGTCATTCCGAAAATGTTATTAATACTATTAAAGAAATAAA
AACAATGTATCCAAGTACGCAGCTAGTTGCTGGAAATATTACAACAAAAGAAGCTGCTGAAGCRTTGA
TTAACGCTGGTGCTGATGC

**>wLsig_tig1_912000**

AATTGGTGGGTTACAAGCTGCAATGGGGTATACCGGTAATAGAAATATAGAAGAGATGAAGAAAAAT
TGTAAGTTTACAGTTATGACTGCATCAGGATTAAGAGAAAGCCATGTTCATGACATAACTATCACACA
GGAATCTCCAAATTATGTTTATCAAGTATCCAACAACTTGTCAAGTGATTCAGACATCTAATTGTTAAT
TGCCAGCTAATACAGTAAAAGTAAACAGGATAAAGCAGAAATTTTTGCACTAATACAAAAGTCCCTGC
TGTAGTTAAAAATGCATTATTCCTTAATTTGTCAAGTTATAATGCACGTTTGTTGATCTATGTAATATAT
ATTATACCTGTACTTGCCTGTACATCCTACTACATCTACCCCGCATCATTATCAATTACCCACCTTCACC
ATCTTCTGCTCTATCTTCATTATAAGACCAATTTCTACTATATAAAGAACGGATAGACAATAATGAACA
AGATAAA

**>wLsig_tig1_926000**

GATATAAAATTTCATTTTATGCATATAATGATGTATAGACTTGCTGACATTTAAGTTAAATTTTTCCTA
AATTCTTATTCTCATTTCTTTAATGGTGATATTAGTTTTTTGTATTCATATTTCAACTTGTTTAAATTGAC
TCTGATTCAATCTAATTATTTTCCGATGTTGAGGAGAAGCAAATAAATTTCTTTTCTTTCAAATTTTAGT
ACTTTATCTGTGTAGTTAGTACTGTTCGCAAAATGAGACATATTTTTACTACAGCTGTTATAGTGTATTT
TTCTGCAATTACAGCATTTAGTTTTTTTGCAACATATACATTATCTCGCATTTTCTTCAGCATTTCTTGT
GCTGATTTCATGACTTTTTCATTTAATAATTTTGATTTCAATATCATTTAAACCTCGTTATTTACTTGCTT
CAGTTTGGCAAGAATTAGCATAAGATTTTCCCTGAAGAGATCTACATACAAAAGGCCTTTGAACCTAA
AAAATTACCAAGATCTCACATATATAATGCCCATACTGTTAAAAATGCTTTTGTACTATTTTATCTATTT
CTTCTAAATATTTATCACTAGAGTTACTTGTATTACAAGACCTATTTATAAAATATAAAATTCTTAAA

**>wLsig_tig1_966000**

GAATGAGAAAGAGAGGTAATATGTTTATATAAAGTGCTTCCAAGCAAAGCAATTATGTTATAAACTTT
ATTATGCAAAATGGTCTGTTTATATAGATAACCAAAGTCTCTCAATAGGATTGAATTTAGGTAAGTATG
TGATGGTAGATATATAGTCTTAATATTTTTAGGCACATTAAAATTTTTCTACTTATACTAACTAGTACA
ATTCATGACAAGAAATGCTCTTCATGTTTCTAAATACTGCAACATTTGTTCAAGAAACATATTTATACA

ATCAGTGTTGACATTCGATGCAAATAAGATAGAATTCTTCCCATTTCTAAAATGGGCTGCATTGTAGAA
GTAAAAACTTTATCTACCTAATTTTACTTTAATCTGCGTTTTAACATCCTTTTTTAATCACCCGTATCCA
ACTTCTGAATGTGTGCTAAACCACAATTCATTAAAGAAAAGTAACTTTTTTTGAGATACTTACTAATAA
TCTCGTCAAGTTTTTTTAACTCCTCTTGTCCACCTTTATCTTGTTCATTGCAAGTTAGTCTTGGTGTAATG
TATAACATTTTTTGCATATTAAG
